# MetaLab 2.0 enables accurate post-translational modifications profiling in metaproteomics

**DOI:** 10.1101/753996

**Authors:** Kai Cheng, Zhibin Ning, Xu Zhang, Leyuan Li, Bo Liao, Janice Mayne, Daniel Figeys

## Abstract

Studying the structure and function of microbiomes is an emerging research field. Metaproteomic approaches focusing on the characterization of expressed proteins and post-translational modifications (PTMs) provide a deeper understanding of microbial communities. Previous research has highlighted the value of examining microbiome-wide protein expression in studying the roles of the microbiome in human diseases. Nevertheless, the regulation of protein functions in complex microbiomes remains under-explored. This is mainly due to the lack of efficient bioinformatics tools to identify and quantify PTMs in the microbiome. We have developed a comprehensive software termed MetaLab for the data analysis of metaproteomic datasets. Here we build an open search workflow within MetaLab for unbiased identification and quantification of PTMs from microbiome samples. This bioinformatics platform provides information about proteins, PTMs, taxa, functions, and pathways of microbial communities. The performance of the workflow was evaluated using conventional proteomics, metaproteomics from mouse and human gut microbiomes, and modification-specific enriched datasets. Superior accuracy and sensitivity were obtained simultaneously by using our method comparing with the traditional closed search strategy.

## Introduction

The relationship between the microbiome and human health has been a very active subject of research in recent years^1, 2^. The study of host-microbiome relationships has been primarily focused on the composition of the microbial community and the relative abundance of microbes using DNA sequencing-based technologies, as well as functions of the microbiome using metagenomics and metatranscriptomics^3^. Metaproteomics provides direct assessments of the microbiome functions by qualitative and quantitative investigating the expressed proteins products^4–8^, which plays an important role in microbiota researches^9^.

Currently, the data interpretation is still a bottleneck of metaproteomic studies. To overcome the challenges in data analysis, we have built MetaLab, which is an integrated software platform specifically designed for the metaproteomic data analysis^10^. MetaLab provides a complete workflow including sample-specific protein database generation, peptide/protein identification and quantification, taxonomy analysis and functional annotation. MetaLab is written in Java with a friendly graphical user interface that can run on all platforms. For the convenience of the researchers, MetaLab also provides a web-based version for the online data analysis^11^.

Post-translational modifications (PTMs), which are important for the dynamic regulations of the biological functions in the microbiome, can also be identified by mass spectrometry-based metaproteomics^12^. Yet, the characterization of the PTM profile from microbial communities brings additional computational challenges in data interpretation. Conventional protein identification strategies only allow a few pre-selected PTMs to be identified (variable modifications in database search). In contrast, the open search strategy enables unbiased identification of potential PTMs and amino acid variants^13–15^. Unfortunately, the application of open search strategies in metaproteomics is still hampered by the lack of effective bioinformatics tools. Open search strategies increase the search space when considering multiple possible mass differences for modified peptides. This is compounded for metaproteomics as protein sequence databases are already very large. Generally, an increase of several hundred-fold in processing time occurs during open searches. Fortunately, several recently developed software tools have significantly improved open search speeds^14, 15^, making open search feasible on a metaproteomic scale. Nevertheless, considering the complexity of microbial communities, ensuring the accuracy of peptide and PTM identifications is difficult. Several hundred potential PTMs can be found from one metaproteomic dataset. Calculation of the false discovery rate (FDR) at the peptide-spectra match (PSM) level provides only an overall estimation of the confidence of identified peptides. Considering the complexity of microbial communities, ensuring the accuracy of peptide and PTM identifications is difficult.

To solve this problem we built an updated version of MetaLab (MetaLab 2.0) which incorporated an open search workflow to enable the study of PTMs in microbiota samples. We developed a novel multistage filtering strategy in our workflow, which can efficiently validate the open search results. The profile of PTMs of microbiota samples, as well as relationships with taxa, functions, and pathways, can be obtained easily by our software. Comparing with the general used closed search strategy, the accuracy and sensitivity of identifications were improved simultaneously. Traditional proteomic studies using single species are also supported by this strategy. MetaLab 2.0 is free for academic use. Both the online and local versions of the tools are available at https://imetalab.ca/.

## Results and discussion

### Open search workflow of MetaLab 2.0

The open search workflow of MetaLab 2.0 was shown in Figure 1. In MetaLab 2.0, MSFragger^14^ was incorporated as the open search engine into the workflow. Then we developed a multistage filtering strategy for the evaluation of open search results. Because only controlling the global FDR of whole identification results at the PSM level is not sufficient for the investigation of PTMs, we evaluated the confident level of each potential modification in our workflow. Then multiple strict filtering criteria were applied in all of the PTM, PSM, peptide and protein levels, which significantly improved the sensitivity and accuracy of the open search results. Since quantitative analysis is an indispensable step in proteomic data processing workflows, we incorporated a quantitative tool, FlashLFQ^16^ into our pipeline. By combining these modules with our previously developed taxonomy analysis and functional annotation modules^10, 11^, we built a complete open search data analysis workflow in MetaLab 2.0, which enables researchers to study the profile of PTMs of microbiota samples, as well as relationships with taxa, functions and pathways. Each step of the process was detailed in **Materials and methods**.

**Figure 1.**
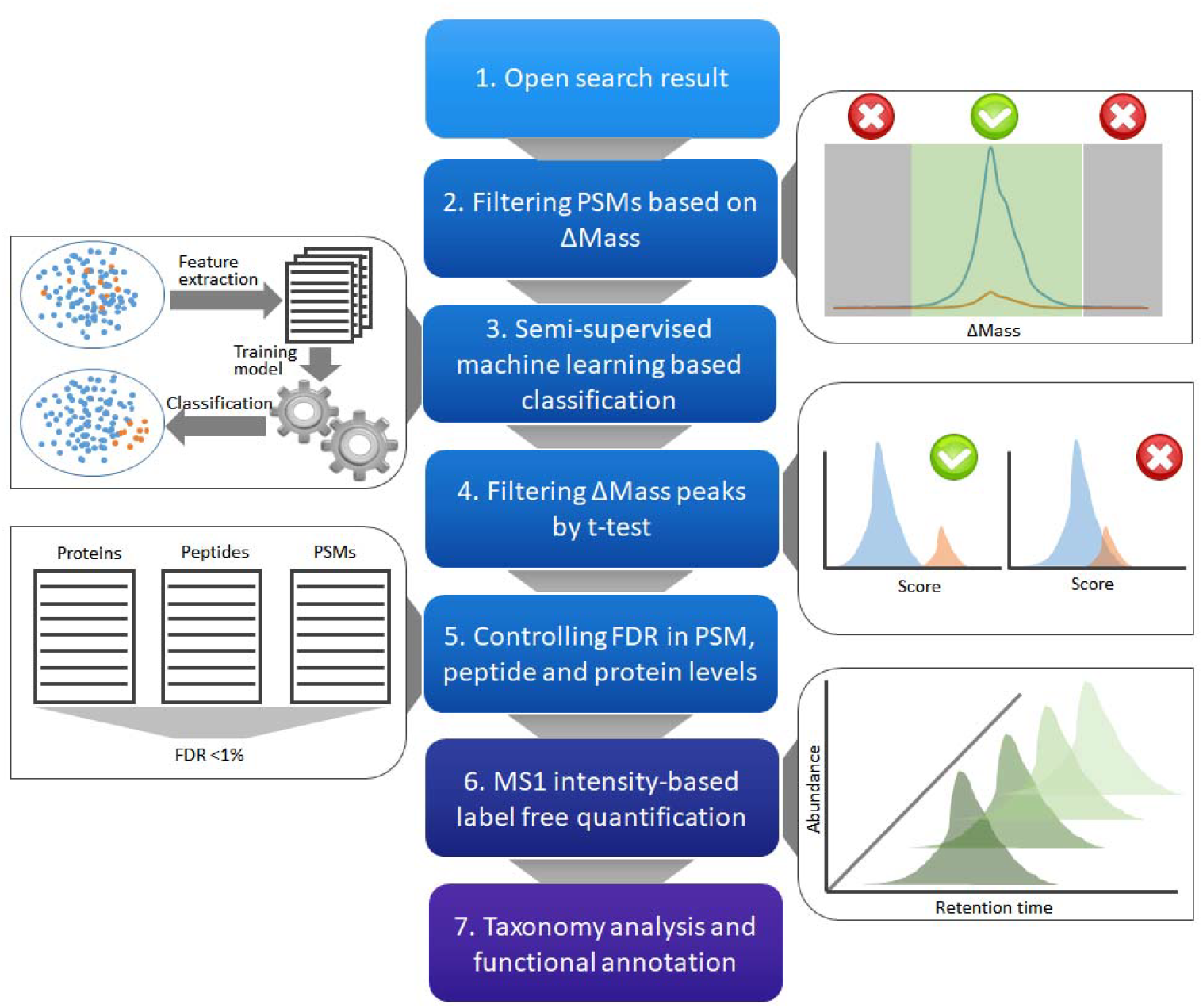
The workflow for the post-analysis processes of open search datasets. After the open search result was obtained, a multi-stage filtering strategy in steps 2-5 was adopted for the validation of the results. Then MS1 intensity-based label-free quantification was performed. The final step was taxonomy analysis and functional annotation for metaproteomics studies.

### Tackling the problem of false discovery rate control with a multistage filtering strategy

The validation of peptide identification is a tricky problem in proteomics. The most prevalent strategy for validation of methods in proteomic software is the use of a target-decoy database search, which provides an estimate of the probability of random matches. Generally a FDR cutoff at 1% is accepted, however, it has been proposed that a lower FDR threshold (of about 0.1%) is required for open search results, which would ensure a comparable confidence level with the conventional closed search strategy^17^. Moreover, only controlling the global FDR of whole identification results at the PSM level is not sufficient for the investigation of PTMs.

Herein we developed the multistage filtering strategy for the confident identification of peptides and PTMs using open search. We applied this strategy to a large-scale proteomic dataset from the HEK293 cell line^13^ (**Supplementary Data 1**). After the PSM list was obtained and the Gaussian peaks along the ∆Mass range were fitted, we investigated the score distribution of the four types of PSMs: unmodified target PSMs (PSMs belonged to the Gaussian peak located at ∆Mass=0, annotated as “peak 0”); target PSMs belonged to any other Gaussian peaks than the “peak 0” (“peak other”), e.g., the 4 peaks annotated in Figure 2a; target PSMs outside of any Gaussian peaks (“no peak”); and all of the decoy PSMs (“decoy”) (Figure 2b). Overall the unmodified target PSMs were assigned with higher scores. The difference between “peak other” and “no peak” was major. However, the medians of “no peak” and “decoy” were quite close. This result showed that PSMs that did not belong to any ∆Mass peak were more inclined to be random matches. It was observed that PSMs that belonged to the fitted Gaussian peaks had higher scores than the PSMs outside of the Gaussian peaks (Figure 2c-m). For this dataset, 44.7% (463,673/1,038,110) PSMs did not belong to any Gaussian peaks. After this portion was removed, a FDR of about 1.1% (6,268/568,169) was obtained from the remaining PSMs. In this step 207 ∆Mass peaks were kept, among which 160 peaks were identified as monoisotopic peptide peaks. As a result, aside from the peak with ∆Mass of 0 representing unmodified peptides, 159 types of potential modifications were obtained. To further validate the result, we compared it with the closed search result. In the closed search 46,868 decoy PSMs were obtained in total, with 10,236 of the same spectra also identified as decoy matches in open search (**Supplementary Fig. 1**). This portion can represent random matches. We found these decoy matches were generally distributed evenly in the ∆Mass range (about 10% were concentrated in ∆Mass=0 because unmodified peptides had higher priority). In the filtering step 84.2% (8,619/10,236) of them were removed because they didn’t belong to any ∆Mass peak. This result proved that retaining only those PSMs belonging to high quality Gaussian peaks in the ∆Mass range can effectively eliminate random matches.

**Figure 2.**
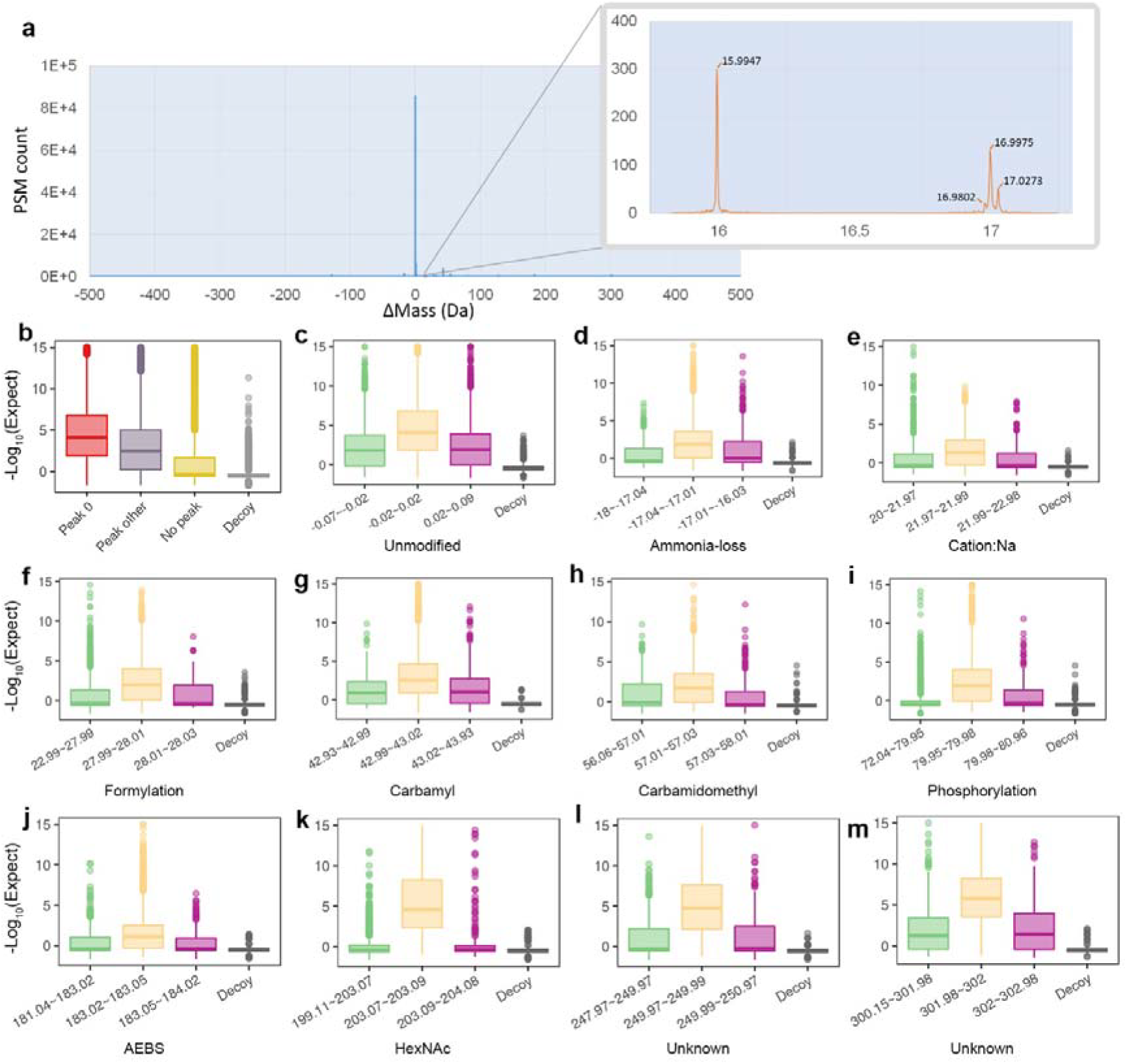
The PSMs whose ∆Mass falled in a specific range (detected ∆Mass peaks) will have higher scores than the PSMs out of the range. **(a)** The density of ∆Mass distribution of the identified PSMs. The zoomed-in part shows four ∆Mass peaks which were detected by Gaussian fitting along the ∆Mass range. **(b-m)** Score distributions of PSMs identified by open search. The score was calculated as -Log_10_(Expect) and higher scores represented better matches. The bottom and top of the boxes are the first and third quartiles, respectively, the middle lines represent the sample median. **(b)** Peak 0: the scores of the PSMs in the fitted Gaussian peak on the position ∆Mass=0, which consisted of PSMs without modifications; Peak other: the PSMs in other fitted Gaussian peaks than Peak 0, which consisted of modified PSMs; No peak: the scores of the PSMs with modifications but do not belong to any ∆Mass peaks; Decoy: the scores of all the decoy PSMs. **(c)** first box: the scores of the PSMs with ∆Mass from −0.07 to −0.02. −0.07 was the right edge of a Gaussian peak and −0.02 was the left edge of the next Gaussian peak, therefore the PSMs in this range did not belong to any Gaussian peak; second box: the scores of the PSMs with ∆Mass from −0.02 to 0.02. −0.02 and 0.02 were the left and right edges of a fitted Gaussian peak respectively, which consisted of unmodified PSMs; third box: the scores of the PSMs with ∆Mass from 0.02 to 0.09. 0.09 was the left edge of the next Gaussian peak, so the PSMs in this range also do not belong to any Gaussian peak; fourth box: the scores of the decoy PSMs with ∆Mass from −0.07 to 0.09. **(d-m)** examples of the scores of the PSMs with other potential modifications. Similarly, the first box represented the scores of the PSMs between this Gaussian peak and the previous Gaussian peak; the second box represented the scores of the PSMs belong to this Gaussian peak; the third box represented the scores of the PSMs between this Gaussian peak and the next Gaussian peak; the fourth box represented the scores of all the decoy PSMs in this range.

After the first filtering step the FDR was 1.1%, but it should be noted that if we calculated the modification-specific FDR, i.e., calculating the FDR of the corresponding PSMs for each ∆Mass Gaussian peak, the FDRs were below 1% for only 12 potential modifications. The FDR of the unmodified peptides which accounted for 80.2% (460,546/574,437) of the total amount of PSMs was only 0.28% (1,290/460,546). This meant that the FDR was higher for the modified portion. Only controlling the FDR at the level of PSM cannot assure the confidence of PSMs with certain modifications. Therefore, we added filtering at the level of modification. Firstly a classification score was assigned to each PSM based on a semi-supervised machine learning strategy^18^ (**Supplementary Table 1**). Then for each potential modification, an unpaired two-sample t-test was performed between target PSMs and decoy PSMs using the assigned classification score. If the two sets of score values showed significant difference (p-value < 0.01), this modification was retained. Following this step, 17 ∆Mass peaks were discarded. The remaining modifications were considered “true” with PSMs having significant differences between target and decoy matches. The last filtering process was adopted at the protein and peptide levels. The protein inference algorithm was utilized at this step to calculate the scores of the proteins and peptides^19^. Then the protein and peptide lists were filtered to ensure that, at both levels, the FDRs were less than 1%. For this dataset the final FDRs were only 0.08% (441/562,302) at the PSM level and 0.15% (179/120,037) at the peptide level, respectively (**Supplementary Data 1**). In total 142 possible modifications were detected, with 50 modifications matched with items in Unimod (http://www.unimod.org/) and 40 unknown modifications that can be constructed by combining two modifications from the 50 known modifications (**Supplementary Table 2**). We analyzed the compositions of 40 unknown modifications and found that 72% (29/40) of them contained at least one modification listed below: −156.1019, 156.1014, 128.0951 and −128.0958 Da, which corresponded to addition/loss of Arg or Lys. Loss of the terminal amino acid by in-source fragmentation was common in ESI-MS. Addition of Arg or Lys could be caused by additional missed cleavage sites (consecutive Arg or Lys in the peptide C-term). In addition, 25% (10/40) of the combinations contained −71.0375 Da (Gln➔Gly), −71.0743 Da (Lys➔Gly) or −57.0221 Da (Asn➔Gly or Gln➔Ala), which were related to amino acid substitutions. Therefore, from the combination of information we can determine that potential modifications without Unimod matched items, can be explained by known modifications to a large extent.

Using this strategy 562,302 PSMs were identified with an ultra-low false discovery rate (FDR) 0.08% and protein level FDR less than 1% (**Supplementary Fig. 2**). In a closed search, 385,990 PSMs were obtained in the same situation which is 45.7% fewer PSMs than our strategy. The significant improvement of sensitivity using our strategy was not only due to the identification of modified peptides, which cannot be identified by the conventional method, but also because of the accurate match of spectra to peptides. In the above dataset 36.2% (16,969/46,868) decoy PSMs in the closed search were matched to more credible targets in our open search. Therefore, some of the false negatives from close search were recovers in our open search while additional expense for the elimination of these random matches in FDR control was avoided. Moreover, the random matches in the open search were evenly distributed over a wider range, which made the delta mass (∆Mass) a significant feature to distinguish random matches effectively. If this strategy was not applied and the open search result was filtered directly, only 388,512 PSMs were identified with FDR less than 0.11%.

### Identification of a group of phosphorylation related modifications from enriched samples

To further benchmark the post-validation strategy, we re-analyzed *E. coli* phospho-enriched samples^20^. Authenticating our strategy, phosphorylation was identified as the most abundant modification from the phospho-enriched *E. coli* sample. Even more substantiating, the 2^nd^-4^th^ potential modifications, *i.e.*, phosphate-ribosylation (∆Mass=212.0091), PhosphoHex (∆Mass=242.0195) and 3-phosphoglyceryl (∆Mass=167.983), were all phosphorylation related modifications (**Supplementary Notes**). Actually, by manual interpretation of the spectra from the top 30 abundant mass differences, we found that neutral loss of phosphorylation was observed from 14 types of potential modifications (**Supplementary Data 2, Supplementary Material 1**). Aside from phosphorylation, pyrophosphorylation (∆Mass=159.9328), and the abundant phospho-related modifications mentioned above, other known phospho-related modifications also include phosphogluconoylation (∆Mass=258.0149) and pyridoxal phosphate (∆Mass=229.0134). The possible structures of the identified phospho-related modifications were illustrated (Figure 3).

**Figure 3.**
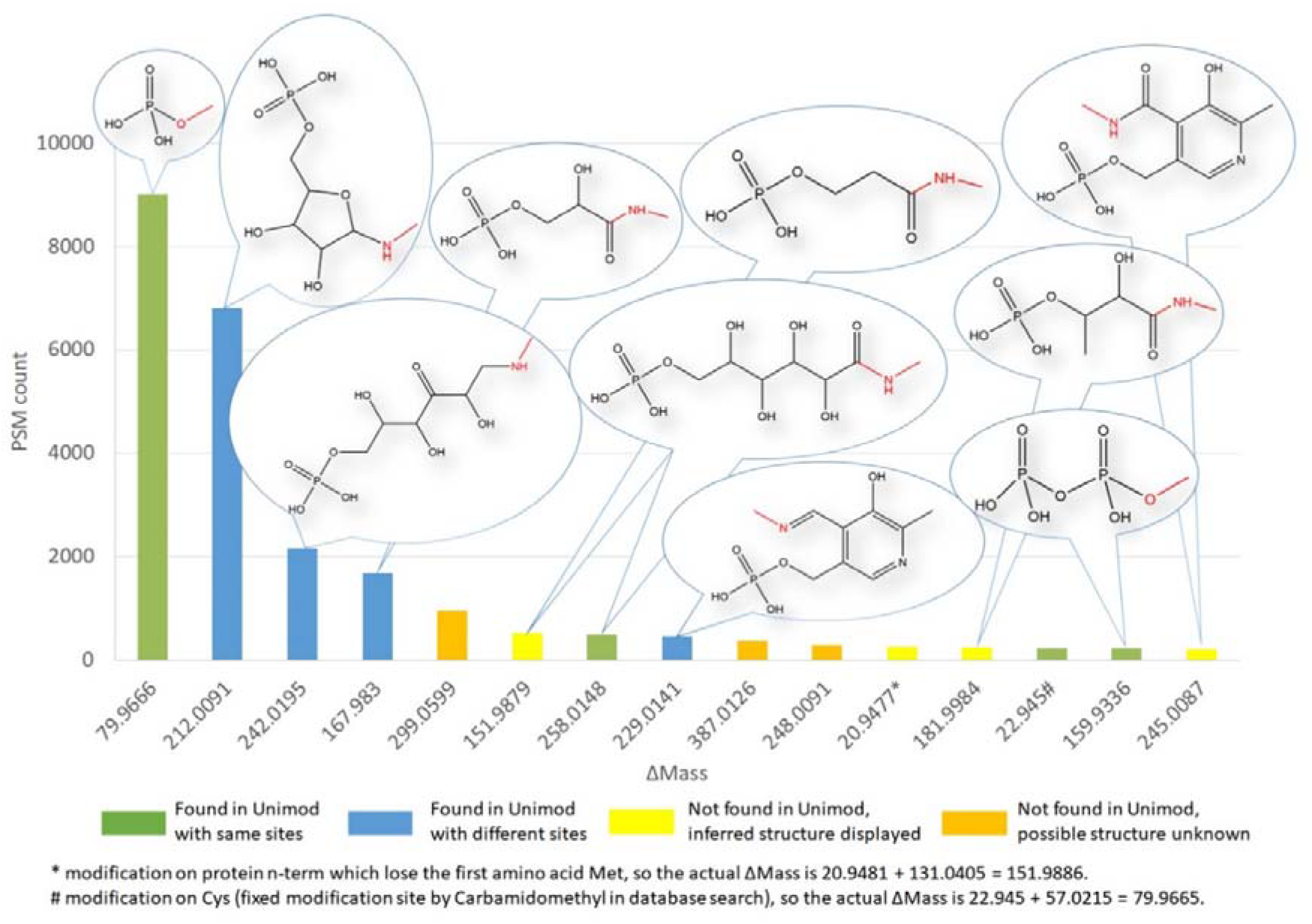
Phospho-related modifications identified from phosphorylation enriched *E.coli* samples. The possible structures and the corresponding PSM counts of phospho-related modifications were showed.

Encouraged by this success we investigated phosphorylation enriched mammalian samples from mouse kidney (Dataset ID: mouse_phos)^21^. Surprisingly, only two of the top 30 ∆Masses were matched with known modifications in Unimod, namely phosphorylation and pyrophosphorylation (**Supplementary Data 3**). It’s easy to find ∆Masses 239.8978 and 319.8637 that represented multiple phosphorylation sites occurring on one peptide. By manually interpretation of the spectra we found that all of the unknown ∆Masses were combinations of multiple modifications, and in most cases, were combinations of phosphorylation and others (**Supplementary material 2**). In fact, among the top 30 abundant ∆Masses, 26 contained at least one phosphate group, which strongly confirmed the confidence of this result. Four of them were combinations of phosphorylation and cations (protons replaced by Na^+^ or K^+^). This type was not observed for *Ecoli*_phos (but common in mouse_phos). While 3-phosphoglyceryl and phosphate-ribosylation were found with few PSMs counts and phosphoHex was not observed. Apart from this, peptides from mouse_phos tended to contain multiple modifications, which was a significant difference between these two datasets.

We also studied two human cell lines, HeLa^22^ (Dataset ID: Hela_phos, **Supplementary Data 4, Supplementary material 3**) and U-87 glioblastoma cells (Dataset ID: U87_phos, **Supplementary Data 5, Supplementary material 4**)^23^. Similar to the mouse_phos dataset, multiple phosphorylation and combinations of phosphorylation and other modifications were prevalent. But complex phospho-related modifications as these existed in *Ecoli*_phos data were very rarely observed. Our results clearly showed the different features of phosphopeptides identified from bacteria and mammal cells.

In this part we investigated the performance of our workflow using datasets derived from samples enriched for phosphopeptides. These results showed the confidence of the potential modifications identified by MetaLab 2.0. It also indicated that by the application of open search in the analysis of enriched samples, co-enriched modifications can be identified in one step, which could be helpful for the study of the interactions of various modifications.

### Characterizing the modification profile from microbiome samples

We then used MetaLab 2.0 to analyze a microbiome dataset which was constructed by assembling 32 species/strains of Archaea, Bacteria, Eukaryotes and Bacteriophages^24^ (**Supplementary Data 6, Supplementary Note**). Given the advantage of known microbial composition in this dataset, we also used this dataset to benchmark the quantitative accuracy of our method, to investigate if the quantitation will be affected by the additionally identified peptides from open search. In total, we identified 1,048,759 PSMs with FDR less than 0.1%, which was 17% more than from conventional closed searches (**Supplementary Fig. 3**). Through quantitative analysis of the mock microbiota community, satisfactory quantitative accuracy was obtained at the species-level, which also confirmed the confidence of the identifications of modified peptides (**Supplementary Fig. 4**).

To demonstrate the capability of open searches for microbiome sample analyses we applied MetaLab 2.0 to analyze a mouse gut microbiome dataset^25^. In total 918,132 PSMs were identified with an estimated FDR of 0.17%. The PSM count increased significantly by 28.3%, 202,682 additional PSMs were identified when compared with our previous work ^10^. At the peptide level the FDR was 0.33% and 127,218 unique sequences (37,544 peptides contained at least one modified form) were obtained (**Supplementary Data 7**). In closed search 115,832 peptide identifications were obtained and the open search strategy resulted in a 9.8% increase in peptide identifications (**Supplementary Fig. 5**). The number of confidently recognized species (with at least three unique peptides) increased to 163 species compared to 143 in our previous close search, with 134 species common across the two methods. Most of the newly found species were from phyla *Firmicutes* and *Bacteroidetes*, the two most common phyla in mammalian gut microbiota. From this result we can conclude that through our strategy, similar accuracy and optimized sensitivity can be obtained by open searches when compared with traditional closed searches for the analysis of real microbiota samples.

In this experiment the mouse gut microbiome were collected from both high-fat diet (HFD) and low-fat diet (LFD) fed mice for the interpretation of diet-microbiota-host interactions^25^. In this work, based on the quantitative information of the potential modifications, we investigated the response of the determined modifications to diet in mouse microbiomes. The abundances of various modifications were calculated in each identified taxon node. Both the absolute and relative values were considered, that is, for each taxon the summation of the intensity of peptides with specific modifications were calculated as the absolute abundance of the modification in this taxon, and the ratio between the absolute value and the total abundance of this taxon were calculated as the relative abundance. Firstly, we tested if there were modifications that showed significant differences between HFD and LFD. We found that the modification ∆Mass=71.0368 (Acrylamide adduct/Gly->Gln substitution) showed significant increases in the HFD samples using both absolute (*p*=0.00044 by a two-sample, two-tailed t-test) and relative values (*p*=0.00072 by a two-sample, two-tailed t-test), and the modification ∆Mass=52.1229 (replacement of 3 protons by iron) decreased significantly in relative value (*p*=0.0044 by a two-sample, two-tailed t-test) (Figure 4). This result confirmed the ability of our strategy for the identification of quantitative changes of PTMs from real gut microbiome samples.

**Figure 4.**
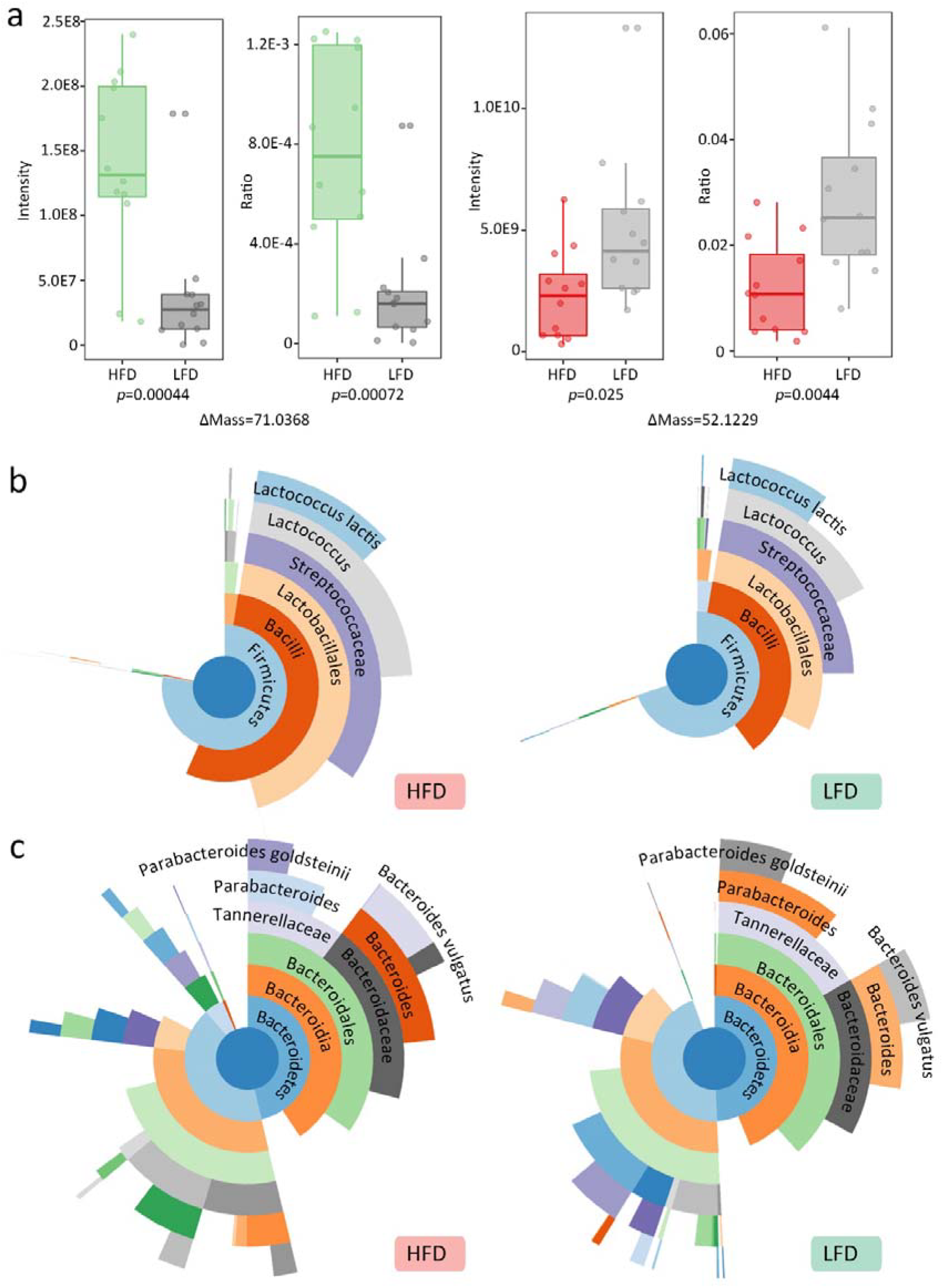
(a) The absolute and relative abundances of potential modifications ΔMass=71.0368 and 437 ΔMass=52.1229 identified in mouse gut microbiome. The bottom and top of the boxes are the first and third quartiles, respectively, the middle lines represent the sample median. (b-c) The relative abundance of each taxon determined by (b) acrylamide adduct and (c) cation:Fe[III] modified peptides in HFD and LFD samples.

Since there were two possibilities for the modification ∆Mass=71.0368, we investigated the information of localization sites to determine which was validate. Acrylamide adducts may occur on Cys, Lys and any amino acid on peptide N-term whereas Gly->Gln substitution only exists on Gly. Considering all the possible sites for localization analysis, we found Lys the most frequent modification site for ∆Mass=71.0368 (**Supplementary Fig. 6**), which suggested that the most probable modification was an acrylamide adduct. The relative abundance of modified peptides in each taxonomy in both HFD and LFD samples were illustrated respectively (Figure 4b). The compositions and relative abundances were almost constant in these two conditions. The lineage *Bacteria->Firmicutes->Bacilli->Lactobacillales->Streptococcaceae->Lactococcus->Lactococcus lactis* was the main source for acrylamide adduct modifications and contributed to the high abundance in HFD samples directly. At the protein level the difference was derived from the single site K264 from protein glyceraldehyde-3-phosphate dehydrogenase (GAPDH) in *Lactococcus lactis*. GAPDH is involved in the glucose metabolic process and related to the function NAD binding, NADP binding and GAPDH (NAD+) (phosphorylating) activity. Acrylamide adducts have been deemed as potentially toxic compounds in food^26^. The degradation of acrylamide by bacteria was widely studied^27, 28^ and *Bacillus cereus* has been reported to efficiently degrade acrylamide^29^. Molecular mechanisms of the addition of acrylamide adducts on GAPDH was studied by incubating purified human GAPDH with acrylamide and the kinetics of Cys acrylamide adduct formation was characterized, but the modification on Lys was not described^30^. This work presented the novel discovery of Lys acrylamide adducts on GAPDH from *Lactococcus lactis*, and the potential relationship with HFDs.

For the modification replacement of three protons by iron (cation:Fe[III] for short), the most abundant species were *Bacteroides vulgatus* and *Parabacteroides goldsteinii* from the same order *Bacteroidales* (Figure 4c). Significant change was only observed in species *Parabacteroides goldsteinii*. The involved proteins included TonB-dependent receptor, RagB/SusD family nutrient uptake outer membrane protein, SusC/RagA family TonB-linked outer membrane protein and hypothetical proteins. The first three were all membrane proteins and TonB-dependent receptor was a known iron transporter^31^. The identification of cation:Fe[III] modification on these proteins could help researchers further their understanding of the transport mechanism of bacteria.

### Identification of multiple types of glycosylation from human gut microbiome dataset

Finally, we used MetaLab 2.0 for the analysis of a human gut microbiome dataset from a study of type 1 diabetes^32^. Two groups of data from healthy control (CO, n=22) and new-onset patients (NO, n=33) were analyzed. From this dataset we identified 829,458 PSMs at FDR about 0.16% (**Supplementary Data 8**). Only 676,281 PSMs were obtained at the FDR about 4.1% by the conventional method (**Supplementary Fig. 7**). The MS2 spectra identification rate was increased from 35.8% to 44.0%, and while a much lower FDR was obtained. For each sample, 10,305 ± 2,205 (mean ± standard deviation) unique peptide sequences and 4,763 ± 1,027 proteins were quantified. In total 43,475 non-human microbiome proteins and 629 human proteins were identified. We quantified 611 species, among which 240 species were identified with more than three unique peptides. Annotations with KEGG database allowed us to identify 459 ± 69 microbial and 100 ± 27 human KEGG orthologies (KOs) per sample.

Here we identified 288 types of potential modifications and 102 were successfully matched to records in Unimod. Those known modifications covered 82% (78,931/97,102) of the modified PSMs. In general, human proteins had more PTMs than microbial proteins (**Supplementary Fig. 8**). PSMs from human proteins accounting for 17.7% (146,734/829,458) of the total number but contributed to 41.8% (40,608/97,102) of the PSMs with PTMs. The dynamic range of various modifications was high. The top 10 modifications covered 56.6% of the total number of modified PSMs identified in this study (**Supplementary Fig. 9**).

Currently the glycosylation in the microbial community is an understudied area. In this human fecal metaproteomic dataset, five types of glycosylation including Hex (hexose, ∆Mass = 162.053), HexNAc (N-Acetylhexosamine, ∆Mass = 203.079), HexNAc1dHex1 (∆Mass = 349.137), HexNAc2 (∆Mass = 406.159) and bacillosamine (2,4-diacetamido-2,4,6-trideoxyglucopyranose, ∆Mass = 228.1110) were identified. An important feature of the glycopeptide spectra were that the diagnostic ions could be observed in low mass-to-charge ratios (m/z), which were called glycan oxonium ions^33^. In our results, we detected at least one glycan oxonium ion whose intensity exceeded 50% of the peaks in the corresponding MS2 spectra, in 92.0% (5,424/5,895), 94.6% (5,750/6,078), 93.4% (157/168) and 69.5% (91/131) of the HexNAc, HexNAc1dHex1, HexNAc2, and bacillosamine modified PSMs, respectively (**Supplementary Table 3**). Manual verifications were also adopted for part of the glycopeptide spectra, and generally successive y-ions were observed in these spectra (**Supplementary material 5**). These results ensured the reliability of the identification of glycopeptides. Interestingly, we found the distributions of these glycosylations between human and microbial proteins differed (Figure 5a). Two types of glycosylation with apparently higher abundances, HexNAc1dHex1 and HexNAc were only observed on human proteins. In this dataset, decreases in the abundance of HexNAc modified peptides were detected in NO patients compared to CO subjects (*p*=0.048 by a two-sample, two-tailed t-test) (Figure 5b, **Supplementary Note**). For Hex and HexNAc2, there were 38.1% (45/118) and 28.6% (48/168) of the PSMs that were identified from microbial proteins, respectively. Bacillosamine was a rare monosaccharide which was found in bacterial glycoproteins^34^. And in our results 93.1% (122/131) of the bacillosamine modified PSMs were from bacteria proteins. Various types of glycosylation, as well as the corresponding information about taxa and functions can be identified in our strategy (**Supplementary Note**), which will greatly improve our understanding of gut microbiome-host interactions.

**Figure 5.**
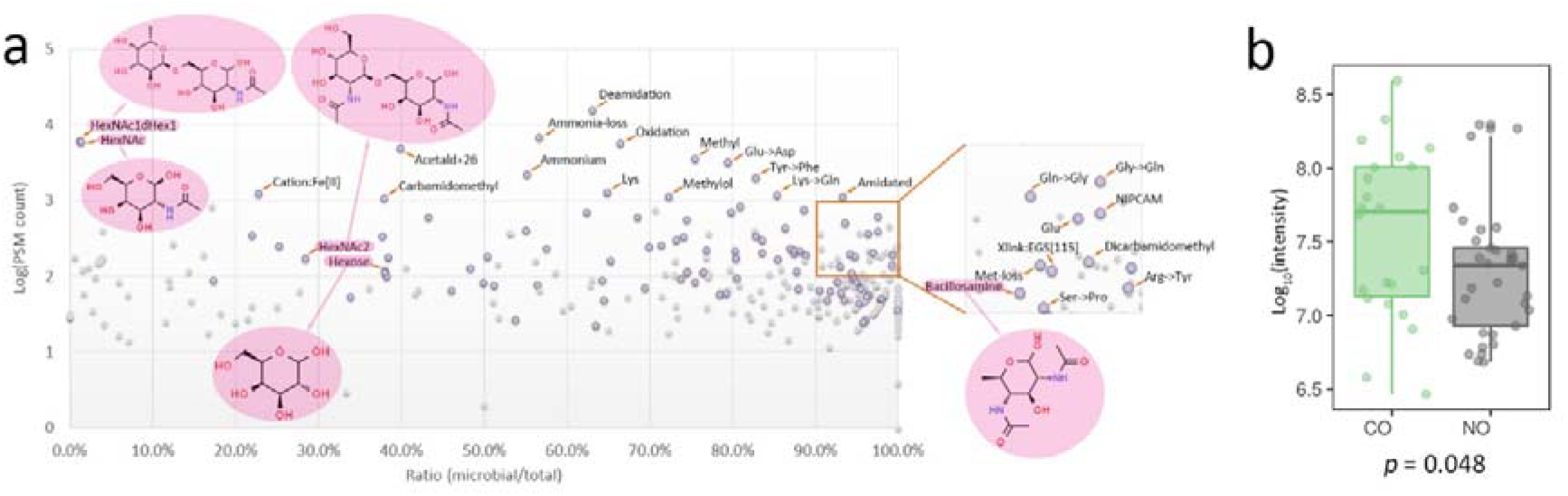
The PTMs identified from the human gut microbiome samples. **(a)** The identified potential modifications and their distribution between human and microbial proteins. Purple dots: modifications matched in Unimod; Grey dots: unknown modifications. The annotated modifications fulfilled one of the following two conditions: 1) with above 1,000 PSM counts; 2) with above 100 PSM counts and more than 90% were from microbial proteins. The possible structures of five glycosylation were illustrated. **(b)** HexNAc modified glycopeptides showed a significant difference between CO and NO samples. The bottom and top of the boxes are the first and third quartiles, respectively, the middle lines represent the sample median.

## Conclusion

In this work, we developed a metaproteomic data analysis tool MetaLab 2.0 which enabled PTM profiling from microbiome samples. Modifications of peptides bring mass variations which can increase the number of unidentified spectra in traditional database searches. The open search strategy is a method capable of identifying modifications in an unrestricted manner, however, the validation of the results remains challenging, especially for the studies of microbial communities. In this method, rigorous filters are utilized to remove potential modifications with less confidence. The analysis of the benchmark datasets (Homo_HEK293, Mock_micro, Mouse_gut and Human_gut) suggests that high accuracy is achieved without the sacrifice of sensitivity. It is worth noting that the identification rates are greatly improved in the analysis of complex mouse/human gut microbiota samples using the open search strategy. Combined with the previously developed metaproteomics data analysis tools by our group, a novel bioinformatic workflow is provided for the comprehensive analysis of PTM profiles from microbial communities. Our approach opens the door to better understand the roles that PTMs play in the biological processes of microorganisms and host-microbiome interaction. Rich and quantitative information can be obtained easily and with high accuracy, which is the major reason we recommend this strategy to be a standard workflow for metaproteomics studies.

## Materials and methods

### Detailed information of the open search workflow in MetaLab 2.0

The first step is the database search by the open search strategy. Only mass spectrometer raw files and searched protein databases were required as the input. The input raw files were converted to mzXML and MGF format for the post-analysis. The converted mzXML files were subject to MSFragger for the peptide identification. A target-decoy database was utilized for the FDR estimation. After the identification results were obtained, all the PSMs were binned according to the determined ∆Mass. Through the ∆Mass range, the Gaussian peaks with R^2^ above 0.9 were detected (users can change this parameter in our software). PSMs will be discarded if they don’t belong to any ∆Mass Gaussian peak. The centroids of the fitted Gaussian peaks were determined. For each PSM, the mass of their potential modification was set as the mass of the calculated centroid, instead of the mass deviation determined directly from the open search result. If the relative mass difference between the original mass deviation and the calculated centroid was above 10 ppm, this PSM was also discarded. After the mass of the modification of each PSM was determined, deisotoping was performed. The theory mass difference between two isotope peaks was set as 1.00286864 Da^35^. When we found two Gaussian peaks, Peak_i_ (∆Mass=M_i_) and Peak_j_ (∆Mass=M_j_), which fulfilled the conditions that |M_j_-M_i_-1.00286864|≤0.01, and at the same time Peak_i_ had more PSMs than Peak_j_, Peak_j_ was deemed as an isotope peak of Peak_i_. The mass of modifications of PSMs in Peak_j_ was set as M_i_ in this step.

Then a semi-supervised machine learning strategy was performed (**Supplementary Table 1**). The 10% top-scored PSMs were randomly distributed into five datasets, and all the decoy PSMs were appended to each of the datasets for the construction of training datasets. Five naive Bayes classifiers were built based on the training datasets and used to calculate the probability for each PSM to be a target/decoy one. A classification score was assigned for each PSM, which was calculated by the average of the probabilities to be a target match determined by the five classifiers. Then the PSMs were sorted by the newly assigned classification score, and the top 10% were selected for another loop of the classification. After five iterations, the average of the probabilities was determined as the final classification score.

The next step was filtering PSMs at the modification level. Unpaired t-tests were performed between the classification scores of target and decoy PSMs. We also created a universal decoy sample that consisted of the classification scores of all the decoy PSMs. For each of the modifications, if there were significant differences (p-value<0.01) between the classification scores of target PSMs and decoy PSMs, and at the same time, between the target PSMs and the universal decoy PSMs, this modification was deemed as validated. Modifications which did not satisfy any of the two conditions were filtered out.

The final step of the multi-stage strategy was performing filtering at all of the three levels: protein, peptide and PSM. A protein inference algorithm was adopted in this step^19^. Then according to the determined protein and peptide scores, the FDRs were controlled at less than 1% for all of the three levels. Usually the protein level FDR was about 1% and the other levels much less than 1%.

According to the filtered PSMs, MS1 peak intensity based label-free quantitation was adopted^16^. Then the qualitative and quantitative information of PSMs, peptides, proteins and modifications were exported. For conventional proteomics studies, the post-analysis workflow ended here. For metaproteomic studies, taxonomy analysis and functional annotation were then performed consecutively.

### MS/MS datasets

The information of the analyzed benchmark dataset was described in **Supplementary Table 4**. For each dataset refer to the corresponding supplementary data for the detailed information about the raw files used, the identified potential modifications, PSMs, peptides and proteins. Some examples of manually annotated spectra of PSMs with most frequently identified modifications (usually the top 30) were provided for four datasets, including *Ecoli*_phos (**Supplementary Material 1**), Mouse_phos (**Supplementary Material 2**), Hela_phos (**Supplementary Material 3**), U87_phos (**Supplementary Material 4**). Examples of manually annotated spectra of glycopeptides were provided for Human_gut dataset (**Supplementary Material 5**).

### Database search

All the open search in this work were used in the same parameter setting. Carboxyamidomethylation on C was set as a fixed modification. Although an open search can identify modifications unrestrictedly, considering that oxidation on Met and acetylation on protein N-term commonly exist, these two were still set as variable modification. Oxidation and acetylation that occur on other amino acids would be identified by the open search. The precursor mass tolerance was set as 500 Da and the fragment mass tolerance was set as 10 ppm. The enzyme is trypsin (KR/P) and digestion mode is fully enzymatic digestion. The number of missed cleavage sites is 2. In the analysis of Homo_HEK293 samples, as a comparison, a closed search by MSFragger was performed. The same parameter setting was utilized except the precursor mass tolerance is ±10 PPM. In the result validation step, if the multistage filtering strategy was not used, we applied the protein inference algorithm^19^ directly based on the search results. Then the results were refined to keep the FDRs on both of the protein, peptide and PSM levels less than 1%. The closed search results of the Mouse_gut and Human_gut datasets were from the paper directly but not a re-analysis result by MSFragger.

### Data availability

All the mass spectrometry data are available at the PRIDE Archive (Homo_HEK293: https://www.ebi.ac.uk/pride/archive/projects/PXD001468; Ecoli_phos: https://www.ebi.ac.uk/pride/archive/projects/PXD008289; Mouse_phos: https://www.ebi.ac.uk/pride/archive/projects/PXD001792; Hela_phos: https://www.ebi.ac.uk/pride/archive/projects/PXD004940; U87_phos: https://www.ebi.ac.uk/pride/archive/projects/PXD009227; Mock_micro: https://www.ebi.ac.uk/pride/archive/projects/PXD006118; Mouse_gut: https://www.ebi.ac.uk/pride/archive/projects/PXD003527; Human_gut: https://www.ebi.ac.uk/pride/archive/projects/PXD008870).

### Software availability

MetaLab 2.0 is free for academic use and has a user-friendly Graphic User Interface (GUI). Both online and local versions of the tool are available at https://imetalab.ca/. The source code of MetaLab 2.0 is available at https://bitbucket.org/cksakura/metalab/src/master/.

## Acknowledgement

This work was supported by the Government of Canada through Genome Canada and the Ontario Genomics Institute (OGI-114), CIHR grant (ECD-144627), the Natural Sciences and Engineering Research Council of Canada (NSERC, grant no. 210034), the Ontario Ministry of Economic Development and Innovation (REG1-4450) and The University of Ottawa.

## Authors’ contributions

KC, DF, ZN, XZ and JM designed the study. KC wrote the software and performed data analysis. ZN, BL and LL also contributed to the software. KC, ZN, XZ, LL and BL tested the software. KC and DF wrote the manuscript, and all the authors contributed to the editing and revision of the manuscript. All authors read and approved the final manuscript.

