## Supplementary material 1 for "MetaLab 2.0 enables accurate post-translational modifications profiling in metaproteomics"

### Supplementary material 1: Manually annotated spectra of the top 30 identified potential modifications

Dataset: Ecoli_phos

# 1

- Experimental mass: 79.9666
- Unimod information: phosphorylation
- Possible composition: H O(3) P
- Sites in Unimod: S;T;Y;D;H;C;R;K
- Sites observed: S;T;Y;H

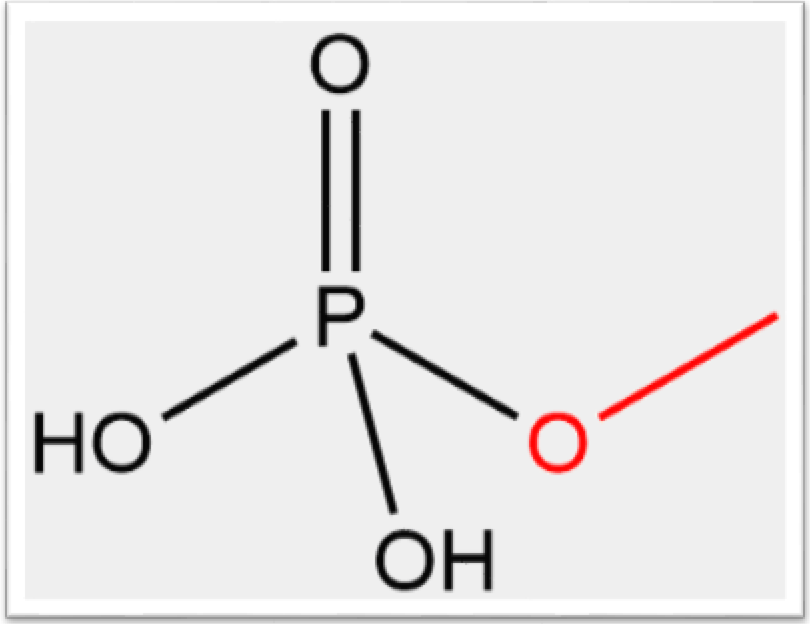

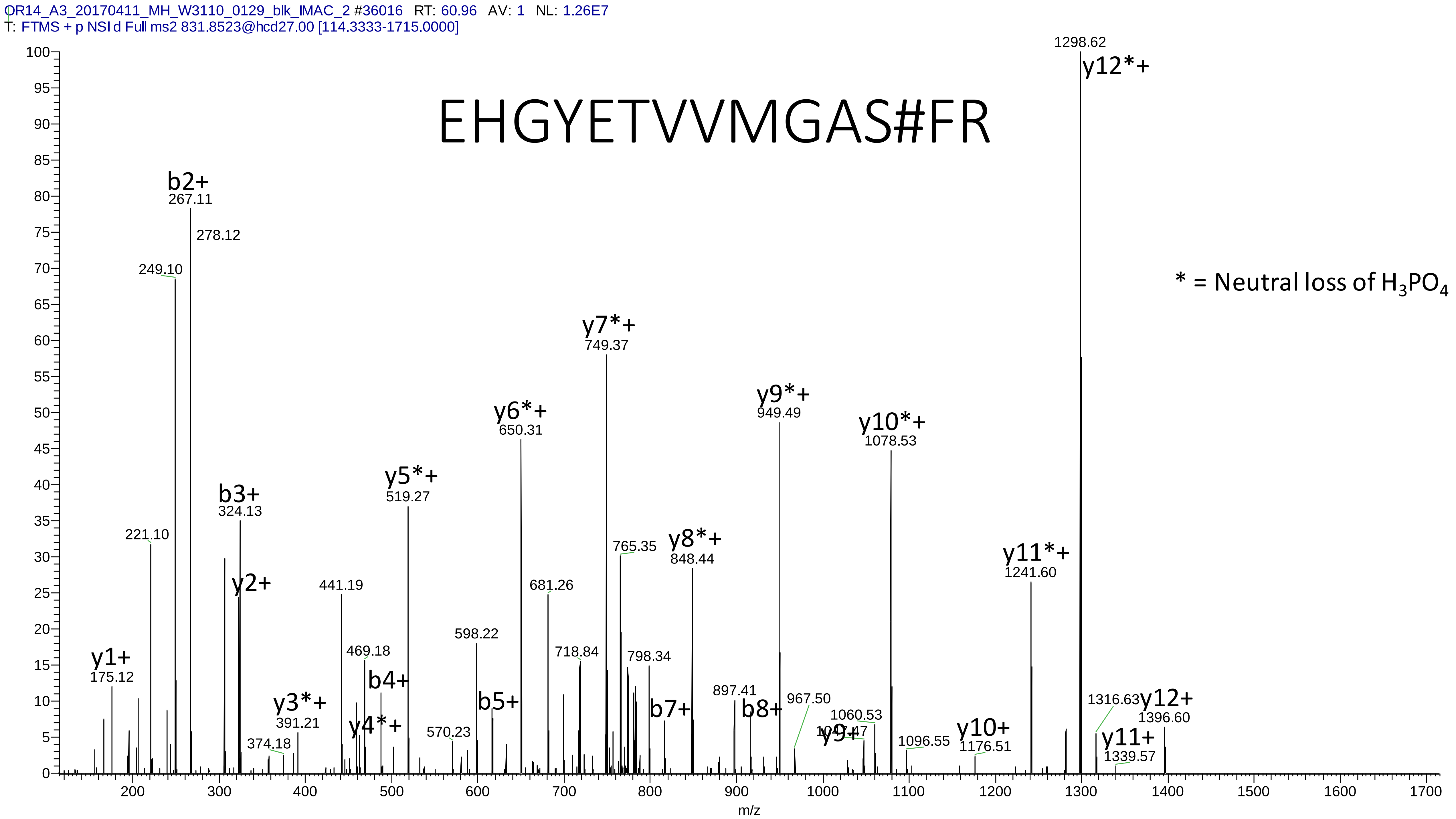

OR14_A3_20170411_MH_W3110_0129_blk_IMAC_2 #36016 RT: 60.96 AV: 1 NL: 1.26E7
T: FTMS + p NSI d Full ms2 831.8523@hcd27.00 [114.3333-1715.0000]
EHGYETVVMGAS#FR
* = Neutral loss of H3PO4
y1+ 175.12
221.10
249.10
b2+ 267.11
278.12
b3+ 324.13
y2+
374.18
y3*+ 391.21
441.19
469.18
y4*+
b4+
y5*+ 519.27
570.23
598.22
b5+
y6*+ 650.31
681.26
718.84
y7*+ 749.37
765.35
798.34
b7+
y8*+ 848.44
897.41
b8+
y9*+ 949.49
967.50
1047.47
y9+
1060.53
y10*+ 1078.53
1096.55
y10+ 1176.51
y11*+ 1241.60
1298.62
y12*+
1316.63
y11+ 1339.57
y12+ 1396.60
m/z

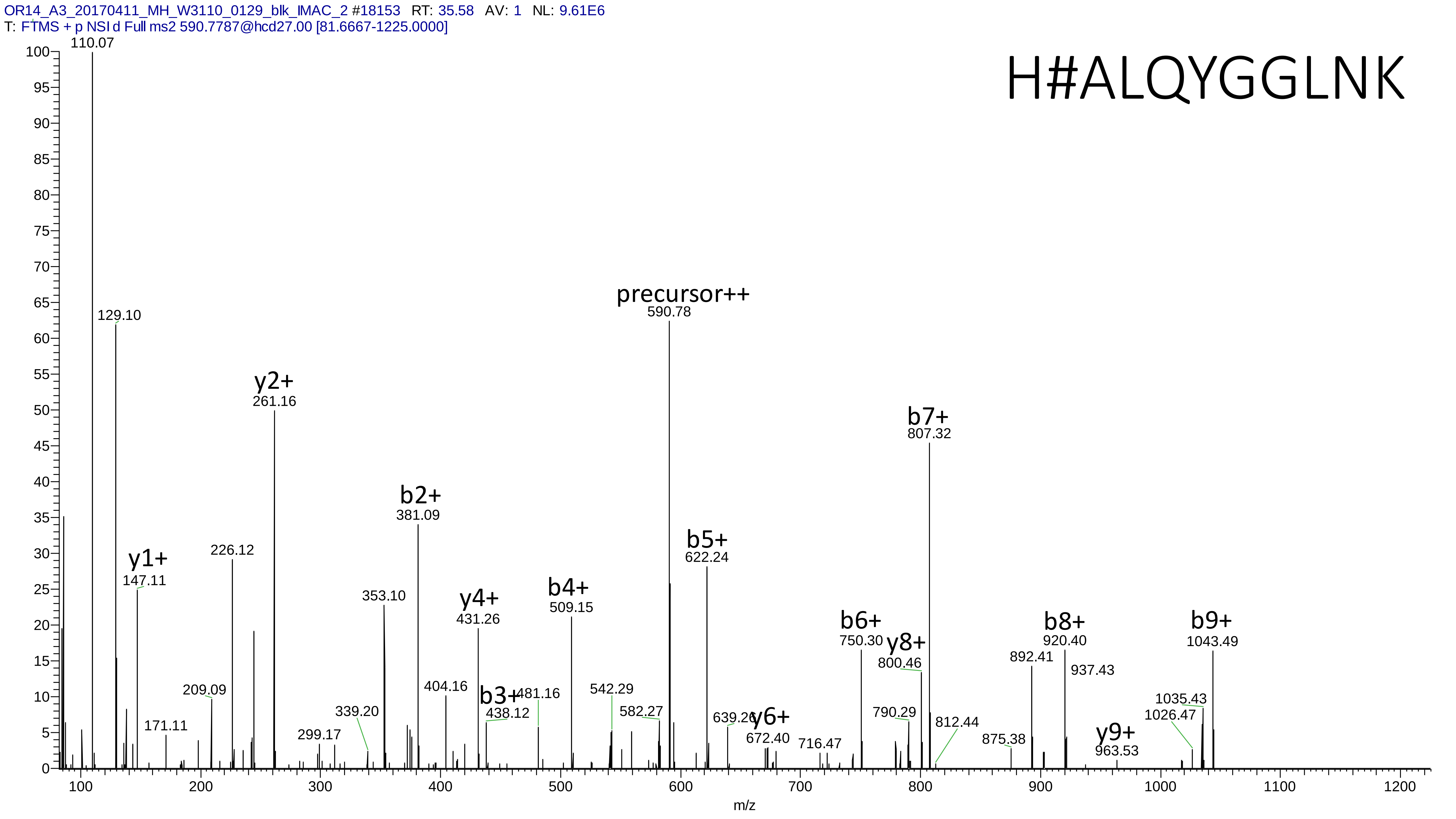
OR14_A3_20170411_MH_W3110_0129_blk_IMAC_2 #18153 RT: 35.58 AV: 1 NL: 9.61E6
T: FTMS + p NSI d Full ms2 590.7787@hcd27.00 [81.6667-1225.0000]
H#ALQYGGLNK
precursor++
590.78
y2+
261.16
b2+
381.09
b7+
807.32
b5+
622.24
y1+
147.11
226.12
353.10
y4+
431.26
b4+
509.15
b6+
750.30
y8+
800.46
b8+
920.40
b9+
1043.49
110.07
129.10
209.09
171.11
299.17
339.20
404.16
b3+
438.12
481.16
542.29
582.27
639.26
y6+
672.40
716.47
790.29
812.44
875.38
892.41
937.43
y9+
963.53
1035.43
1026.47
m/z

# 2

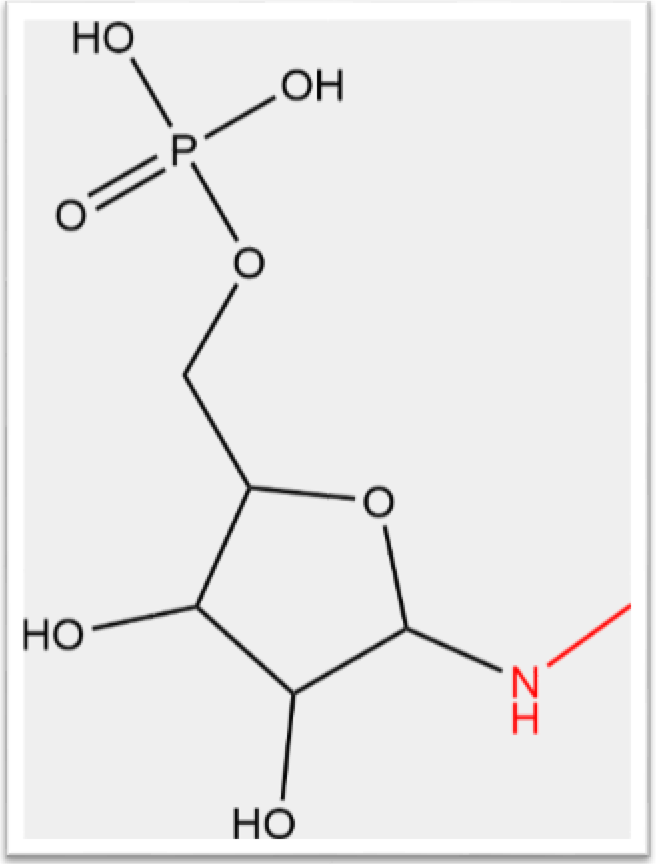


- Experimental mass: 212.0091
- Unimod information: phosphate-ribosylation
- Possible composition: H(9) C(5) O(7) P
- Sites in Unimod: D;E;R
- Sites observed: K; protein n-term
- Features: 86.3% (864/1001) peptides contain a miss cleavage site K, 7.4% (74/1001) peptides start with the first amino acid in the proteins, 8.9% (89/1001) peptides start with the second amino acid in the proteins, in total 16.3% (163/1001) peptides contains protein n-term; neutral loss of $H_3PO_4$ are observed at both precursor and fragment ions

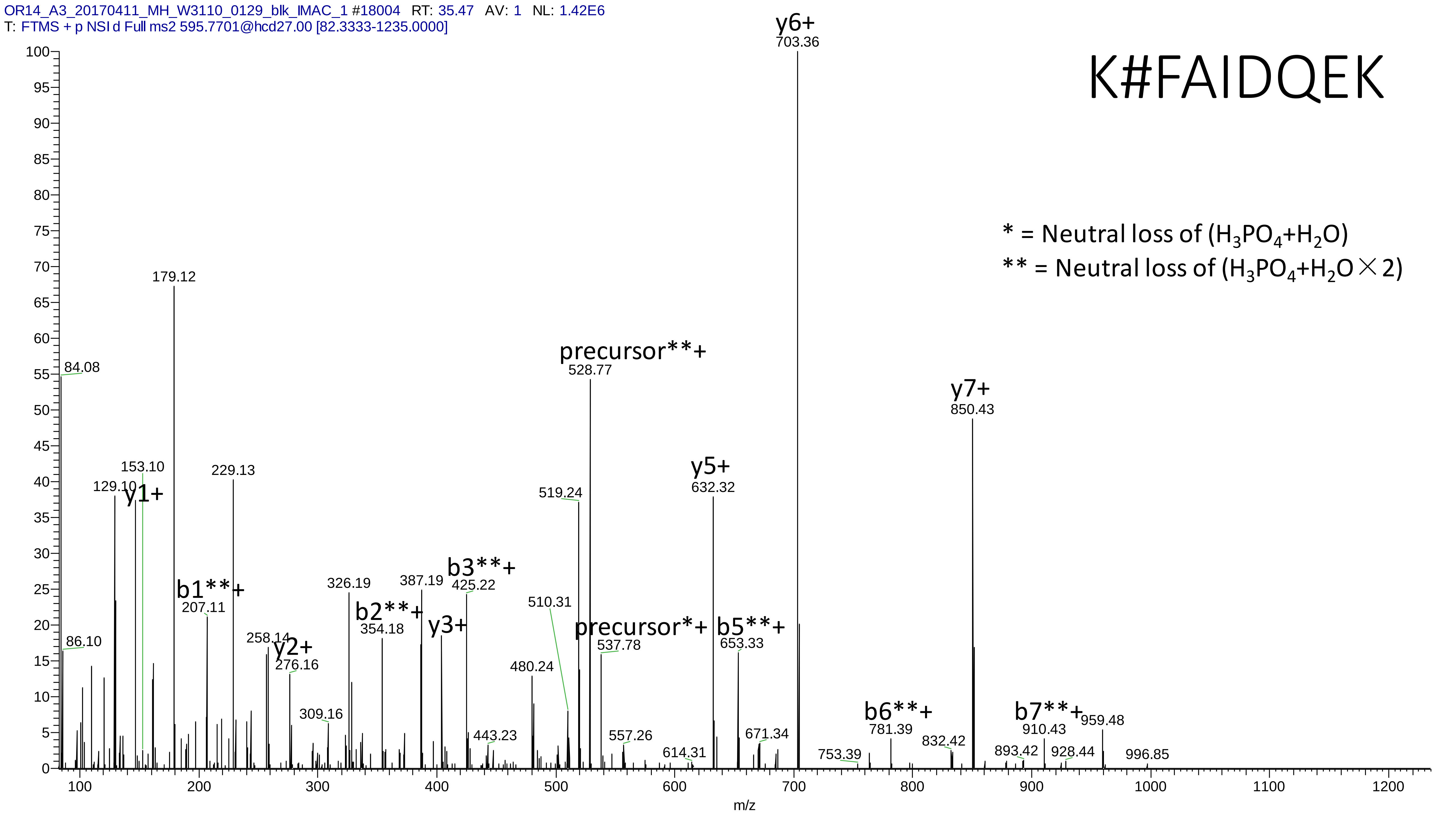
OR14_A3_20170411_MH_W3110_0129_blk_IMAC_1 #18004 RT: 35.47 AV: 1 NL: 1.42E6
T: FTMS + p NSI d Full ms2 595.7701@hcd27.00 [82.3333-1235.0000]
K#FAIDQEK
* = Neutral loss of (H3PO4+H2O)
** = Neutral loss of (H3PO4+H2O×2)
y6+
703.36
179.12
precursor**+
528.77
84.08
y7+
850.43
153.10
129.10
y1+
229.13
y5+
632.32
519.24
b1**+
207.11
326.19
387.19
b3**+
425.22
b2**+
354.18
y3+
510.31
precursor*+
537.78
b5**+
653.33
86.10
258.14
y2+
276.16
480.24
309.16
443.23
557.26
614.31
671.34
753.39
b6**+
781.39
832.42
b7**+
910.43
959.48
893.42
928.44
996.85
m/z

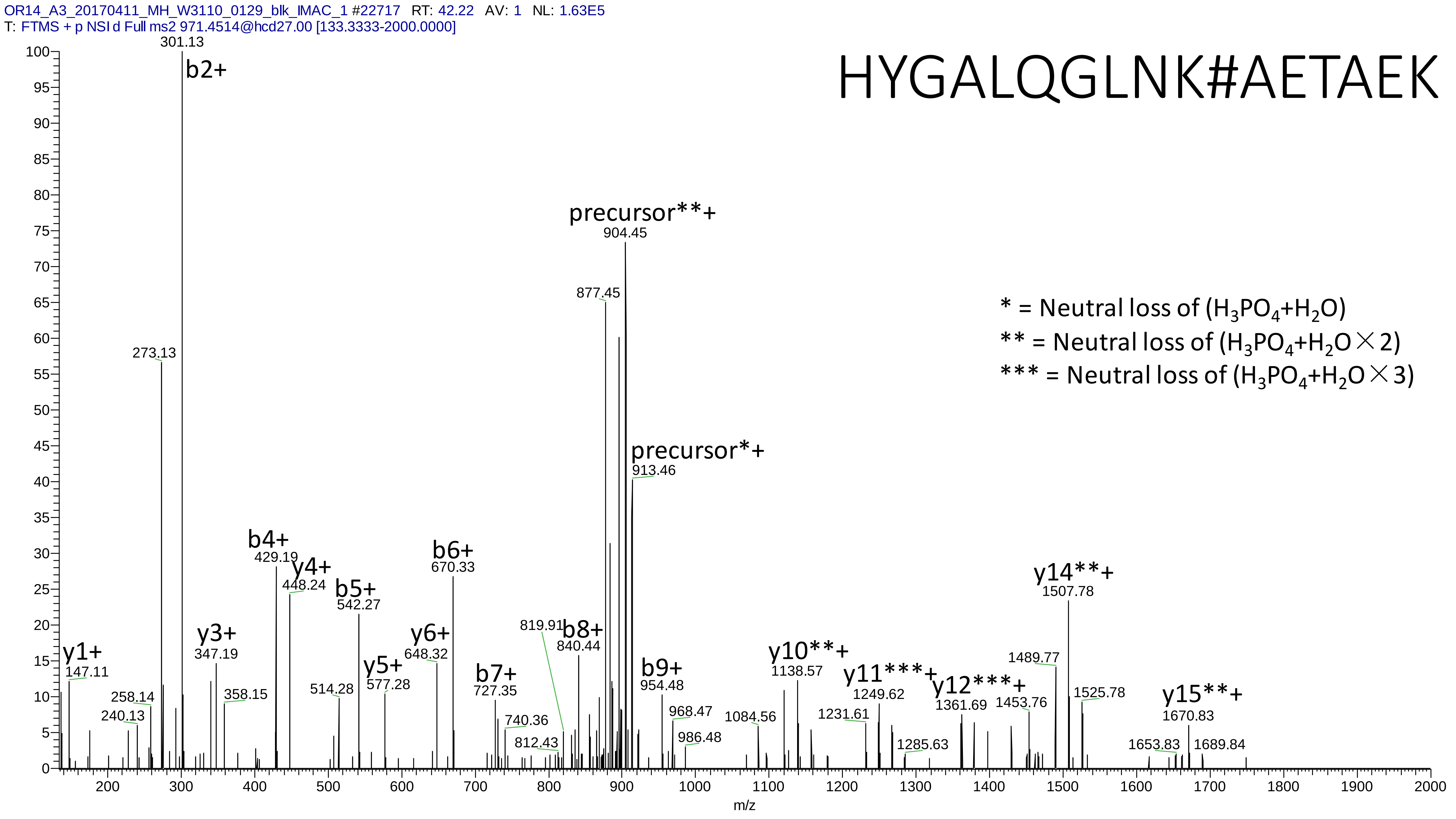
OR14_A3_20170411_MH_W3110_0129_blk_IMAC_1 #22717 RT: 42.22 AV: 1 NL: 1.63E5
T: FTMS + p NSI d Full ms2 971.4514@hcd27.00 [133.3333-2000.0000]
HYGALQGLNK#AETAEK
* = Neutral loss of (H3PO4+H2O)
** = Neutral loss of (H3PO4+H2O×2)
*** = Neutral loss of (H3PO4+H2O×3)
b2+ 301.13
273.13
precursor**+ 904.45
877.45
precursor*+ 913.46
y1+ 147.11
240.13
258.14
y3+ 347.19
358.15
b4+ 429.19
y4+ 448.24
514.28
b5+ 542.27
y5+ 577.28
y6+ 648.32
b6+ 670.33
b7+ 727.35
740.36
812.43
819.91
b8+ 840.44
b9+ 954.48
968.47
986.48
1084.56
y10**+ 1138.57
1231.61
y11***+ 1249.62
1285.63
y12***+ 1361.69
1453.76
1489.77
y14**+ 1507.78
1525.78
1653.83
y15**+ 1670.83
1689.84
m/z

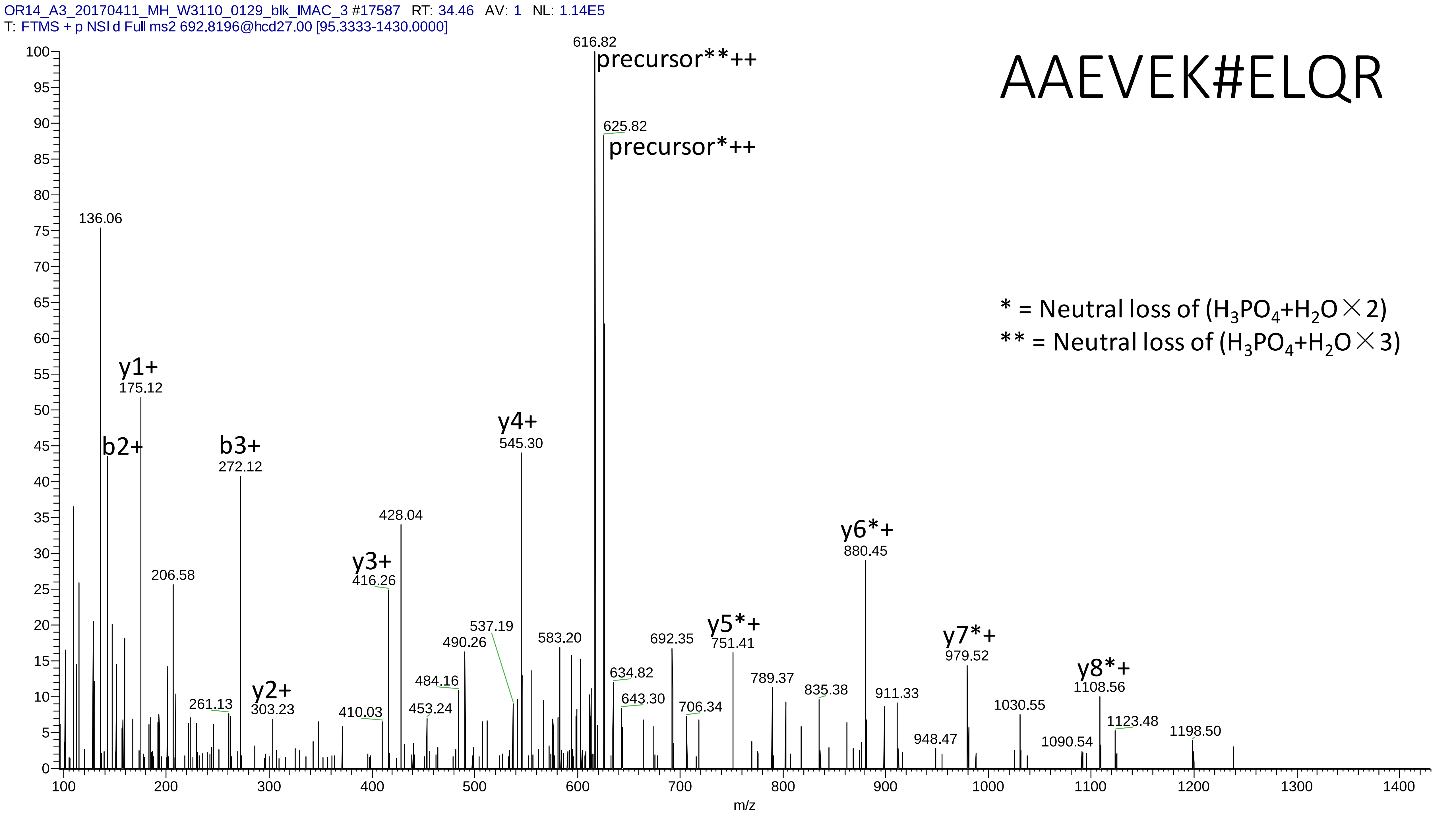
OR14_A3_20170411_MH_W3110_0129_blk_IMAC_3 #17587 RT: 34.46 AV: 1 NL: 1.14E5
T: FTMS + p NSI d Full ms2 692.8196@hcd27.00 [95.3333-1430.0000]
AAEVEK#ELQR
* = Neutral loss of (H3PO4+H2O×2)
** = Neutral loss of (H3PO4+H2O×3)
precursor**++
616.82
precursor*++
625.82
136.06
y1+
175.12
b2+
b3+
272.12
206.58
261.13
y2+
303.23
410.03
y3+
416.26
428.04
453.24
484.16
490.26
537.19
y4+
545.30
583.20
634.82
643.30
692.35
706.34
y5*+
751.41
789.37
835.38
y6*+
880.45
911.33
948.47
y7*+
979.52
1030.55
1090.54
y8*+
1108.56
1123.48
1198.50
m/z

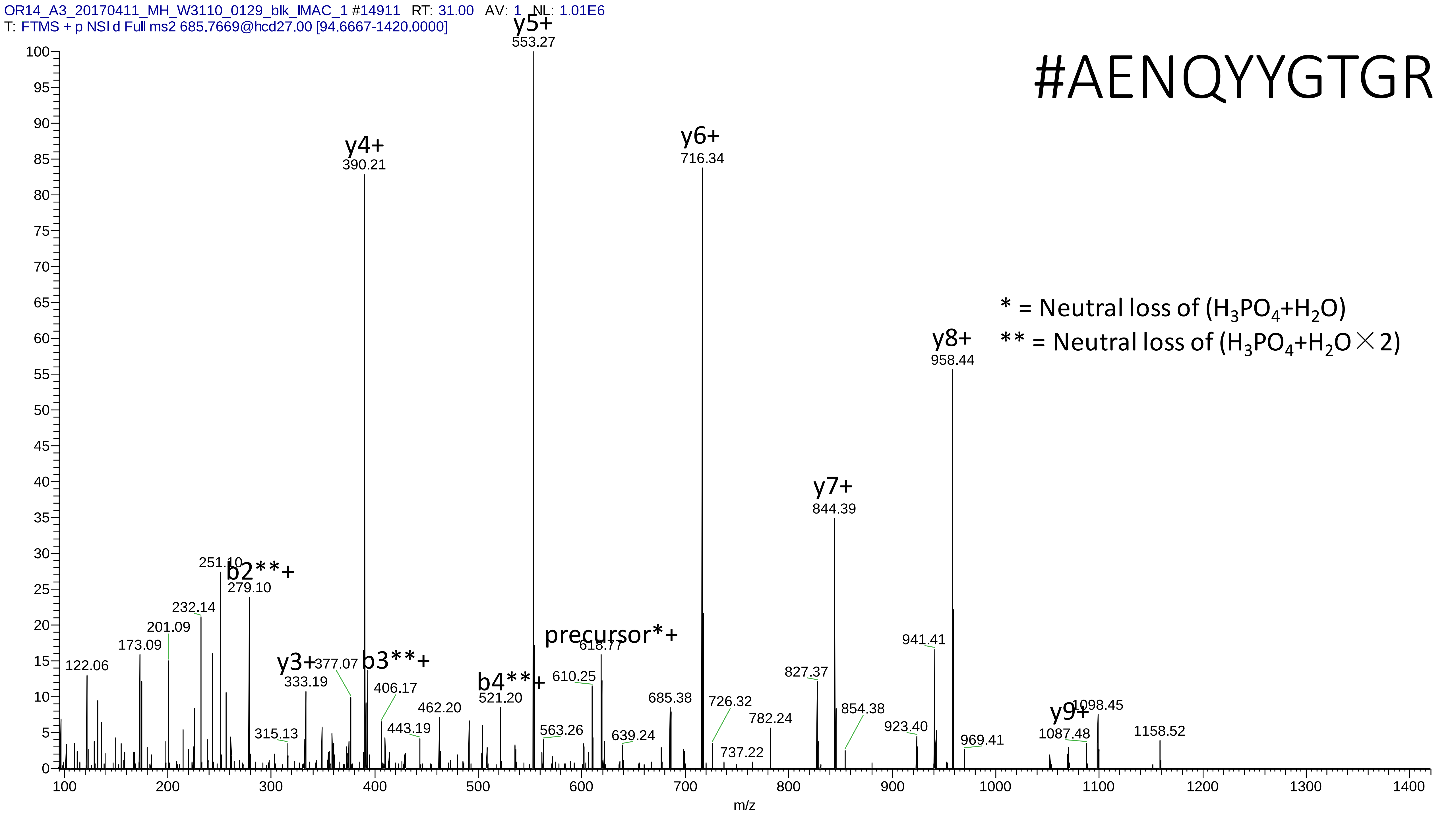
OR14_A3_20170411_MH_W3110_0129_blk_IMAC_1 #14911 RT: 31.00 AV: 1 NL: 1.01E6
T: FTMS + p NSI d Full ms2 685.7669@hcd27.00 [94.6667-1420.0000]
#AENQYYGTGR
* = Neutral loss of ($H_3PO_4$+$H_2O$)
** = Neutral loss of ($H_3PO_4$+$H_2O$×2)
y5+ 553.27
y4+ 390.21
y6+ 716.34
y8+ 958.44
y7+ 844.39
b2**+ 279.10
251.10
232.14
201.09
173.09
122.06
y3+ 333.19
377.07
b3**+
406.17
315.13
443.19
462.20
b4**+ 521.20
precursor*+ 618.77
610.25
563.26
639.24
685.38
726.32
737.22
782.24
827.37
854.38
941.41
923.40
969.41
y9+ 1087.48
1098.45
1158.52
m/z

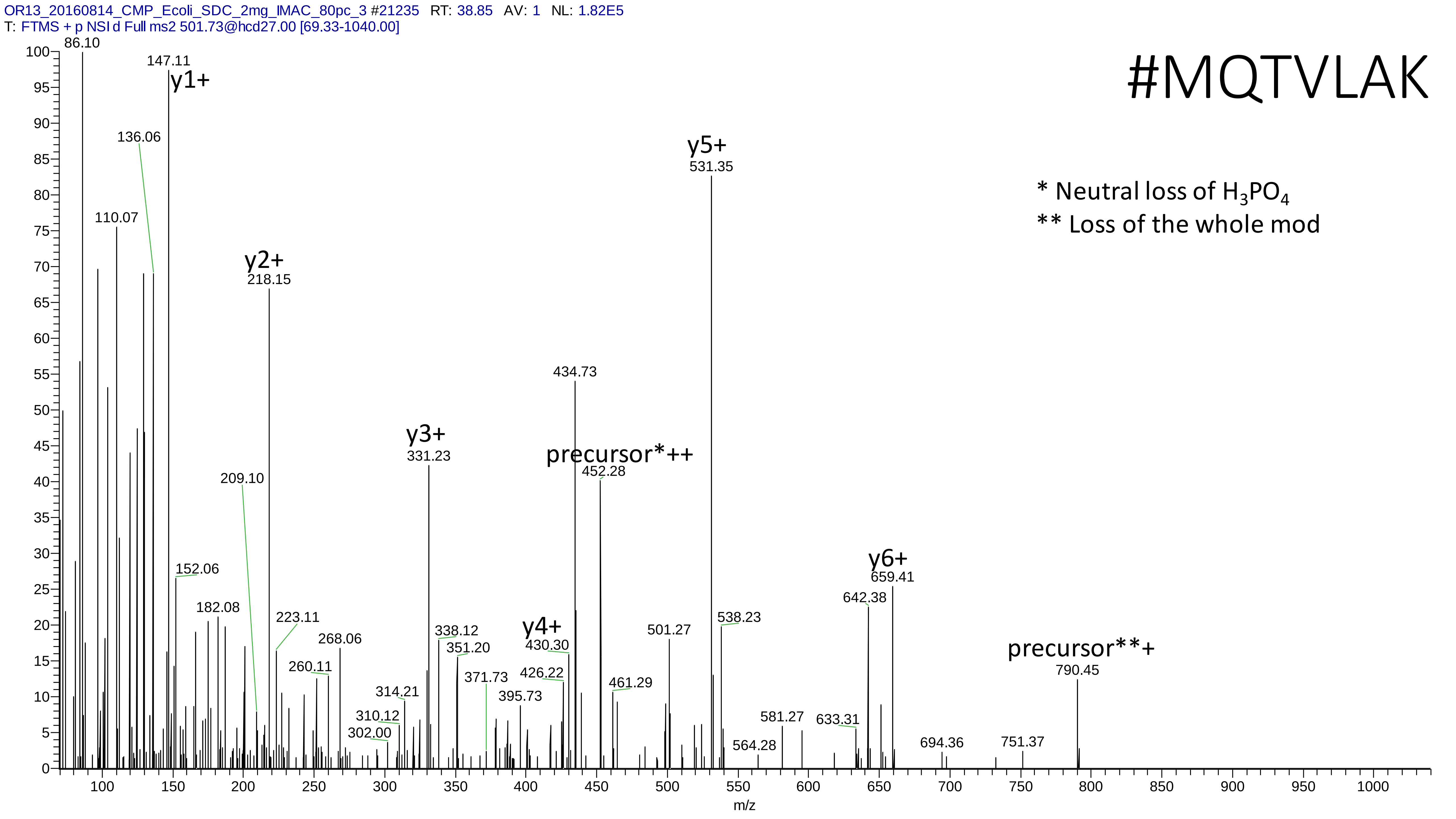
OR13_20160814_CMP_Ecoli_SDC_2mg_IMAC_80pc_3 #21235 RT: 38.85 AV: 1 NL: 1.82E5
T: FTMS + p NSI d Full ms2 501.73@hcd27.00 [69.33-1040.00]
#MQTVLAK
* Neutral loss of H3PO4
** Loss of the whole mod
86.10
147.11
y1+
136.06
110.07
y2+
218.15
209.10
152.06
182.08
223.11
268.06
260.11
y3+
331.23
338.12
351.20
371.73
314.21
310.12
302.00
395.73
y4+
430.30
426.22
434.73
precursor*++
452.28
461.29
501.27
y5+
531.35
538.23
564.28
581.27
633.31
642.38
y6+
659.41
694.36
751.37
precursor**+
790.45
m/z

# 3

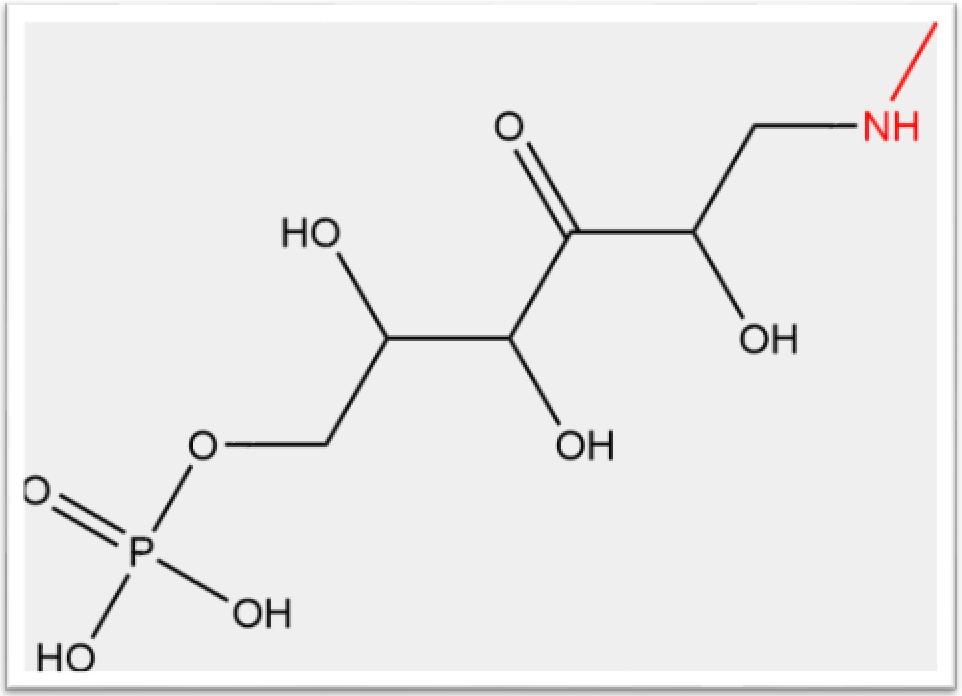


- Experimental mass: 242.0195
- Unimod information: PhosphoHex
- Possible composition: H O(3) P Hex
- Sites in Unimod: S;T
- Sites observed: K;N-term
- Features: 97.6% (495/507) peptides contain a miss cleavage site K; 2.4% (12/507) peptides start with the first amino acid in the proteins, 3.2% (16/507) peptides start with the second amino acid in the proteins, in total 5.5% (28/507) peptides contains protein n-term; neutral loss of $H_3PO_4$ are observed at both precursor and fragment ions

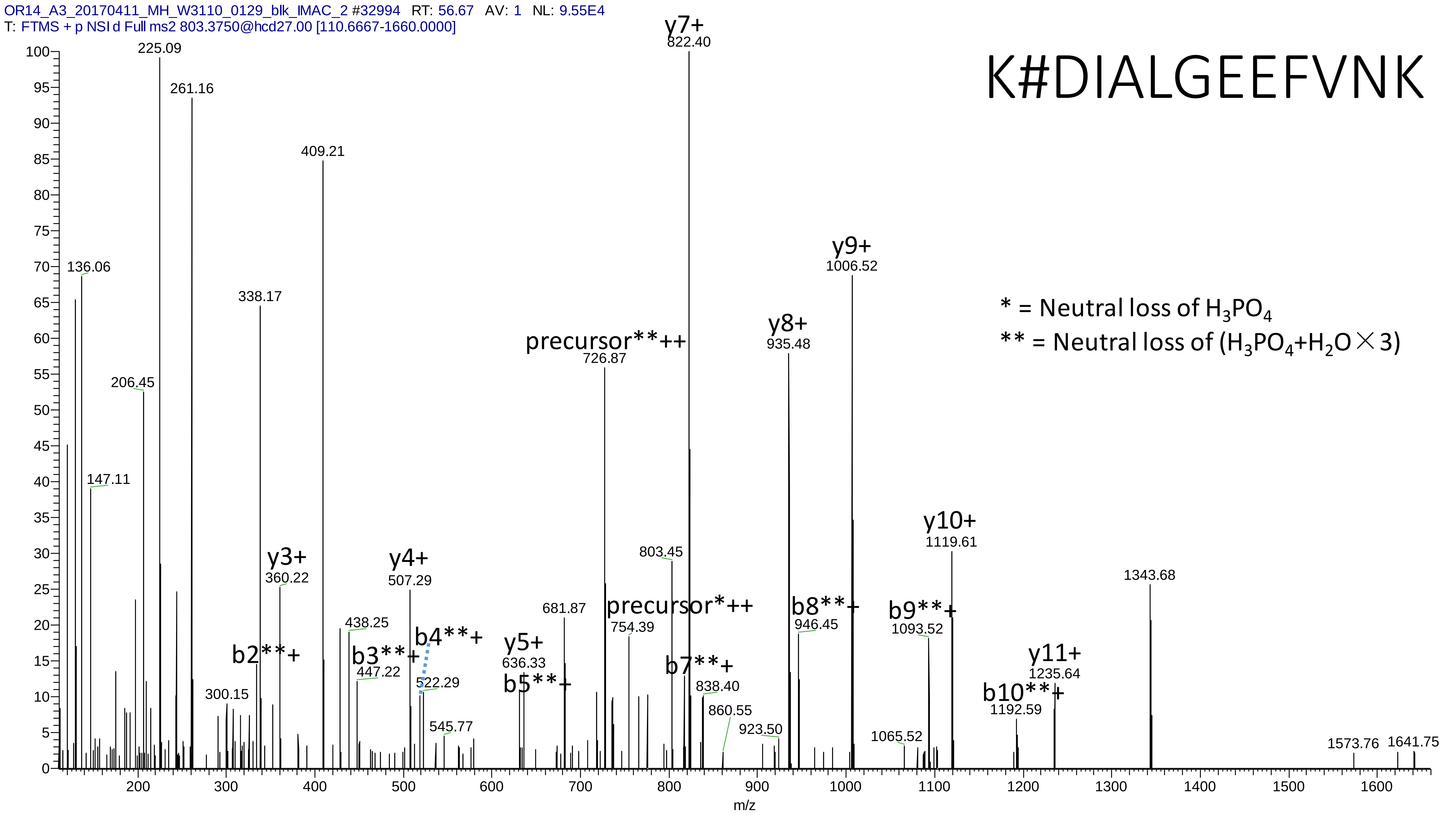

OR14_A3_20170411_MH_W3110_0129_blk_IMAC_2 #32994 RT: 56.67 AV: 1 NL: 9.55E4
T: FTMS + p NSI d Full ms2 803.3750@hcd27.00 [110.6667-1660.0000]
K#DIALGEEFVNK
* = Neutral loss of $H_3PO_4$
** = Neutral loss of ($H_3PO_4$+$H_2O$×3)
y7+
822.40
225.09
261.16
409.21
136.06
338.17
206.45
147.11
y9+
1006.52
y8+
935.48
precursor**++
726.87
y3+
360.22
y4+
507.29
803.45
y10+
1119.61
1343.68
precursor*++
754.39
b8**+
946.45
b9**+
1093.52
438.25
b4**+
b2**+
b3**+
447.22
522.29
y5+
636.33
b5**+
681.87
300.15
b7**+
838.40
y11+
1235.64
b10**+
1192.59
860.55
545.77
923.50
1065.52
1573.76
1641.75
m/z

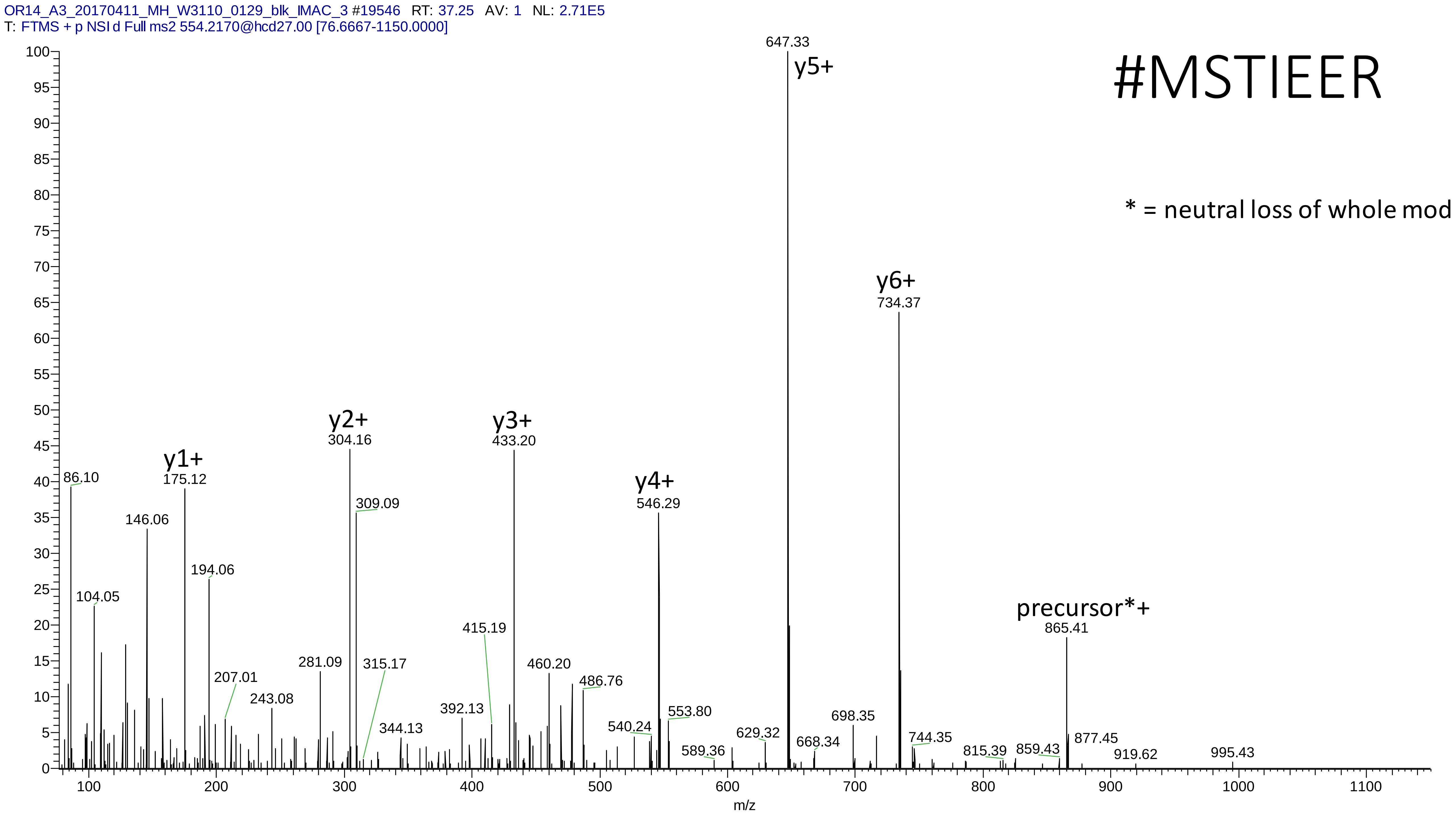
OR14_A3_20170411_MH_W3110_0129_blk_IMAC_3 #19546 RT: 37.25 AV: 1 NL: 2.71E5
T: FTMS + p NSI d Full ms2 554.2170@hcd27.00 [76.6667-1150.0000]
#MSTIEER
* = neutral loss of whole mod
y1+
y2+
y3+
y4+
y5+
y6+
precursor*+
86.10
104.05
146.06
175.12
194.06
207.01
243.08
281.09
304.16
309.09
315.17
344.13
392.13
415.19
433.20
460.20
486.76
540.24
546.29
553.80
589.36
629.32
647.33
668.34
698.35
734.37
744.35
815.39
859.43
865.41
877.45
919.62
995.43
m/z

# 4

- Experimental mass: 167.983
- Unimod information: 3-phosphoglyceryl
- Possible composition: H(5) C(3) O(6) P
- Sites in Unimod: K
- Sites observed: K;N-term
- Features: 89.1% (351/394) peptides contain a miss cleavage site K; 4.3% (17/394) peptides start with the first amino acid in the proteins, 2.5% (10/394) peptides start with the second amino acid in the proteins, in total 6.8% (27/408) peptides contains protein n-term; neutral loss of $H_3PO_4$ are observed at precursor ions

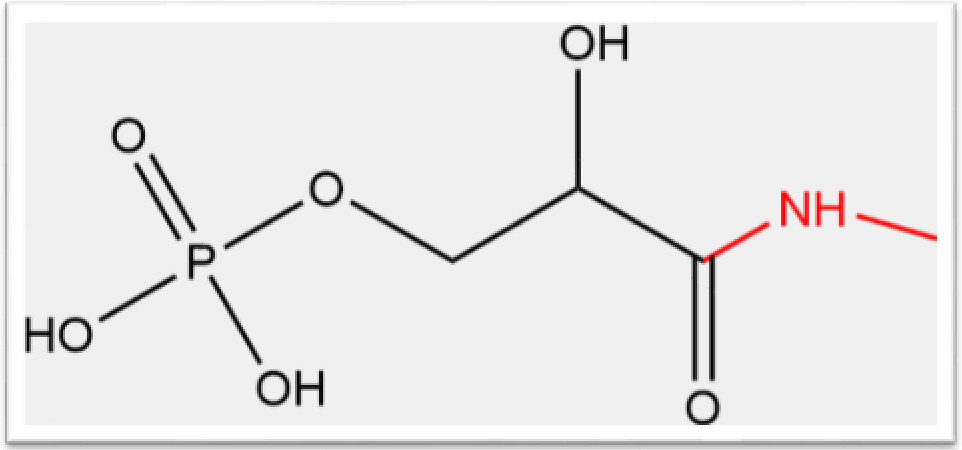

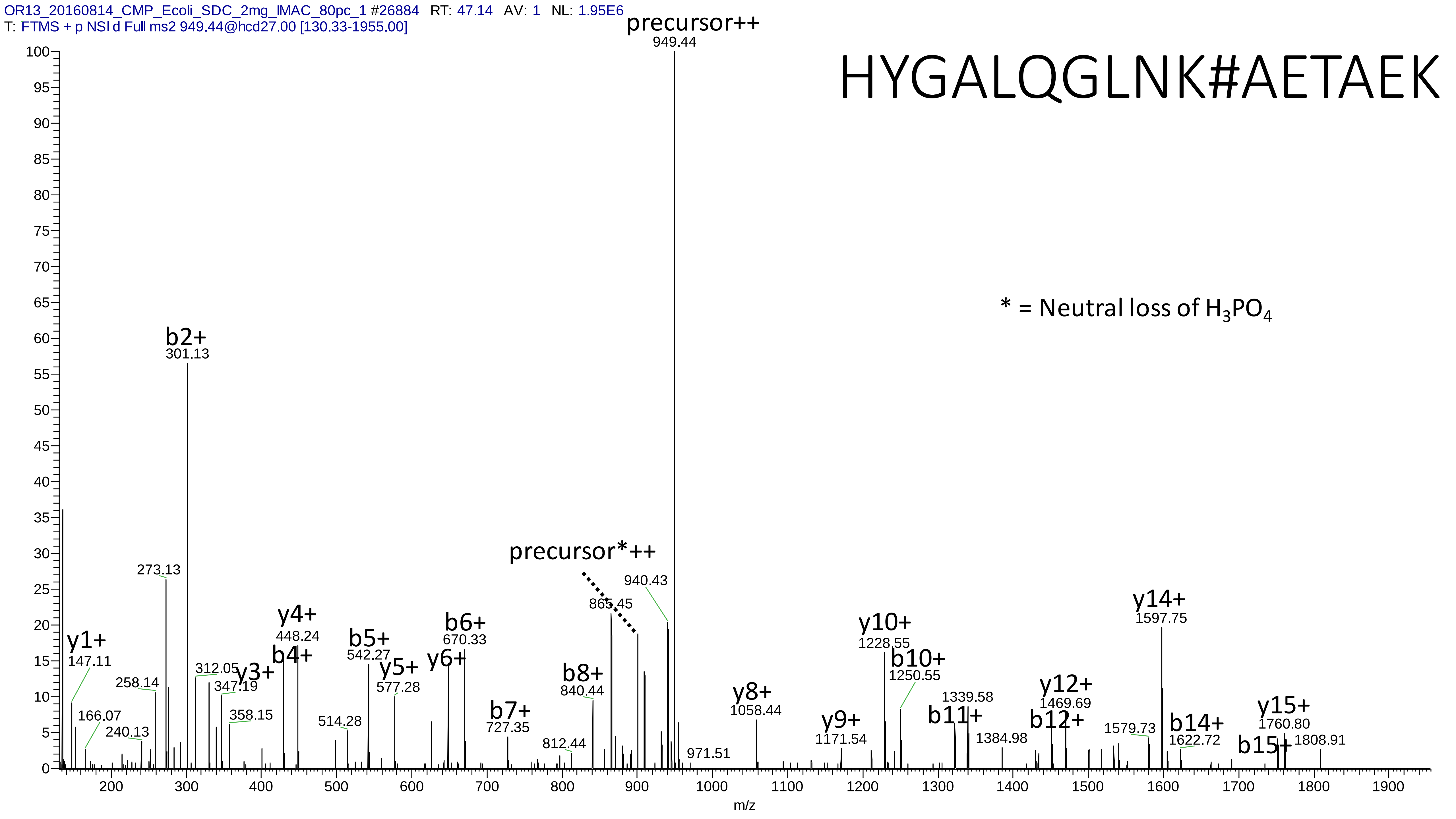
OR13_20160814_CMP_Ecoli_SDC_2mg_IMAC_80pc_1 #26884 RT: 47.14 AV: 1 NL: 1.95E6
T: FTMS + p NSI d Full ms2 949.44@hcd27.00 [130.33-1955.00]
HYGALQGLNK#AETAEK
* = Neutral loss of $H_3PO_4$
precursor++
949.44
precursor*++
b2+
301.13
273.13
y1+
147.11
166.07
240.13
258.14
312.05
y3+
347.19
358.15
y4+
448.24
b4+
b5+
542.27
514.28
y5+
577.28
y6+
b6+
670.33
b7+
727.35
812.44
b8+
840.44
865.45
940.43
971.51
y8+
1058.44
y9+
1171.54
y10+
1228.55
b10+
1250.55
1339.58
b11+
1384.98
y12+
1469.69
b12+
1579.73
y14+
1597.75
b14+
1622.72
b15+
y15+
1760.80
1808.91
m/z

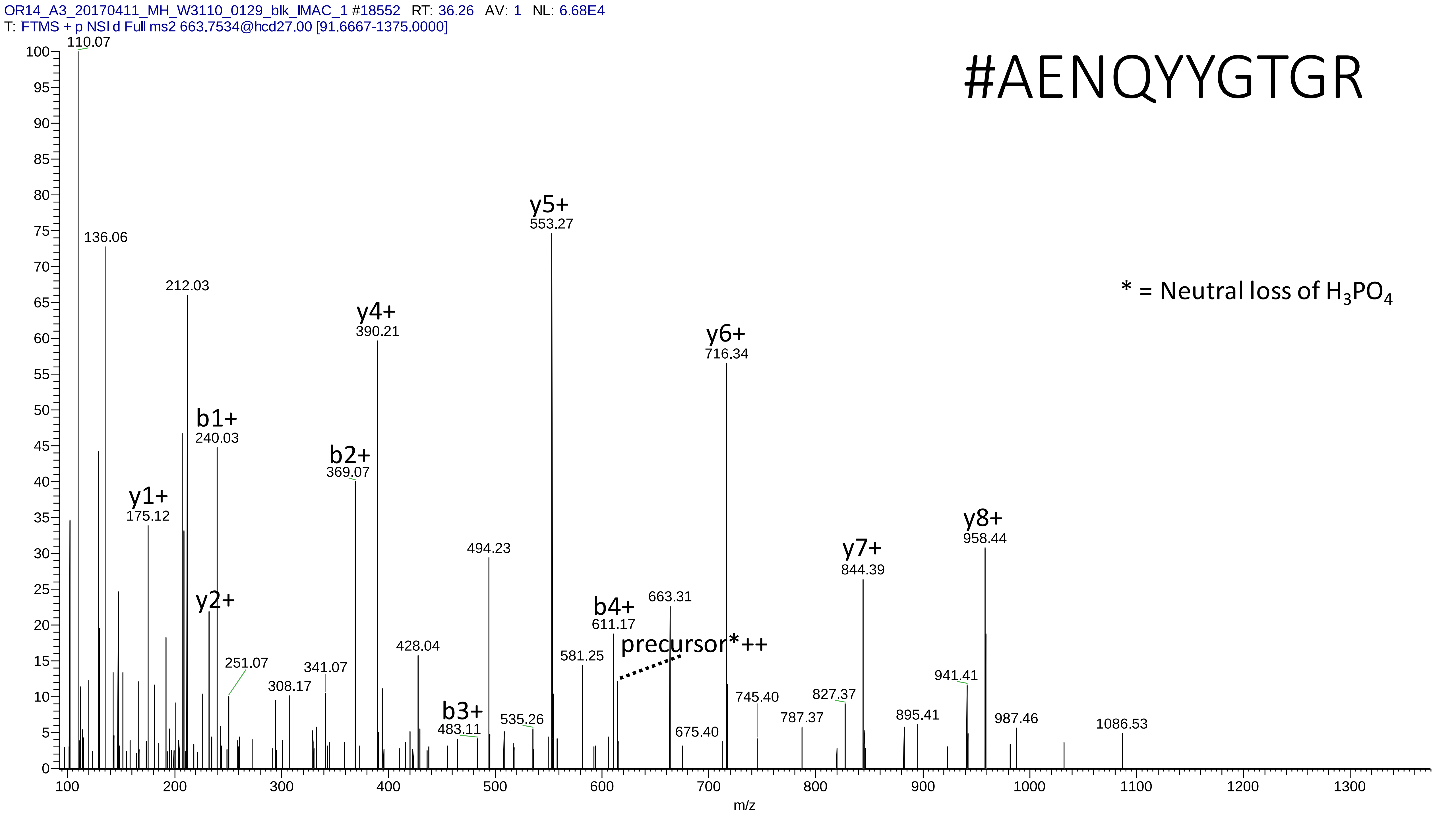
OR14_A3_20170411_MH_W3110_0129_blk_IMAC_1 #18552 RT: 36.26 AV: 1 NL: 6.68E4
T: FTMS + p NSI d Full ms2 663.7534@hcd27.00 [91.6667-1375.0000]
#AENQYYGTGR
* = Neutral loss of $H_3PO_4$
110.07
136.06
212.03
y1+
175.12
y2+
b1+
240.03
251.07
308.17
341.07
b2+
369.07
y4+
390.21
428.04
b3+
483.11
494.23
535.26
y5+
553.27
581.25
b4+
611.17
precursor*++
663.31
675.40
y6+
716.34
745.40
787.37
827.37
y7+
844.39
895.41
941.41
y8+
958.44
987.46
1086.53
m/z

# 5

- Experimental mass: 299.0599
- Unimod information: not found
- Possible composition: H(16) C(8) N(2) O(8) P
- Sites in Unimod: not found
- Sites observed: S;T;C
- Features: neutral loss of $H_3PO_4$ are observed at both precursor and fragment ions; structure unknown

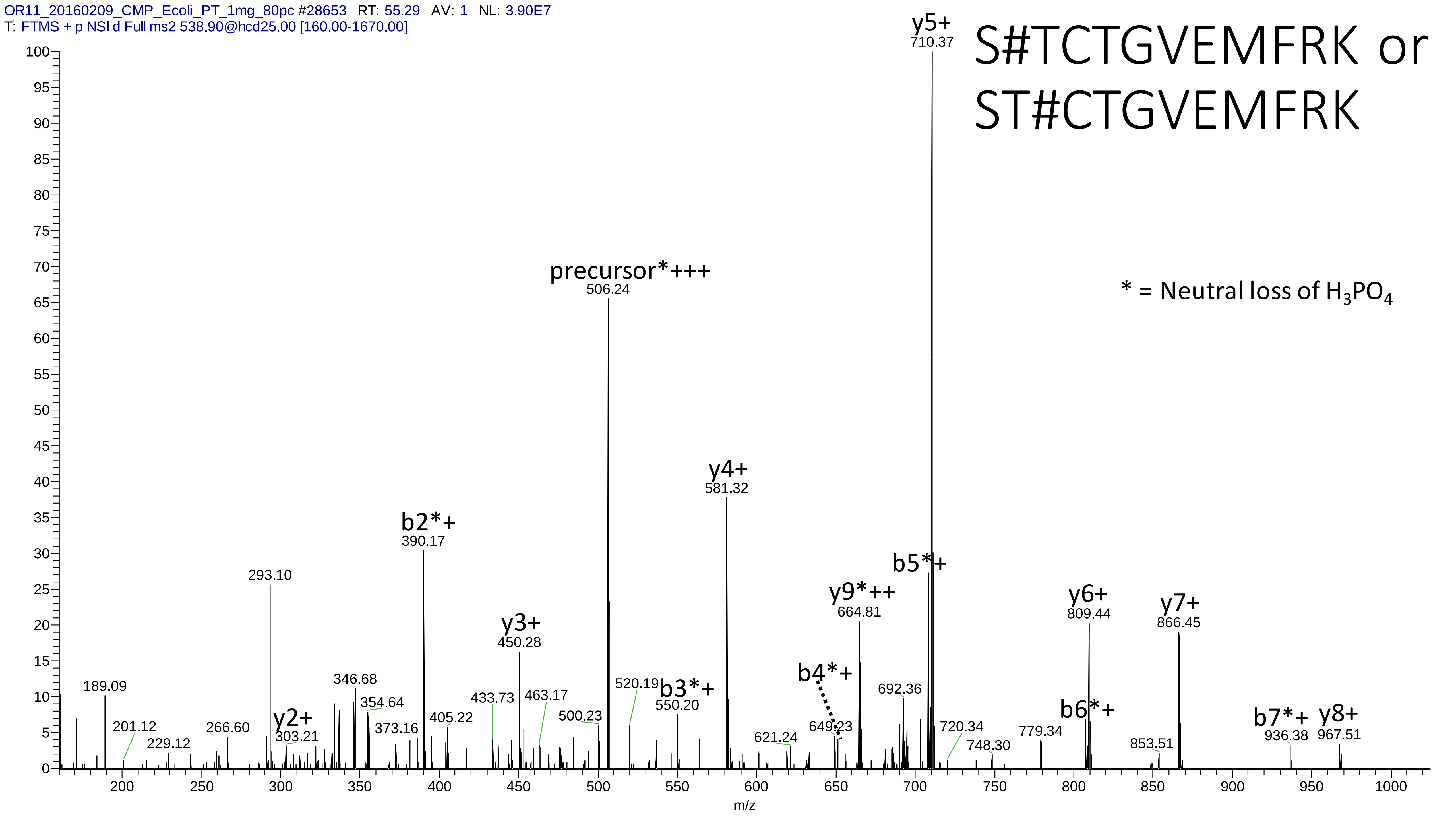
OR11_20160209_CMP_Ecoli_PT_1mg_80pc #28653 RT: 55.29 AV: 1 NL: 3.90E7
T: FTMS + p NSI d Full ms2 538.90@hcd25.00 [160.00-1670.00]
S#TCTGVEMFRK or
ST#CTGVEMFRK
* = Neutral loss of $H_3PO_4$
y5+ 710.37
precursor*+++ 506.24
y4+ 581.32
b2*+ 390.17
293.10
y3+ 450.28
b5*+
y9*++ 664.81
y6+ 809.44
y7+ 866.45
189.09
201.12
229.12
266.60
y2+ 303.21
346.68
354.64
373.16
405.22
433.73
463.17
500.23
520.19
b3*+ 550.20
b4*+
621.24
649.23
692.36
720.34
748.30
779.34
b6*+
853.51
b7*+ 936.38
y8+ 967.51
m/z

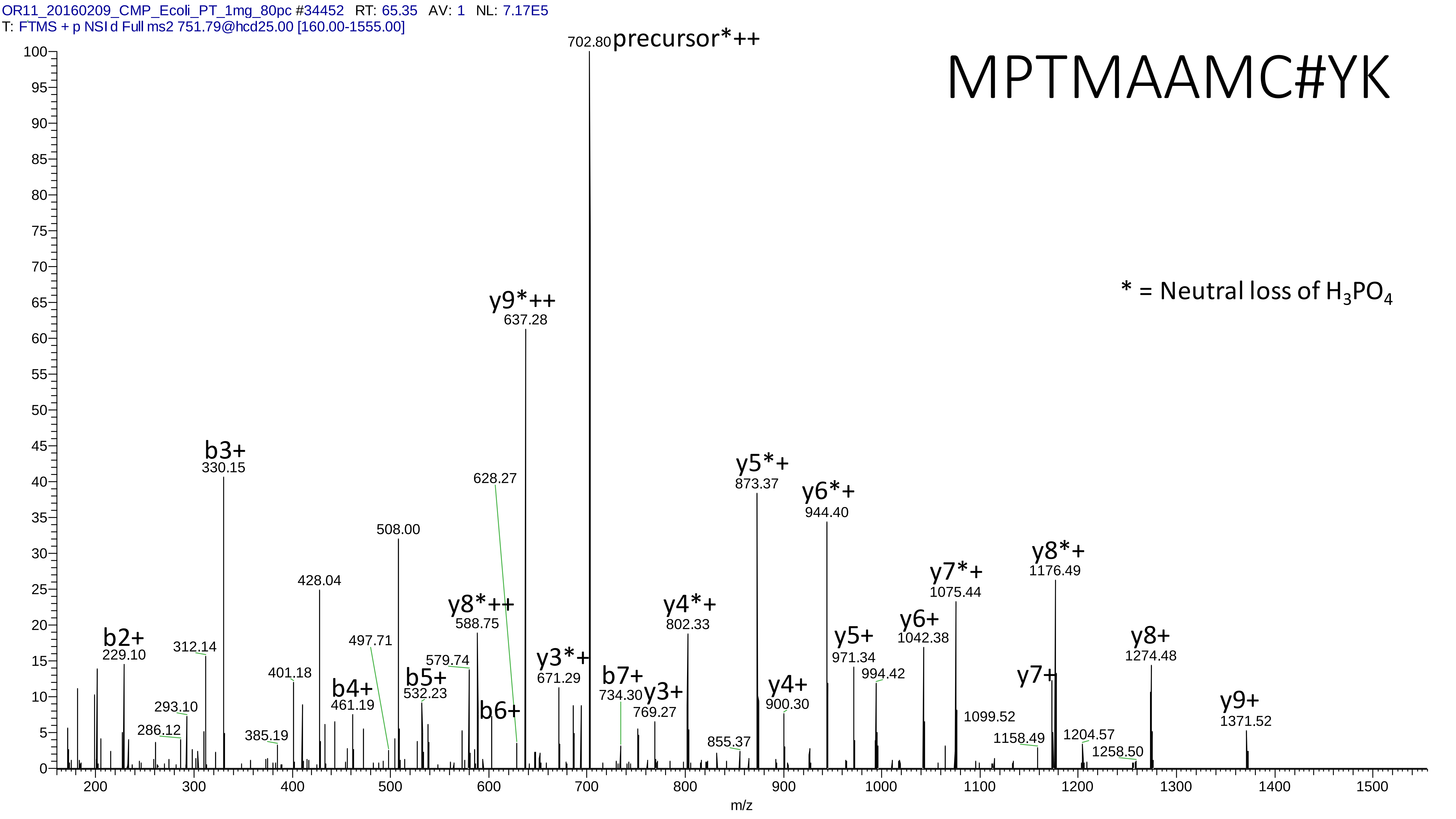
OR11_20160209_CMP_Ecoli_PT_1mg_80pc #34452 RT: 65.35 AV: 1 NL: 7.17E5
T: FTMS + p NSI d Full ms2 751.79@hcd25.00 [160.00-1555.00]
MPTMAAMC#YK
* = Neutral loss of H3PO4
702.80 precursor*++
y9*++ 637.28
b3+ 330.15
628.27
508.00
428.04
y8*++ 588.75
b2+ 229.10
312.14
401.18
497.71
579.74
b5+ 532.23
b4+ 461.19
b6+
293.10
286.12
385.19
y3*+ 671.29
b7+ 734.30
y3+ 769.27
y4*+ 802.33
y5*+ 873.37
y4+ 900.30
855.37
y6*+ 944.40
y5+ 971.34
994.42
y6+ 1042.38
y7*+ 1075.44
1099.52
y8*+ 1176.49
y7+
1158.49
1204.57
1258.50
y8+ 1274.48
y9+ 1371.52
m/z

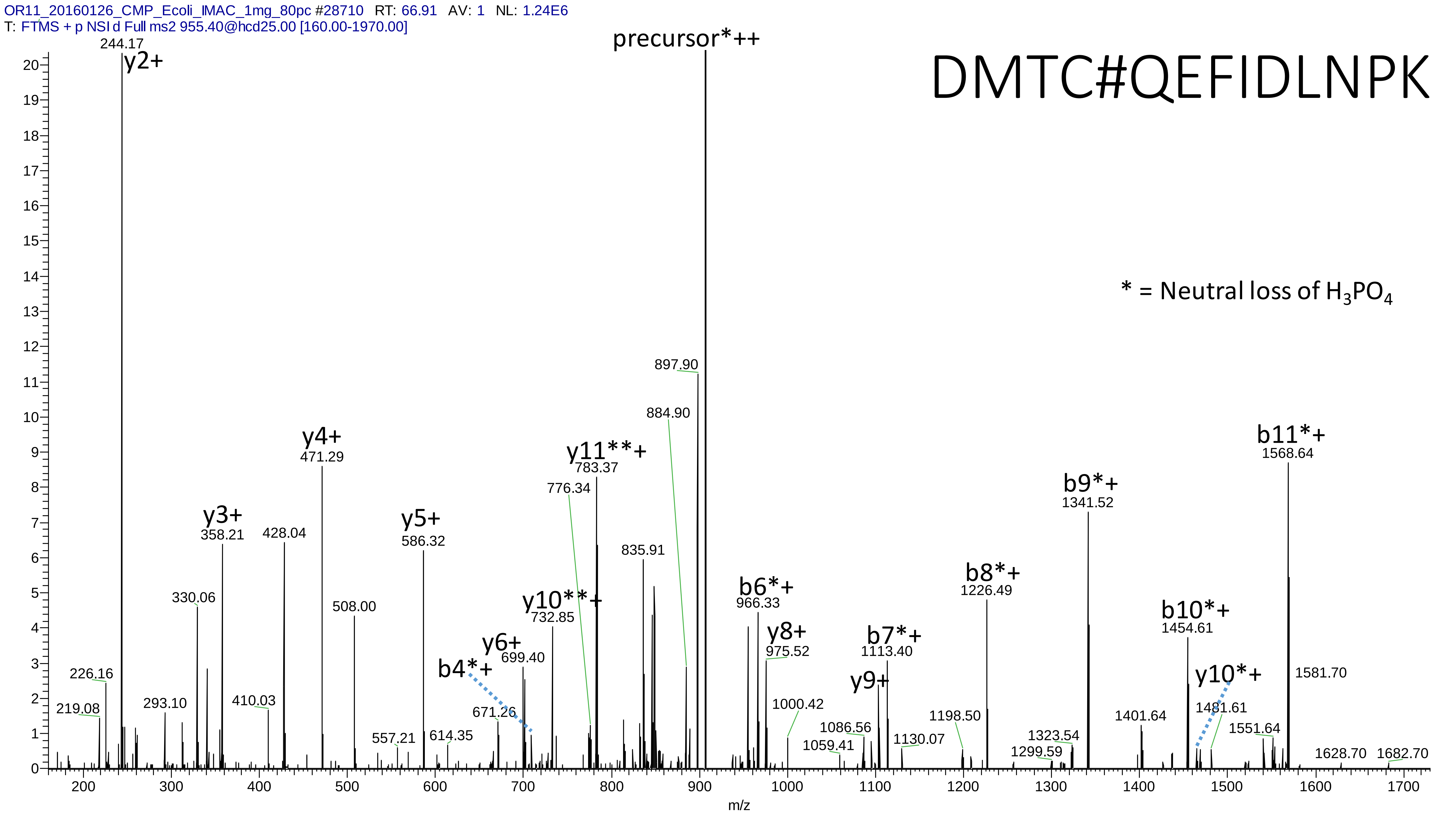

OR11_20160126_CMP_Ecoli_IMAC_1mg_80pc #28710 RT: 66.91 AV: 1 NL: 1.24E6
T: FTMS + p NSI d Full ms2 955.40@hcd25.00 [160.00-1970.00]
DMTC#QEFIDLNPK
* = Neutral loss of H3PO4
precursor*++
y2+
244.17
y3+
358.21
y4+
471.29
y5+
586.32
y6+
699.40
y8+
975.52
y9+
y10**+
732.85
y11**+
783.37
y10*+
b4*+
b6*+
966.33
b7*+
1113.40
b8*+
1226.49
b9*+
1341.52
b10*+
1454.61
b11*+
1568.64
219.08
226.16
293.10
330.06
410.03
428.04
508.00
557.21
614.35
671.26
776.34
835.91
884.90
897.90
1000.42
1059.41
1086.56
1130.07
1198.50
1299.59
1323.54
1401.64
1481.61
1551.64
1581.70
1628.70
1682.70
m/z

# 6

- Experimental mass: -57.0211
- Unimod information: Asn->Gly / Gln->Ala
- Possible composition: Gly-loss or Cys not modified by Carbamidomethyl, [H(-3) C(-2) N(-1) O(-1)]
- Sites in Unimod: N;Q
- Sites observed: G;C

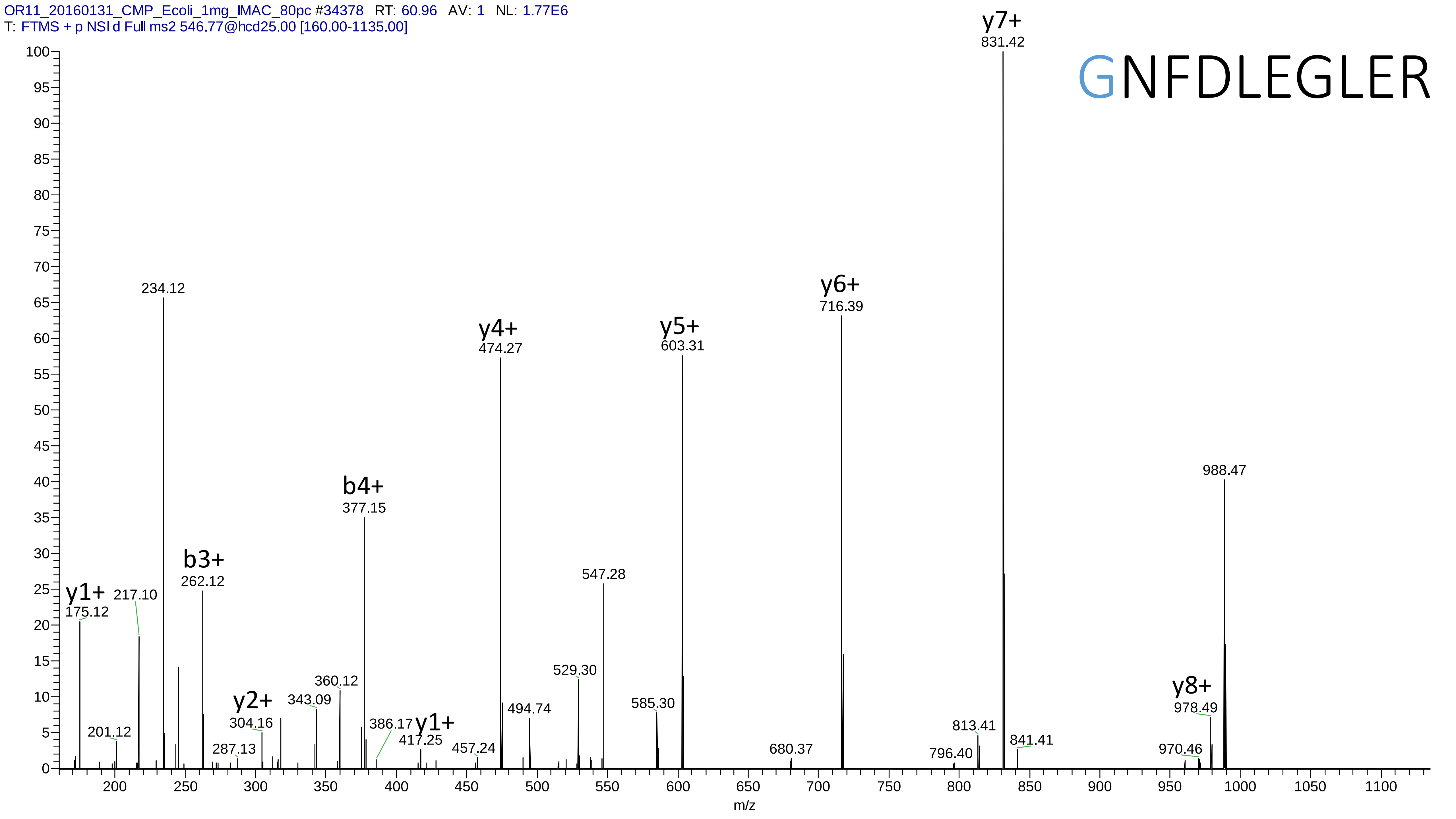
OR11_20160131_CMP_Ecoli_1mg_IMAC_80pc #34378 RT: 60.96 AV: 1 NL: 1.77E6
T: FTMS + p NSI d Full ms2 546.77@hcd25.00 [160.00-1135.00]
GNFDLEGLER
y7+ 831.42
y6+ 716.39
y5+ 603.31
y4+ 474.27
b4+ 377.15
b3+ 262.12
y1+ 175.12
y2+ 304.16
y1+ 417.25
y8+ 978.49
234.12
217.10
201.12
287.13
343.09
360.12
386.17
457.24
494.74
529.30
547.28
585.30
680.37
796.40
813.41
841.41
970.46
988.47
m/z

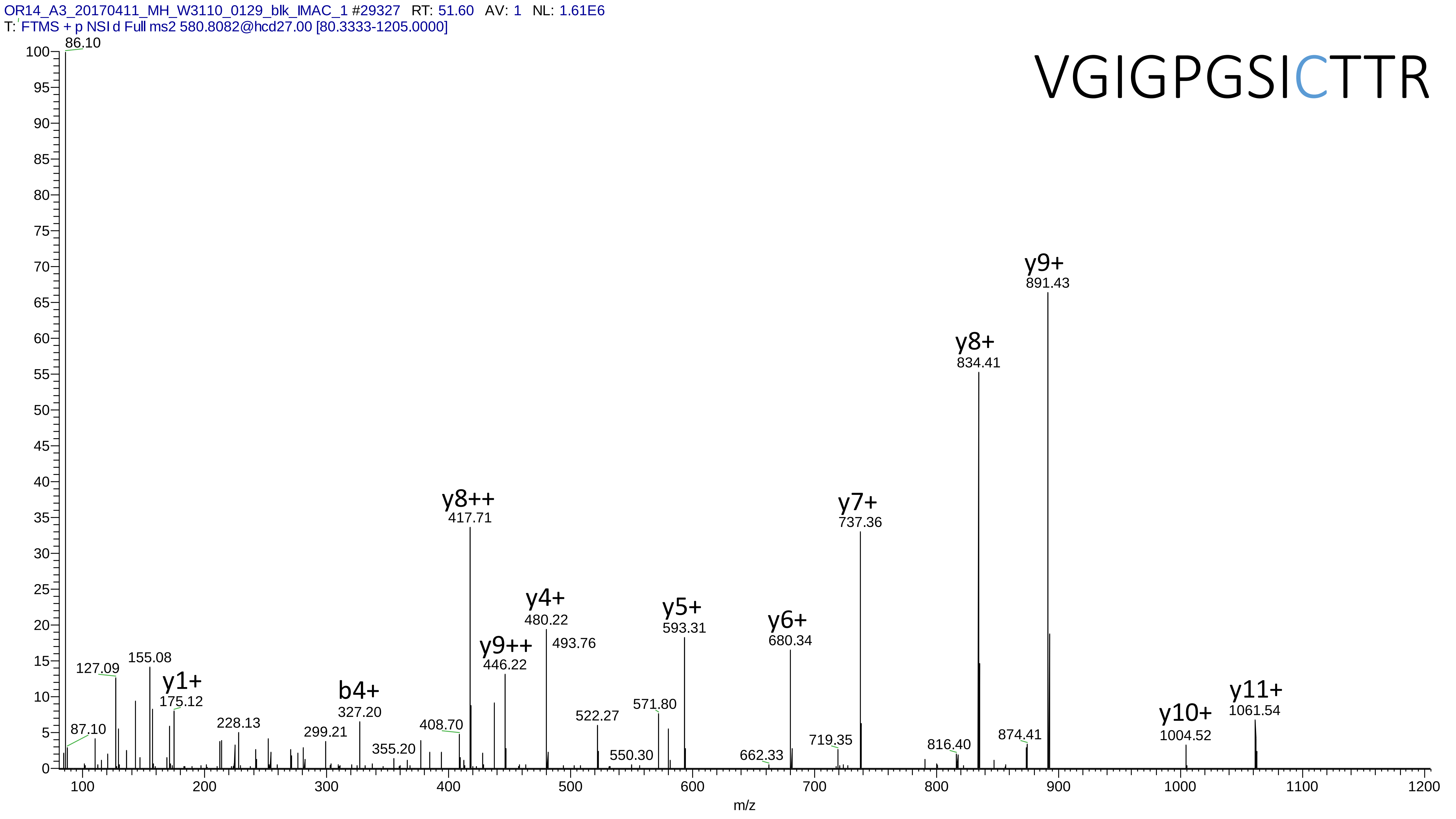
OR14_A3_20170411_MH_W3110_0129_blk_IMAC_1 #29327 RT: 51.60 AV: 1 NL: 1.61E6
T: FTMS + p NSI d Full ms2 580.8082@hcd27.00 [80.3333-1205.0000]
VGIGPGSICTTR
86.10
87.10
127.09
155.08
y1+
175.12
228.13
299.21
b4+
327.20
355.20
408.70
y8++
417.71
y9++
446.22
y4+
480.22
493.76
522.27
550.30
571.80
y5+
593.31
662.33
y6+
680.34
719.35
y7+
737.36
816.40
y8+
834.41
874.41
y9+
891.43
y10+
1004.52
y11+
1061.54
m/z

# 7

- Experimental mass: 0.9843
- Unimod information: Deamidation / Asn->Asp / Gln->Glu
- Possible composition: H(-1) N(-1) O
- Sites in Unimod: N;Q;R;F
- Sites observed: N
- Features: 81.7% (232/284) peptides contain Asn/Gln

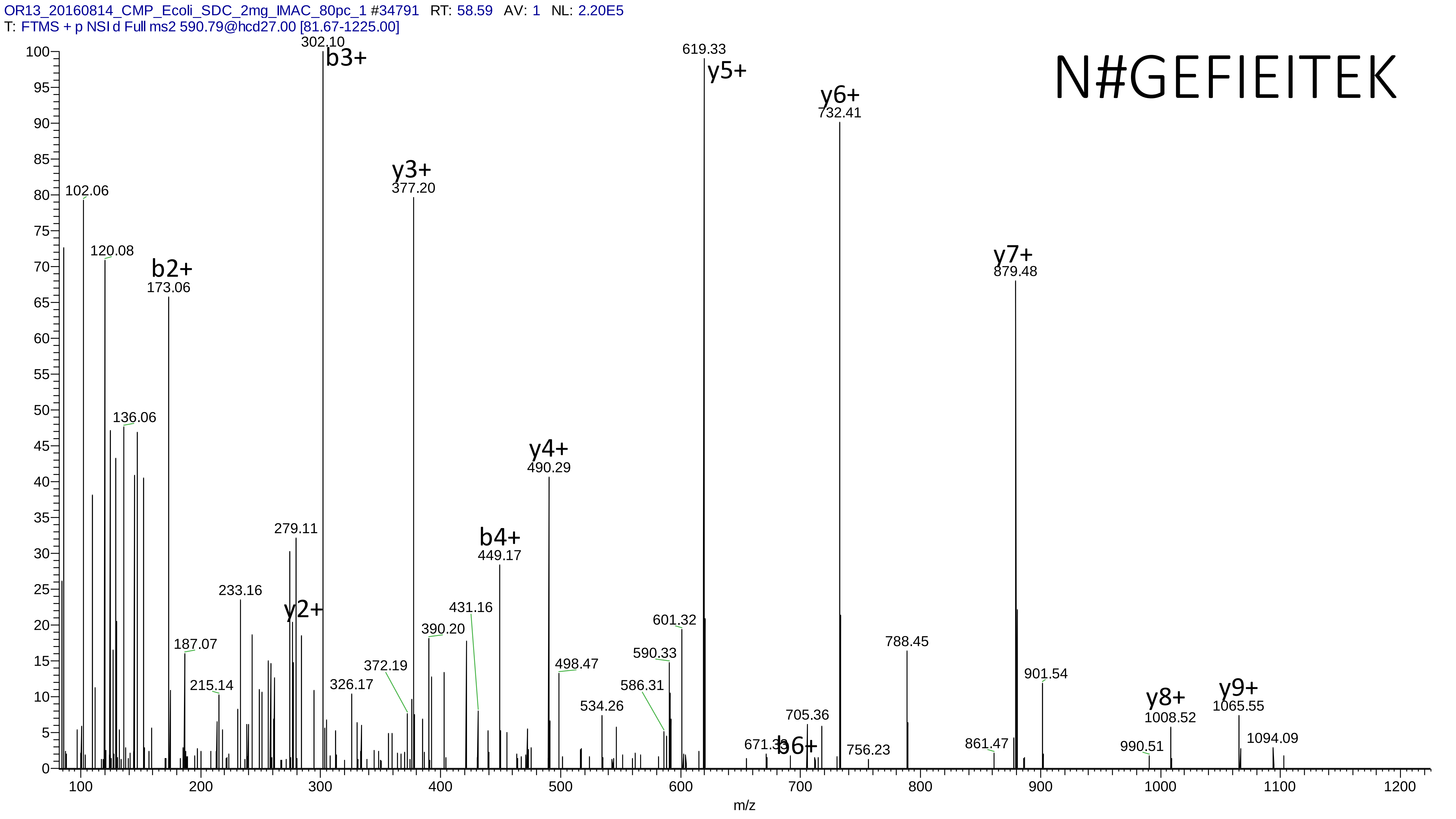
OR13_20160814_CMP_Ecoli_SDC_2mg_IMAC_80pc_1 #34791 RT: 58.59 AV: 1 NL: 2.20E5
T: FTMS + p NSI d Full ms2 590.79@hcd27.00 [81.67-1225.00]
N#GEFIEITEK
302.10 b3+
619.33 y5+
y6+ 732.41
y3+ 377.20
102.06
120.08
b2+ 173.06
y7+ 879.48
136.06
y4+ 490.29
279.11
b4+ 449.17
233.16
y2+
431.16
390.20
601.32
187.07
372.19
590.33
498.47
788.45
215.14
326.17
586.31
901.54
534.26
705.36
y8+ 1008.52
y9+ 1065.55
671.33
b6+
756.23
861.47
990.51
1094.09
m/z

# 8

- Experimental mass: 122.9616
- Unimod information: not found
- Possible composition: no Carbamidomethyl, then add H(5)C(4)O(6)P(1)
- Sites in Unimod: not found
- Sites observed: C
- Features: 100% (9/9) peptides contain C, 88.9% (8/9) peptides contain C at n-term, 94.5% (549/581) PSMs are from the same peptide WTGMCGELAGDER, neutral loss of $H_3PO_4$ are **not** observed, structure unknown

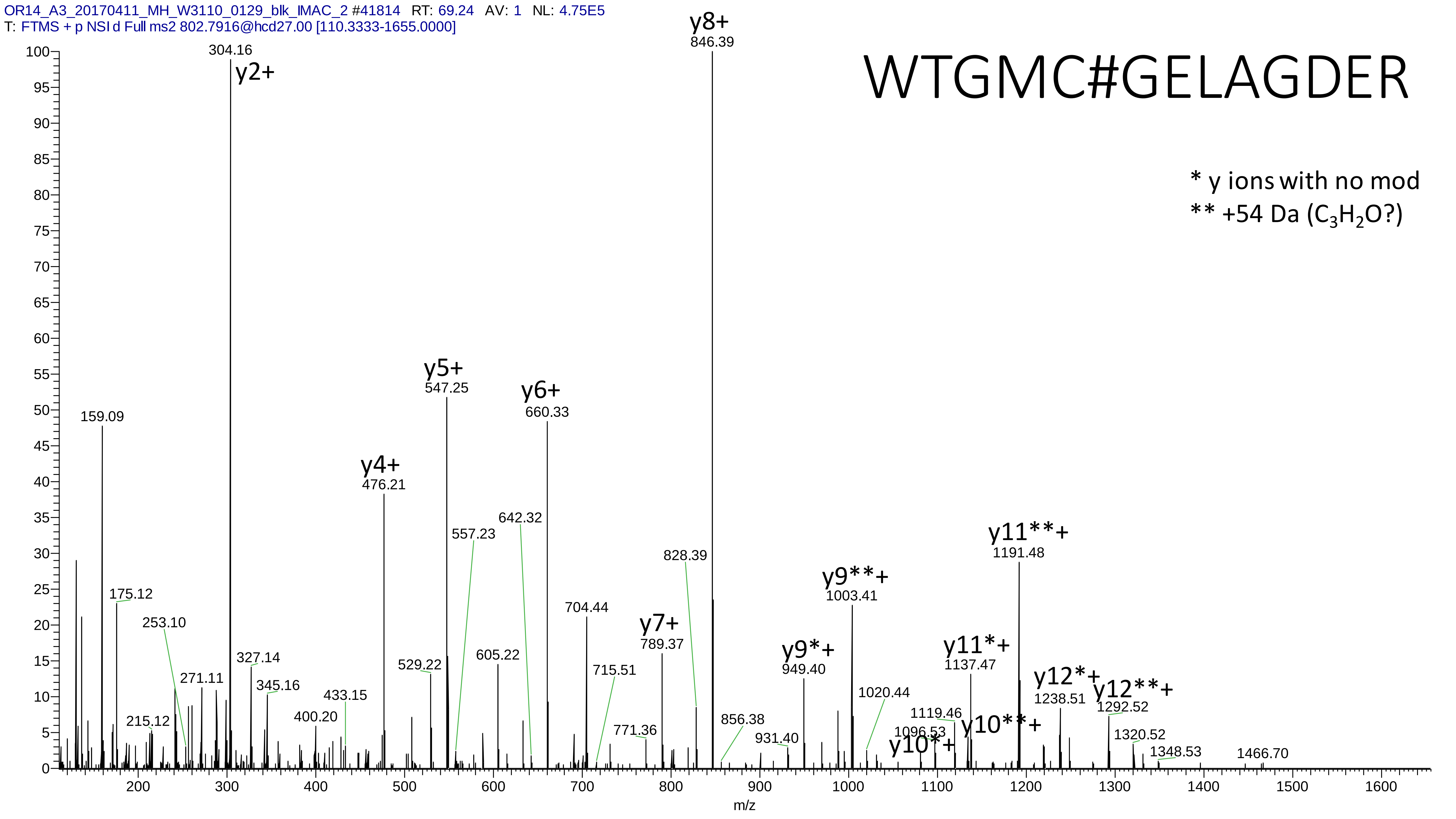
OR14_A3_20170411_MH_W3110_0129_blk_IMAC_2 #41814 RT: 69.24 AV: 1 NL: 4.75E5
T: FTMS + p NSI d Full ms2 802.7916@hcd27.00 [110.3333-1655.0000]
WTGMC#GELAGDER
* y ions with no mod
** +54 Da (C3H2O?)
y2+ 304.16
159.09
175.12
253.10
215.12
271.11
327.14
345.16
400.20
433.15
y4+ 476.21
529.22
y5+ 547.25
557.23
605.22
642.32
y6+ 660.33
704.44
715.51
771.36
y7+ 789.37
828.39
y8+ 846.39
856.38
931.40
y9*+ 949.40
y9**+ 1003.41
1020.44
1096.53
y10*+
1119.46
y10**+
y11*+ 1137.47
y11**+ 1191.48
y12*+ 1238.51
y12**+ 1292.52
1320.52
1348.53
1466.70
m/z

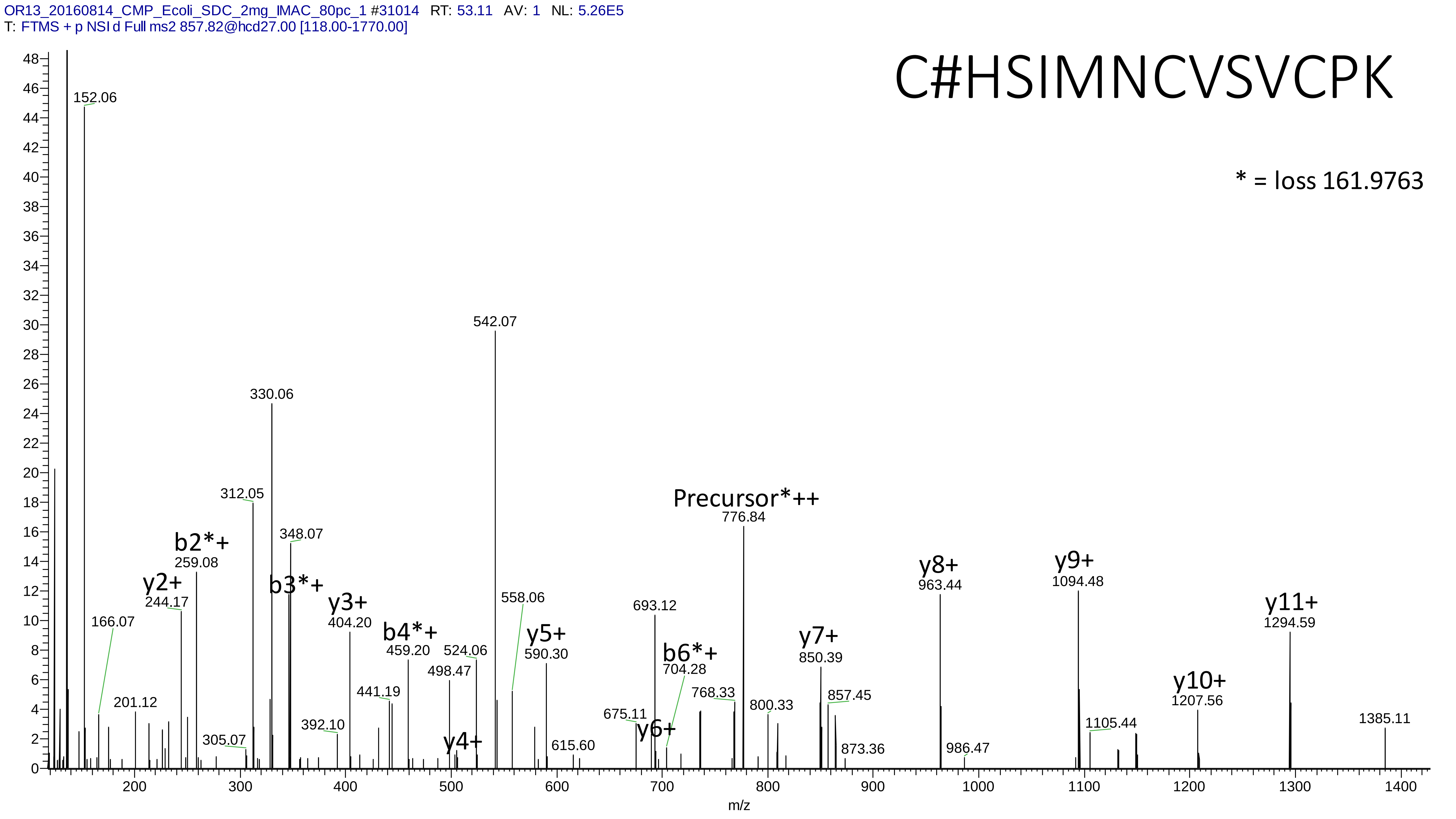
OR13_20160814_CMP_Ecoli_SDC_2mg_IMAC_80pc_1 #31014 RT: 53.11 AV: 1 NL: 5.26E5
T: FTMS + p NSI d Full ms2 857.82@hcd27.00 [118.00-1770.00]
C#HSIMNCVSVCPK
* = loss 161.9763
152.06
542.07
330.06
312.05
348.07
b2*+
259.08
y2+
244.17
b3*+
y3+
404.20
166.07
b4*+
459.20
524.06
498.47
558.06
y5+
590.30
201.12
441.19
392.10
305.07
y4+
615.60
675.11
y6+
693.12
b6*+
704.28
768.33
Precursor*++
776.84
800.33
y7+
850.39
857.45
873.36
y8+
963.44
986.47
y9+
1094.48
1105.44
y10+
1207.56
y11+
1294.59
1385.11
m/z

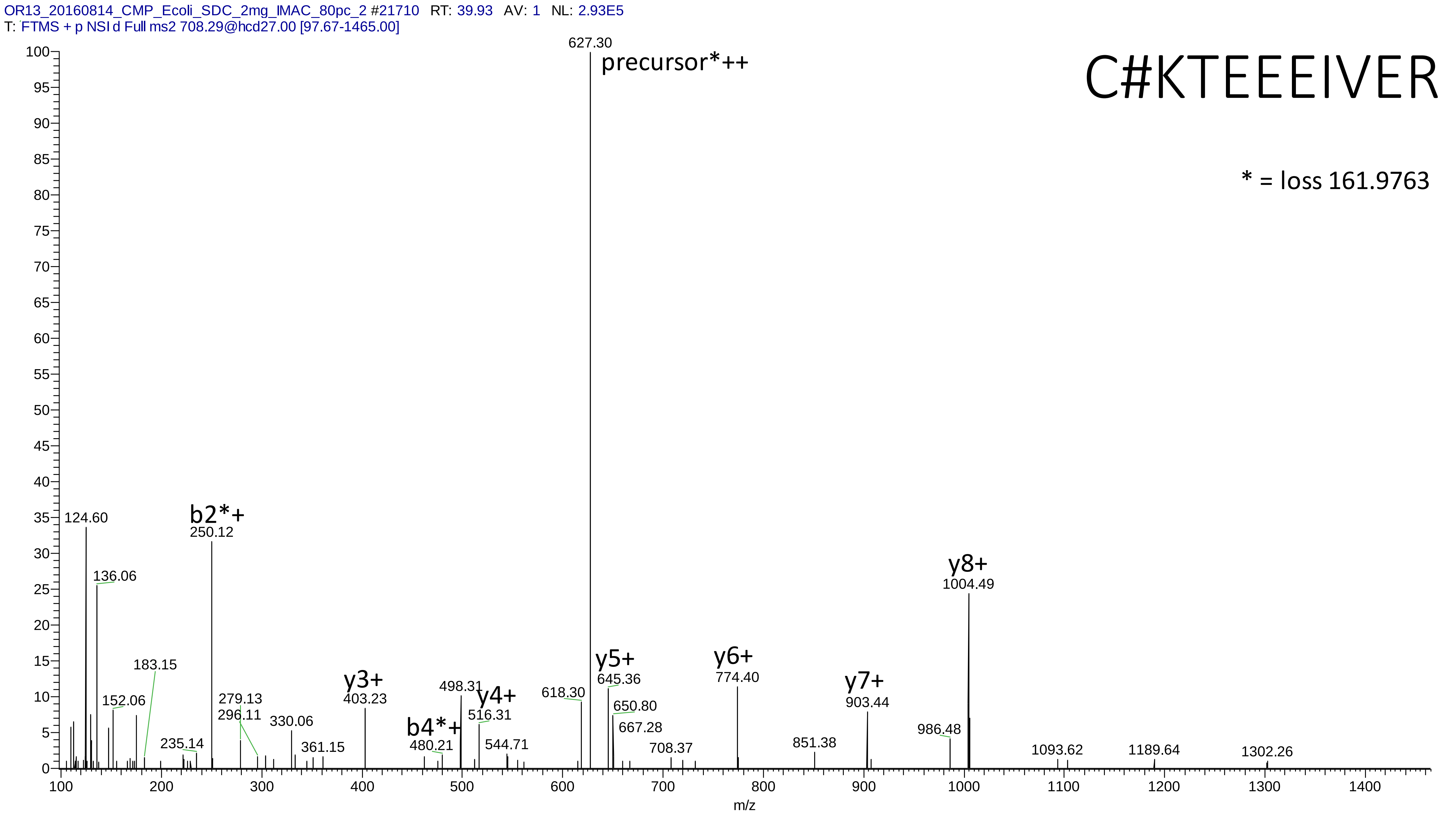

OR13_20160814_CMP_Ecoli_SDC_2mg_IMAC_80pc_2 #21710 RT: 39.93 AV: 1 NL: 2.93E5
T: FTMS + p NSI d Full ms2 708.29@hcd27.00 [97.67-1465.00]
C#KTEEEIVER
* = loss 161.9763
627.30
precursor*++
124.60
b2*+
250.12
136.06
y8+
1004.49
183.15
152.06
279.13
296.11
235.14
330.06
361.15
y3+
403.23
498.31
y4+
516.31
b4*+
480.21
544.71
618.30
y5+
645.36
650.80
667.28
708.37
y6+
774.40
851.38
y7+
903.44
986.48
1093.62
1189.64
1302.26
m/z

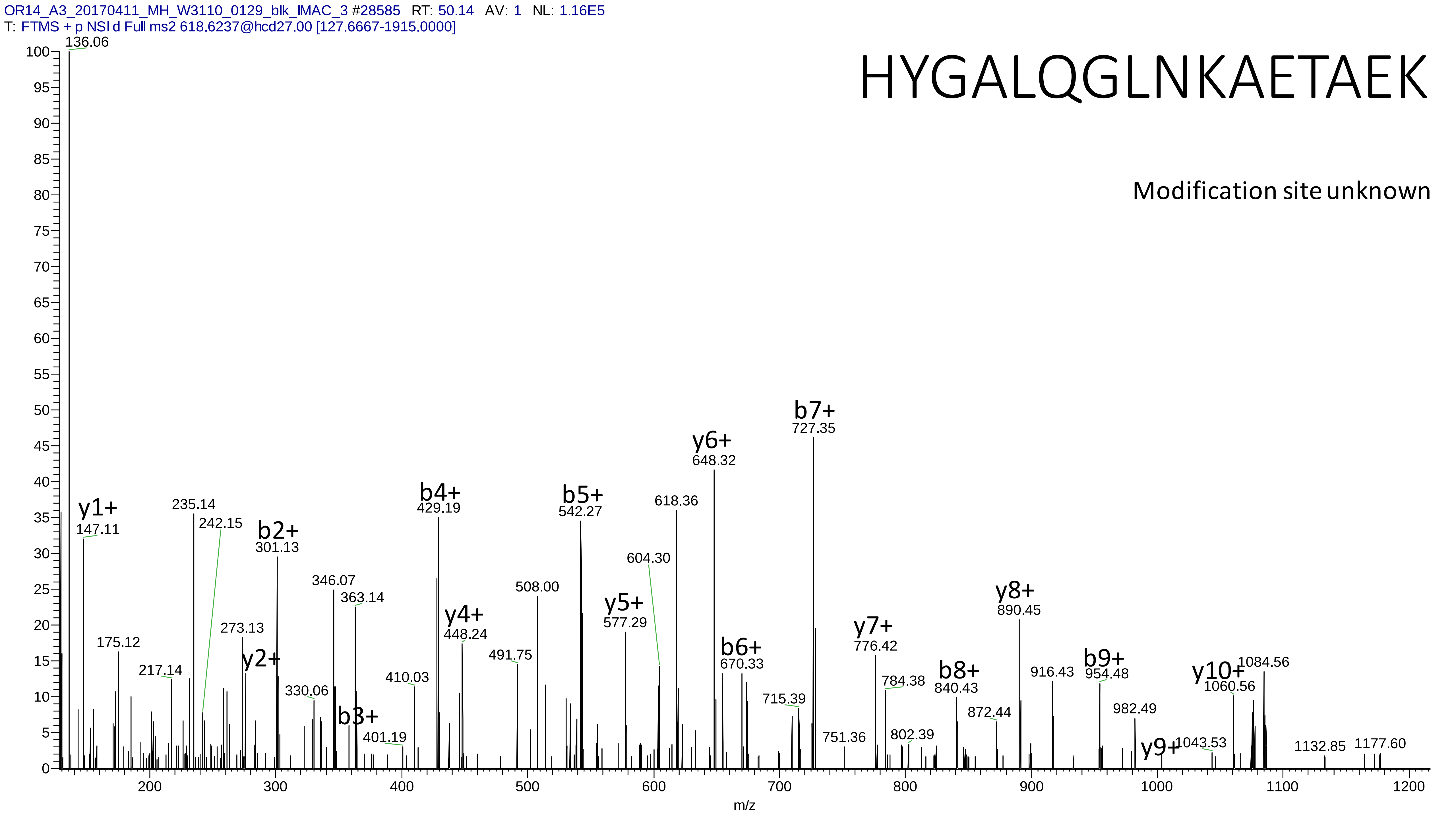
OR14_A3_20170411_MH_W3110_0129_blk_IMAC_3 #28585 RT: 50.14 AV: 1 NL: 1.16E5
T: FTMS + p NSI d Full ms2 618.6237@hcd27.00 [127.6667-1915.0000]
HYGALQGLNKAETAEK
Modification site unknown
136.06
y1+
147.11
175.12
217.14
235.14
242.15
273.13
y2+
b2+
301.13
330.06
346.07
b3+
363.14
401.19
410.03
b4+
429.19
y4+
448.24
491.75
508.00
b5+
542.27
y5+
577.29
604.30
618.36
y6+
648.32
b6+
670.33
715.39
b7+
727.35
751.36
y7+
776.42
784.38
802.39
b8+
840.43
872.44
y8+
890.45
916.43
b9+
954.48
982.49
y9+
1043.53
y10+
1060.56
1084.56
1132.85
1177.60
m/z

# 9

- Experimental mass: 151.9879
- Unimod information: not found
- Possible composition: H(5) C(3) O(5) P
- Sites in Unimod: not found
- Sites observed: K; protein n-term
- Features: 58.4% (66/113) peptides start with the first amino acid in the proteins, 32.7% (37/113) peptides start with the second amino acid in the proteins, in total 91.2% (103/113) peptides contains protein n-term; neutral loss of $H_3PO_4$ are observed at both precursor and fragment ions

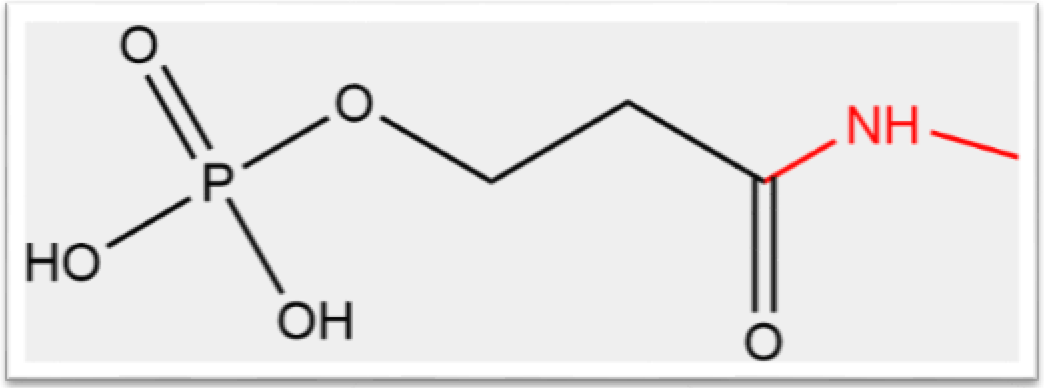

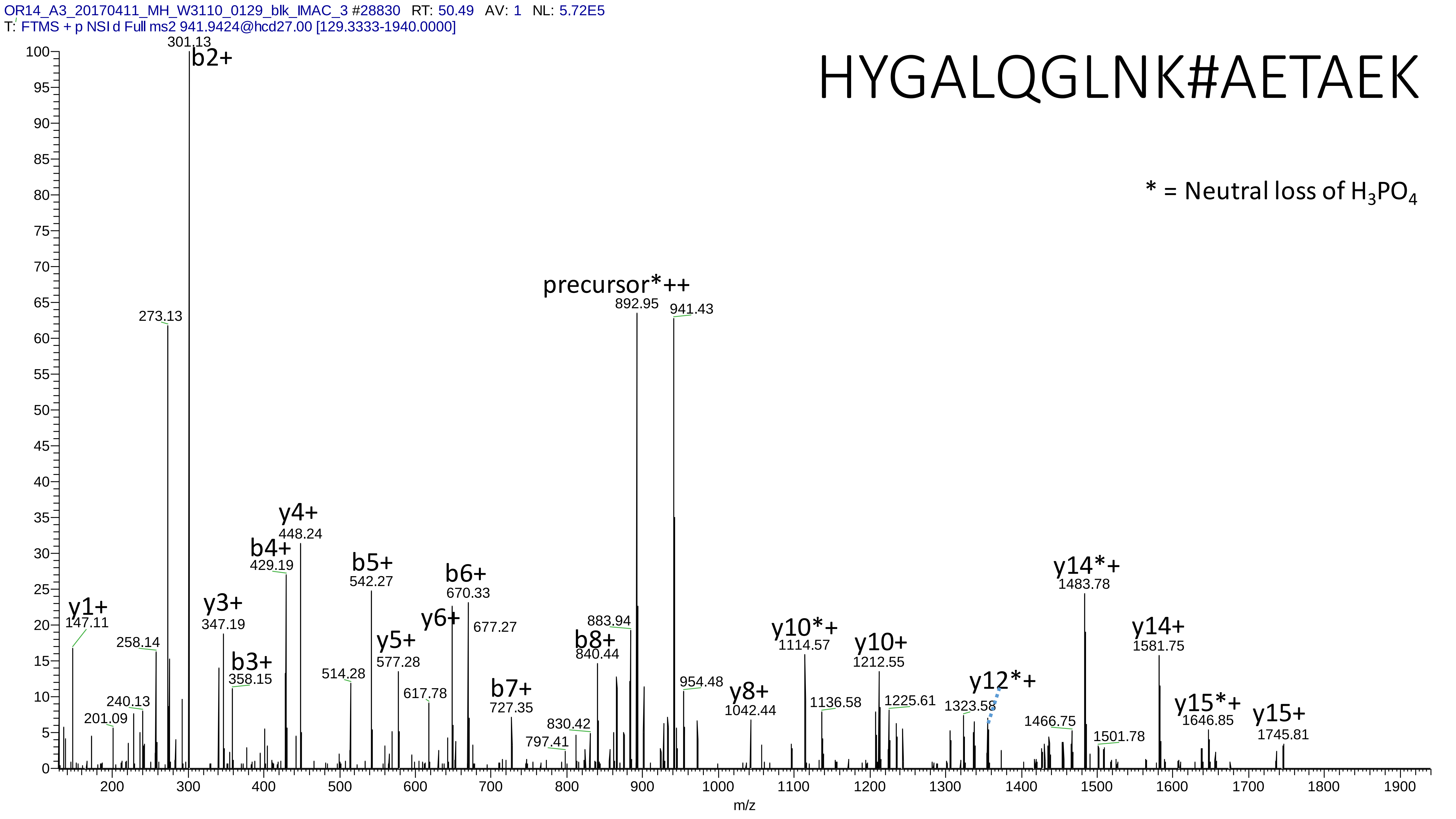
OR14_A3_20170411_MH_W3110_0129_blk_IMAC_3 #28830 RT: 50.49 AV: 1 NL: 5.72E5
T: FTMS + p NSI d Full ms2 941.9424@hcd27.00 [129.3333-1940.0000]
HYGALQGLNK#AETAEK
* = Neutral loss of H3PO4
301.13
b2+
273.13
precursor*++
892.95
941.43
y1+
147.11
258.14
240.13
201.09
y3+
347.19
b3+
358.15
b4+
429.19
y4+
448.24
514.28
b5+
542.27
y5+
577.28
617.78
y6+
b6+
670.33
677.27
b7+
727.35
797.41
830.42
b8+
840.44
883.94
954.48
y8+
1042.44
y10*+
1114.57
1136.58
y10+
1212.55
1225.61
1323.58
y12*+
1466.75
y14*+
1483.78
1501.78
y14+
1581.75
y15*+
1646.85
y15+
1745.81
m/z

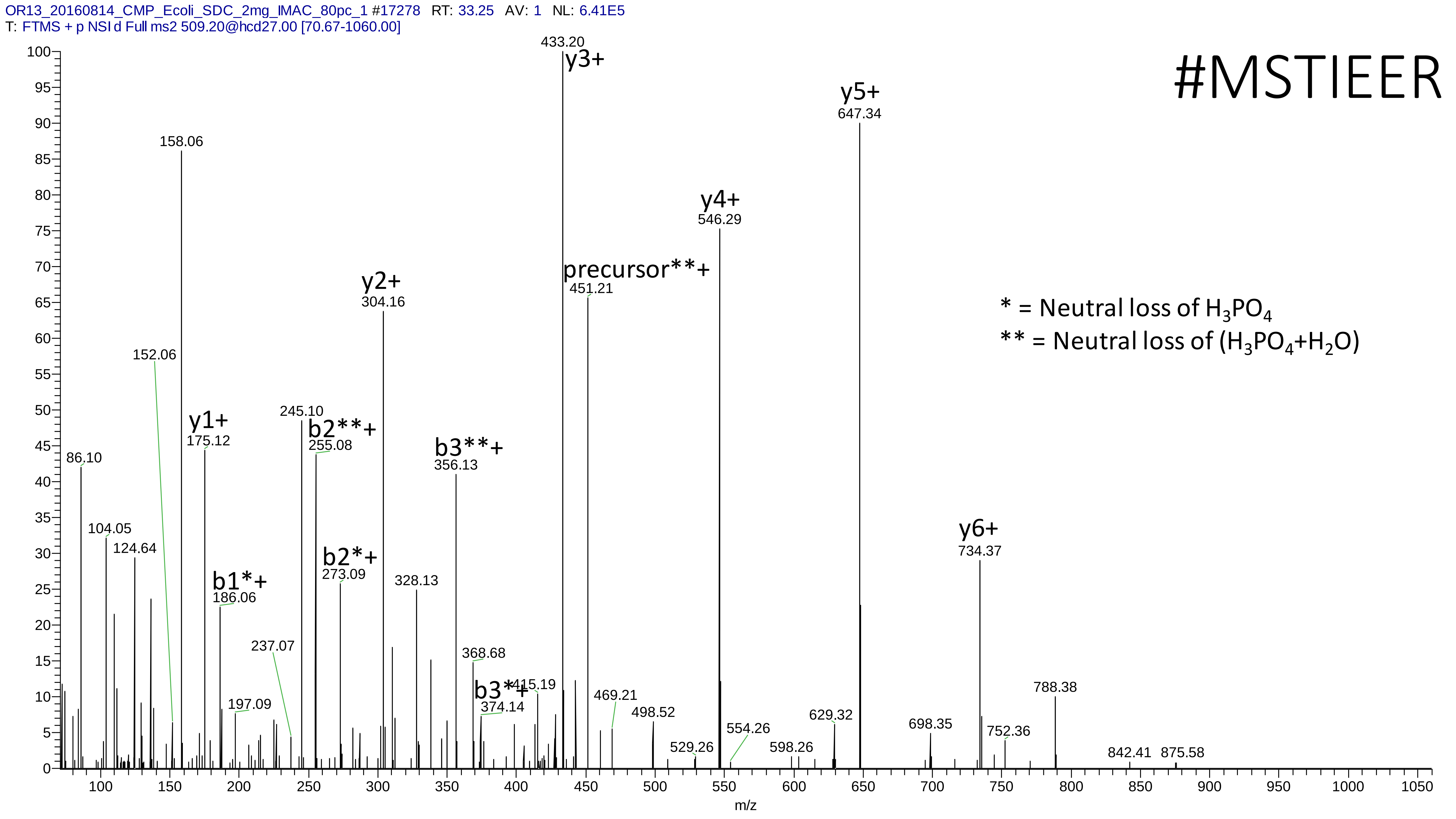

OR13_20160814_CMP_Ecoli_SDC_2mg_IMAC_80pc_1 #17278 RT: 33.25 AV: 1 NL: 6.41E5
T: FTMS + p NSI d Full ms2 509.20@hcd27.00 [70.67-1060.00]
#MSTIEER
* = Neutral loss of H3PO4
** = Neutral loss of (H3PO4+H2O)
y3+ 433.20
y5+ 647.34
158.06
y4+ 546.29
precursor**+ 451.21
y2+ 304.16
152.06
y1+ 175.12
245.10
b2**+ 255.08
b3**+ 356.13
86.10
104.05
124.64
b1*+ 186.06
b2*+ 273.09
328.13
y6+ 734.37
237.07
368.68
197.09
b3*+ 374.14
415.19
469.21
498.52
529.26
554.26
598.26
629.32
698.35
752.36
788.38
842.41
875.58
m/z

# 10

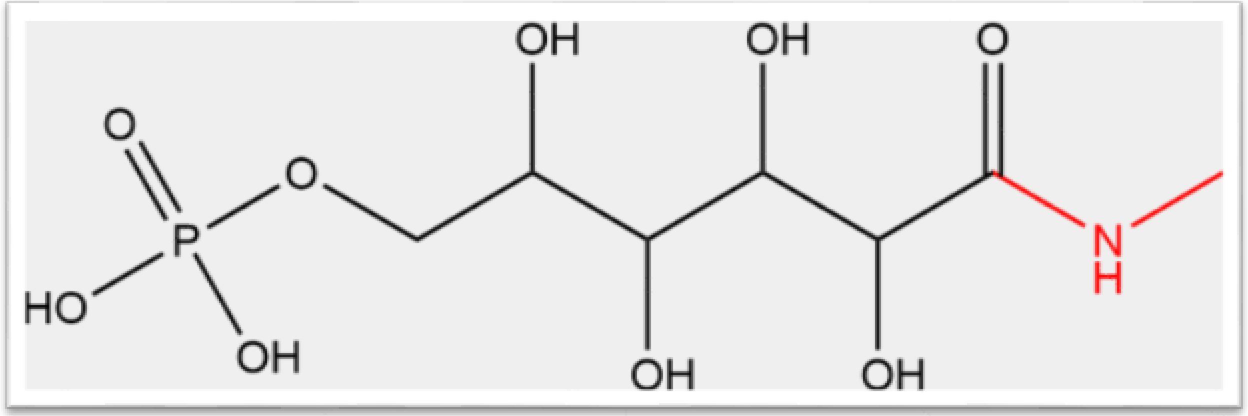


- Experimental mass: 258.0148
- Unimod information: Phosphogluconoylation
- Possible composition: H(11) C(6) O(9) P
- Sites in Unimod: K;N-term
- Sites observed: K;N-term
- Features: 74.8% (95/127) peptides contain a miss cleavage site K, 11.8% (15/127) peptides start with the first amino acid in the proteins, 17.3% (22/127) peptides start with the second amino acid in the proteins, in total 29.1% (37/127) peptides contains protein n-term; neutral loss of $H_3PO_4$ from precursor ions are observed

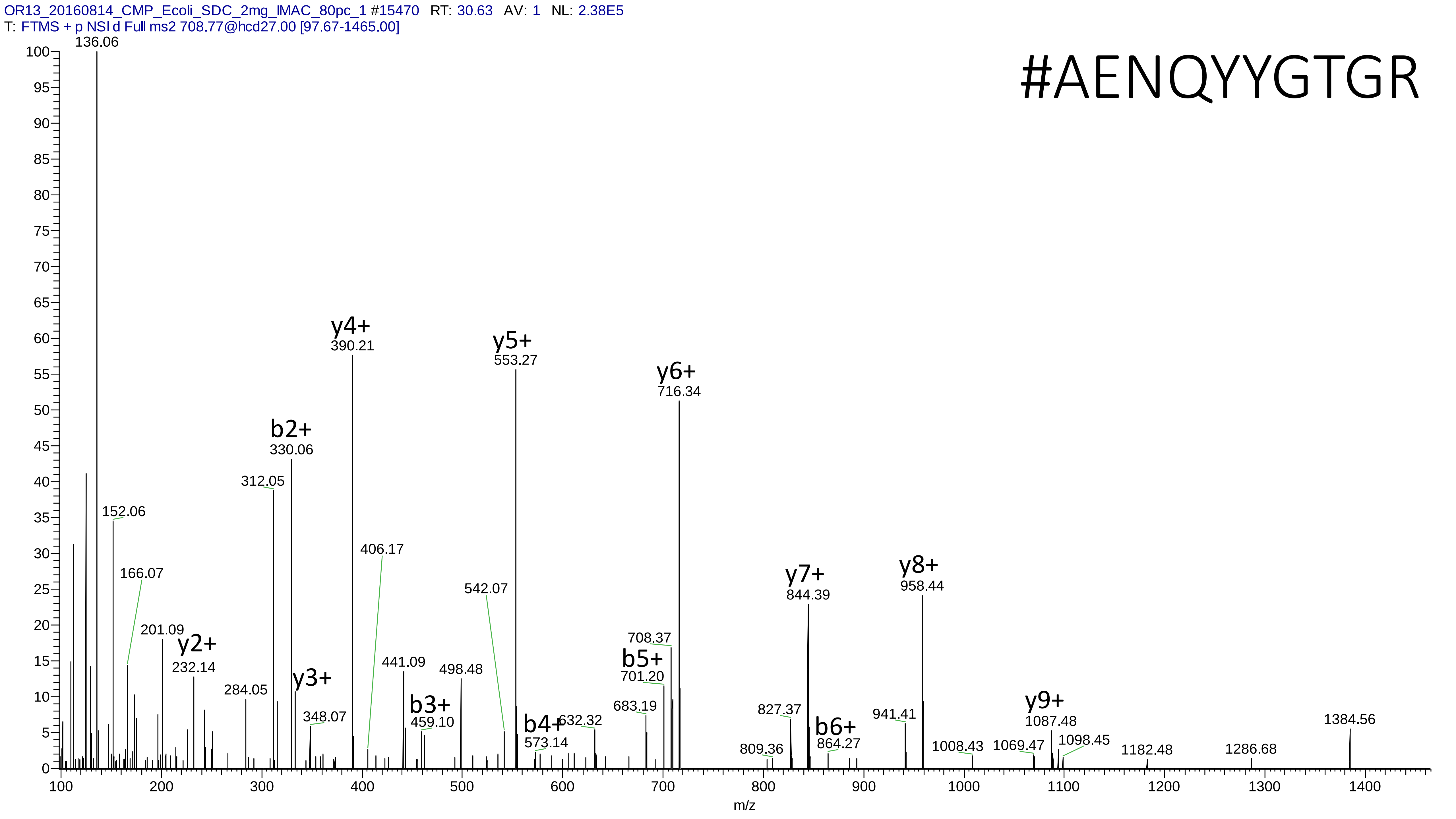

OR13_20160814_CMP_Ecoli_SDC_2mg_IMAC_80pc_1 #15470 RT: 30.63 AV: 1 NL: 2.38E5
T: FTMS + p NSI d Full ms2 708.77@hcd27.00 [97.67-1465.00]
#AENQYYGTGR
136.06
152.06
166.07
201.09
y2+ 232.14
284.05
312.05
b2+ 330.06
y3+
348.07
y4+ 390.21
406.17
441.09
b3+ 459.10
498.48
542.07
y5+ 553.27
b4+ 573.14
632.32
683.19
b5+ 701.20
708.37
y6+ 716.34
809.36
827.37
y7+ 844.39
b6+ 864.27
941.41
y8+ 958.44
1008.43
1069.47
y9+ 1087.48
1098.45
1182.48
1286.68
1384.56
m/z

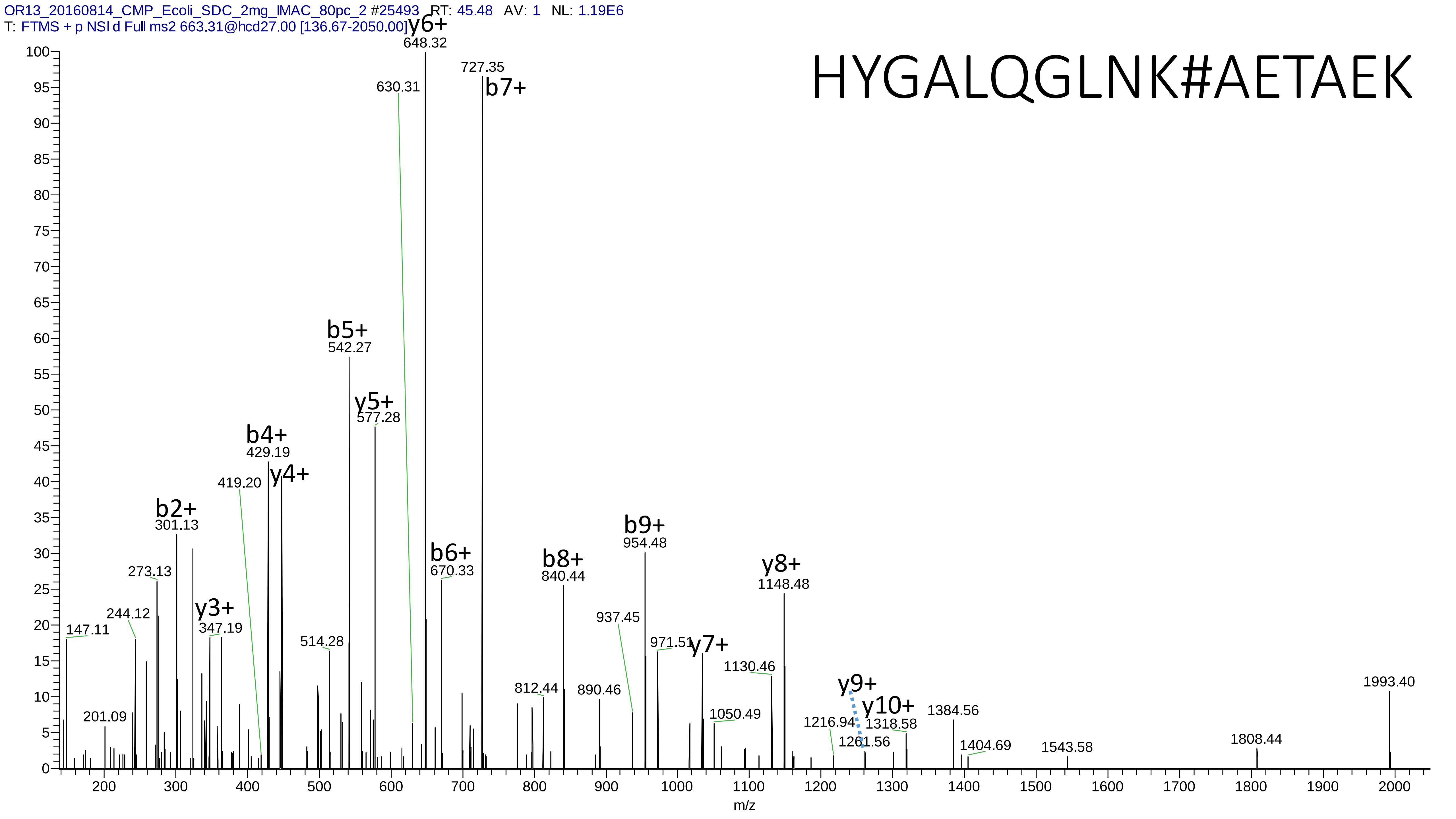
OR13_20160814_CMP_Ecoli_SDC_2mg_IMAC_80pc_2 #25493 RT: 45.48 AV: 1 NL: 1.19E6
T: FTMS + p NSI d Full ms2 663.31@hcd27.00 [136.67-2050.00]
HYGALQGLNK#AETAEK
y6+ 648.32
727.35 b7+
630.31
b5+ 542.27
y5+ 577.28
b4+ 429.19
y4+
419.20
b2+ 301.13
273.13
y3+ 347.19
244.12
147.11
201.09
514.28
b6+ 670.33
b8+ 840.44
b9+ 954.48
y8+ 1148.48
937.45
971.51
y7+
1130.46
812.44
890.46
1050.49
y9+
y10+
1216.94
1261.56
1318.58
1384.56
1404.69
1543.58
1808.44
1993.40
m/z

# 11

- Experimental mass: 229.0141
- Unimod information: PyridoxalPhosphate
- Possible composition: H(8) C(8) N O(5) P
- Sites in Unimod: K
- Sites observed: H;W;L;M
- Features: 61.8% (94/152) peptide contains H at the n-term; neutral loss of $H_3PO_4$ are observed at precursor and fragment ions

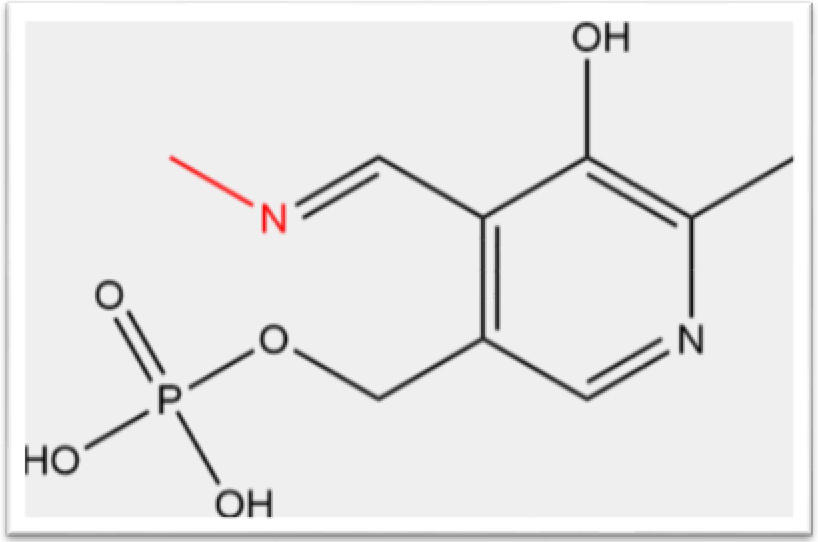

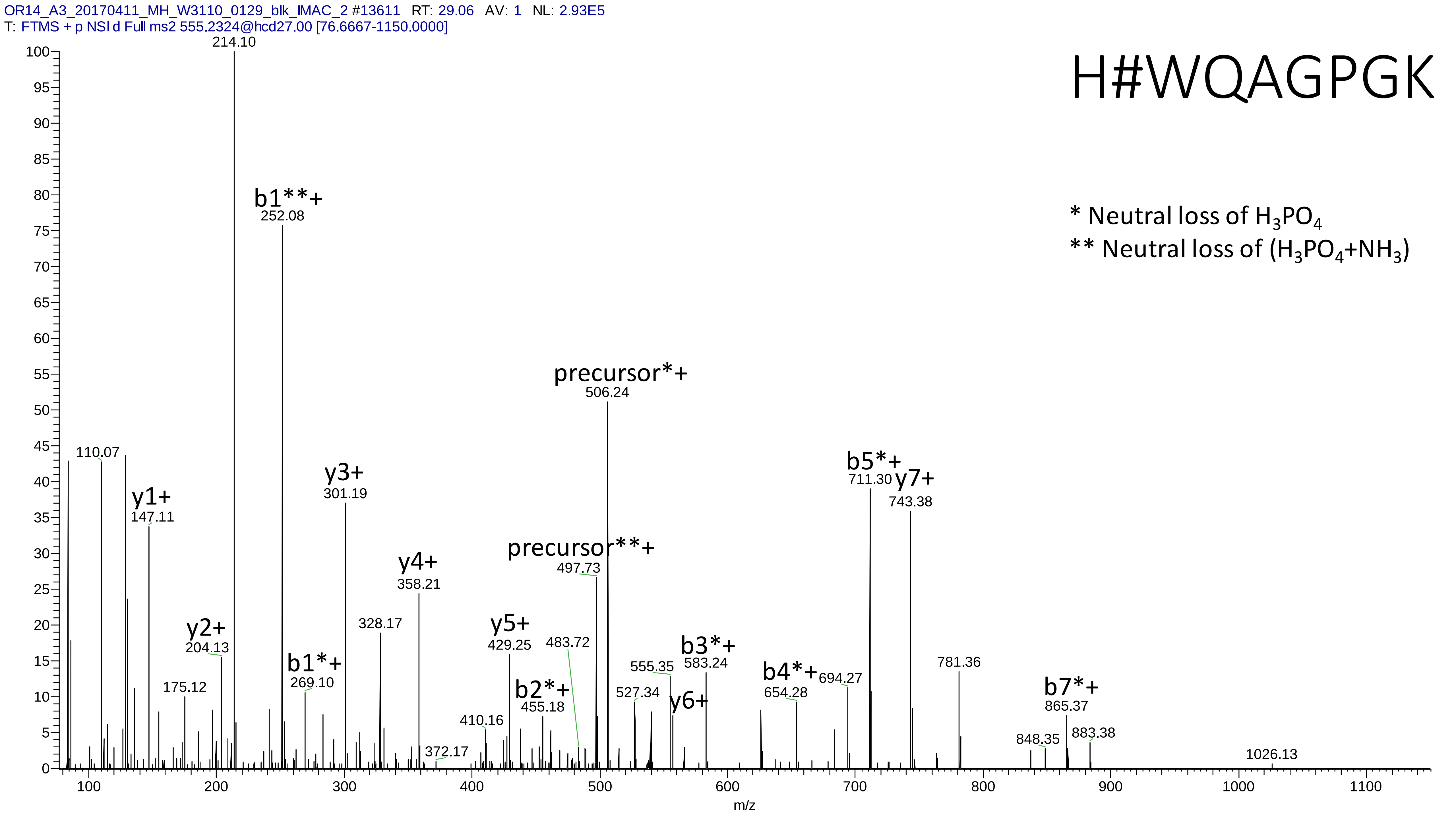
OR14_A3_20170411_MH_W3110_0129_blk_IMAC_2 #13611 RT: 29.06 AV: 1 NL: 2.93E5
T: FTMS + p NSI d Full ms2 555.2324@hcd27.00 [76.6667-1150.0000]
H#WQAGPGK
* Neutral loss of $H_3PO_4$
** Neutral loss of ($H_3PO_4$+$NH_3$)
214.10
b1**+ 252.08
110.07
y1+ 147.11
y2+ 204.13
175.12
b1*+ 269.10
y3+ 301.19
328.17
y4+ 358.21
372.17
410.16
y5+ 429.25
b2*+ 455.18
483.72
precursor**+ 497.73
precursor*+ 506.24
527.34
555.35
y6+
b3*+ 583.24
b4*+ 654.28
694.27
b5*+ 711.30
y7+ 743.38
781.36
848.35
b7*+ 865.37
883.38
1026.13
m/z

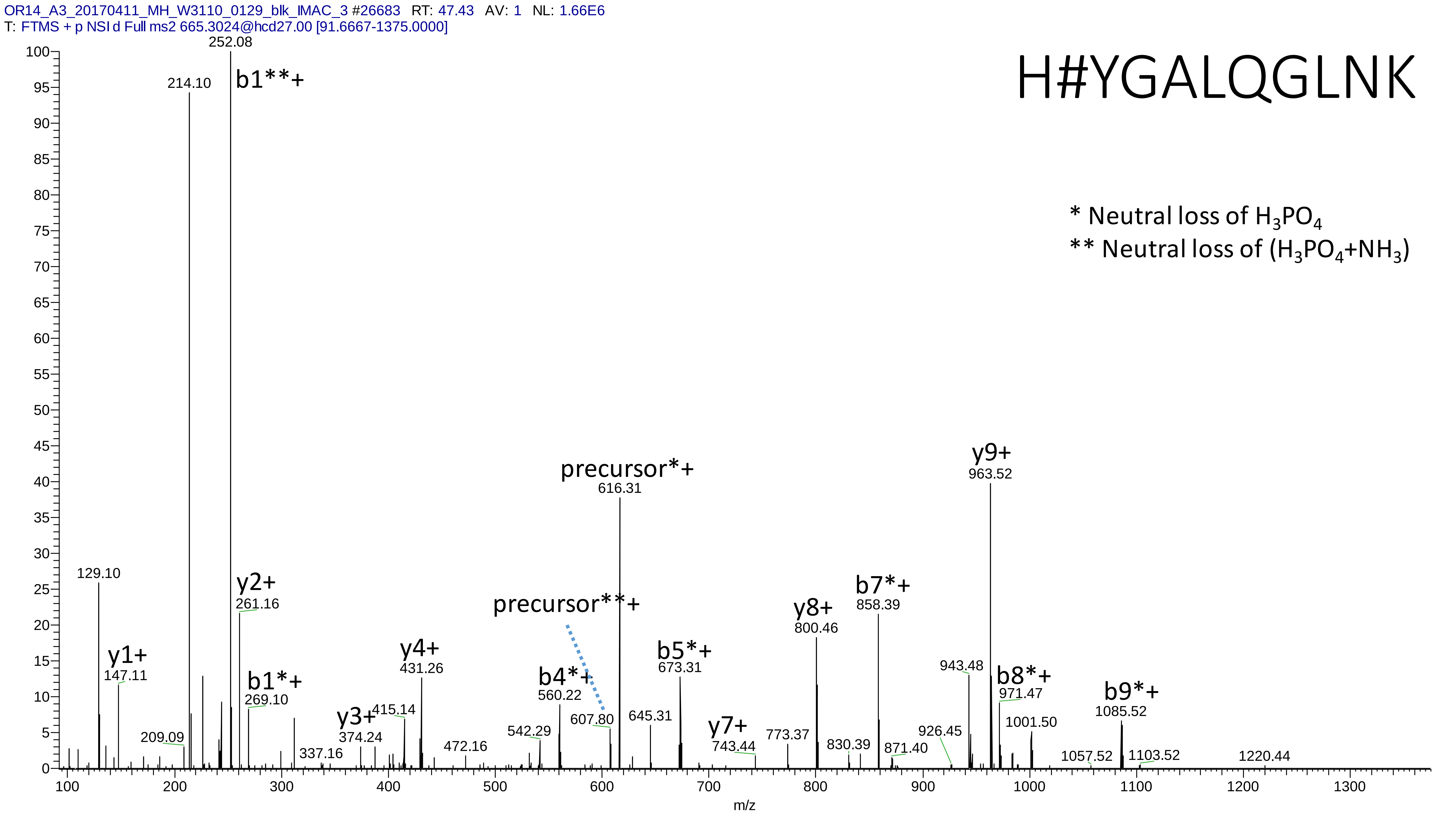
OR14_A3_20170411_MH_W3110_0129_blk_IMAC_3 #26683 RT: 47.43 AV: 1 NL: 1.66E6
T: FTMS + p NSI d Full ms2 665.3024@hcd27.00 [91.6667-1375.0000]
H#YGALQGLNK
* Neutral loss of H3PO4
** Neutral loss of (H3PO4+NH3)
252.08
b1**+
214.10
129.10
y1+
147.11
209.09
y2+
261.16
b1*+
269.10
337.16
y3+
374.24
415.14
y4+
431.26
472.16
542.29
b4*+
560.22
precursor**+
607.80
precursor*+
616.31
645.31
b5*+
673.31
y7+
743.44
773.37
y8+
800.46
830.39
b7*+
858.39
871.40
926.45
943.48
y9+
963.52
b8*+
971.47
1001.50
1057.52
b9*+
1085.52
1103.52
1220.44
m/z

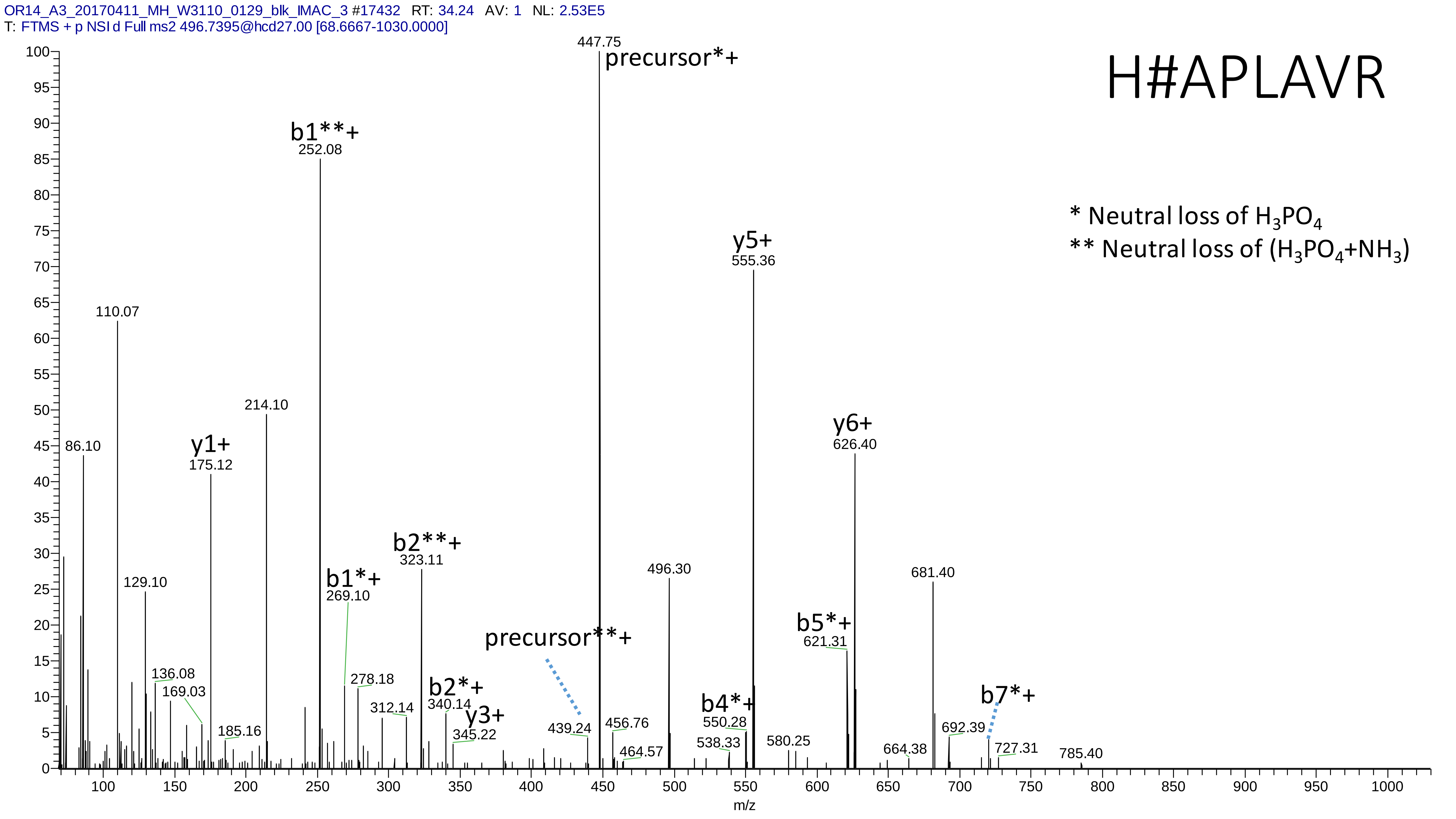
OR14_A3_20170411_MH_W3110_0129_blk_IMAC_3 #17432 RT: 34.24 AV: 1 NL: 2.53E5
T: FTMS + p NSI d Full ms2 496.7395@hcd27.00 [68.6667-1030.0000]
H#APLAVR
* Neutral loss of $H_3PO_4$
** Neutral loss of ($H_3PO_4$+$NH_3$)
precursor*+
447.75
b1**+
252.08
y5+
555.36
y6+
626.40
214.10
110.07
86.10
y1+
175.12
129.10
b1*+
269.10
b2**+
323.11
496.30
681.40
b5*+
621.31
precursor**+
136.08
169.03
185.16
278.18
312.14
b2*+
340.14
y3+
345.22
439.24
456.76
464.57
b4*+
550.28
538.33
580.25
664.38
b7*+
692.39
727.31
785.40
m/z

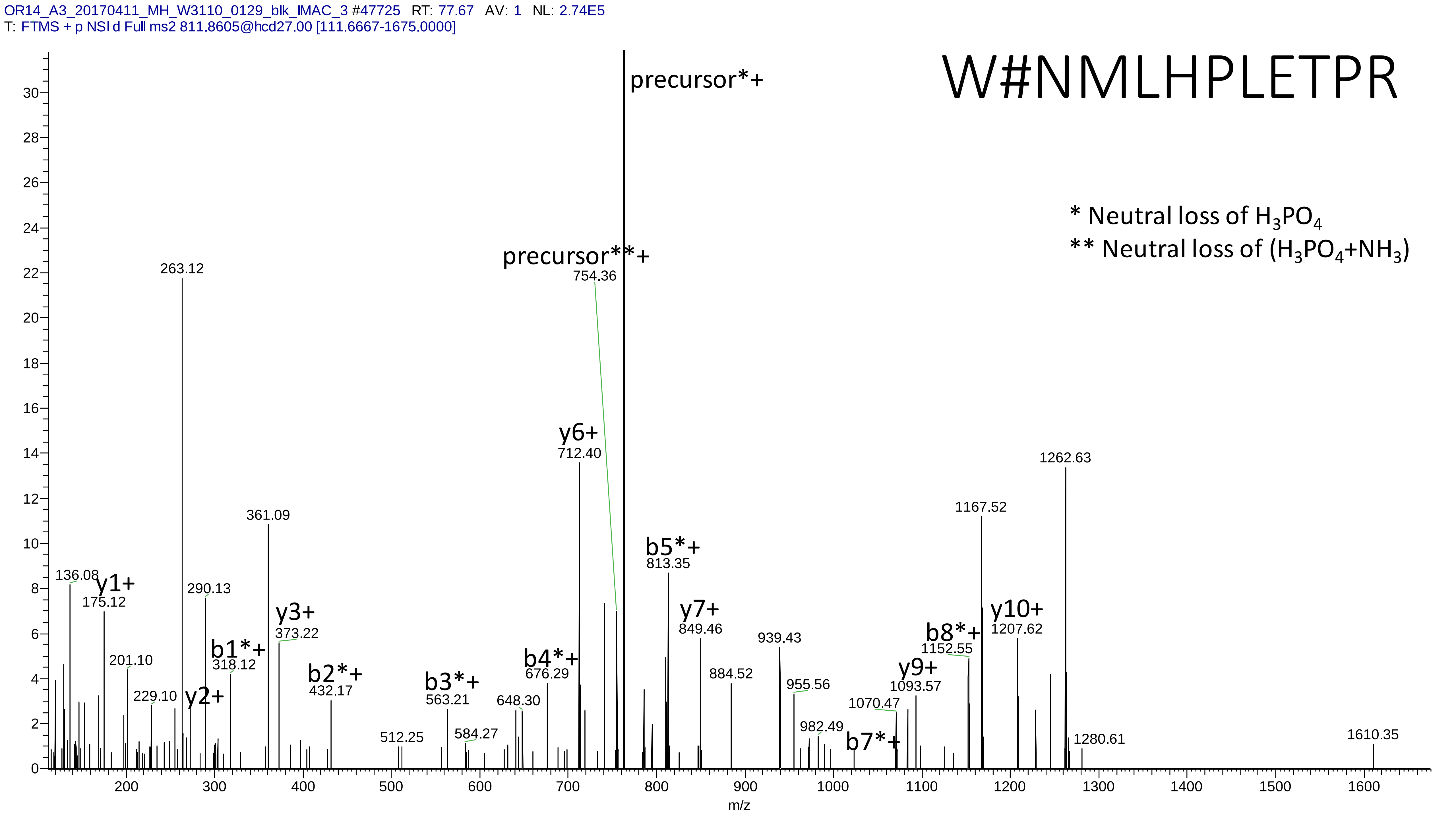
OR14_A3_20170411_MH_W3110_0129_blk_IMAC_3 #47725 RT: 77.67 AV: 1 NL: 2.74E5
T: FTMS + p NSI d Full ms2 811.8605@hcd27.00 [111.6667-1675.0000]
W#NMLHPLETPR
* Neutral loss of H3PO4
** Neutral loss of (H3PO4+NH3)
precursor*+
precursor**+
754.36
263.12
361.09
136.08
y1+
175.12
290.13
y3+
373.22
b1*+
318.12
201.10
229.10
y2+
b2*+
432.17
512.25
b3*+
563.21
584.27
648.30
b4*+
676.29
y6+
712.40
b5*+
813.35
y7+
849.46
884.52
939.43
955.56
982.49
1070.47
b7*+
y9+
1093.57
b8*+
1152.55
1167.52
y10+
1207.62
1262.63
1280.61
1610.35
m/z

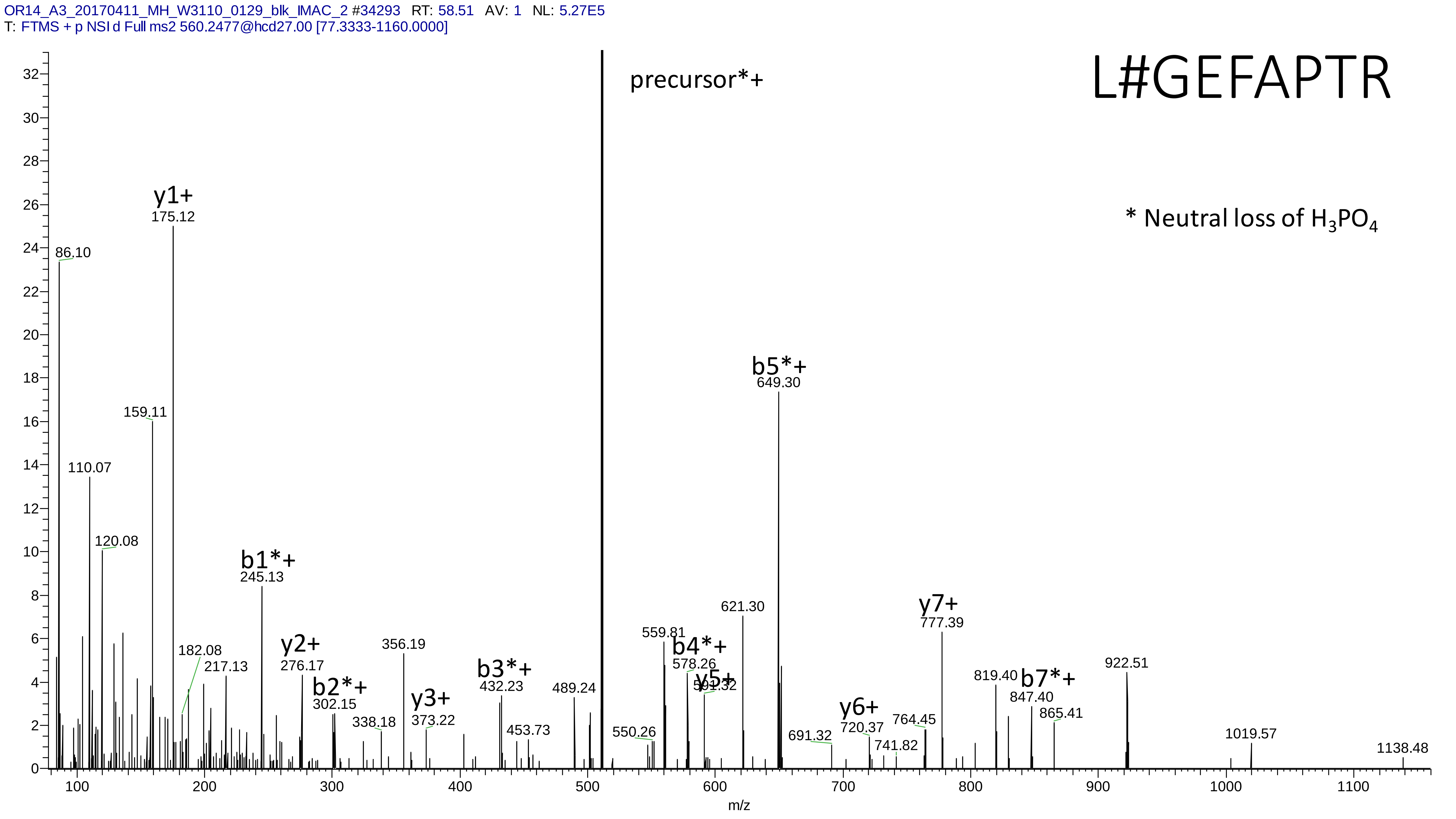
OR14_A3_20170411_MH_W3110_0129_blk_IMAC_2 #34293 RT: 58.51 AV: 1 NL: 5.27E5
T: FTMS + p NSI d Full ms2 560.2477@hcd27.00 [77.3333-1160.0000]
L#GEFAPTR
precursor*+
* Neutral loss of $H_3PO_4$
y1+
175.12
86.10
159.11
110.07
120.08
b1*+
245.13
182.08
217.13
y2+
276.17
b2*+
302.15
338.18
356.19
y3+
373.22
b3*+
432.23
453.73
489.24
550.26
559.81
b4*+
578.26
y5+
621.30
b5*+
649.30
691.32
y6+
720.37
741.82
764.45
y7+
777.39
819.40
b7*+
847.40
865.41
922.51
1019.57
1138.48
m/z

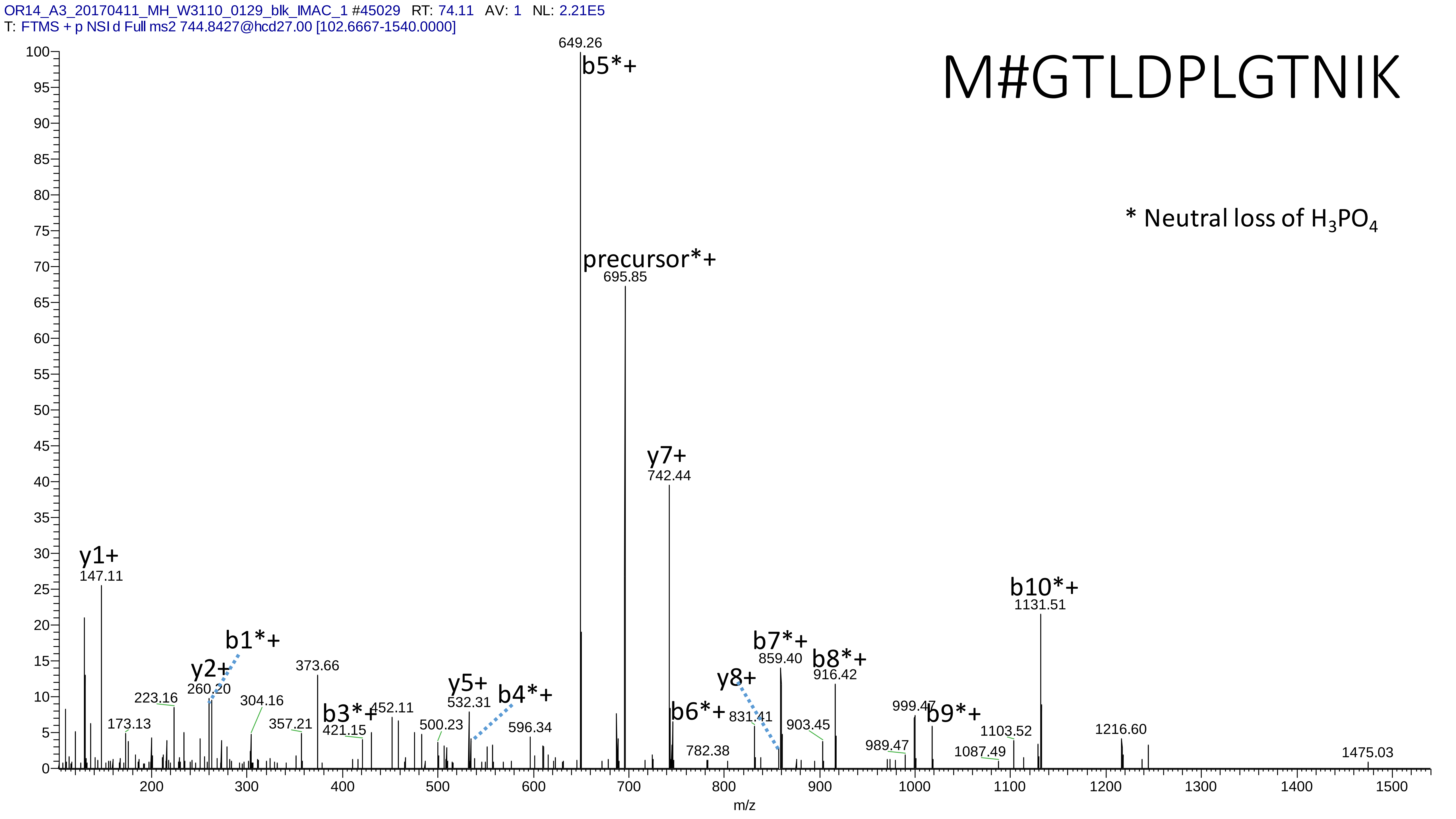

OR14_A3_20170411_MH_W3110_0129_blk_IMAC_1 #45029 RT: 74.11 AV: 1 NL: 2.21E5
T: FTMS + p NSI d Full ms2 744.8427@hcd27.00 [102.6667-1540.0000]
M#GTLDPLGTNIK
* Neutral loss of $H_3PO_4$
649.26
b5*+
precursor*+
695.85
y7+
742.44
y1+
147.11
b10*+
1131.51
b1*+
y2+
260.20
223.16
173.13
304.16
373.66
357.21
b3*+
421.15
452.11
y5+
532.31
b4*+
500.23
596.34
b6*+
y8+
831.41
782.38
b7*+
859.40
b8*+
916.42
903.45
999.47
b9*+
989.47
1103.52
1087.49
1216.60
1475.03
m/z

# 12

- Experimental mass: 57.0206
- Unimod information: Carbamidomethyl
- Possible composition: H(3) C(2) N O
- Sites in Unimod: C;K;N-term;H;D;E;S;T;Y;U;M
- Sites observed: H

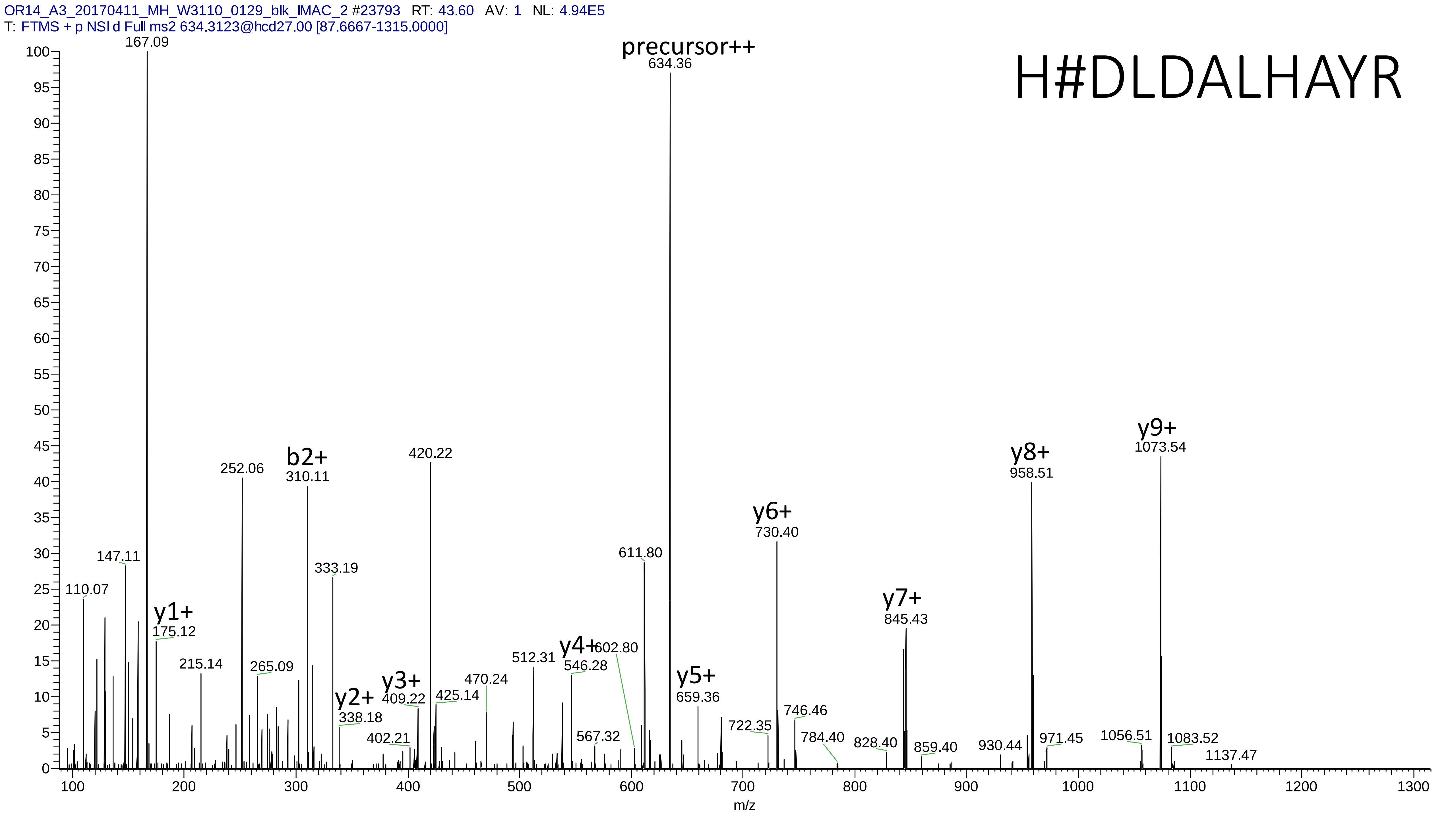

OR14_A3_20170411_MH_W3110_0129_blk_IMAC_2 #23793 RT: 43.60 AV: 1 NL: 4.94E5
T: FTMS + p NSI d Full ms2 634.3123@hcd27.00 [87.6667-1315.0000]
H#DLDALHAYR
precursor++
634.36
167.09
y9+
1073.54
y8+
958.51
b2+
310.11
252.06
420.22
y6+
730.40
611.80
333.19
147.11
110.07
y7+
845.43
y1+
175.12
215.14
265.09
y4+
546.28
602.80
512.31
470.24
y5+
659.36
y3+
409.22
425.14
y2+
338.18
746.46
722.35
402.21
567.32
784.40
828.40
859.40
930.44
971.45
1056.51
1083.52
1137.47
m/z

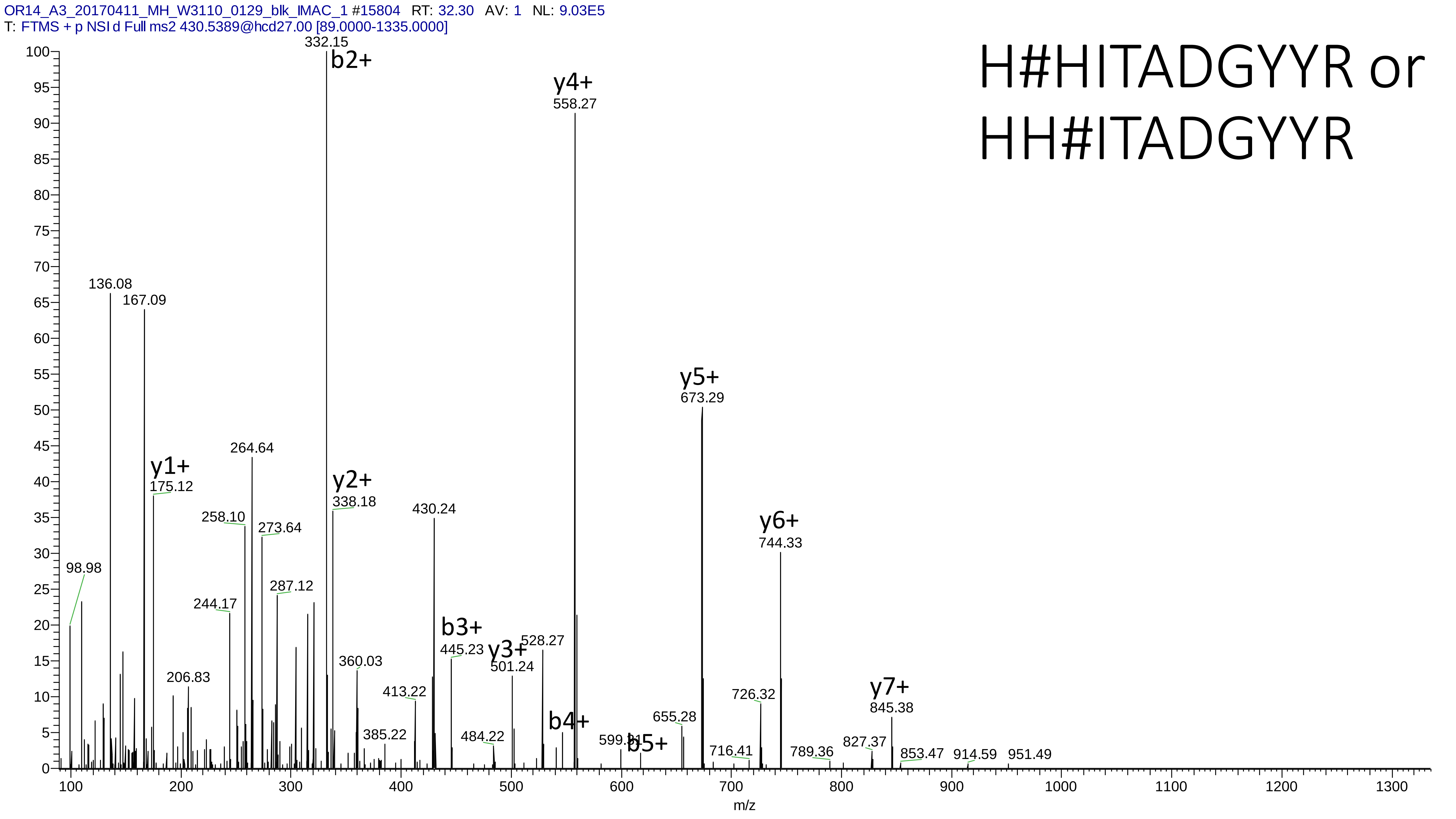
OR14_A3_20170411_MH_W3110_0129_blk_IMAC_1 #15804 RT: 32.30 AV: 1 NL: 9.03E5
T: FTMS + p NSI d Full ms2 430.5389@hcd27.00 [89.0000-1335.0000]
H#HITADGYYR or
HH#ITADGYYR
332.15
b2+
y4+
558.27
136.08
167.09
264.64
y1+
175.12
y2+
338.18
430.24
258.10
273.64
98.98
287.12
244.17
b3+
445.23
y3+
528.27
501.24
360.03
206.83
413.22
385.22
484.22
b4+
599.31
b5+
y5+
673.29
655.28
716.41
y6+
744.33
726.32
789.36
y7+
845.38
827.37
853.47
914.59
951.49
m/z

# 13

- Experimental mass: 387.0126
- Unimod information: not found
- Possible composition:
- Sites in Unimod: not found
- Sites observed: K
- Features: 97.8% (91/93) peptides contain a miss cleavage site of K; neutral loss of $H_3PO_4$ are observed at both precursor and fragment ions, structure unknown

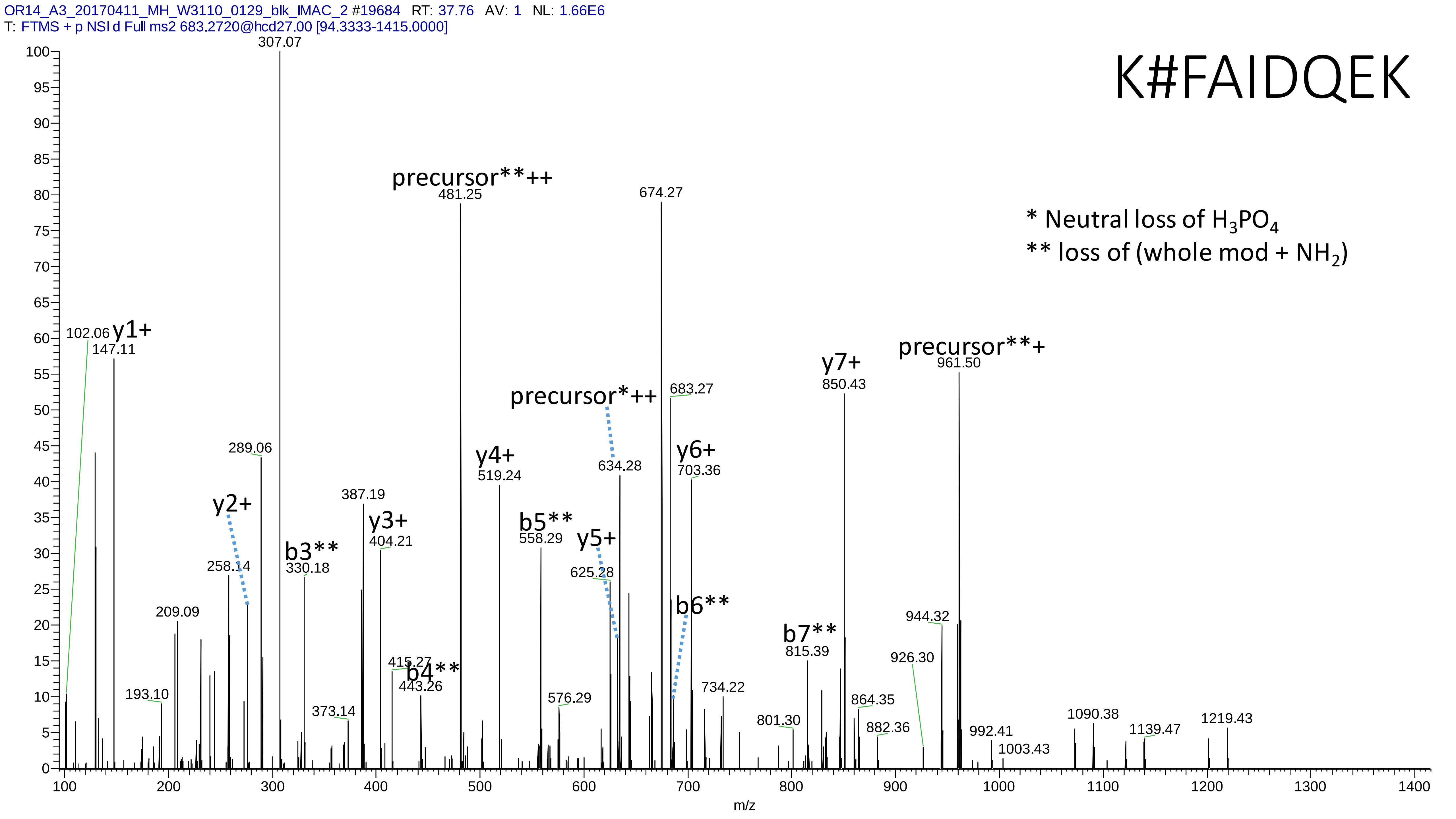
OR14_A3_20170411_MH_W3110_0129_blk_IMAC_2 #19684 RT: 37.76 AV: 1 NL: 1.66E6
T: FTMS + p NSI d Full ms2 683.2720@hcd27.00 [94.3333-1415.0000]
K#FAIDQEK
* Neutral loss of $H_3PO_4$
** loss of (whole mod + $NH_2$)
precursor**++
481.25
precursor**+
961.50
precursor*++
634.28
y1+
147.11
y2+
y3+
404.21
y4+
519.24
y5+
y6+
703.36
y7+
850.43
b3**
330.18
b4**
443.26
b5**
558.29
b6**
b7**
815.39
307.07
674.27
683.27
289.06
387.19
258.14
625.28
209.09
102.06
193.10
373.14
415.27
576.29
734.22
801.30
864.35
882.36
926.30
944.32
992.41
1003.43
1090.38
1139.47
1219.43
m/z

# 14

- Experimental mass: -113.0843
- Unimod information: not found
- Possible composition: Ile-loss or Leu-loss, [H(-11) C(-6) N(-1) O(-1)]
- Sites in Unimod: not found
- Sites observed: loss of L or I at n-term
- Features: 100.0% (131/131) peptides contain L or I at the peptide n-term

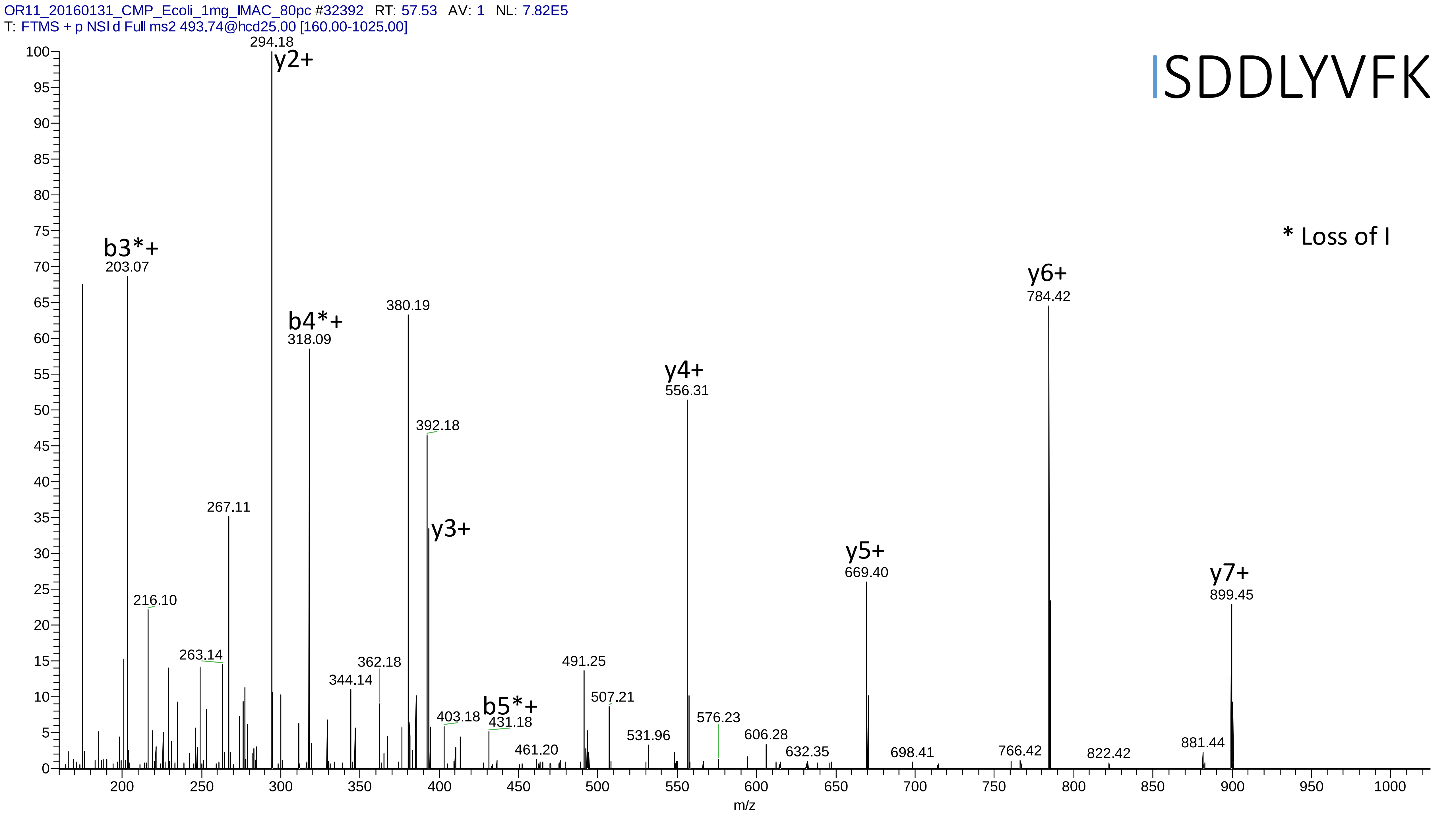
OR11_20160131_CMP_Ecoli_1mg_IMAC_80pc #32392 RT: 57.53 AV: 1 NL: 7.82E5
T: FTMS + p NSI d Full ms2 493.74@hcd25.00 [160.00-1025.00]
ISDDLYVFK
* Loss of I
y2+
294.18
b3*+
203.07
b4*+
318.09
380.19
392.18
y3+
267.11
216.10
263.14
362.18
344.14
b5*+
431.18
403.18
461.20
491.25
507.21
531.96
y4+
556.31
576.23
606.28
632.35
y5+
669.40
698.41
y6+
784.42
766.42
822.42
y7+
899.45
881.44
m/z

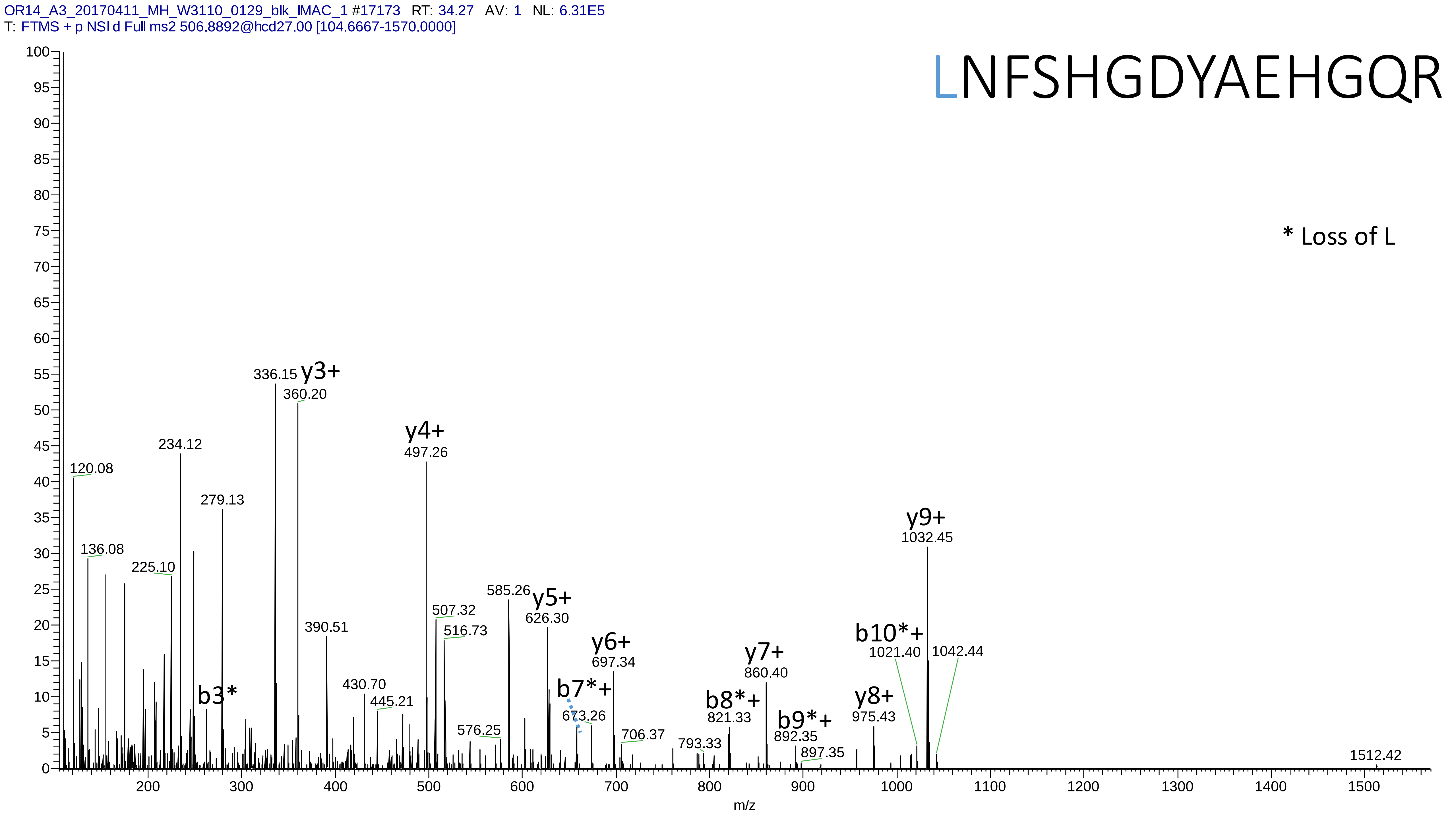

OR14_A3_20170411_MH_W3110_0129_blk_IMAC_1 #17173 RT: 34.27 AV: 1 NL: 6.31E5
T: FTMS + p NSI d Full ms2 506.8892@hcd27.00 [104.6667-1570.0000]
LNFSHGDYAEHGQR
* Loss of L
120.08
136.08
225.10
234.12
279.13
b3*
336.15
y3+
360.20
390.51
430.70
445.21
y4+
497.26
507.32
516.73
576.25
585.26
y5+
626.30
b7*+
673.26
y6+
697.34
706.37
793.33
b8*+
821.33
y7+
860.40
b9*+
892.35
897.35
y8+
975.43
b10*+
1021.40
y9+
1032.45
1042.44
1512.42
m/z

# 15

- Experimental mass: 248.0091
- Unimod information: not found
- Possible composition: H(12) C(4) N(1) O(7) P(2)
- Sites in Unimod: not found
- Sites observed: K
- Features: 98.7% (74/75) peptides contain a miss cleavage site K, neutral loss of $H_3PO_4$ are observed in precursor ions, neutral loss of $H_3PO_4$ are **not** observed in fragment ions, structure unknown

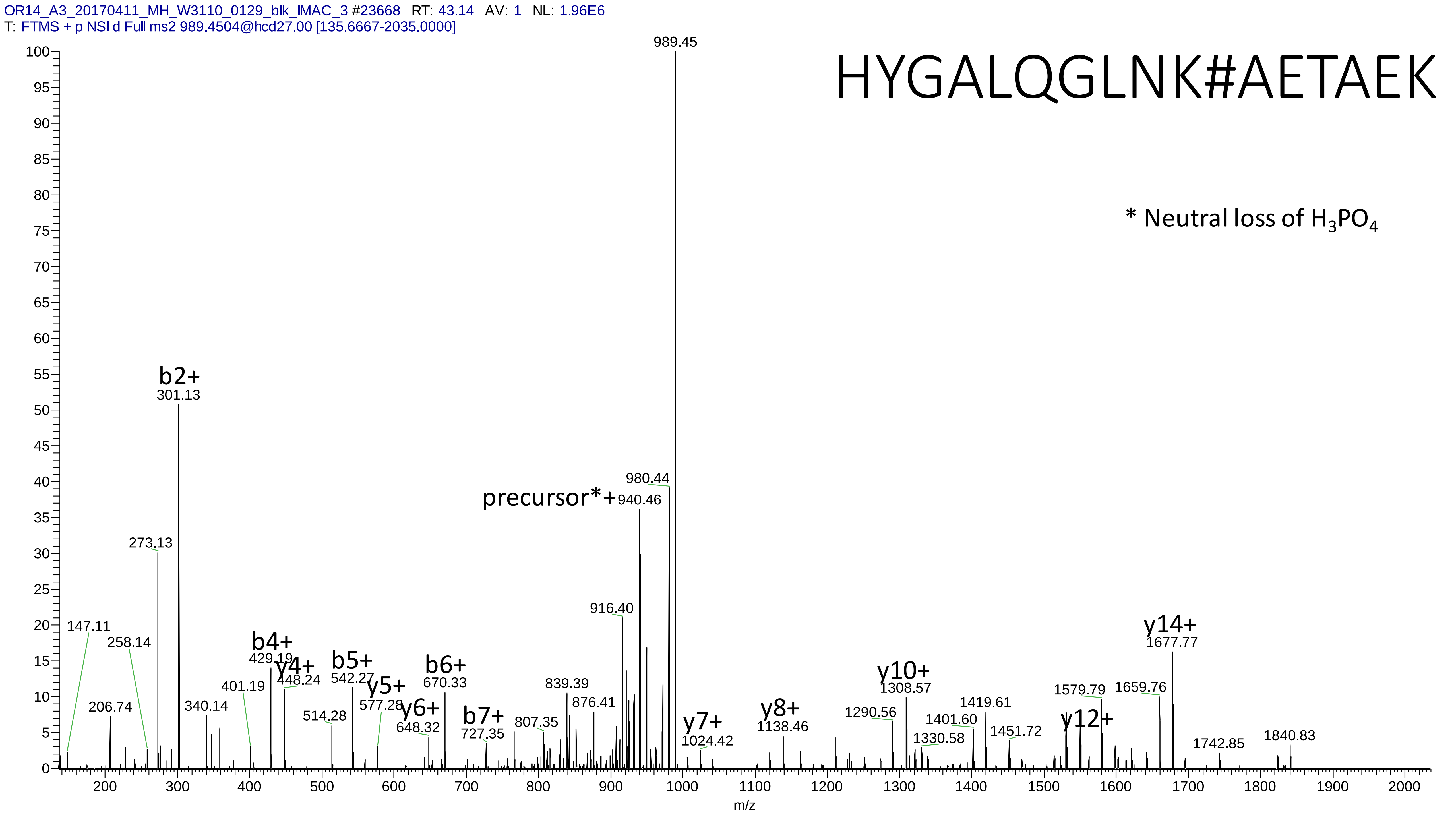
OR14_A3_20170411_MH_W3110_0129_blk_IMAC_3 #23668 RT: 43.14 AV: 1 NL: 1.96E6
T: FTMS + p NSI d Full ms2 989.4504@hcd27.00 [135.6667-2035.0000]
HYGALQGLNK#AETAEK
* Neutral loss of H3PO4
b2+ 301.13
273.13
147.11
258.14
206.74
340.14
401.19
b4+ 429.19
y4+ 448.24
514.28
b5+ 542.27
y5+ 577.28
y6+ 648.32
b6+ 670.33
b7+ 727.35
807.35
839.39
876.41
916.40
precursor*+ 940.46
980.44
989.45
y7+ 1024.42
y8+ 1138.46
1290.56
y10+ 1308.57
1330.58
1401.60
1419.61
1451.72
y12+
1579.79
1659.76
y14+ 1677.77
1742.85
1840.83
m/z

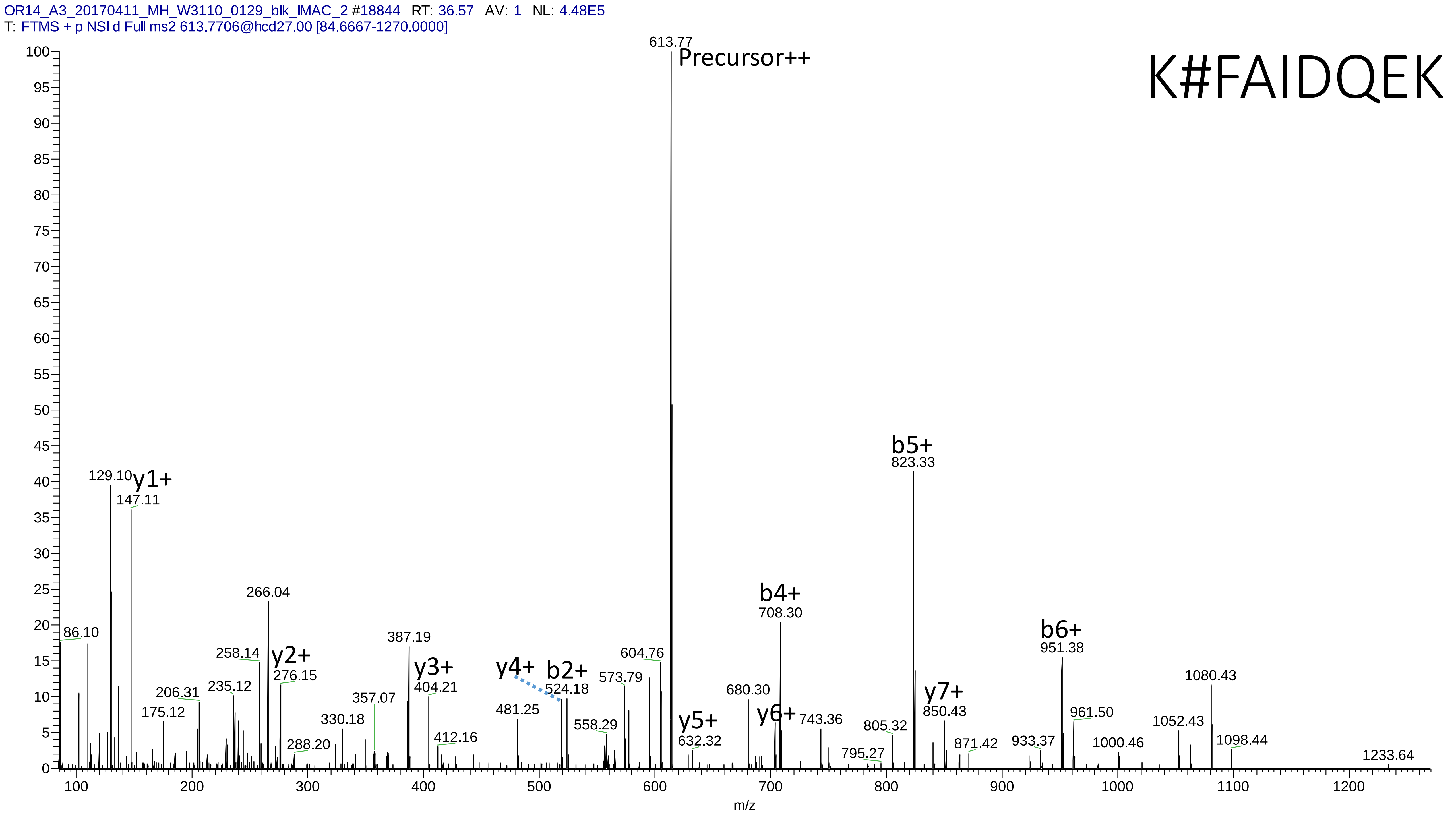

OR14_A3_20170411_MH_W3110_0129_blk_IMAC_2 #18844 RT: 36.57 AV: 1 NL: 4.48E5
T: FTMS + p NSI d Full ms2 613.7706@hcd27.00 [84.6667-1270.0000]
K#FAIDQEK
Precursor++
613.77
y1+
129.10
147.11
86.10
266.04
258.14
y2+
276.15
206.31
235.12
175.12
288.20
330.18
357.07
387.19
y3+
404.21
412.16
y4+
481.25
b2+
524.18
558.29
573.79
604.76
y5+
632.32
680.30
y6+
b4+
708.30
743.36
795.27
805.32
b5+
823.33
y7+
850.43
871.42
933.37
b6+
951.38
961.50
1000.46
1052.43
1080.43
1098.44
1233.64
m/z

# 16

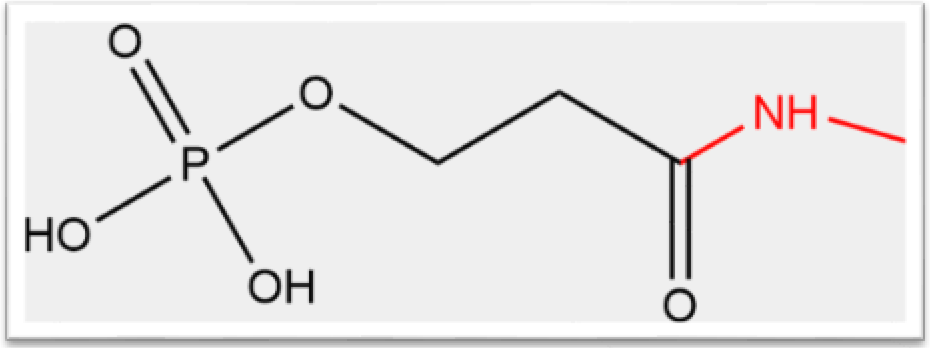


- Experimental mass: 20.9477 (151.98858 - 131.04048)
- Unimod information: not found
- Possible composition: loss of M at n-term, then add H(5) C(3) O(5) P
- Sites in Unimod: not found
- Sites observed: M at n-term
- Features: 98.0% (50/51) peptides start with M at protein n-term

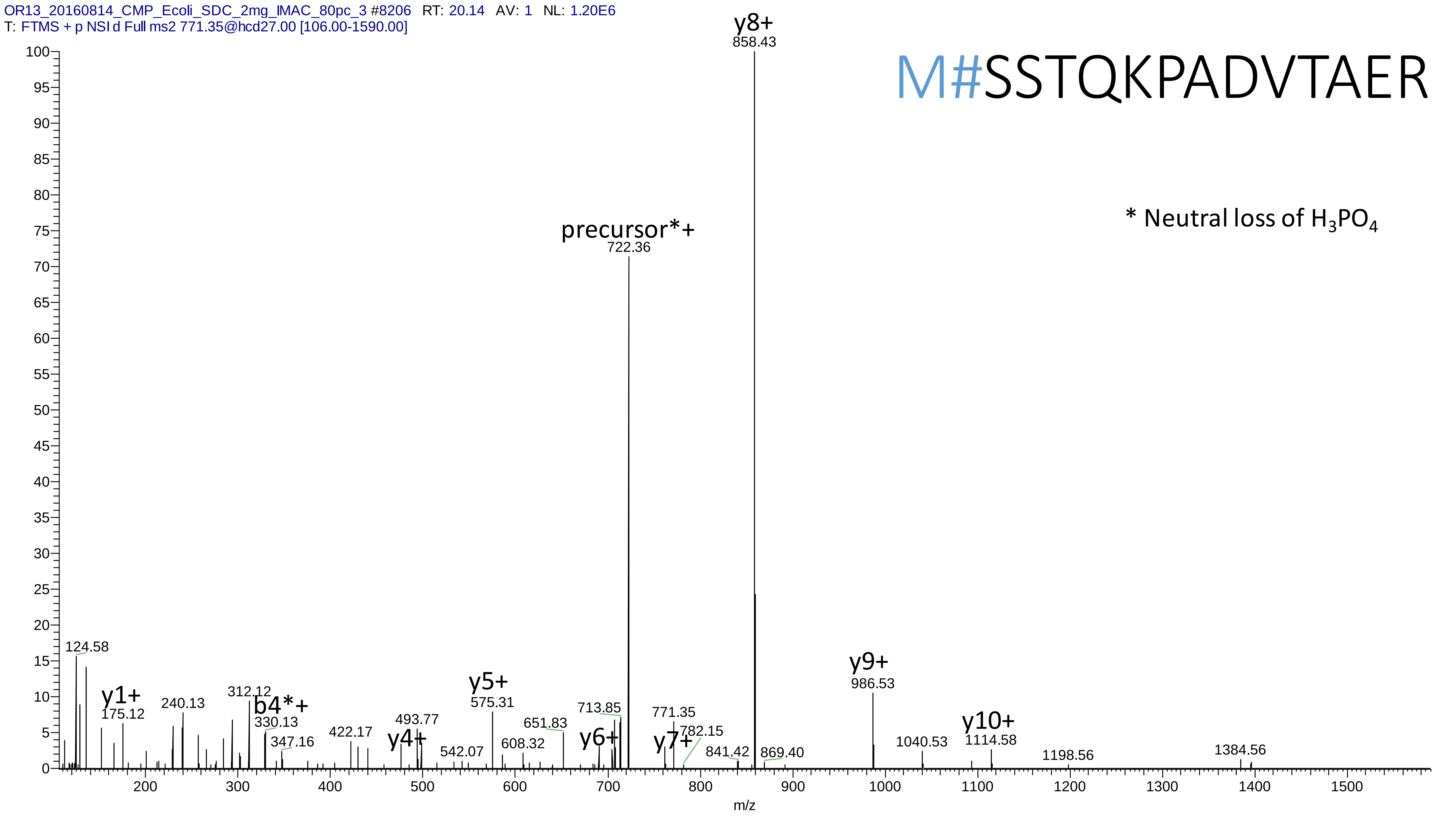
OR13_20160814_CMP_Ecoli_SDC_2mg_IMAC_80pc_3 #8206 RT: 20.14 AV: 1 NL: 1.20E6
T: FTMS + p NSI d Full ms2 771.35@hcd27.00 [106.00-1590.00]
M#SSTQKPADVTAER
* Neutral loss of H3PO4
y8+
858.43
precursor*+
722.36
124.58
y1+
175.12
240.13
312.12
b4*+
330.13
347.16
422.17
y4+
493.77
542.07
y5+
575.31
608.32
651.83
y6+
713.85
y7+
771.35
782.15
841.42
869.40
y9+
986.53
1040.53
y10+
1114.58
1198.56
1384.56
m/z

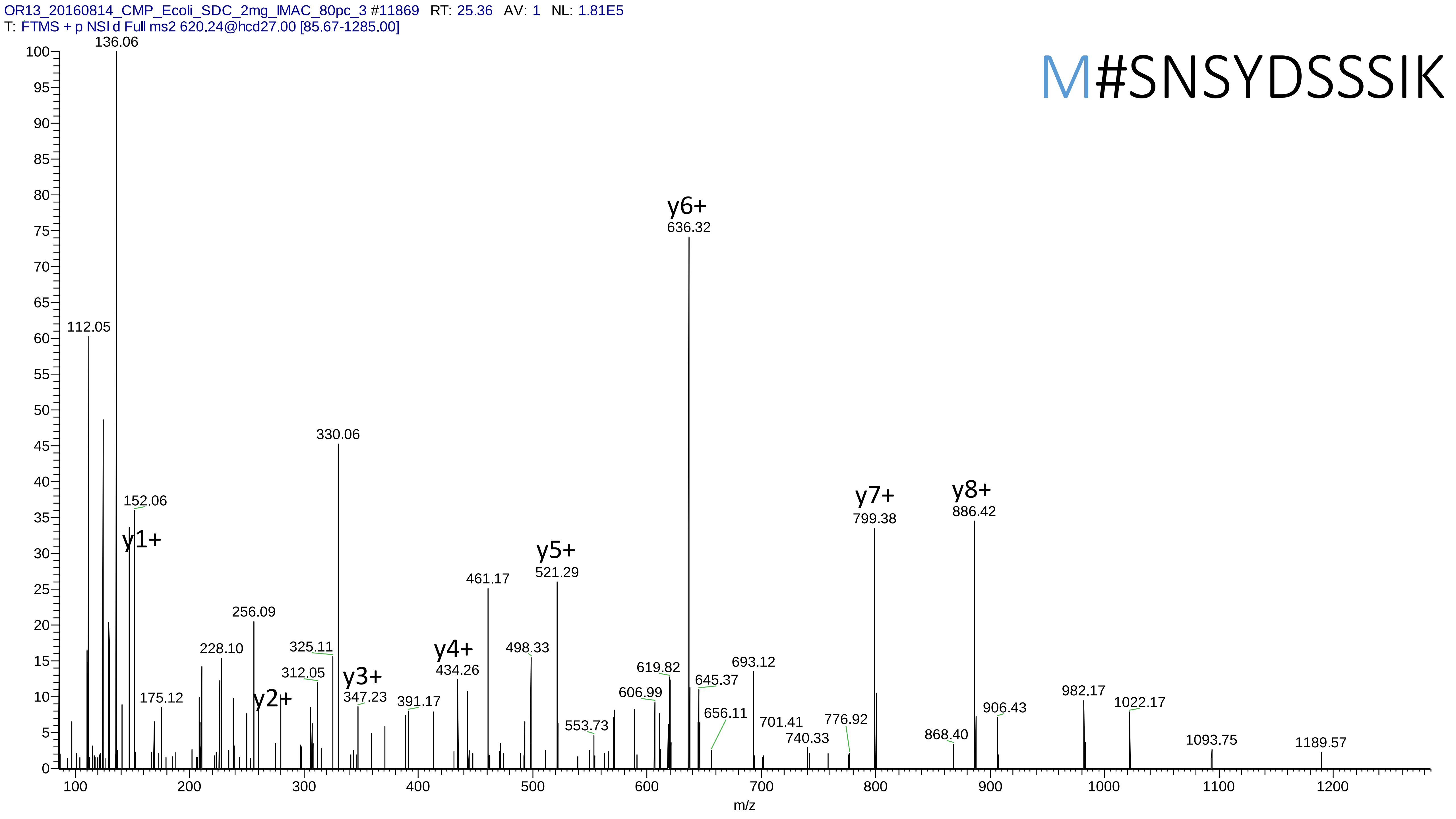
OR13_20160814_CMP_Ecoli_SDC_2mg_IMAC_80pc_3 #11869 RT: 25.36 AV: 1 NL: 1.81E5
T: FTMS + p NSI d Full ms2 620.24@hcd27.00 [85.67-1285.00]
M#SNSYDSSSIK
136.06
112.05
152.06
y1+
175.12
228.10
256.09
y2+
312.05
325.11
330.06
y3+
347.23
391.17
y4+
434.26
461.17
498.33
y5+
521.29
553.73
606.99
619.82
y6+
636.32
645.37
656.11
693.12
701.41
740.33
776.92
y7+
799.38
868.40
y8+
886.42
906.43
982.17
1022.17
1093.75
1189.57
m/z

# 17

- Experimental mass: 15.994
- Unimod information: oxidation
- Possible composition: O
- Sites in Unimod: D;K;N;P;F;Y;R;M;C;H;W;G;U;E;I;L;Q;S;T;V
- Sites observed: Y;W

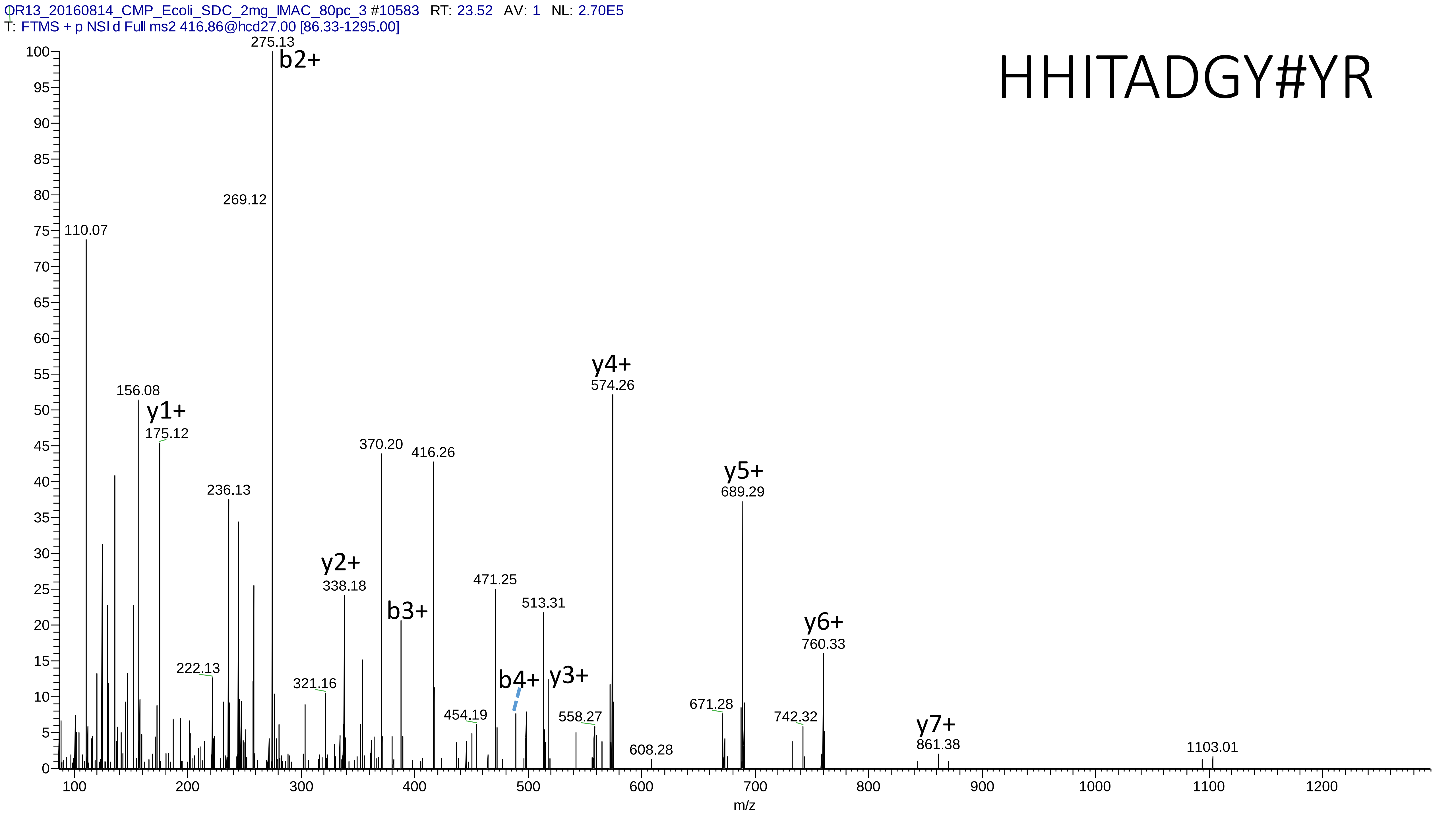

OR13_20160814_CMP_Ecoli_SDC_2mg_IMAC_80pc_3 #10583 RT: 23.52 AV: 1 NL: 2.70E5
T: FTMS + p NSI d Full ms2 416.86@hcd27.00 [86.33-1295.00]
HHITADGY#YR
b2+
y1+
y2+
b3+
b4+
y3+
y4+
y5+
y6+
y7+
275.13
269.12
110.07
156.08
175.12
236.13
222.13
321.16
338.18
370.20
416.26
454.19
471.25
513.31
558.27
574.26
608.28
671.28
689.29
742.32
760.33
861.38
1103.01
m/z

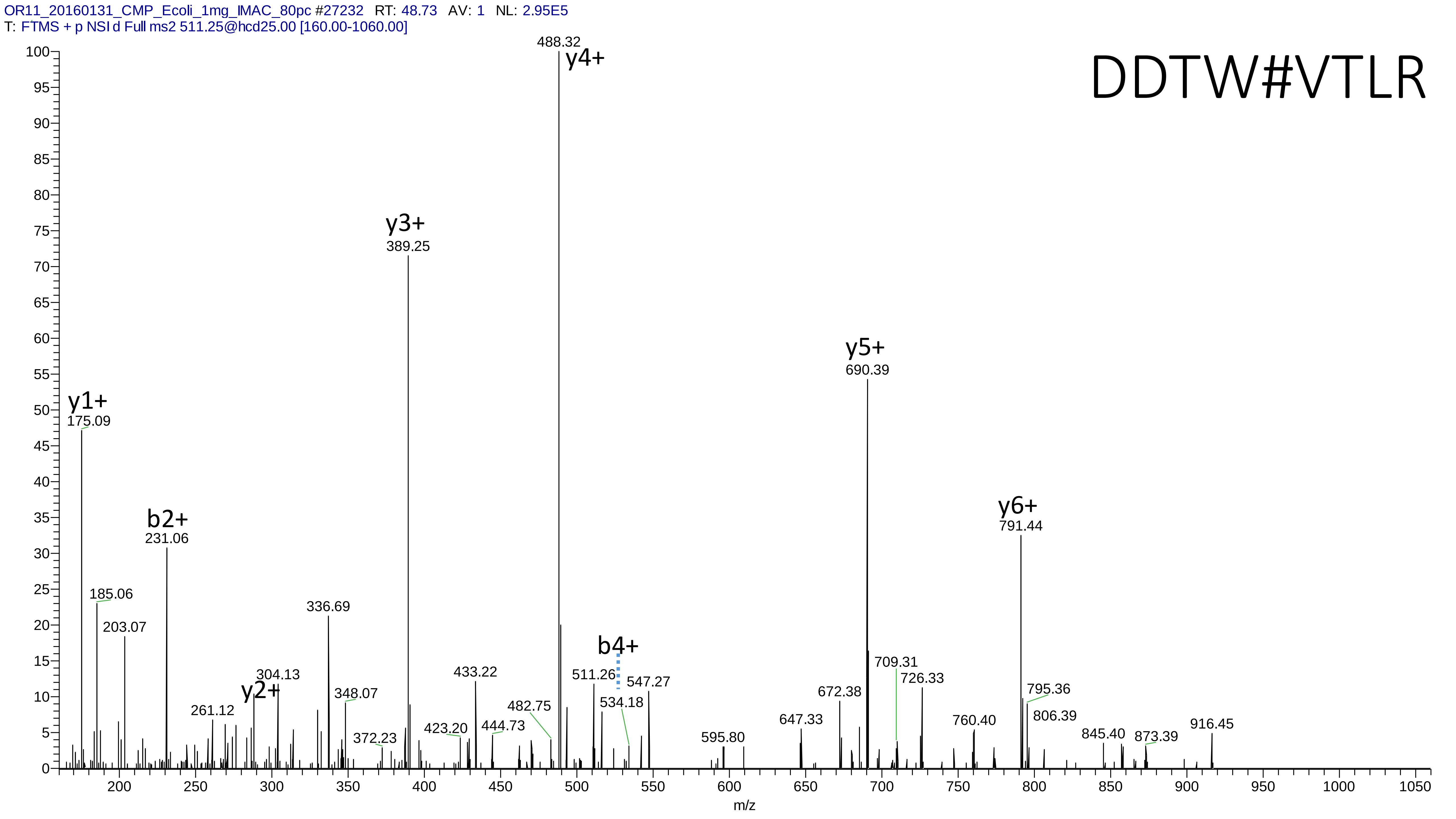

OR11_20160131_CMP_Ecoli_1mg_IMAC_80pc #27232 RT: 48.73 AV: 1 NL: 2.95E5
T: FTMS + p NSI d Full ms2 511.25@hcd25.00 [160.00-1060.00]
DDTW#VTLR
488.32
y4+
y3+
389.25
y5+
690.39
y1+
175.09
y6+
791.44
b2+
231.06
185.06
203.07
336.69
b4+
304.13
y2+
348.07
433.22
511.26
547.27
709.31
726.33
672.38
795.36
534.18
482.75
261.12
806.39
647.33
760.40
444.73
423.20
372.23
595.80
845.40
873.39
916.45
m/z

# 18-1

- Experimental mass: 183.034
- Unimod information: Aminoethylbenzenesulfonylation (AEBS)
- Possible composition: H(9) C(8) N O(2) S
- Sites in Unimod: H;K;S;Y;n-term
- Sites observed: Y
- Features: neutral loss of $H_3PO_4$ are **not** observed, so it's possible that two different modifications exist with very close Δmass

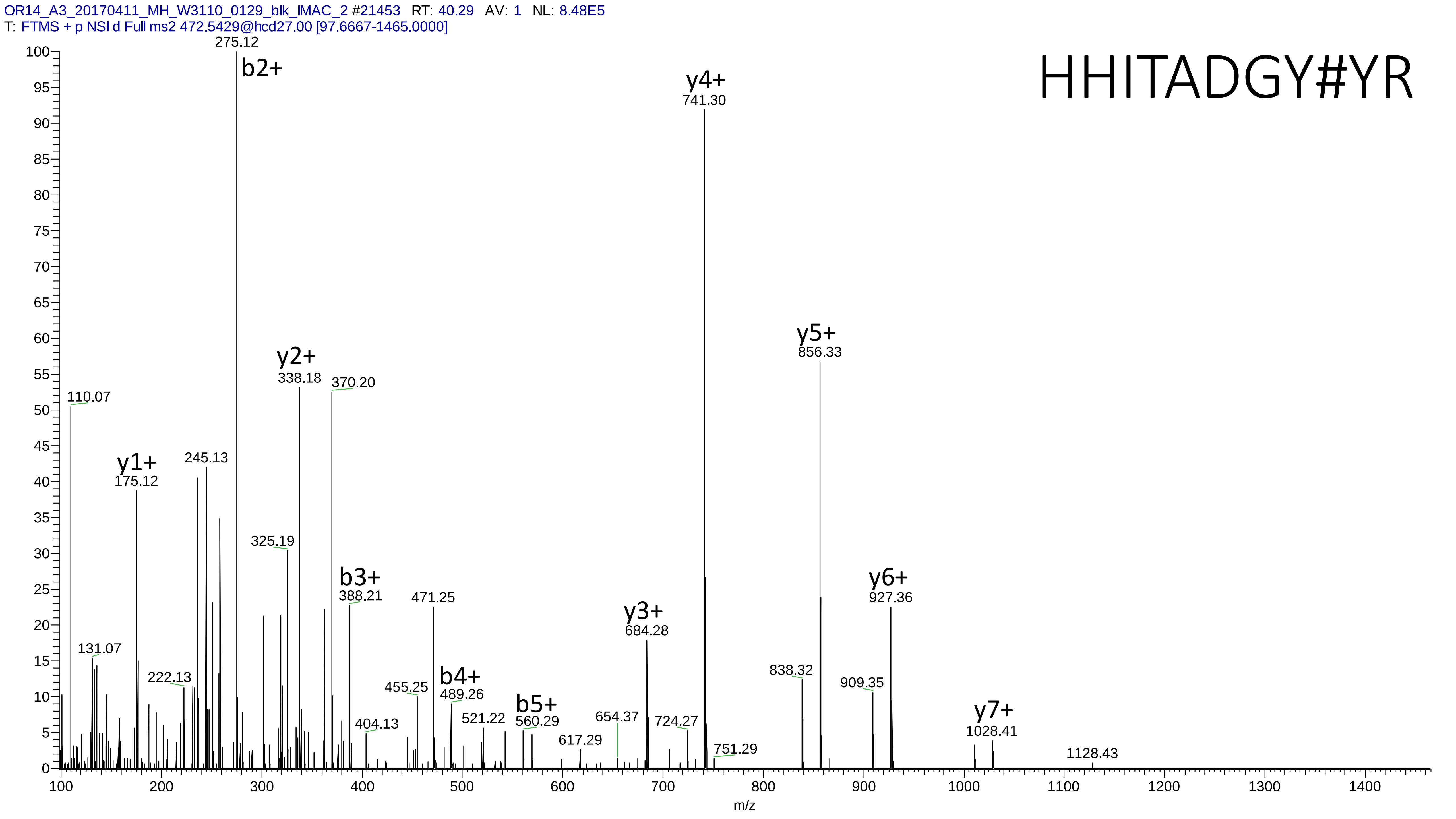

OR14_A3_20170411_MH_W3110_0129_blk_IMAC_2 #21453 RT: 40.29 AV: 1 NL: 8.48E5
T: FTMS + p NSI d Full ms2 472.5429@hcd27.00 [97.6667-1465.0000]
HHITADGY#YR
275.12
b2+
y4+
741.30
y5+
856.33
y2+
338.18
370.20
110.07
y1+
175.12
245.13
325.19
b3+
388.21
471.25
y6+
927.36
y3+
684.28
131.07
222.13
455.25
b4+
489.26
838.32
909.35
b5+
560.29
521.22
654.37
724.27
617.29
404.13
751.29
y7+
1028.41
1128.43
m/z

# 18-2

- Experimental mass: 183.034
- Unimod information: not found
- Possible composition: H(10) C(4) N O(5) P
- Sites in Unimod: not found
- Sites observed: H;K
- Features: neutral loss of $H_3PO_4$ are observed in both precursor and fragment ions, so it's possible that two different modifications exist with very close Δmass

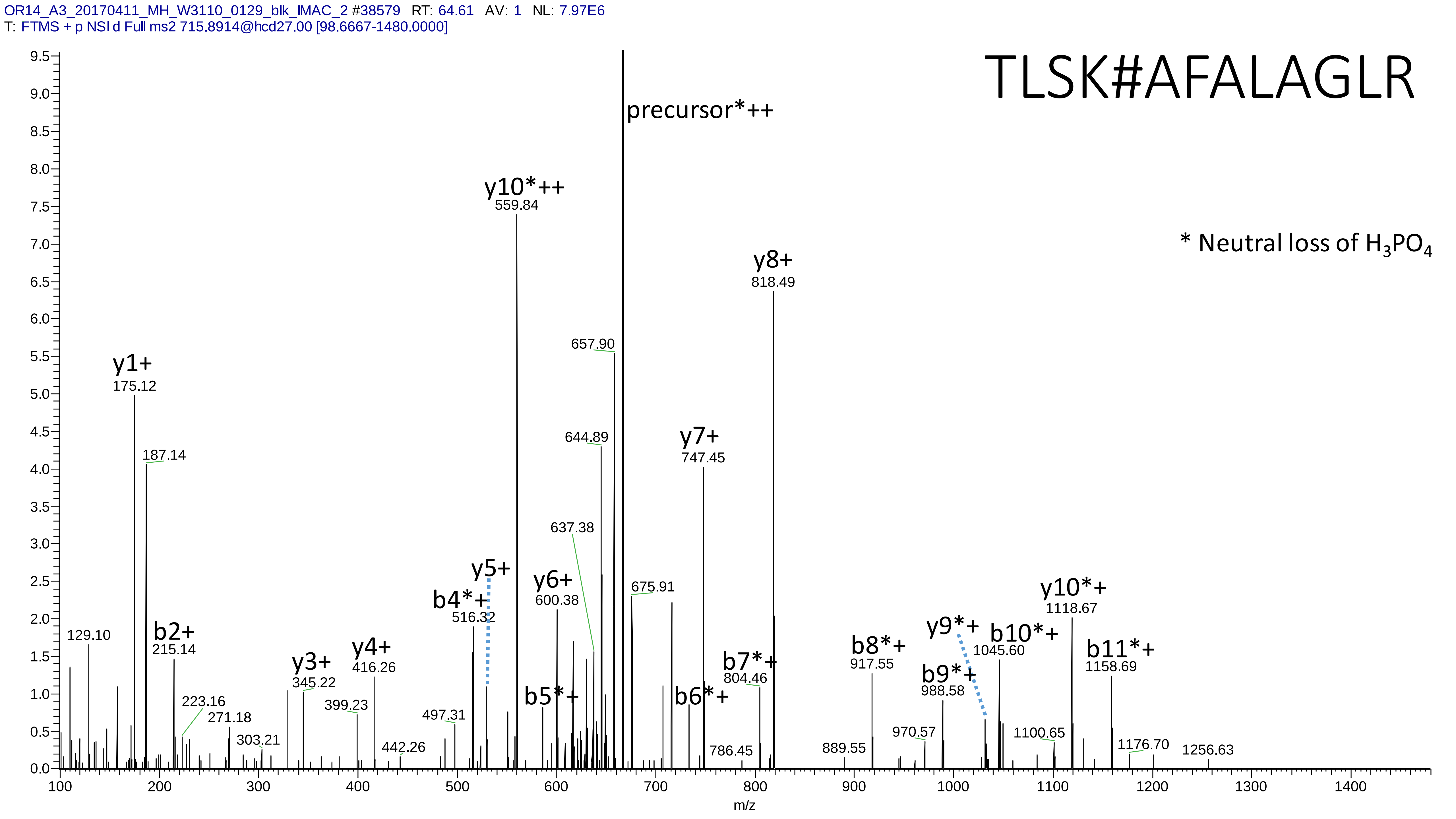
OR14_A3_20170411_MH_W3110_0129_blk_IMAC_2 #38579 RT: 64.61 AV: 1 NL: 7.97E6
T: FTMS + p NSI d Full ms2 715.8914@hcd27.00 [98.6667-1480.0000]
TLSK#AFALAGLR
* Neutral loss of H3PO4
precursor*++
y10*++ 559.84
y8+ 818.49
657.90
644.89
y7+ 747.45
y1+ 175.12
187.14
637.38
y5+
y6+ 600.38
b4*+ 516.32
675.91
y10*+ 1118.67
y9*+
b10*+ 1045.60
b11*+ 1158.69
b8*+ 917.55
b9*+ 988.58
b7*+ 804.46
b6*+
b5*+
b2+ 215.14
129.10
y4+ 416.26
y3+ 345.22
223.16
271.18
399.23
303.21
497.31
442.26
786.45
889.55
970.57
1100.65
1176.70
1256.63
m/z

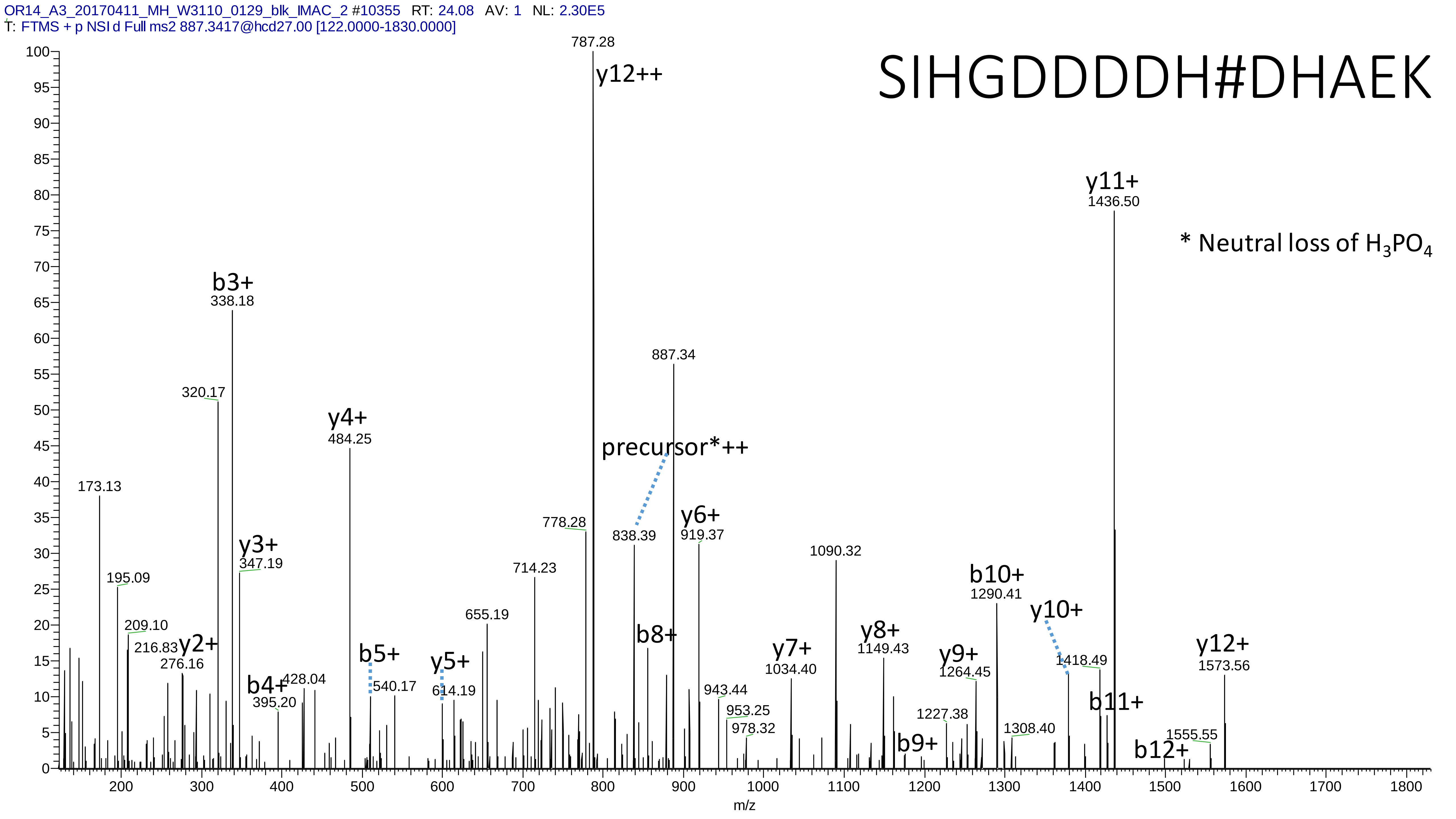

OR14_A3_20170411_MH_W3110_0129_blk_IMAC_2 #10355 RT: 24.08 AV: 1 NL: 2.30E5
T: FTMS + p NSI d Full ms2 887.3417@hcd27.00 [122.0000-1830.0000]
SIHGDDDDH#DHAEK
* Neutral loss of H3PO4
787.28
y12++
y11+
1436.50
b3+
338.18
320.17
y4+
484.25
887.34
precursor*++
173.13
778.28
838.39
y6+
919.37
y3+
347.19
195.09
714.23
1090.32
b10+
1290.41
y10+
209.10
216.83
y2+
276.16
655.19
b8+
y8+
1149.43
y7+
1034.40
y12+
1573.56
b5+
y5+
y9+
1264.45
1418.49
b4+
428.04
540.17
614.19
943.44
395.20
953.25
978.32
b11+
1227.38
1308.40
b9+
1555.55
b12+
m/z

# 19

- Experimental mass: 181.9984
- Unimod information: not found
- Possible composition: H(7)C(4)O(6)P(1)
- Sites in Unimod: not found
- Sites observed: K;N-term
- Features: 19.7% (12/61) peptides contain a miss cleavage site K, 41.0% (25/61) peptides start with the first amino acid in the proteins, 34.4% (21/61) peptides start with the second amino acid in the proteins, in total 75.4% (46/61) peptides contains protein n-term; neutral loss of $H_3PO_4$ are observed

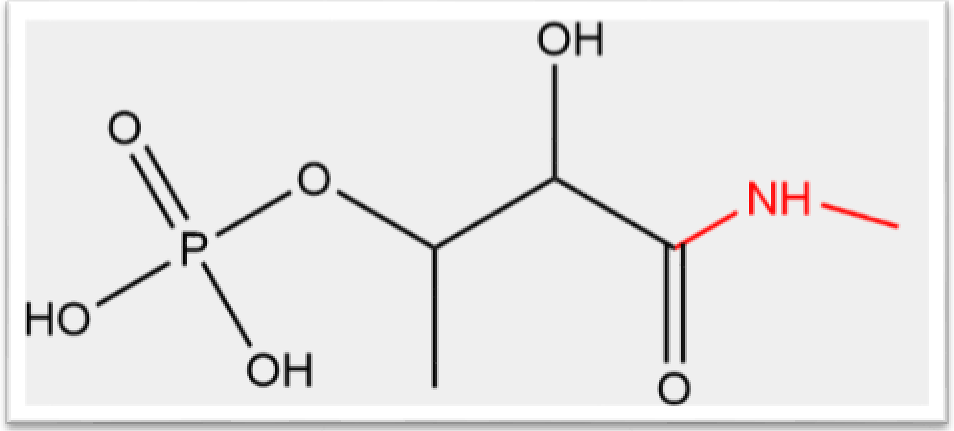

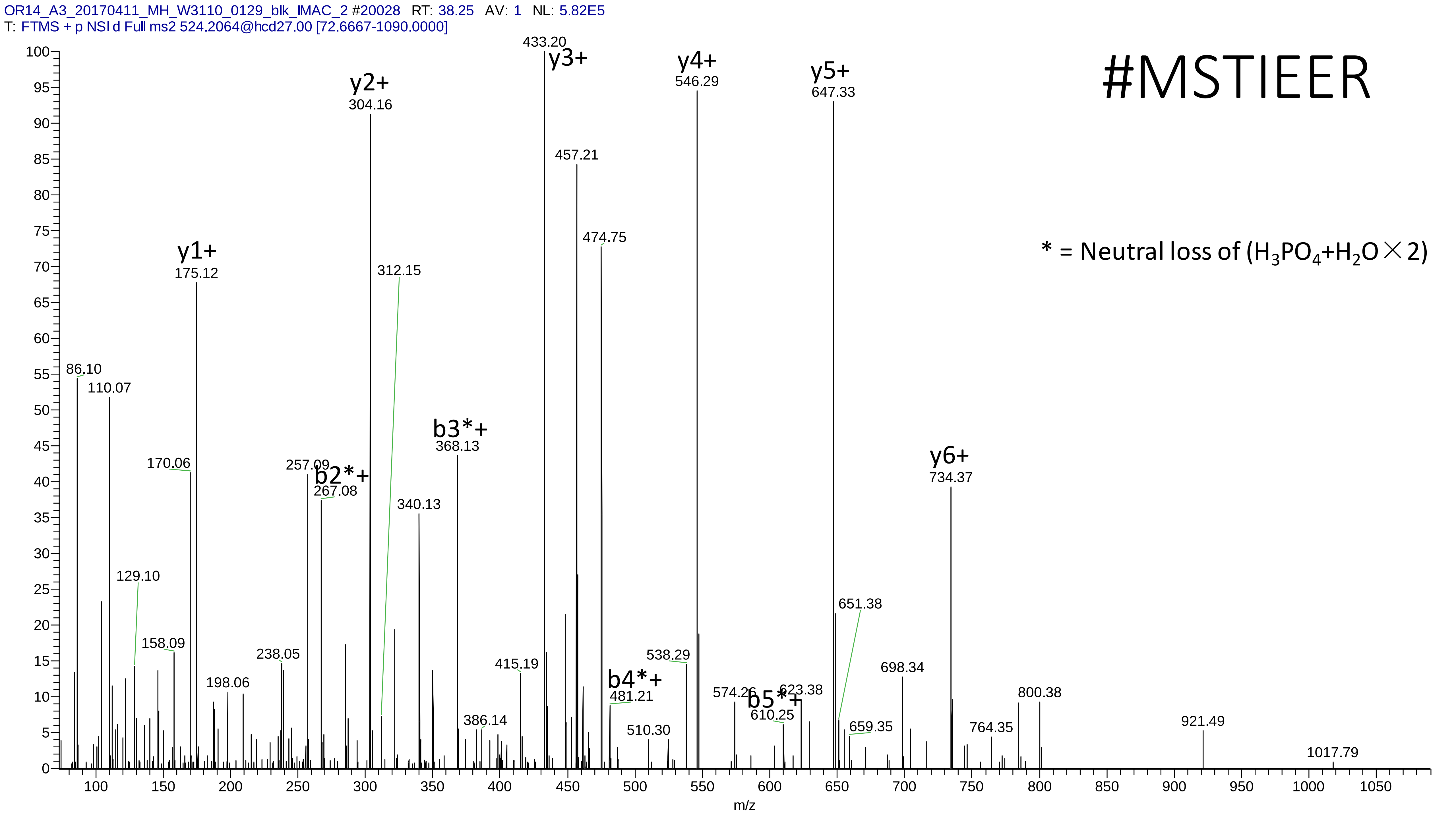
OR14_A3_20170411_MH_W3110_0129_blk_IMAC_2 #20028 RT: 38.25 AV: 1 NL: 5.82E5
T: FTMS + p NSI d Full ms2 524.2064@hcd27.00 [72.6667-1090.0000]
#MSTIEER
* = Neutral loss of ($H_3PO_4$+$H_2O$ × 2)
y1+ 175.12
y2+ 304.16
y3+ 433.20
y4+ 546.29
y5+ 647.33
y6+ 734.37
b2*+ 267.08
b3*+ 368.13
b4*+ 481.21
b5*+ 610.25
86.10
110.07
129.10
158.09
170.06
198.06
238.05
257.09
312.15
340.13
386.14
415.19
457.21
474.75
510.30
538.29
574.26
623.38
651.38
659.35
698.34
764.35
800.38
921.49
1017.79
m/z

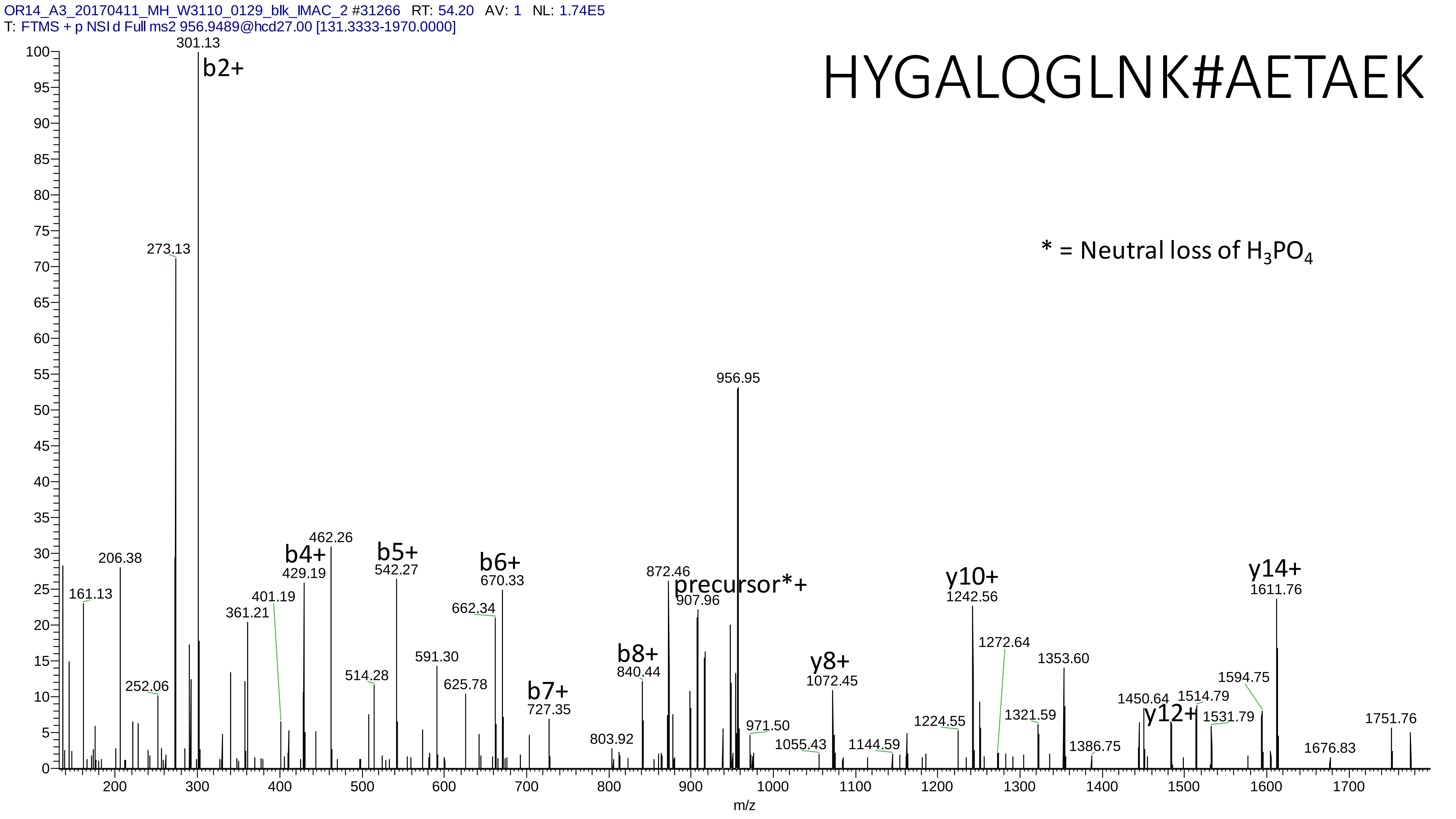
OR14_A3_20170411_MH_W3110_0129_blk_IMAC_2 #31266 RT: 54.20 AV: 1 NL: 1.74E5
T: FTMS + p NSI d Full ms2 956.9489@hcd27.00 [131.3333-1970.0000]
HYGALQGLNK#AETAEK
* = Neutral loss of $H_3PO_4$
b2+
301.13
273.13
206.38
161.13
252.06
361.21
401.19
b4+
429.19
462.26
514.28
b5+
542.27
591.30
625.78
662.34
b6+
670.33
b7+
727.35
803.92
b8+
840.44
872.46
precursor*+
907.96
956.95
971.50
1055.43
y8+
1072.45
1144.59
1224.55
y10+
1242.56
1272.64
1321.59
1353.60
1386.75
1450.64
y12+
1514.79
1531.79
1594.75
y14+
1611.76
1676.83
1751.76
m/z

# 20

- Experimental mass: 128.0945
- Unimod information: Lys
- Possible composition: H(12) C(6) N(2) O
- Sites in Unimod: n-term
- Sites observed: c-term

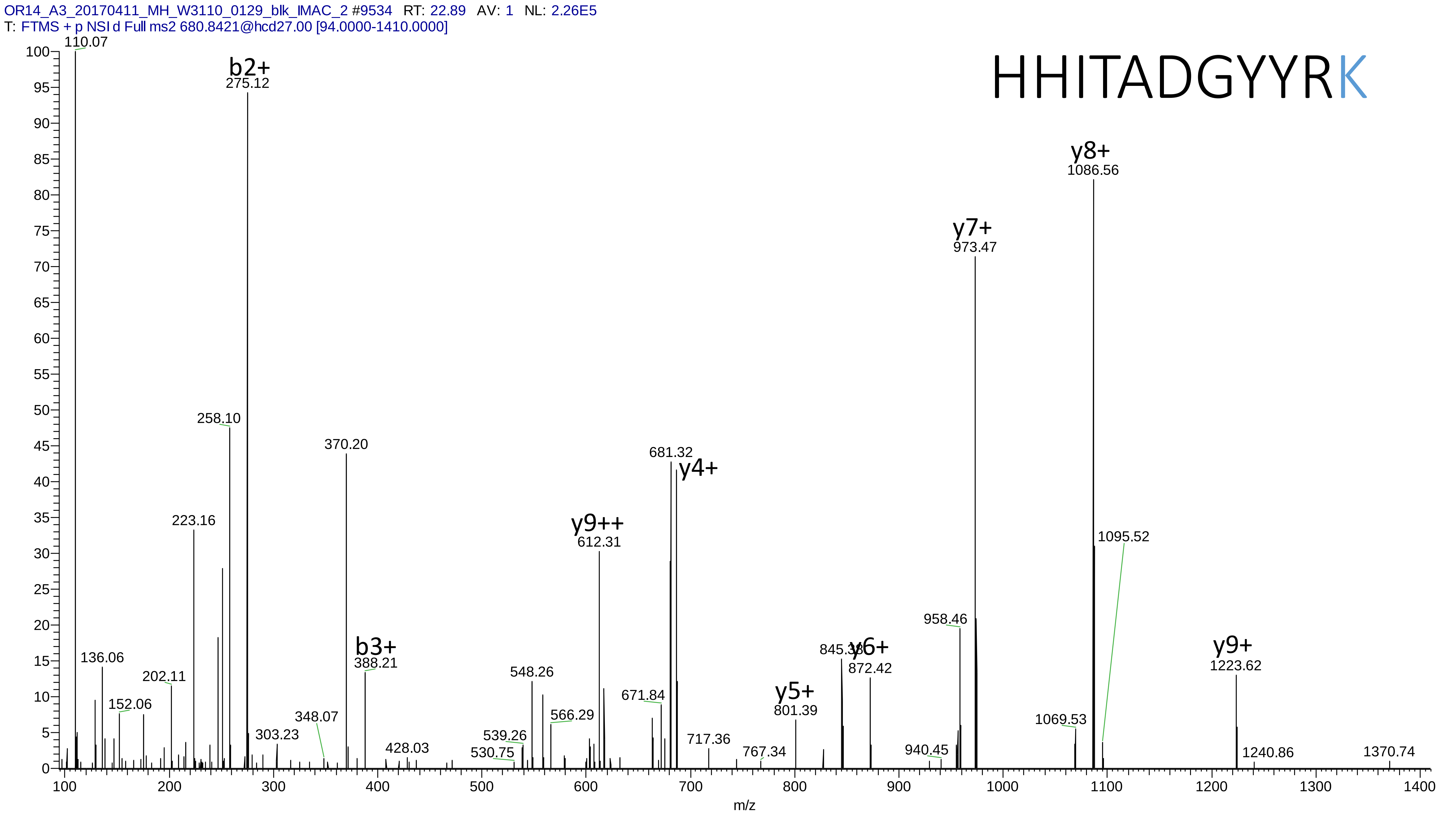
OR14_A3_20170411_MH_W3110_0129_blk_IMAC_2 #9534 RT: 22.89 AV: 1 NL: 2.26E5
T: FTMS + p NSI d Full ms2 680.8421@hcd27.00 [94.0000-1410.0000]
HHITADGYYRK
110.07
b2+
275.12
y8+
1086.56
y7+
973.47
258.10
370.20
681.32
y4+
223.16
y9++
612.31
1095.52
958.46
136.06
202.11
152.06
b3+
388.21
548.26
845.38
y6+
872.42
y9+
1223.62
671.84
y5+
801.39
566.29
348.07
303.23
539.26
428.03
530.75
717.36
767.34
940.45
1069.53
1240.86
1370.74
m/z

# 21

- Experimental mass: 283.028
- Unimod information: s-GlcNAc
- Possible composition: H(13)C(8)N(1)O(8)S(1)
- Sites in Unimod: T;S
- Sites observed: peptide n-term
- Features: only y ions are observed

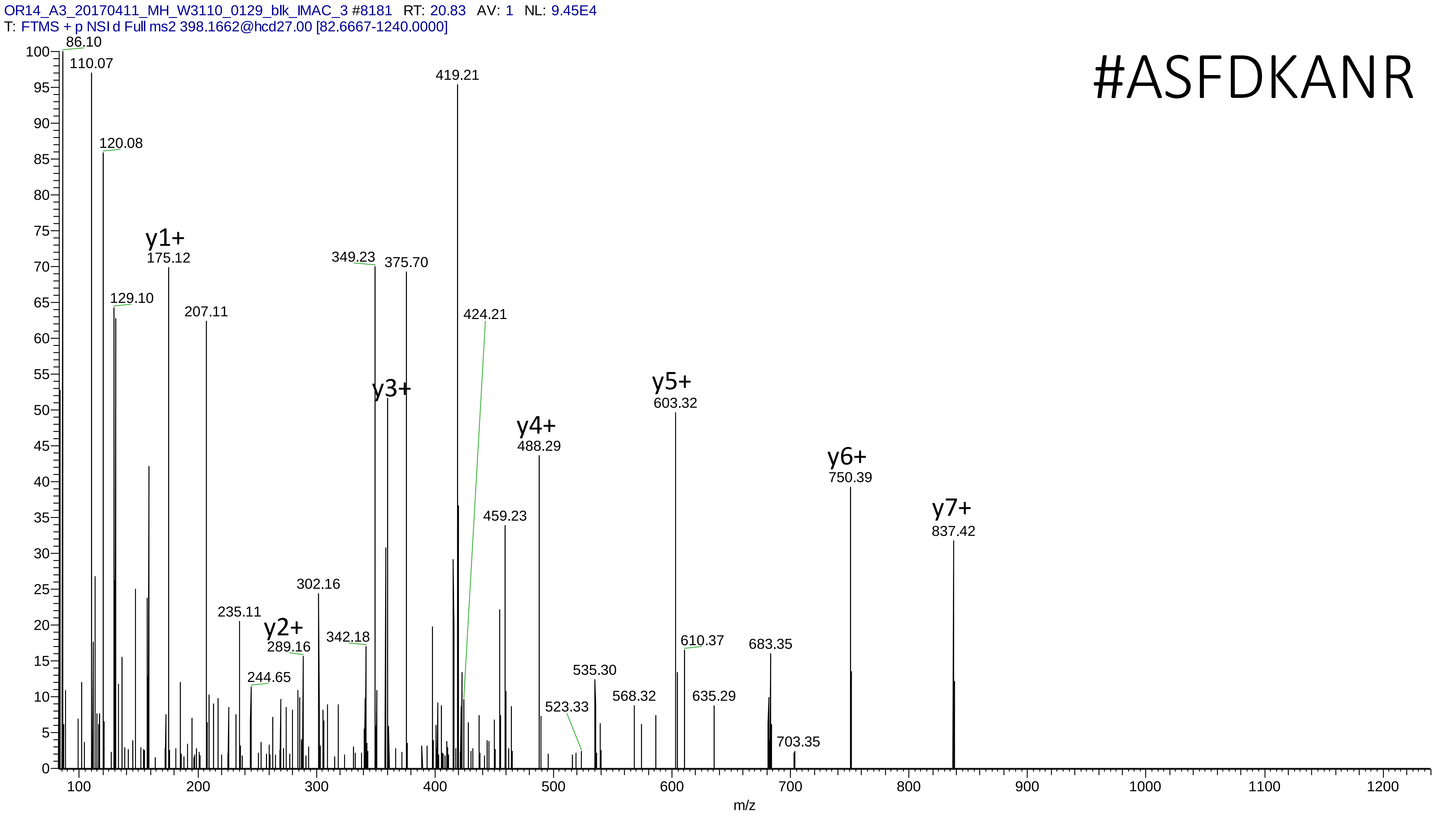
OR14_A3_20170411_MH_W3110_0129_blk_IMAC_3 #8181 RT: 20.83 AV: 1 NL: 9.45E4
T: FTMS + p NSI d Full ms2 398.1662@hcd27.00 [82.6667-1240.0000]
#ASFDKANR
86.10
110.07
120.08
129.10
y1+
175.12
207.11
235.11
244.65
y2+
289.16
302.16
342.18
349.23
y3+
375.70
419.21
424.21
459.23
y4+
488.29
523.33
535.30
568.32
y5+
603.32
610.37
635.29
683.35
703.35
y6+
750.39
y7+
837.42
m/z

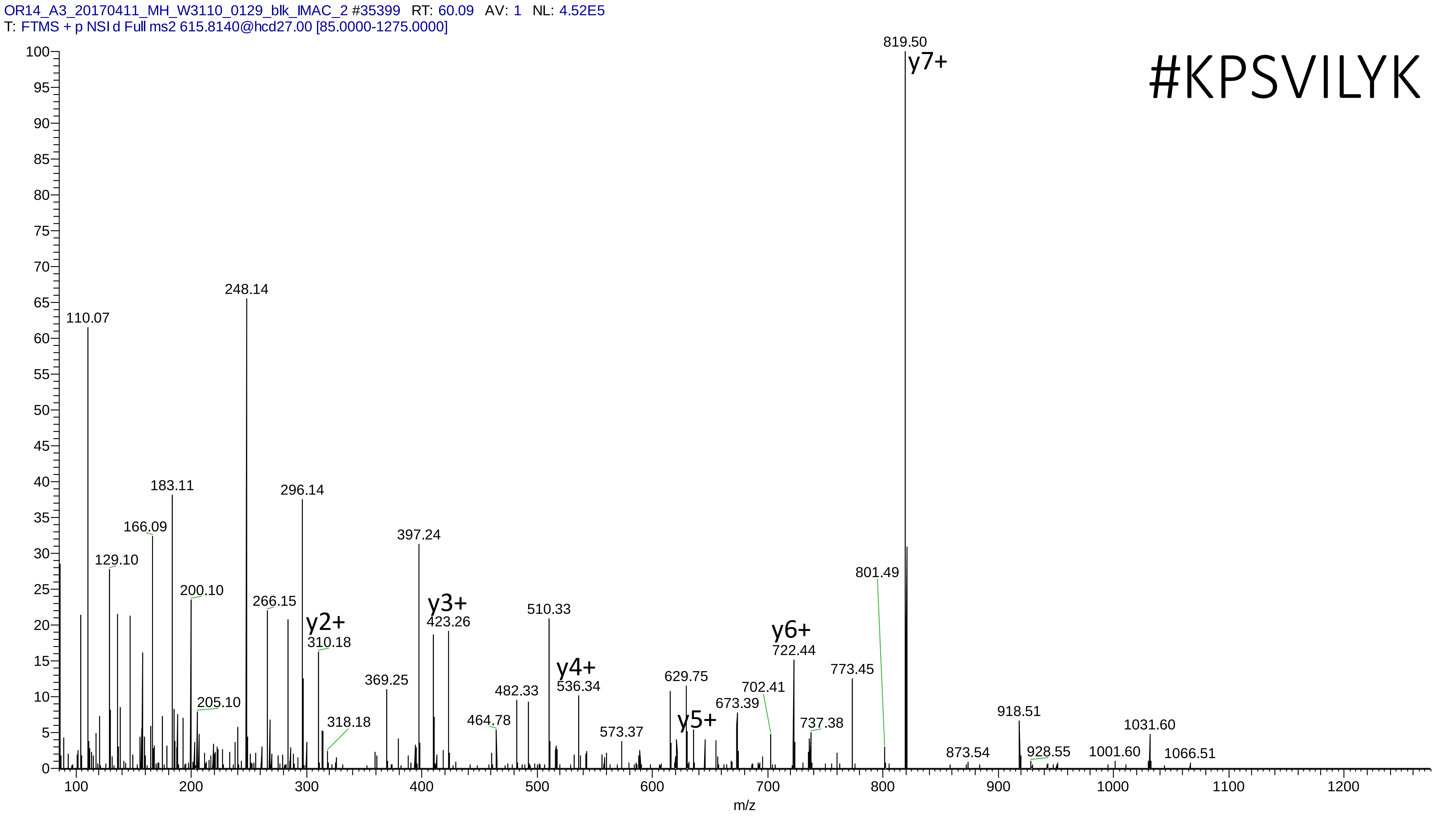
OR14_A3_20170411_MH_W3110_0129_blk_IMAC_2 #35399 RT: 60.09 AV: 1 NL: 4.52E5
T: FTMS + p NSI d Full ms2 615.8140@hcd27.00 [85.0000-1275.0000]
#KPSVILYK
819.50
y7+
248.14
110.07
183.11
296.14
166.09
129.10
397.24
801.49
200.10
266.15
y3+
423.26
510.33
y2+
310.18
y6+
722.44
773.45
y4+
536.34
629.75
369.25
482.33
702.41
205.10
673.39
918.51
1031.60
464.78
318.18
y5+
737.38
573.37
873.54
928.55
1001.60
1066.51
m/z

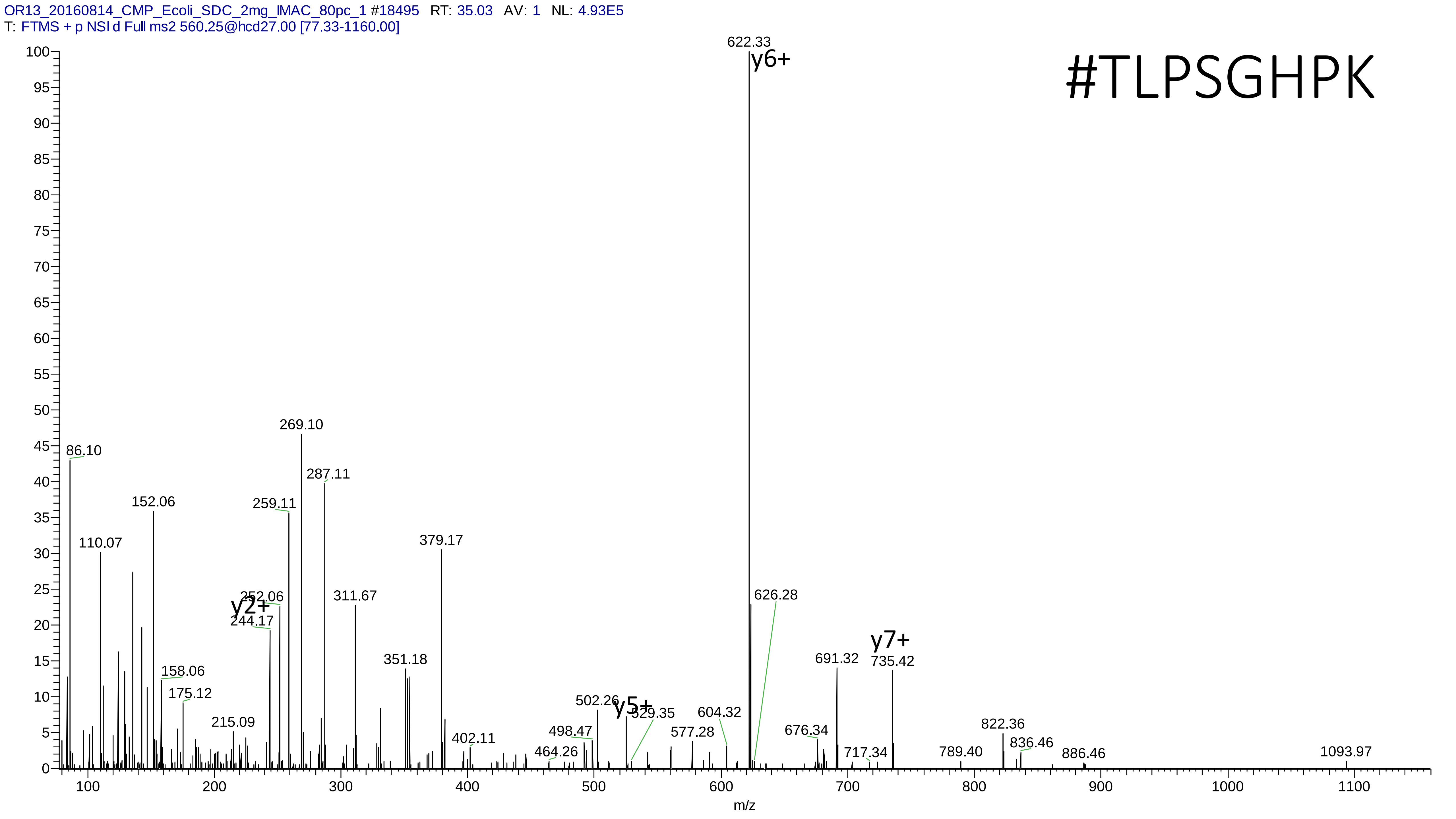
OR13_20160814_CMP_Ecoli_SDC_2mg_IMAC_80pc_1 #18495 RT: 35.03 AV: 1 NL: 4.93E5
T: FTMS + p NSI d Full ms2 560.25@hcd27.00 [77.33-1160.00]
#TLPSGHPK
622.33
y6+
269.10
86.10
287.11
152.06
259.11
110.07
379.17
252.06
y2+
244.17
311.67
626.28
y7+
691.32
735.42
351.18
158.06
175.12
215.09
502.26
y5+
529.35
604.32
577.28
498.47
402.11
464.26
676.34
717.34
789.40
822.36
836.46
886.46
1093.97
m/z

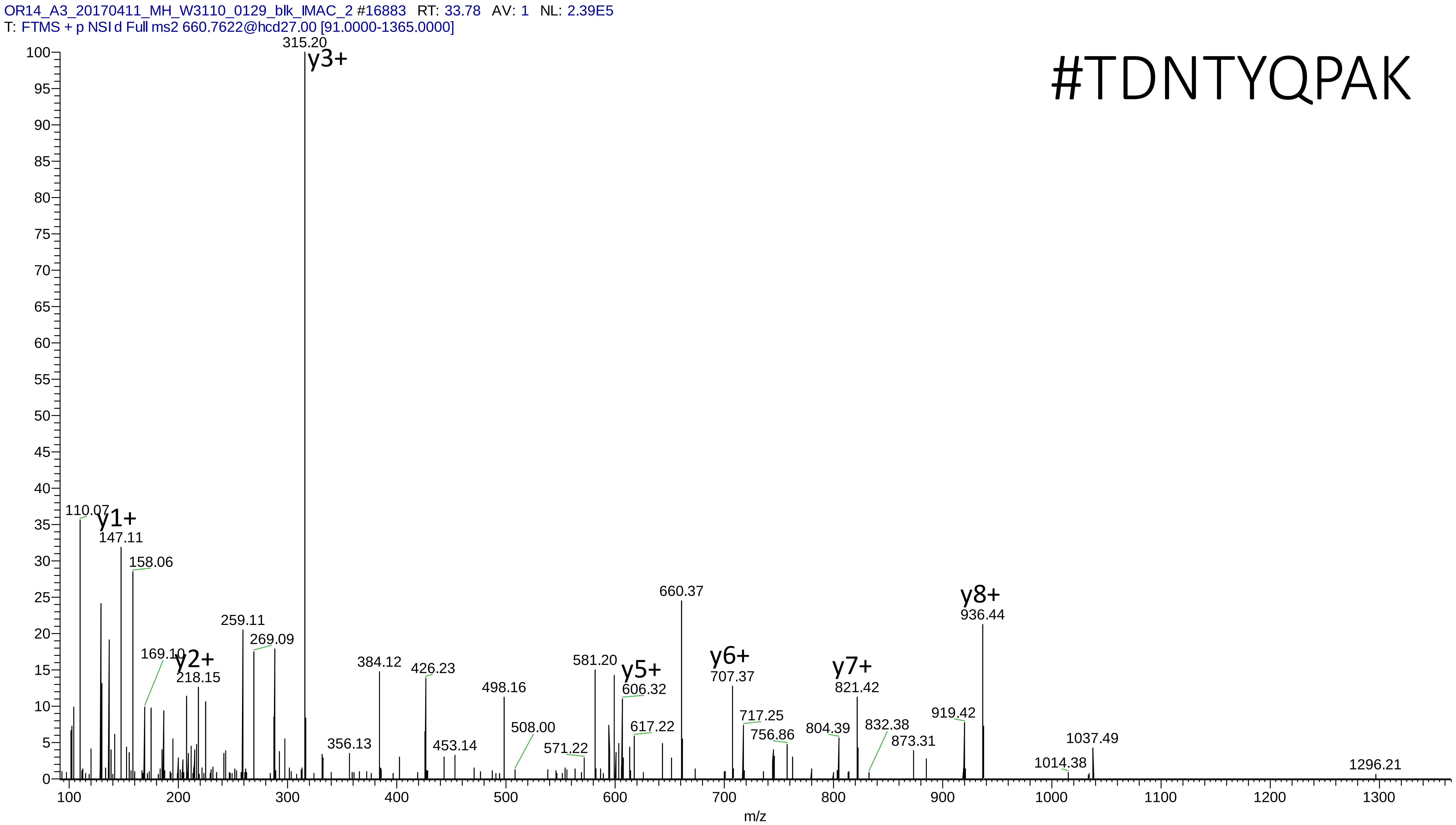
OR14_A3_20170411_MH_W3110_0129_blk_IMAC_2 #16883 RT: 33.78 AV: 1 NL: 2.39E5
T: FTMS + p NSI d Full ms2 660.7622@hcd27.00 [91.0000-1365.0000]
#TDNTYQPAK
315.20
y3+
110.07
y1+
147.11
158.06
169.10
y2+
218.15
259.11
269.09
356.13
384.12
426.23
453.14
498.16
508.00
571.22
581.20
y5+
606.32
617.22
660.37
y6+
707.37
717.25
756.86
804.39
y7+
821.42
832.38
873.31
919.42
y8+
936.44
1014.38
1037.49
1296.21
m/z

# 22

- Experimental mass: -128.0953
- Unimod information: Lys-loss
- Possible composition: H(-12) C(-6) N(-2) O(-1)
- Sites in Unimod: K
- Sites observed: loss of K at n-term
- Features: 94.5% (86/91) peptides contain K at the peptide n-term

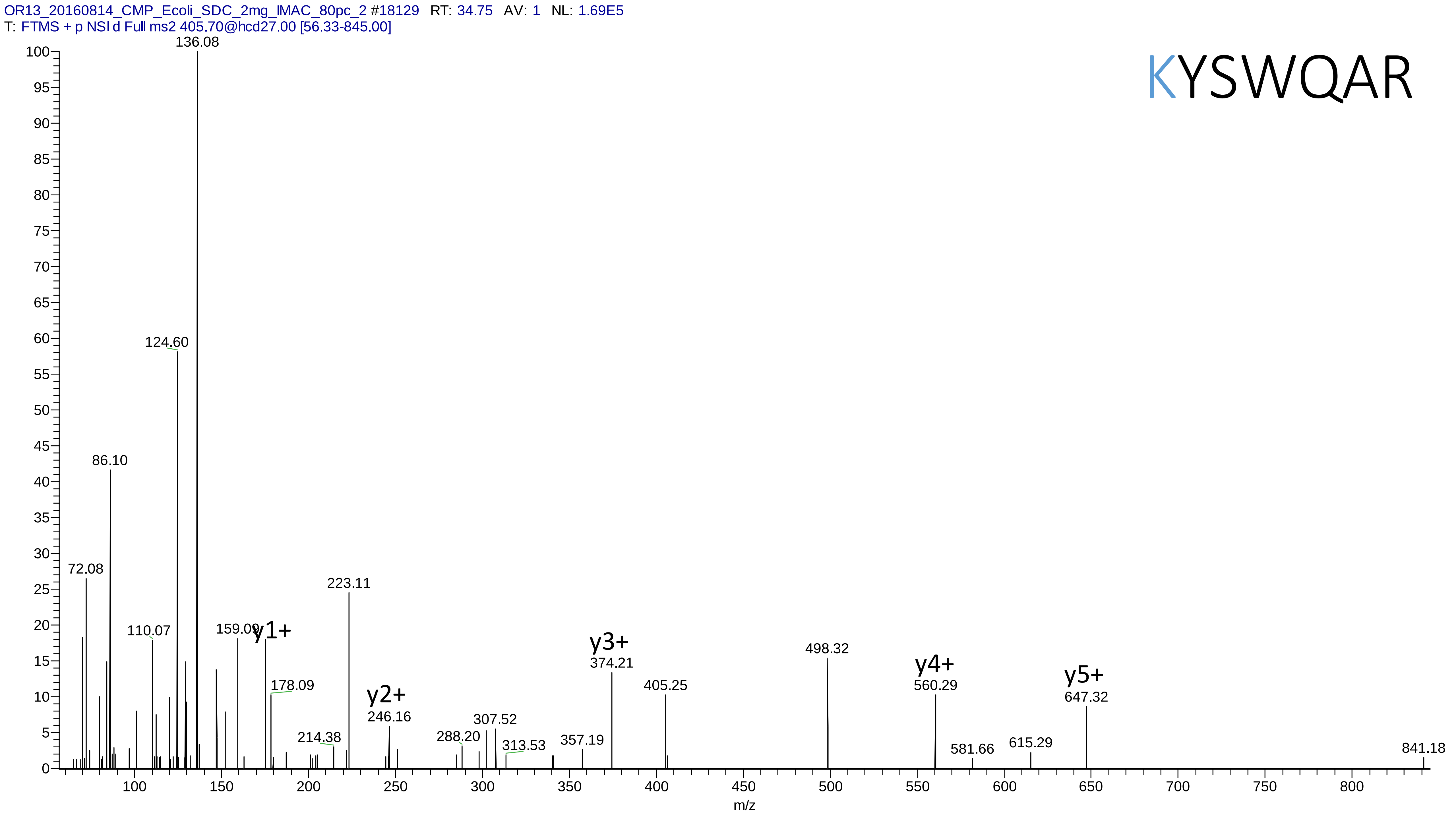

OR13_20160814_CMP_Ecoli_SDC_2mg_IMAC_80pc_2 #18129 RT: 34.75 AV: 1 NL: 1.69E5
T: FTMS + p NSI d Full ms2 405.70@hcd27.00 [56.33-845.00]
KYSWQAR
136.08
124.60
86.10
72.08
110.07
159.09
y1+
178.09
223.11
214.38
y2+
246.16
288.20
307.52
313.53
357.19
y3+
374.21
405.25
498.32
y4+
560.29
581.66
615.29
y5+
647.32
841.18
m/z

# 23

- Experimental mass: 22.945
- Unimod information: not found
- Possible composition: Phospho + loss of Carbamidomethyl, [H O(3) P + H(-3)C(-2)N(-1)O(-1)]
- Sites in Unimod: not found
- Sites observed: C
- Features: 100.0% (23/23) peptides contain C; neutral loss of $HPO_3$ and $H_3PO_4$(or $HPO_3$+$H_2O$) are observed in both precursor and fragment ions

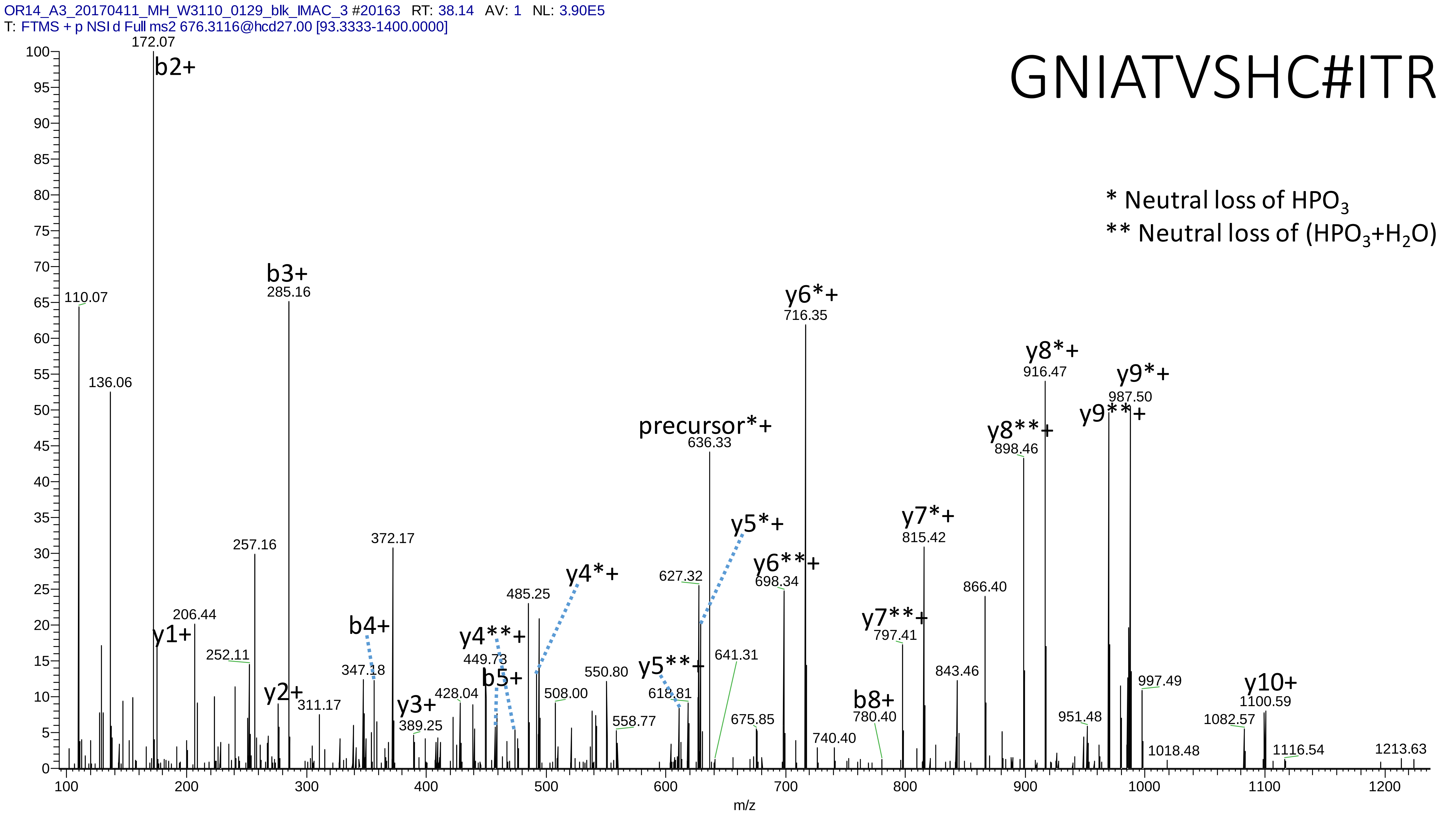
OR14_A3_20170411_MH_W3110_0129_blk_IMAC_3 #20163 RT: 38.14 AV: 1 NL: 3.90E5
T: FTMS + p NSI d Full ms2 676.3116@hcd27.00 [93.3333-1400.0000]
GNIATVSHC#ITR
* Neutral loss of HPO3
** Neutral loss of (HPO3+H2O)
172.07
b2+
110.07
136.06
b3+
285.16
257.16
206.44
y1+
252.11
y2+
311.17
b4+
347.18
372.17
y3+
389.25
428.04
y4**+
449.73
b5+
485.25
y4*+
508.00
550.80
558.77
precursor*+
636.33
627.32
y5*+
y5**+
618.81
641.31
675.85
y6*+
716.35
y6**+
698.34
740.40
b8+
780.40
y7*+
815.42
y7**+
797.41
843.46
866.40
y8*+
916.47
y8**+
898.46
y9*+
987.50
y9**+
951.48
997.49
1018.48
1082.57
y10+
1100.59
1116.54
1213.63
m/z

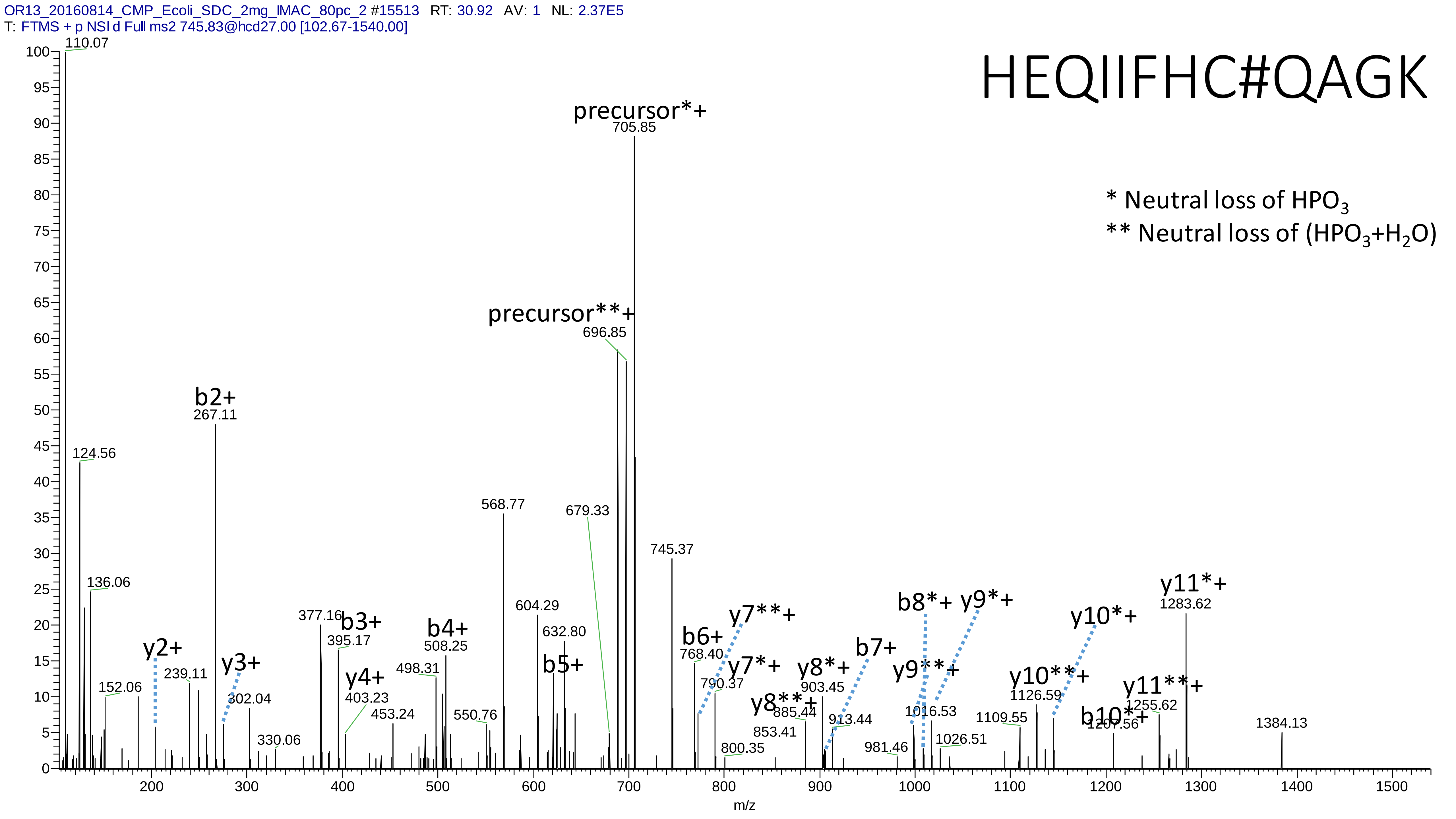
OR13_20160814_CMP_Ecoli_SDC_2mg_IMAC_80pc_2 #15513 RT: 30.92 AV: 1 NL: 2.37E5
T: FTMS + p NSI d Full ms2 745.83@hcd27.00 [102.67-1540.00]
HEQIIFHC#QAGK
* Neutral loss of HPO3
** Neutral loss of (HPO3+H2O)
precursor*+
705.85
precursor**+
696.85
b2+
267.11
110.07
124.56
136.06
152.06
y2+
239.11
y3+
302.04
330.06
377.16
b3+
395.17
y4+
403.23
453.24
b4+
508.25
498.31
550.76
568.77
604.29
632.80
b5+
679.33
745.37
b6+
768.40
y7**+
y7*+
790.37
800.35
y8**+
853.41
885.44
y8*+
903.45
913.44
b7+
b8*+
y9*+
y9**+
981.46
1016.53
1026.51
1109.55
y10**+
1126.59
y10*+
b10*+
1207.56
y11**+
1255.62
y11*+
1283.62
1384.13
m/z

# 24

- Experimental mass: 159.9336
- Unimod information: pyrophospho
- Possible composition: H(2) O(6) P(2)
- Sites in Unimod: S;T
- Sites observed: S;T
- Features: Neutral loss of $H_3PO_4$ are observed in both precursor and fragment ions

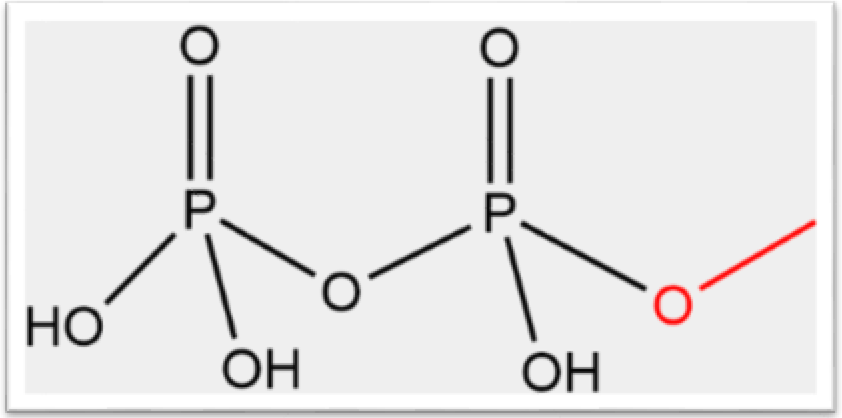

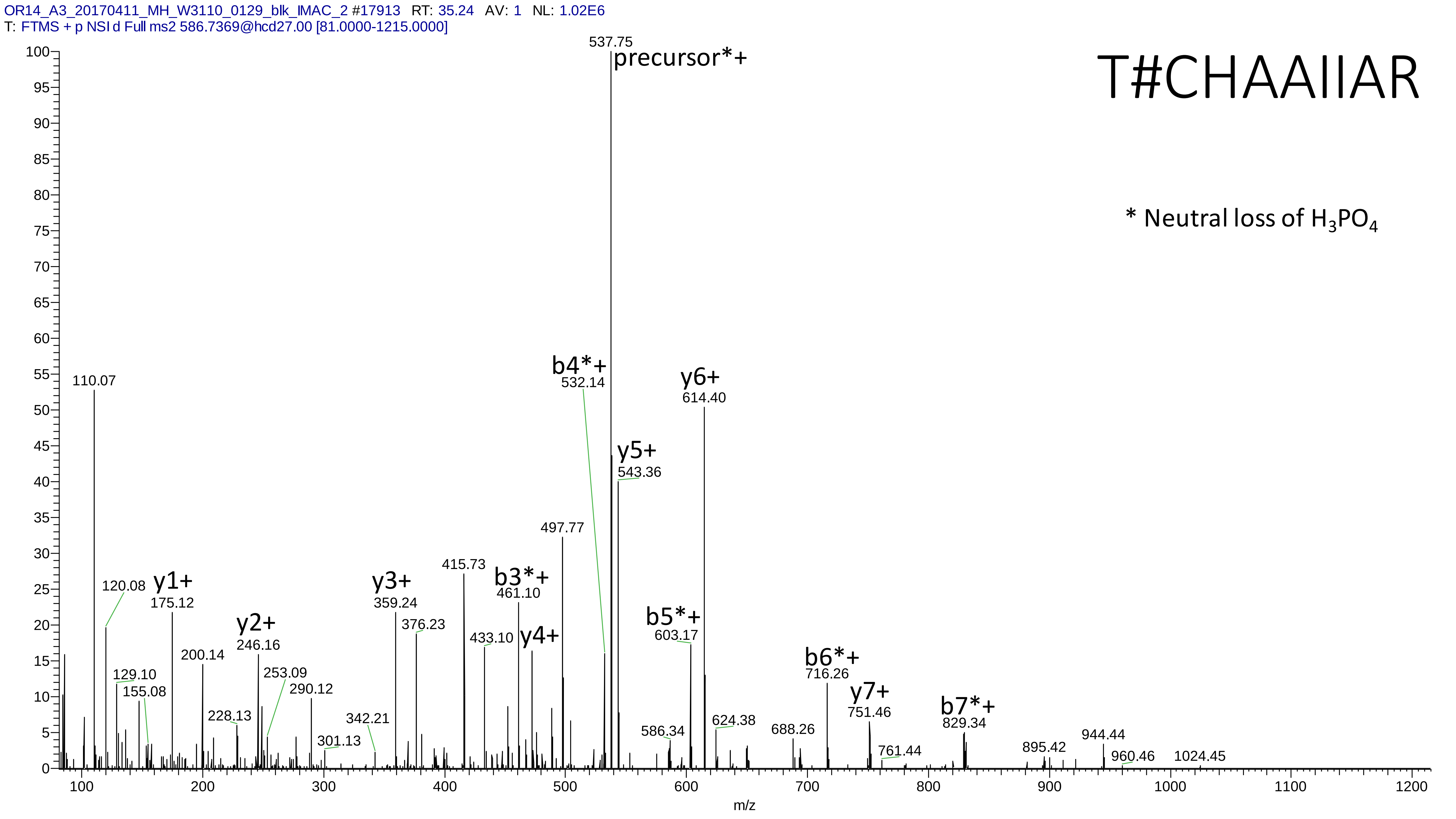

OR14_A3_20170411_MH_W3110_0129_blk_IMAC_2 #17913 RT: 35.24 AV: 1 NL: 1.02E6
T: FTMS + p NSI d Full ms2 586.7369@hcd27.00 [81.0000-1215.0000]
T#CHAAIIAR
* Neutral loss of H3PO4
precursor*+
537.75
110.07
120.08
y1+
175.12
129.10
155.08
200.14
228.13
y2+
246.16
253.09
290.12
301.13
342.21
y3+
359.24
376.23
415.73
433.10
b3*+
461.10
y4+
497.77
b4*+
532.14
y5+
543.36
586.34
b5*+
603.17
y6+
614.40
624.38
688.26
b6*+
716.26
y7+
751.46
761.44
b7*+
829.34
895.42
944.44
960.46
1024.45
m/z

# 25

- Experimental mass: -229.1431
- Unimod information: not found
- Possible composition: loss of amino acid T and K
- Sites in Unimod: not found
- Sites observed: “TK” at peptide n-term
- Features: 99.5% (213/214) peptide spectra matches contain “TK” at peptide n-term or c-term

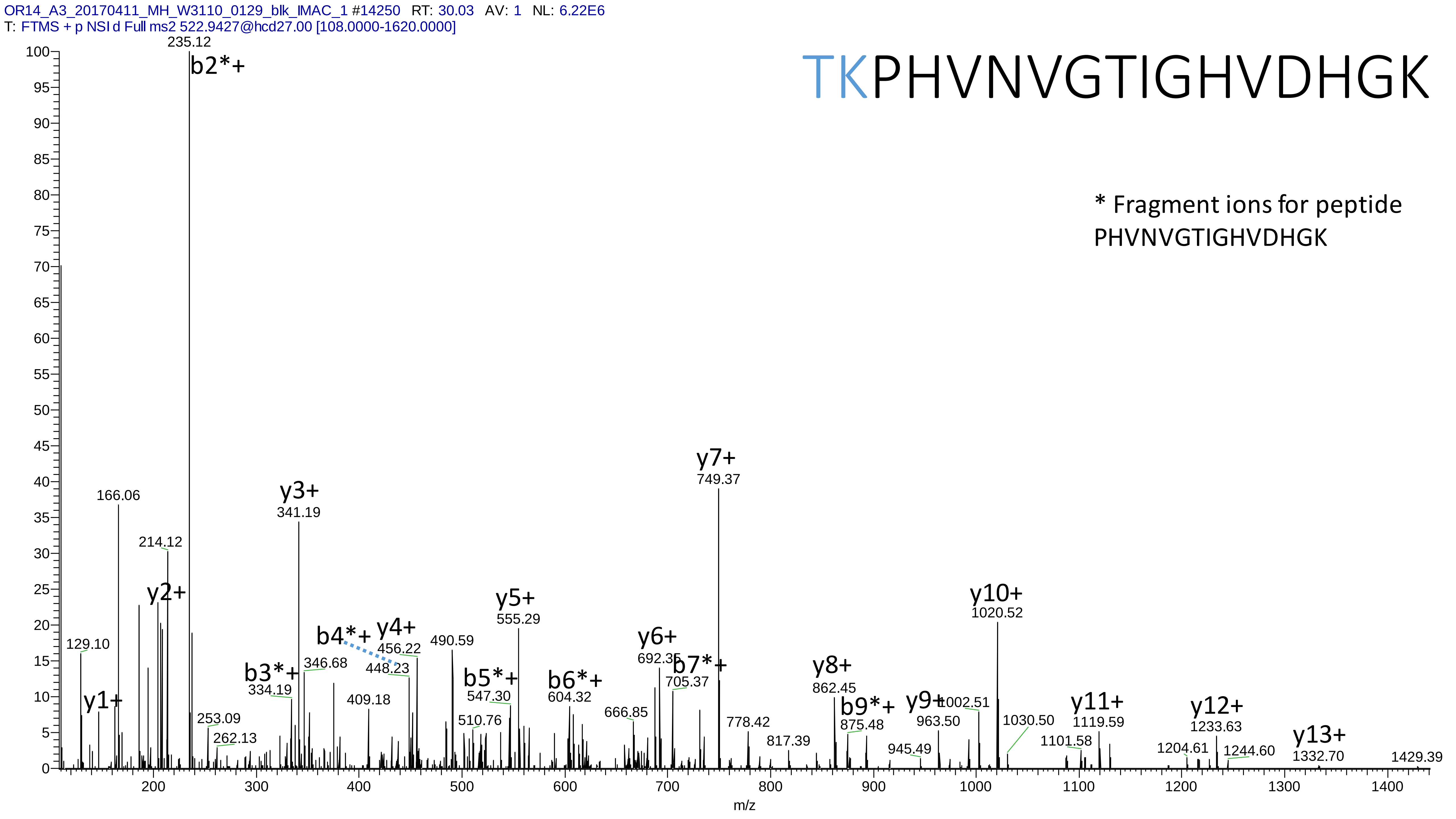

OR14_A3_20170411_MH_W3110_0129_blk_IMAC_1 #14250 RT: 30.03 AV: 1 NL: 6.22E6
T: FTMS + p NSI d Full ms2 522.9427@hcd27.00 [108.0000-1620.0000]
TKPHVNVGTIGHVDHGK
* Fragment ions for peptide PHVNVGTIGHVDHGK
235.12
b2*+
166.06
214.12
y2+
y1+
129.10
y3+
341.19
b3*+
334.19
346.68
b4*+
y4+
456.22
448.23
409.18
490.59
y5+
555.29
b5*+
547.30
510.76
b6*+
604.32
666.85
253.09
262.13
y6+
692.35
b7*+
705.37
y7+
749.37
778.42
817.39
y8+
862.45
b9*+
875.48
y9+
963.50
945.49
1002.51
y10+
1020.52
1030.50
1101.58
y11+
1119.59
1204.61
y12+
1233.63
1244.60
y13+
1332.70
1429.39
m/z

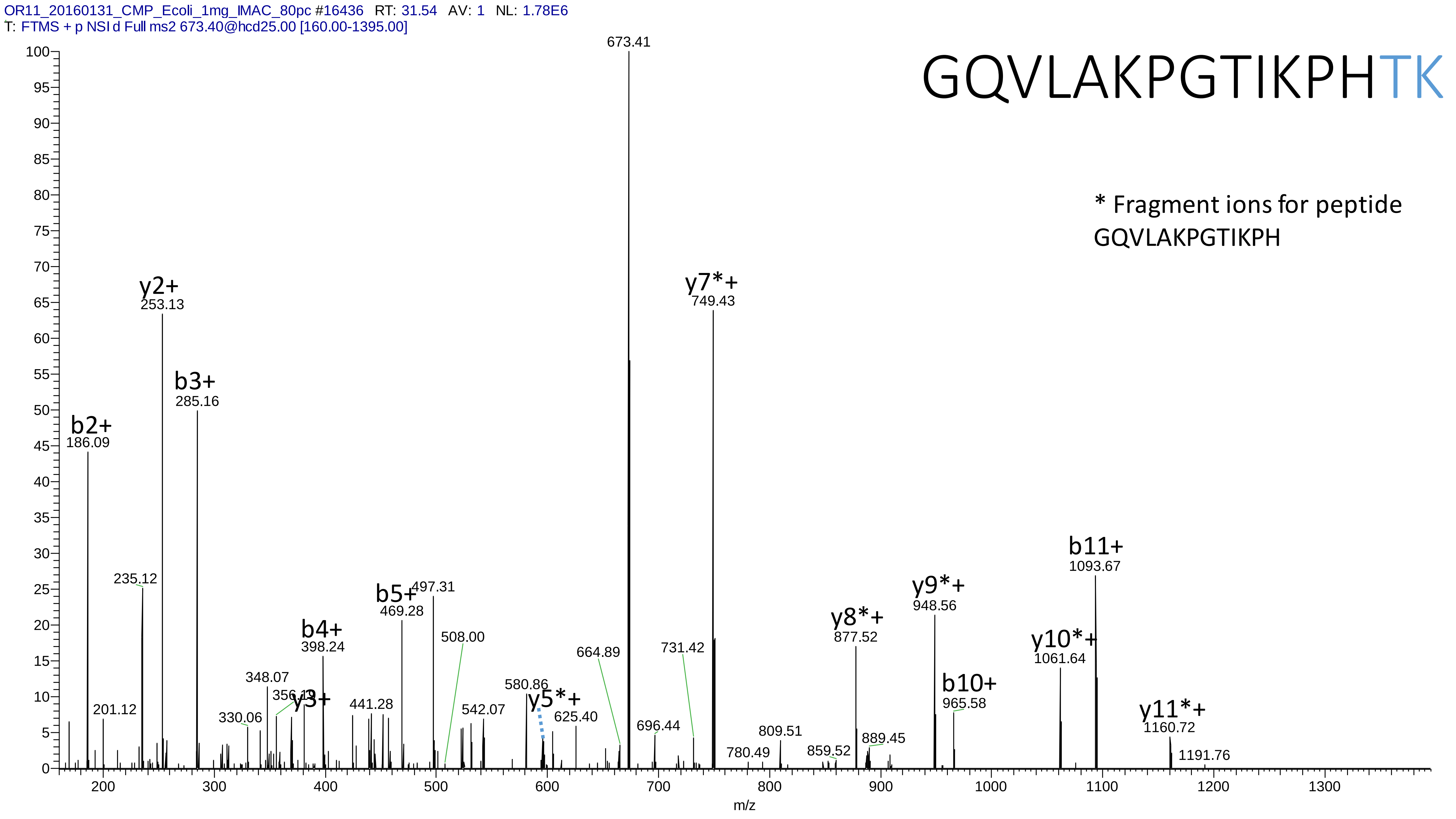
OR11_20160131_CMP_Ecoli_1mg_IMAC_80pc #16436 RT: 31.54 AV: 1 NL: 1.78E6
T: FTMS + p NSI d Full ms2 673.40@hcd25.00 [160.00-1395.00]
GQVLAKPGTIKPHTK
* Fragment ions for peptide GQVLAKPGTIKPH
673.41
y2+ 253.13
b3+ 285.16
b2+ 186.09
y7*+ 749.43
b11+ 1093.67
y9*+ 948.56
y8*+ 877.52
y10*+ 1061.64
b10+ 965.58
y11*+ 1160.72
b5+ 469.28
497.31
b4+ 398.24
y3+
y5*+
235.12
201.12
348.07
356.13
330.06
441.28
508.00
542.07
580.86
625.40
664.89
696.44
731.42
780.49
809.51
859.52
889.45
1191.76
m/z

# 26

- Experimental mass: -327.151
- Unimod information: not found
- Possible composition:
- Sites in Unimod: not found
- Sites observed:
- Features: 99.5% (212/213) PSMs are from the same peptide VPYGAVLAKGDGEQVAGGETVANWDPHTMPVITEVSGFVR, spectra are too complex to be interpreted

# 27

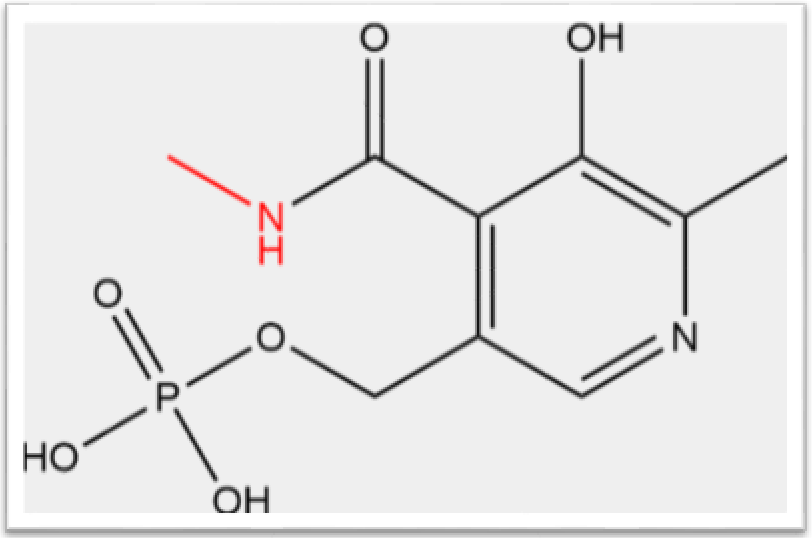


- Experimental mass: 245.0087
- Unimod information: not found
- Possible composition: H(8) C(8) N(1) O(6) P(1)
- Sites in Unimod: not found
- Sites observed: peptide N-term
- Features: neutral loss of $H_3PO_4$ is observed in both precursor and fragment ions

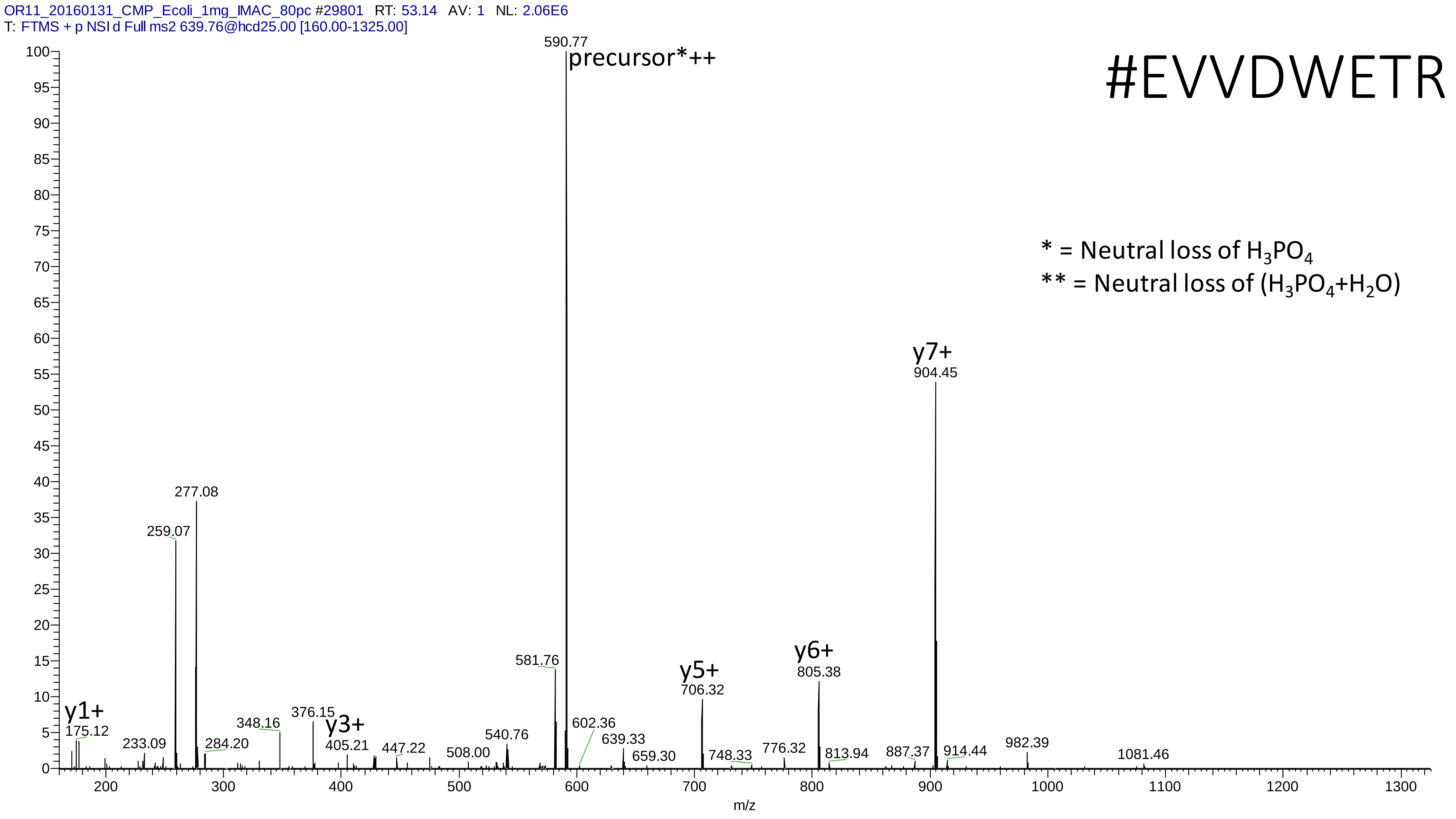

OR11_20160131_CMP_Ecoli_1mg_IMAC_80pc #29801 RT: 53.14 AV: 1 NL: 2.06E6
T: FTMS + p NSI d Full ms2 639.76@hcd25.00 [160.00-1325.00]
#EVVDWETR
* = Neutral loss of H3PO4
** = Neutral loss of (H3PO4+H2O)
precursor*++
590.77
y7+
904.45
y6+
805.38
y5+
706.32
y3+
405.21
y1+
175.12
277.08
259.07
581.76
376.15
348.16
284.20
233.09
447.22
508.00
540.76
602.36
639.33
659.30
748.33
776.32
813.94
887.37
914.44
982.39
1081.46
m/z

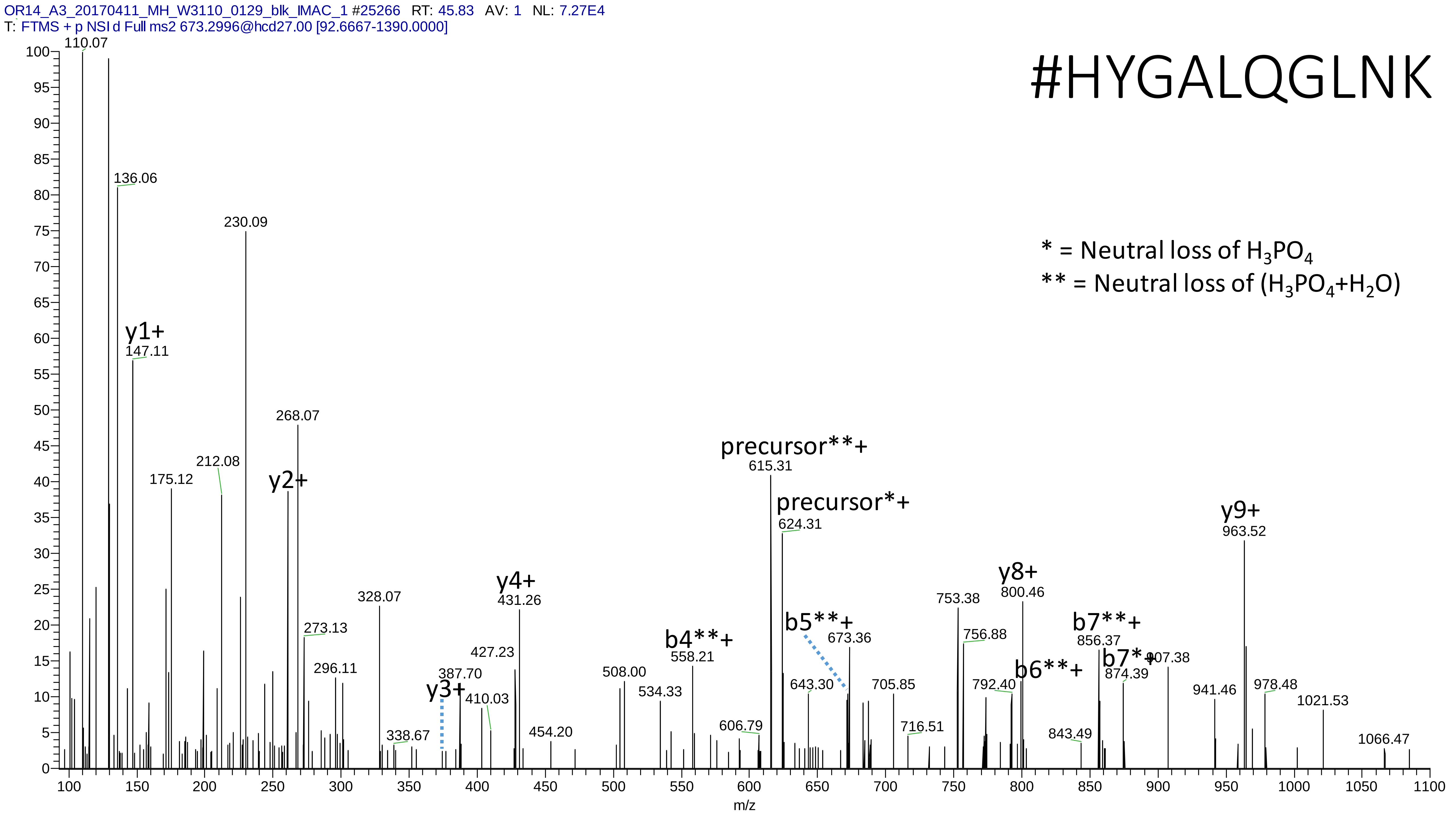

OR14_A3_20170411_MH_W3110_0129_blk_IMAC_1 #25266 RT: 45.83 AV: 1 NL: 7.27E4
T: FTMS + p NSI d Full ms2 673.2996@hcd27.00 [92.6667-1390.0000]
#HYGALQGLNK
* = Neutral loss of H3PO4
** = Neutral loss of (H3PO4+H2O)
y1+
y2+
y3+
y4+
b4**+
b5**+
precursor**+
precursor*+
b6**+
b7**+
b7*+
y8+
y9+
110.07
136.06
147.11
175.12
212.08
230.09
268.07
273.13
296.11
328.07
338.67
387.70
410.03
427.23
431.26
454.20
508.00
534.33
558.21
606.79
615.31
624.31
643.30
673.36
705.85
716.51
753.38
756.88
792.40
800.46
843.49
856.37
874.39
907.38
941.46
963.52
978.48
1021.53
1066.47
m/z

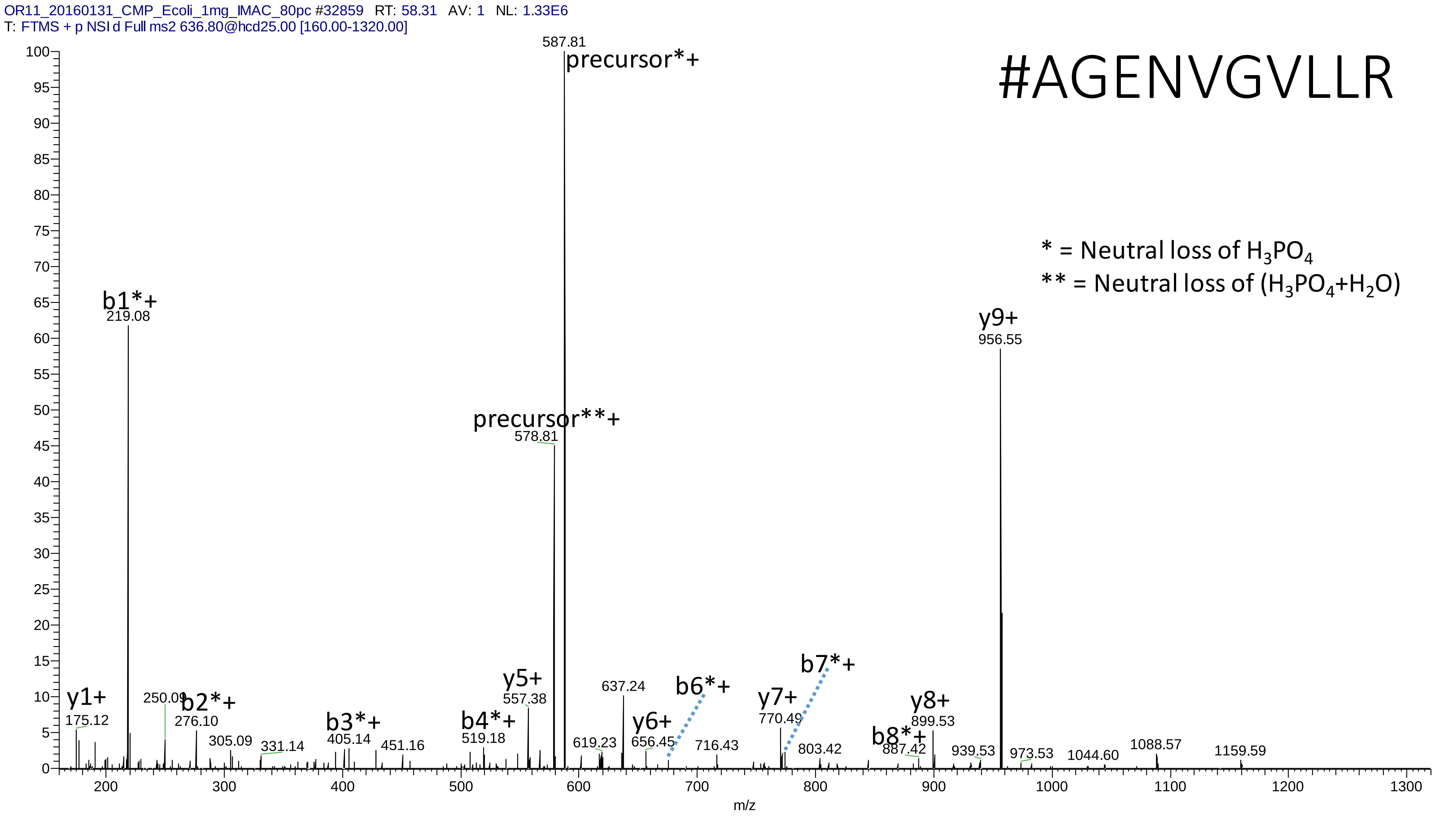
OR11_20160131_CMP_Ecoli_1mg_IMAC_80pc #32859 RT: 58.31 AV: 1 NL: 1.33E6
T: FTMS + p NSI d Full ms2 636.80@hcd25.00 [160.00-1320.00]
#AGENVGVLLR
* = Neutral loss of H3PO4
** = Neutral loss of (H3PO4+H2O)
587.81
precursor*+
precursor**+
578.81
b1*+
219.08
y9+
956.55
y1+
175.12
250.09
b2*+
276.10
305.09
331.14
b3*+
405.14
451.16
b4*+
519.18
y5+
557.38
619.23
637.24
y6+
656.45
b6*+
716.43
y7+
770.49
b7*+
803.42
b8*+
887.42
y8+
899.53
939.53
973.53
1044.60
1088.57
1159.59
m/z

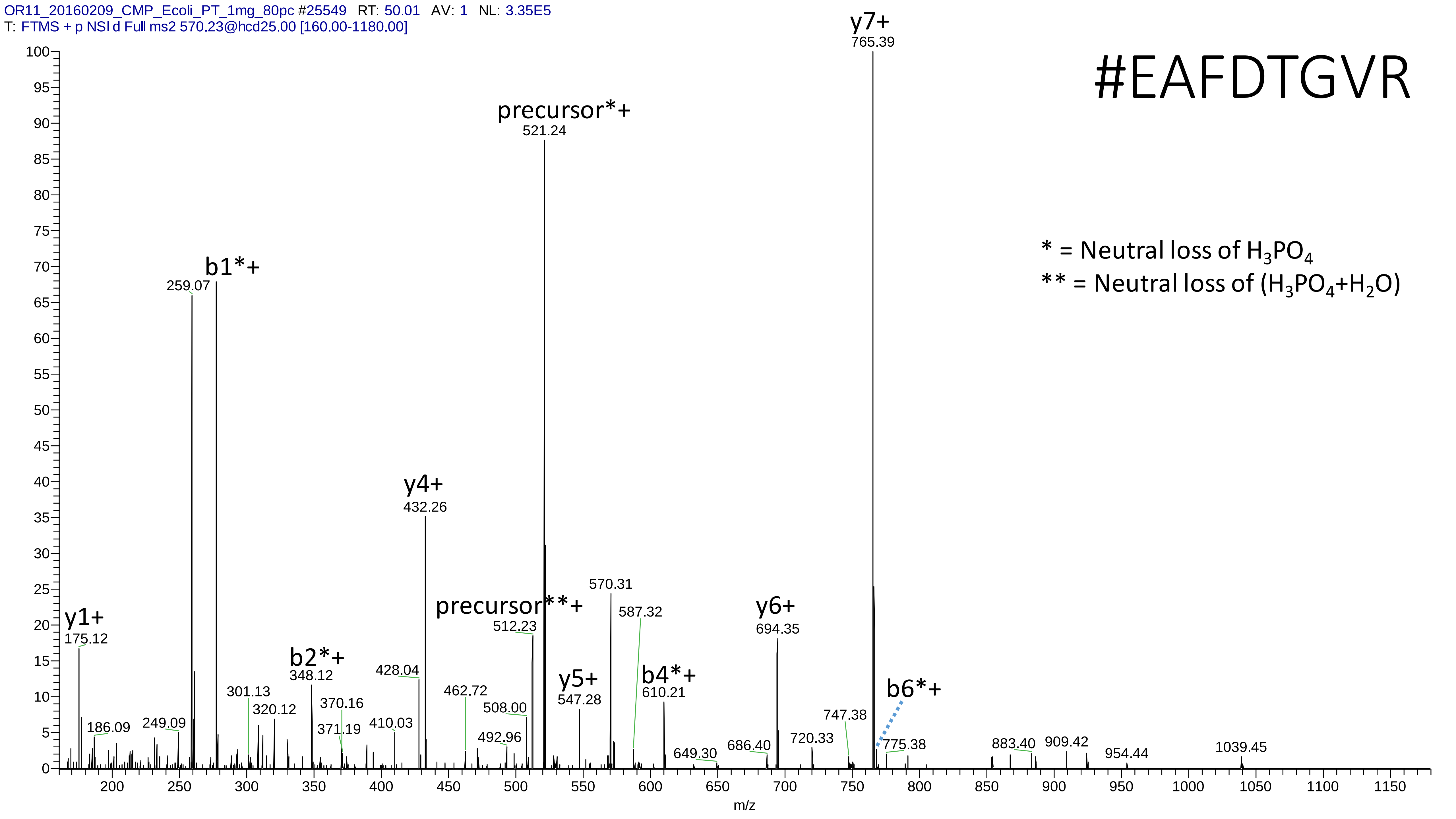

OR11_20160209_CMP_Ecoli_PT_1mg_80pc #25549 RT: 50.01 AV: 1 NL: 3.35E5
T: FTMS + p NSI d Full ms2 570.23@hcd25.00 [160.00-1180.00]
#EAFDTGVR
* = Neutral loss of $H_3PO_4$
** = Neutral loss of ($H_3PO_4$+$H_2O$)
y7+ 765.39
precursor*+ 521.24
b1*+ 259.07
y4+ 432.26
precursor**+ 512.23
570.31
587.32
y6+ 694.35
y1+ 175.12
b2*+ 348.12
428.04
y5+ 547.28
b4*+ 610.21
b6*+
301.13
320.12
370.16
371.19
410.03
462.72
508.00
492.96
186.09
249.09
649.30
686.40
720.33
747.38
775.38
883.40
909.42
954.44
1039.45
m/z

# 28

- Experimental mass: 14.0156
- Unimod information: methyl; Val->Xle; Asp->Glu; Ser->Thr; Asn->Gln; Gly->Ala
- Possible composition: H(2) C
- Sites in Unimod: C;H;K;N;Q;R;I;L;D;E;S;T;n-term;c-term
- Sites observed: I;K

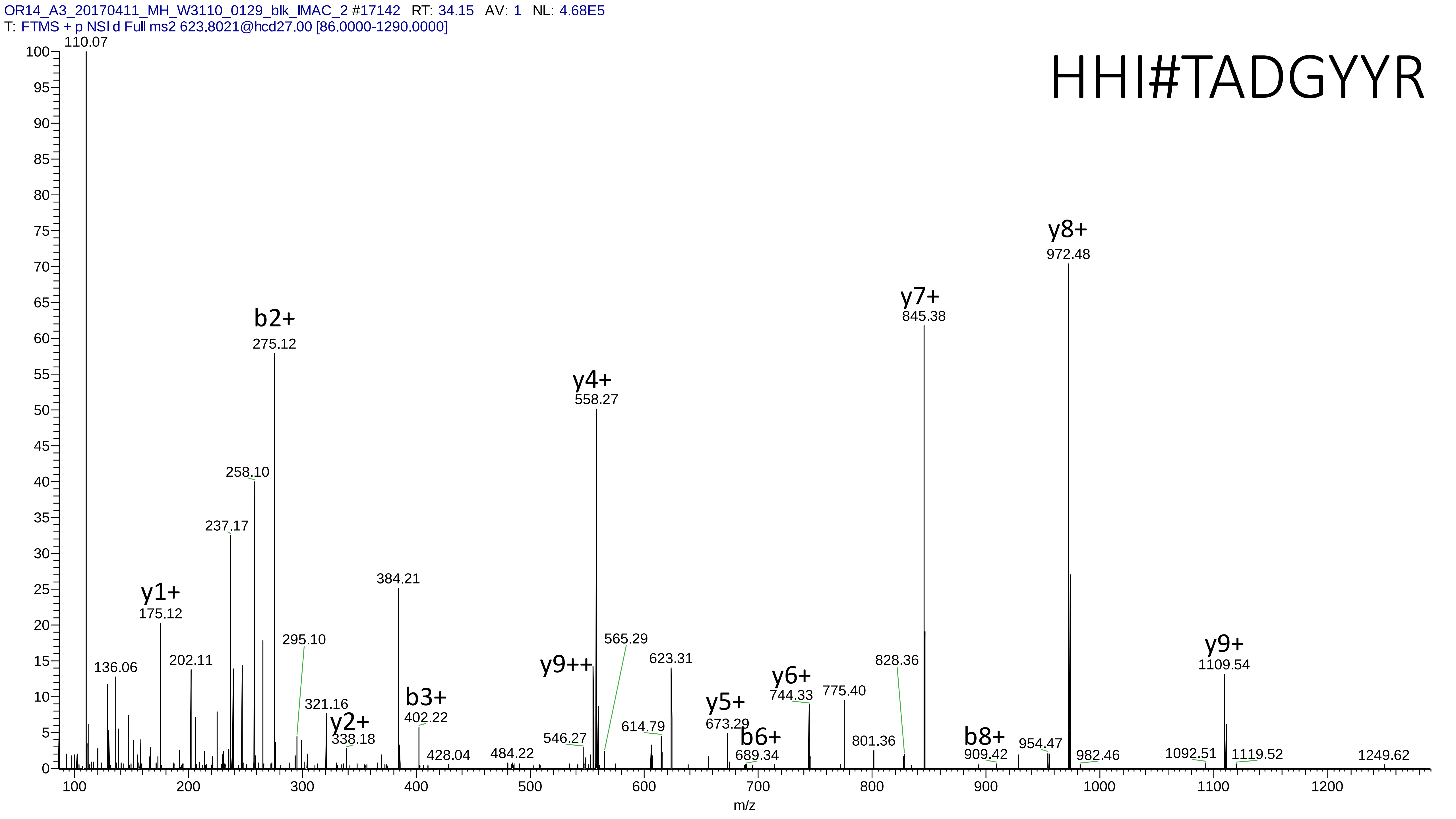
OR14_A3_20170411_MH_W3110_0129_blk_IMAC_2 #17142 RT: 34.15 AV: 1 NL: 4.68E5
T: FTMS + p NSI d Full ms2 623.8021@hcd27.00 [86.0000-1290.0000]
HHI#TADGYYR
110.07
y1+
175.12
136.06
202.11
237.17
258.10
b2+
275.12
295.10
321.16
y2+
338.18
384.21
b3+
402.22
428.04
484.22
546.27
y9++
y4+
558.27
565.29
614.79
623.31
y5+
673.29
b6+
689.34
y6+
744.33
775.40
801.36
828.36
y7+
845.38
b8+
909.42
954.47
y8+
972.48
982.46
1092.51
y9+
1109.54
1119.52
1249.62
m/z

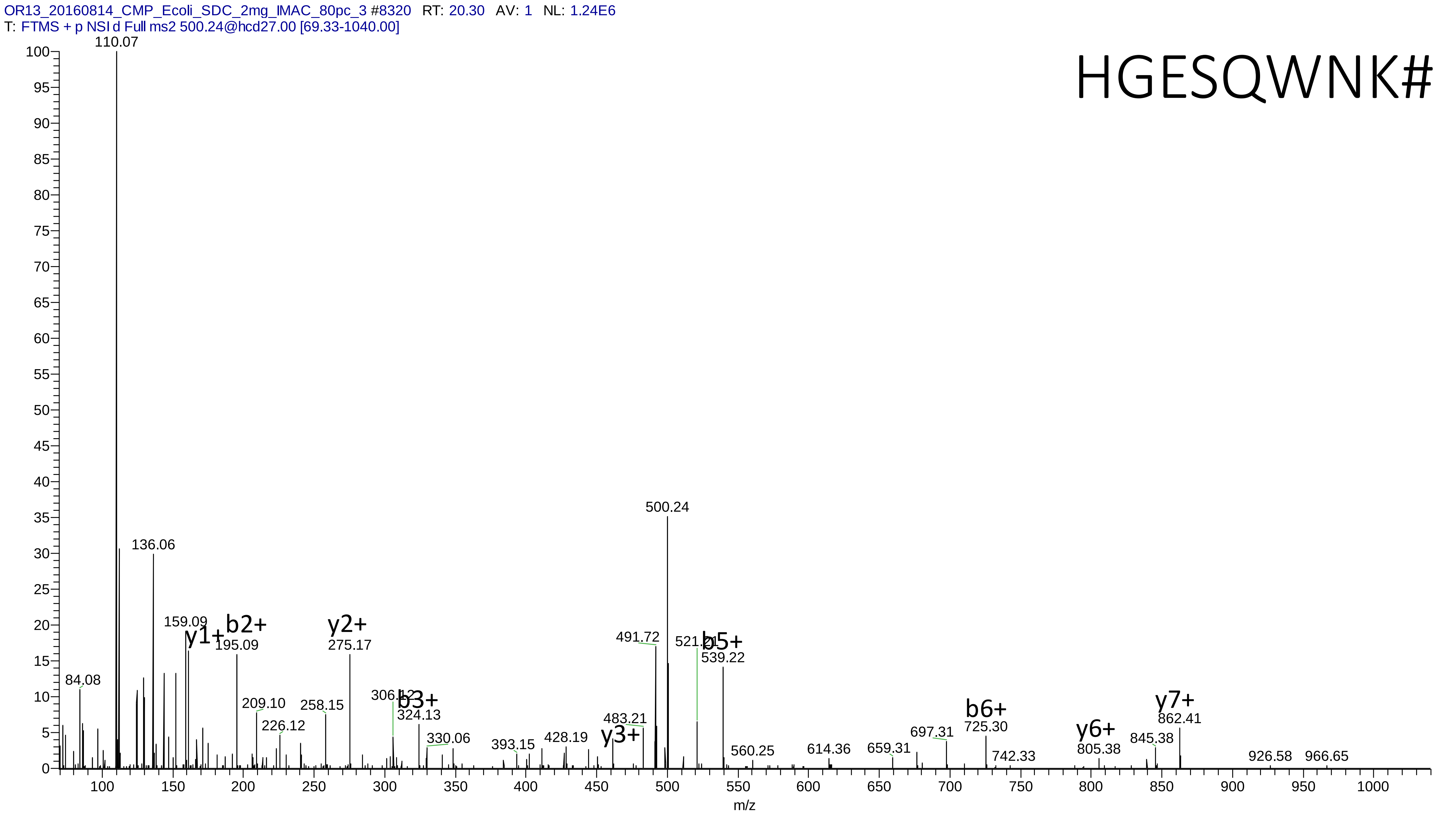
OR13_20160814_CMP_Ecoli_SDC_2mg_IMAC_80pc_3 #8320 RT: 20.30 AV: 1 NL: 1.24E6
T: FTMS + p NSI d Full ms2 500.24@hcd27.00 [69.33-1040.00]
HGESQWNK#
110.07
136.06
159.09
y1+
b2+
195.09
y2+
275.17
84.08
209.10
226.12
258.15
306.12
b3+
324.13
330.06
393.15
428.19
483.21
y3+
491.72
500.24
521.21
b5+
539.22
560.25
614.36
659.31
697.31
b6+
725.30
742.33
y6+
805.38
845.38
y7+
862.41
926.58
966.65
m/z

# 29

- Experimental mass: 17.0276
- Unimod information: Ammonium
- Possible composition: [H(3)N(1)]
- Sites in Unimod: E;D;c*
- Sites observed: D;E

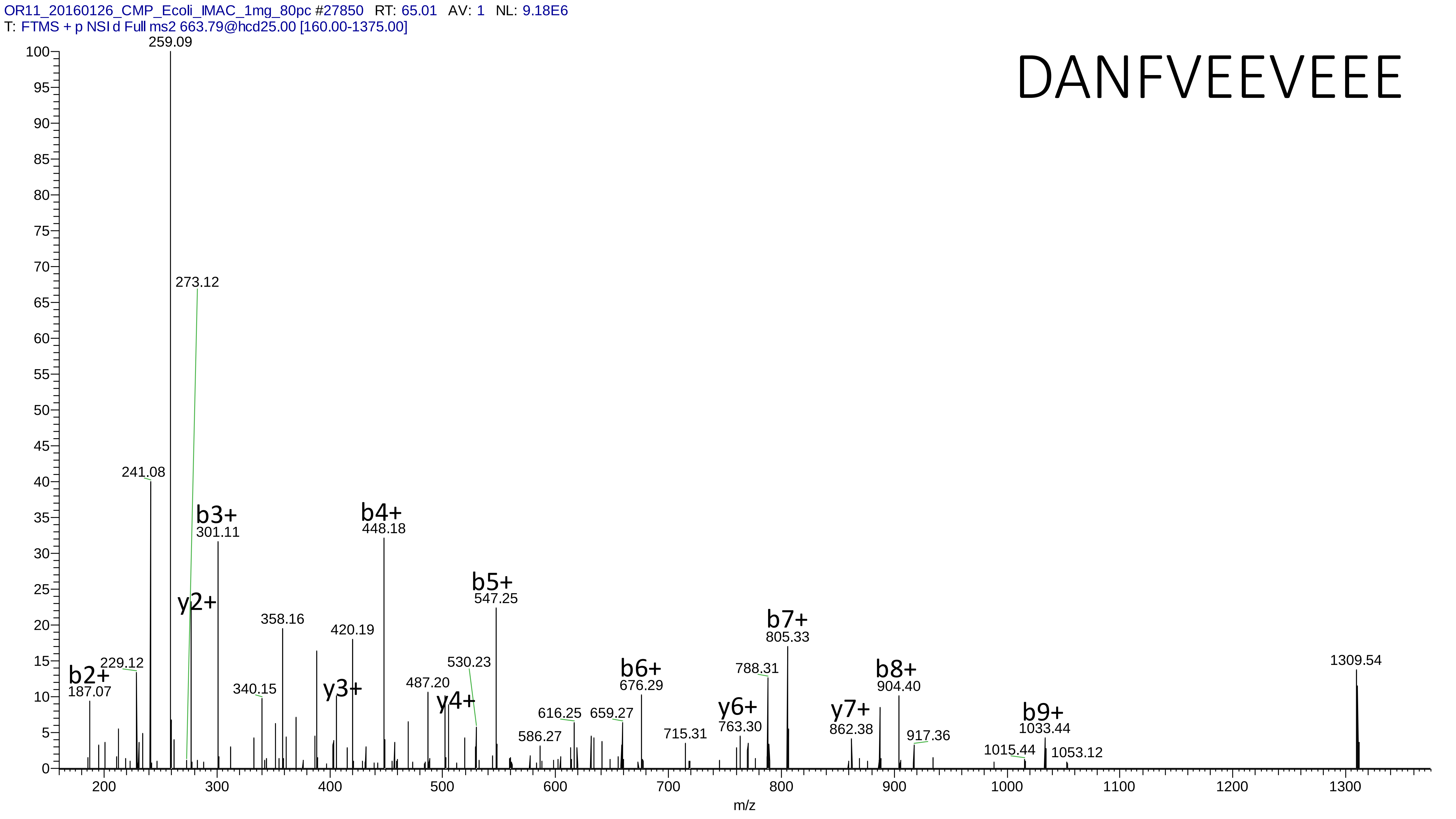
OR11_20160126_CMP_Ecoli_IMAC_1mg_80pc #27850 RT: 65.01 AV: 1 NL: 9.18E6
T: FTMS + p NSI d Full ms2 663.79@hcd25.00 [160.00-1375.00]
DANFVEEVEEE
259.09
273.12
241.08
b3+
301.11
b4+
448.18
y2+
b5+
547.25
358.16
420.19
b2+
187.07
229.12
340.15
y3+
487.20
y4+
530.23
b6+
676.29
616.25
659.27
586.27
715.31
y6+
763.30
788.31
b7+
805.33
y7+
862.38
b8+
904.40
917.36
b9+
1033.44
1015.44
1053.12
1309.54
m/z

# 30

- Experimental mass: 344.0406
- Unimod information: not found
- Possible composition:
- Sites in Unimod: not found
- Sites observed:
- Features: 49.2% (98/199) PSMs are from the peptide EDLLHGGAHKTNQVLGQALLAK and 43.7% (87/199) PSMs are from the peptide REDLLHGGAHKTNQVLGQALLAK, spectra are too complex to be interpreted, 99.0% (197/199) PSMs contain miss cleavage site K

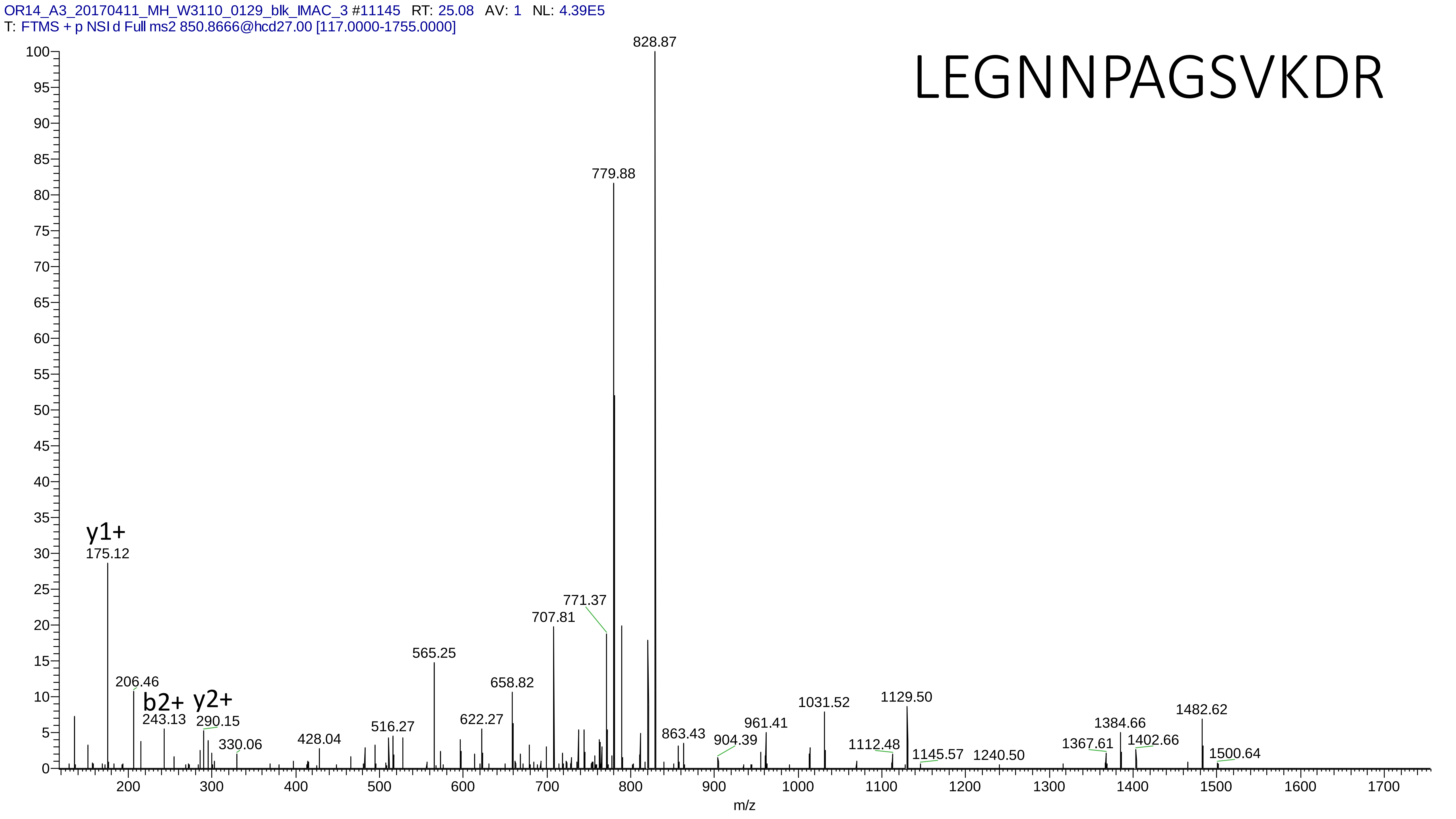
OR14_A3_20170411_MH_W3110_0129_blk_IMAC_3 #11145 RT: 25.08 AV: 1 NL: 4.39E5
T: FTMS + p NSI d Full ms2 850.8666@hcd27.00 [117.0000-1755.0000]
LEGNNPAGSVKDR
y1+
175.12
206.46
b2+
243.13
y2+
290.15
330.06
428.04
516.27
565.25
622.27
658.82
707.81
771.37
779.88
828.87
863.43
904.39
961.41
1031.52
1112.48
1129.50
1145.57
1240.50
1367.61
1384.66
1402.66
1482.62
1500.64
m/z

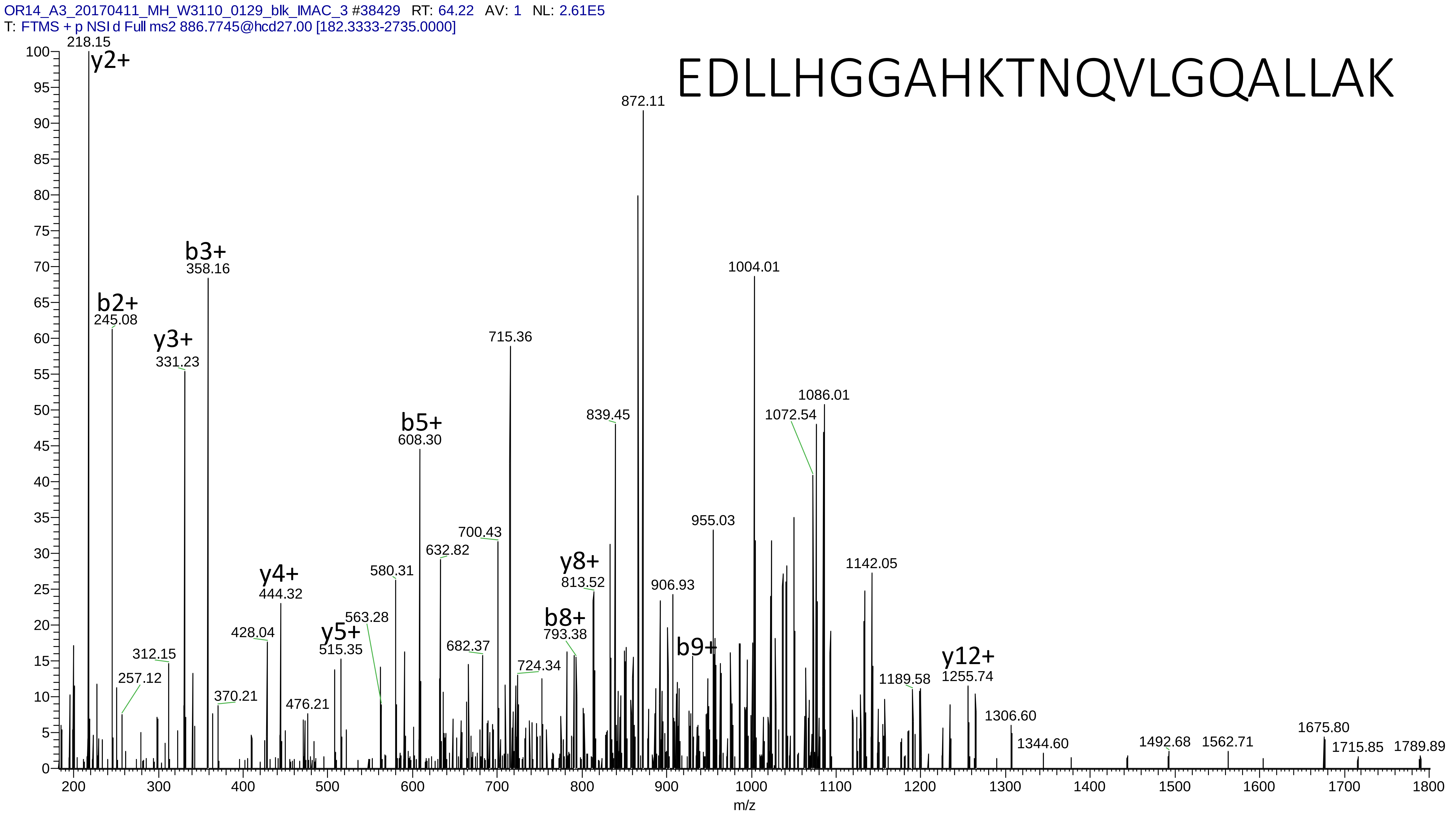
OR14_A3_20170411_MH_W3110_0129_blk_IMAC_3 #38429 RT: 64.22 AV: 1 NL: 2.61E5
T: FTMS + p NSI d Full ms2 886.7745@hcd27.00 [182.3333-2735.0000]
EDLLHGGAHKTNQVLGQALLAK
218.15 y2+
b2+ 245.08
y3+ 331.23
b3+ 358.16
257.12
312.15
370.21
428.04
y4+ 444.32
476.21
y5+ 515.35
563.28
580.31
b5+ 608.30
632.82
682.37
700.43
715.36
724.34
b8+ 793.38
y8+ 813.52
839.45
872.11
906.93
955.03
b9+
1004.01
1072.54
1086.01
1142.05
1189.58
y12+ 1255.74
1306.60
1344.60
1492.68
1562.71
1675.80
1715.85
1789.89
m/z
