## Supplementary material 2 for "MetaLab 2.0 enables accurate post-translational modifications profiling in metaproteomics"

### Supplementary material 2: Manually annotated spectra of the top 30 identified potential modifications

Dataset: Mouse_phos

# 1

- Experimental mass: 79.9662
- Unimod information: phosphorylation
- Possible composition: H O(3) P
- Sites in Unimod: S;T;Y;D;H;C;R;K
- Sites observed:
- Since phosphorylation is common in this dataset, spectra are not showed

# 2

- Experimental mass: 159.9319
- Unimod information: pyrophospho
- Possible composition: H(2)O(6)P(2)
- Sites in Unimod: S;T
- Sites observed: S;T;Y;H
- Features:  100% (1886/1886) peptides contain equal or more than two Ser/Thr

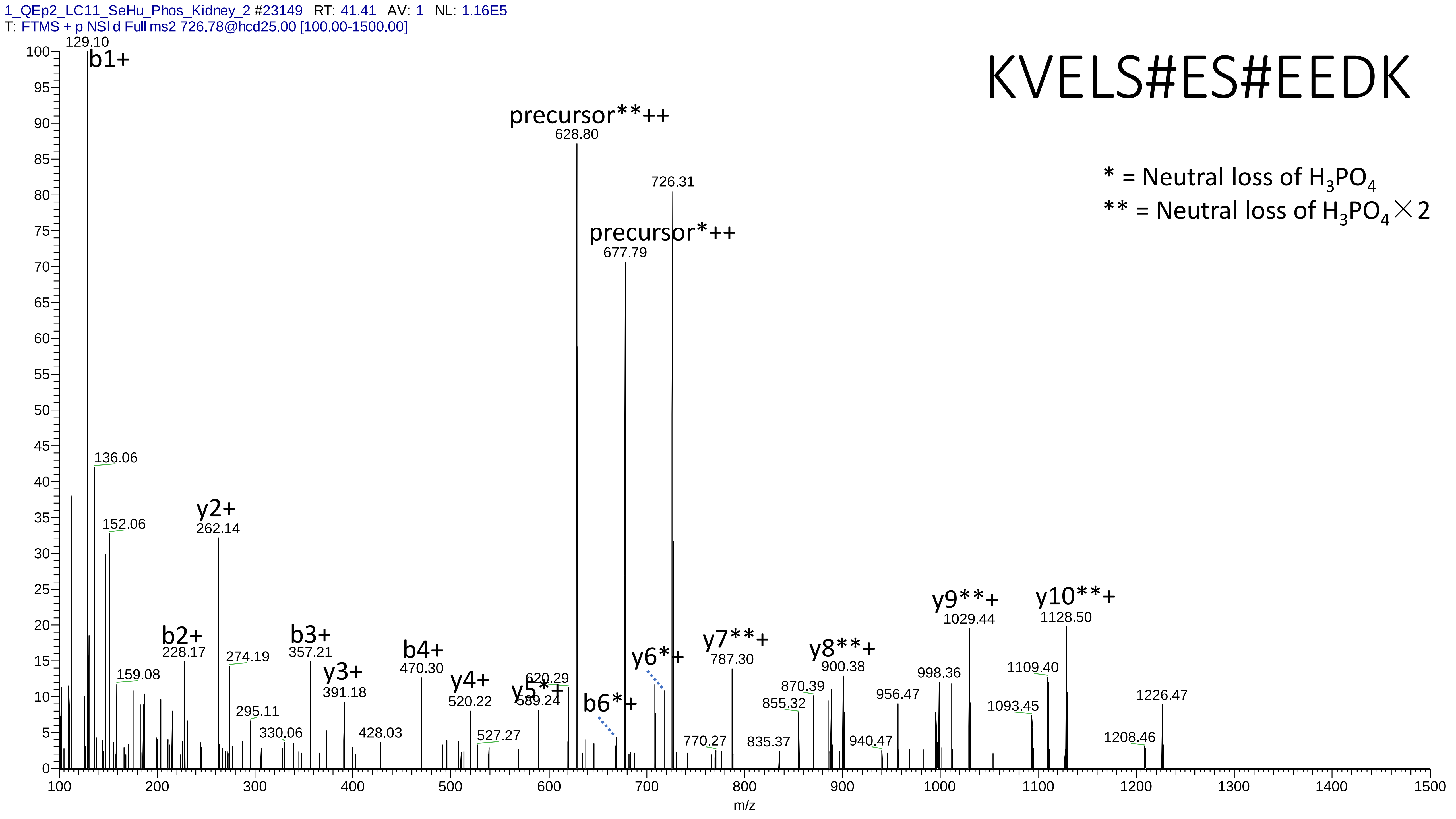
1_QEp2_LC11_SeHu_Phos_Kidney_2 #23149 RT: 41.41 AV: 1 NL: 1.16E5
T: FTMS + p NSI d Full ms2 726.78@hcd25.00 [100.00-1500.00]
KVELS#ES#EEDK
* = Neutral loss of H3PO4
** = Neutral loss of H3PO4×2
b1+ 129.10
136.06
152.06
159.08
b2+ 228.17
y2+ 262.14
274.19
295.11
330.06
b3+ 357.21
y3+ 391.18
428.03
b4+ 470.30
y4+ 520.22
527.27
y5*+ 589.24
620.29
precursor**++ 628.80
b6*+
precursor*++ 677.79
y6*+
726.31
770.27
y7**+ 787.30
835.37
855.32
870.39
y8**+ 900.38
940.47
956.47
998.36
y9**+ 1029.44
1093.45
1109.40
y10**+ 1128.50
1208.46
1226.47
m/z

# 3

- Experimental mass: 239.8979
- Unimod information:
- Possible composition: Phospho * 3, [H(1)O(3)P(1) * 3]
- Sites in Unimod:
- Sites observed:
- Features:  99.8% (482/483) peptides contain equal or more than three Ser/Thr

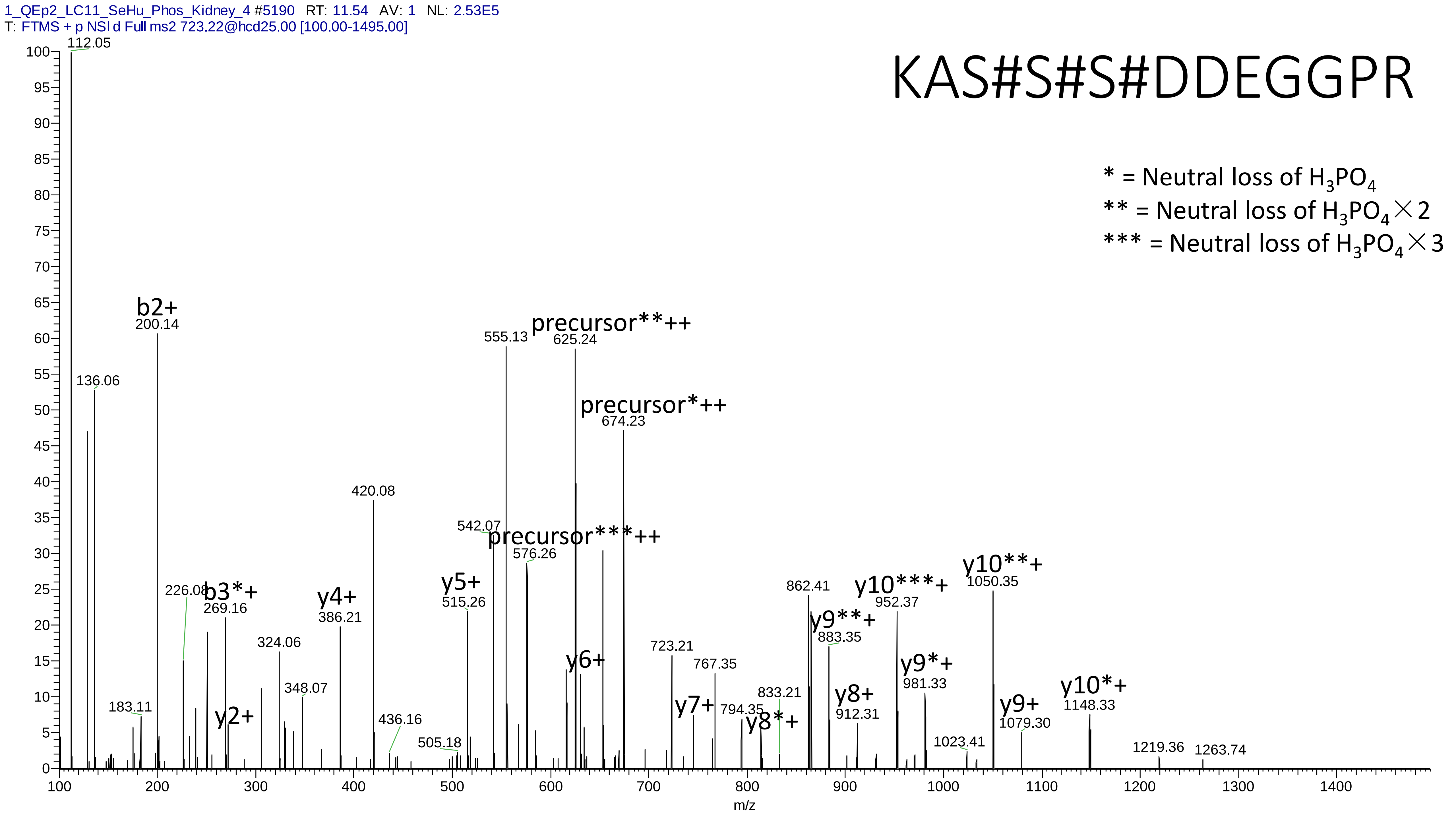
1_QEp2_LC11_SeHu_Phos_Kidney_4 #5190 RT: 11.54 AV: 1 NL: 2.53E5
T: FTMS + p NSI d Full ms2 723.22@hcd25.00 [100.00-1495.00]
KAS#S#S#DDEGGPR
* = Neutral loss of H3PO4
** = Neutral loss of H3PO4×2
*** = Neutral loss of H3PO4×3
112.05
136.06
b2+
200.14
183.11
226.08
b3*+
269.16
y2+
324.06
348.07
y4+
386.21
420.08
436.16
505.18
y5+
515.26
542.07
555.13
precursor***++
576.26
precursor**++
625.24
y6+
precursor*++
674.23
723.21
y7+
767.35
794.35
y8*+
833.21
862.41
y9**+
883.35
y8+
912.31
y10***+
952.37
y9*+
981.33
1023.41
y10**+
1050.35
y9+
1079.30
y10*+
1148.33
1219.36
1263.74
m/z

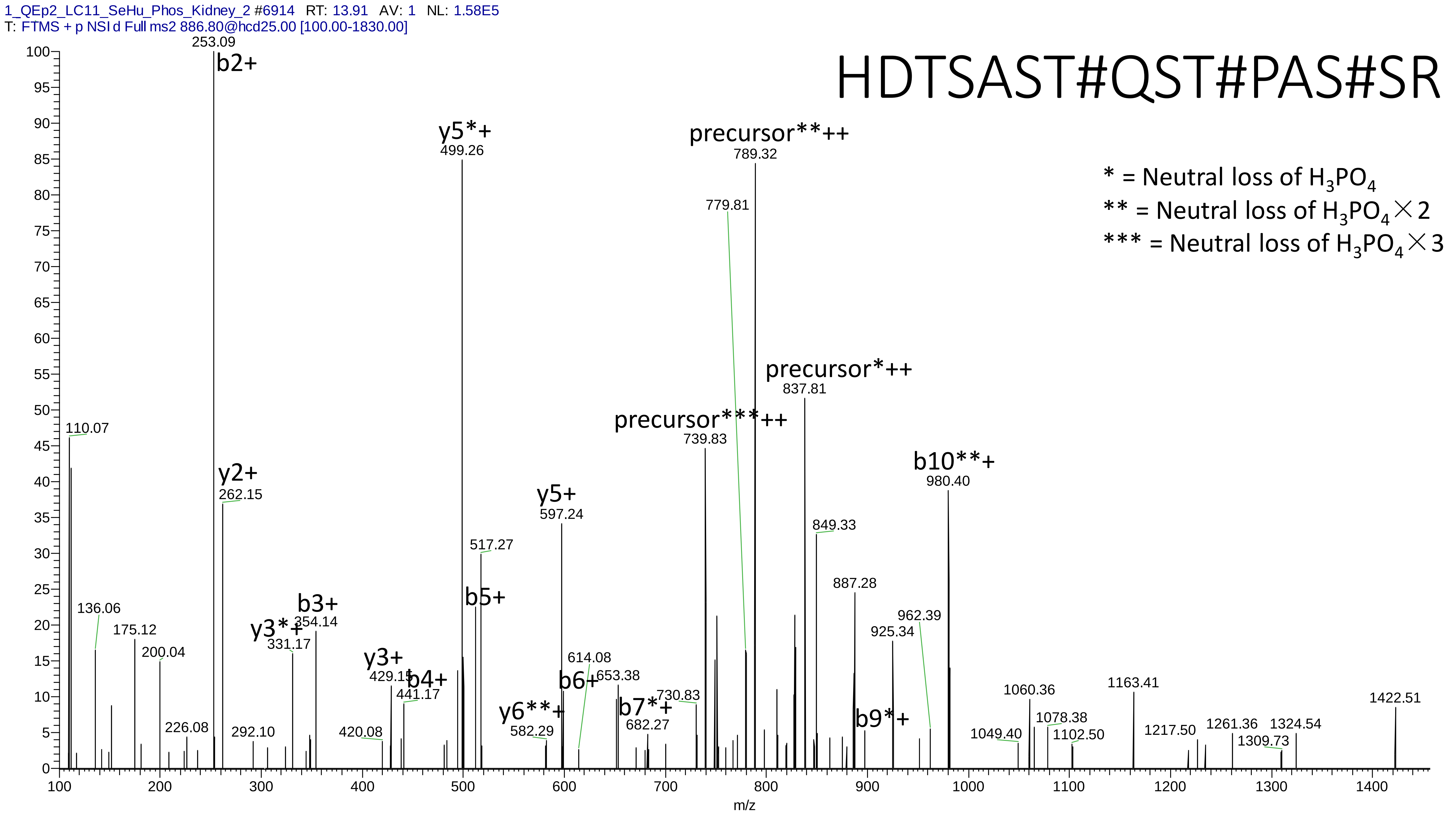
1_QEp2_LC11_SeHu_Phos_Kidney_2 #6914 RT: 13.91 AV: 1 NL: 1.58E5
T: FTMS + p NSI d Full ms2 886.80@hcd25.00 [100.00-1830.00]
HDTSAST#QST#PAS#SR
* = Neutral loss of H3PO4
** = Neutral loss of H3PO4×2
*** = Neutral loss of H3PO4×3
253.09
b2+
y5*+
499.26
precursor**++
789.32
779.81
precursor*++
837.81
precursor***++
739.83
b10**+
980.40
110.07
y2+
262.15
y5+
597.24
849.33
517.27
887.28
b5+
b3+
354.14
y3*+
331.17
136.06
175.12
200.04
962.39
925.34
y3+
429.15
b4+
441.17
614.08
653.38
b6+
730.83
b7*+
682.27
y6**+
582.29
b9*+
1060.36
1163.41
1422.51
1078.38
1102.50
1049.40
1217.50
1261.36
1324.54
1309.73
226.08
292.10
420.08
m/z

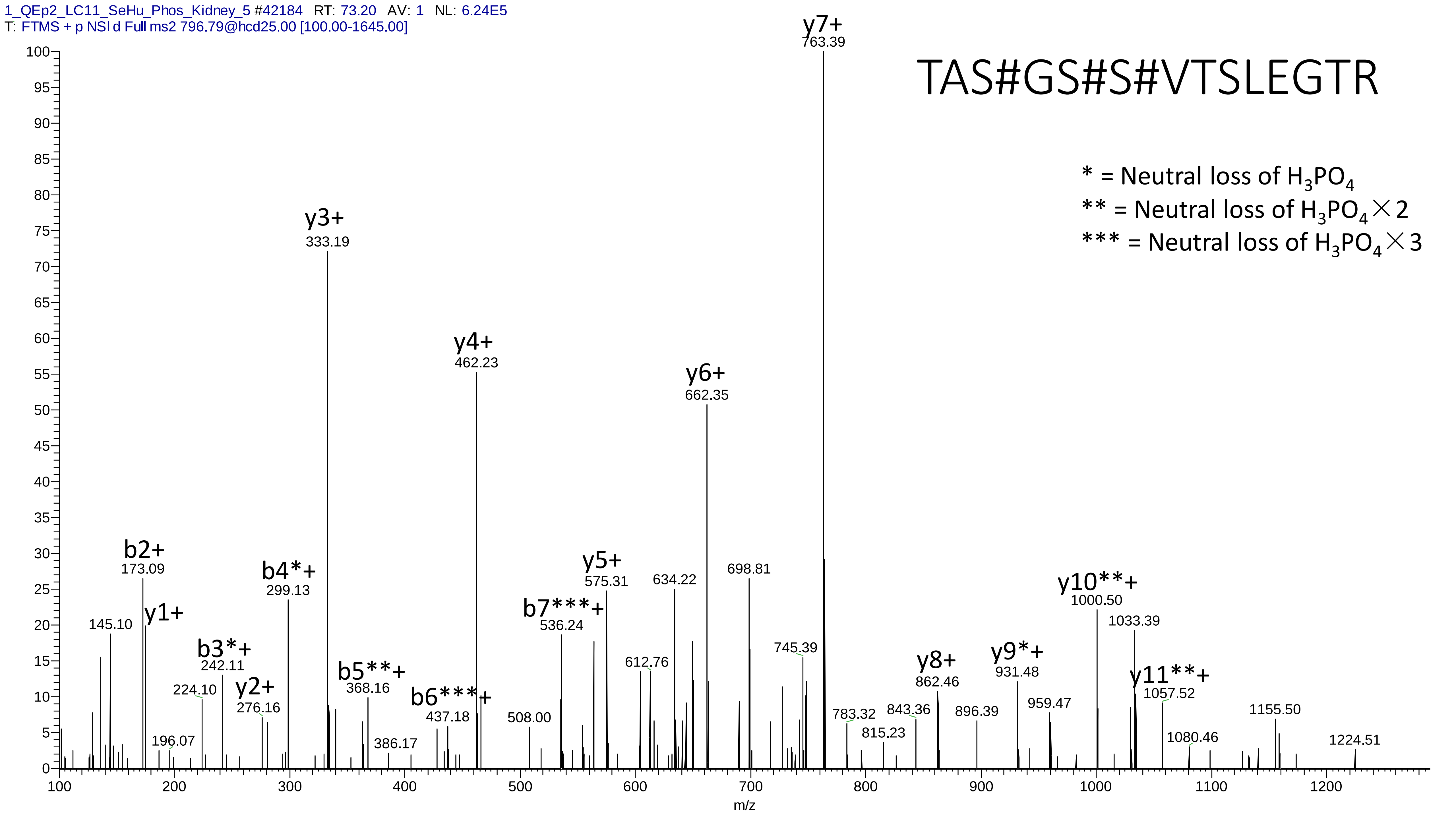
1_QEp2_LC11_SeHu_Phos_Kidney_5 #42184 RT: 73.20 AV: 1 NL: 6.24E5
T: FTMS + p NSI d Full ms2 796.79@hcd25.00 [100.00-1645.00]
TAS#GS#S#VTSLEGTR
* = Neutral loss of H3PO4
** = Neutral loss of H3PO4×2
*** = Neutral loss of H3PO4×3
y7+ 763.39
y3+ 333.19
y4+ 462.23
y6+ 662.35
b2+ 173.09
y1+
145.10
b4*+ 299.13
b3*+ 242.11
224.10
y2+ 276.16
196.07
b5**+ 368.16
386.17
b6***+ 437.18
508.00
b7***+ 536.24
y5+ 575.31
634.22
612.76
698.81
745.39
783.32
815.23
843.36
y8+ 862.46
896.39
y9*+ 931.48
959.47
y10**+ 1000.50
1033.39
y11**+ 1057.52
1080.46
1155.50
1224.51
m/z

# 4

- Experimental mass: 80.9523
- Unimod information:
- Possible composition: Phosphorylation + Deamidation, [H(1)O(3)P(1) + H(-1) N(-1) O]; Phosphorylation + Carboxymethyl replace of Carbamidomethyl on Cys
- Sites in Unimod:
- Sites observed:
- Features:  88.1% (540/613) peptides contain Asn/Gln

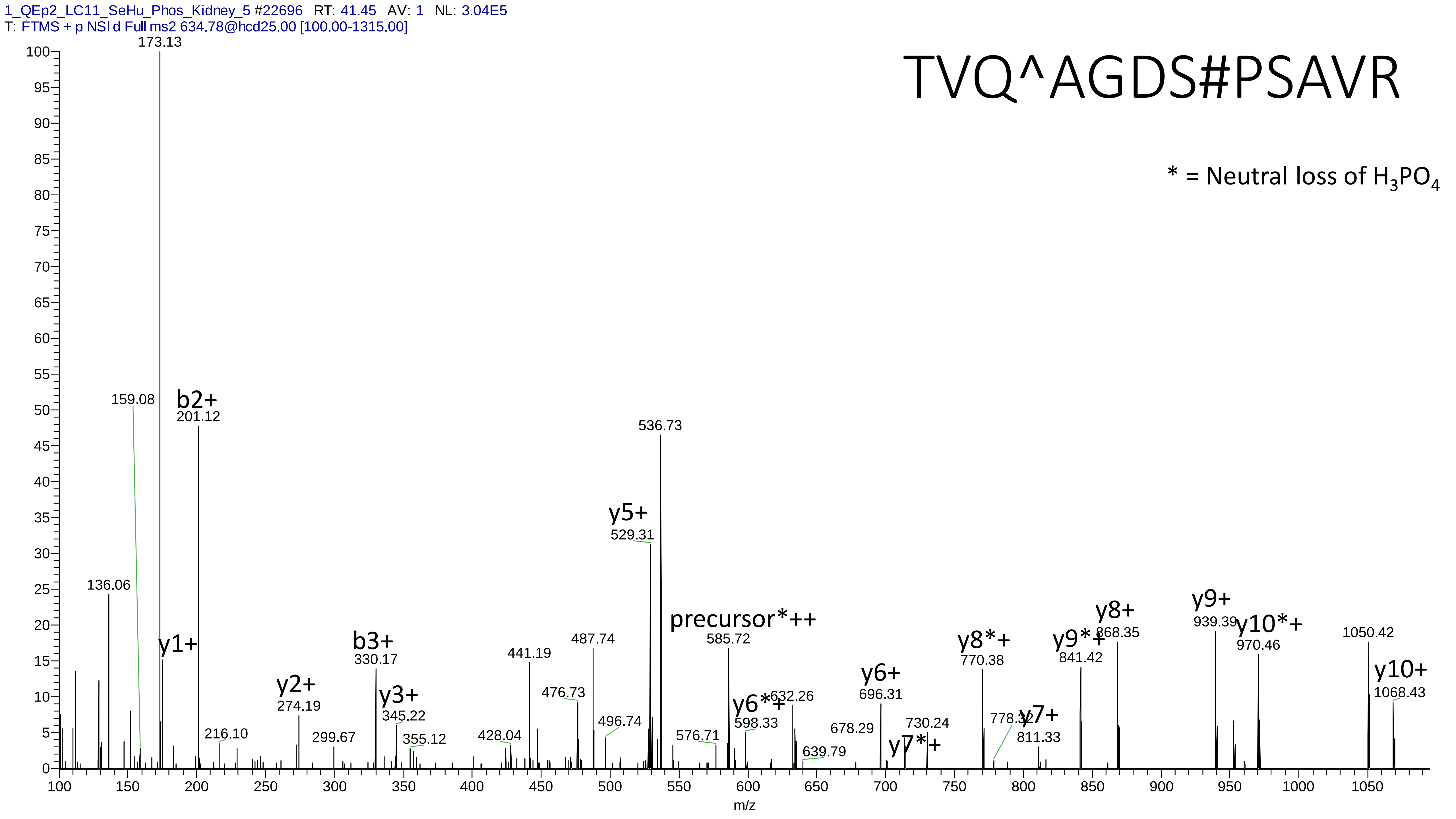
1_QEp2_LC11_SeHu_Phos_Kidney_5 #22696 RT: 41.45 AV: 1 NL: 3.04E5
T: FTMS + p NSI d Full ms2 634.78@hcd25.00 [100.00-1315.00]
TVQ^AGDS#PSAVR
* = Neutral loss of H3PO4
173.13
159.08
b2+
201.12
136.06
y1+
216.10
y2+
274.19
299.67
b3+
330.17
y3+
345.22
355.12
428.04
441.19
476.73
487.74
496.74
y5+
529.31
536.73
576.71
precursor*++
585.72
y6*+
598.33
632.26
639.79
678.29
y6+
696.31
y7*+
730.24
y8*+
770.38
778.32
y7+
811.33
y9*+
841.42
y8+
868.35
y9+
939.39
y10*+
970.46
1050.42
y10+
1068.43
m/z

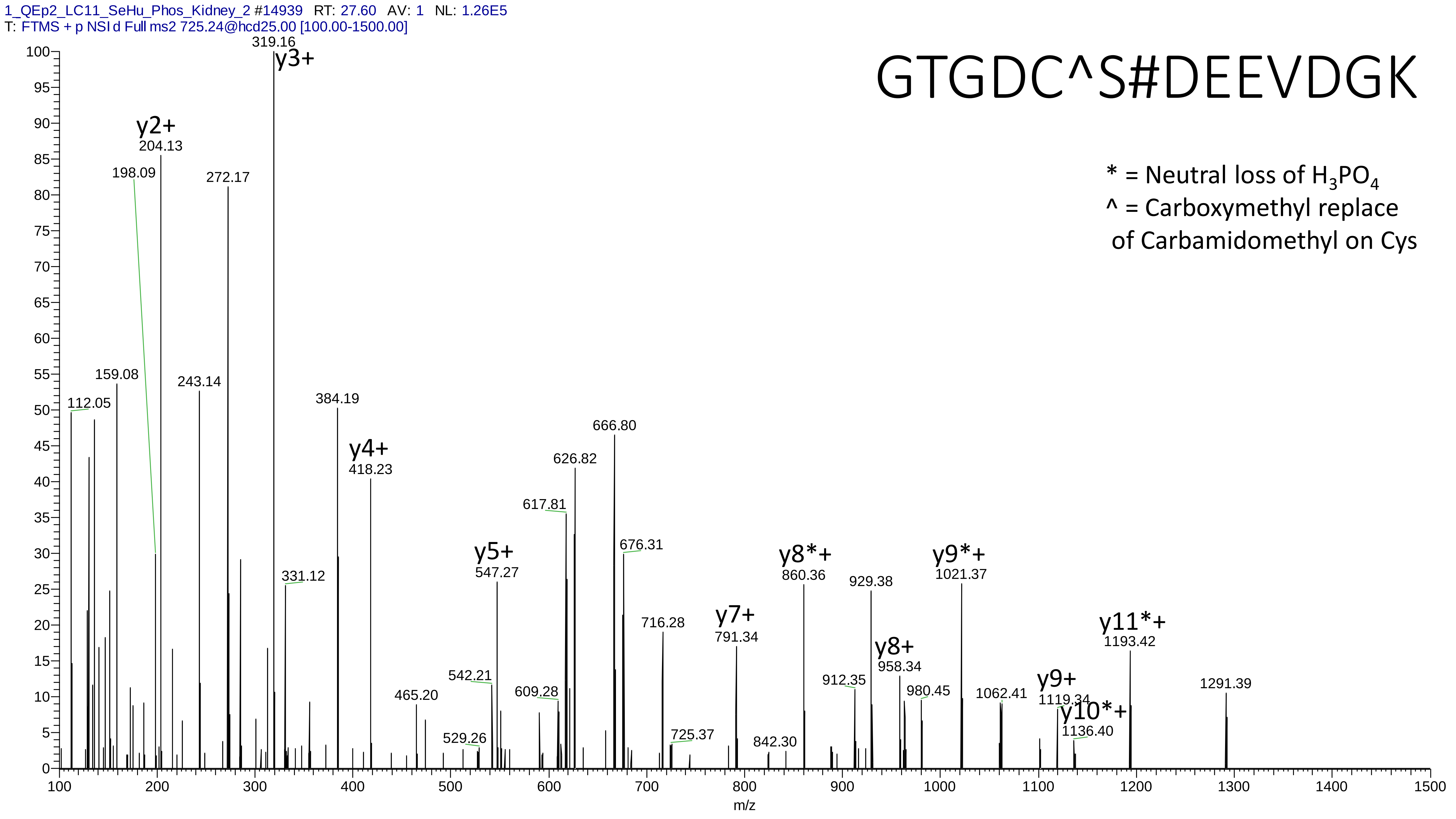
1_QEp2_LC11_SeHu_Phos_Kidney_2 #14939 RT: 27.60 AV: 1 NL: 1.26E5
T: FTMS + p NSI d Full ms2 725.24@hcd25.00 [100.00-1500.00]
GTGDC^S#DEEVDGK
* = Neutral loss of $H_3PO_4$
^ = Carboxymethyl replace
of Carbamidomethyl on Cys
y3+
319.16
y2+
204.13
198.09
272.17
159.08
243.14
112.05
384.19
y4+
418.23
331.12
465.20
542.21
529.26
y5+
547.27
617.81
626.82
666.80
609.28
676.31
716.28
725.37
y7+
791.34
842.30
y8*+
860.36
912.35
929.38
y8+
958.34
980.45
y9*+
1021.37
1062.41
y9+
1119.34
y10*+
1136.40
y11*+
1193.42
1291.39
m/z

# 5

- Experimental mass: 95.9605
- Unimod information:
- Possible composition: Phosphorylation + Oxidation, [H(1)O(3)P(1) + O]
- Sites in Unimod:
- Sites observed:

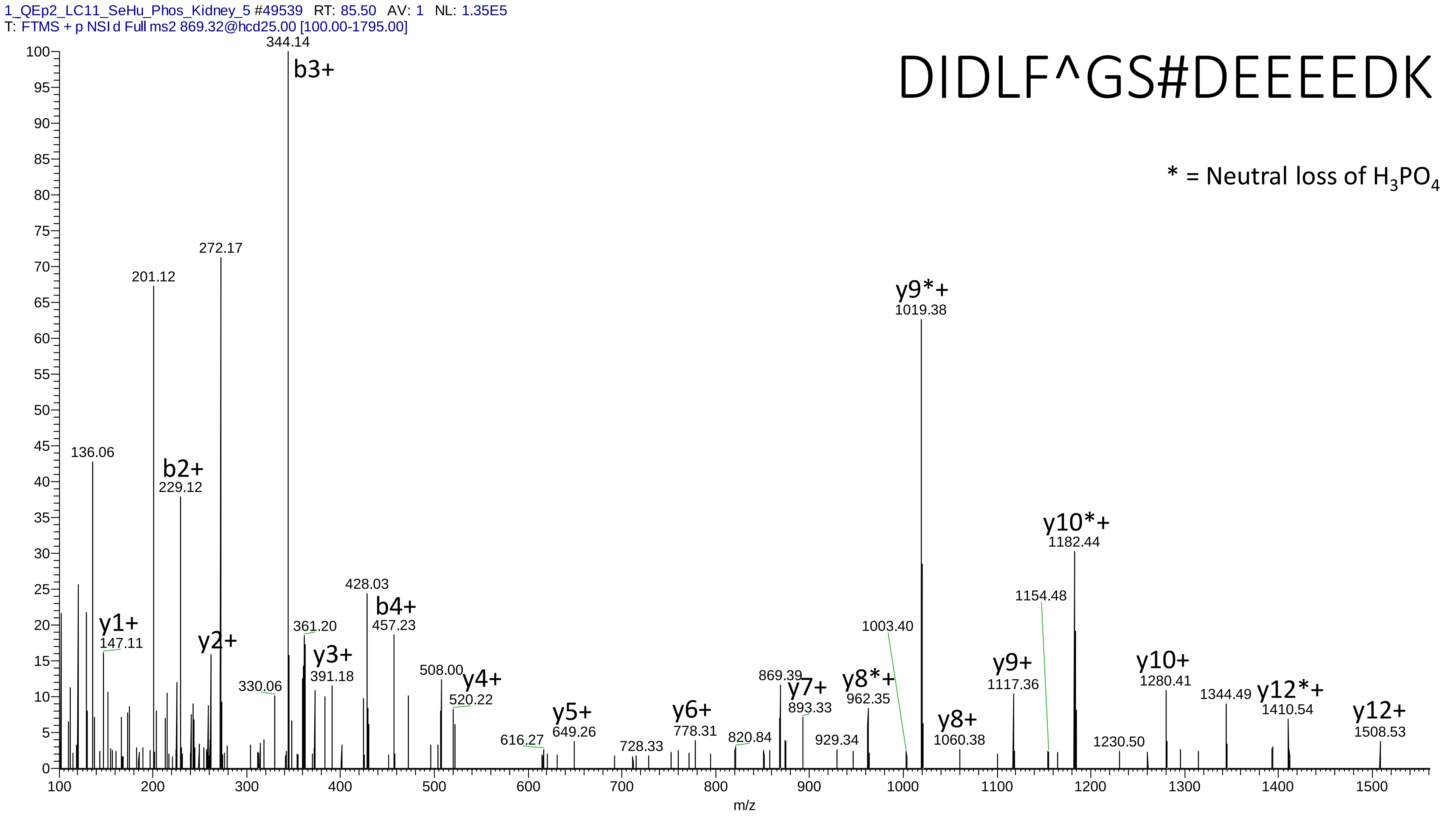
1_QEp2_LC11_SeHu_Phos_Kidney_5 #49539 RT: 85.50 AV: 1 NL: 1.35E5
T: FTMS + p NSI d Full ms2 869.32@hcd25.00 [100.00-1795.00]
DIDLF^GS#DEEEEDK
* = Neutral loss of $H_3PO_4$
344.14
b3+
272.17
201.12
136.06
b2+
229.12
y1+
147.11
y2+
361.20
y3+
391.18
330.06
428.03
b4+
457.23
508.00
y4+
520.22
616.27
y5+
649.26
728.33
y6+
778.31
820.84
869.39
y7+
893.33
929.34
y8*+
962.35
1003.40
y9*+
1019.38
y8+
1060.38
y9+
1117.36
1154.48
y10*+
1182.44
1230.50
y10+
1280.41
1344.49
y12*+
1410.54
y12+
1508.53
m/z

# 6

- Experimental mass: 319.8637
- Unimod information:
- Possible composition: Phospho * 4, [H(1)O(3)P(1) * 4]
- Sites in Unimod:
- Sites observed:
- Features:  98.5% (67/68) peptides contain equal or more than four Ser/Thr

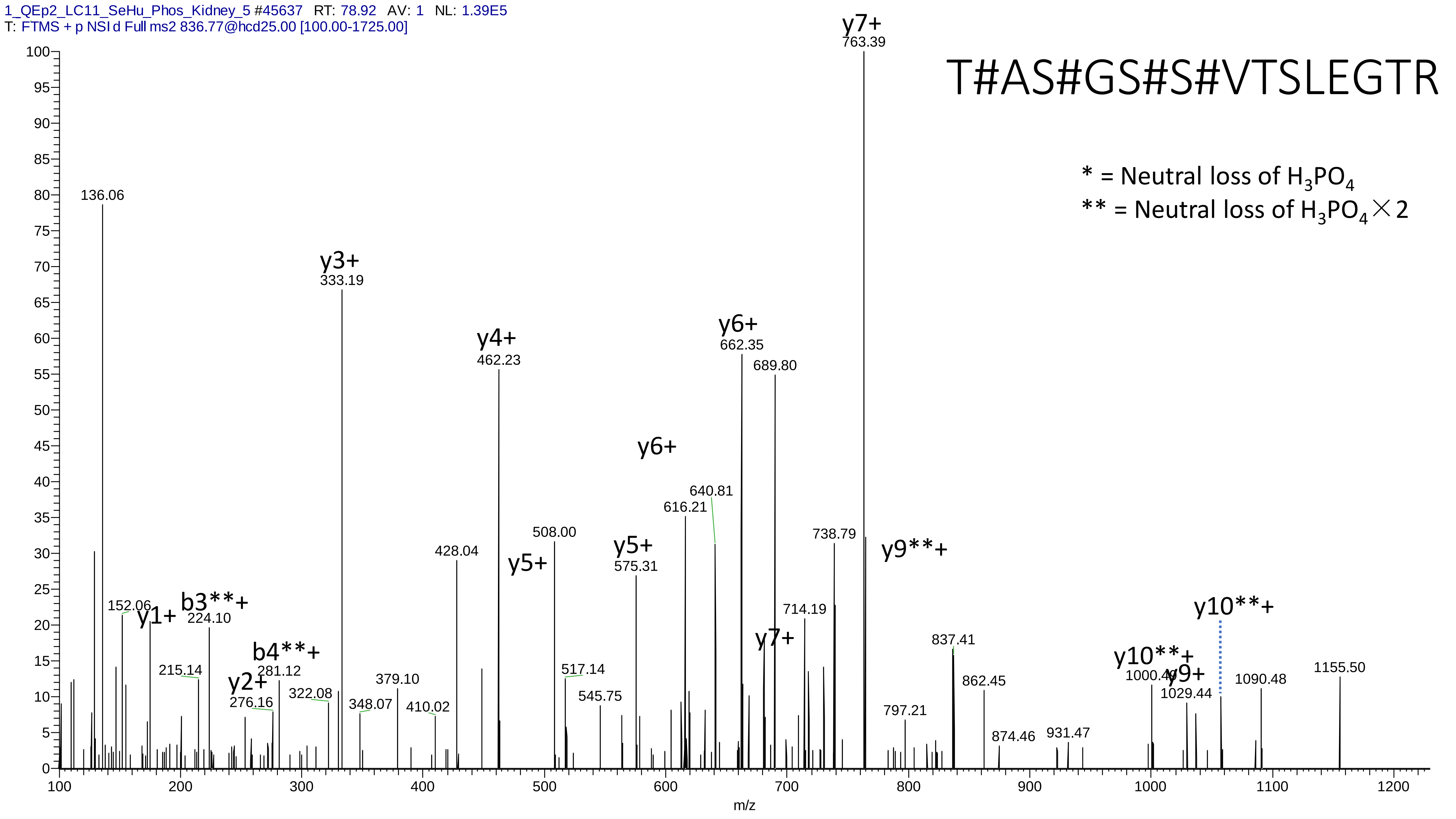
1_QEp2_LC11_SeHu_Phos_Kidney_5 #45637 RT: 78.92 AV: 1 NL: 1.39E5
T: FTMS + p NSI d Full ms2 836.77@hcd25.00 [100.00-1725.00]
T#AS#GS#S#VTSLEGTR
* = Neutral loss of H3PO4
** = Neutral loss of H3PO4 × 2
y7+ 763.39
136.06
y3+ 333.19
y4+ 462.23
y6+ 662.35
689.80
y6+
640.81
616.21
508.00
428.04
y5+
y5+ 575.31
738.79
y9**+
152.06
y1+
b3**+ 224.10
714.19
y10**+
y7+
837.41
215.14
b4**+ 281.12
517.14
y10**+
y2+ 276.16
322.08
379.10
348.07
410.02
545.75
862.45
1000.49
y9+ 1029.44
1090.48
1155.50
797.21
874.46
931.47
m/z

# 7

- Experimental mass: 119.9974
- Unimod information:
- Possible composition: Phosphorylation + Gly->Pro, [H(1)O(3)P(1) + H(4) C(3)]
- Sites in Unimod:
- Sites observed:
- Features: 97.7% (130/133) peptides contain Gly (Gly->Pro)

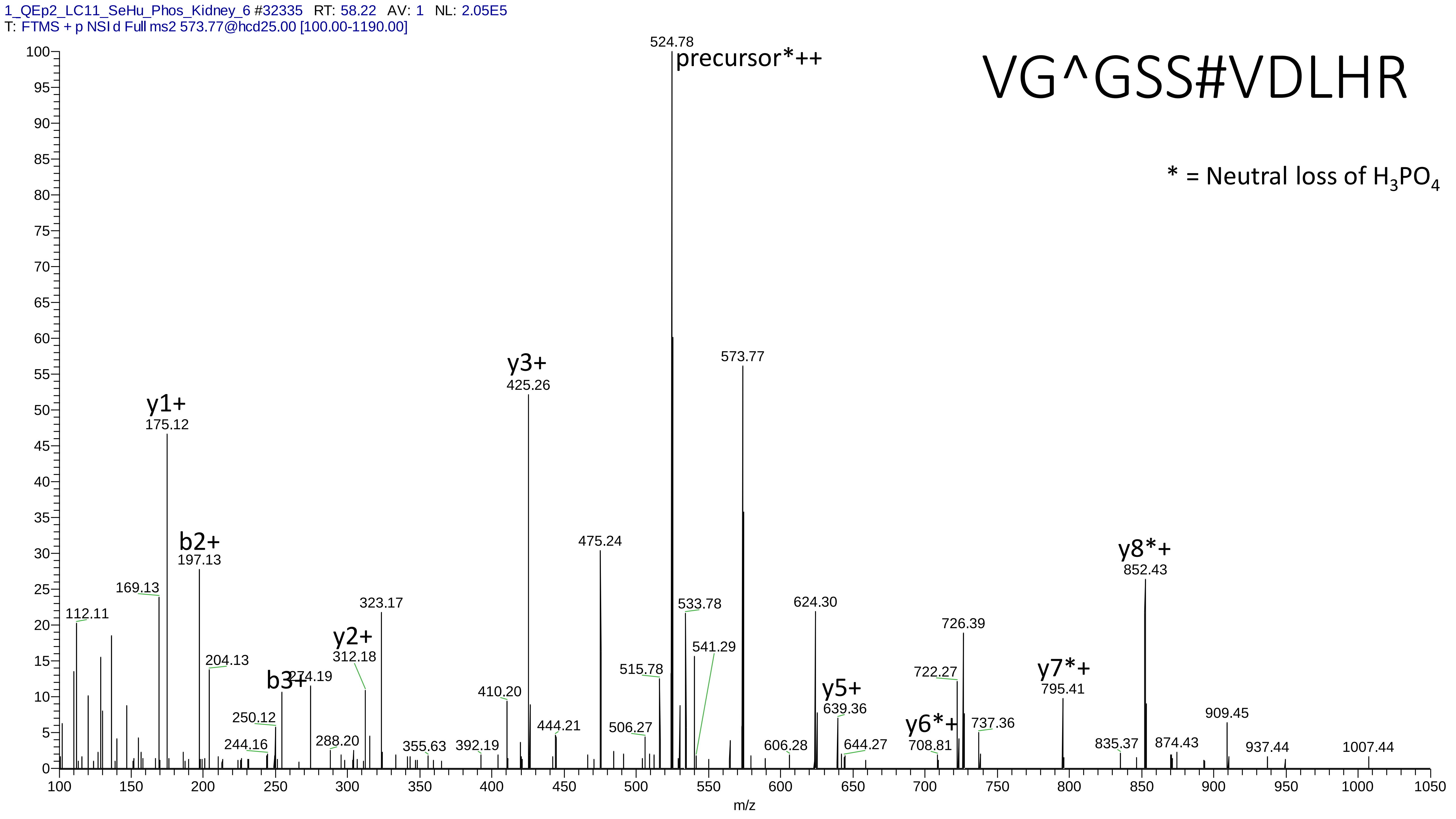
1_QEp2_LC11_SeHu_Phos_Kidney_6 #32335 RT: 58.22 AV: 1 NL: 2.05E5
T: FTMS + p NSI d Full ms2 573.77@hcd25.00 [100.00-1190.00]
VG^GSS#VDLHR
* = Neutral loss of $H_3PO_4$
precursor*++
524.78
573.77
y3+
425.26
y1+
175.12
b2+
197.13
475.24
y8*+
852.43
169.13
112.11
323.17
533.78
624.30
726.39
y2+
312.18
204.13
541.29
515.78
b3+
274.19
y5+
639.36
722.27
y7*+
795.41
410.20
250.12
444.21
506.27
y6*+
737.36
909.45
288.20
244.16
355.63
392.19
606.28
644.27
708.81
835.37
874.43
937.44
1007.44
m/z

# 8

- Experimental mass: -48.1284
- Unimod information:
- Possible composition: Phosphorylation + Lys-loss, [H(1)O(3)P(1) + H(-12) C(-6) N(-2) O(-1)]
- Sites in Unimod:
- Sites observed:
- Features:  95.8% (92/96) peptides contain Lys at peptide n-term

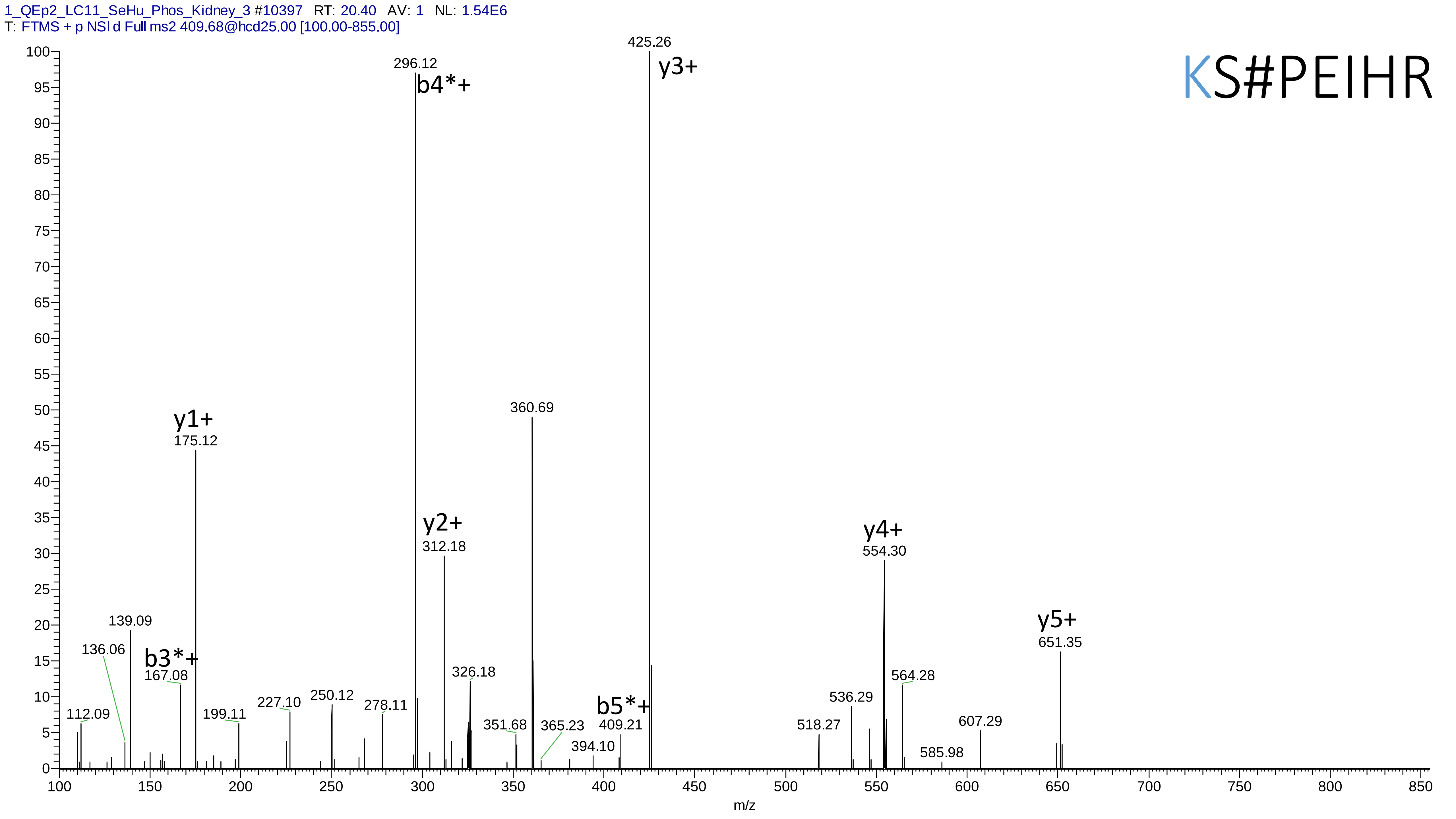
1_QEp2_LC11_SeHu_Phos_Kidney_3 #10397 RT: 20.40 AV: 1 NL: 1.54E6
T: FTMS + p NSI d Full ms2 409.68@hcd25.00 [100.00-855.00]
KS#PEIHR
425.26
y3+
296.12
b4*+
360.69
y1+
175.12
y2+
312.18
y4+
554.30
y5+
651.35
139.09
136.06
b3*+
167.08
326.18
564.28
536.29
250.12
227.10
278.11
112.09
199.11
b5*+
409.21
351.68
365.23
518.27
607.29
394.10
585.98
m/z

# 9

- Experimental mass: 62.9401
- Unimod information:
- Possible composition: Phosphorylation + Ammonia-loss, [H(1)O(3)P(1) + H(-3)N(-1)]
- Sites in Unimod:
- Sites observed:
- Features:  70.4% (81/115) peptides contain Gln at peptide n-term (Pyro-glu from Q)

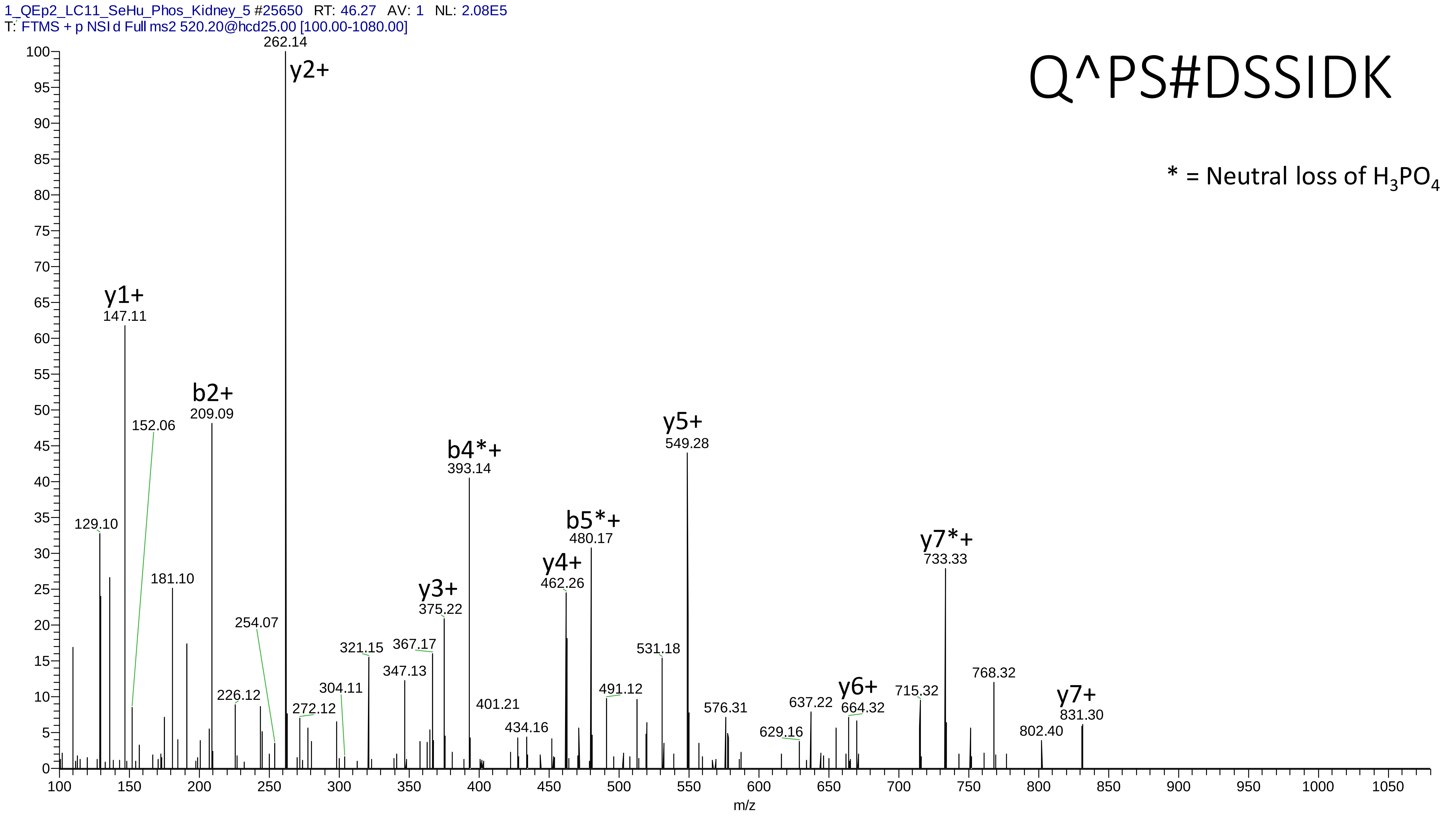
1_QEp2_LC11_SeHu_Phos_Kidney_5 #25650 RT: 46.27 AV: 1 NL: 2.08E5
T: FTMS + p NSI d Full ms2 520.20@hcd25.00 [100.00-1080.00]
Q^PS#DSSIDK
* = Neutral loss of $H_3PO_4$
y2+ 262.14
y1+ 147.11
b2+ 209.09
152.06
129.10
181.10
254.07
226.12
272.12
304.11
321.15
347.13
367.17
y3+ 375.22
b4*+ 393.14
401.21
434.16
y4+ 462.26
b5*+ 480.17
491.12
531.18
y5+ 549.28
576.31
629.16
637.22
y6+ 664.32
715.32
y7*+ 733.33
768.32
802.40
y7+ 831.30
m/z

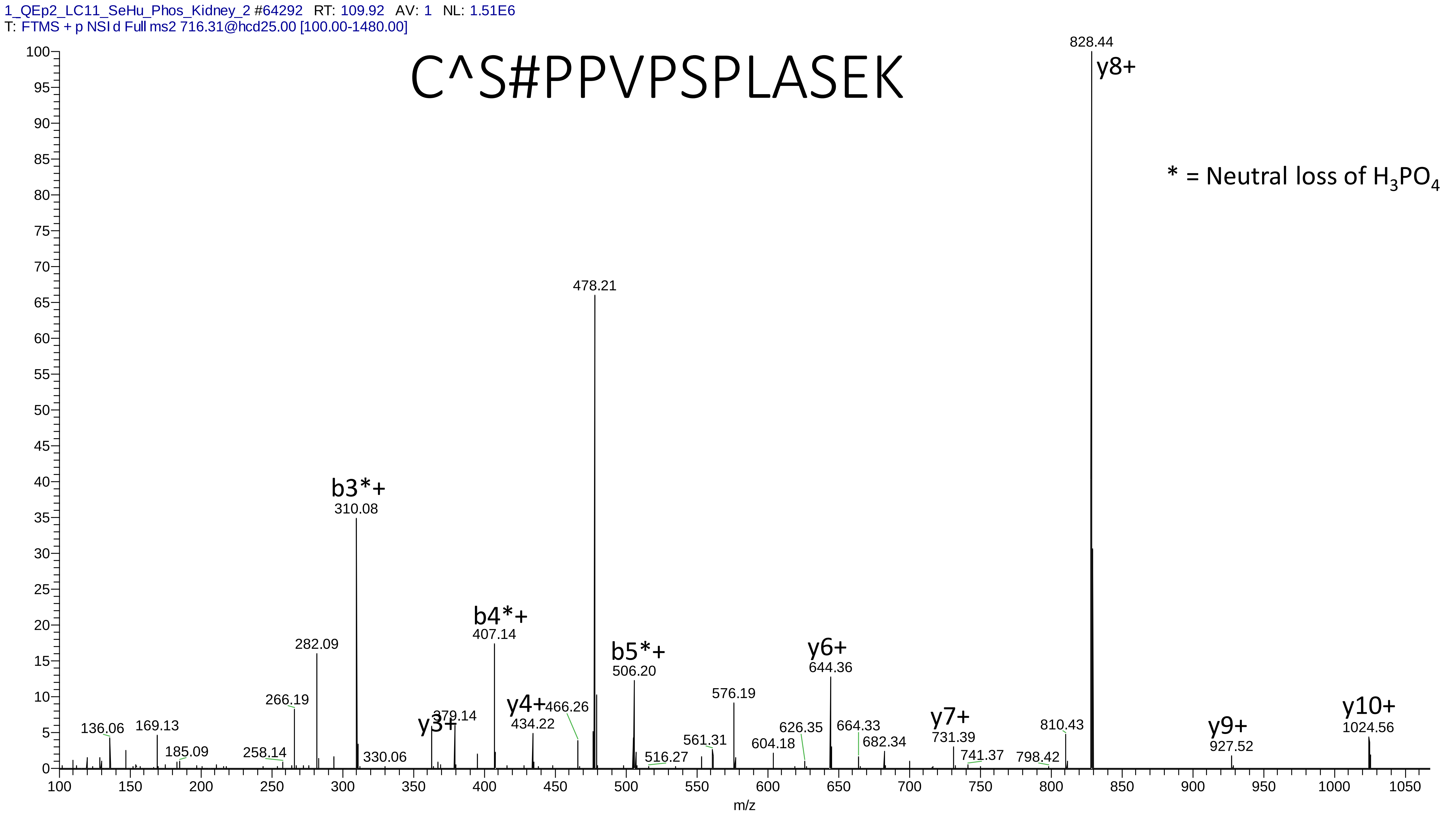

1_QEp2_LC11_SeHu_Phos_Kidney_2 #64292 RT: 109.92 AV: 1 NL: 1.51E6
T: FTMS + p NSI d Full ms2 716.31@hcd25.00 [100.00-1480.00]
C^S#PPVPSPLASEK
* = Neutral loss of $H_3PO_4$
828.44
y8+
478.21
b3*+
310.08
b4*+
407.14
282.09
b5*+
506.20
y6+
644.36
576.19
266.19
y10+
1024.56
y7+
731.39
y9+
927.52
y4+
434.22
466.26
y3+
379.14
136.06
169.13
185.09
258.14
330.06
516.27
561.31
604.18
626.35
664.33
682.34
741.37
798.42
810.43
m/z

# 10

- Experimental mass: 101.9447
- Unimod information:
- Possible composition: Phosphorylation + Cation:Na, [H(1)O(3)P(1) + H(-1) Na]
- Sites in Unimod:
- Sites observed:
- Features: phosphorylation sites determined, cation modified sites are different to be determined

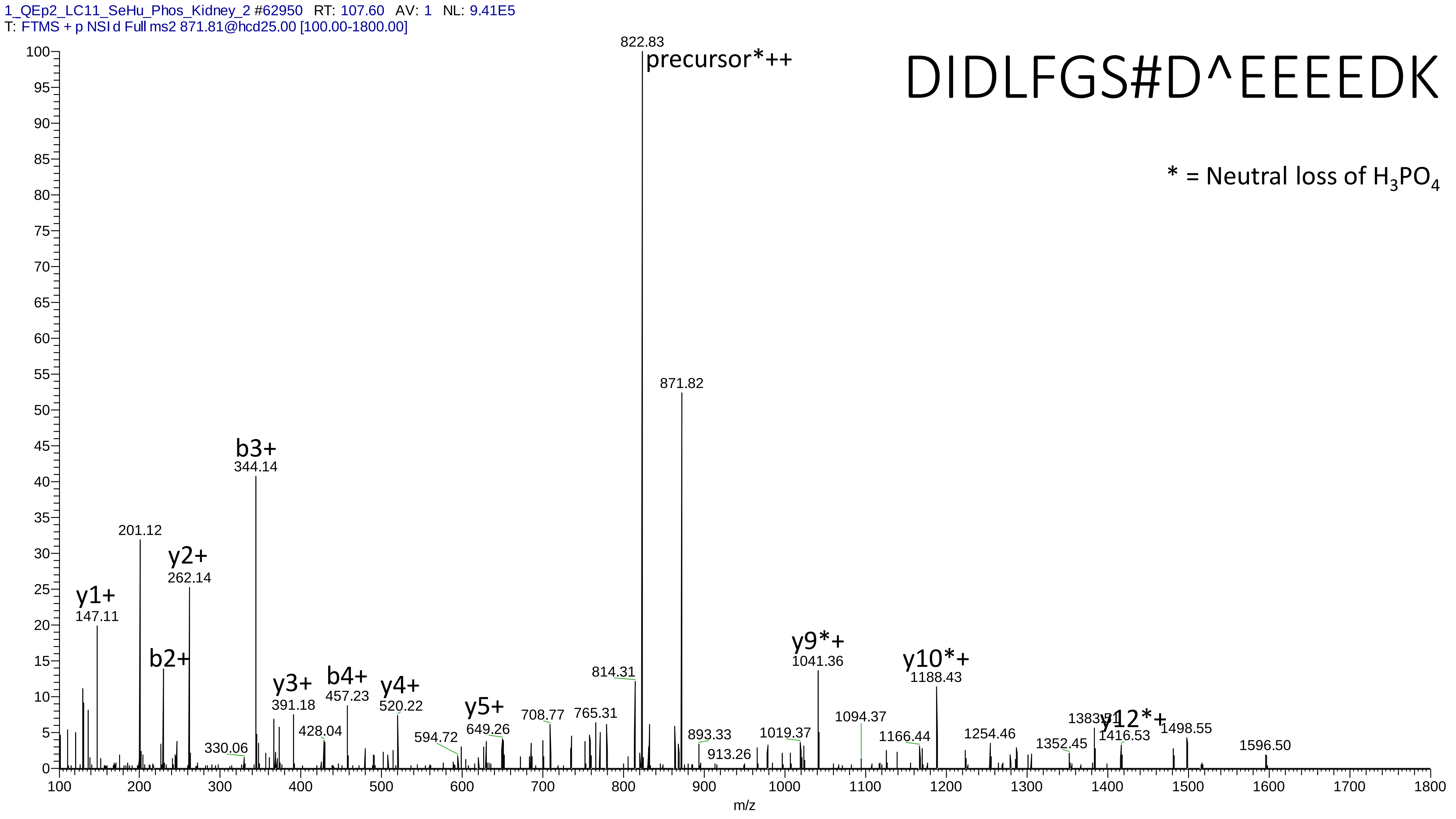
1_QEp2_LC11_SeHu_Phos_Kidney_2 #62950 RT: 107.60 AV: 1 NL: 9.41E5
T: FTMS + p NSI d Full ms2 871.81@hcd25.00 [100.00-1800.00]
DIDLFGS#D^EEEEDK
* = Neutral loss of $H_3PO_4$
822.83
precursor*++
871.82
b3+
344.14
201.12
y2+
262.14
y1+
147.11
b2+
y3+
391.18
b4+
457.23
y4+
520.22
y5+
649.26
y9*+
1041.36
y10*+
1188.43
y12*+
814.31
330.06
428.04
594.72
708.77
765.31
893.33
913.26
1019.37
1094.37
1166.44
1254.46
1352.45
1383.51
1416.53
1498.55
1596.50
m/z

# 11

- Experimental mass: 117.9104
- Unimod information:
- Possible composition: Phosphorylation + Cation:K, [H(1)O(3)P(1) + H(-1) K]
- Sites in Unimod:
- Sites observed:
- Features: phosphorylation sites determined, cation modified sites are different to be determined

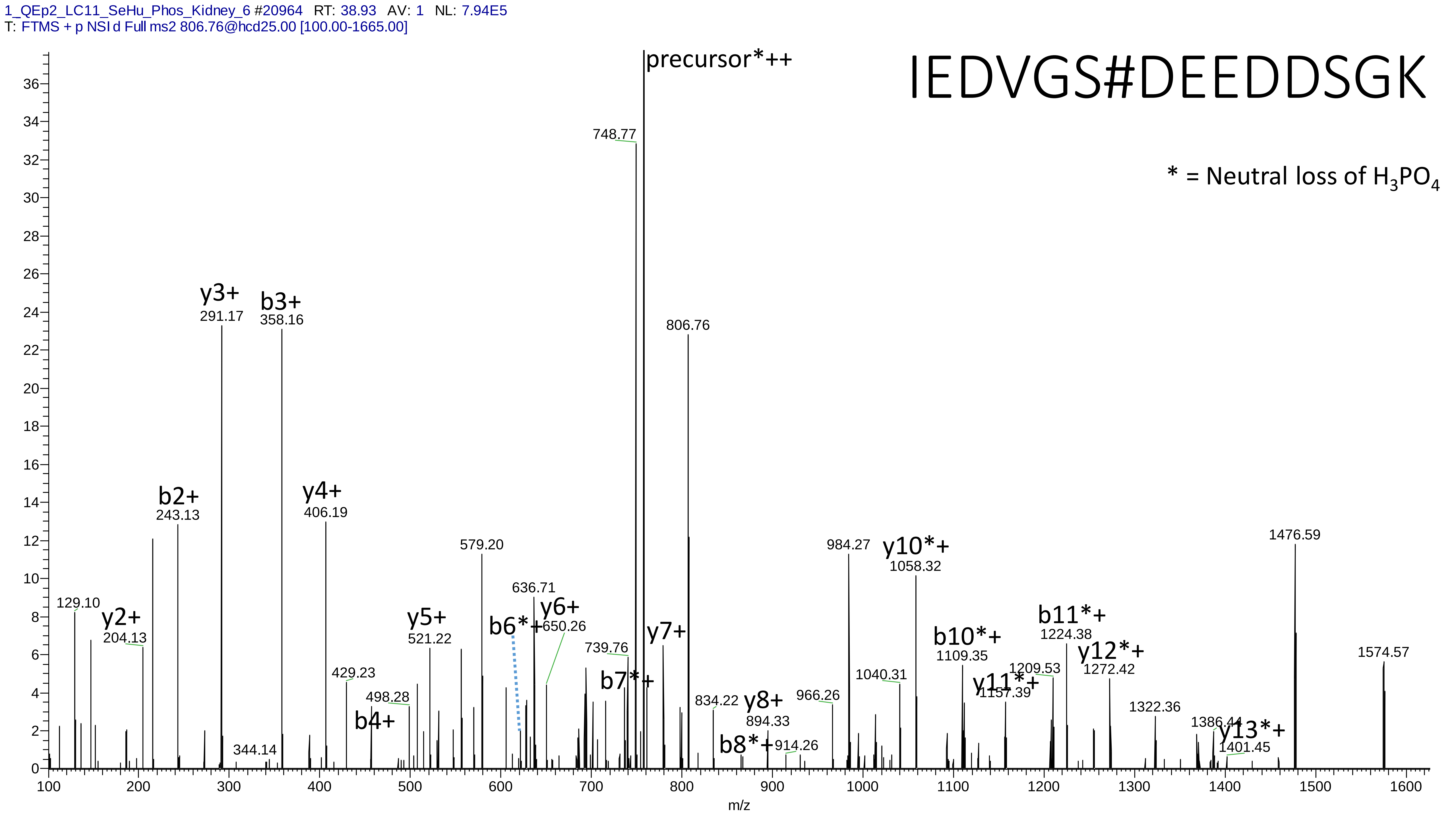
1_QEp2_LC11_SeHu_Phos_Kidney_6 #20964 RT: 38.93 AV: 1 NL: 7.94E5
T: FTMS + p NSI d Full ms2 806.76@hcd25.00 [100.00-1665.00]
IEDVGS#DEEDDSGK
* = Neutral loss of $H_3PO_4$
precursor*++
748.77
806.76
y3+
291.17
b3+
358.16
b2+
243.13
y4+
406.19
579.20
984.27
y10*+
1058.32
1476.59
1574.57
129.10
y2+
204.13
y5+
521.22
636.71
y6+
650.26
b6*+
739.76
y7+
b7*+
429.23
498.28
b4+
344.14
834.22
y8+
894.33
b8*+
914.26
966.26
1040.31
b10*+
1109.35
y11*+
1157.39
1209.53
b11*+
1224.38
y12*+
1272.42
1322.36
1386.44
y13*+
1401.45
m/z

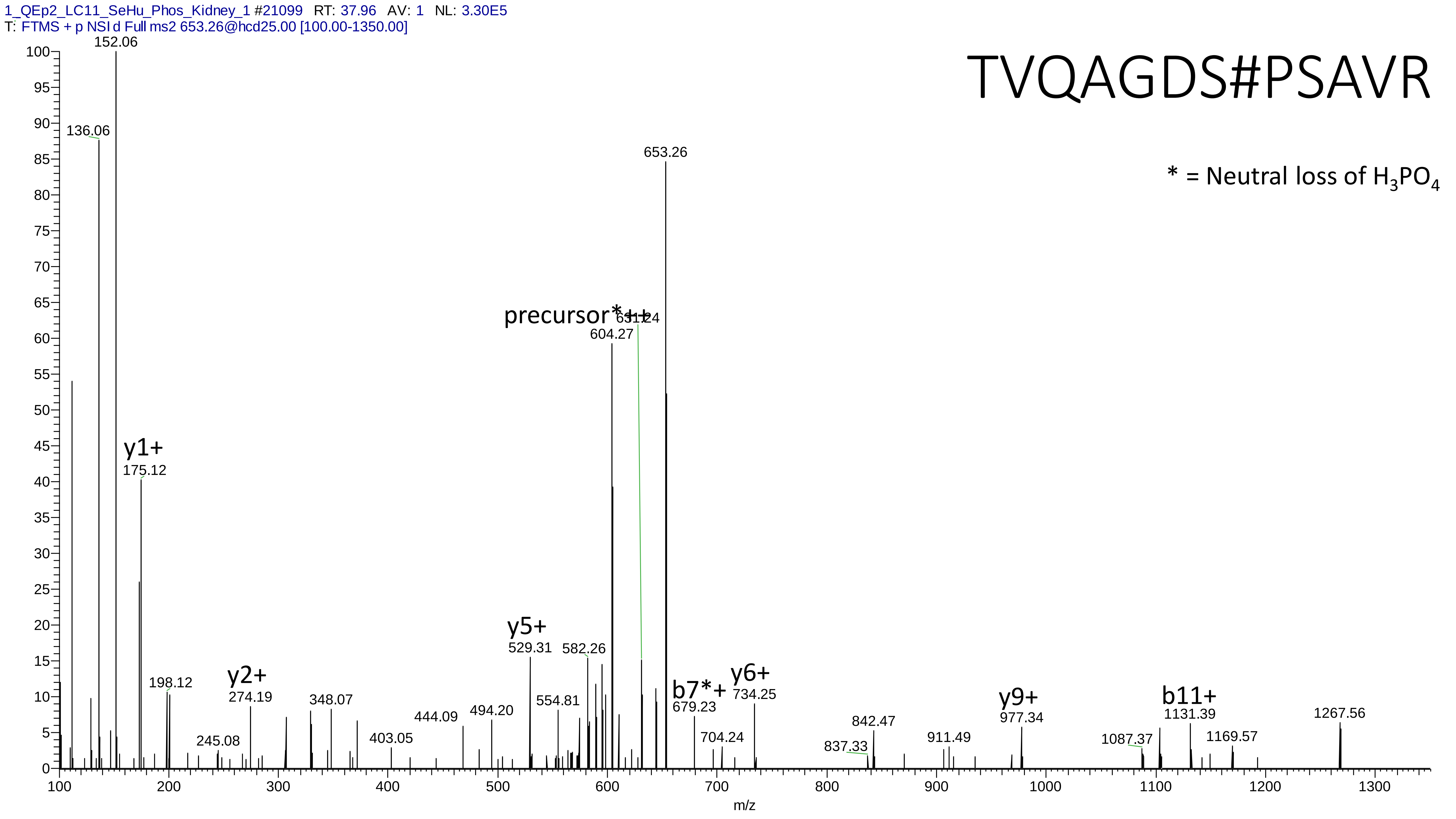
1_QEp2_LC11_SeHu_Phos_Kidney_1 #21099 RT: 37.96 AV: 1 NL: 3.30E5
T: FTMS + p NSI d Full ms2 653.26@hcd25.00 [100.00-1350.00]
TVQAGDS#PSAVR
* = Neutral loss of $H_3PO_4$
152.06
136.06
653.26
precursor*++
604.27
y1+
175.12
198.12
y2+
274.19
245.08
348.07
403.05
444.09
494.20
y5+
529.31
554.81
582.26
b7*+
679.23
704.24
y6+
734.25
837.33
842.47
911.49
y9+
977.34
1087.37
b11+
1131.39
1169.57
1267.56
m/z

# 12

- Experimental mass: 111.9548
- Unimod information:
- Possible composition: Phosphorylation + Dioxidation, [H(1)O(3)P(1) + O(2)]
- Sites in Unimod:
- Sites observed:

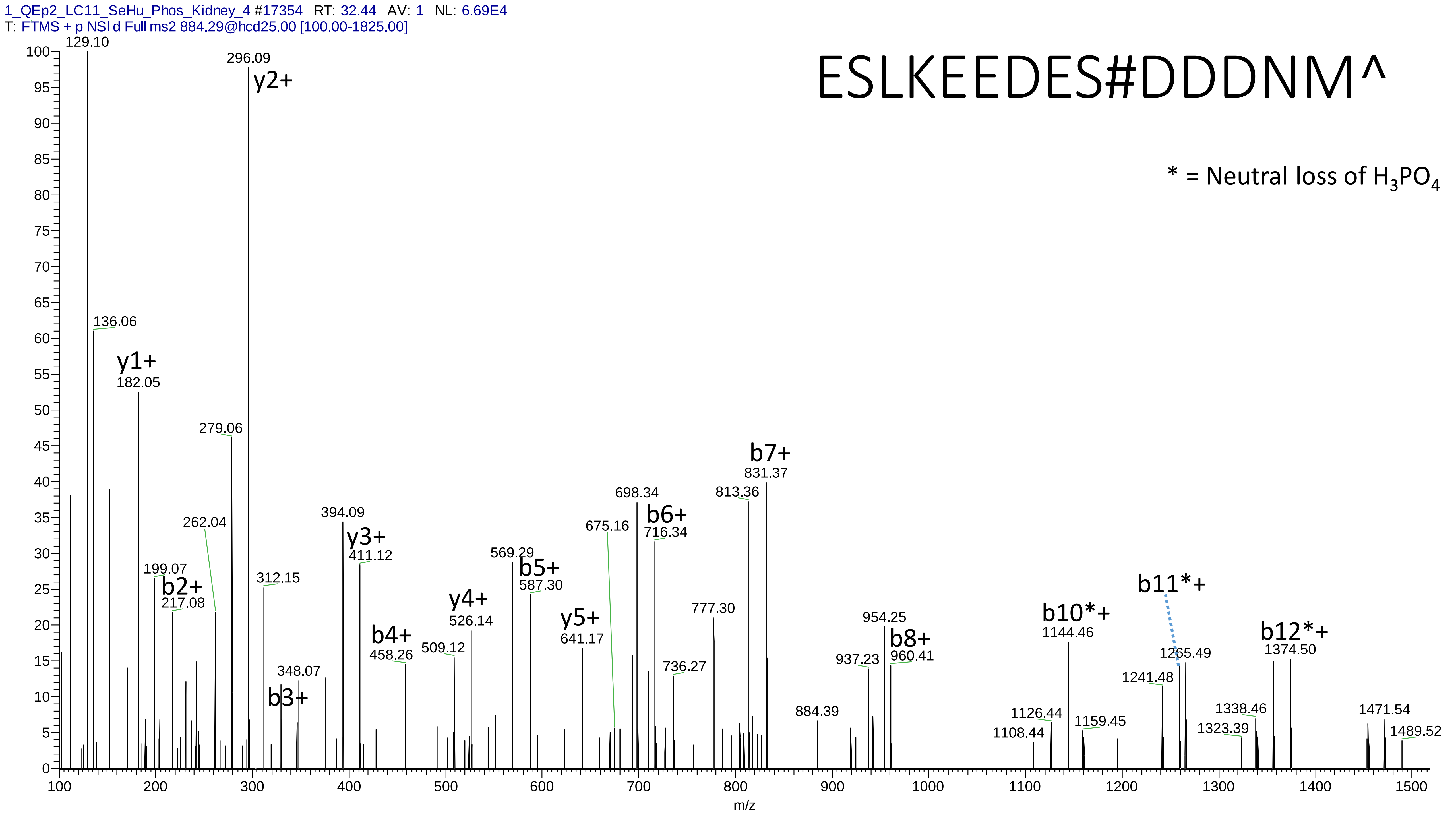
1_QEp2_LC11_SeHu_Phos_Kidney_4 #17354 RT: 32.44 AV: 1 NL: 6.69E4
T: FTMS + p NSI d Full ms2 884.29@hcd25.00 [100.00-1825.00]
ESLKEEDES#DDDNM^
* = Neutral loss of H3PO4
129.10
296.09
y2+
136.06
y1+
182.05
279.06
262.04
199.07
b2+
217.08
312.15
348.07
b3+
394.09
y3+
411.12
b4+
458.26
509.12
y4+
526.14
569.29
b5+
587.30
y5+
641.17
675.16
698.34
b6+
716.34
736.27
777.30
813.36
b7+
831.37
884.39
937.23
954.25
b8+
960.41
1108.44
1126.44
b10*+
1144.46
1159.45
1241.48
b11*+
1265.49
1323.39
1338.46
b12*+
1374.50
1471.54
1489.52
m/z

# 13

- Experimental mass: 175.9278
- Unimod information:
- Possible composition: Phosphorylation * 2 + Oxidation, [H(1)O(3)P(1) * 2 + O]
- Sites in Unimod:
- Sites observed:

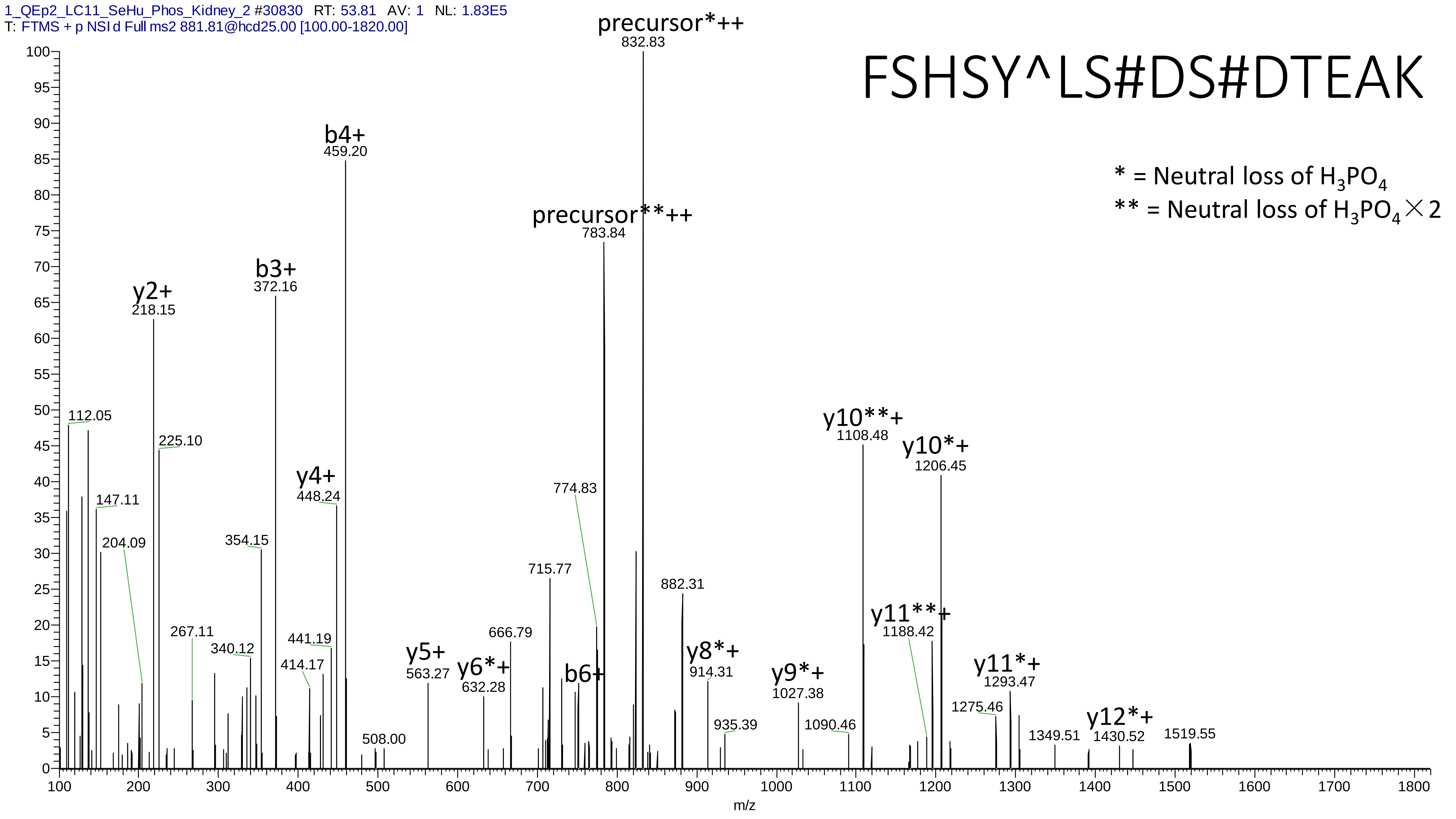

1_QEp2_LC11_SeHu_Phos_Kidney_2 #30830 RT: 53.81 AV: 1 NL: 1.83E5
T: FTMS + p NSI d Full ms2 881.81@hcd25.00 [100.00-1820.00]
FSHSY^LS#DS#DTEAK
* = Neutral loss of $H_3PO_4$
** = Neutral loss of $H_3PO_4 \times 2$
precursor*++ 832.83
precursor**++ 783.84
b4+ 459.20
b3+ 372.16
y2+ 218.15
112.05
225.10
147.11
204.09
354.15
y4+ 448.24
774.83
715.77
882.31
y10**+ 1108.48
y10*+ 1206.45
y11**+ 1188.42
267.11
340.12
441.19
414.17
y5+ 563.27
y6*+ 632.28
666.79
b6+
y8*+ 914.31
y9*+ 1027.38
y11*+ 1293.47
1275.46
935.39
1090.46
508.00
1349.51
y12*+ 1430.52
1519.55
m/z

# 14

- Experimental mass: 181.9112
- Unimod information:
- Possible composition: Phosphorylation * 2 + Cation:Na, [H(1)O(3)P(1) * 2 + H(-1) Na ]
- Sites in Unimod:
- Sites observed:
- Features: phosphorylation sites determined, cation modified sites are different to be determined

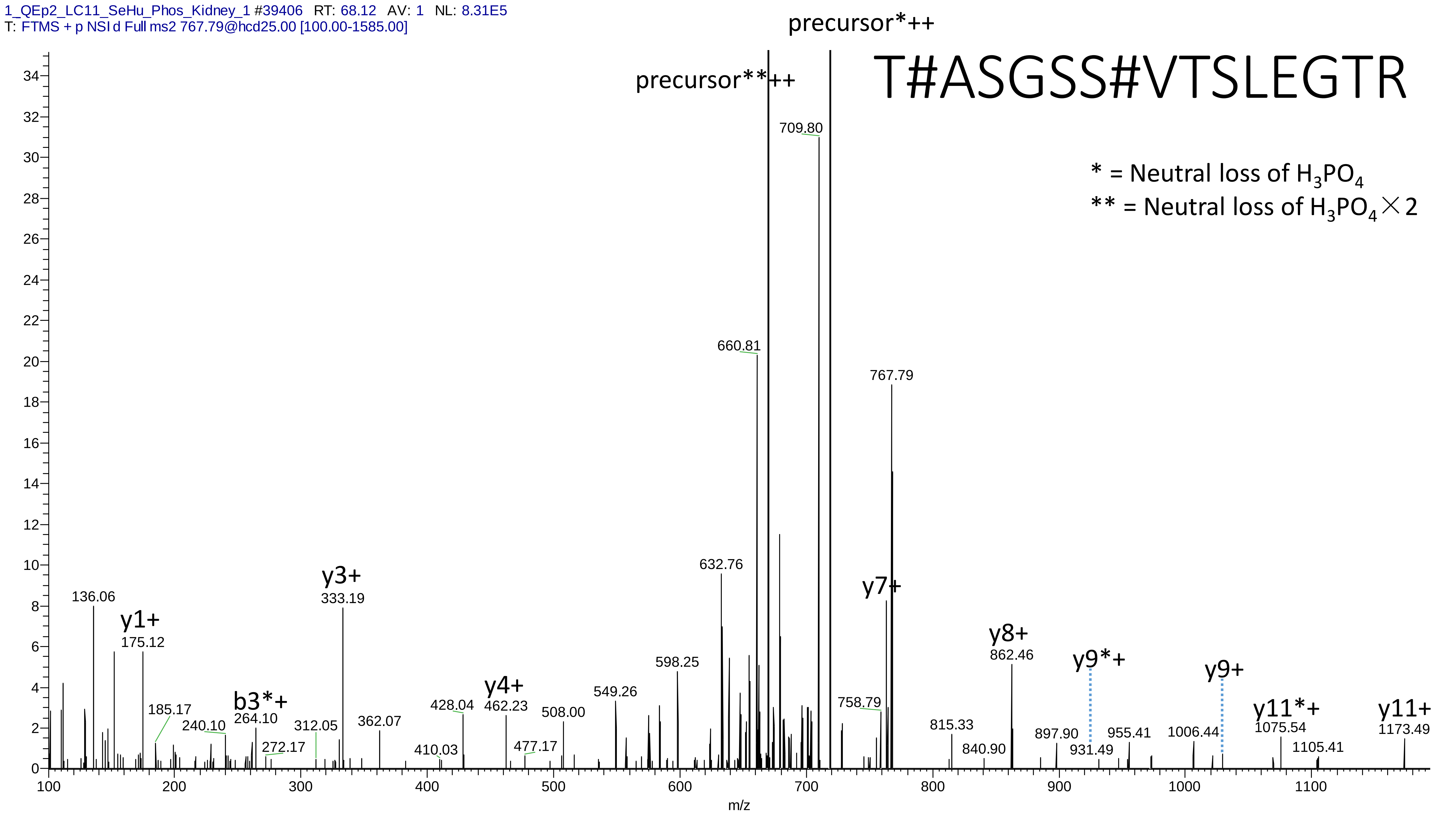
1_QEp2_LC11_SeHu_Phos_Kidney_1 #39406 RT: 68.12 AV: 1 NL: 8.31E5
T: FTMS + p NSI d Full ms2 767.79@hcd25.00 [100.00-1585.00]
T#ASGSS#VTSLEGTR
* = Neutral loss of $H_3PO_4$
** = Neutral loss of $H_3PO_4 \times 2$
precursor*++
precursor**++
709.80
660.81
767.79
632.76
y7+
136.06
y1+
175.12
y3+
333.19
185.17
240.10
b3*+
264.10
272.17
312.05
362.07
410.03
428.04
y4+
462.23
477.17
508.00
549.26
598.25
758.79
815.33
840.90
y8+
862.46
897.90
y9*+
931.49
955.41
1006.44
y9+
y11*+
1075.54
1105.41
y11+
1173.49
m/z

# 15

- Experimental mass: 31.8376
- Unimod information:
- Possible composition: Phosphorylation * 2 + Lys-loss, [H(1)O(3)P(1) * 2 + H(-12) C(-6) N(-2) O(-1)]
- Sites in Unimod:
- Sites observed: 97.8% (44/45) peptides contain Lys at peptide n-term;

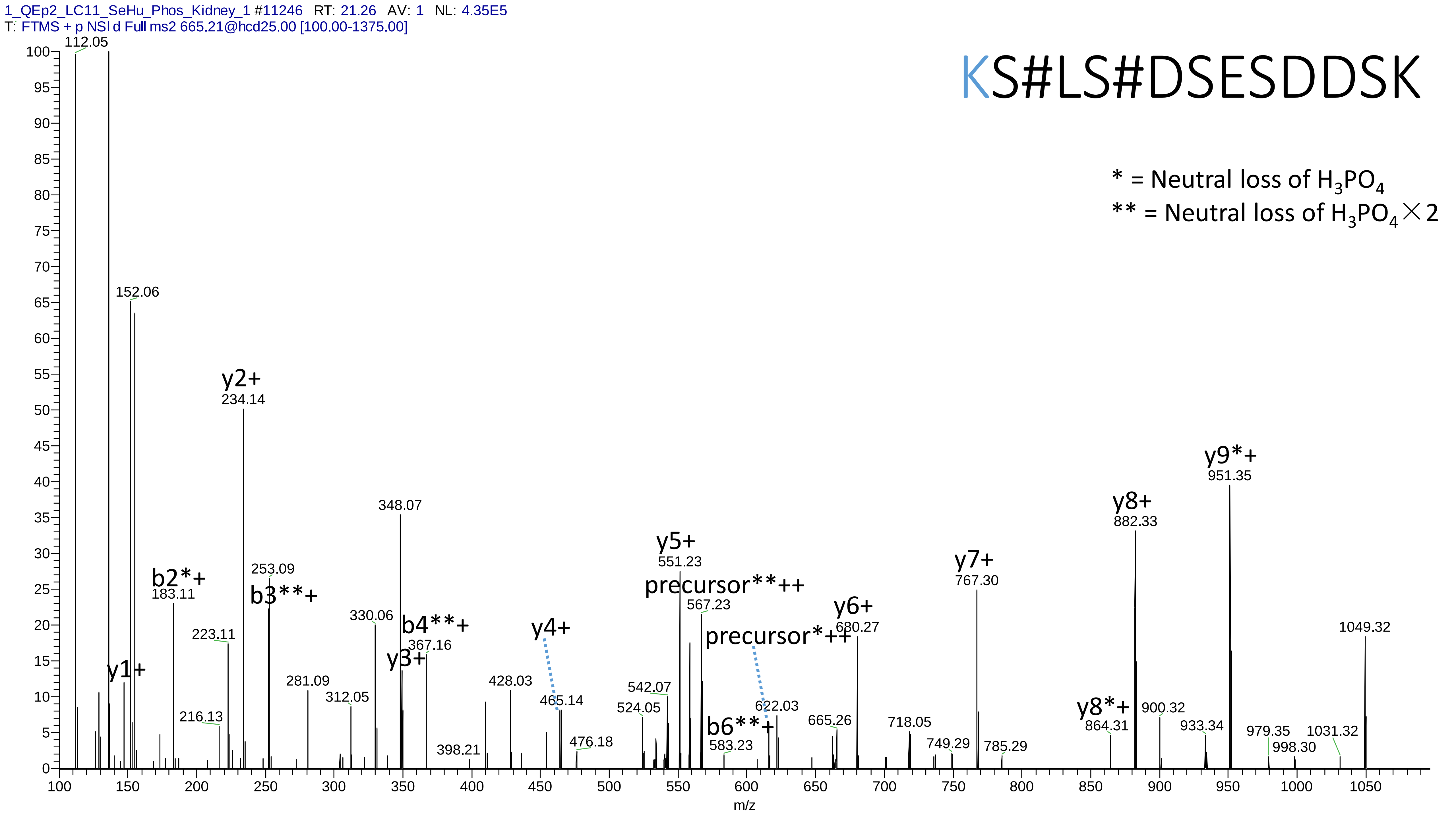

1_QEp2_LC11_SeHu_Phos_Kidney_1 #11246 RT: 21.26 AV: 1 NL: 4.35E5
T: FTMS + p NSI d Full ms2 665.21@hcd25.00 [100.00-1375.00]
KS#LS#DSESDDSK
* = Neutral loss of $H_3PO_4$
** = Neutral loss of $H_3PO_4 \times 2$
112.05
152.06
y2+
234.14
348.07
y9*+
951.35
y8+
882.33
y5+
551.23
y7+
767.30
precursor**++
567.23
b2*+
183.11
253.09
b3**+
330.06
b4**+
367.16
y3+
y1+
y4+
y6+
680.27
precursor*++
1049.32
223.11
281.09
312.05
428.03
465.14
542.07
524.05
622.03
b6**+
583.23
665.26
718.05
y8*+
864.31
900.32
933.34
979.35
998.30
1031.32
216.13
398.21
476.18
749.29
785.29
m/z

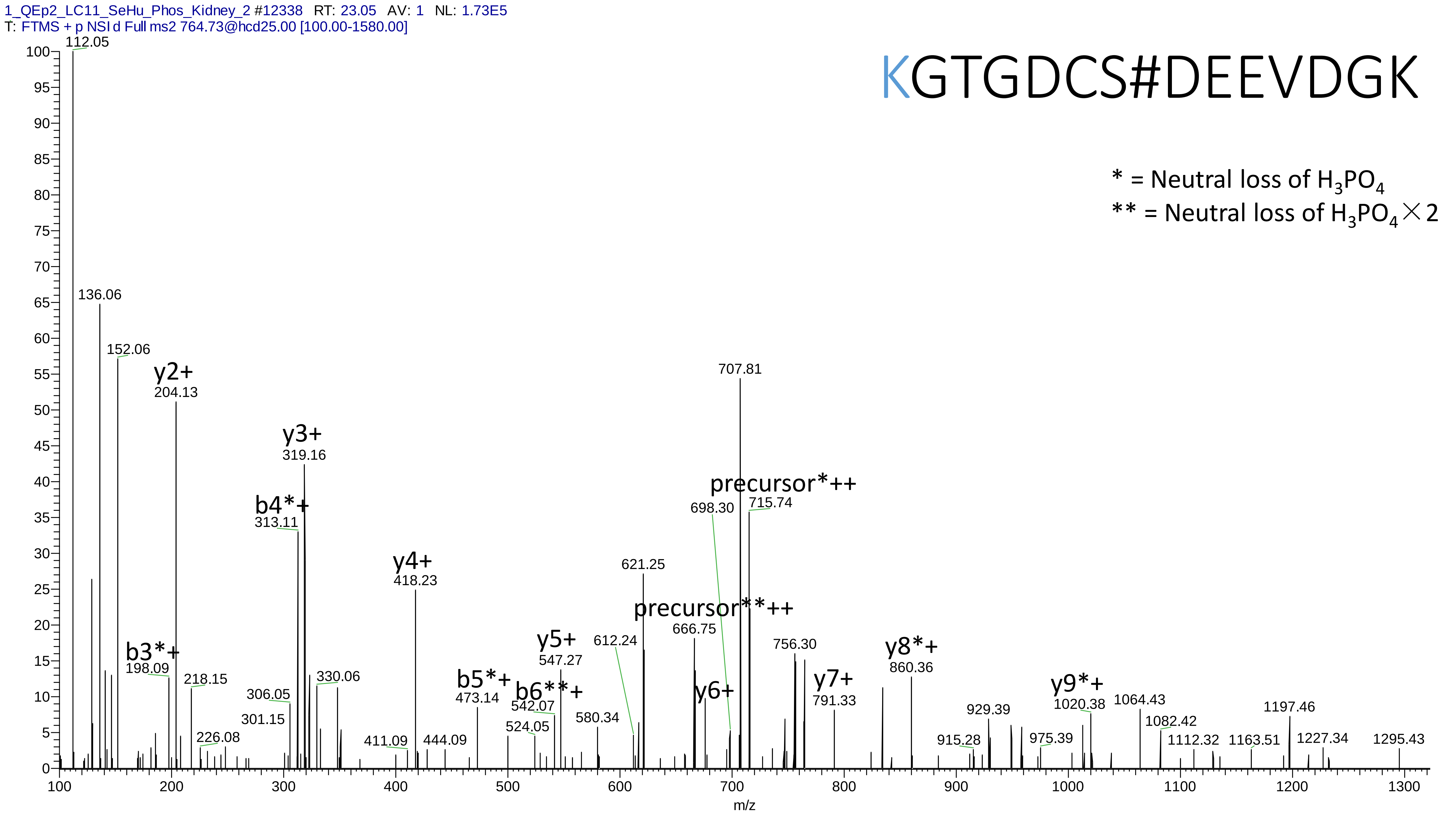
1_QEp2_LC11_SeHu_Phos_Kidney_2 #12338 RT: 23.05 AV: 1 NL: 1.73E5
T: FTMS + p NSI d Full ms2 764.73@hcd25.00 [100.00-1580.00]
KGTGDCS#DEEVDGK
* = Neutral loss of H3PO4
** = Neutral loss of H3PO4×2
y2+
y3+
b4*+
y4+
b3*+
b5*+
b6**+
y5+
precursor*++
precursor**++
y6+
y7+
y8*+
y9*+
m/z

# 16

- Experimental mass: 71.9849
- Unimod information:
- Possible composition: Formyl + Carboxy, [C O + C O(2)]
- Sites in Unimod:
- Sites observed:
- Features: 24.4% (10/41) peptides contain Lys at peptide n-term;

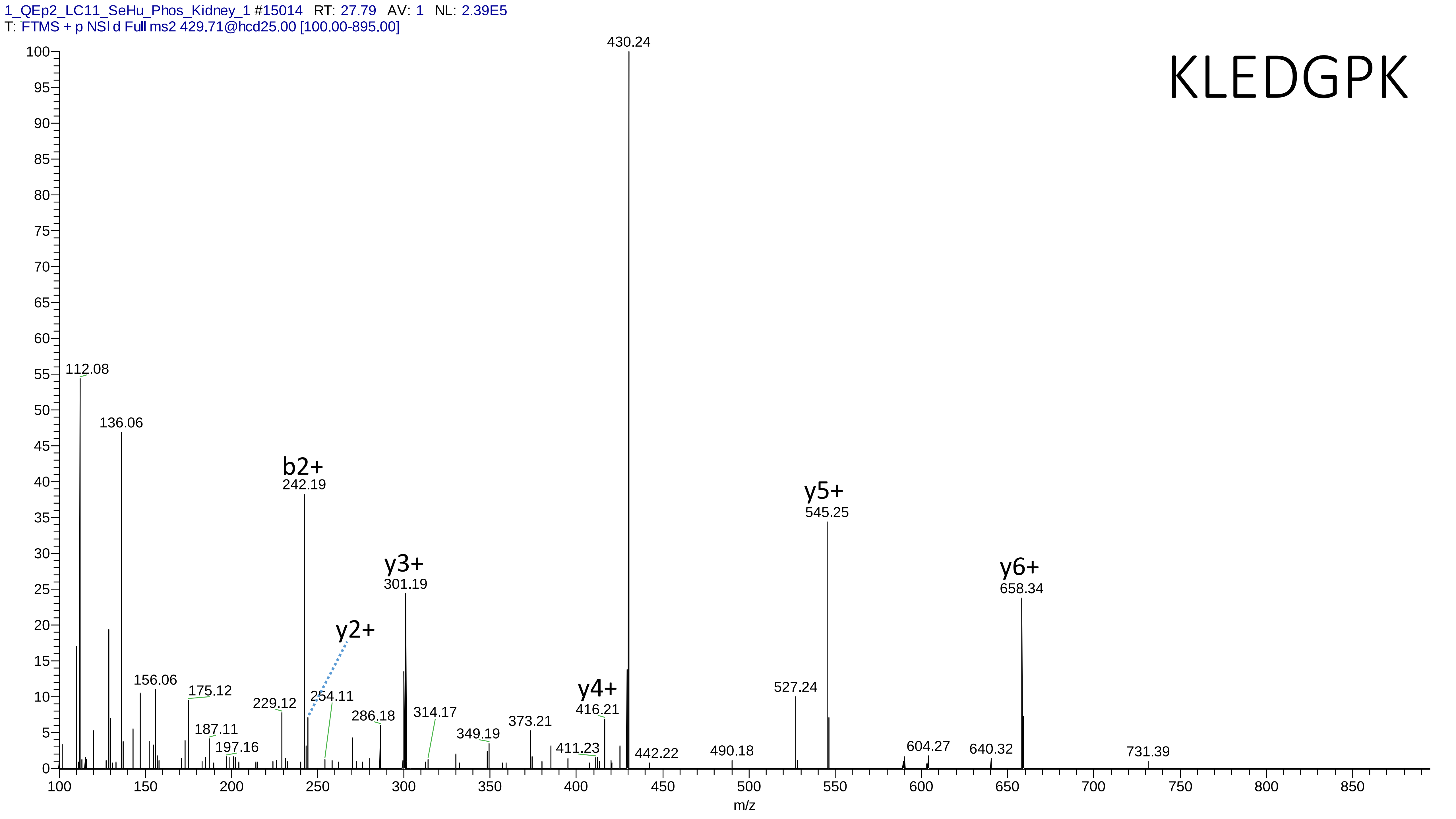

1_QEp2_LC11_SeHu_Phos_Kidney_1 #15014 RT: 27.79 AV: 1 NL: 2.39E5
T: FTMS + p NSI d Full ms2 429.71@hcd25.00 [100.00-895.00]
KLEDGPK
430.24
112.08
136.06
b2+
242.19
y5+
545.25
y3+
301.19
y6+
658.34
y2+
156.06
175.12
229.12
254.11
y4+
416.21
527.24
286.18
314.17
187.11
349.19
373.21
197.16
411.23
442.22
490.18
604.27
640.32
731.39
m/z

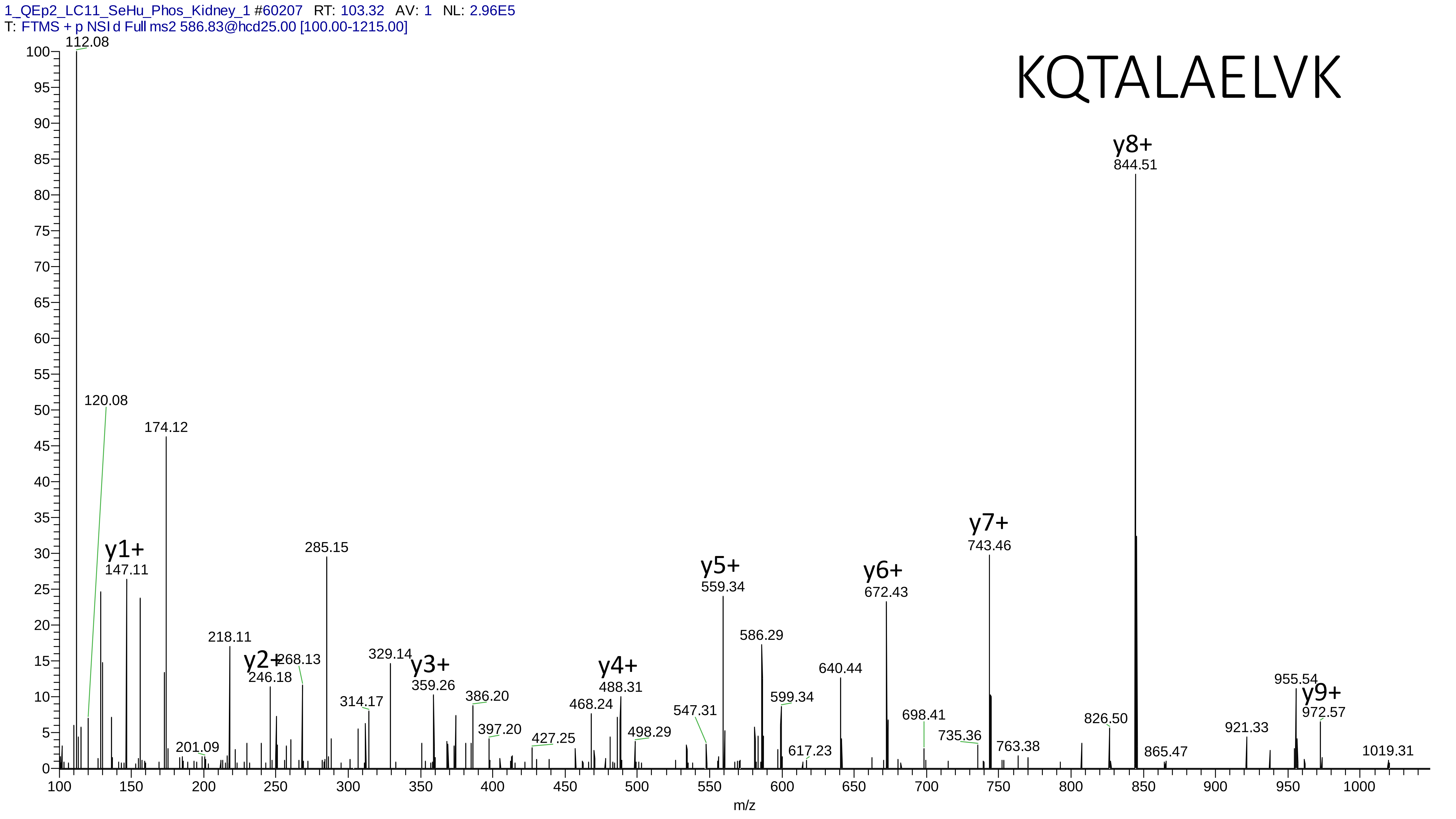
1_QEp2_LC11_SeHu_Phos_Kidney_1 #60207 RT: 103.32 AV: 1 NL: 2.96E5
T: FTMS + p NSI d Full ms2 586.83@hcd25.00 [100.00-1215.00]
KQTALAELVK
y8+
844.51
112.08
120.08
174.12
y1+
147.11
285.15
y7+
743.46
y5+
559.34
y6+
672.43
218.11
y2+
246.18
268.13
329.14
y3+
359.26
y4+
488.31
586.29
640.44
955.54
y9+
972.57
314.17
386.20
468.24
547.31
599.34
698.41
826.50
921.33
397.20
427.25
498.29
735.36
763.38
201.09
617.23
865.47
1019.31
m/z

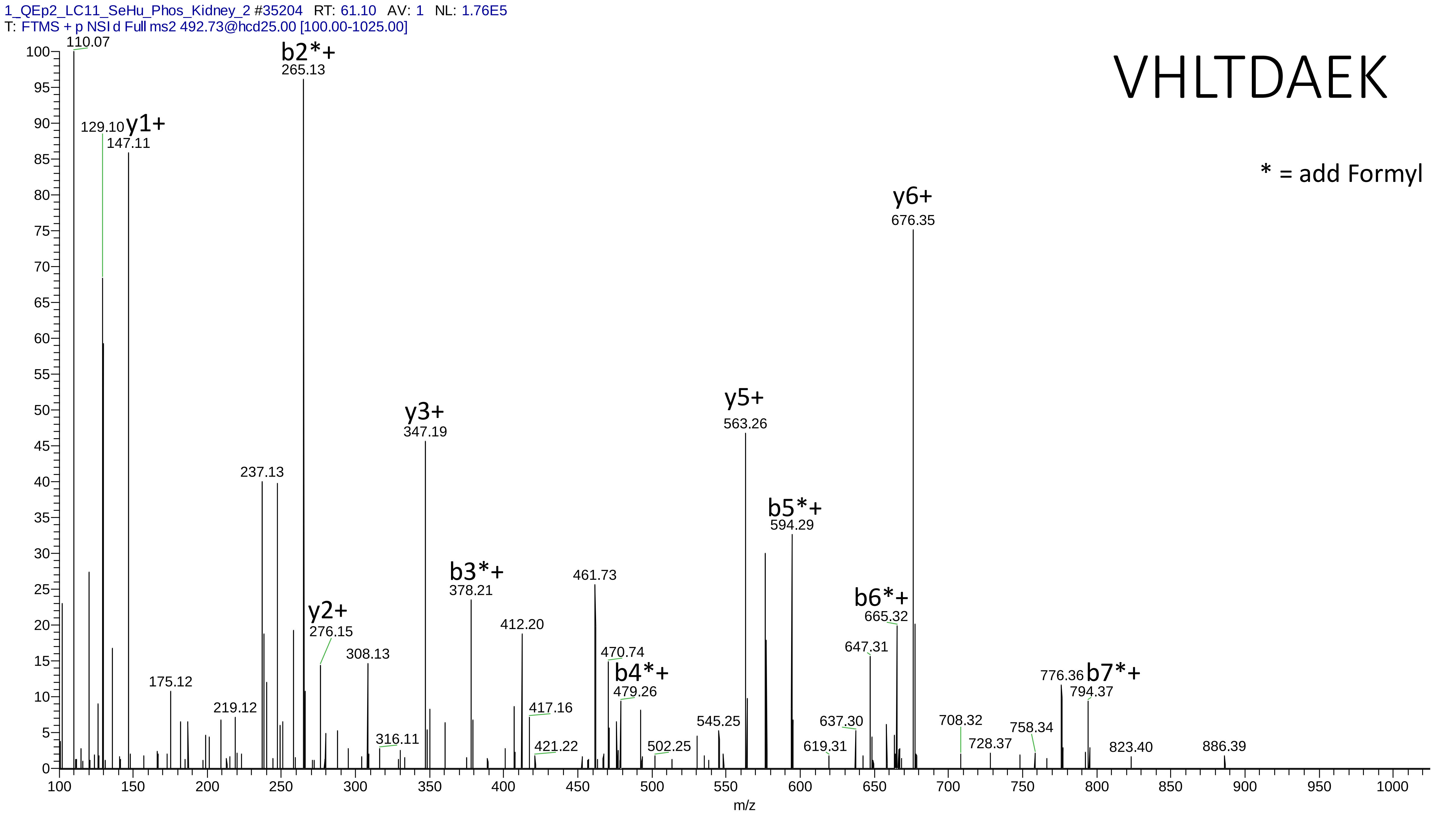

1_QEp2_LC11_SeHu_Phos_Kidney_2 #35204 RT: 61.10 AV: 1 NL: 1.76E5
T: FTMS + p NSI d Full ms2 492.73@hcd25.00 [100.00-1025.00]
VHLTDAEK
* = add Formyl
110.07
b2*+
265.13
129.10
y1+
147.11
y6+
676.35
y5+
563.26
y3+
347.19
237.13
b5*+
594.29
b3*+
378.21
461.73
b6*+
665.32
y2+
276.15
412.20
647.31
308.13
470.74
b4*+
479.26
175.12
776.36
b7*+
794.37
219.12
417.16
545.25
637.30
708.32
316.11
758.34
421.22
502.25
619.31
728.37
823.40
886.39
m/z

# 17

- Experimental mass: 197.8788
- Unimod information:
- Possible composition: Phosphorylation * 2 + Cation:K, [H(1)O(3)P(1) * 2 + H(-2) K ]
- Sites in Unimod:
- Sites observed:
- Features: 100% (70/70) peptides contain equal or more than two Ser/Thr

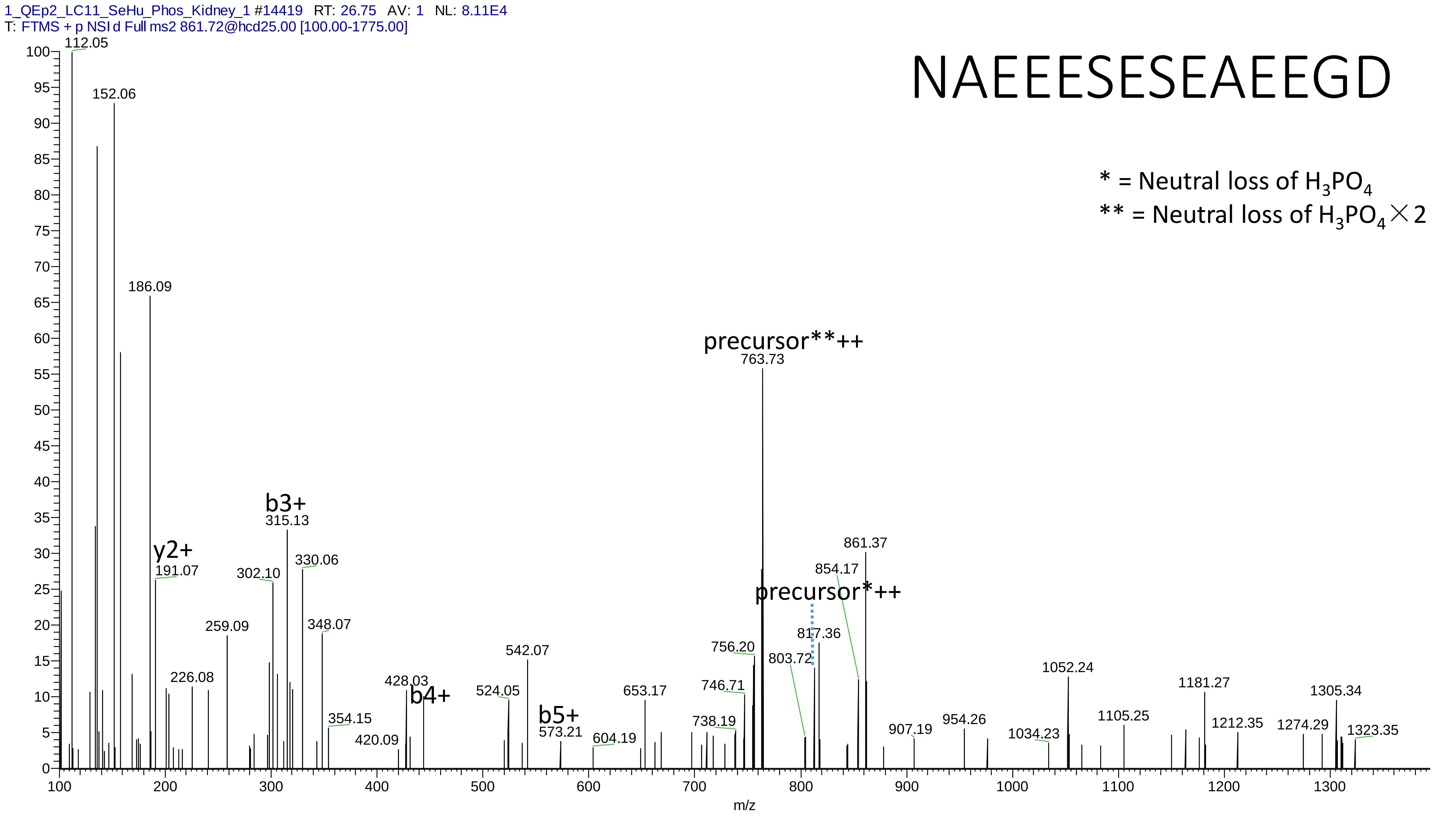

1_QEp2_LC11_SeHu_Phos_Kidney_1 #14419 RT: 26.75 AV: 1 NL: 8.11E4
T: FTMS + p NSI d Full ms2 861.72@hcd25.00 [100.00-1775.00]
NAEEESESEAEEGD
* = Neutral loss of H3PO4
** = Neutral loss of H3PO4×2
precursor**++
763.73
precursor*++
y2+
191.07
b3+
315.13
b4+
b5+
573.21
112.05
152.06
186.09
226.08
259.09
302.10
330.06
348.07
354.15
420.09
428.03
524.05
542.07
604.19
653.17
738.19
746.71
756.20
803.72
817.36
854.17
861.37
907.19
954.26
1034.23
1052.24
1105.25
1181.27
1212.35
1274.29
1305.34
1323.35
m/z

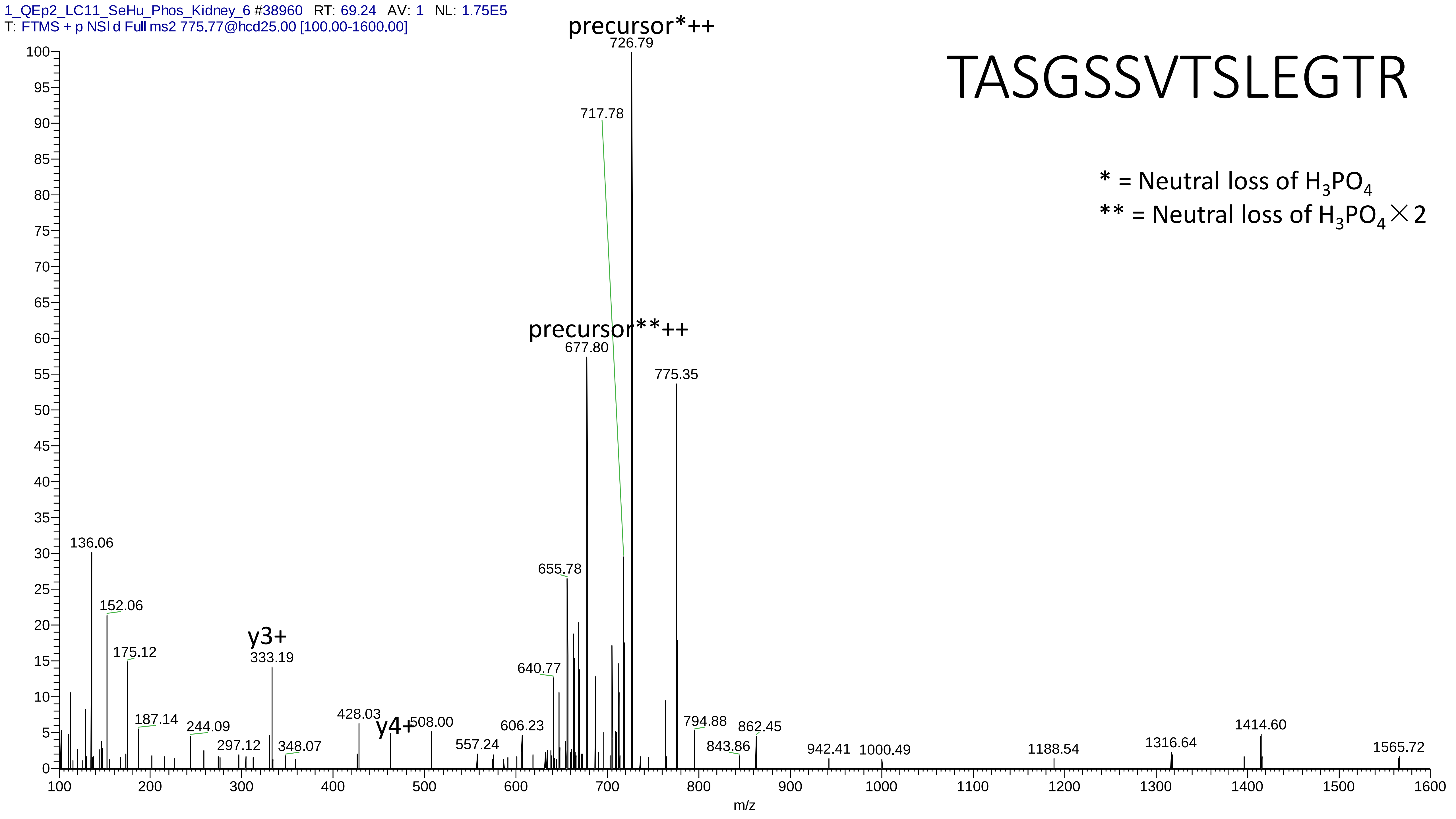

1_QEp2_LC11_SeHu_Phos_Kidney_6 #38960 RT: 69.24 AV: 1 NL: 1.75E5
T: FTMS + p NSI d Full ms2 775.77@hcd25.00 [100.00-1600.00]
TASGSSVTSLEGTR
* = Neutral loss of $H_3PO_4$
** = Neutral loss of $H_3PO_4 \times 2$
precursor*++
726.79
717.78
precursor**++
677.80
775.35
655.78
640.77
136.06
152.06
175.12
y3+
333.19
187.14
244.09
297.12
348.07
428.03
y4+
508.00
557.24
606.23
794.88
862.45
843.86
942.41
1000.49
1188.54
1316.64
1414.60
1565.72
m/z

# 18

- Experimental mass: 96.9928
- Unimod information:
- Possible composition: Phosphorylation + Ammonium, [H(1)O(3)P(1) + H(3) N ]
- Sites in Unimod:
- Sites observed:

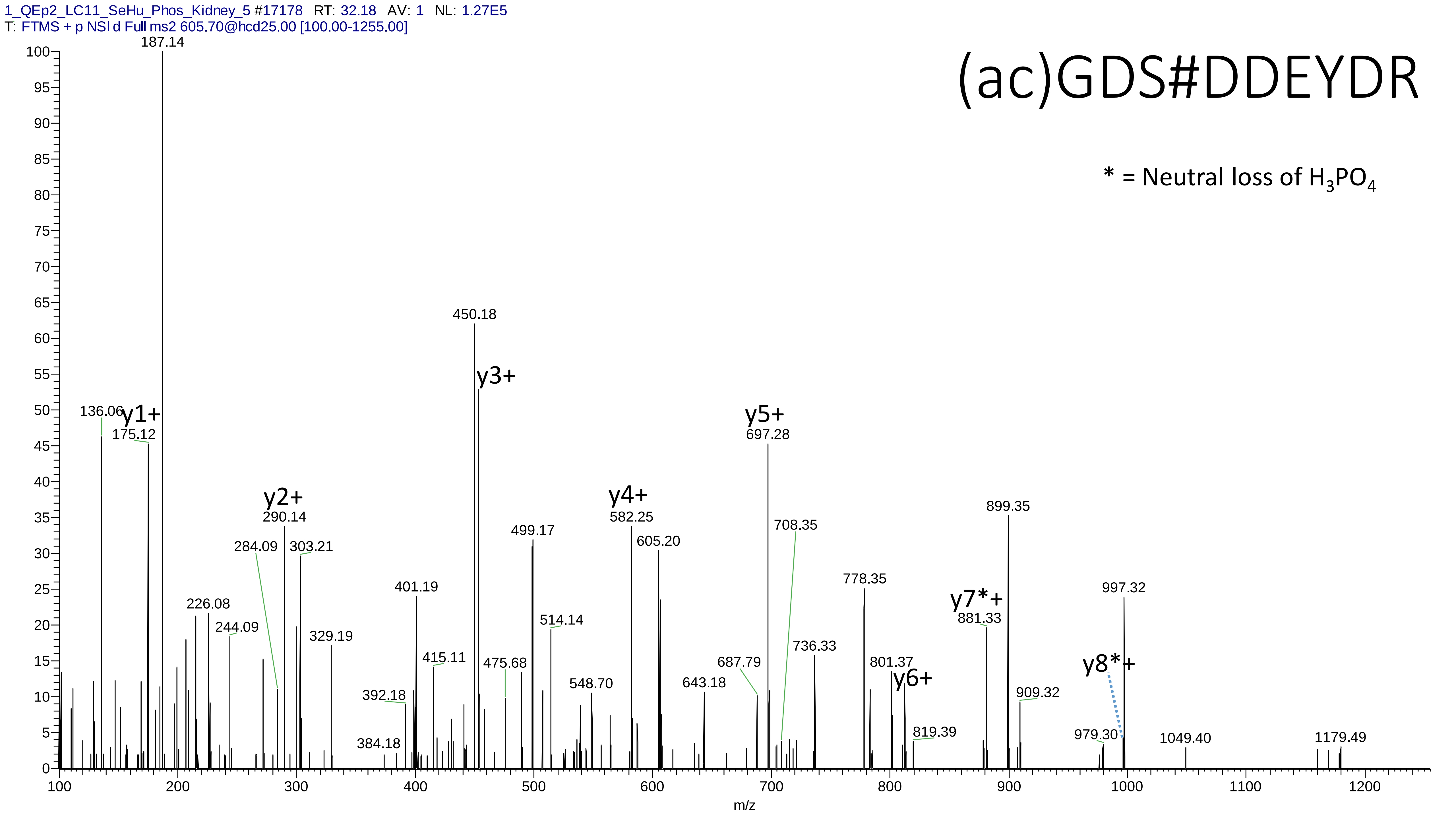
1_QEp2_LC11_SeHu_Phos_Kidney_5 #17178 RT: 32.18 AV: 1 NL: 1.27E5
T: FTMS + p NSI d Full ms2 605.70@hcd25.00 [100.00-1255.00]
(ac)GDS#DDEYDR
* = Neutral loss of H3PO4
187.14
450.18
y3+
136.06
y1+
175.12
y5+
697.28
y2+
290.14
y4+
582.25
899.35
708.35
605.20
499.17
284.09
303.21
401.19
778.35
997.32
y7*+
881.33
226.08
244.09
329.19
514.14
415.11
736.33
475.68
687.79
801.37
y8*+
y6+
392.18
548.70
643.18
909.32
819.39
384.18
979.30
1049.40
1179.49
m/z

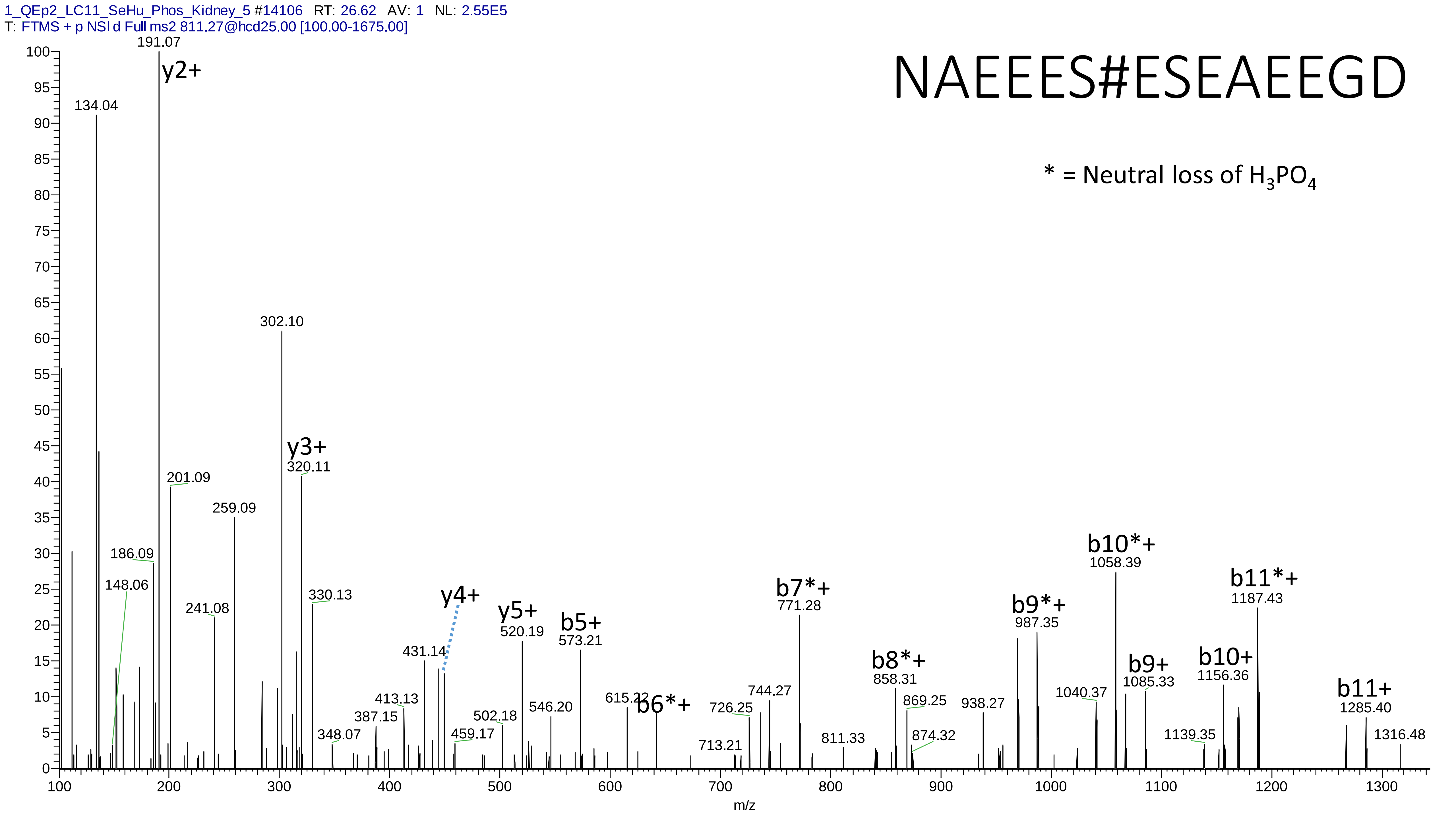
1_QEp2_LC11_SeHu_Phos_Kidney_5 #14106 RT: 26.62 AV: 1 NL: 2.55E5
T: FTMS + p NSI d Full ms2 811.27@hcd25.00 [100.00-1675.00]
NAEEES#ESEAEEGD
* = Neutral loss of H3PO4
y2+
191.07
134.04
302.10
y3+
320.11
201.09
259.09
186.09
148.06
241.08
330.13
y4+
y5+
520.19
b5+
573.21
b6*+
b7*+
771.28
b8*+
858.31
b9*+
987.35
b10*+
1058.39
b9+
1085.33
b10+
1156.36
b11*+
1187.43
b11+
1285.40
1316.48
431.14
413.13
387.15
348.07
459.17
502.18
546.20
615.22
726.25
744.27
713.21
811.33
869.25
874.32
938.27
1040.37
1139.35
m/z

# 19

- Experimental mass: 97.9898
- Unimod information:
- Possible composition:
- Sites in Unimod:
- Sites observed:

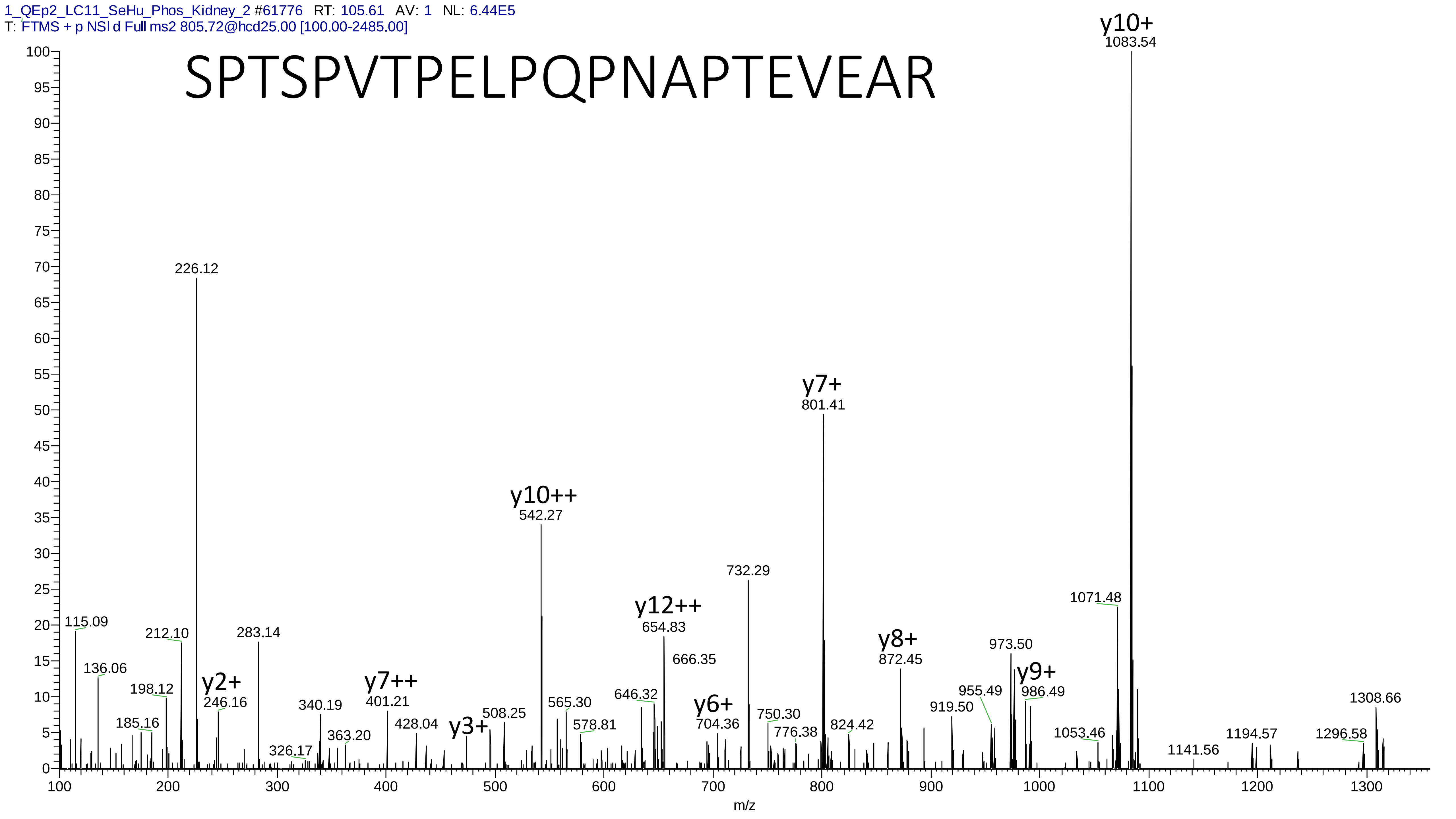
1_QEp2_LC11_SeHu_Phos_Kidney_2 #61776 RT: 105.61 AV: 1 NL: 6.44E5
T: FTMS + p NSI d Full ms2 805.72@hcd25.00 [100.00-2485.00]
SPTSPVTPELPQPNAPTEVEAR
y10+
1083.54
226.12
y7+
801.41
y10++
542.27
732.29
1071.48
y12++
654.83
666.35
115.09
212.10
283.14
136.06
198.12
185.16
y2+
246.16
340.19
363.20
326.17
y7++
401.21
428.04
y3+
508.25
565.30
578.81
646.32
y6+
704.36
750.30
776.38
824.42
y8+
872.45
919.50
955.49
973.50
y9+
986.49
1053.46
1141.56
1194.57
1296.58
1308.66
m/z

# 20

- Experimental mass: 199.9627
- Unimod information:
- Possible composition: Phosphorylation * 2 + Gly->Pro, [H(1)O(3)P(1) * 2 + H(4) C(3)]
- Sites in Unimod:
- Sites observed:
- Features: 100% (41/41) peptides contain equal or more than two Ser/Thr; 92.7% (38/41) peptides contain Gly

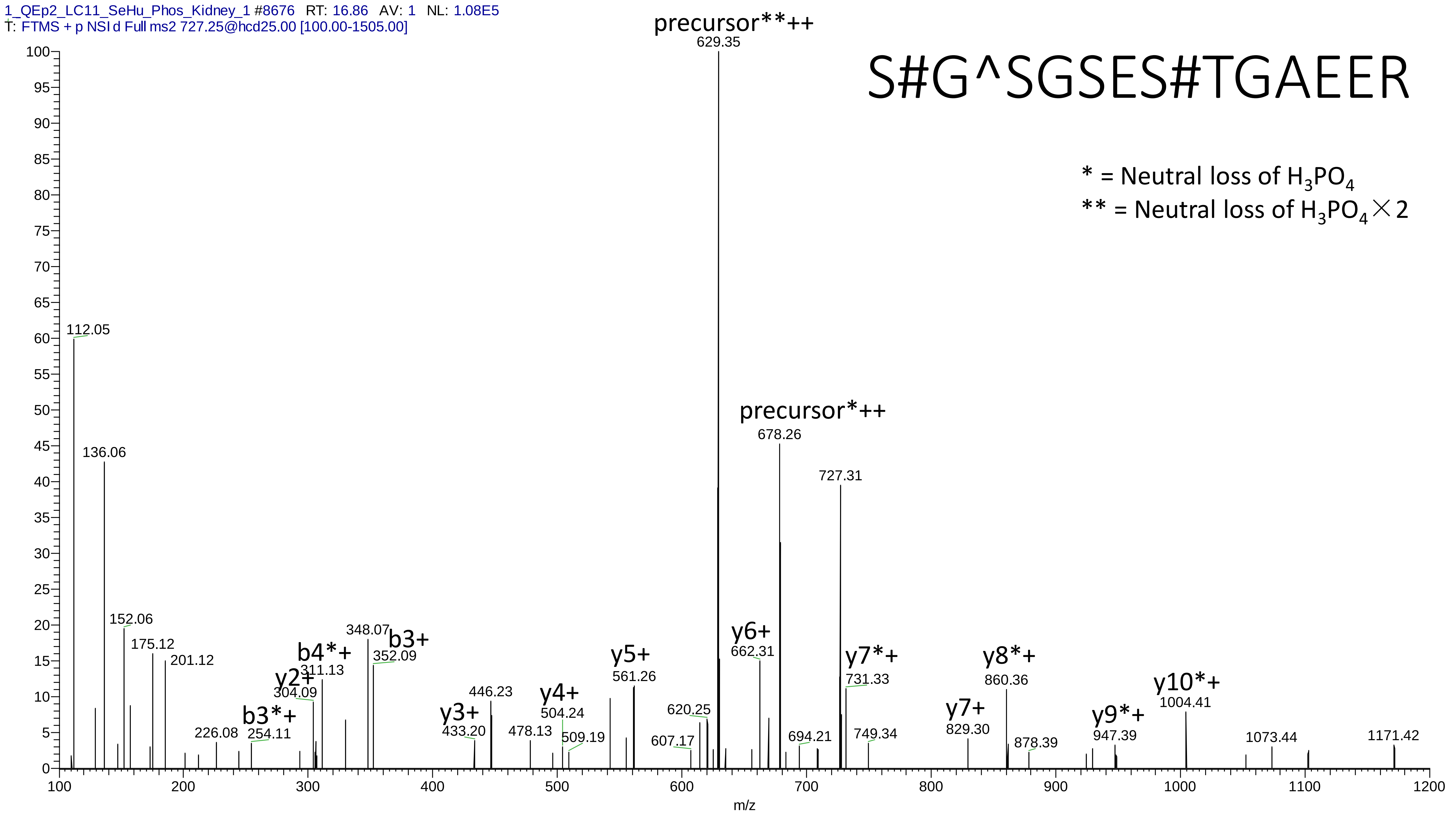
1_QEp2_LC11_SeHu_Phos_Kidney_1 #8676 RT: 16.86 AV: 1 NL: 1.08E5
T: FTMS + p NSI d Full ms2 727.25@hcd25.00 [100.00-1505.00]
S#G^SGSES#TGAEER
* = Neutral loss of H3PO4
** = Neutral loss of H3PO4×2
precursor**++
629.35
precursor*++
678.26
727.31
112.05
136.06
152.06
175.12
201.12
226.08
b3*+
254.11
y2+
304.09
b4*+
311.13
348.07
b3+
352.09
y3+
433.20
446.23
478.13
y4+
504.24
509.19
y5+
561.26
607.17
620.25
y6+
662.31
694.21
y7*+
731.33
749.34
y7+
829.30
y8*+
860.36
878.39
y9*+
947.39
y10*+
1004.41
1073.44
1171.42
m/z

# 21

- Experimental mass: 403.0656
- Unimod information:
- Possible composition: Phosphorylation * 2 + Ser-Arg, [H(1)O(3)P(1) * 2 + H(5)C(3)N(1)O(2) + H(12)C(6)N(4)O(1)]
- Sites in Unimod:
- Sites observed:
- Features: 100% (11/11) peptides contain equal or more than two Ser/Thr; for 100.0% (11/11) peptides with this modification, two amino acids before the peptides are Ser-Arg

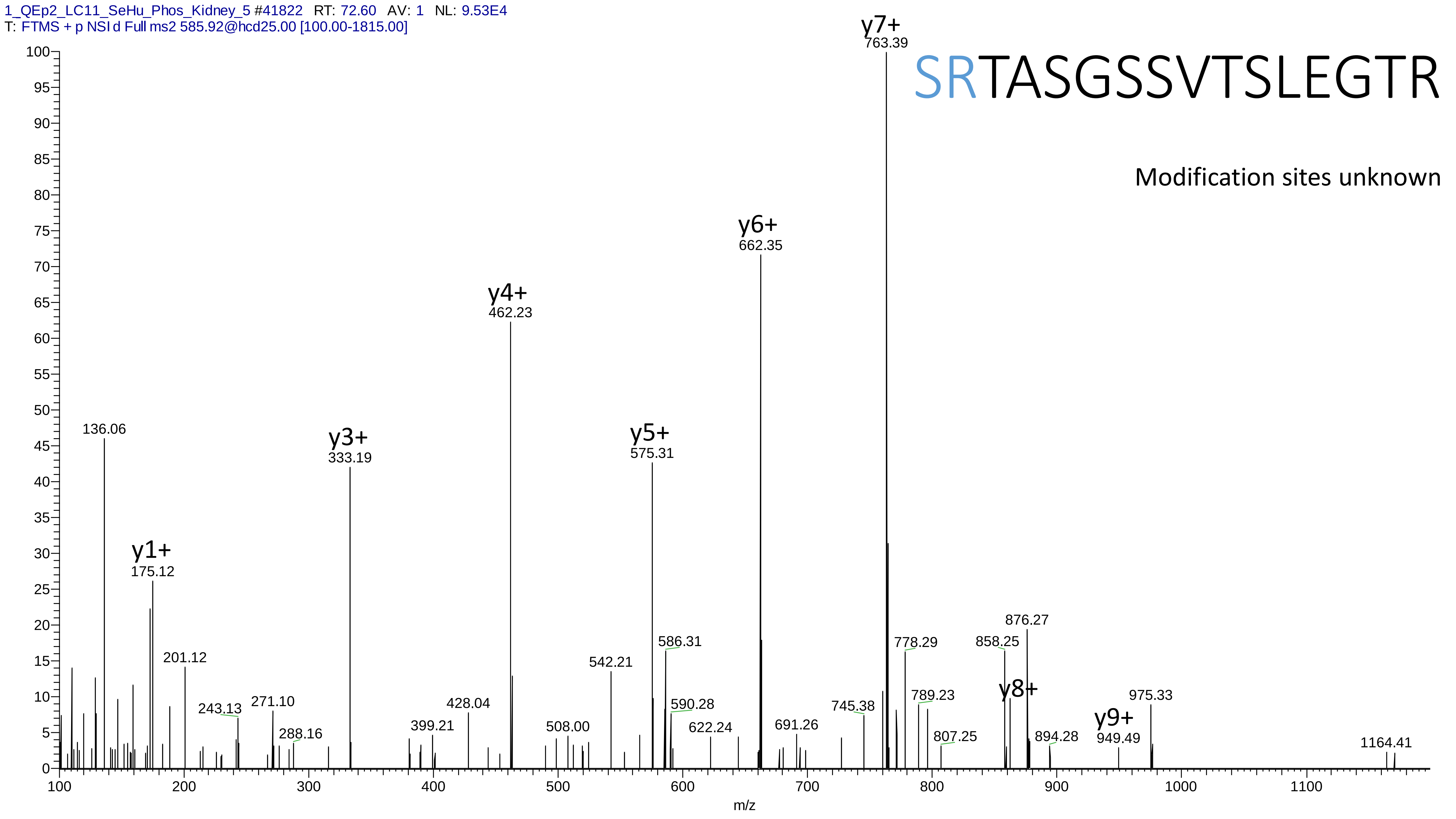
1_QEp2_LC11_SeHu_Phos_Kidney_5 #41822 RT: 72.60 AV: 1 NL: 9.53E4
T: FTMS + p NSI d Full ms2 585.92@hcd25.00 [100.00-1815.00]
SRTASGSSVTSLEGTR
Modification sites unknown
y1+ 175.12
y3+ 333.19
y4+ 462.23
y5+ 575.31
y6+ 662.35
y7+ 763.39
y8+
y9+ 949.49
136.06
201.12
243.13
271.10
288.16
399.21
428.04
508.00
542.21
586.31
590.28
622.24
691.26
745.38
778.29
789.23
807.25
858.25
876.27
894.28
975.33
1164.41
m/z

# 22

- Experimental mass: 97.978
- Unimod information:
- Possible composition:
- Sites in Unimod:
- Sites observed:
- Features:

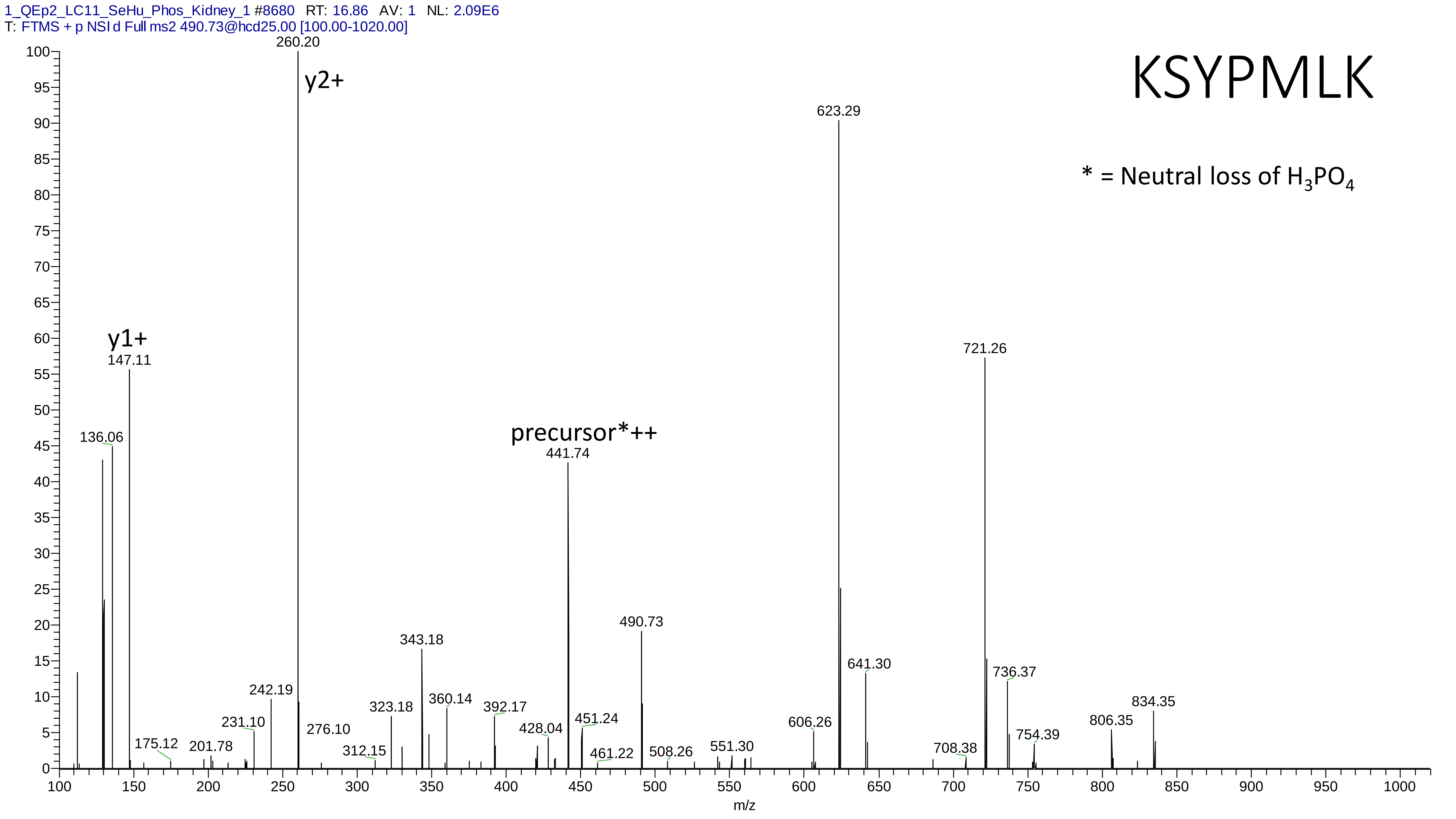
1_QEp2_LC11_SeHu_Phos_Kidney_1 #8680 RT: 16.86 AV: 1 NL: 2.09E6
T: FTMS + p NSI d Full ms2 490.73@hcd25.00 [100.00-1020.00]
KSYPMLK
* = Neutral loss of $H_3PO_4$
y2+
260.20
623.29
721.26
y1+
147.11
136.06
precursor*++
441.74
490.73
343.18
641.30
736.37
834.35
242.19
323.18
360.14
392.17
806.35
606.26
451.24
428.04
231.10
276.10
754.39
175.12
201.78
551.30
312.15
708.38
461.22
508.26
m/z

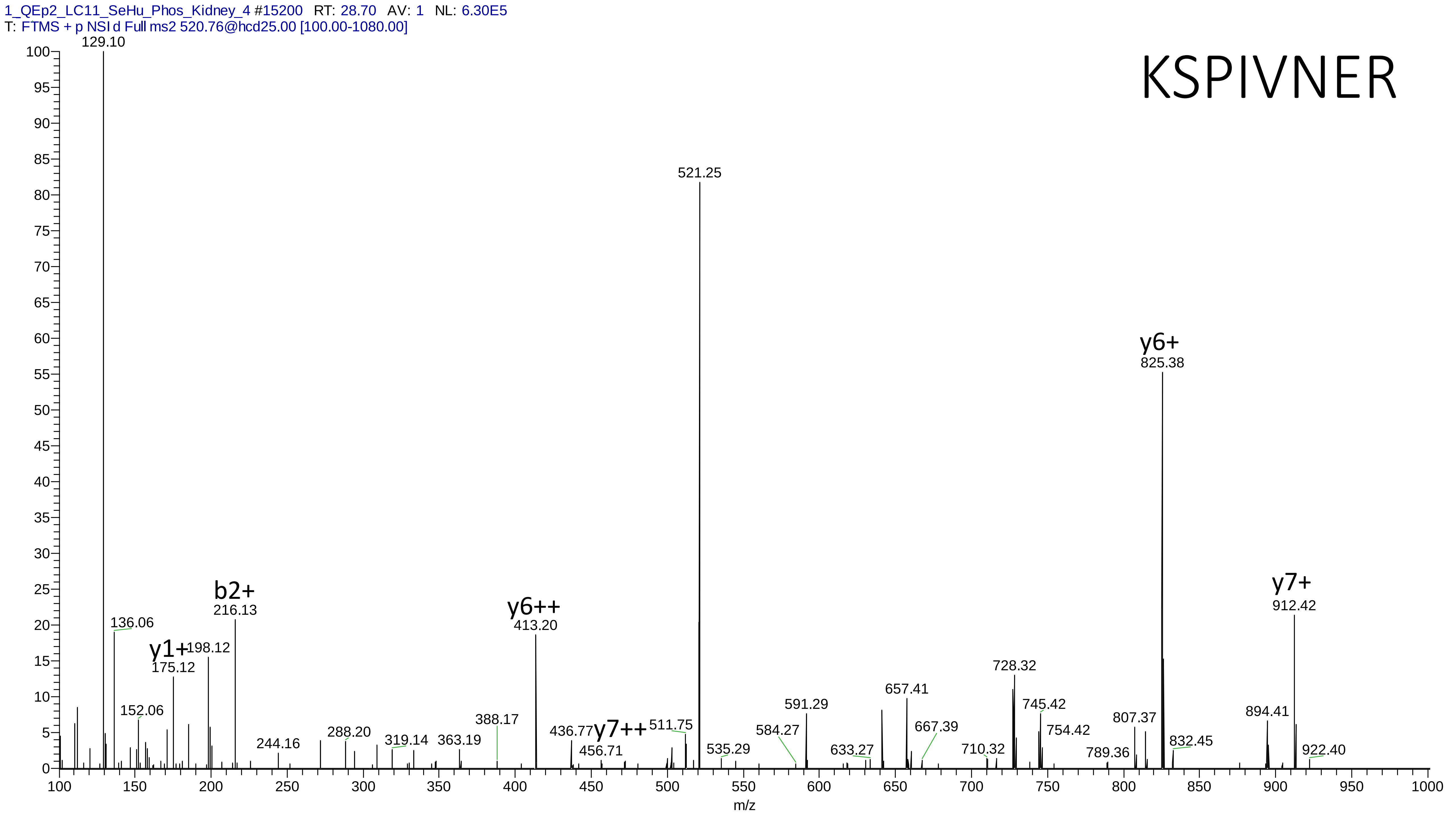
1_QEp2_LC11_SeHu_Phos_Kidney_4 #15200 RT: 28.70 AV: 1 NL: 6.30E5
T: FTMS + p NSI d Full ms2 520.76@hcd25.00 [100.00-1080.00]
KSPIVNER
129.10
521.25
y6+
825.38
y7+
912.42
b2+
216.13
y6++
413.20
136.06
y1+
175.12
198.12
152.06
244.16
288.20
319.14
363.19
388.17
436.77
y7++
456.71
511.75
535.29
584.27
591.29
633.27
657.41
667.39
710.32
728.32
745.42
754.42
789.36
807.37
832.45
894.41
922.40
m/z

# 23

- Experimental mass: -177.1697
- Unimod information:
- Possible composition: Phosphorylation + Glu-Lys-loss, [H(1)O(3)P(1) + H(-12) C(-6) N(-4) O(-1)]
- Sites in Unimod:
- Sites observed:
- Features: 97.5% (117/120) peptide spectra matches contain EK at peptide n-term

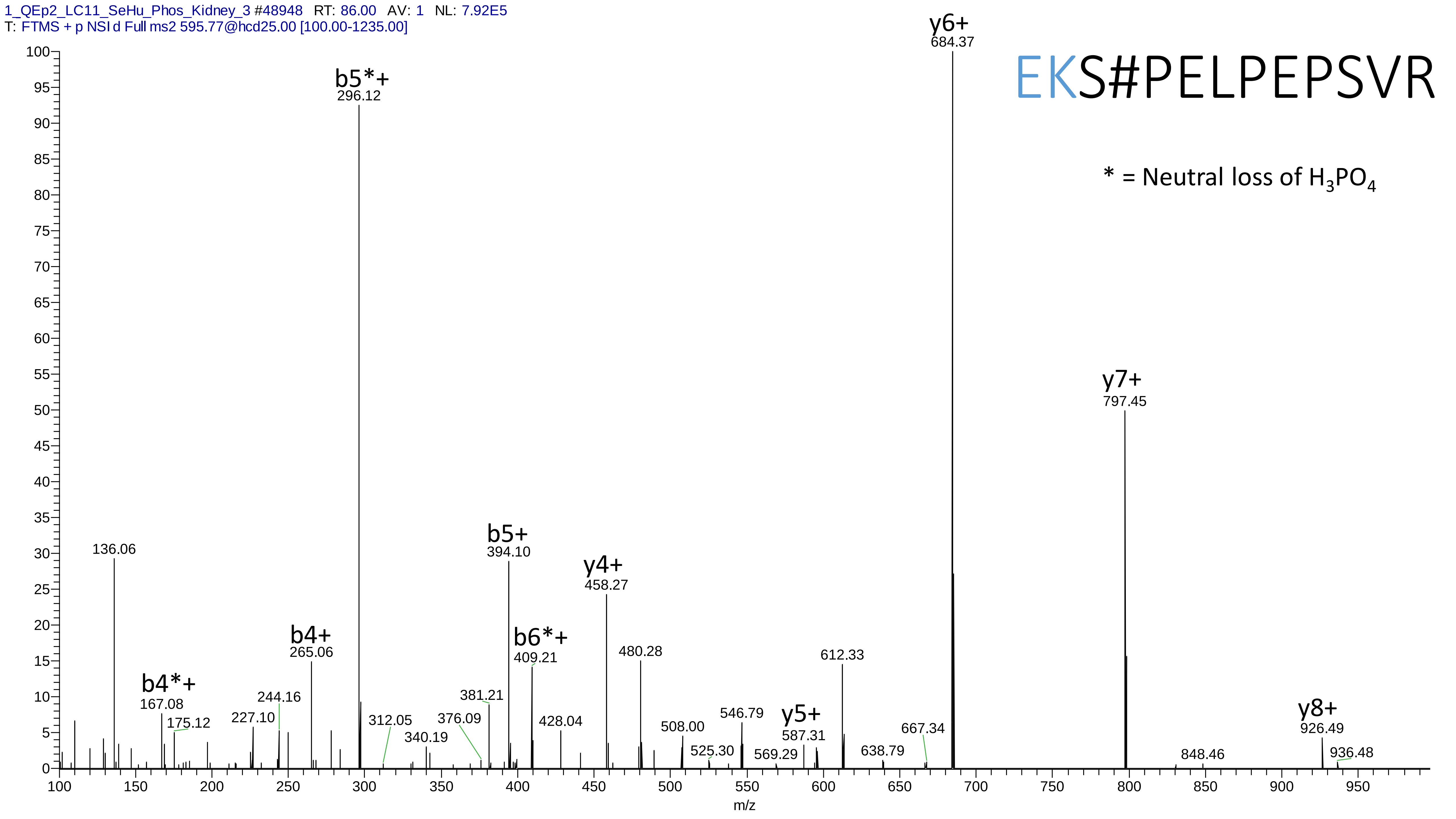

1_QEp2_LC11_SeHu_Phos_Kidney_3 #48948 RT: 86.00 AV: 1 NL: 7.92E5
T: FTMS + p NSI d Full ms2 595.77@hcd25.00 [100.00-1235.00]
EKS#PELPEPSVR
* = Neutral loss of $H_3PO_4$
y6+ 684.37
b5*+ 296.12
y7+ 797.45
b5+ 394.10
y4+ 458.27
136.06
b4+ 265.06
b6*+ 409.21
480.28
612.33
b4*+ 167.08
244.16
381.21
175.12
227.10
312.05
376.09
428.04
546.79
y5+ 587.31
y8+ 926.49
340.19
508.00
667.34
525.30
569.29
638.79
848.46
936.48
m/z

# 24

- Experimental mass: -76.1348
- Unimod information:
- Possible composition: Phosphorylation + Arg-loss, [H(1)O(3)P(1) + H(-12) C(-6) N(-4) O(-1)]
- Sites in Unimod:
- Sites observed:
- Features:  91.2% (31/34) peptides contain Arg at peptide n-term

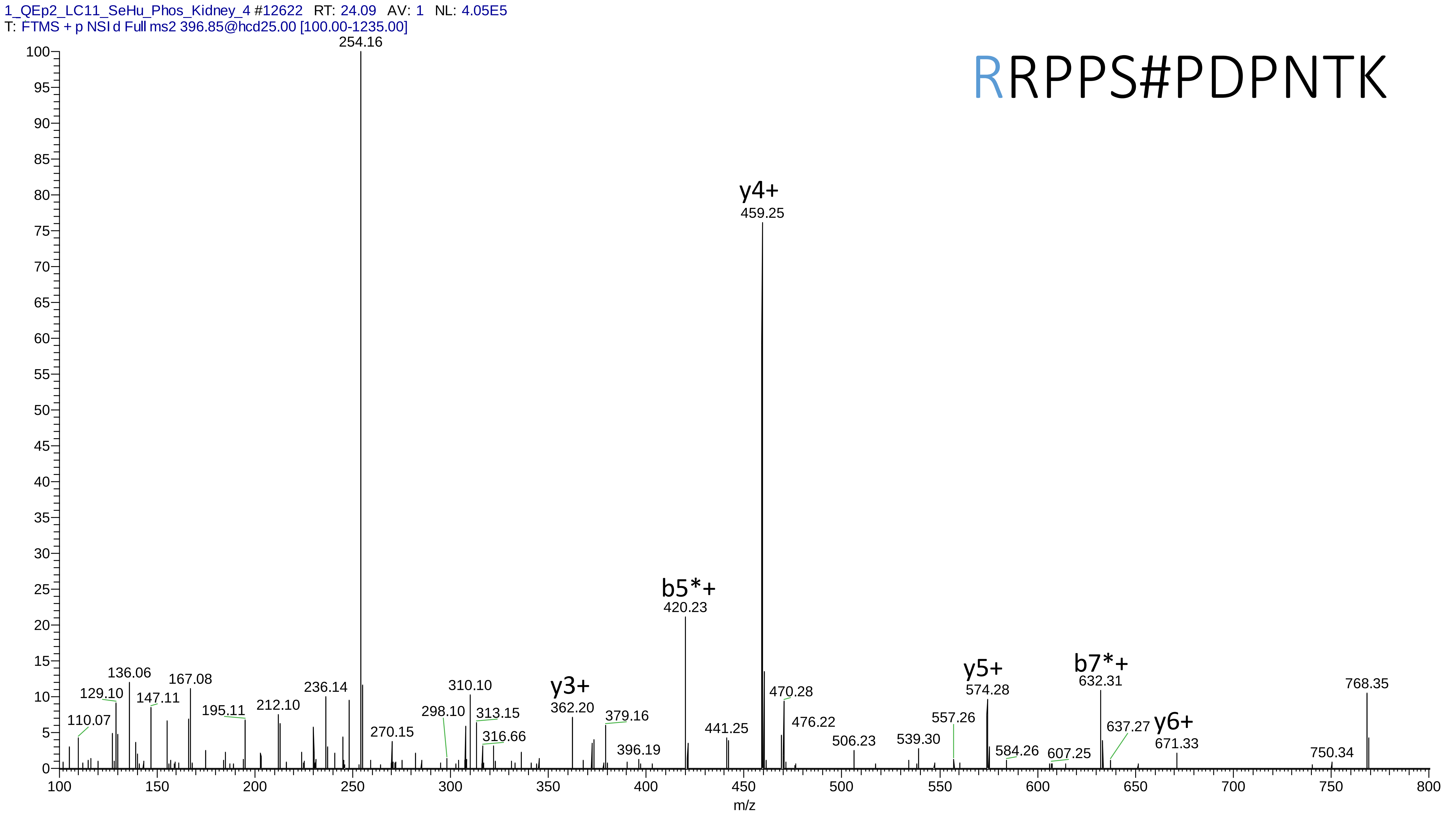
1_QEp2_LC11_SeHu_Phos_Kidney_4 #12622 RT: 24.09 AV: 1 NL: 4.05E5
T: FTMS + p NSI d Full ms2 396.85@hcd25.00 [100.00-1235.00]
RRPPS#PDPNTK
y4+
b5*+
y3+
y5+
b7*+
y6+
254.16
459.25
420.23
136.06
167.08
129.10
147.11
110.07
195.11
212.10
236.14
310.10
298.10
313.15
270.15
316.66
362.20
379.16
396.19
441.25
470.28
476.22
506.23
539.30
557.26
574.28
584.26
607.25
632.31
637.27
671.33
750.34
768.35
m/z

# 25

- Experimental mass: -91.0769
- Unimod information:
- Possible composition:
- Sites in Unimod:
- Sites observed:
- Features: 100% (11/11) peptides contain Tyr at peptide n-term

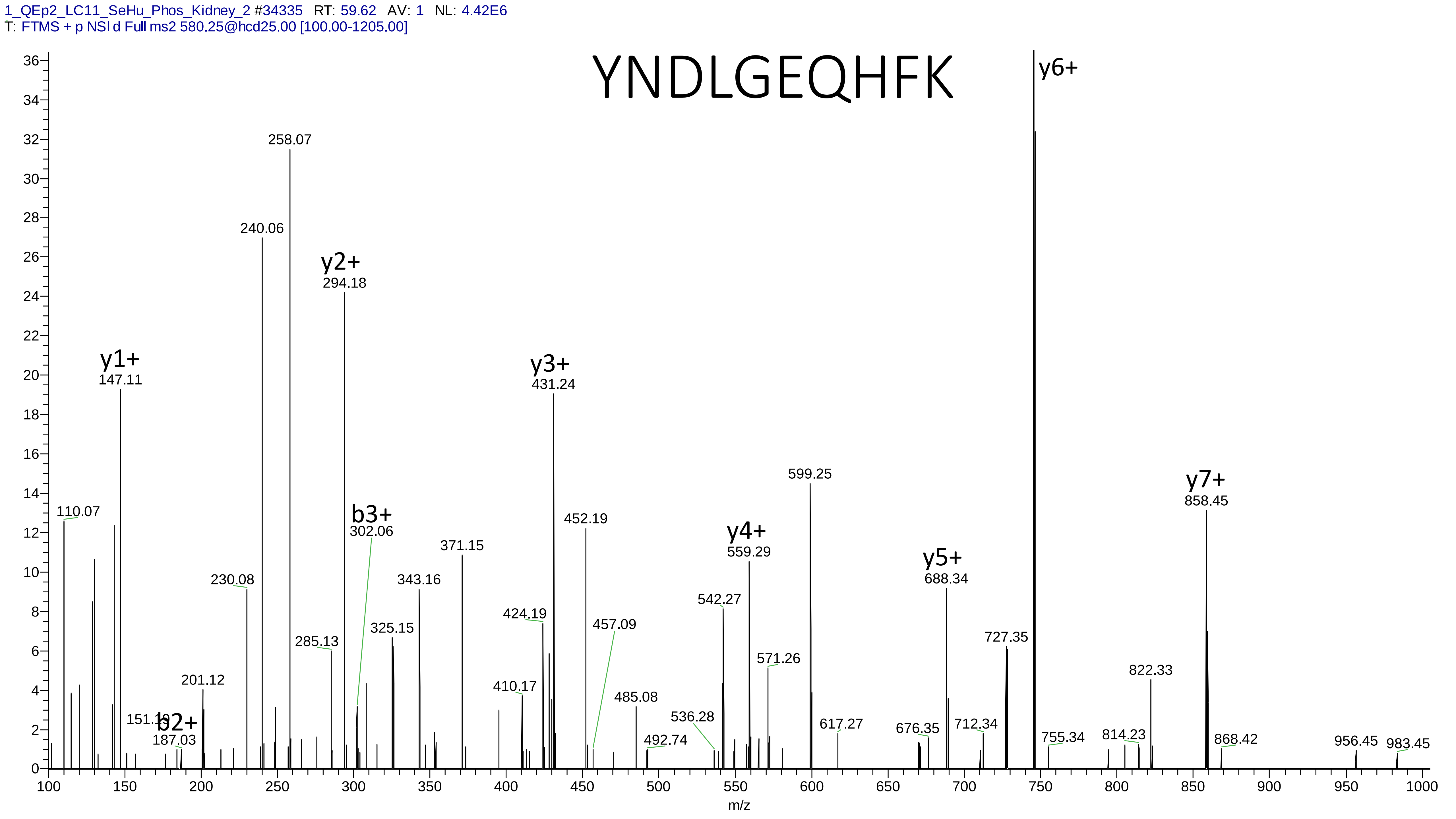
1_QEp2_LC11_SeHu_Phos_Kidney_2 #34335 RT: 59.62 AV: 1 NL: 4.42E6
T: FTMS + p NSI d Full ms2 580.25@hcd25.00 [100.00-1205.00]
YNDLGEQHFK
y6+
258.07
240.06
y2+
294.18
y1+
147.11
y3+
431.24
599.25
y7+
858.45
110.07
b3+
302.06
452.19
371.15
y4+
559.29
230.08
343.16
y5+
688.34
424.19
542.27
457.09
325.15
285.13
727.35
571.26
822.33
201.12
410.17
485.08
536.28
151.10
b2+
187.03
617.27
676.35
712.34
755.34
814.23
868.42
956.45
983.45
492.74
m/z

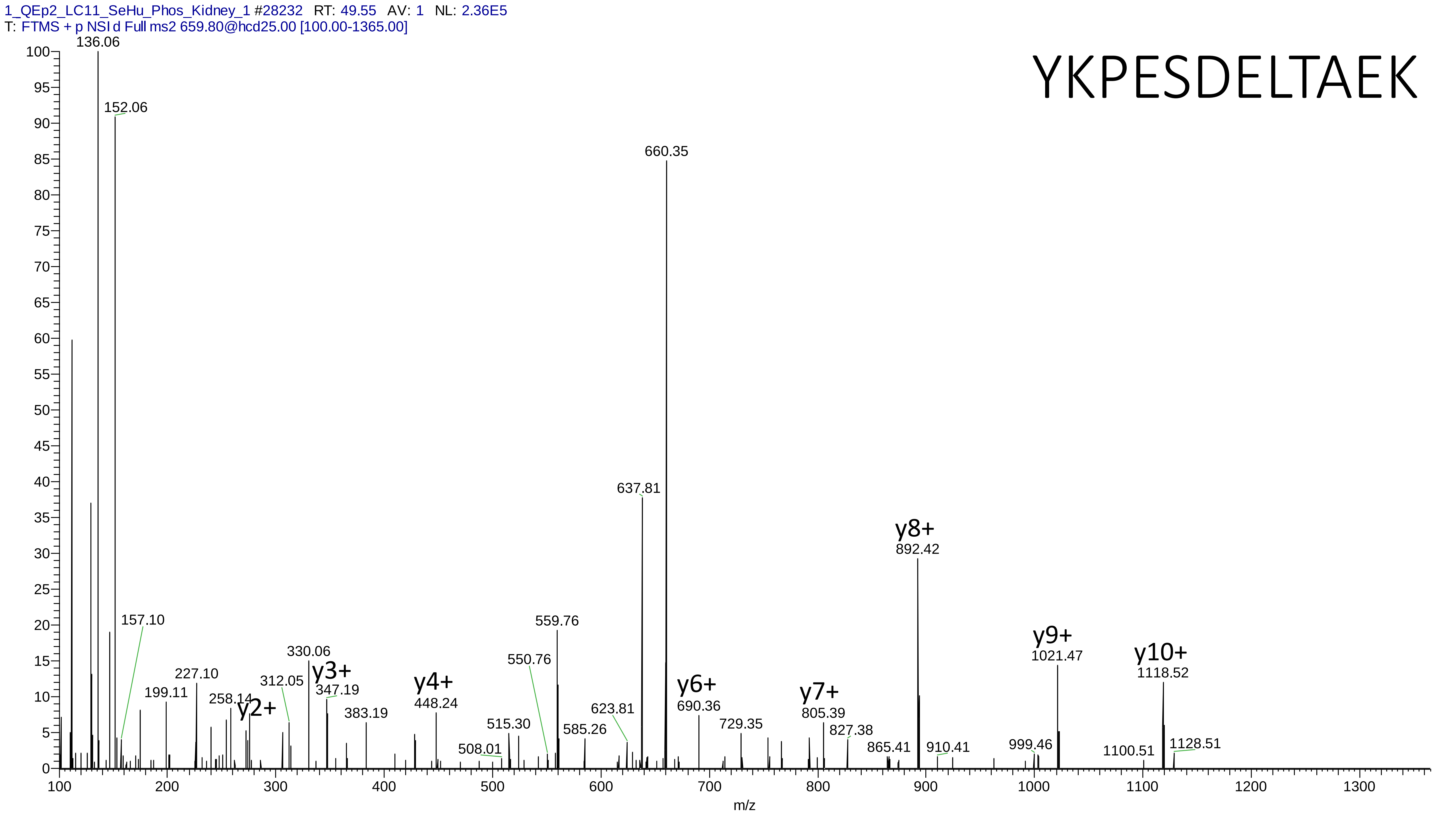
1_QEp2_LC11_SeHu_Phos_Kidney_1 #28232 RT: 49.55 AV: 1 NL: 2.36E5
T: FTMS + p NSI d Full ms2 659.80@hcd25.00 [100.00-1365.00]
YKPESDELTAEK
136.06
152.06
660.35
637.81
y8+
892.42
157.10
559.76
330.06
y3+
347.19
227.10
199.11
258.14
y2+
312.05
550.76
y4+
448.24
y6+
690.36
y7+
805.39
y9+
1021.47
y10+
1118.52
383.19
515.30
508.01
585.26
623.81
729.35
827.38
865.41
910.41
999.46
1100.51
1128.51
m/z

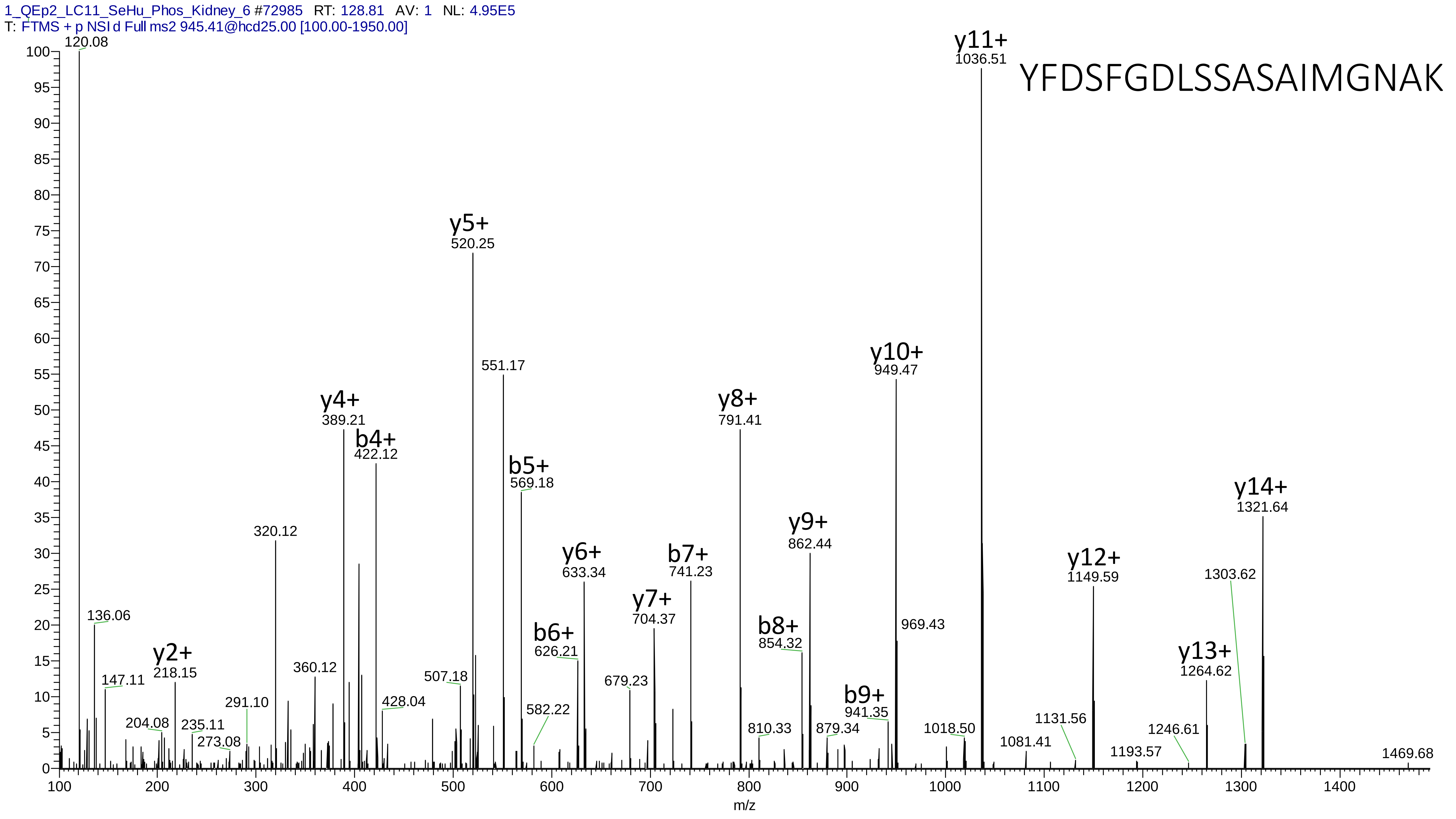
1_QEp2_LC11_SeHu_Phos_Kidney_6 #72985 RT: 128.81 AV: 1 NL: 4.95E5
T: FTMS + p NSI d Full ms2 945.41@hcd25.00 [100.00-1950.00]
YFDSFGDLSSASAIMGNAK
120.08
136.06
147.11
y2+ 218.15
204.08
235.11
273.08
291.10
320.12
360.12
y4+ 389.21
b4+ 422.12
428.04
507.18
y5+ 520.25
551.17
b5+ 569.18
582.22
b6+ 626.21
y6+ 633.34
679.23
y7+ 704.37
b7+ 741.23
y8+ 791.41
810.33
b8+ 854.32
y9+ 862.44
879.34
b9+ 941.35
y10+ 949.47
969.43
1018.50
y11+ 1036.51
1081.41
1131.56
y12+ 1149.59
1193.57
1246.61
y13+ 1264.62
1303.62
y14+ 1321.64
1469.68
m/z

# 26

- Experimental mass: 236.0658
- Unimod information:
- Possible composition: Phosphorylation + Arg, [H(1)O(3)P(1) + H(12) C(6) N(4) O ]
- Sites in Unimod:
- Sites observed:

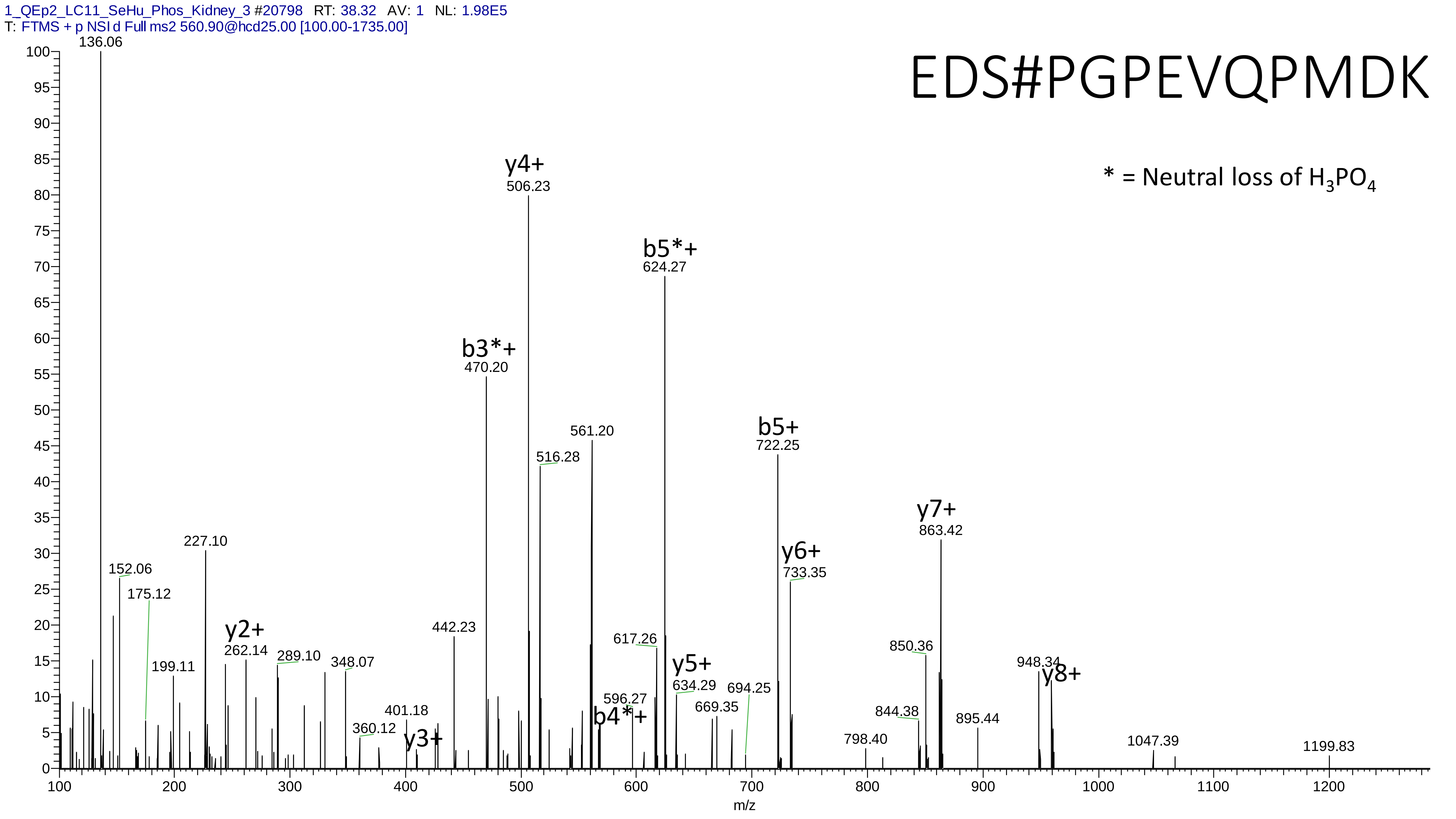
1_QEp2_LC11_SeHu_Phos_Kidney_3 #20798 RT: 38.32 AV: 1 NL: 1.98E5
T: FTMS + p NSI d Full ms2 560.90@hcd25.00 [100.00-1735.00]
EDS#PGPEVQPMDK
* = Neutral loss of H3PO4
136.06
y4+
506.23
b5*+
624.27
b3*+
470.20
561.20
516.28
b5+
722.25
y7+
863.42
y6+
733.35
227.10
152.06
175.12
y2+
262.14
442.23
617.26
y5+
634.29
850.36
948.34
y8+
199.11
289.10
348.07
596.27
b4*+
694.25
669.35
401.18
844.38
895.44
360.12
y3+
798.40
1047.39
1199.83
m/z

# 27

- Experimental mass: -195.1808
- Unimod information:
- Possible composition: Phosphorylation + Glu-Lys-loss+ Dehydrated, [H(1)O(3)P(1) + H(-12) C(-6) N(-4) O(-1) + N(-2) O(-1) ]
- Sites in Unimod:
- Sites observed:
- Features: 99.0% (103/104) peptide spectra matches contain EK at peptide n-term

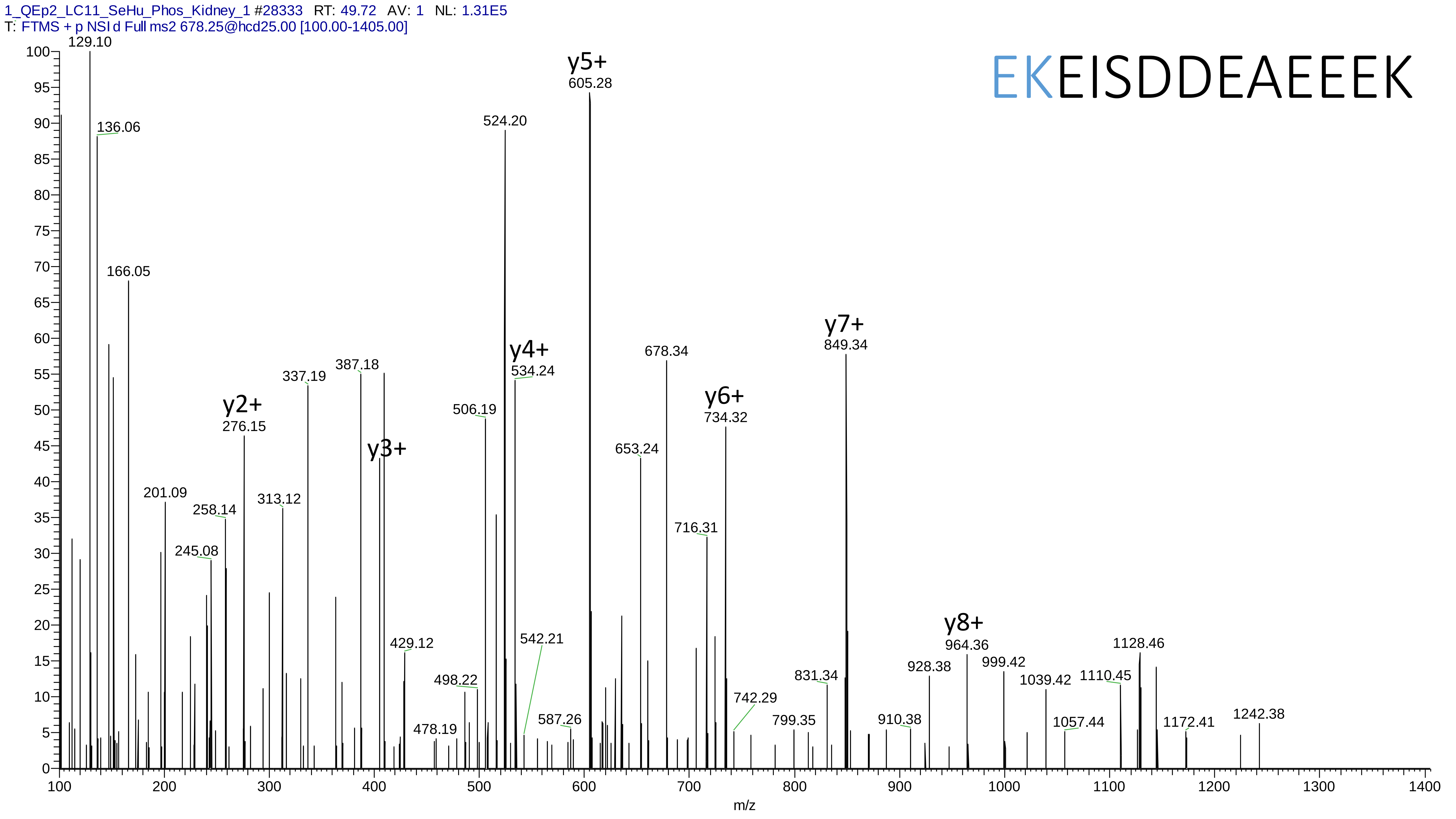
1_QEp2_LC11_SeHu_Phos_Kidney_1 #28333 RT: 49.72 AV: 1 NL: 1.31E5
T: FTMS + p NSI d Full ms2 678.25@hcd25.00 [100.00-1405.00]
EKEISDDEAEEEK
y2+
276.15
y3+
y4+
534.24
y5+
605.28
y6+
734.32
y7+
849.34
y8+
964.36
129.10
136.06
166.05
201.09
245.08
258.14
313.12
337.19
387.18
429.12
478.19
498.22
506.19
524.20
542.21
587.26
653.24
678.34
716.31
742.29
799.35
831.34
910.38
928.38
999.42
1039.42
1057.44
1110.45
1128.46
1172.41
1242.38
m/z

# 28

- Experimental mass: -9.0631
- Unimod information:
- Possible composition: Phosphorylation + (Met-loss + Acetyl), [H(1)O(3)P(1) + H(-7) C(-3) N(-1) S(-1)]
- Sites in Unimod:
- Sites observed:
- Features: 100.0% (22/22) peptide spectra matches contain Met at **protein** n-term

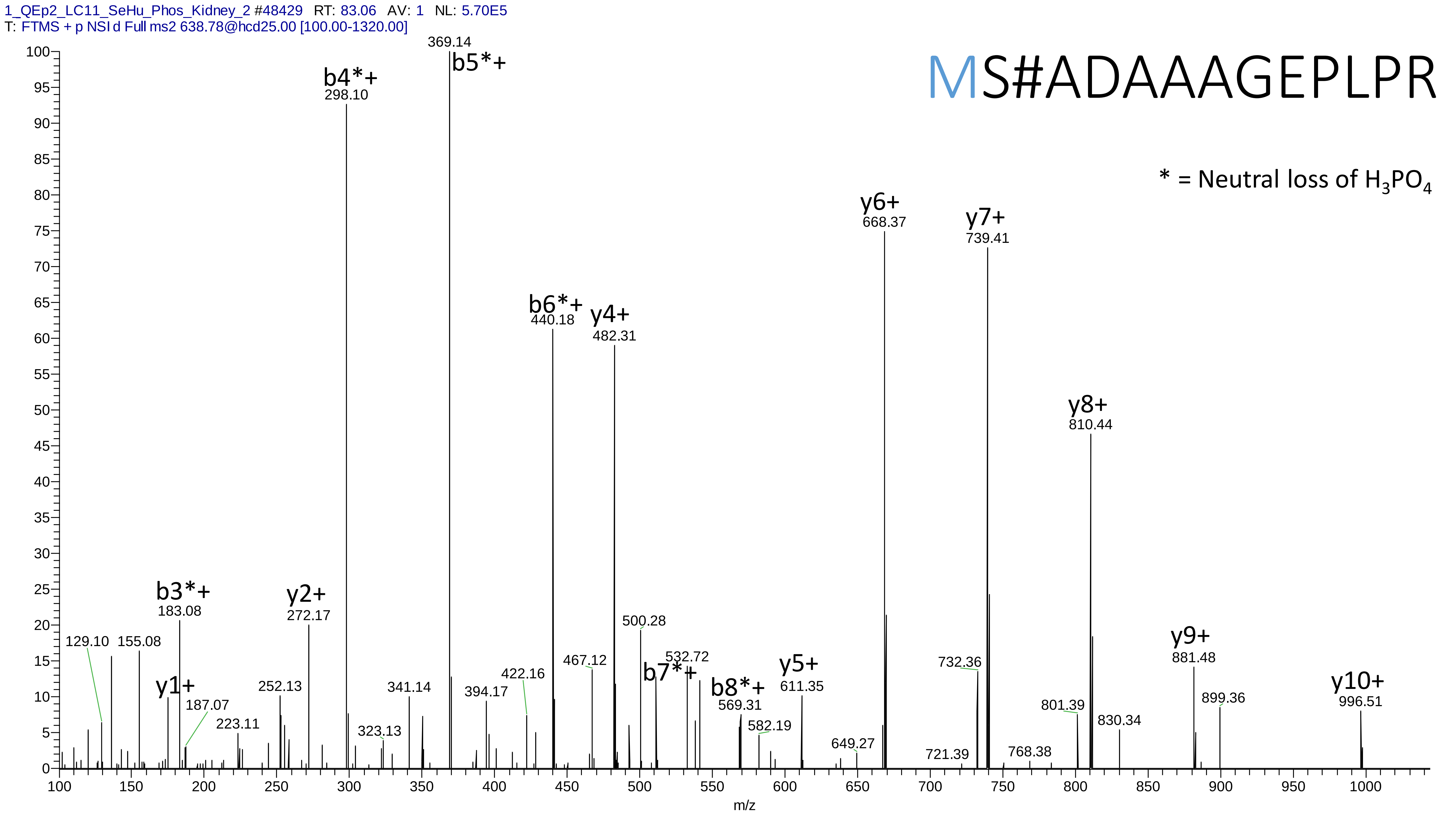
1_QEp2_LC11_SeHu_Phos_Kidney_2 #48429 RT: 83.06 AV: 1 NL: 5.70E5
T: FTMS + p NSI d Full ms2 638.78@hcd25.00 [100.00-1320.00]
MS#ADAAAGEPLPR
* = Neutral loss of $H_3PO_4$
369.14
b5*+
b4*+
298.10
y6+
668.37
y7+
739.41
b6*+
440.18
y4+
482.31
y8+
810.44
b3*+
183.08
y2+
272.17
500.28
129.10
155.08
532.72
y1+
187.07
252.13
341.14
394.17
422.16
467.12
b7*+
b8*+
569.31
y5+
611.35
732.36
y9+
881.48
y10+
996.51
899.36
801.39
830.34
223.11
323.13
582.19
649.27
721.39
768.38
m/z

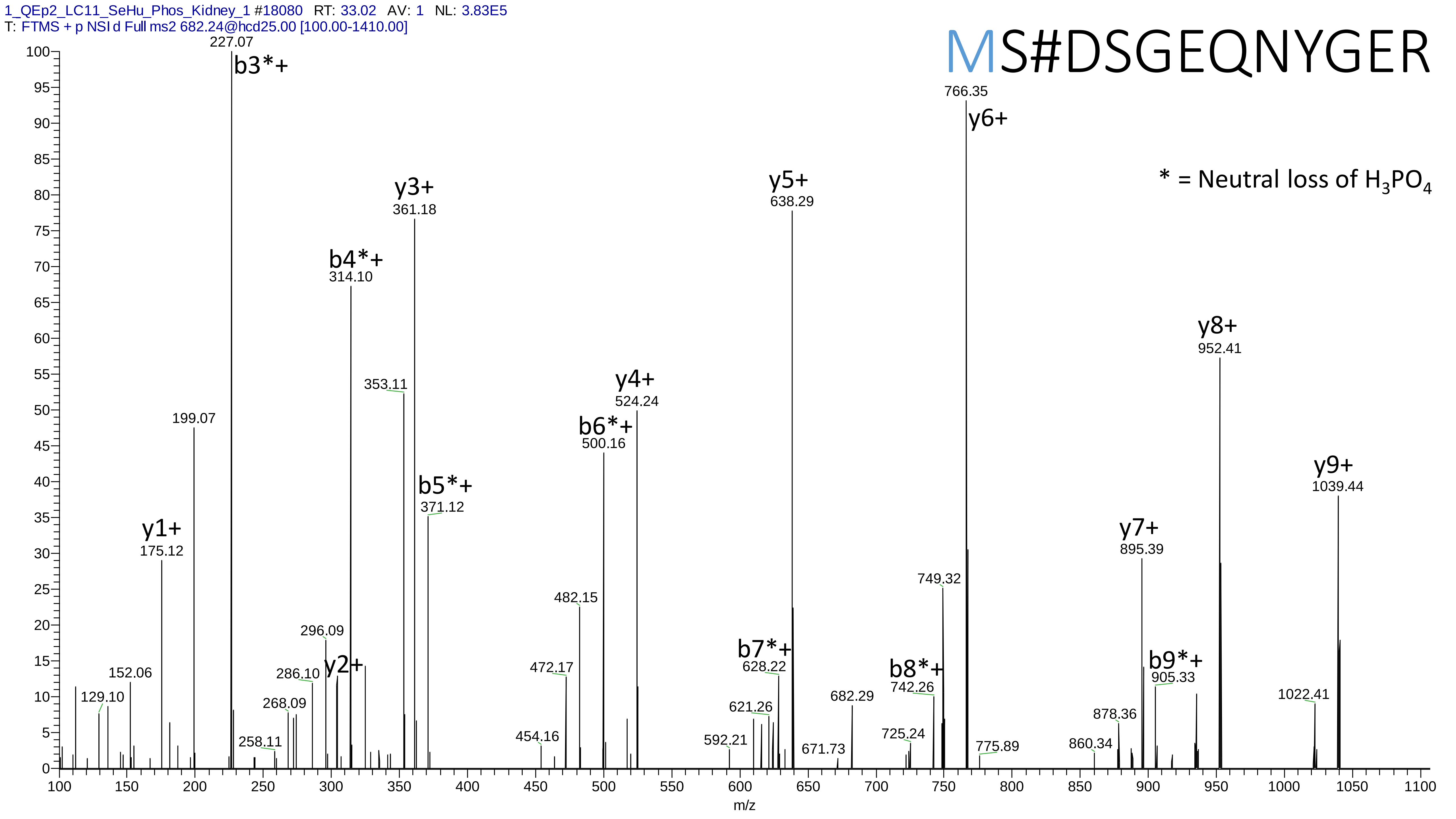

1_QEp2_LC11_SeHu_Phos_Kidney_1 #18080 RT: 33.02 AV: 1 NL: 3.83E5
T: FTMS + p NSI d Full ms2 682.24@hcd25.00 [100.00-1410.00]
MS#DSGEQNYGER
* = Neutral loss of H3PO4
227.07 b3*+
766.35 y6+
y5+ 638.29
y3+ 361.18
b4*+ 314.10
y8+ 952.41
353.11
y4+ 524.24
b6*+ 500.16
199.07
y9+ 1039.44
b5*+ 371.12
y1+ 175.12
y7+ 895.39
749.32
482.15
296.09
b7*+ 628.22
y2+
472.17
286.10
152.06
b8*+ 742.26
b9*+ 905.33
129.10
268.09
682.29
1022.41
621.26
878.36
725.24
454.16
258.11
592.21
671.73
775.89
860.34
m/z

# 29

- Experimental mass: -147.1727
- Unimod information:
- Possible composition: Phosphorylation + Ala-Arg-loss, [H(1)O(3)P(1) + H(-5) C(-3) N(-1) O(-1) + H(-12) C(-6) N(-4) O(-1) ]
- Sites in Unimod:
- Sites observed:
- Features: 98.9% (92/93) peptide spectra matches contain AR at peptide n-term

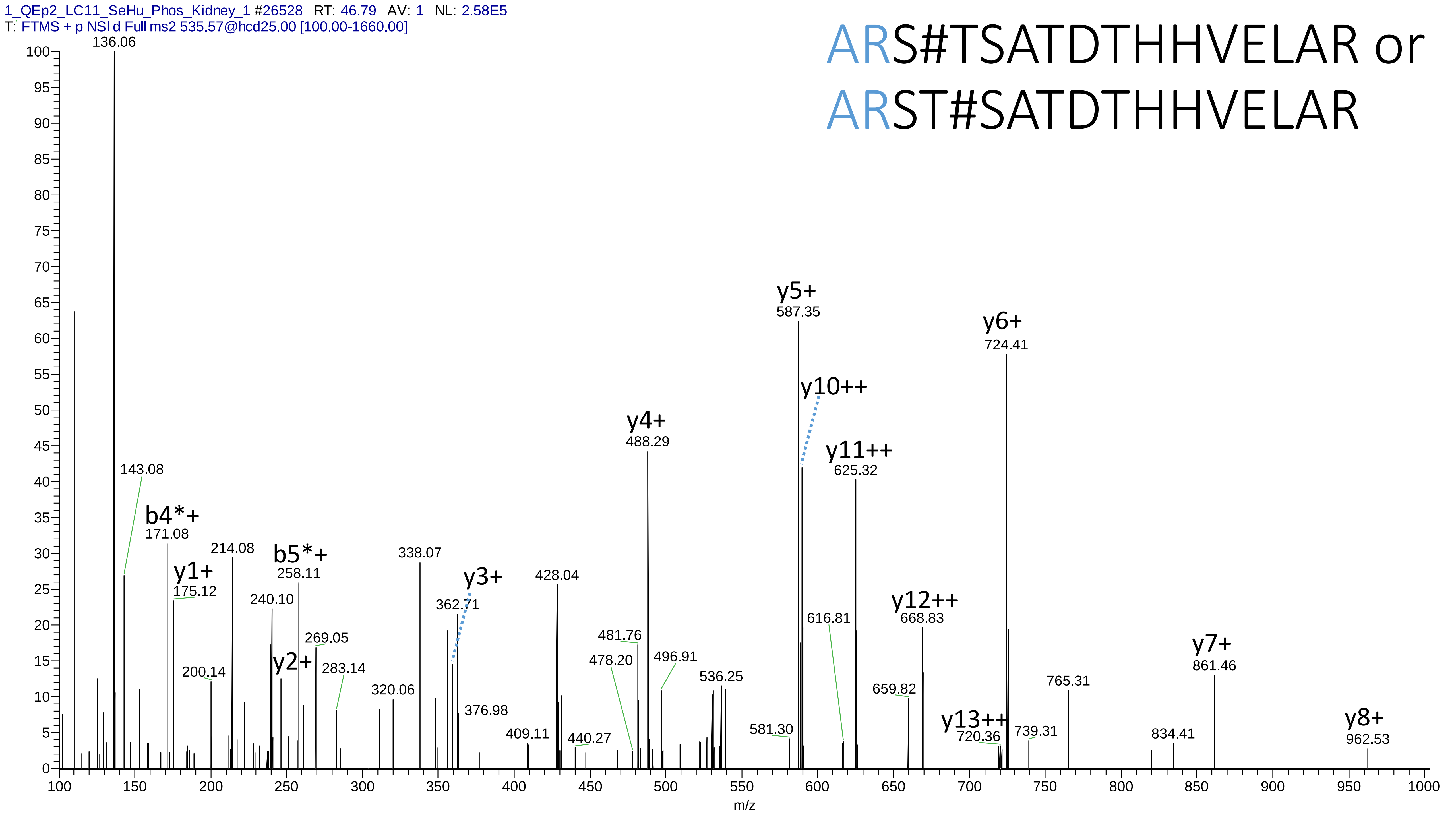
1_QEp2_LC11_SeHu_Phos_Kidney_1 #26528 RT: 46.79 AV: 1 NL: 2.58E5
T: FTMS + p NSI d Full ms2 535.57@hcd25.00 [100.00-1660.00]
ARS#TSATDTHHVELAR or
ARST#SATDTHHVELAR
136.06
143.08
b4*+
171.08
y1+
175.12
200.14
214.08
240.10
y2+
b5*+
258.11
269.05
283.14
320.06
338.07
y3+
362.71
376.98
409.11
428.04
440.27
478.20
481.76
y4+
488.29
496.91
536.25
581.30
y5+
587.35
y10++
y11++
625.32
616.81
659.82
y12++
668.83
y13++
720.36
y6+
724.41
739.31
765.31
834.41
y7+
861.46
y8+
962.53
m/z

# 30

- Experimental mass: -240.2926
- Unimod information:
- Possible composition: Phosphorylation * 2 + Leu-Met-Arg-loss, [H(1)O(3)P(1) + H(-11) C(-6) N(-1) O(-1) + H(-9) C(-5) N(-1) O(-1) S(-1) + H(-12) C(-6) N(-4) O(-1)]
- Sites in Unimod:
- Sites observed:
- Features: 100.0% (84/84) peptide spectra matches contain LMR at peptide n-term

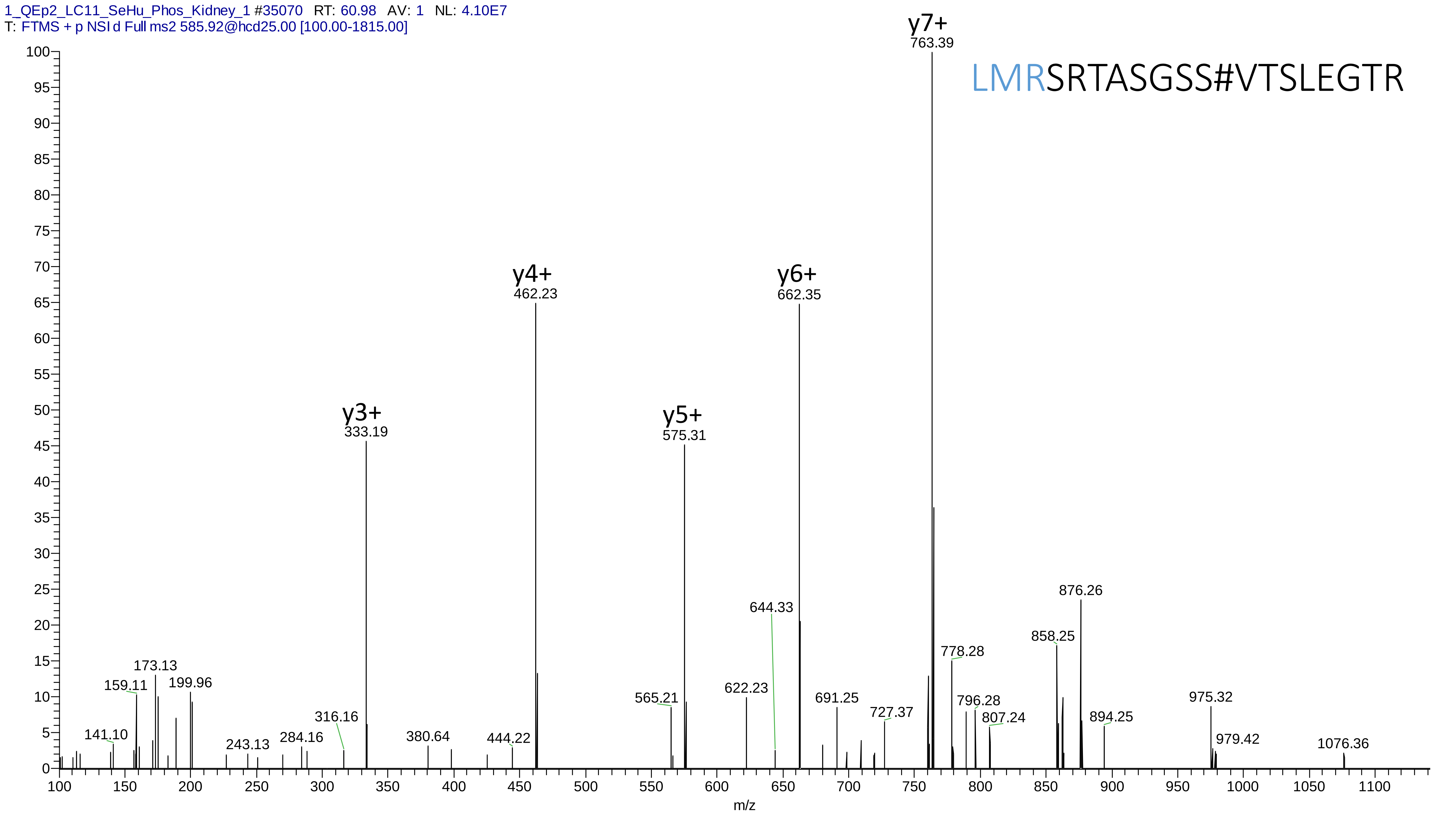
1_QEp2_LC11_SeHu_Phos_Kidney_1 #35070 RT: 60.98 AV: 1 NL: 4.10E7
T: FTMS + p NSI d Full ms2 585.92@hcd25.00 [100.00-1815.00]
LMRSRTASGSS#VTSLEGTR
y7+ 763.39
y6+ 662.35
y4+ 462.23
y5+ 575.31
y3+ 333.19
876.26
644.33
858.25
778.28
173.13
159.11
199.96
622.23
565.21
691.25
796.28
975.32
727.37
807.24
894.25
316.16
141.10
380.64
444.22
243.13
284.16
979.42
1076.36
m/z
