## Supplementary material 3 for "MetaLab 2.0 enables accurate post-translational modifications profiling in metaproteomics"

### Supplementary material 3: Manually annotated spectra of the top 30 identified potential modifications

Dataset: Hela_phos

# 1

- Experimental mass: 79.9659
- Unimod information: phosphorylation
- Possible composition: H O(3) P
- Sites in Unimod: S;T;Y;D;H;C;R;K
- Sites observed:
- Since phosphorylation is common in this dataset, spectra are not showed

# 2

- Experimental mass: 80.9497
- Unimod information:
- Possible composition: Phosphorylation + Deamidation, [H(1)O(3)P(1) + H(-1) N(-1) O]; could also be Phosphorylation + Carboxymethyl replace of Carbamidomethyl on Cys
- Sites in Unimod:
- Sites observed:
- Features:  87.6% (560/639) peptides contain Asn/Gln

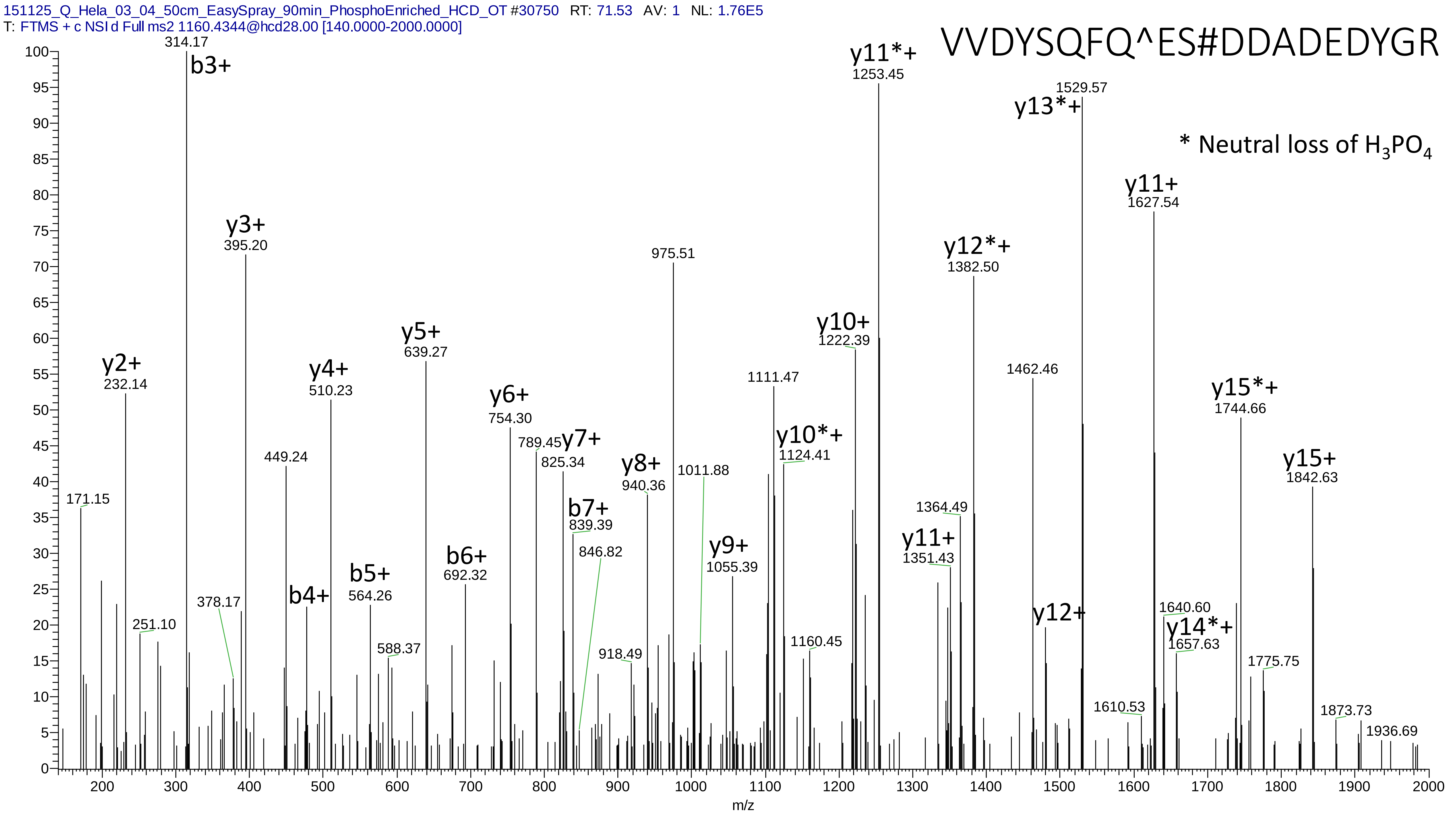
151125_Q_Hela_03_04_50cm_EasySpray_90min_PhosphoEnriched_HCD_OT #30750 RT: 71.53 AV: 1 NL: 1.76E5
T: FTMS + c NSI d Full ms2 1160.4344@hcd28.00 [140.0000-2000.0000]
VVDYSQFQ^ES#DDADEDYGR
* Neutral loss of H3PO4
314.17
b3+
y3+
395.20
y2+
232.14
y4+
510.23
y5+
639.27
y6+
754.30
789.45
y7+
825.34
449.24
171.15
251.10
378.17
b4+
b5+
564.26
588.37
b6+
692.32
b7+
839.39
846.82
y8+
940.36
918.49
975.51
1011.88
y9+
1055.39
1111.47
y10*+
1124.41
1160.45
y10+
1222.39
y11*+
1253.45
y11+
1351.43
1364.49
y12*+
1382.50
1462.46
y12+
1529.57
y13*+
y11+
1627.54
1640.60
y14*+
1657.63
1610.53
y15*+
1744.66
1775.75
y15+
1842.63
1873.73
1936.69
m/z

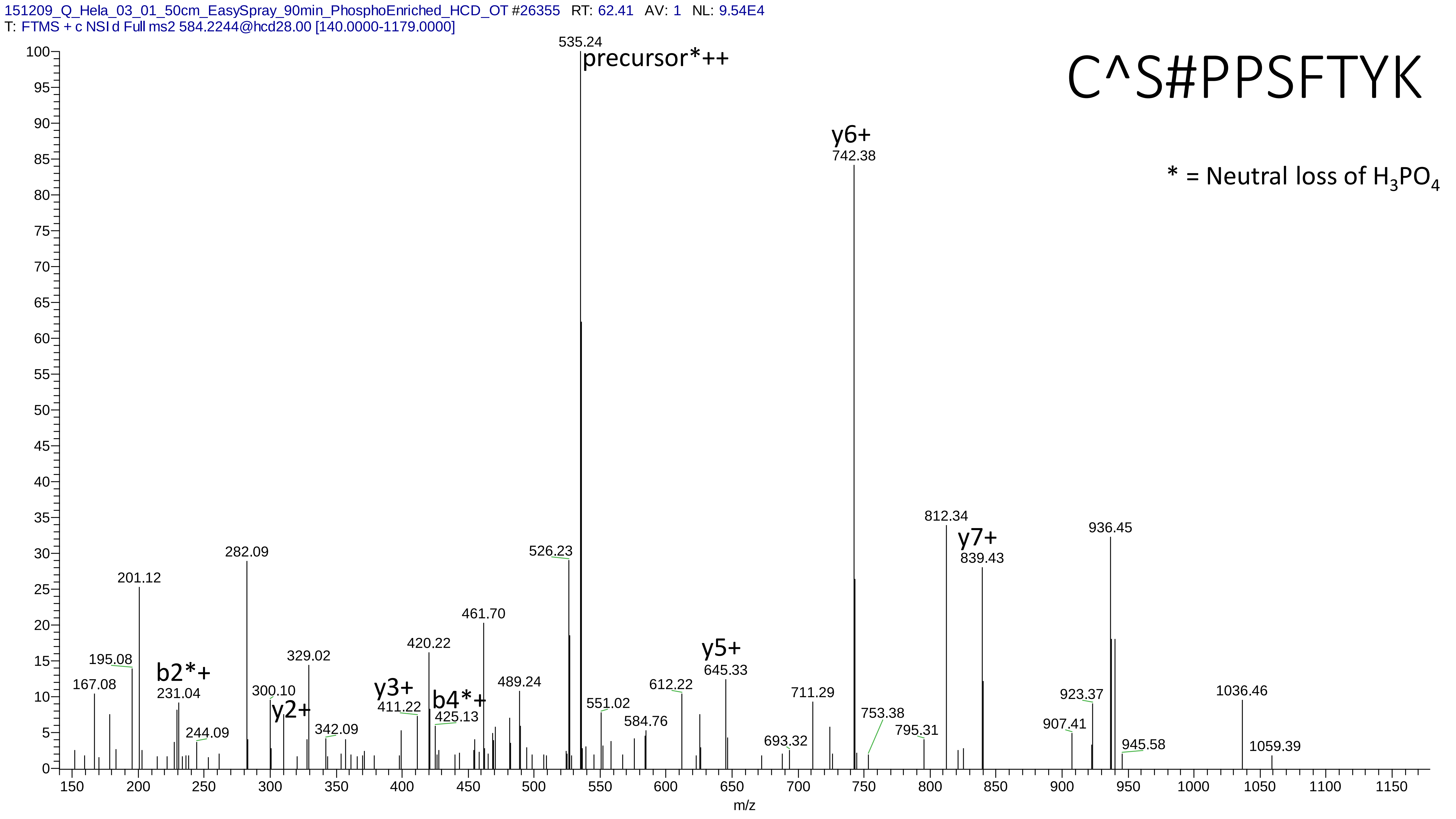
151209_Q_Hela_03_01_50cm_EasySpray_90min_PhosphoEnriched_HCD_OT #26355 RT: 62.41 AV: 1 NL: 9.54E4
T: FTMS + c NSI d Full ms2 584.2244@hcd28.00 [140.0000-1179.0000]
C^S#PPSFTYK
* = Neutral loss of H3PO4
precursor*++
535.24
y6+
742.38
812.34
y7+
839.43
936.45
282.09
526.23
201.12
461.70
420.22
329.02
195.08
b2*+
231.04
167.08
300.10
y2+
y3+
411.22
b4*+
425.13
489.24
244.09
342.09
y5+
645.33
612.22
551.02
584.76
711.29
693.32
753.38
795.31
923.37
907.41
945.58
1036.46
1059.39
m/z

# 3

- Experimental mass: 0.9834
- Unimod information: deamidation
- Possible composition: H(-1) N(-1) O
- Sites in Unimod: N;Q;R;F
- Sites observed:

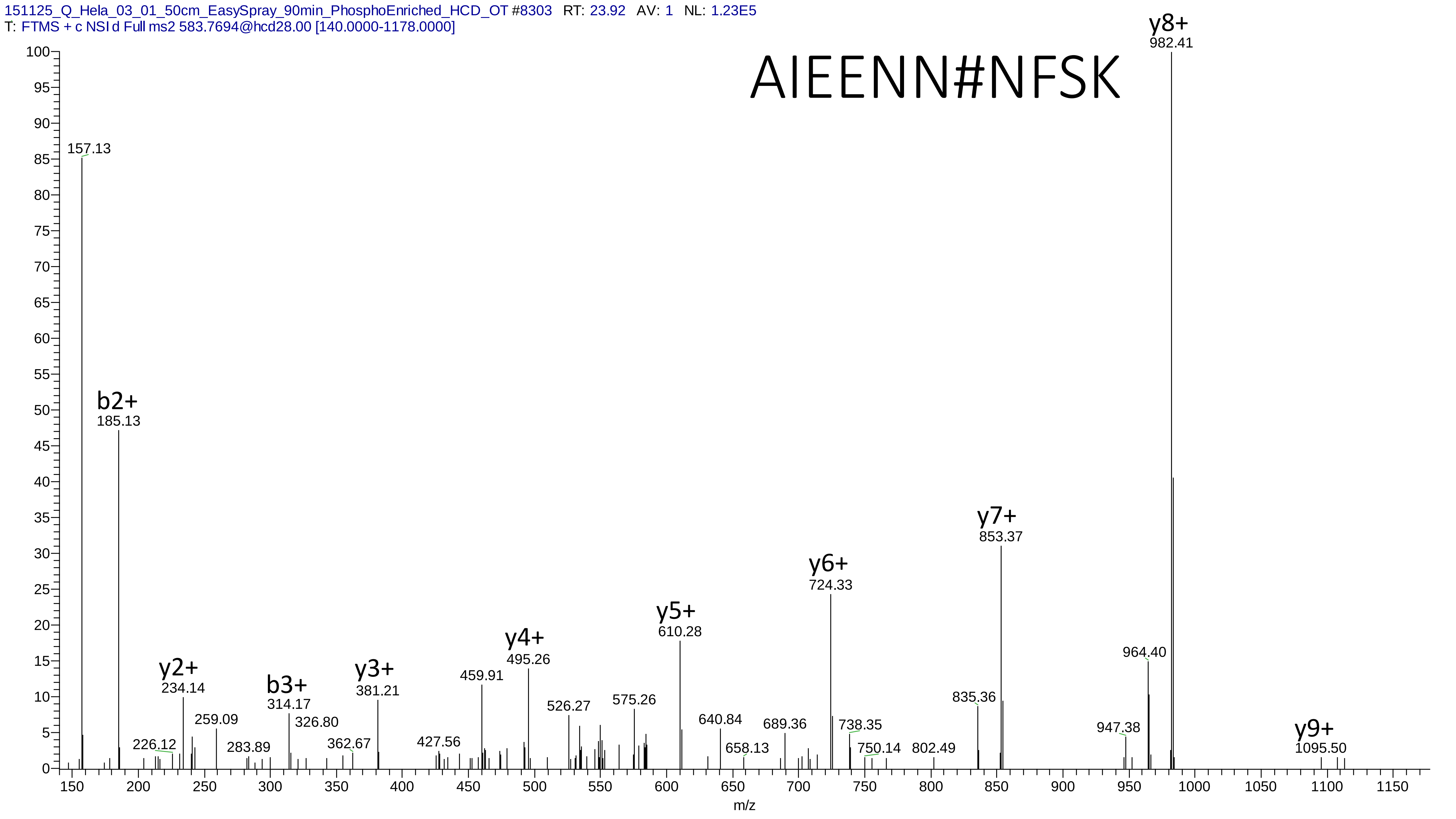

151125_Q_Hela_03_01_50cm_EasySpray_90min_PhosphoEnriched_HCD_OT #8303 RT: 23.92 AV: 1 NL: 1.23E5
T: FTMS + c NSI d Full ms2 583.7694@hcd28.00 [140.0000-1178.0000]
AIEENN#NFSK
157.13
b2+
185.13
y2+
234.14
226.12
259.09
283.89
b3+
314.17
326.80
362.67
y3+
381.21
427.56
459.91
y4+
495.26
526.27
575.26
y5+
610.28
640.84
658.13
689.36
y6+
724.33
738.35
750.14
802.49
835.36
y7+
853.37
947.38
964.40
y8+
982.41
y9+
1095.50
m/z

# 4

- Experimental mass: 133.8816
- Unimod information:
- Possible composition: Phosphorylation + Cation:Fe[II], [H(1)O(3)P(1) + H(-2) Fe]
- Sites in Unimod:
- Sites observed: S;T
- Features: phosphorylation sites determined, cation modified sites are not observed

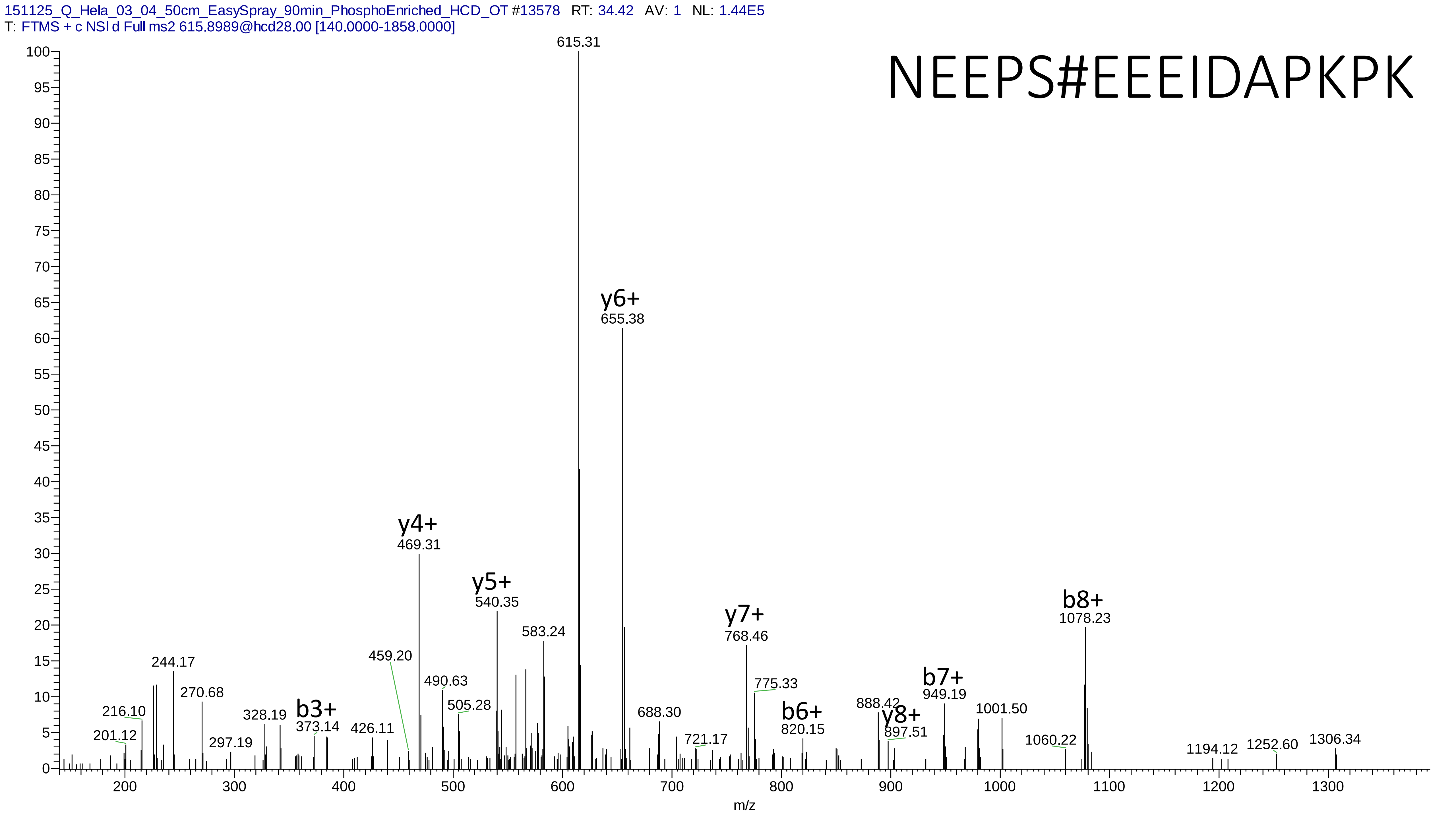
151125_Q_Hela_03_04_50cm_EasySpray_90min_PhosphoEnriched_HCD_OT #13578 RT: 34.42 AV: 1 NL: 1.44E5
T: FTMS + c NSI d Full ms2 615.8989@hcd28.00 [140.0000-1858.0000]
NEEPS#EEEIDAPKPK
615.31
y6+
655.38
y4+
469.31
y5+
540.35
583.24
y7+
768.46
b8+
1078.23
459.20
244.17
270.68
216.10
201.12
297.19
328.19
b3+
373.14
426.11
490.63
505.28
688.30
721.17
775.33
b6+
820.15
888.42
y8+
897.51
b7+
949.19
1001.50
1060.22
1194.12
1252.60
1306.34
m/z

# 5

- Experimental mass: 159.9316
- Unimod information: pyrophospho
- Possible composition: H(2)O(6)P(2)
- Sites in Unimod: S;T
- Sites observed:
- Features:  100% (199/199) peptides contain equal or more than two Ser/Thr

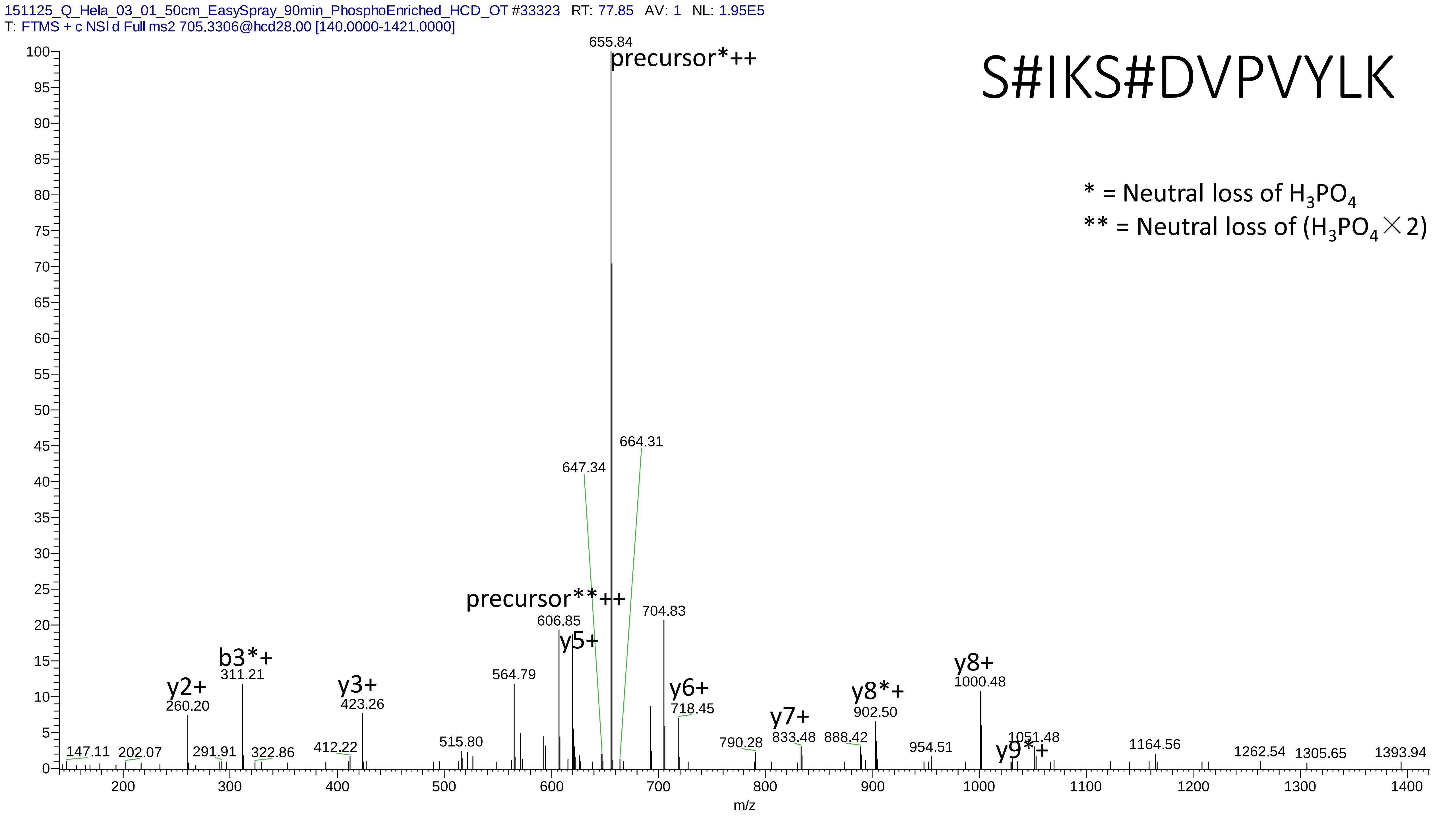
151125_Q_Hela_03_01_50cm_EasySpray_90min_PhosphoEnriched_HCD_OT #33323 RT: 77.85 AV: 1 NL: 1.95E5
T: FTMS + c NSI d Full ms2 705.3306@hcd28.00 [140.0000-1421.0000]
S#IKS#DVPVYLK
* = Neutral loss of H3PO4
** = Neutral loss of (H3PO4×2)
655.84
precursor*++
664.31
647.34
precursor**++
606.85
y5+
704.83
y6+
718.45
b3*+
311.21
y2+
260.20
y3+
423.26
564.79
515.80
412.22
147.11
202.07
291.91
322.86
790.28
y7+
833.48
888.42
y8*+
902.50
y8+
1000.48
954.51
1051.48
y9*+
1164.56
1262.54
1305.65
1393.94
m/z

# 6

- Experimental mass: -17.0266
- Unimod information: Ammonia-loss or Gln->pyro-Glu
- Possible composition: H(-3)N(-1)
- Sites in Unimod: nC;[S;[T;N; Q
- Sites observed: nQ; nC
- Features:  75.2% (115/153) peptides contain Gln at peptide n-term (Pyro-glu from Q), 12.4% (19/153) peptides contain Cys at peptide n-term

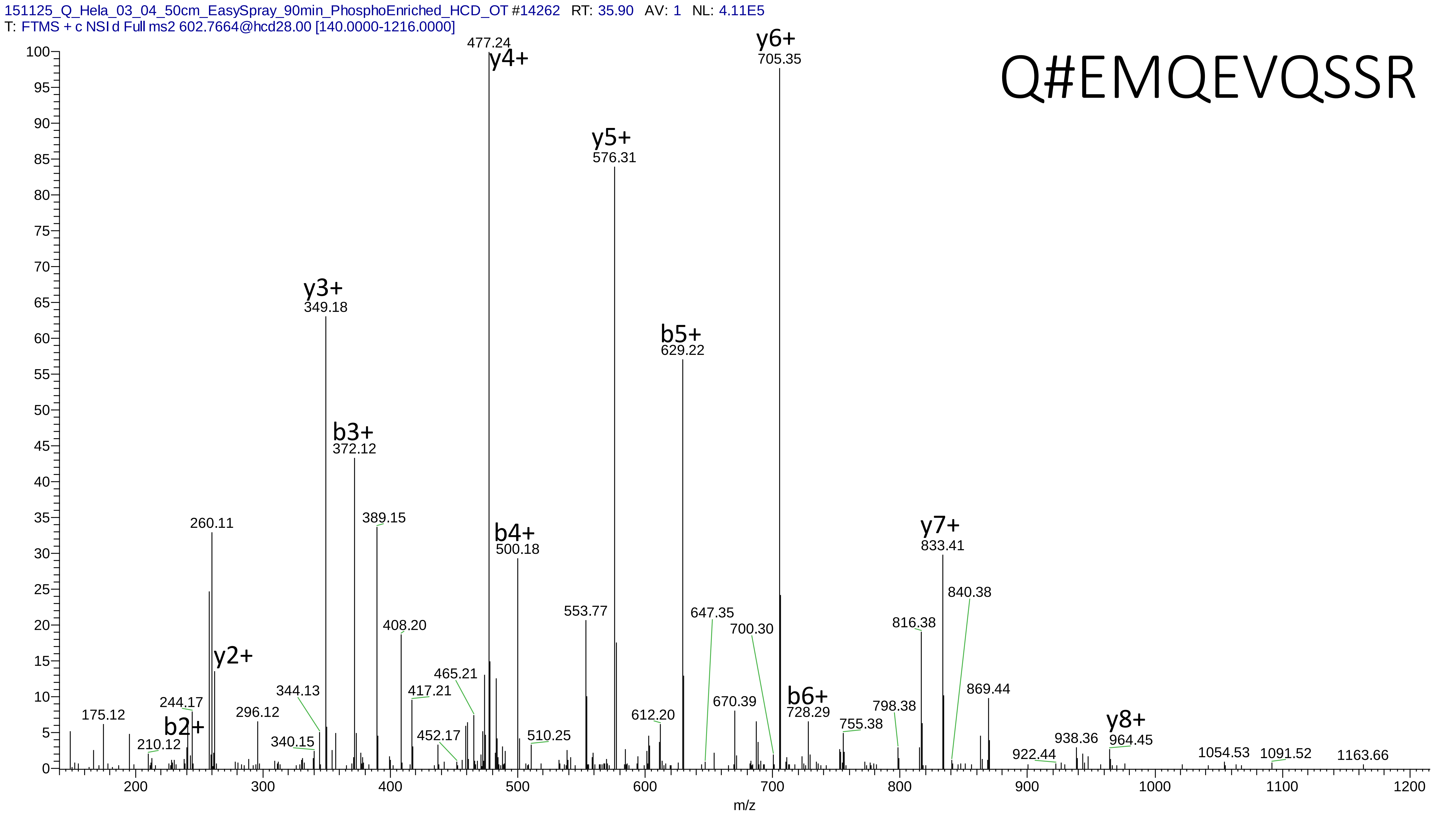
151125_Q_Hela_03_04_50cm_EasySpray_90min_PhosphoEnriched_HCD_OT #14262 RT: 35.90 AV: 1 NL: 4.11E5
T: FTMS + c NSI d Full ms2 602.7664@hcd28.00 [140.0000-1216.0000]
Q#EMQEVQSSR
y6+ 705.35
477.24 y4+
y5+ 576.31
b5+ 629.22
y3+ 349.18
b3+ 372.12
389.15
260.11
b4+ 500.18
y7+ 833.41
840.38
553.77
647.35
700.30
408.20
816.38
y2+
465.21
344.13
417.21
b6+ 728.29
869.44
670.39
798.38
175.12
244.17
b2+
296.12
612.20
755.38
y8+ 964.45
210.12
340.15
452.17
510.25
938.36
922.44
1054.53
1091.52
1163.66
m/z

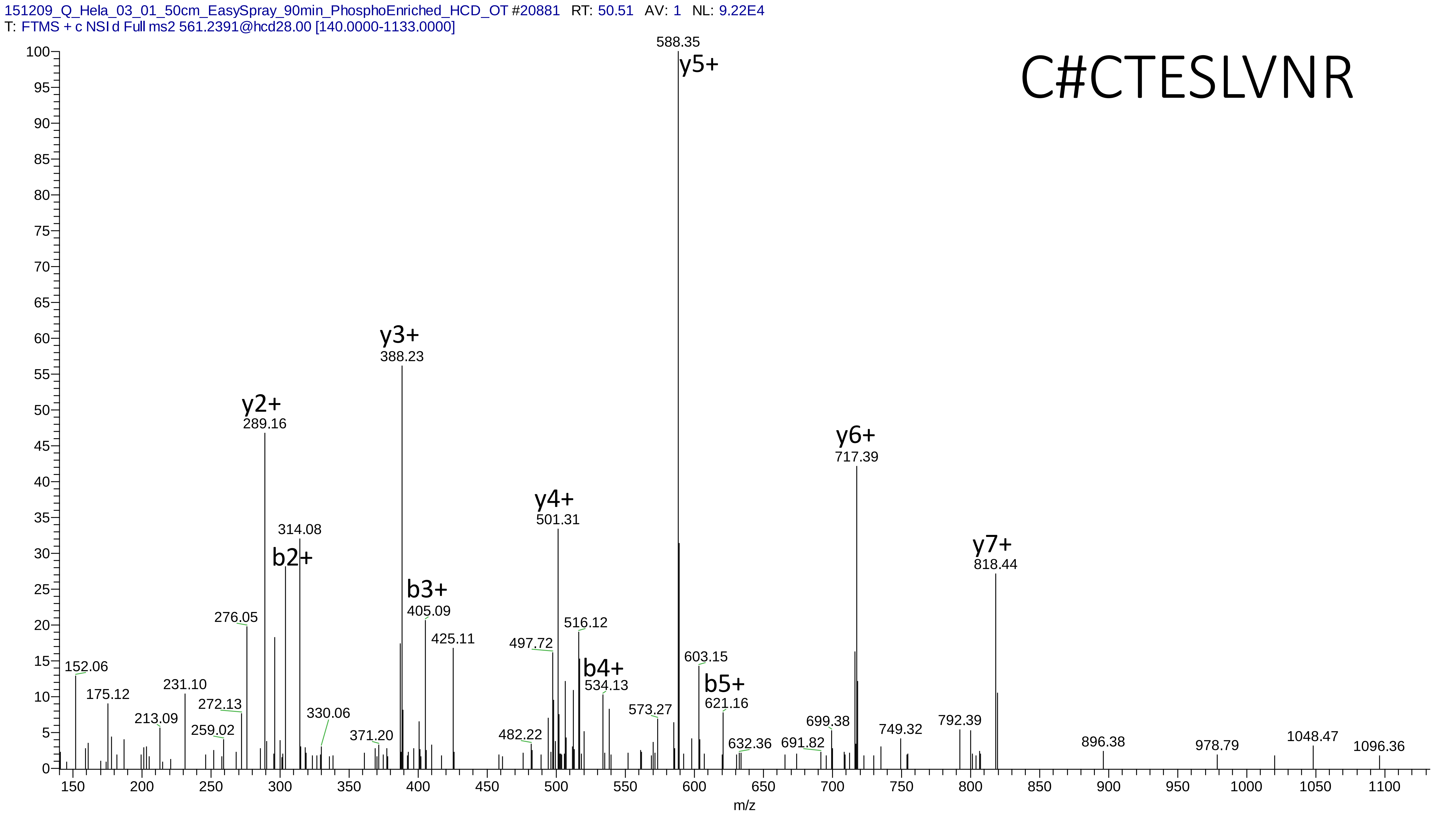
151209_Q_Hela_03_01_50cm_EasySpray_90min_PhosphoEnriched_HCD_OT #20881 RT: 50.51 AV: 1 NL: 9.22E4
T: FTMS + c NSI d Full ms2 561.2391@hcd28.00 [140.0000-1133.0000]
C#CTESLVNR
588.35
y5+
y3+
388.23
y2+
289.16
314.08
b2+
y4+
501.31
y6+
717.39
y7+
818.44
b3+
405.09
425.11
276.05
497.72
516.12
b4+
534.13
603.15
b5+
621.16
152.06
175.12
231.10
213.09
272.13
259.02
330.06
371.20
482.22
573.27
632.36
691.82
699.38
749.32
792.39
896.38
978.79
1048.47
1096.36
m/z

# 7

- Experimental mass: 62.9394
- Unimod information:
- Possible composition: Phosphorylation + Ammonia-loss, [H(1)O(3)P(1) + H(-3)N(-1)]
- Sites in Unimod:
- Sites observed:
- Features:  76.2% (96/126) peptides contain Gln at peptide n-term (Pyro-glu from Q), 15.1% (19/126) peptides contain Cys at peptide n-term

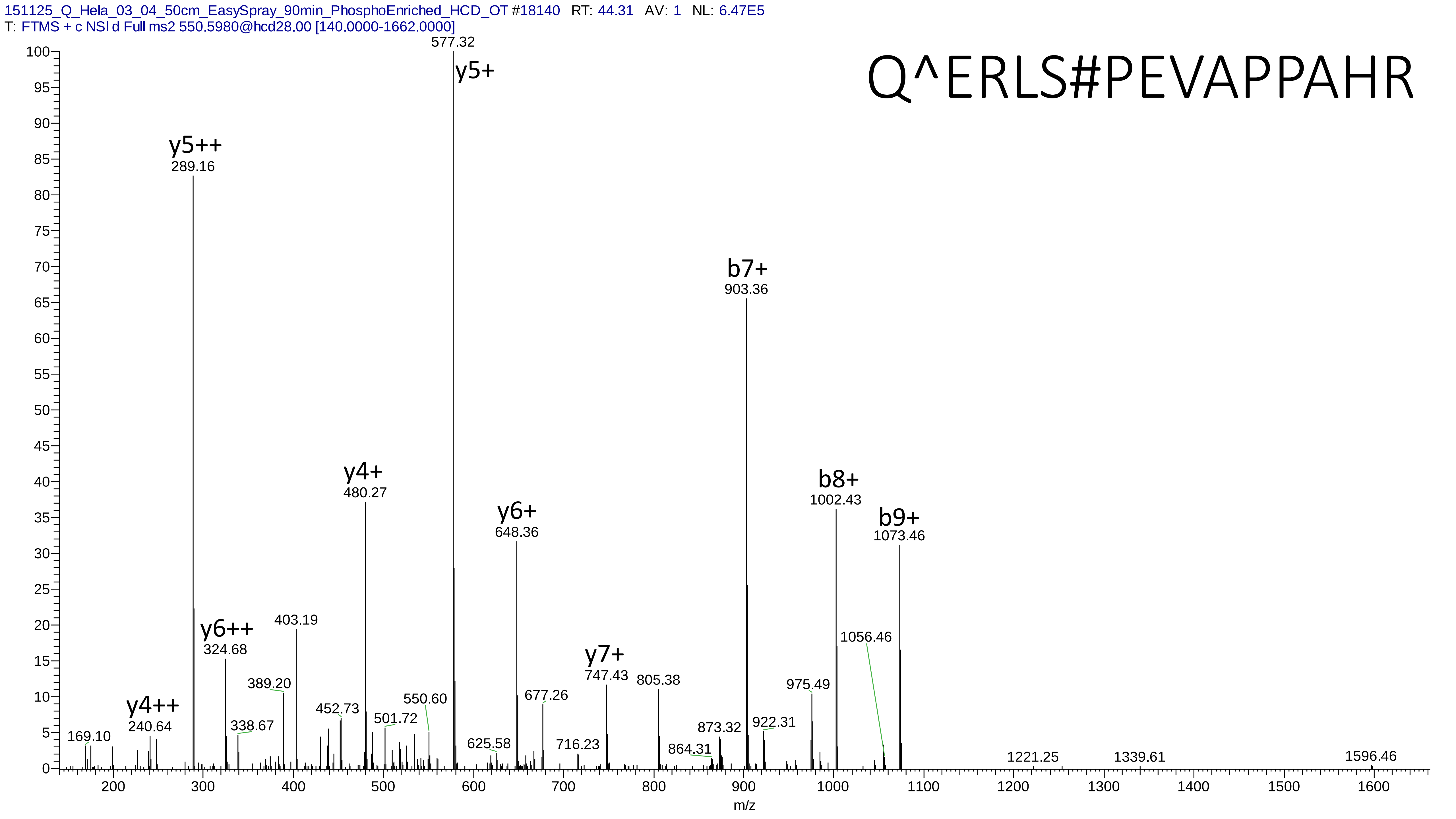
151125_Q_Hela_03_04_50cm_EasySpray_90min_PhosphoEnriched_HCD_OT #18140 RT: 44.31 AV: 1 NL: 6.47E5
T: FTMS + c NSI d Full ms2 550.5980@hcd28.00 [140.0000-1662.0000]
Q^ERLS#PEVAPPAHR
577.32
y5+
y5++
289.16
b7+
903.36
y4+
480.27
b8+
1002.43
y6+
648.36
b9+
1073.46
1056.46
y6++
324.68
403.19
y7+
747.43
805.38
389.20
975.49
677.26
y4++
240.64
452.73
550.60
501.72
873.32
922.31
338.67
169.10
625.58
716.23
864.31
1221.25
1339.61
1596.46
m/z

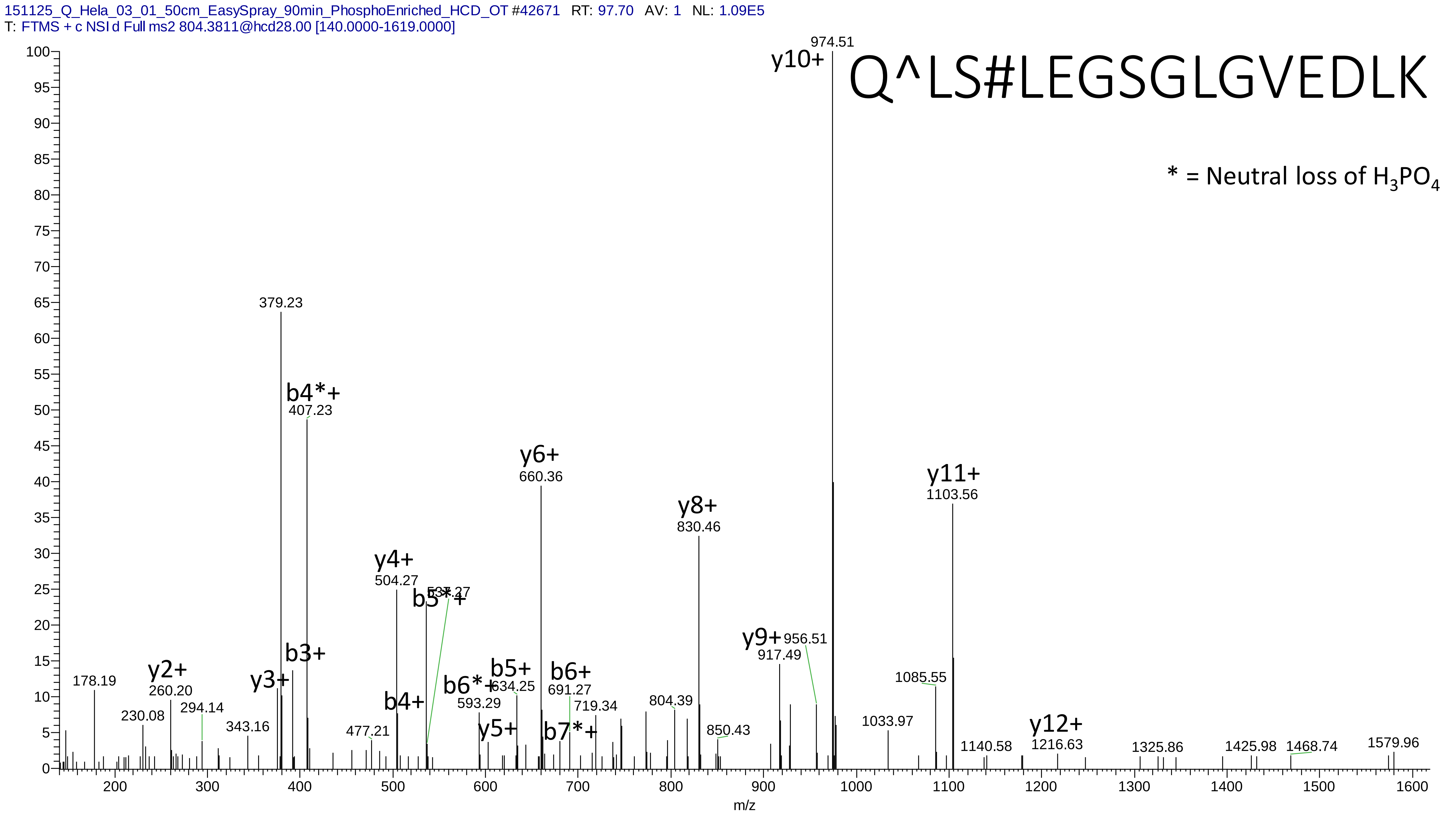
151125_Q_Hela_03_01_50cm_EasySpray_90min_PhosphoEnriched_HCD_OT #42671 RT: 97.70 AV: 1 NL: 1.09E5
T: FTMS + c NSI d Full ms2 804.3811@hcd28.00 [140.0000-1619.0000]
Q^LS#LEGSGLGVEDLK
* = Neutral loss of $H_3PO_4$
y10+ 974.51
379.23
b4*+ 407.23
y6+ 660.36
y11+ 1103.56
y8+ 830.46
y4+ 504.27
b5*+ 537.27
y9+ 917.49
956.51
b3+
178.19
y2+ 260.20
y3+
b5+ 634.25
b6+ 691.27
b6*+ 593.29
b4+
1085.55
230.08
294.14
719.34
804.39
343.16
y5+
b7*+
1033.97
y12+ 1216.63
477.21
850.43
1140.58
1325.86
1425.98
1468.74
1579.96
m/z

# 8

- Experimental mass: 101.947
- Unimod information:
- Possible composition: Phosphorylation + Cation:Na, [H(1)O(3)P(1) + H(-1) Na]
- Sites in Unimod:
- Sites observed:
- Features: phosphorylation sites determined, cation modified sites are not observed

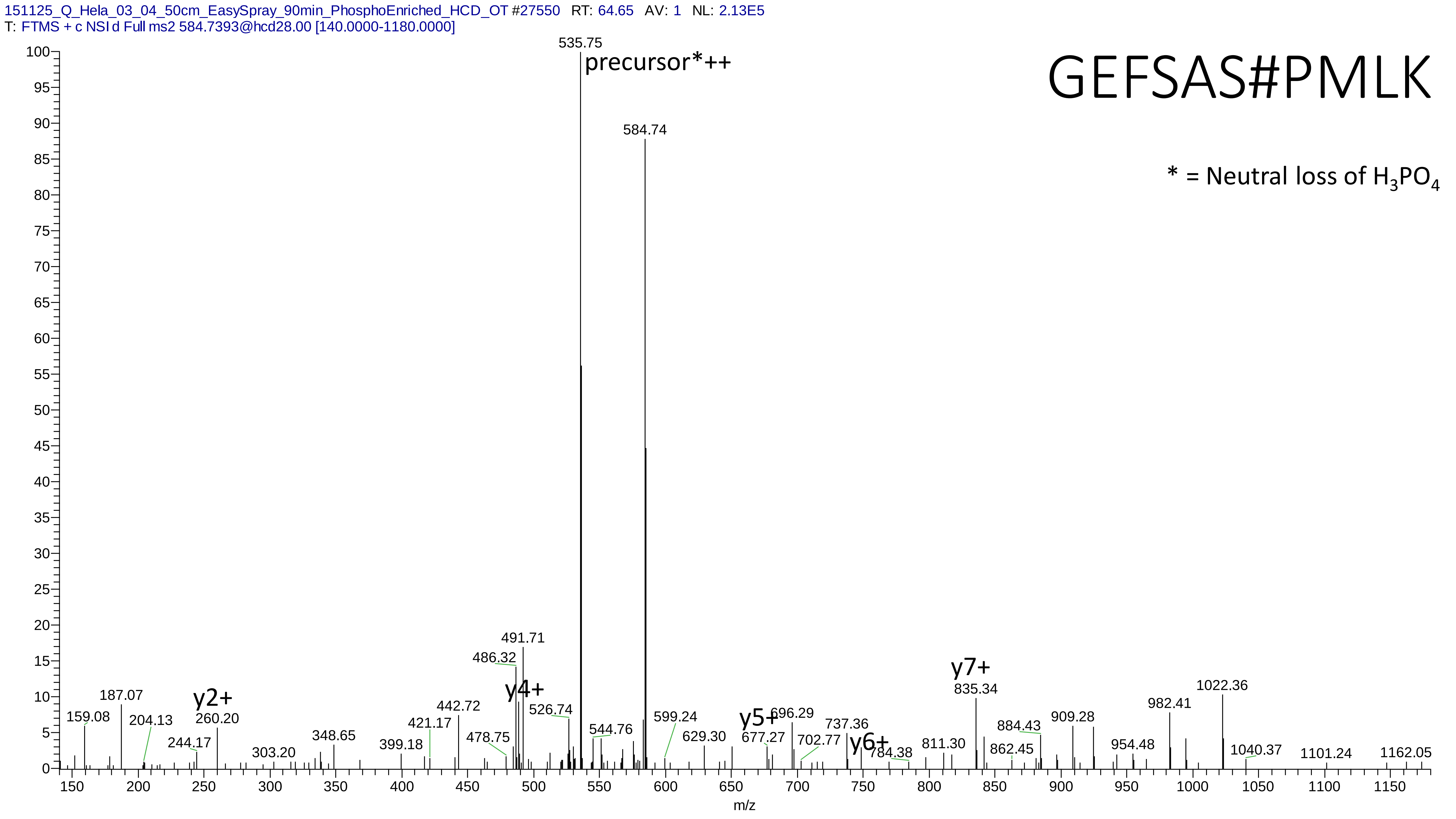

151125_Q_Hela_03_04_50cm_EasySpray_90min_PhosphoEnriched_HCD_OT #27550 RT: 64.65 AV: 1 NL: 2.13E5
T: FTMS + c NSI d Full ms2 584.7393@hcd28.00 [140.0000-1180.0000]
GEFSAS#PMLK
* = Neutral loss of $H_3PO_4$
precursor*++
535.75
584.74
491.71
486.32
y2+
y4+
y5+
y6+
y7+
187.07
159.08
204.13
260.20
244.17
303.20
348.65
399.18
421.17
442.72
478.75
526.74
544.76
599.24
629.30
677.27
696.29
702.77
737.36
784.38
811.30
835.34
862.45
884.43
909.28
954.48
982.41
1022.36
1040.37
1101.24
1162.05
m/z

# 9

- Experimental mass: 31.9881
- Unimod information: Dioxidation
- Possible composition:
- Sites in Unimod: Y;W;F;M;K;R;P;C;U
- Sites observed: W
- Features: 94.1% (16/17) peptides contain Trp

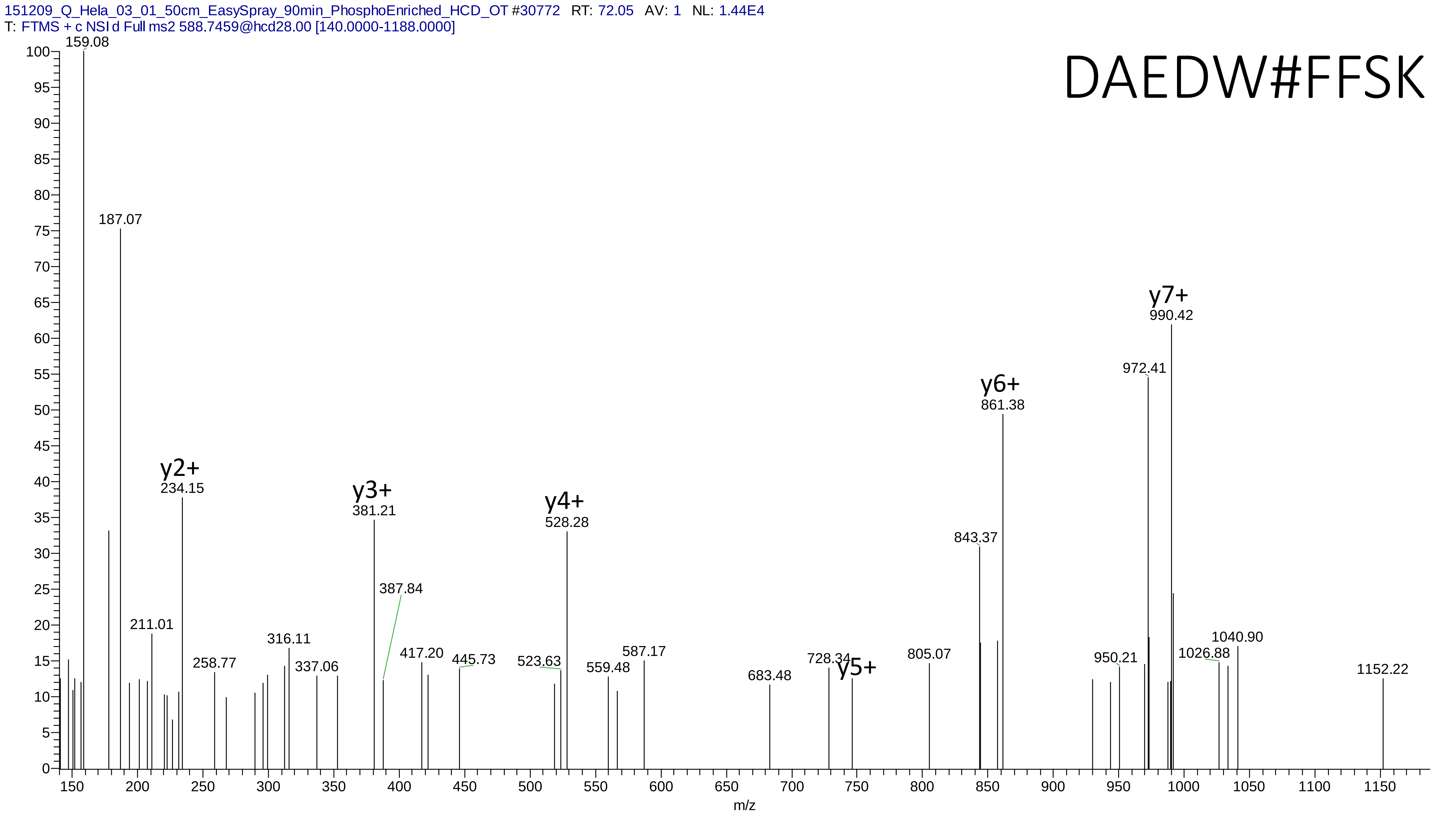
151209_Q_Hela_03_01_50cm_EasySpray_90min_PhosphoEnriched_HCD_OT #30772 RT: 72.05 AV: 1 NL: 1.44E4
T: FTMS + c NSI d Full ms2 588.7459@hcd28.00 [140.0000-1188.0000]
DAEDW#FFSK
y2+
y3+
y4+
y5+
y6+
y7+
159.08
187.07
234.15
211.01
258.77
316.11
337.06
381.21
387.84
417.20
445.73
523.63
528.28
559.48
587.17
683.48
728.34
805.07
843.37
861.38
950.21
972.41
990.42
1026.88
1040.90
1152.22
m/z

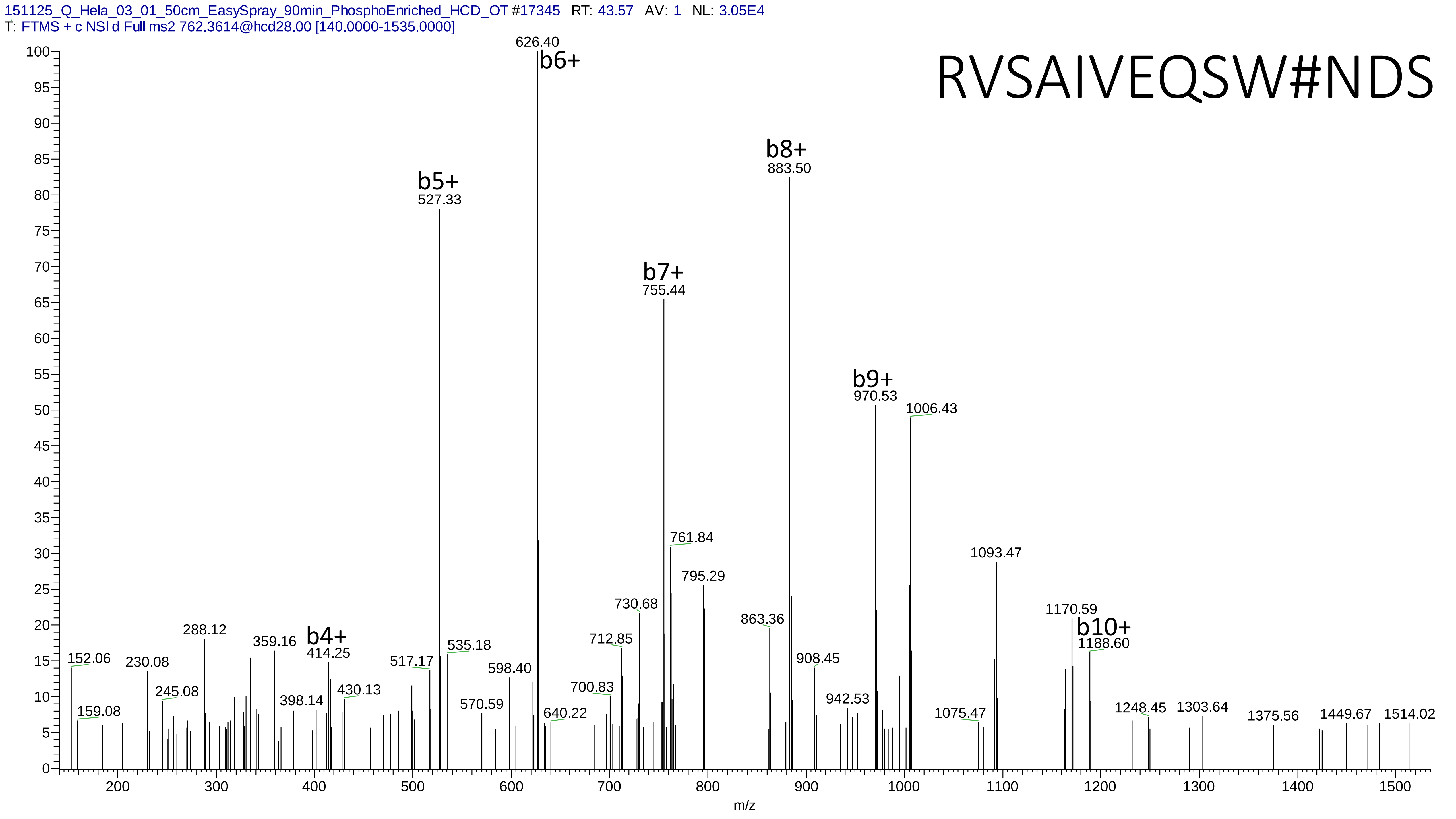
151125_Q_Hela_03_01_50cm_EasySpray_90min_PhosphoEnriched_HCD_OT #17345 RT: 43.57 AV: 1 NL: 3.05E4
T: FTMS + c NSI d Full ms2 762.3614@hcd28.00 [140.0000-1535.0000]
RVSAIVEQSW#NDS
b4+
414.25
b5+
527.33
b6+
626.40
b7+
755.44
b8+
883.50
b9+
970.53
b10+
1188.60
152.06
159.08
230.08
245.08
288.12
359.16
398.14
430.13
517.17
535.18
570.59
598.40
640.22
700.83
712.85
730.68
761.84
795.29
863.36
908.45
942.53
1006.43
1075.47
1093.47
1170.59
1248.45
1303.64
1375.56
1449.67
1514.02
m/z

# 10

- Experimental mass: 117.9126
- Unimod information:
- Possible composition: Phosphorylation + Cation:K, [H(1)O(3)P(1) + H(-1) K]
- Sites in Unimod:
- Sites observed:
- Features: phosphorylation sites determined, cation modified sites are not observed

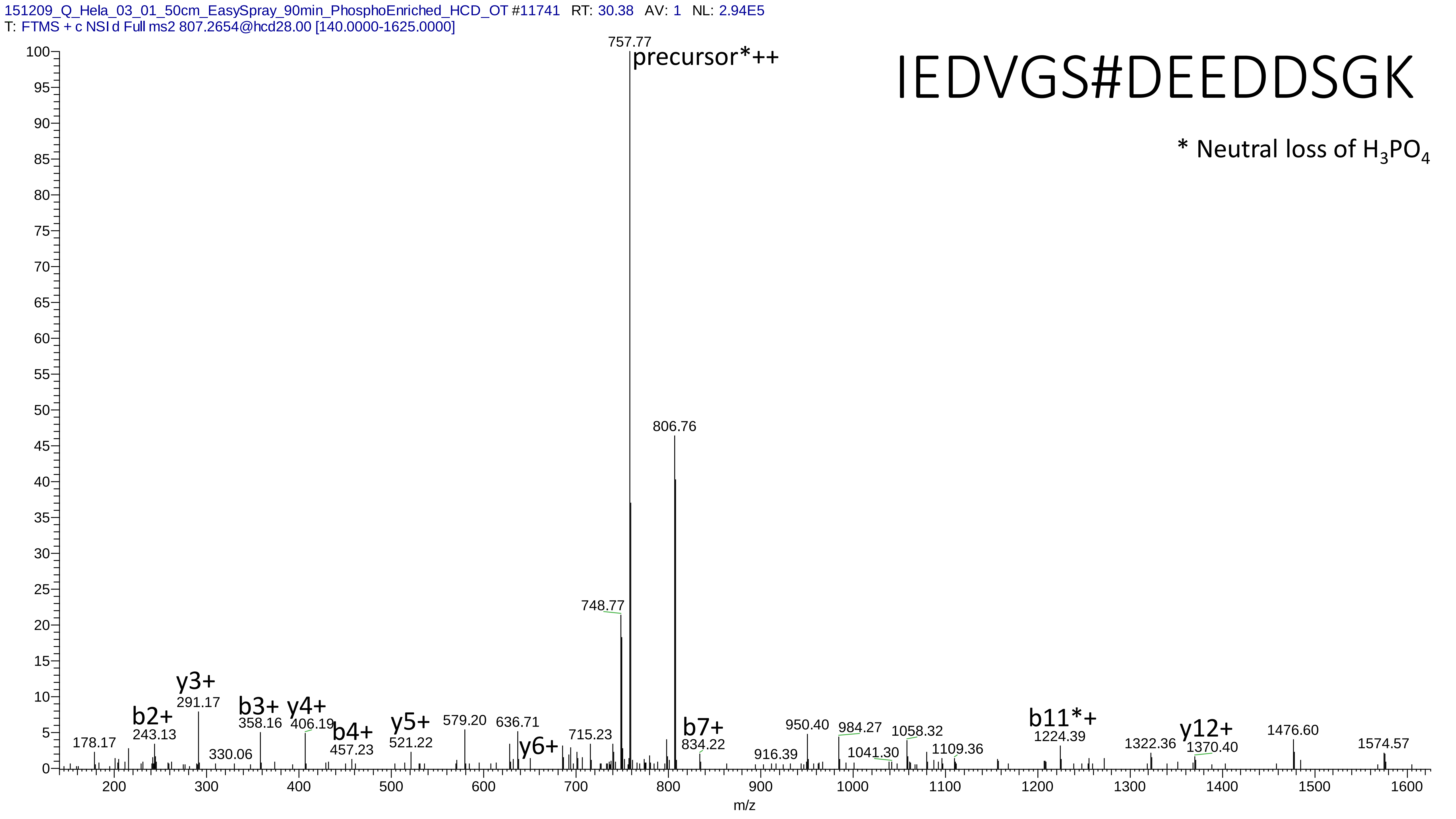
151209_Q_Hela_03_01_50cm_EasySpray_90min_PhosphoEnriched_HCD_OT #11741 RT: 30.38 AV: 1 NL: 2.94E5
T: FTMS + c NSI d Full ms2 807.2654@hcd28.00 [140.0000-1625.0000]
IEDVGS#DEEDDSGK
* Neutral loss of H3PO4
precursor*++
757.77
806.76
748.77
y3+
291.17
b2+
243.13
b3+
358.16
y4+
406.19
b4+
457.23
y5+
521.22
579.20
636.71
y6+
715.23
b7+
834.22
178.17
330.06
916.39
950.40
984.27
1041.30
1058.32
1109.36
b11*+
1224.39
1322.36
y12+
1370.40
1476.60
1574.57
m/z

# 11

- Experimental mass: 17.0254
- Unimod information: Ammonium
- Possible composition: H(3)N(1)
- Sites in Unimod: E;D;c*
- Sites observed: D

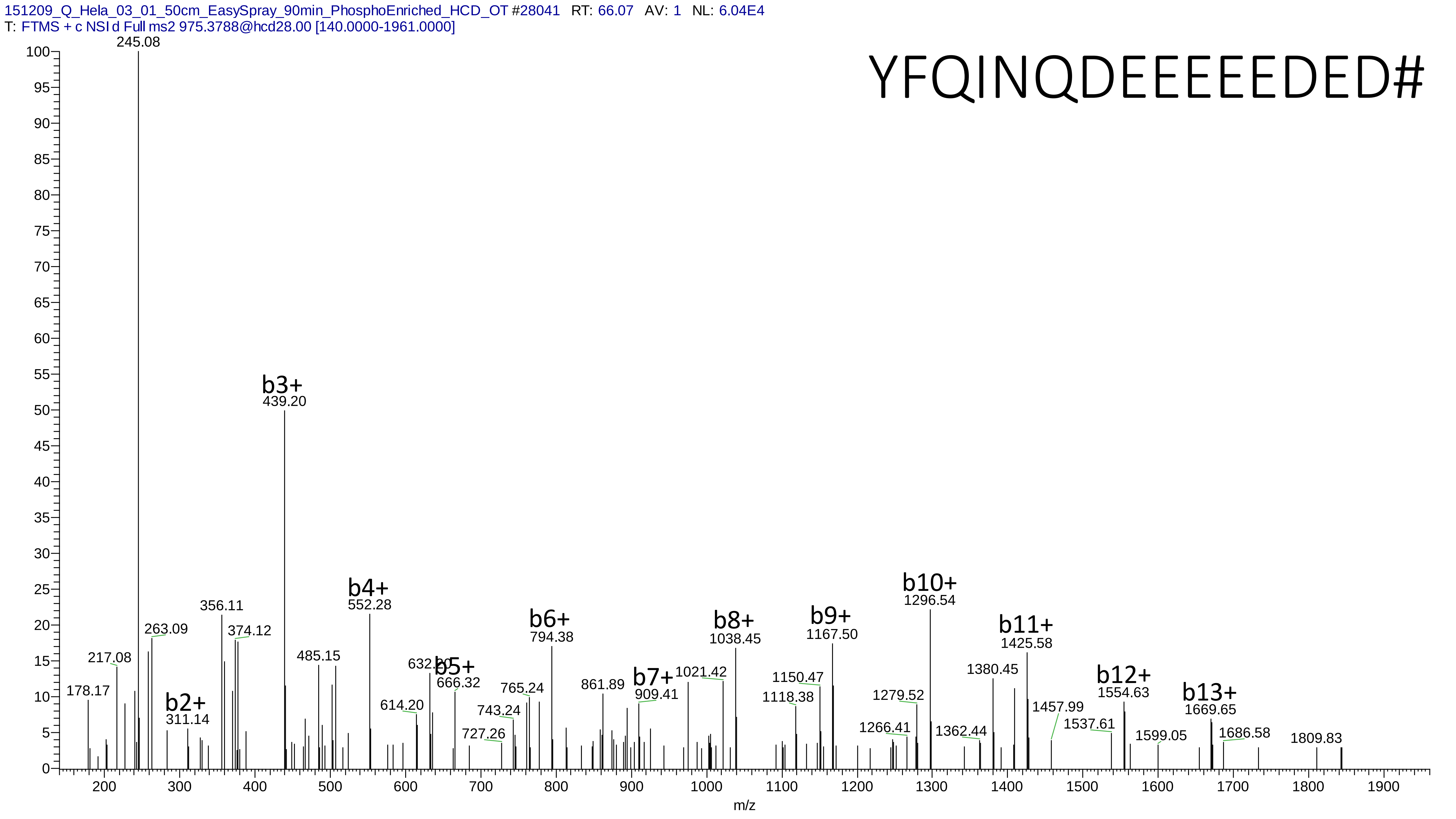
151209_Q_Hela_03_01_50cm_EasySpray_90min_PhosphoEnriched_HCD_OT #28041 RT: 66.07 AV: 1 NL: 6.04E4
T: FTMS + c NSI d Full ms2 975.3788@hcd28.00 [140.0000-1961.0000]
YFQINQDEEEEEDED#
245.08
b2+
311.14
b3+
439.20
b4+
552.28
b5+
666.32
b6+
794.38
b7+
909.41
b8+
1038.45
b9+
1167.50
b10+
1296.54
b11+
1425.58
b12+
1554.63
b13+
1669.65
178.17
217.08
263.09
356.11
374.12
485.15
614.20
632.20
727.26
743.24
765.24
861.89
1021.42
1118.38
1150.47
1266.41
1279.52
1362.44
1380.45
1457.99
1537.61
1599.05
1686.58
1809.83
m/z

# 12

- Experimental mass: 132.8786
- Unimod information:
- Possible composition: Phosphorylation + Cation:Fe[III], [H(1)O(3)P(1) + H(-3) Fe]
- Sites in Unimod:
- Sites observed:
- Features: phosphorylation sites determined, cation modified sites are not observed

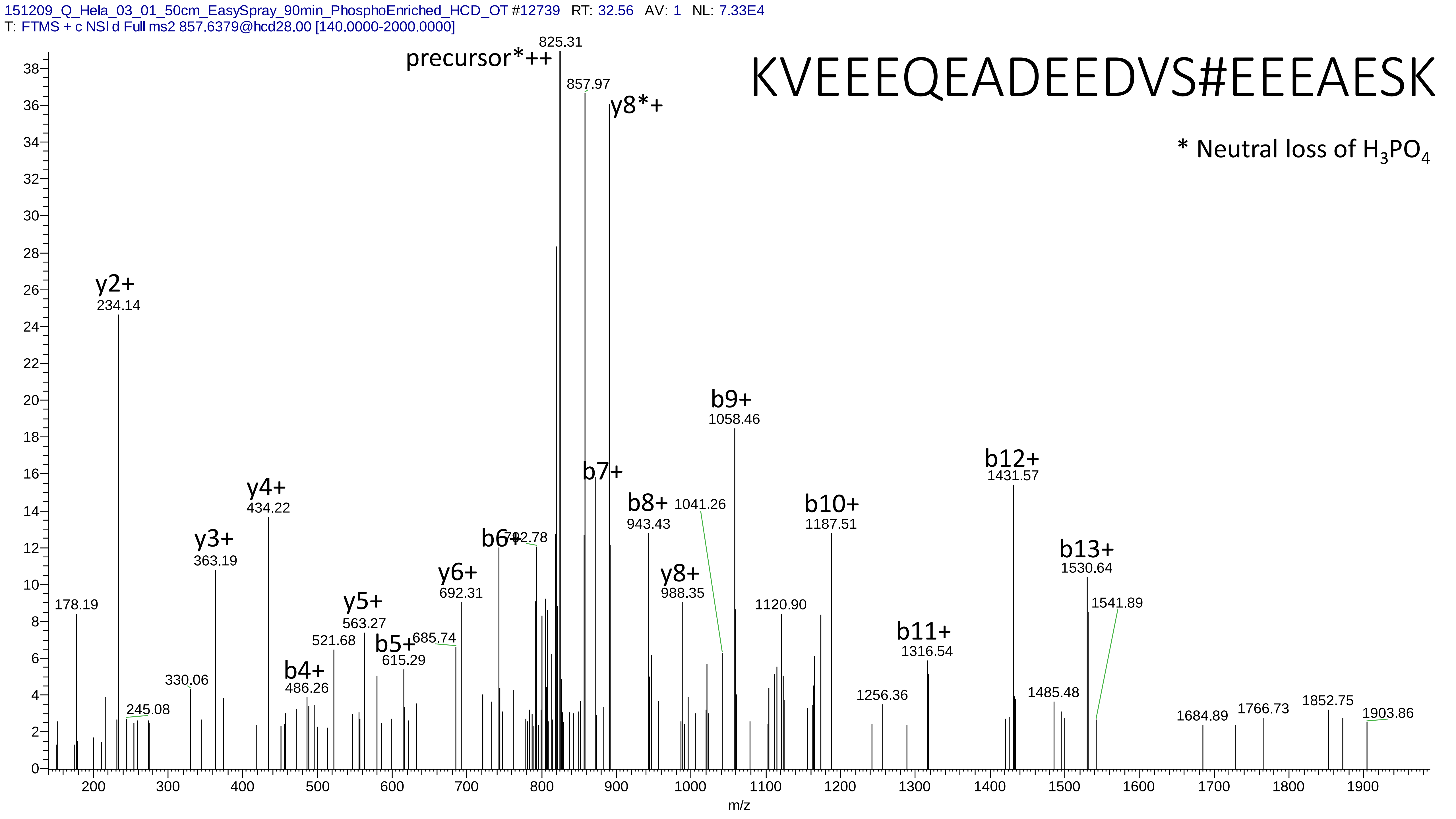
151209_Q_Hela_03_01_50cm_EasySpray_90min_PhosphoEnriched_HCD_OT #12739 RT: 32.56 AV: 1 NL: 7.33E4
T: FTMS + c NSI d Full ms2 857.6379@hcd28.00 [140.0000-2000.0000]
KVEEEQEADEEDVS#EEEAESK
* Neutral loss of H3PO4
precursor*++
825.31
857.97
y8*+
y2+
234.14
y3+
363.19
y4+
434.22
b4+
486.26
y5+
563.27
b5+
615.29
y6+
692.31
b6+
742.78
b7+
b8+
943.43
y8+
988.35
1041.26
b9+
1058.46
1120.90
b10+
1187.51
b11+
1316.54
b12+
1431.57
b13+
1530.64
1541.89
178.19
245.08
330.06
521.68
685.74
1256.36
1485.48
1684.89
1766.73
1852.75
1903.86
m/z

# 13

- Experimental mass: 95.9602
- Unimod information:
- Possible composition: Phosphorylation + Oxidation, [H(1)O(3)P(1) + O]
- Sites in Unimod:
- Sites observed:

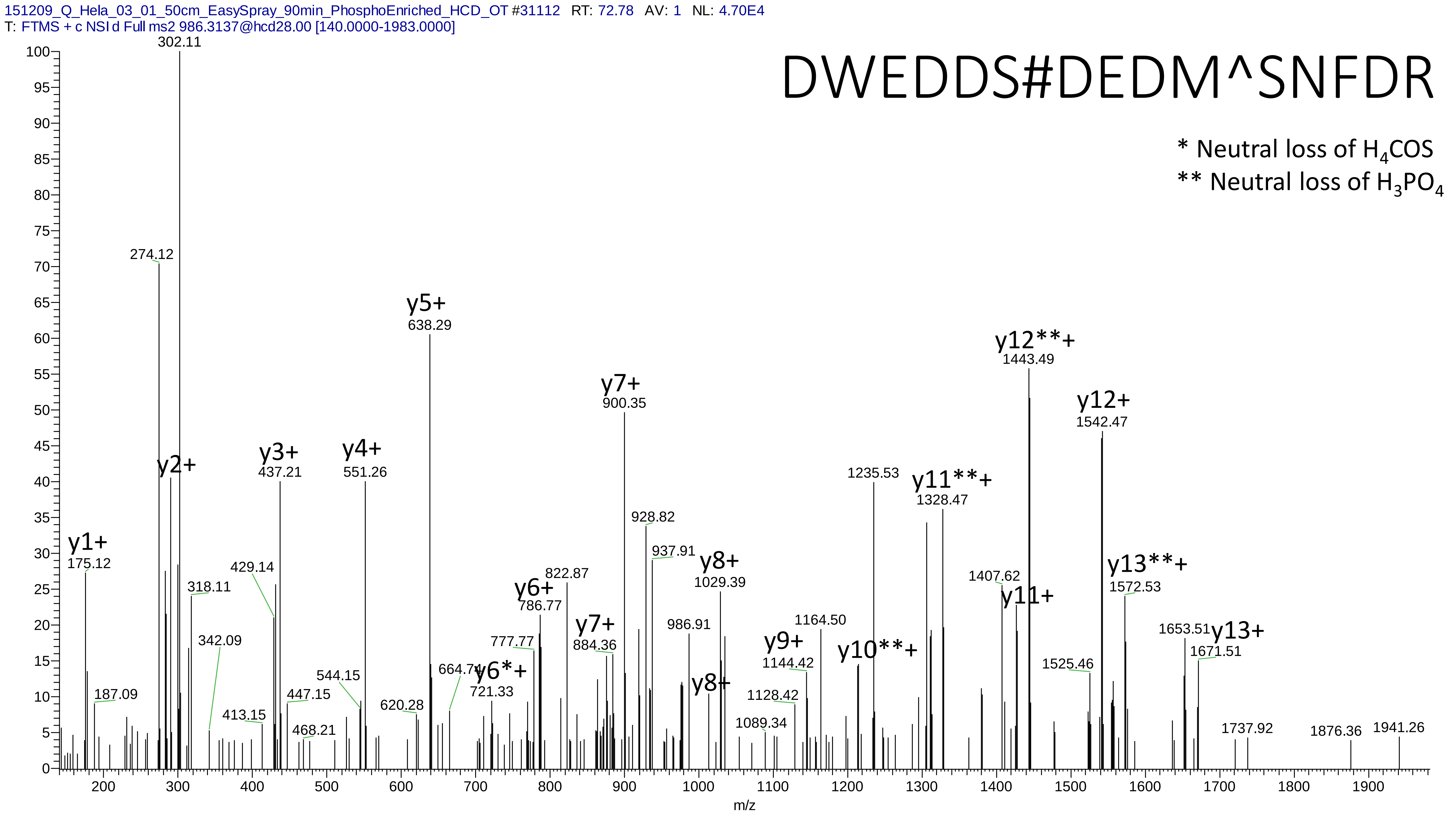
151209_Q_Hela_03_01_50cm_EasySpray_90min_PhosphoEnriched_HCD_OT #31112 RT: 72.78 AV: 1 NL: 4.70E4
T: FTMS + c NSI d Full ms2 986.3137@hcd28.00 [140.0000-1983.0000]
DWEDDS#DEDM^SNFDR
* Neutral loss of H4COS
** Neutral loss of H3PO4
y1+ 175.12
187.09
274.12
y2+
302.11
318.11
342.09
413.15
429.14
y3+ 437.21
447.15
468.21
544.15
y4+ 551.26
620.28
y5+ 638.29
664.74
y6*+ 721.33
777.77
y6+ 786.77
822.87
y7+ 884.36
y7+ 900.35
928.82
937.91
986.91
y8+
y8+ 1029.39
1089.34
1128.42
y9+ 1144.42
1164.50
y10**+
1235.53
y11**+ 1328.47
1407.62
y11+
y12**+ 1443.49
1525.46
y12+ 1542.47
y13**+ 1572.53
1653.51
y13+ 1671.51
1737.92
1876.36
1941.26
m/z

# 14

- Experimental mass: -48.1289
- Unimod information:
- Possible composition: Phosphorylation + Lys-loss, [H(1)O(3)P(1) + H(-12) C(-6) N(-2) O(-1)]
- Sites in Unimod:
- Sites observed:
- Features: 98.1% (52/53) peptides contain Lys at peptide n-term

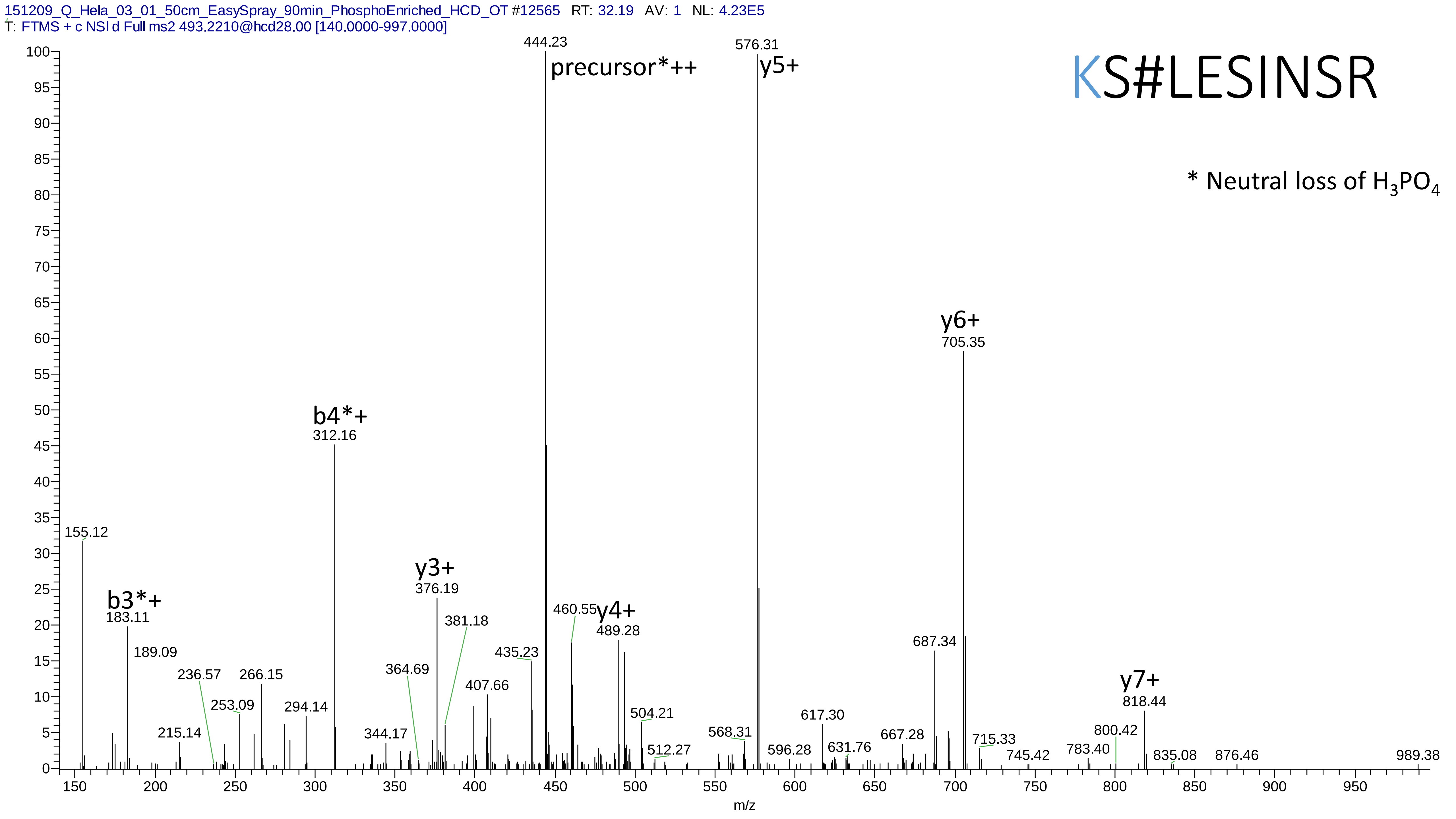
151209_Q_Hela_03_01_50cm_EasySpray_90min_PhosphoEnriched_HCD_OT #12565 RT: 32.19 AV: 1 NL: 4.23E5
T: FTMS + c NSI d Full ms2 493.2210@hcd28.00 [140.0000-997.0000]
KS#LESINSR
* Neutral loss of $H_3PO_4$
precursor*++
444.23
y5+
576.31
y6+
705.35
b4*+
312.16
y3+
376.19
y4+
489.28
b3*+
183.11
y7+
818.44
155.12
189.09
215.14
236.57
253.09
266.15
294.14
344.17
364.69
381.18
407.66
435.23
460.55
504.21
512.27
568.31
596.28
617.30
631.76
667.28
687.34
715.33
745.42
783.40
800.42
835.08
876.46
989.38
m/z

# 15

- Experimental mass: -76.1349
- Unimod information:
- Possible composition: Phosphorylation + Arg-loss, [H(1)O(3)P(1) + H(-12) C(-6) N(-4) O(-1)]
- Sites in Unimod:
- Sites observed:
- Features:  94.9% (56/59) peptides contain Arg at peptide n-term

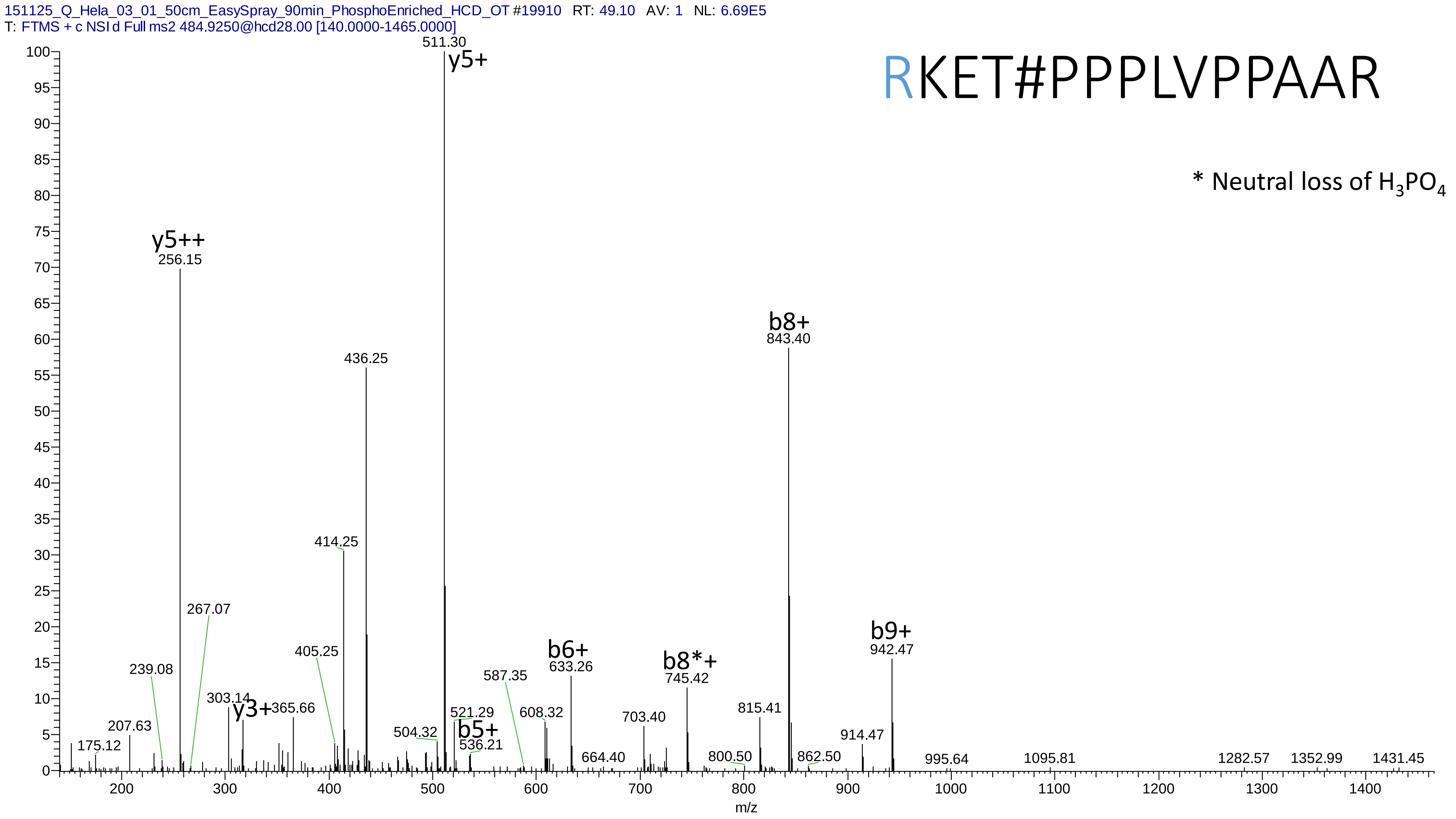
151125_Q_Hela_03_01_50cm_EasySpray_90min_PhosphoEnriched_HCD_OT #19910 RT: 49.10 AV: 1 NL: 6.69E5
T: FTMS + c NSI d Full ms2 484.9250@hcd28.00 [140.0000-1465.0000]
RKET#PPPLVPPAAR
* Neutral loss of H3PO4
y5+
511.30
y5++
256.15
436.25
414.25
267.07
405.25
239.08
207.63
175.12
303.14
y3+
365.66
504.32
521.29
b5+
536.21
587.35
608.32
b6+
633.26
664.40
703.40
b8*+
745.42
800.50
815.41
b8+
843.40
862.50
914.47
b9+
942.47
995.64
1095.81
1282.57
1352.99
1431.45
m/z

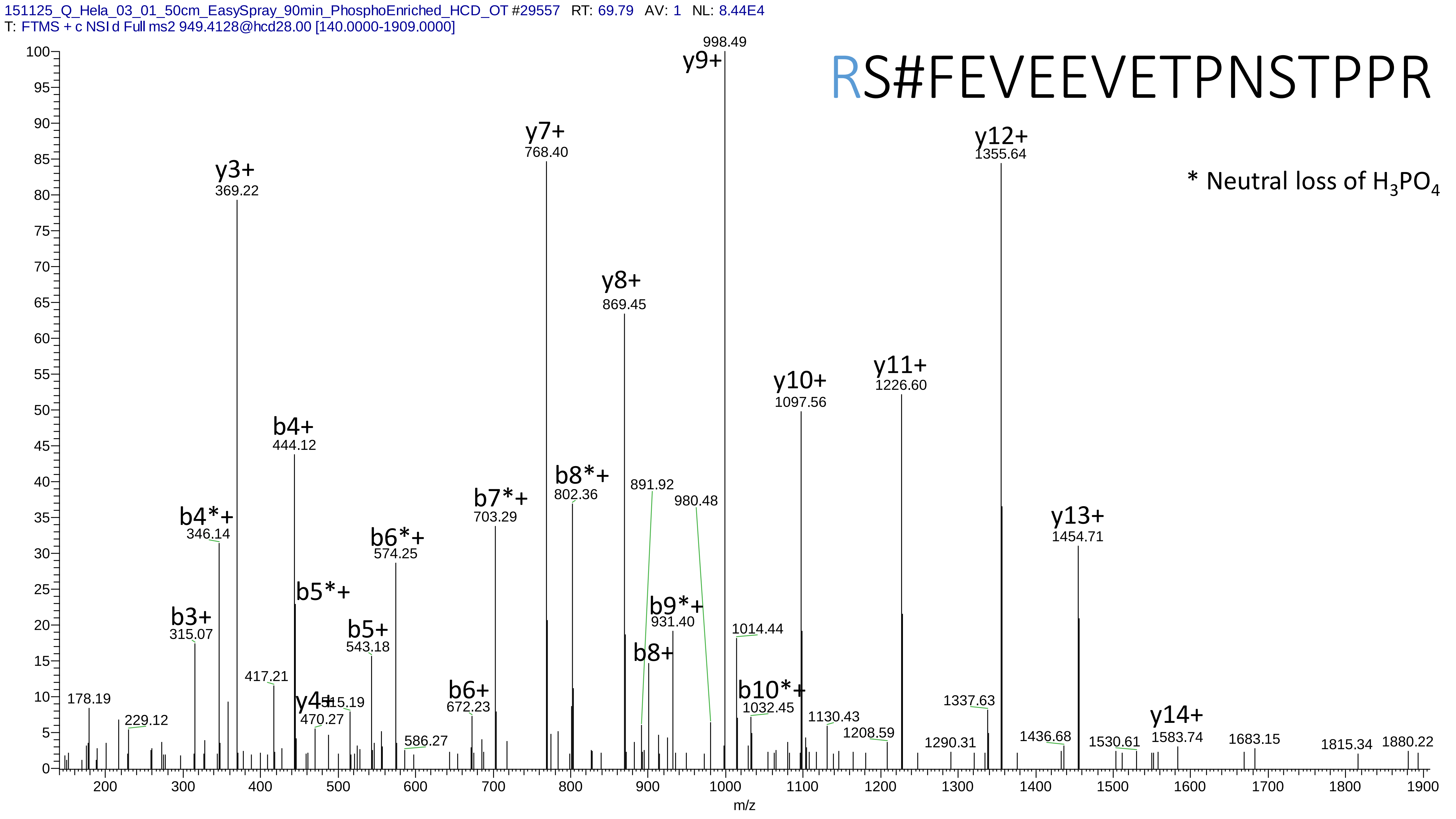

151125_Q_Hela_03_01_50cm_EasySpray_90min_PhosphoEnriched_HCD_OT #29557 RT: 69.79 AV: 1 NL: 8.44E4
T: FTMS + c NSI d Full ms2 949.4128@hcd28.00 [140.0000-1909.0000]
RS#FEVEEVETPNSTPPR
* Neutral loss of $H_3PO_4$
y9+ 998.49
y7+ 768.40
y12+ 1355.64
y3+ 369.22
y8+ 869.45
y11+ 1226.60
y10+ 1097.56
b4+ 444.12
b8*+ 802.36
891.92
980.48
b7*+ 703.29
y13+ 1454.71
b4*+ 346.14
b6*+ 574.25
b5*+
b9*+ 931.40
b3+ 315.07
b5+ 543.18
1014.44
b8+
417.21
b10*+ 1032.45
b6+ 672.23
178.19
y4+ 470.27
515.19
1337.63
229.12
1130.43
y14+ 1583.74
586.27
1208.59
1290.31
1436.68
1530.61
1683.15
1815.34
1880.22
m/z

# 16

- Experimental mass: 43.006
- Unimod information: Carbamyl
- Possible composition: H(1)C(1)N(1)O(1)
- Sites in Unimod: Y;T;S;M;C;R;n*;K;[*
- Sites observed: K;R

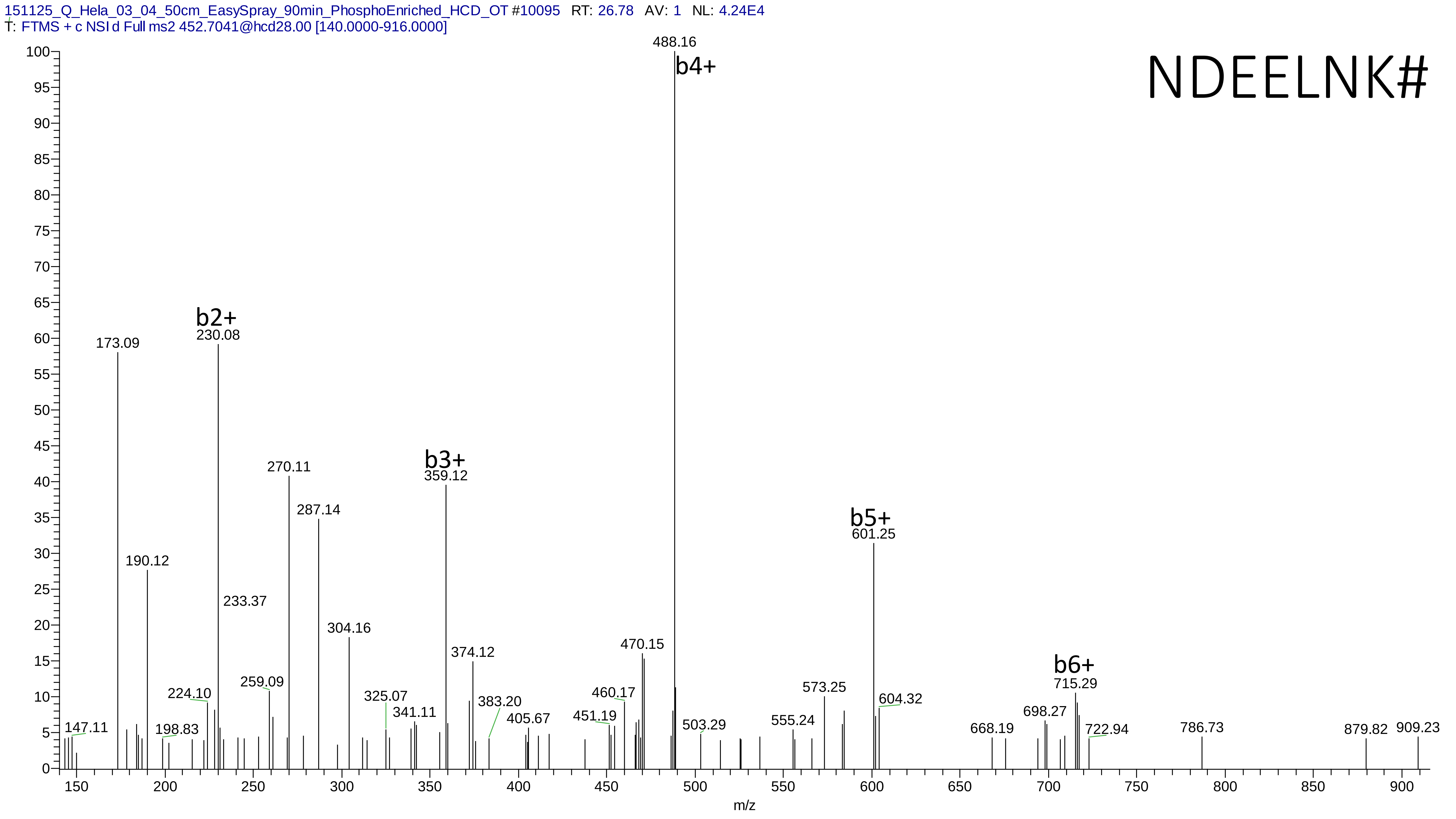
151125_Q_Hela_03_04_50cm_EasySpray_90min_PhosphoEnriched_HCD_OT #10095 RT: 26.78 AV: 1 NL: 4.24E4
T: FTMS + c NSI d Full ms2 452.7041@hcd28.00 [140.0000-916.0000]
NDEELNK#
b2+
b3+
b4+
b5+
b6+
488.16
173.09
230.08
270.11
359.12
287.14
601.25
190.12
233.37
304.16
374.12
470.15
224.10
259.09
325.07
341.11
383.20
405.67
451.19
460.17
503.29
555.24
573.25
604.32
668.19
698.27
715.29
722.94
786.73
879.82
909.23
147.11
198.83
m/z

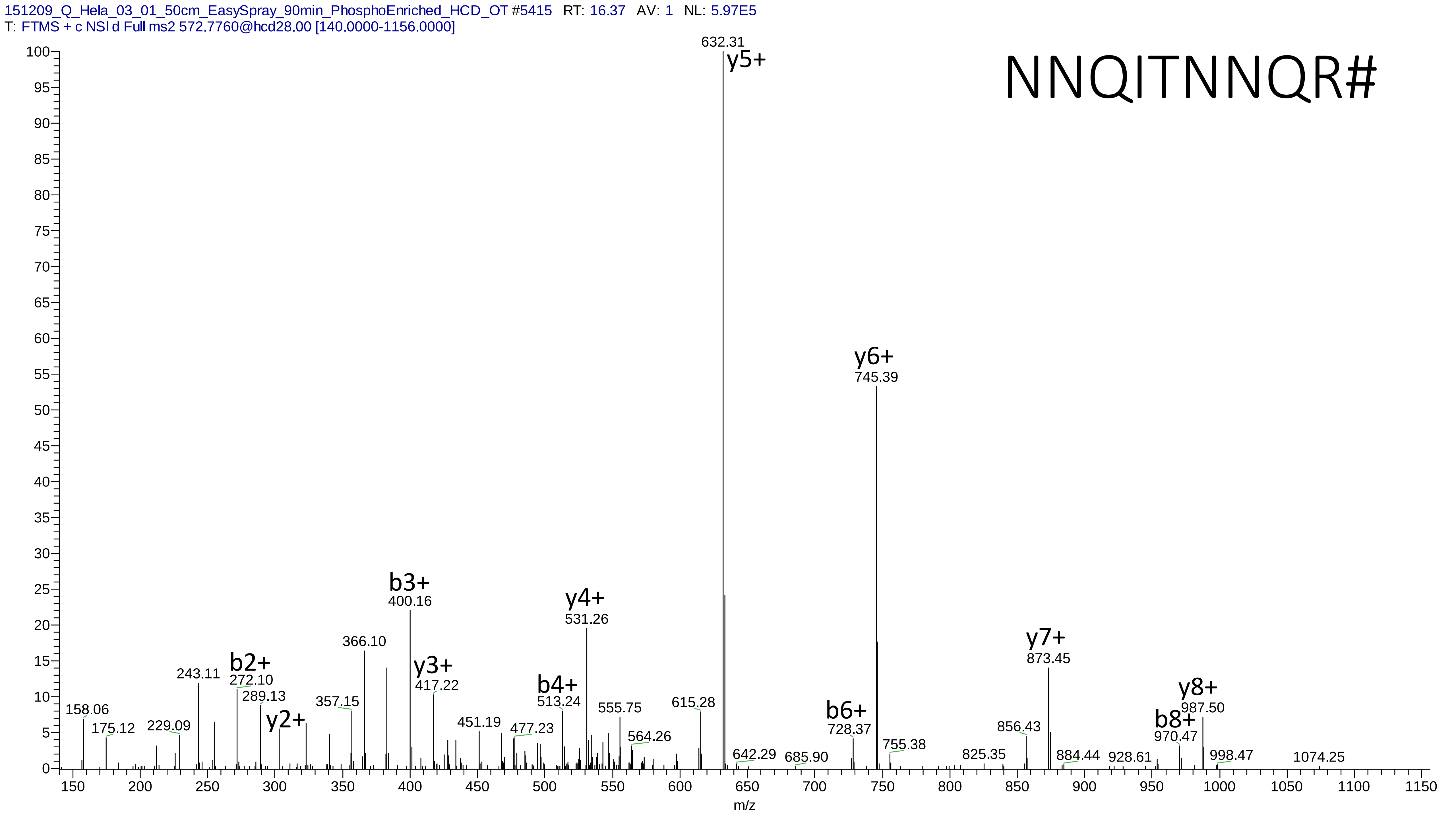
151209_Q_Hela_03_01_50cm_EasySpray_90min_PhosphoEnriched_HCD_OT #5415 RT: 16.37 AV: 1 NL: 5.97E5
T: FTMS + c NSI d Full ms2 572.7760@hcd28.00 [140.0000-1156.0000]
NNQITNNQR#
632.31
y5+
y6+
745.39
b3+
400.16
y4+
531.26
366.10
y7+
873.45
b2+
272.10
243.11
y3+
417.22
289.13
b4+
513.24
y8+
987.50
158.06
357.15
y2+
615.28
555.75
b6+
b8+
728.37
970.47
229.09
175.12
451.19
477.23
564.26
856.43
755.38
642.29
685.90
825.35
884.44
928.61
998.47
1074.25
m/z

# 17

- Experimental mass: -9.0637
- Unimod information:
- Possible composition: Phosphorylation + Met-loss+Acetyl, [H(1)O(3)P(1) + H(-7) C(-3) N(-1) S(-1)]
- Sites in Unimod:
- Sites observed:
- Features: 97.0% (32/33) peptides contain Met at peptide n-term;

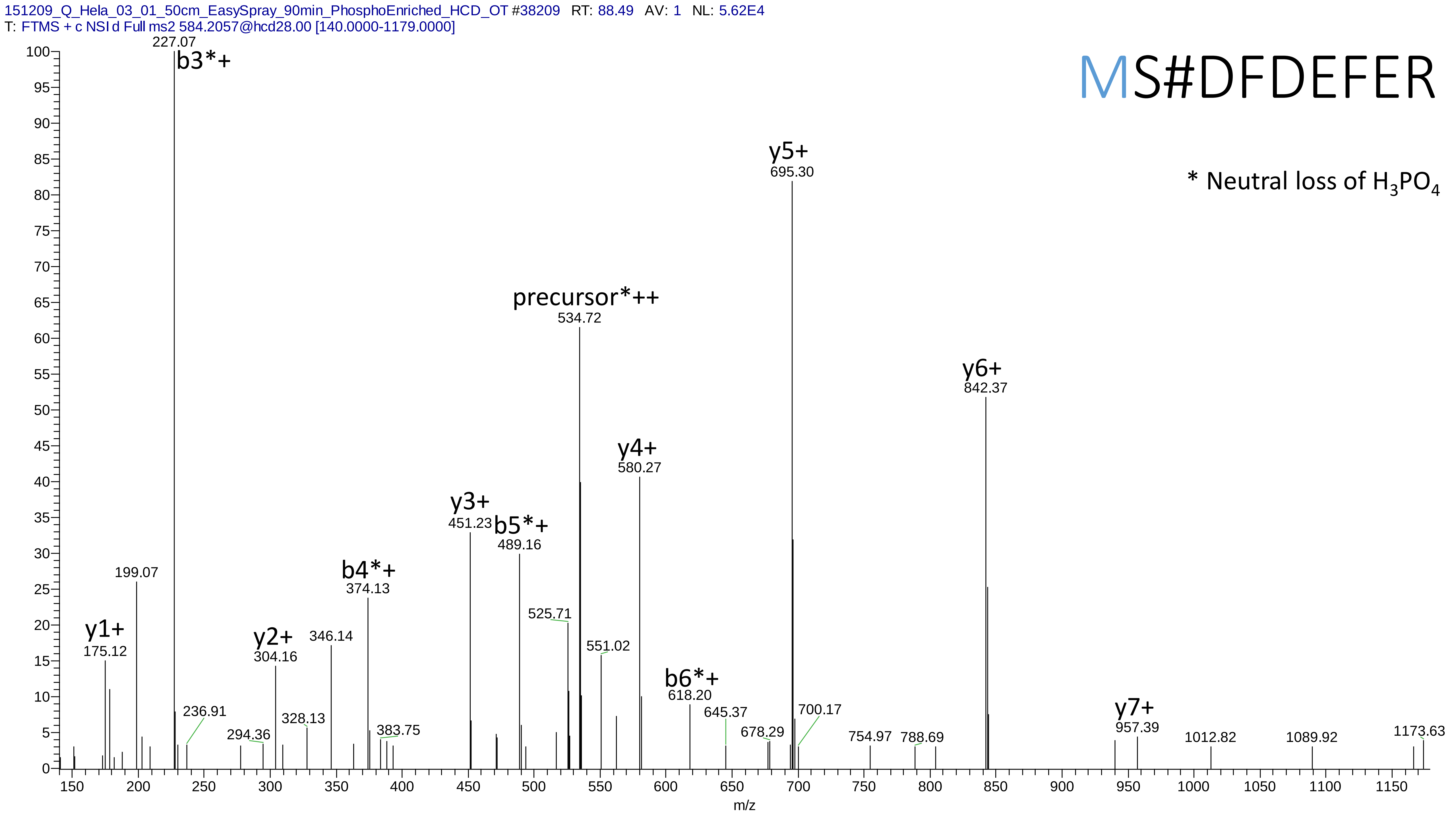
151209_Q_Hela_03_01_50cm_EasySpray_90min_PhosphoEnriched_HCD_OT #38209 RT: 88.49 AV: 1 NL: 5.62E4
T: FTMS + c NSI d Full ms2 584.2057@hcd28.00 [140.0000-1179.0000]
MS#DFDEFER
* Neutral loss of H3PO4
b3*+ 227.07
y5+ 695.30
precursor*++ 534.72
y6+ 842.37
y4+ 580.27
y3+ 451.23
b5*+ 489.16
b4*+ 374.13
199.07
525.71
y1+ 175.12
y2+ 304.16
346.14
551.02
b6*+ 618.20
236.91
294.36
328.13
383.75
645.37
678.29
700.17
754.97
788.69
y7+ 957.39
1012.82
1089.92
1173.63
m/z

# 18

- Experimental mass: 111.9548
- Unimod information:
- Possible composition:Phosphorylation + Dioxidation, [H(1)O(3)P(1) + O(2)]
- Sites in Unimod:
- Sites observed:

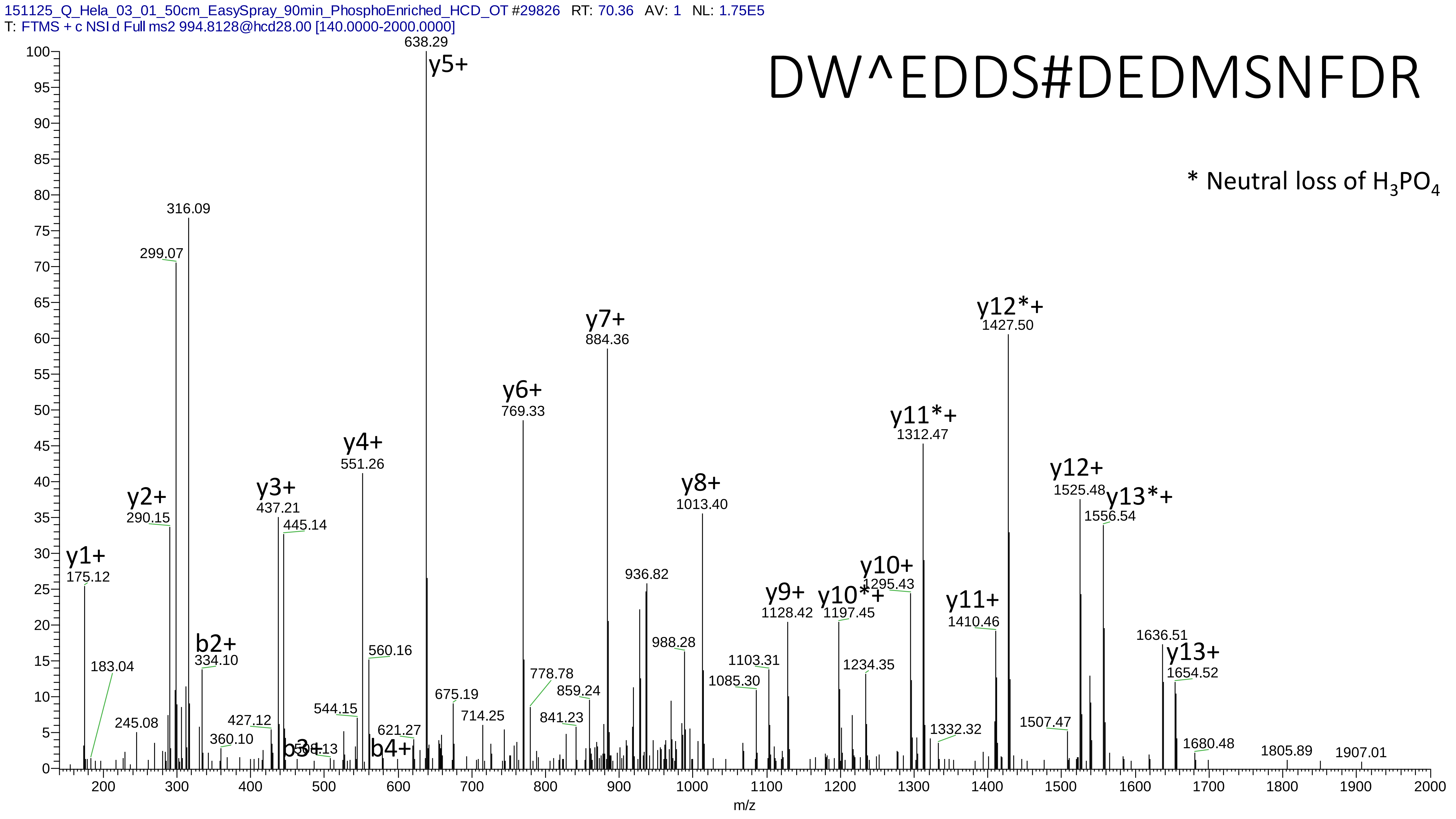

151125_Q_Hela_03_01_50cm_EasySpray_90min_PhosphoEnriched_HCD_OT #29826 RT: 70.36 AV: 1 NL: 1.75E5
T: FTMS + c NSI d Full ms2 994.8128@hcd28.00 [140.0000-2000.0000]
DW^EDDS#DEDMSNFDR
* Neutral loss of $H_3PO_4$
y1+ 175.12
183.04
245.08
y2+ 290.15
299.07
316.09
b2+ 334.10
360.10
427.12
y3+ 437.21
445.14
b3+
544.15
y4+ 551.26
560.16
621.27
b4+
y5+ 638.29
675.19
714.25
y6+ 769.33
778.78
841.23
859.24
y7+ 884.36
936.82
988.28
y8+ 1013.40
1085.30
1103.31
y9+ 1128.42
y10*+ 1197.45
1234.35
y10+ 1295.43
y11*+ 1312.47
1332.32
y11+ 1410.46
y12*+ 1427.50
1507.47
y12+ 1525.48
y13*+ 1556.54
1636.51
y13+ 1654.52
1680.48
1805.89
1907.01
m/z

# 19

- Experimental mass: 61.9549
- Unimod information:
- Possible composition: Phosphorylation + Glu->pyro-Glu, [H(1)O(3)P(1) + H(-2) O(-1)]
- Sites in Unimod:
- Sites observed:
- Features: 100.0% (30/30) peptides contain Glu, 70.0% (21/30) peptides contain Glu at peptide n-term

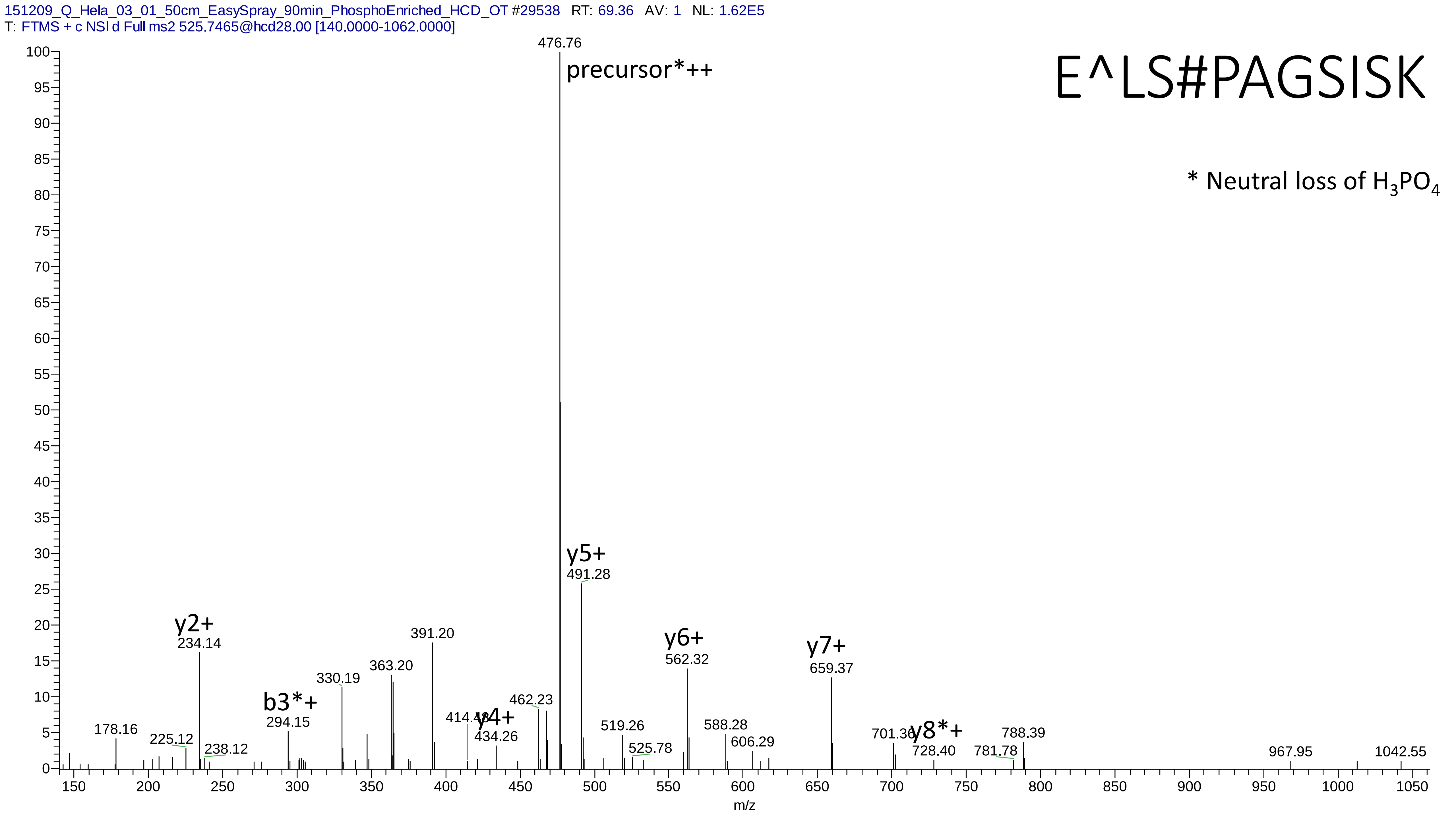

151209_Q_Hela_03_01_50cm_EasySpray_90min_PhosphoEnriched_HCD_OT #29538 RT: 69.36 AV: 1 NL: 1.62E5
T: FTMS + c NSI d Full ms2 525.7465@hcd28.00 [140.0000-1062.0000]
E^LS#PAGSISK
* Neutral loss of H3PO4
476.76
precursor*++
y5+
491.28
y2+
234.14
y6+
562.32
y7+
659.37
391.20
363.20
330.19
b3*+
294.15
462.23
414.48
y4+
434.26
178.16
225.12
238.12
519.26
525.78
588.28
606.29
701.36
y8*+
728.40
781.78
788.39
967.95
1042.55
m/z

# 20

- Experimental mass: 18.0097
- Unimod information:
- Possible composition: Deamidated + Ammonium, [H(-1) N(-1) O + H(3) N]
- Sites in Unimod:
- Sites observed:

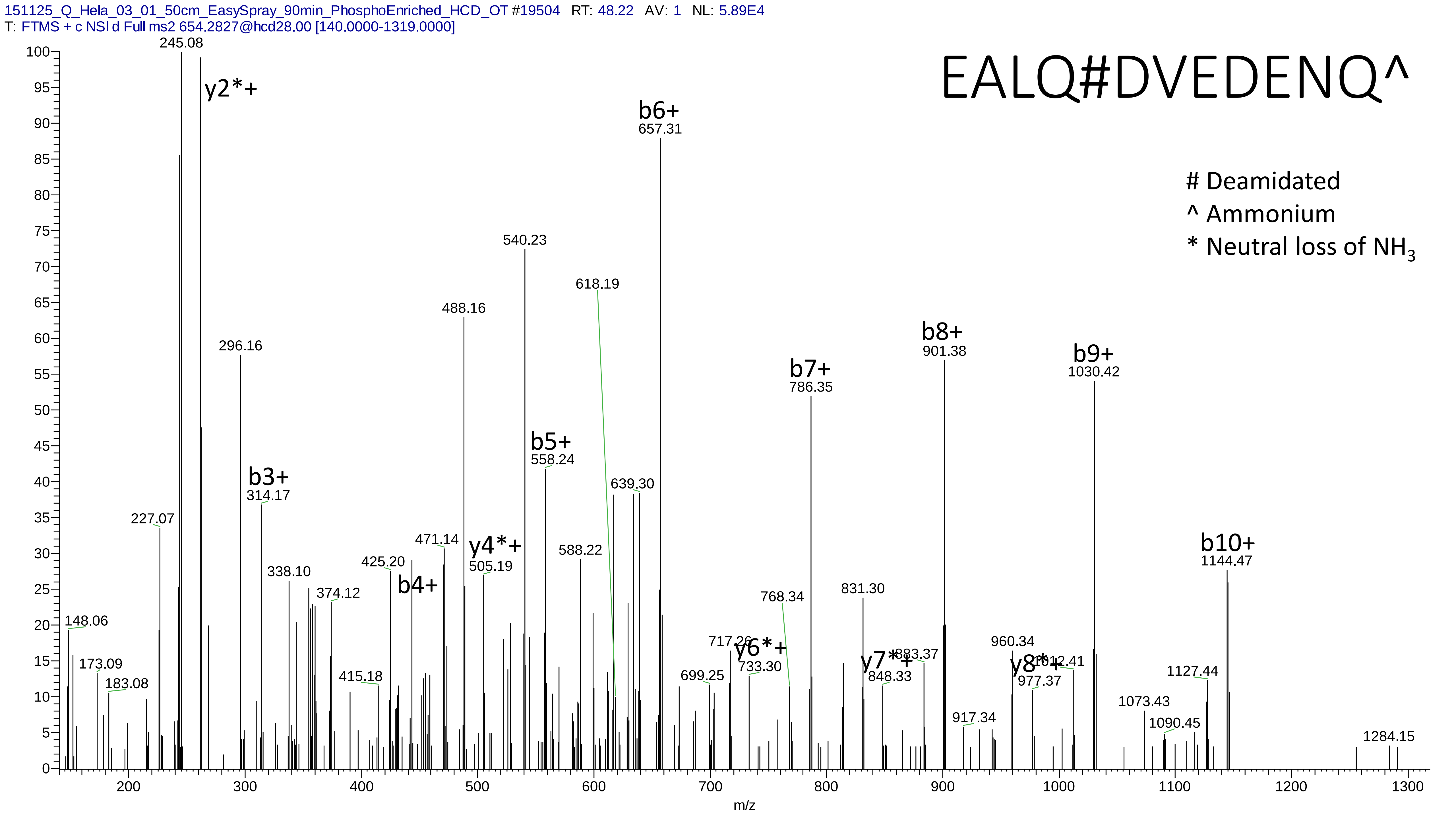
151125_Q_Hela_03_01_50cm_EasySpray_90min_PhosphoEnriched_HCD_OT #19504 RT: 48.22 AV: 1 NL: 5.89E4
T: FTMS + c NSI d Full ms2 654.2827@hcd28.00 [140.0000-1319.0000]
EALQ#DVEDENQ^
### Deamidated
^ Ammonium
* Neutral loss of NH3
y2*+
b3+
b4+
y4*+
b5+
b6+
y6*+
b7+
y7*+
b8+
y8*+
b9+
b10+
245.08
296.16
314.17
488.16
540.23
558.24
618.19
639.30
657.31
786.35
901.38
1030.42
1144.47
m/z

# 21

- Experimental mass: 63.9699
- Unimod information: SulfurDioxide
- Possible composition: Phosphorylation – Oxidation, [H(1)O(3)P(1) - O]
- Sites in Unimod:
- Sites observed:
- Features: MSFragger assigned an incorrect oxidation to each peptide so the rest mass difference is 63.9697, actual mass difference will be 79.9646 (phosphorylation)

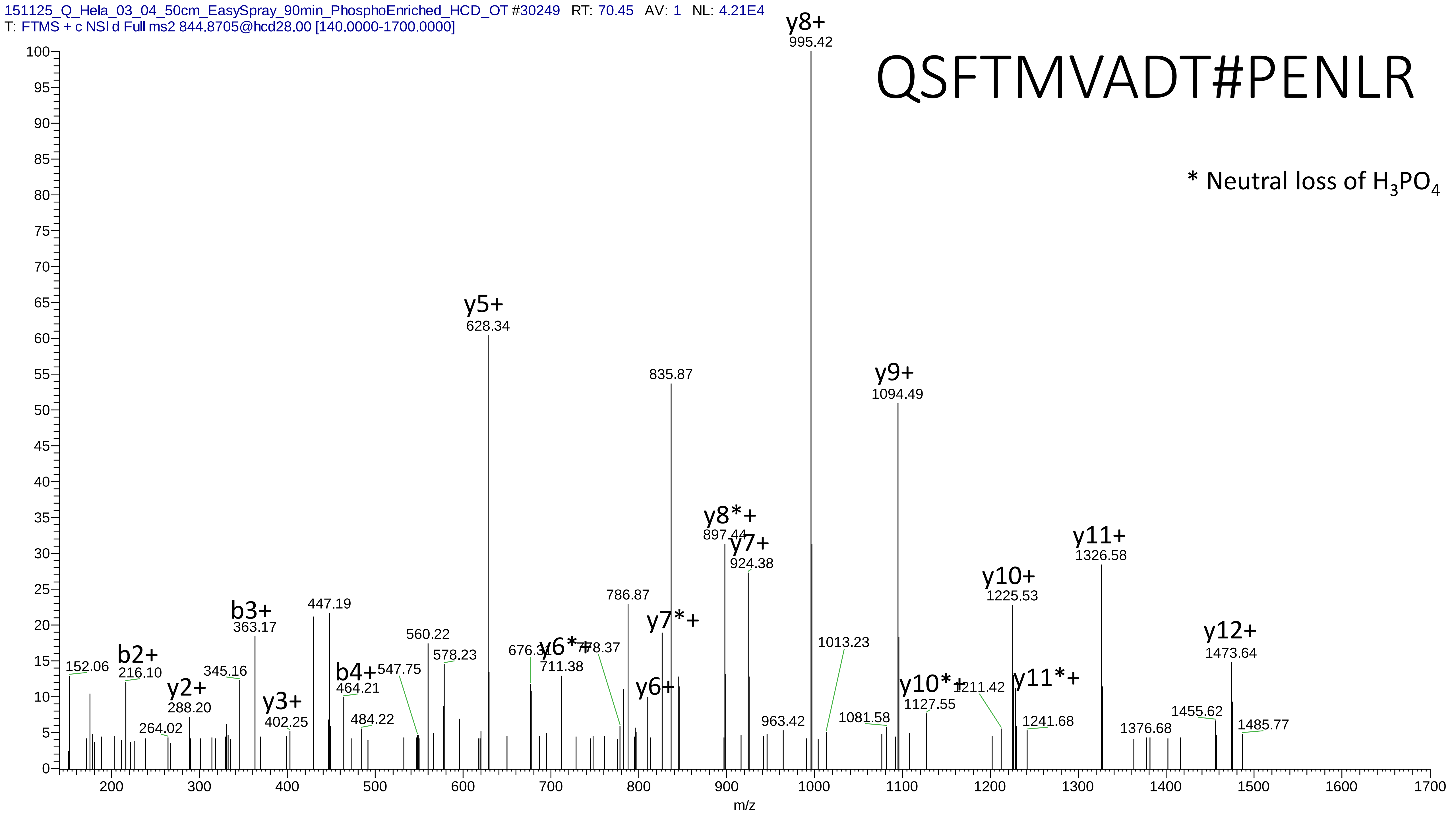
151125_Q_Hela_03_04_50cm_EasySpray_90min_PhosphoEnriched_HCD_OT #30249 RT: 70.45 AV: 1 NL: 4.21E4
T: FTMS + c NSI d Full ms2 844.8705@hcd28.00 [140.0000-1700.0000]
QSFTMVADT#PENLR
* Neutral loss of H3PO4
y8+ 995.42
y5+ 628.34
835.87
y9+ 1094.49
y8*+ 897.44
y7+ 924.38
y11+ 1326.58
y10+ 1225.53
786.87
447.19
b3+ 363.17
y7*+
560.22
y12+ 1473.64
578.23
1013.23
676.31
y6*+ 778.37
711.38
152.06
b2+ 216.10
345.16
b4+ 547.75
464.21
y10*+ 1211.42
y11*+
1127.55
y2+ 288.20
y6+
y3+ 402.25
484.22
963.42
1081.58
1241.68
1455.62
1485.77
264.02
1376.68
m/z

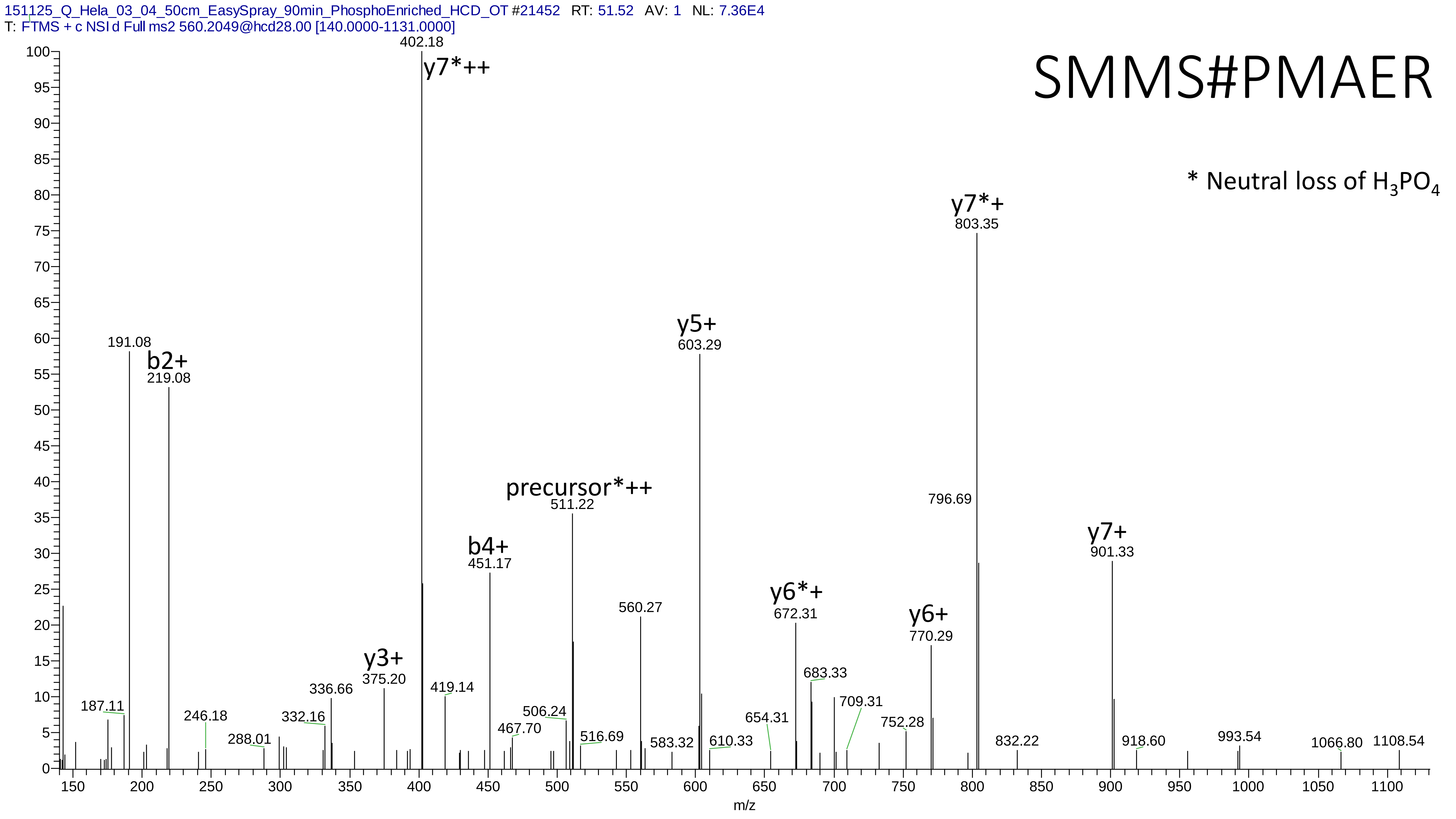
151125_Q_Hela_03_04_50cm_EasySpray_90min_PhosphoEnriched_HCD_OT #21452 RT: 51.52 AV: 1 NL: 7.36E4
T: FTMS + c NSI d Full ms2 560.2049@hcd28.00 [140.0000-1131.0000]
SMMS#PMAER
* Neutral loss of $H_3PO_4$
402.18
y7*++
191.08
b2+
219.08
y7*+
803.35
y5+
603.29
precursor*++
511.22
b4+
451.17
796.69
y7+
901.33
560.27
y6*+
672.31
y6+
770.29
y3+
375.20
683.33
709.31
336.66
419.14
187.11
246.18
332.16
506.24
288.01
467.70
516.69
583.32
610.33
654.31
752.28
832.22
918.60
993.54
1066.80
1108.54
m/z

# 22

- Experimental mass: 81.9330
- Unimod information:
- Possible composition: Phosphorylation + Deamidation * 2, [H(1)O(3)P(1) + H(-1) N(-1) O + H(-1) N(-1) O]
- Sites in Unimod:
- Sites observed:

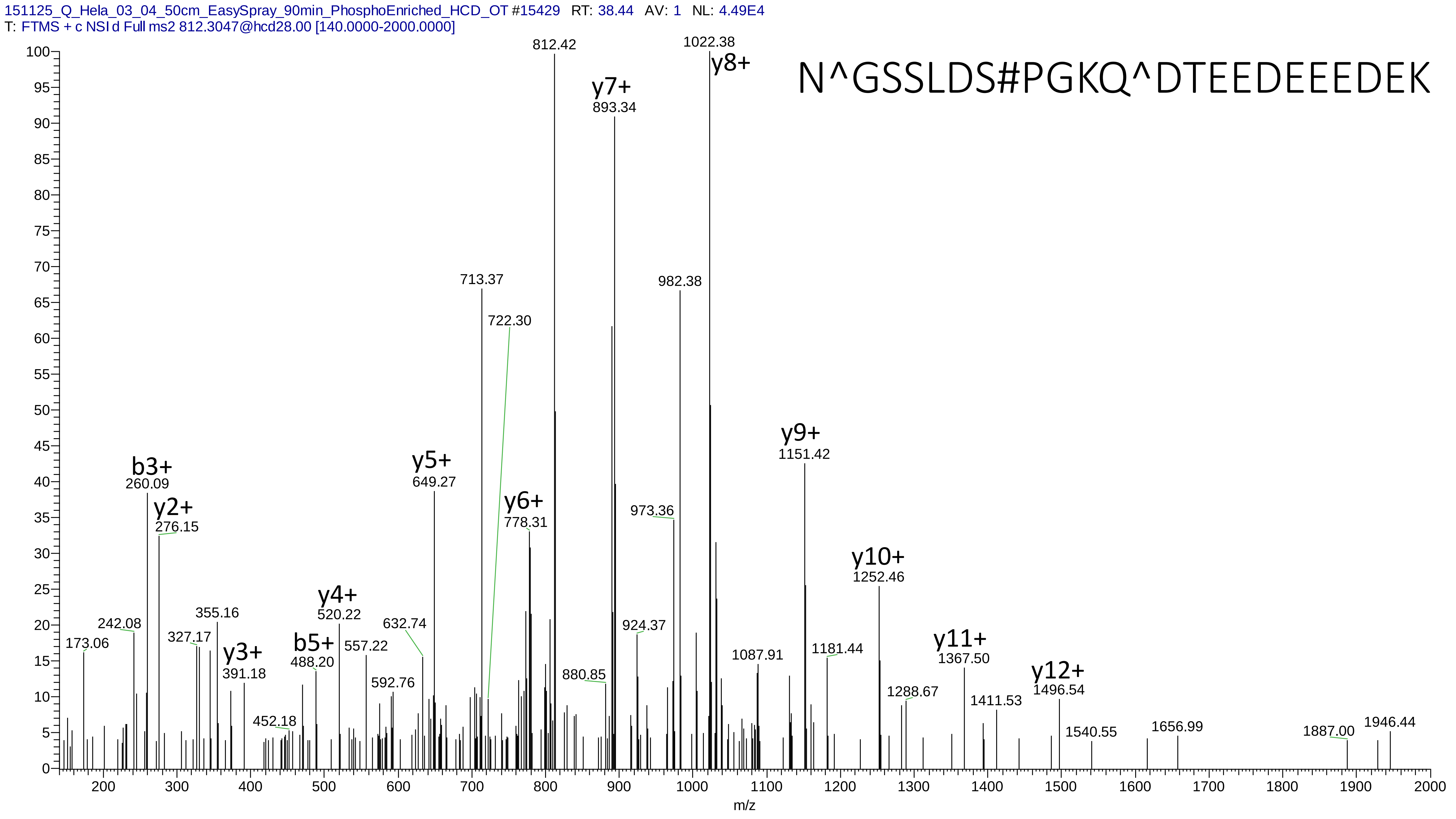
151125_Q_Hela_03_04_50cm_EasySpray_90min_PhosphoEnriched_HCD_OT #15429 RT: 38.44 AV: 1 NL: 4.49E4
T: FTMS + c NSI d Full ms2 812.3047@hcd28.00 [140.0000-2000.0000]
N^GSSLDS#PGKQ^DTEEDEEEDEK
812.42
1022.38
y8+
y7+
893.34
713.37
982.38
722.30
y9+
1151.42
b3+
260.09
y2+
276.15
y5+
649.27
y6+
778.31
973.36
y10+
1252.46
y4+
520.22
355.16
242.08
327.17
173.06
y3+
391.18
b5+
488.20
557.22
632.74
592.76
452.18
924.37
880.85
1087.91
1181.44
y11+
1367.50
y12+
1496.54
1288.67
1411.53
1540.55
1656.99
1887.00
1946.44
m/z

# 23

- Experimental mass: -16.0428
- Unimod information:
- Possible composition: Ammonia-loss + Deamidated, [H(-3)N(-1) + H(-1) N(-1) O]
- Sites in Unimod:
- Sites observed:

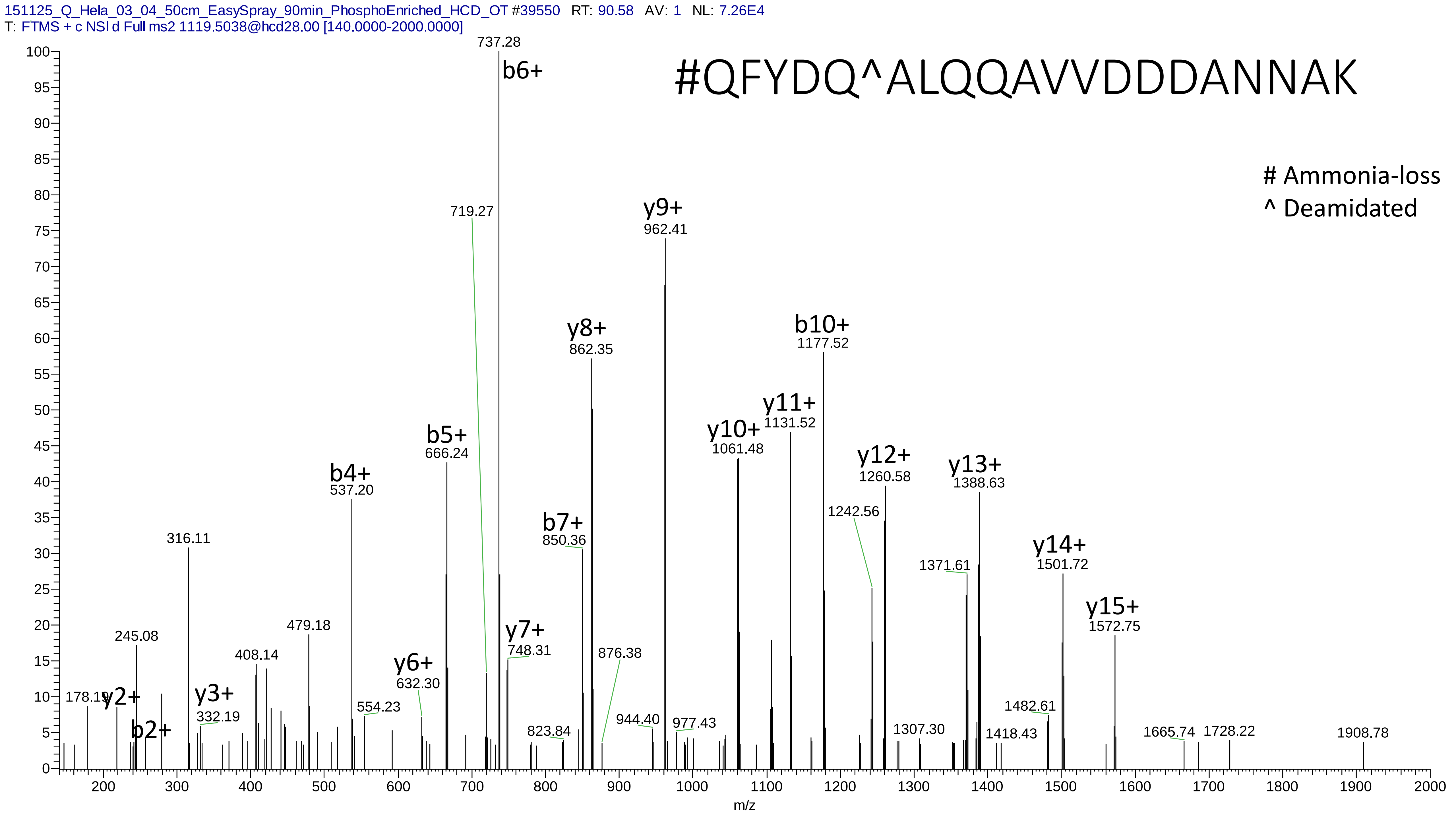
151125_Q_Hela_03_04_50cm_EasySpray_90min_PhosphoEnriched_HCD_OT #39550 RT: 90.58 AV: 1 NL: 7.26E4
T: FTMS + c NSI d Full ms2 1119.5038@hcd28.00 [140.0000-2000.0000]
#QFYDQ^ALQQAVVDDDANNAK
### Ammonia-loss
^ Deamidated
737.28
b6+
719.27
y9+
962.41
y8+
862.35
b10+
1177.52
y11+
1131.52
y10+
1061.48
b5+
666.24
b4+
537.20
y12+
1260.58
y13+
1388.63
b7+
850.36
1242.56
316.11
1371.61
y14+
1501.72
y15+
1572.75
479.18
245.08
y7+
748.31
408.14
876.38
y6+
632.30
178.19
y2+
y3+
332.19
b2+
554.23
823.84
944.40
977.43
1307.30
1482.61
1418.43
1665.74
1728.22
1908.78
m/z

# 24

- Experimental mass: 107.9605
- Unimod information:
- Possible composition: Phosphorylation + Formyl, [H(1)O(3)P(1) +  C O]
- Sites in Unimod:
- Sites observed:

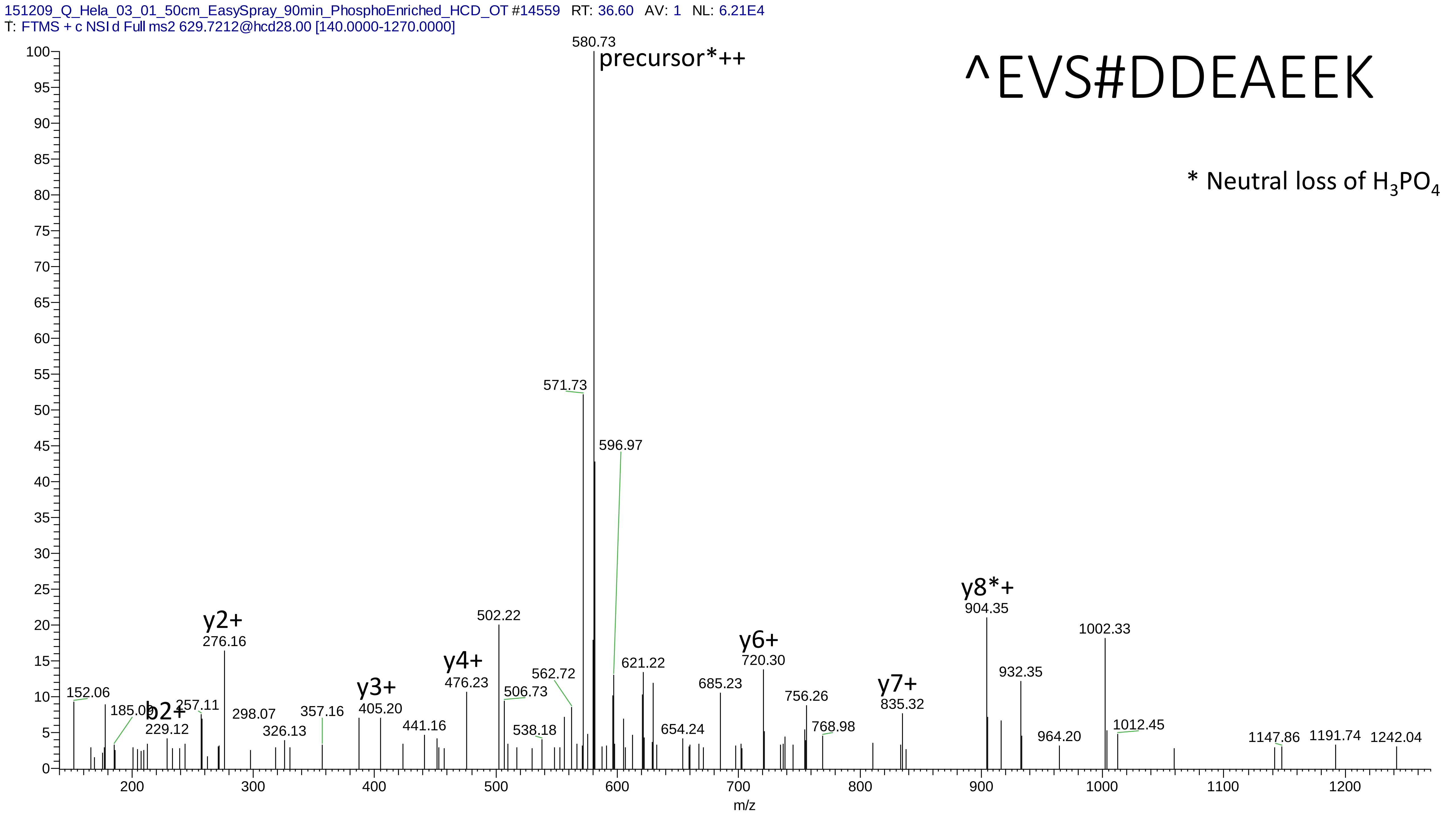
151209_Q_Hela_03_01_50cm_EasySpray_90min_PhosphoEnriched_HCD_OT #14559 RT: 36.60 AV: 1 NL: 6.21E4
T: FTMS + c NSI d Full ms2 629.7212@hcd28.00 [140.0000-1270.0000]
^EVS#DDEAEEK
* Neutral loss of H3PO4
precursor*++
580.73
571.73
596.97
y8*+
904.35
1002.33
y2+
276.16
502.22
y6+
720.30
932.35
621.22
y4+
476.23
562.72
685.23
y3+
y7+
152.06
506.73
756.26
835.32
405.20
b2+
185.09
257.11
357.16
229.12
298.07
441.16
768.98
1012.45
326.13
538.18
654.24
964.20
1147.86
1191.74
1242.04
m/z

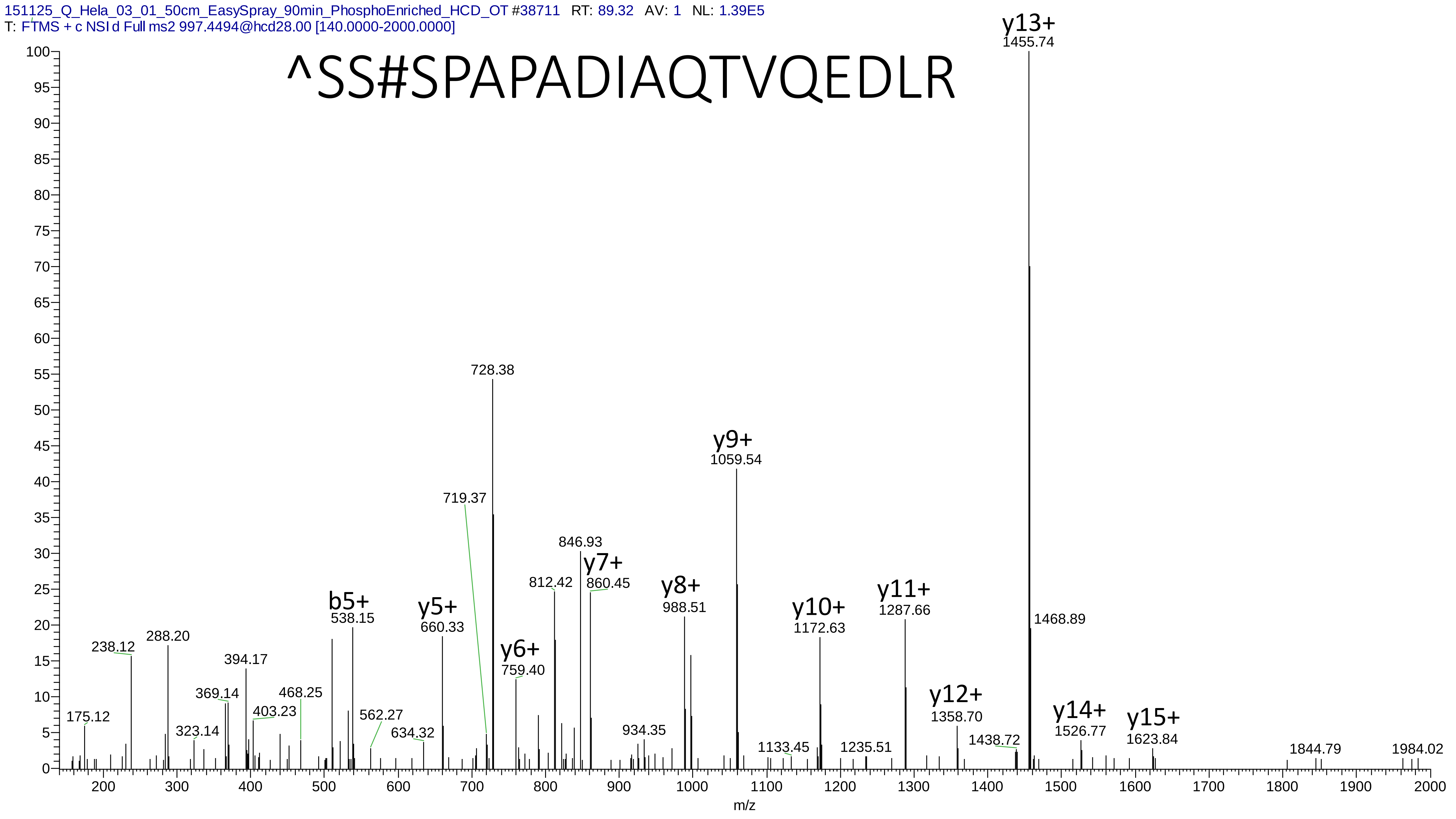
151125_Q_Hela_03_01_50cm_EasySpray_90min_PhosphoEnriched_HCD_OT #38711 RT: 89.32 AV: 1 NL: 1.39E5
T: FTMS + c NSI d Full ms2 997.4494@hcd28.00 [140.0000-2000.0000]
^SS#SPAPADIAQTVQEDLR
y13+
1455.74
728.38
719.37
y9+
1059.54
846.93
y7+
860.45
812.42
y8+
988.51
y10+
1172.63
y11+
1287.66
1468.89
b5+
538.15
y5+
660.33
y6+
759.40
238.12
288.20
394.17
369.14
468.25
403.23
175.12
323.14
562.27
634.32
934.35
1133.45
1235.51
y12+
1358.70
1438.72
y14+
1526.77
y15+
1623.84
1844.79
1984.02
m/z
