## Supplementary material 4 for "MetaLab 2.0 enables accurate post-translational modifications profiling in metaproteomics"

### Supplementary material 4: Manually annotated spectra of the top 30 identified potential modifications

Dataset: U87_phos

# 1

- Experimental mass: 79.967
- Unimod information: phosphorylation
- Possible composition: H O(3) P
- Sites in Unimod: S;T;Y;D;H;C;R;K
- Sites observed:
- Since phosphorylation is common in this dataset, spectra are not showed

# 2

- Experimental mass: 159.9339
- Unimod information: pyrophospho
- Possible composition: H(1)O(3)P(1) * 2
- Sites in Unimod: S;T
- Sites observed:
- Features:  99.8% (2532/2536) peptides contain equal or more than two Ser/Thr

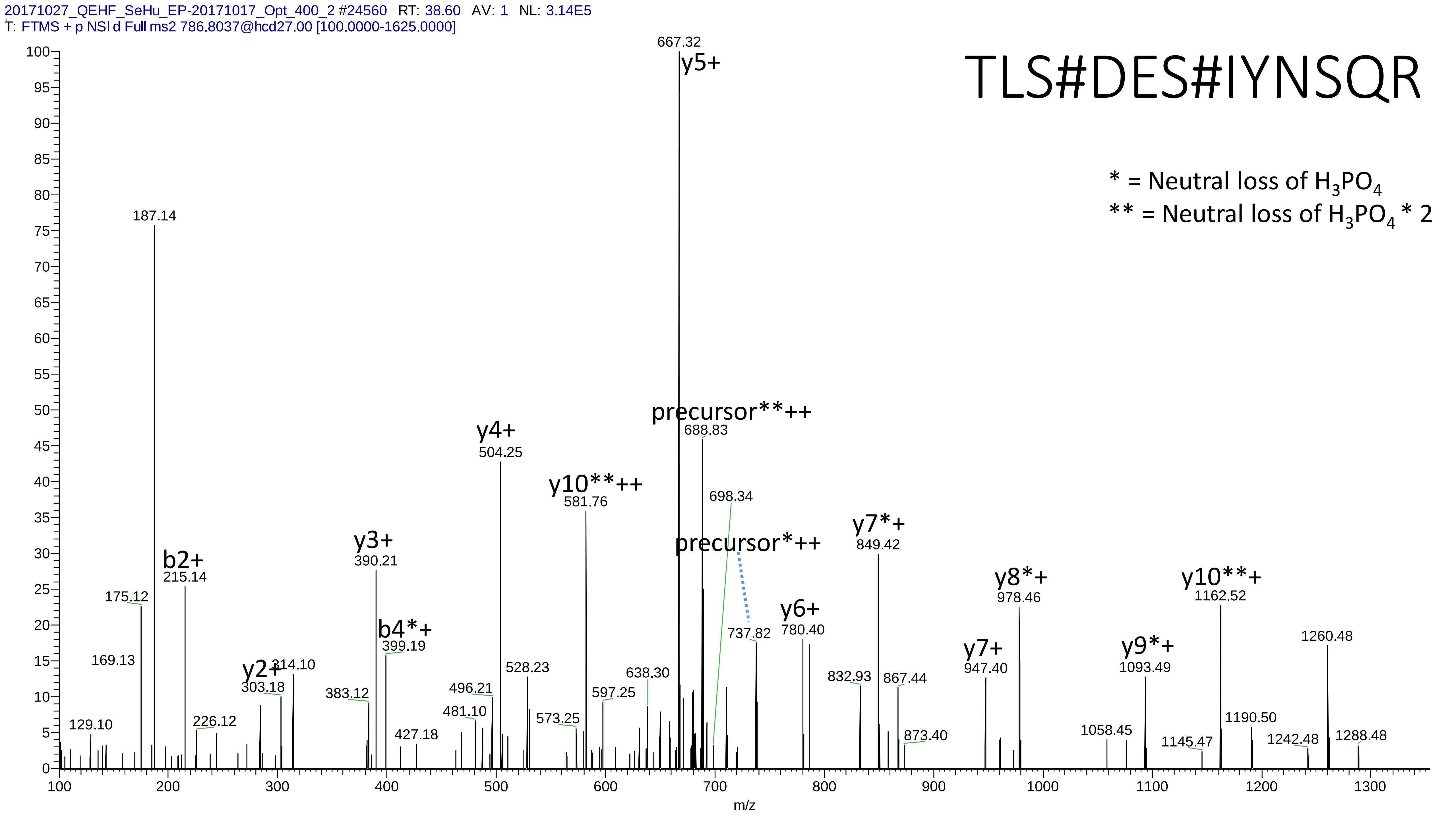

20171027_QEHF_SeHu_EP-20171017_Opt_400_2 #24560 RT: 38.60 AV: 1 NL: 3.14E5
T: FTMS + p NSI d Full ms2 786.8037@hcd27.00 [100.0000-1625.0000]
TLS#DES#IYNSQR
* = Neutral loss of $H_3PO_4$
** = Neutral loss of $H_3PO_4$ * 2
667.32
y5+
187.14
precursor**++
688.83
y4+
504.25
y10**++
581.76
698.34
precursor*++
y7*+
849.42
y3+
390.21
b2+
215.14
175.12
y10**+
1162.52
y8*+
978.46
b4*+
399.19
y6+
780.40
737.82
1260.48
169.13
y2+
303.18
314.10
528.23
638.30
y7+
947.40
y9*+
1093.49
383.12
496.21
597.25
832.93
867.44
481.10
573.25
129.10
226.12
427.18
873.40
1058.45
1145.47
1190.50
1242.48
1288.48
m/z

# 3

- Experimental mass: 95.9629
- Unimod information:
- Possible composition: Phosphorylation + Oxidation, [H(1)O(3)P(1) + O]
- Sites in Unimod:
- Sites observed:

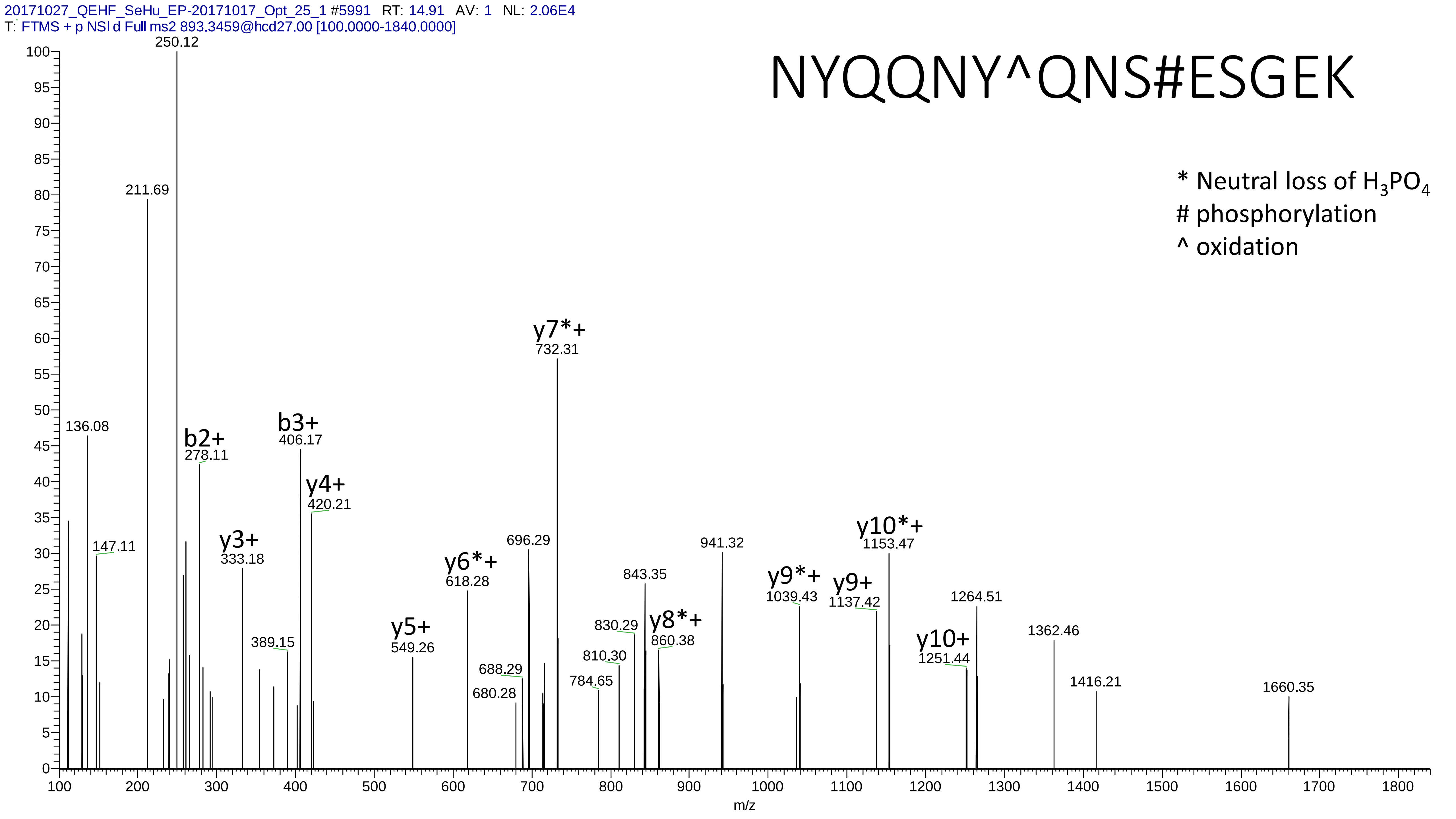

20171027_QEHF_SeHu_EP-20171017_Opt_25_1 #5991 RT: 14.91 AV: 1 NL: 2.06E4
T: FTMS + p NSI d Full ms2 893.3459@hcd27.00 [100.0000-1840.0000]
NYQQNY^QNS#ESGEK
* Neutral loss of $H_3PO_4$
### phosphorylation
^ oxidation
250.12
211.69
136.08
147.11
b2+ 278.11
y3+ 333.18
389.15
b3+ 406.17
y4+ 420.21
y5+ 549.26
y6*+ 618.28
680.28
688.29
696.29
y7*+ 732.31
784.65
810.30
830.29
843.35
y8*+ 860.38
941.32
y9*+ 1039.43
y9+ 1137.42
y10*+ 1153.47
y10+ 1251.44
1264.51
1362.46
1416.21
1660.35
m/z

# 4

- Experimental mass: 101.9485
- Unimod information:
- Possible composition: Phosphorylation + Cation:Na, [H(1)O(3)P(1) + H(-1) Na]
- Sites in Unimod:
- Sites observed:
- Features: phosphorylation sites determined, cation modified sites are not observed

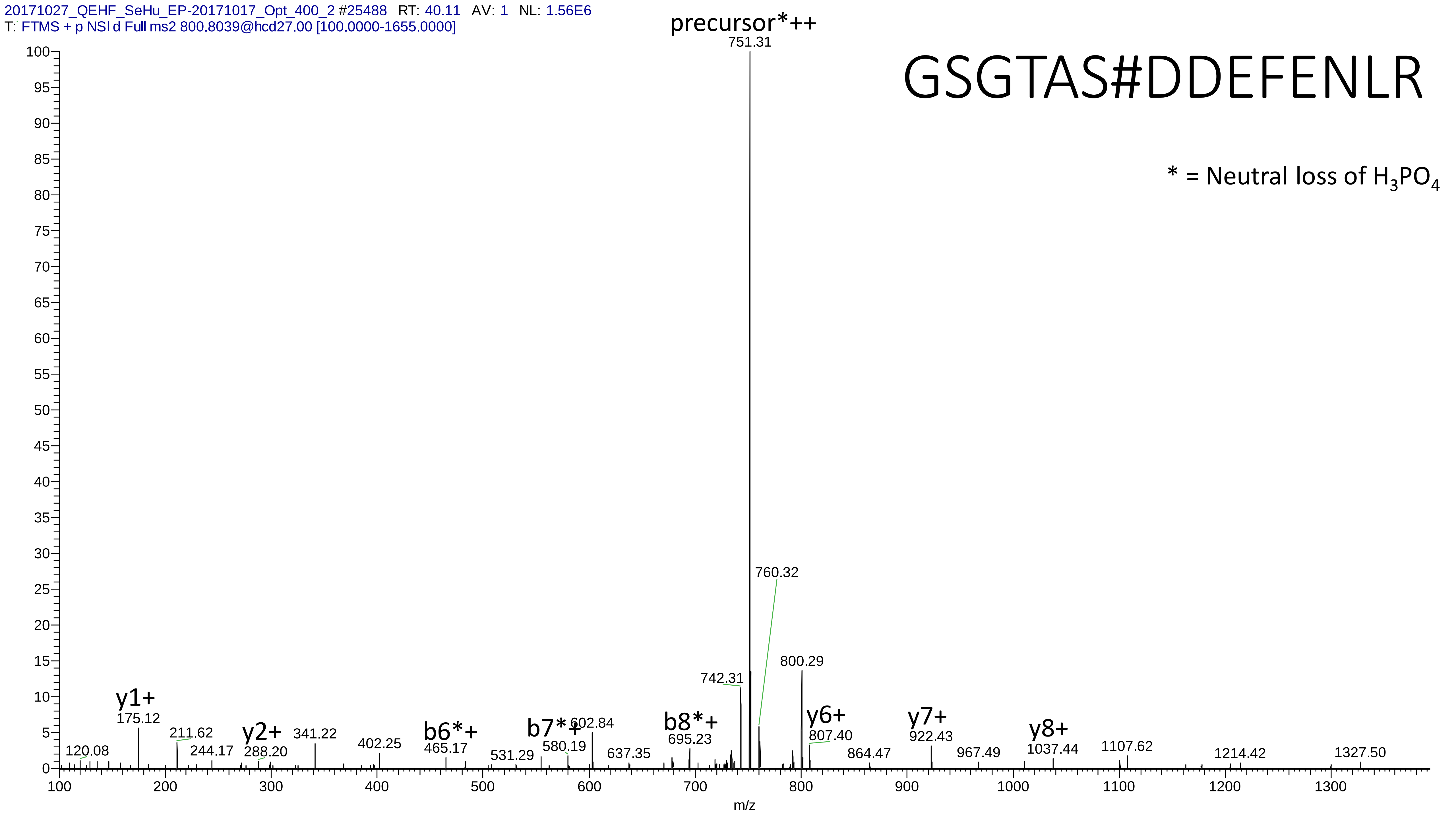

20171027_QEHF_SeHu_EP-20171017_Opt_400_2 #25488 RT: 40.11 AV: 1 NL: 1.56E6
T: FTMS + p NSI d Full ms2 800.8039@hcd27.00 [100.0000-1655.0000]
GSGTAS#DDEFENLR
* = Neutral loss of $H_3PO_4$
precursor*++
751.31
760.32
800.29
742.31
y1+
175.12
211.62
244.17
y2+
288.20
341.22
402.25
b6*+
465.17
531.29
b7*+
580.19
602.84
637.35
b8*+
695.23
y6+
807.40
864.47
y7+
922.43
967.49
y8+
1037.44
1107.62
1214.42
1327.50
120.08
m/z

# 5

- Experimental mass: 80.9524
- Unimod information:
- Possible composition: Phosphorylation + Deamidation, [H(1)O(3)P(1) + H(-1) N(-1) O]; Phosphorylation + Carboxymethyl - Carbamidomethyl, [H(1)O(3)P(1) + H(2) C(2) O(2) - H(3) C(2) N O]
- Sites in Unimod:
- Sites observed: N;Q;C
- Features:  90.6% (739/816) peptides contain Asn/Gln, modification on C is fixed modification Carbamidomethyl replaced by Carboxymethyl

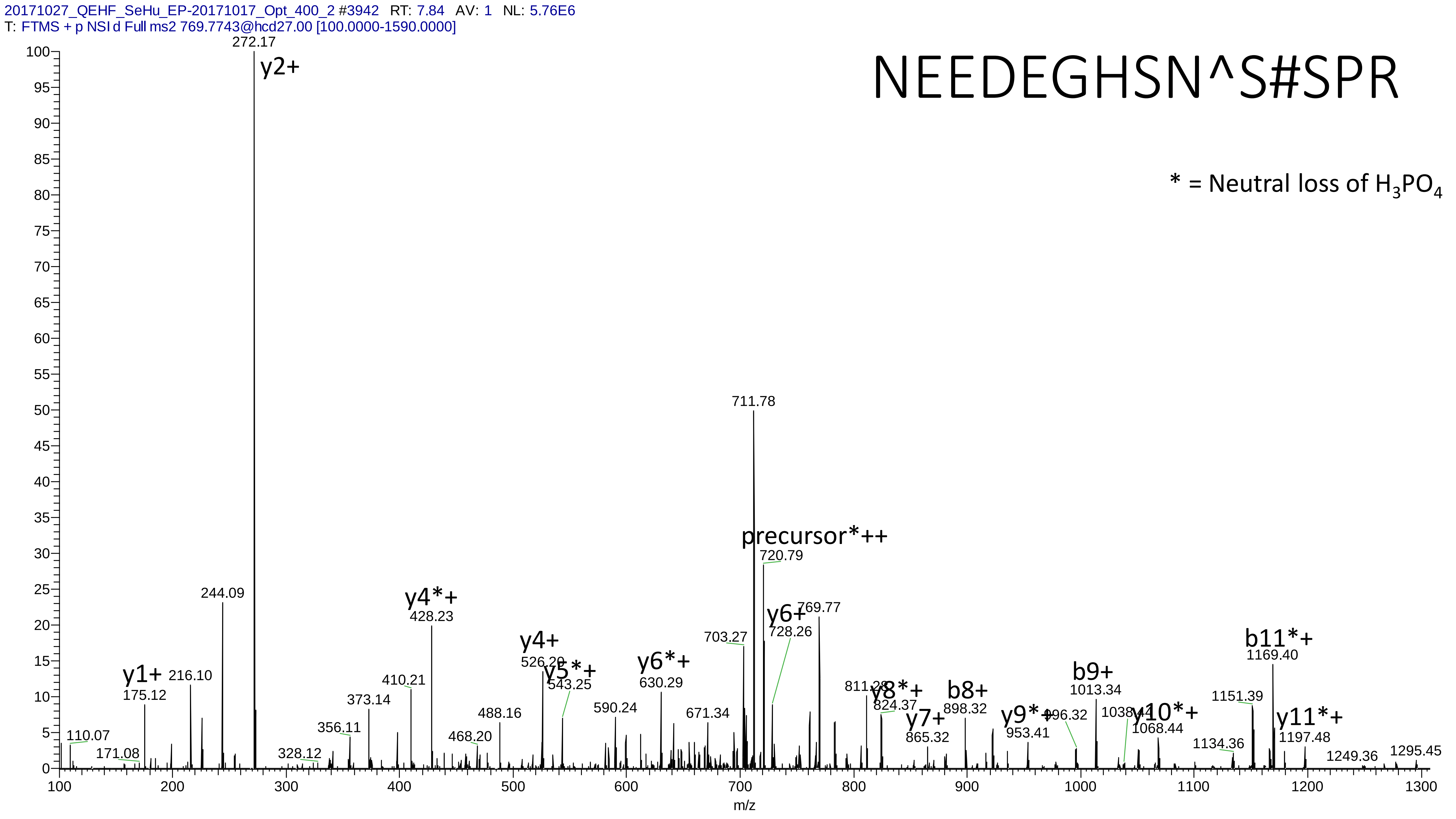

20171027_QEHF_SeHu_EP-20171017_Opt_400_2 #3942 RT: 7.84 AV: 1 NL: 5.76E6
T: FTMS + p NSI d Full ms2 769.7743@hcd27.00 [100.0000-1590.0000]
NEEDEGHSN^S#SPR
* = Neutral loss of H3PO4
272.17
y2+
711.78
precursor*++
720.79
y6+
728.26
769.77
703.27
244.09
y4*+
428.23
y4+
526.29
y5*+
543.25
y6*+
630.29
y1+
175.12
216.10
410.21
373.14
488.16
590.24
671.34
811.28
y8*+
824.37
b8+
898.32
y7+
865.32
y9*+
953.41
996.32
b9+
1013.34
1038.41
y10*+
1068.44
b11*+
1169.40
1151.39
y11*+
1197.48
1134.36
1249.36
1295.45
110.07
171.08
328.12
356.11
468.20
m/z

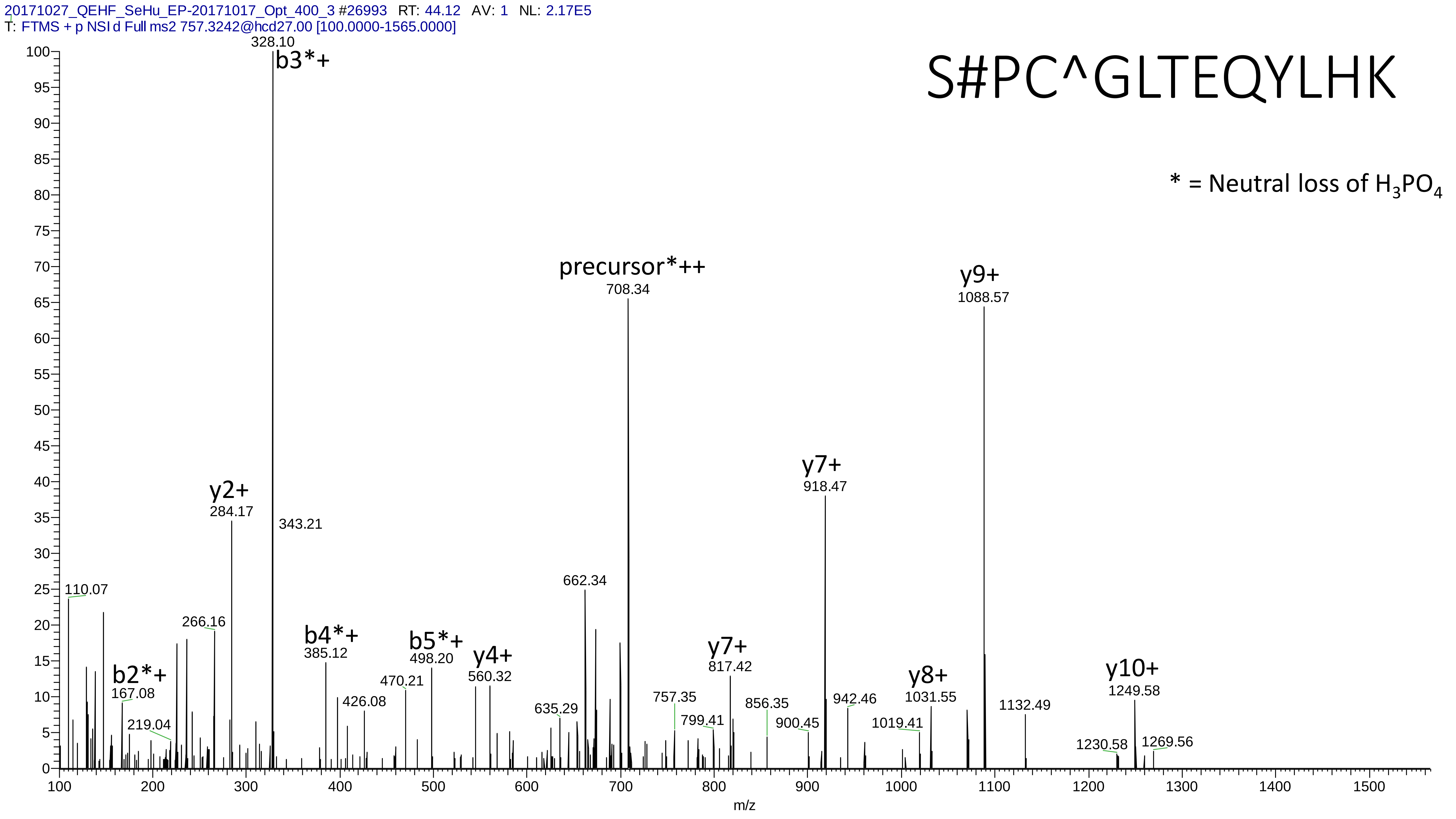

20171027_QEHF_SeHu_EP-20171017_Opt_400_3 #26993 RT: 44.12 AV: 1 NL: 2.17E5
T: FTMS + p NSI d Full ms2 757.3242@hcd27.00 [100.0000-1565.0000]
S#PC^GLTEQYLHK
* = Neutral loss of $H_3PO_4$
b3*+ 328.10
343.21
y2+ 284.17
precursor*++ 708.34
y9+ 1088.57
y7+ 918.47
110.07
266.16
662.34
b4*+ 385.12
b5*+ 498.20
y4+ 560.32
y7+ 817.42
b2*+ 167.08
470.21
426.08
757.35
856.35
942.46
y8+ 1031.55
1132.49
y10+ 1249.58
635.29
799.41
900.45
1019.41
219.04
1230.58
1269.56
m/z

# 6

- Experimental mass: 239.9006
- Unimod information:
- Possible composition: Phosphorylation * 3, [H(1)O(3)P(1) * 3]
- Sites in Unimod:
- Sites observed:
- Features:  99.6% (483/485) peptides contain equal or more than three Ser/Thr

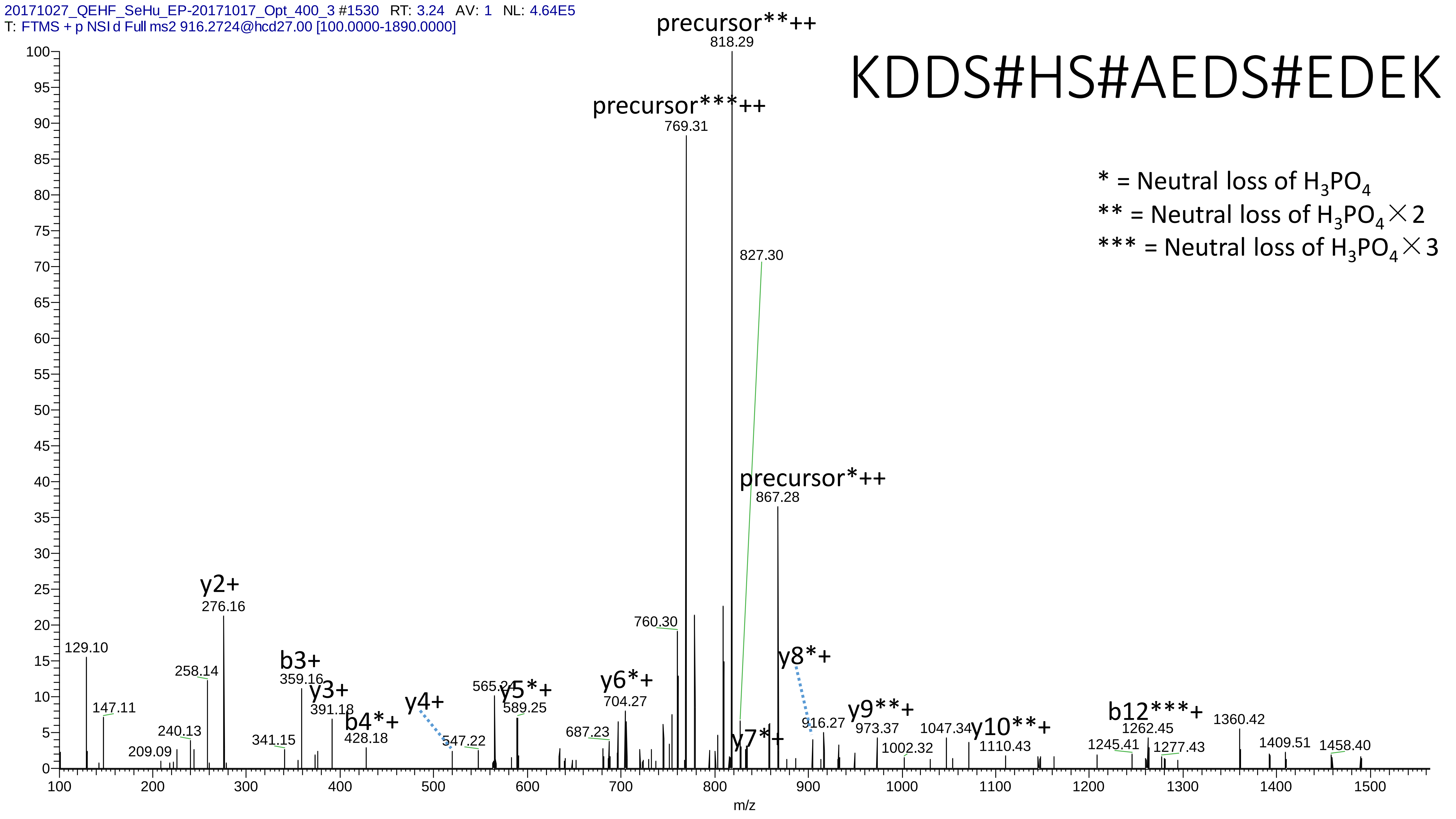

20171027_QEHF_SeHu_EP-20171017_Opt_400_3 #1530 RT: 3.24 AV: 1 NL: 4.64E5
T: FTMS + p NSI d Full ms2 916.2724@hcd27.00 [100.0000-1890.0000]
KDDS#HS#AEDS#EDEK
* = Neutral loss of H3PO4
** = Neutral loss of H3PO4×2
*** = Neutral loss of H3PO4×3
precursor**++
818.29
precursor***++
769.31
827.30
precursor*++
867.28
y2+
276.16
129.10
258.14
147.11
240.13
209.09
b3+
359.16
y3+
391.18
341.15
b4*+
428.18
y4+
547.22
565.24
y5*+
589.25
y6*+
704.27
687.23
760.30
y7*+
y8*+
916.27
y9**+
973.37
1002.32
1047.34
y10**+
1110.43
b12***+
1262.45
1245.41
1277.43
1360.42
1409.51
1458.40
m/z

# 7

- Experimental mass: 136.9889
- Unimod information:
- Possible composition: Phosphorylation + Carbamidomethyl, [H(1)O(3)P(1) + H(3) C(2) N O]
- Sites in Unimod:
- Sites observed:

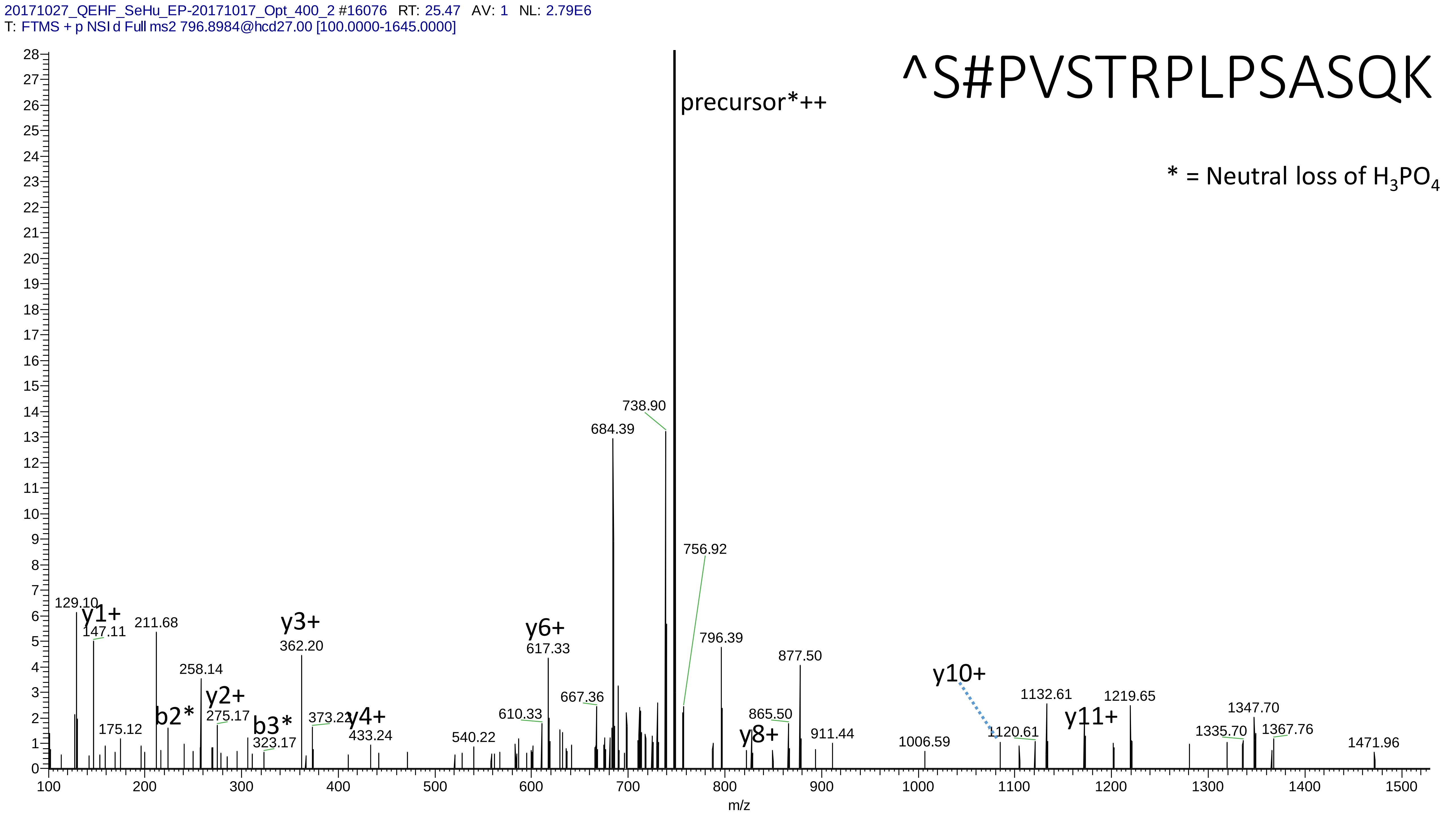
20171027_QEHF_SeHu_EP-20171017_Opt_400_2 #16076 RT: 25.47 AV: 1 NL: 2.79E6
T: FTMS + p NSI d Full ms2 796.8984@hcd27.00 [100.0000-1645.0000]
^S#PVSTRPLPSASQK
* = Neutral loss of $H_3PO_4$
precursor*++
738.90
684.39
756.92
129.10
y1+
147.11
211.68
y3+
362.20
y6+
617.33
796.39
877.50
y10+
1132.61
1219.65
258.14
y2+
275.17
b2*
b3*
323.17
175.12
373.22
y4+
433.24
540.22
610.33
667.36
865.50
y8+
911.44
1006.59
1120.61
y11+
1347.70
1335.70
1367.76
1471.96
m/z

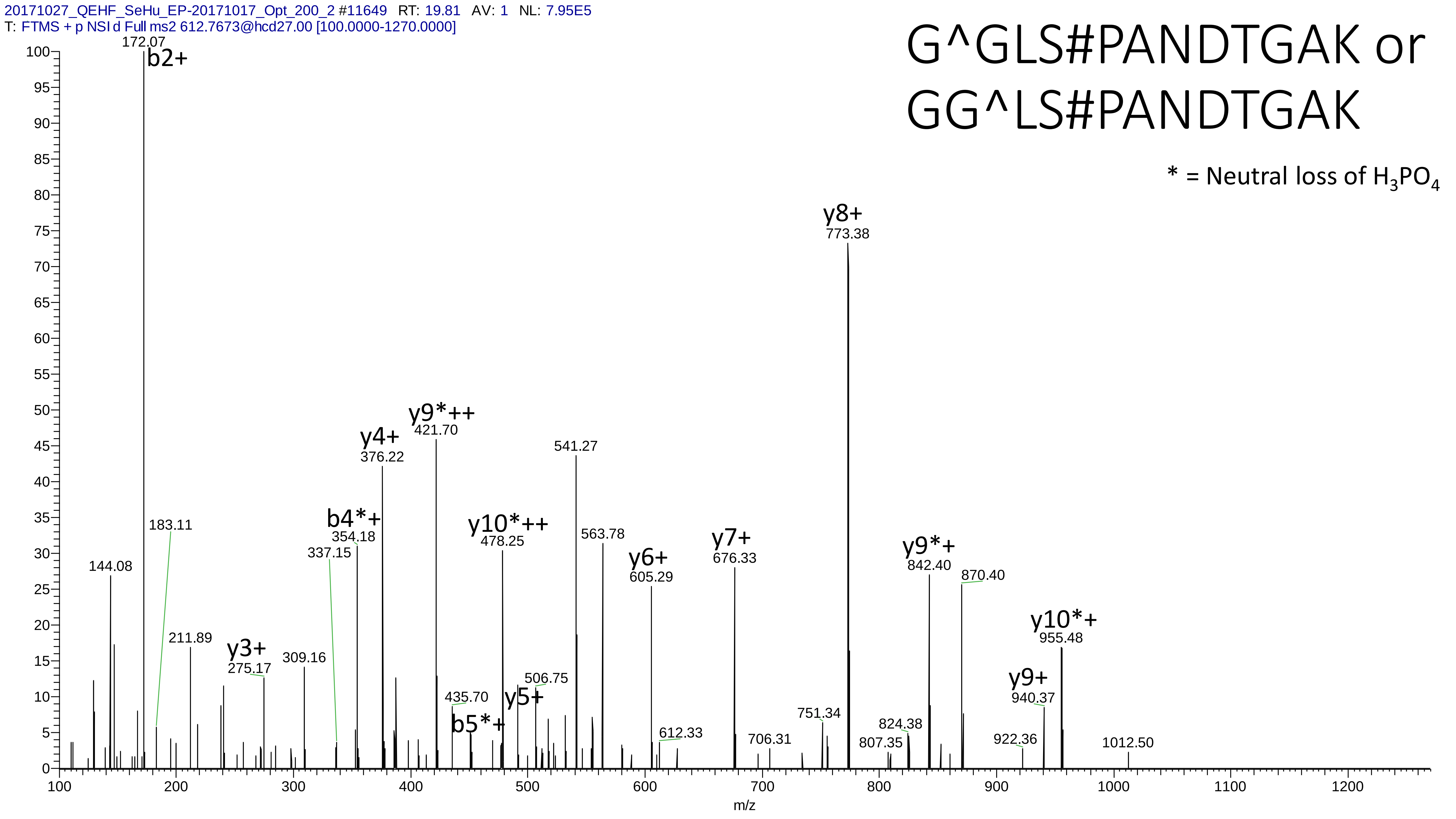
20171027_QEHF_SeHu_EP-20171017_Opt_200_2 #11649 RT: 19.81 AV: 1 NL: 7.95E5
T: FTMS + p NSI d Full ms2 612.7673@hcd27.00 [100.0000-1270.0000]
G^GLS#PANDTGAK or
GG^LS#PANDTGAK
* = Neutral loss of H3PO4
b2+ 172.07
183.11
144.08
211.89
y3+ 275.17
309.16
337.15
b4*+ 354.18
y4+ 376.22
y9*++ 421.70
435.70
b5*+
y10*++ 478.25
506.75
y5+
541.27
563.78
y6+ 605.29
612.33
y7+ 676.33
706.31
751.34
y8+ 773.38
807.35
824.38
y9*+ 842.40
870.40
922.36
y9+ 940.37
y10*+ 955.48
1012.50
m/z

# 8

- Experimental mass: 111.9578
- Unimod information:
- Possible composition:Phosphorylation + Dioxidation, [H(1)O(3)P(1) + O(2)]
- Sites in Unimod:
- Sites observed:

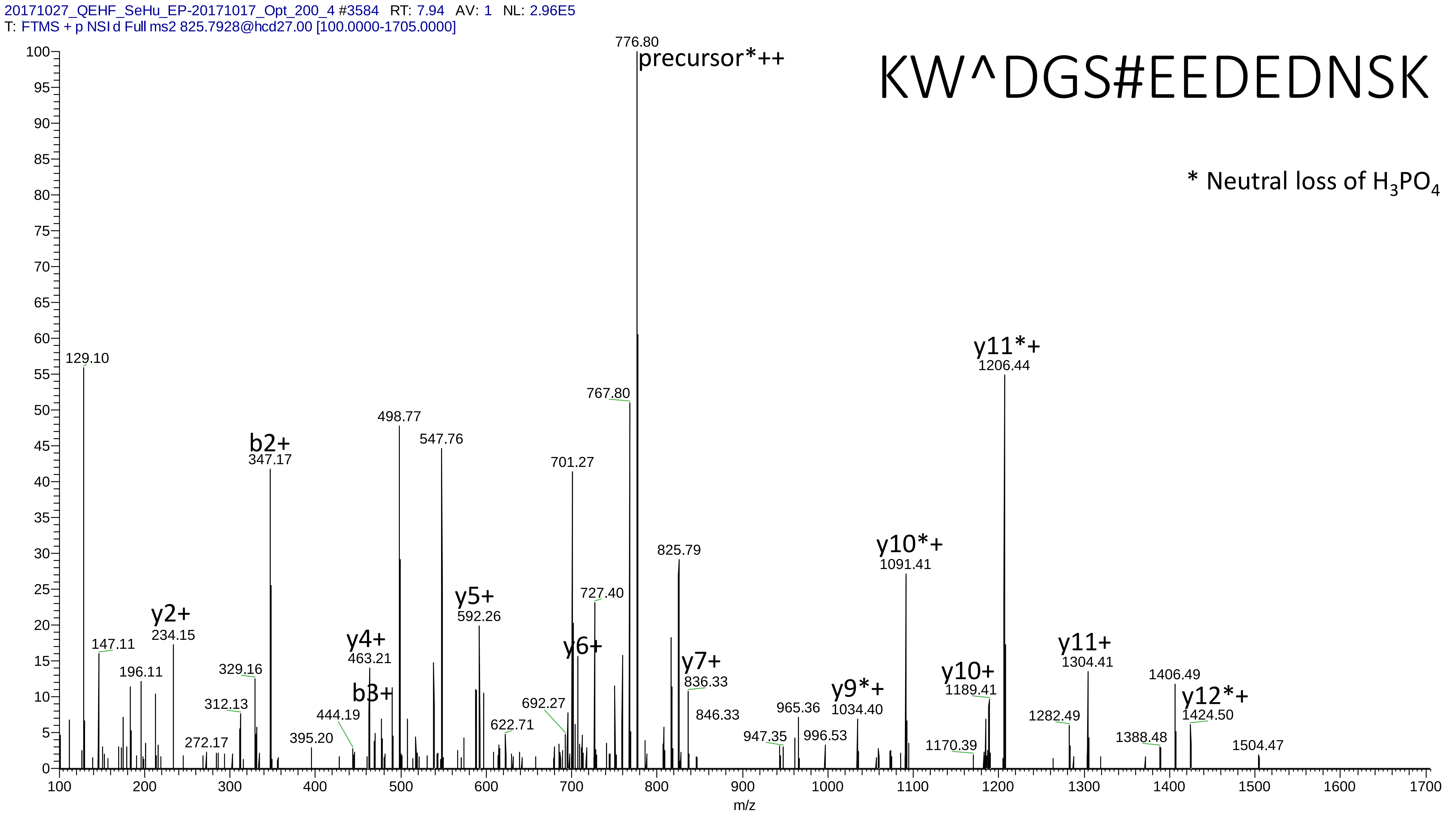
20171027_QEHF_SeHu_EP-20171017_Opt_200_4 #3584 RT: 7.94 AV: 1 NL: 2.96E5
T: FTMS + p NSI d Full ms2 825.7928@hcd27.00 [100.0000-1705.0000]
KW^DGS#EEDEDNSK
* Neutral loss of $H_3PO_4$
776.80
precursor*++
129.10
147.11
196.11
y2+
234.15
272.17
312.13
329.16
b2+
347.17
395.20
444.19
y4+
463.21
b3+
498.77
547.76
y5+
592.26
622.71
692.27
701.27
y6+
727.40
767.80
825.79
y7+
836.33
846.33
947.35
965.36
996.53
y9*+
1034.40
y10*+
1091.41
1170.39
y10+
1189.41
y11*+
1206.44
1282.49
y11+
1304.41
1388.48
1406.49
y12*+
1424.50
1504.47
m/z

20171027_QEHF_SeHu_EP-20171017_Opt_12-5_2 #10064 RT: 25.88 AV: 1 NL: 3.83E4
T: FTMS + p NSI d Full ms2 752.3235@hcd27.00 [100.0000-1555.0000]

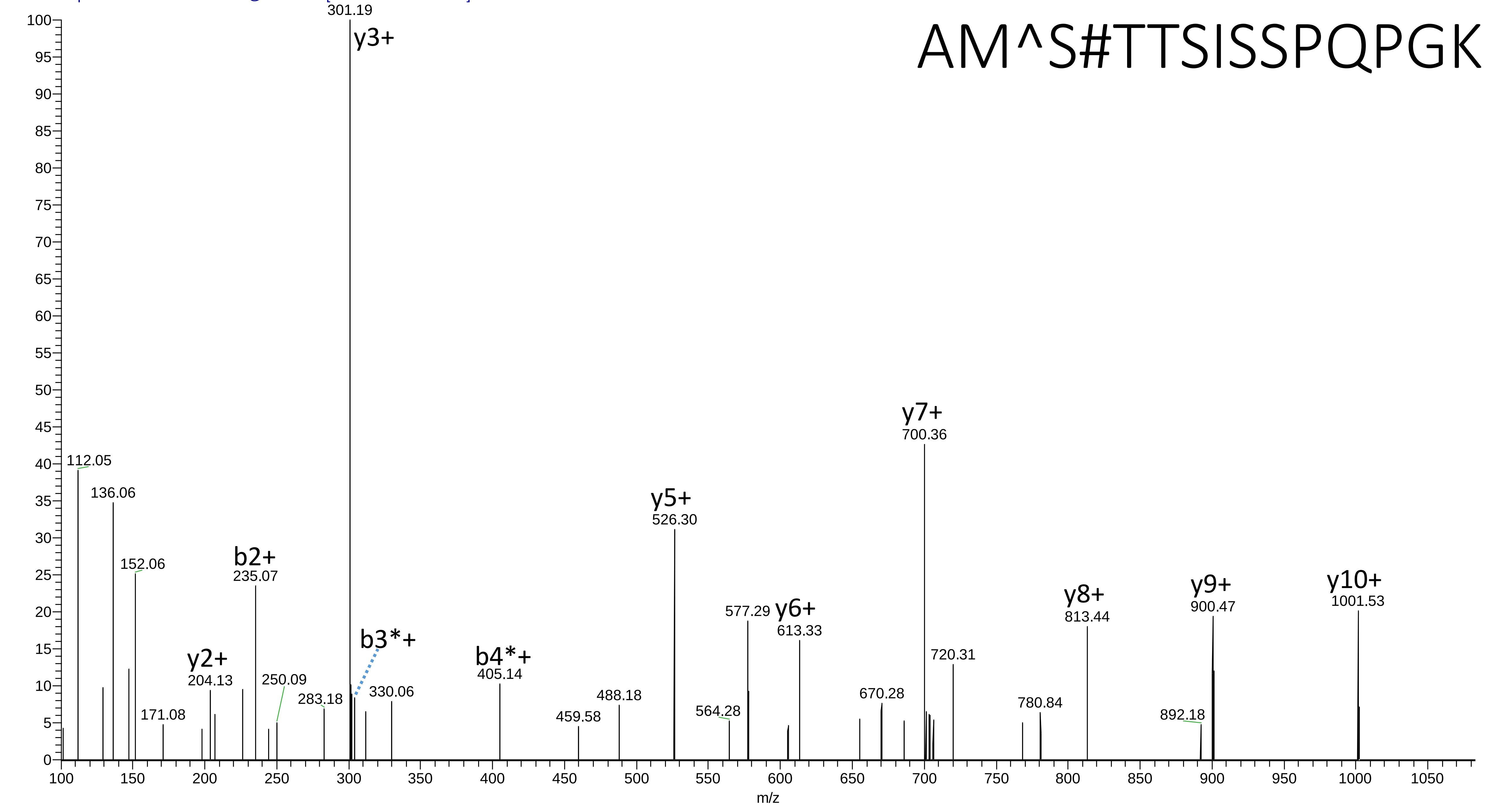

# 9

- Experimental mass: 62.9409
- Unimod information:
- Possible composition: Phosphorylation + Ammonia-loss, [H(1)O(3)P(1) + H(-3)N(-1)]
- Sites in Unimod:
- Sites observed:
- Features:  66.8% (163/244) peptides contain Gln at peptide n-term (Pyro-glu from Q)

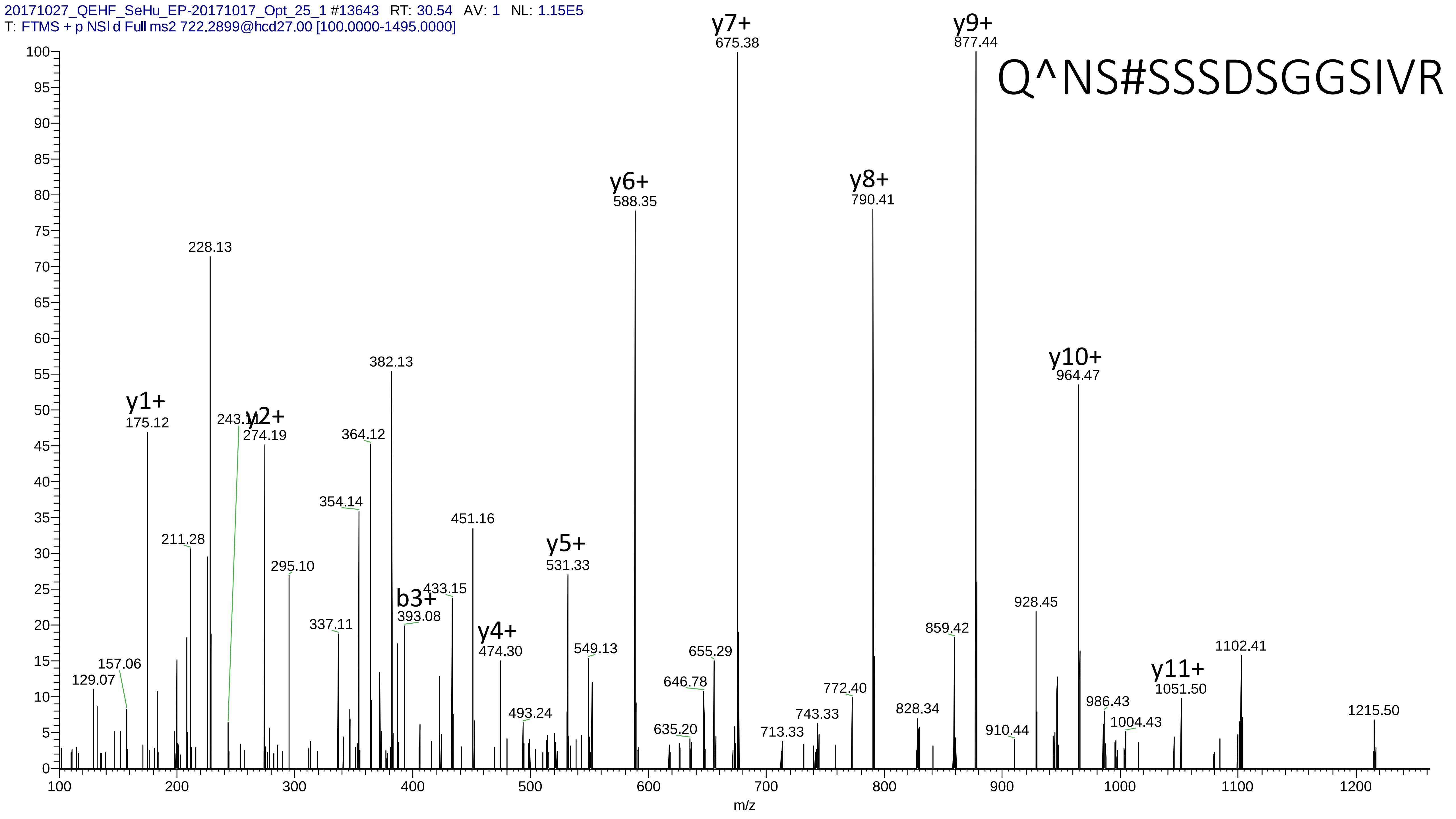

20171027_QEHF_SeHu_EP-20171017_Opt_25_1 #13643 RT: 30.54 AV: 1 NL: 1.15E5
T: FTMS + p NSI d Full ms2 722.2899@hcd27.00 [100.0000-1495.0000]
Q^NS#SSSDSGGSIVR
y1+ 175.12
y2+ 274.19
y4+ 474.30
y5+ 531.33
y6+ 588.35
y7+ 675.38
y8+ 790.41
y9+ 877.44
y10+ 964.47
y11+ 1051.50
b3+ 393.08
129.07
157.06
211.28
228.13
243.11
295.10
337.11
354.14
364.12
382.13
433.15
451.16
493.24
549.13
635.20
646.78
655.29
713.33
743.33
772.40
828.34
859.42
910.44
928.45
986.43
1004.43
1102.41
1215.50
m/z

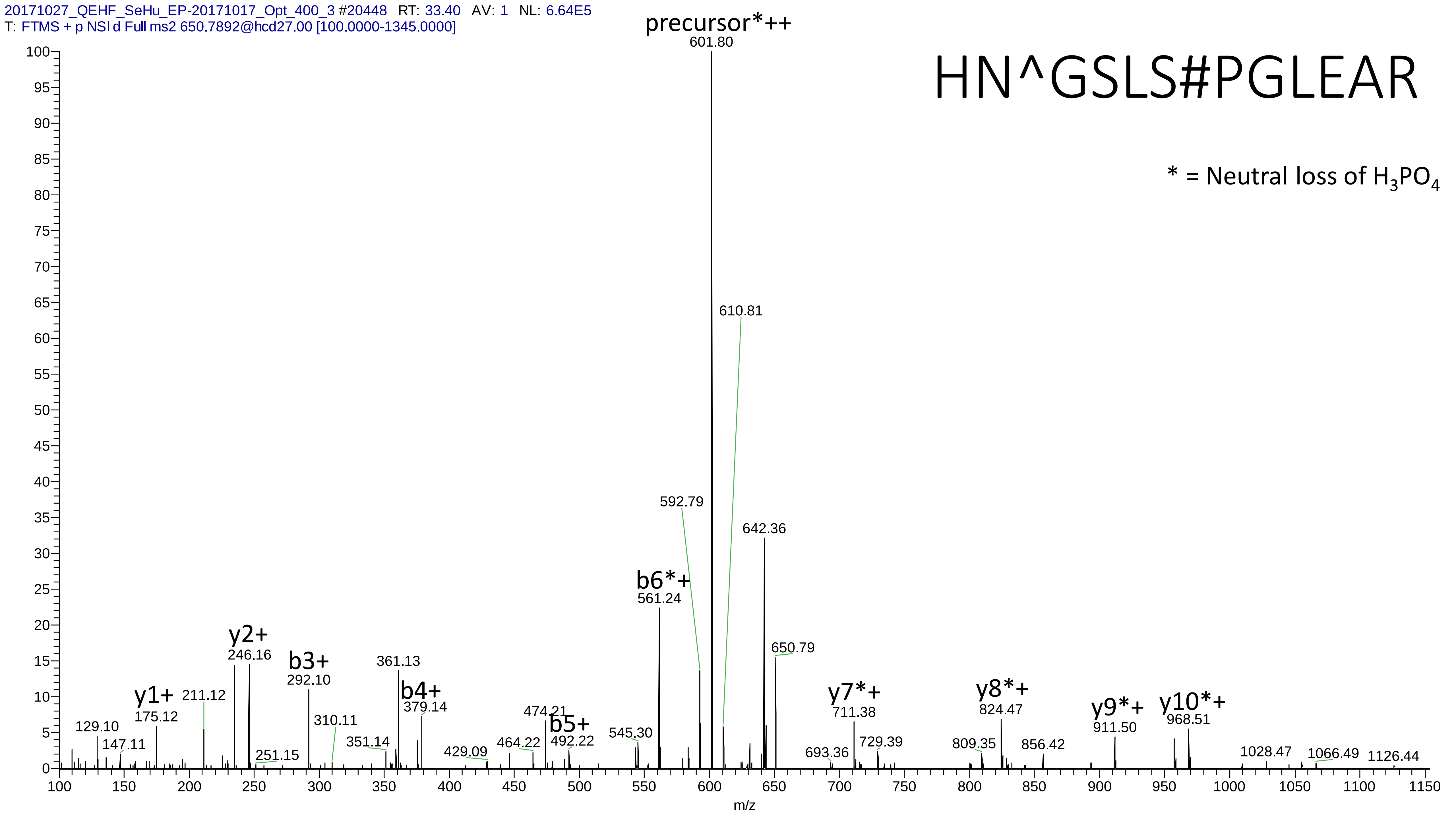

20171027_QEHF_SeHu_EP-20171017_Opt_400_3 #20448 RT: 33.40 AV: 1 NL: 6.64E5
T: FTMS + p NSI d Full ms2 650.7892@hcd27.00 [100.0000-1345.0000]
HN^GSLS#PGLEAR
* = Neutral loss of H3PO4
precursor*++
601.80
610.81
592.79
642.36
b6*+
561.24
650.79
y2+
246.16
b3+
292.10
361.13
b4+
379.14
y1+
175.12
211.12
129.10
147.11
251.15
310.11
351.14
429.09
464.22
474.21
b5+
492.22
545.30
y7*+
711.38
693.36
729.39
y8*+
824.47
809.35
856.42
y9*+
911.50
y10*+
968.51
1028.47
1066.49
1126.44
m/z

# 10

- Experimental mass: 181.915
- Unimod information:
- Possible composition: Phosphorylation * 2 + Cation:Na, [H(1)O(3)P(1) * 2 + H(-1) Na ]
- Sites in Unimod:
- Sites observed:
- Features: phosphorylation sites determined, cation modified sites are not observed

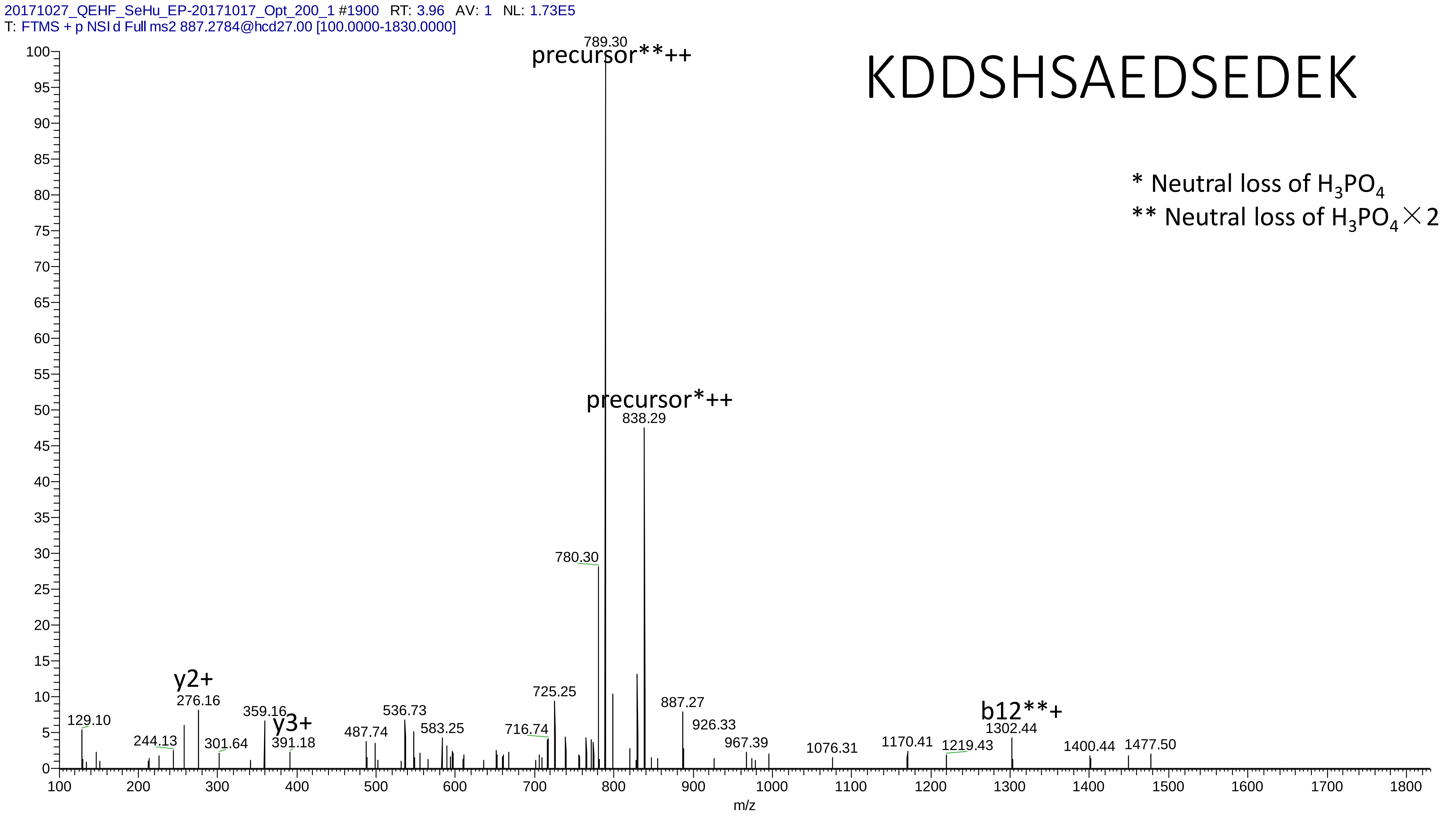
20171027_QEHF_SeHu_EP-20171017_Opt_200_1 #1900 RT: 3.96 AV: 1 NL: 1.73E5
T: FTMS + p NSI d Full ms2 887.2784@hcd27.00 [100.0000-1830.0000]
KDDSHSAEDSEDEK
* Neutral loss of H3PO4
** Neutral loss of H3PO4×2
precursor**++
789.30
precursor*++
838.29
780.30
725.25
887.27
926.33
967.39
1076.31
1170.41
1219.43
b12**+
1302.44
1400.44
1477.50
y2+
276.16
129.10
244.13
301.64
359.16
y3+
391.18
487.74
536.73
583.25
716.74
m/z

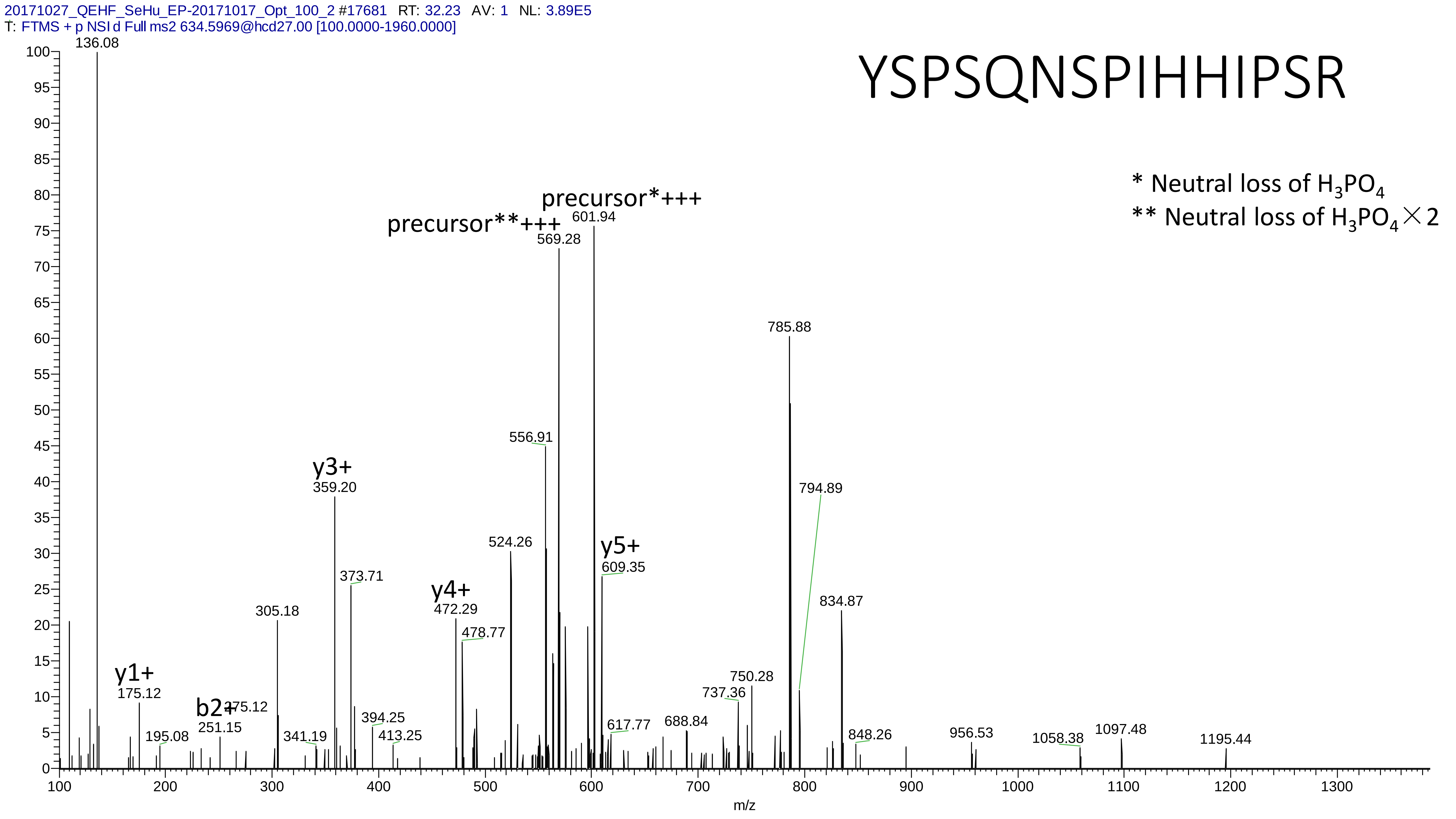

20171027_QEHF_SeHu_EP-20171017_Opt_100_2 #17681 RT: 32.23 AV: 1 NL: 3.89E5
T: FTMS + p NSI d Full ms2 634.5969@hcd27.00 [100.0000-1960.0000]
YSPSQNSPIHHIPSR
* Neutral loss of $H_3PO_4$
** Neutral loss of $H_3PO_4 \times 2$
precursor*+++
precursor**+++
136.08
601.94
569.28
785.88
556.91
794.89
y3+
359.20
524.26
y5+
609.35
373.71
834.87
y4+
472.29
478.77
305.18
750.28
737.36
y1+
175.12
b2+
251.15
175.12
394.25
617.77
688.84
195.08
341.19
413.25
848.26
956.53
1058.38
1097.48
1195.44
m/z

# 11

- Experimental mass: 117.9133
- Unimod information:
- Possible composition: Phosphorylation + Cation:K, [H(1)O(3)P(1) + H(-1) K]
- Sites in Unimod:
- Sites observed:
- Features: phosphorylation sites determined, cation modified sites are not observed

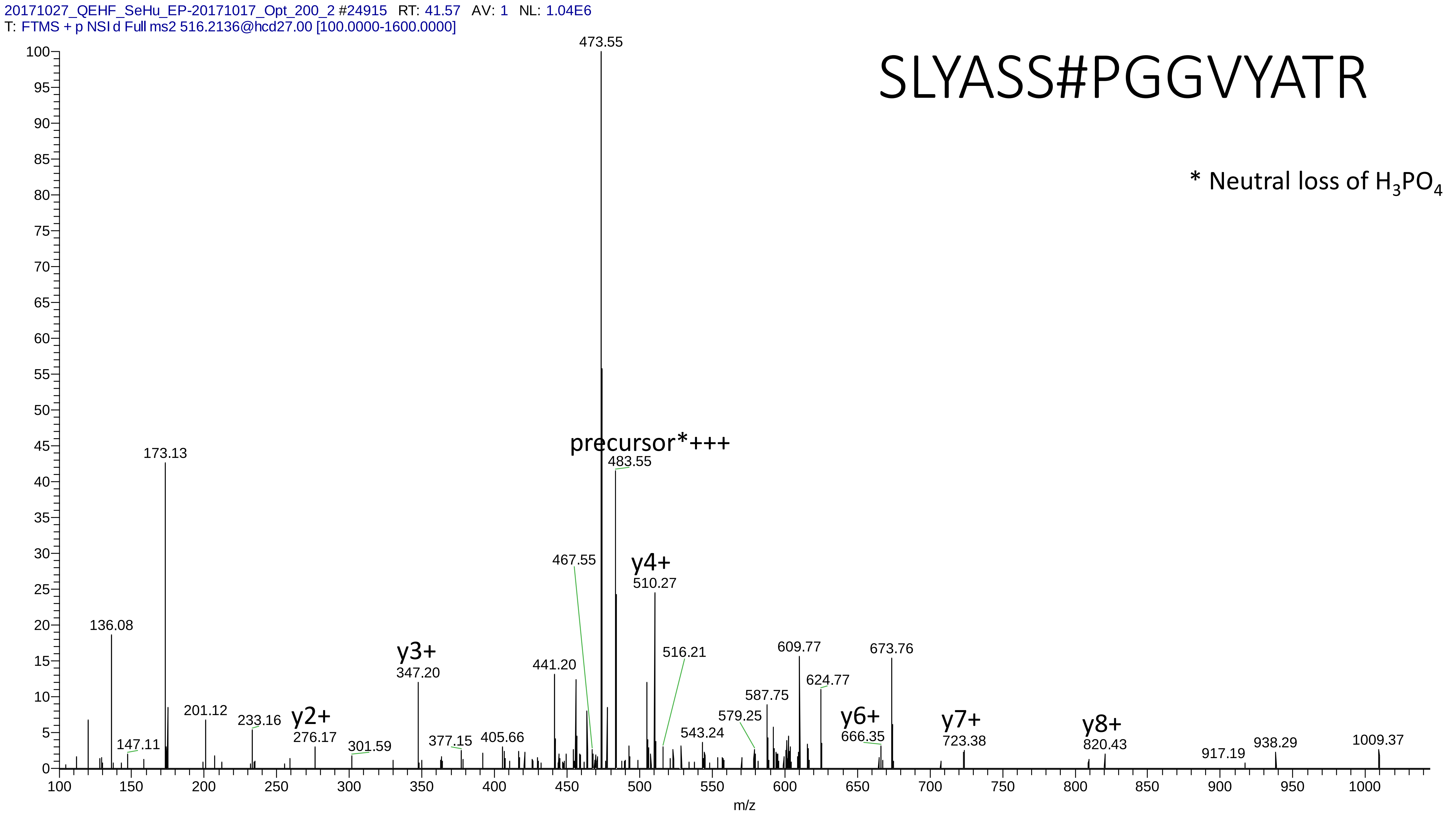
20171027_QEHF_SeHu_EP-20171017_Opt_200_2 #24915 RT: 41.57 AV: 1 NL: 1.04E6
T: FTMS + p NSI d Full ms2 516.2136@hcd27.00 [100.0000-1600.0000]
SLYASS#PGGVYATR
* Neutral loss of H3PO4
precursor*+++
y2+
y3+
y4+
y6+
y7+
y8+
473.55
483.55
173.13
136.08
147.11
201.12
233.16
276.17
301.59
347.20
377.15
405.66
441.20
467.55
510.27
516.21
543.24
579.25
587.75
609.77
624.77
666.35
673.76
723.38
820.43
917.19
938.29
1009.37
m/z

# 12

- Experimental mass: -48.1284
- Unimod information:
- Possible composition: Phosphorylation + Lys-loss, [H(1)O(3)P(1) + H(-12) C(-6) N(-2) O(-1)]
- Sites in Unimod:
- Sites observed:
- Features: 95.3% (164/172) peptides contain Lys at peptide n-term

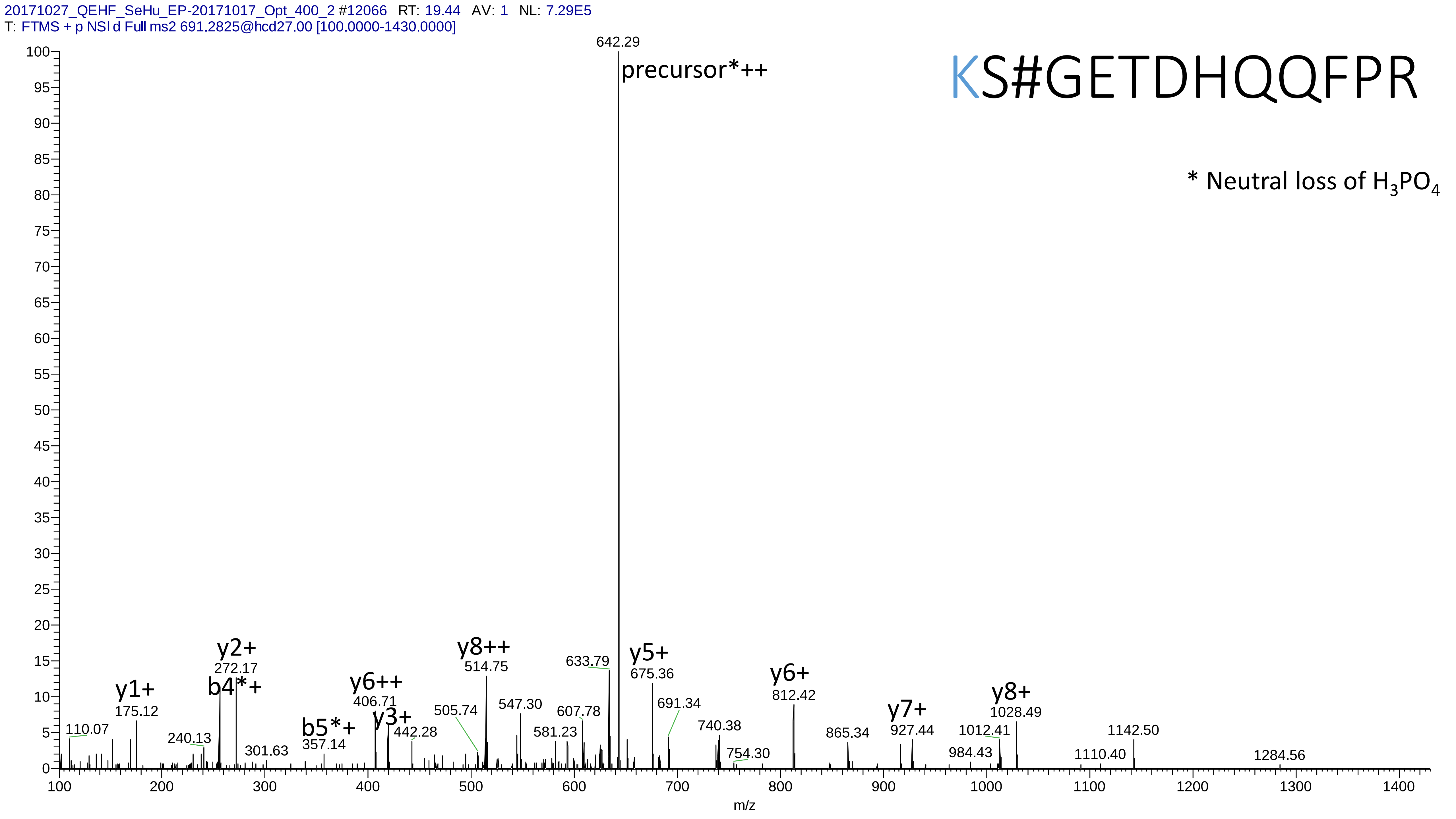

20171027_QEHF_SeHu_EP-20171017_Opt_400_2 #12066 RT: 19.44 AV: 1 NL: 7.29E5
T: FTMS + p NSI d Full ms2 691.2825@hcd27.00 [100.0000-1430.0000]
KS#GETDHQQFPR
* Neutral loss of H3PO4
642.29
precursor*++
y1+
175.12
y2+
272.17
b4*+
b5*+
357.14
y6++
406.71
y3+
442.28
y8++
514.75
633.79
y5+
675.36
y6+
812.42
y7+
927.44
y8+
1028.49
110.07
240.13
301.63
505.74
547.30
581.23
607.78
691.34
740.38
754.30
865.34
984.43
1012.41
1110.40
1142.50
1284.56
m/z

# 13

- Experimental mass: 175.9298
- Unimod information:
- Possible composition: Phospho * 2 + Oxidation, [H(1)O(3)P(1) * 2 + O]
- Sites in Unimod:
- Sites observed:

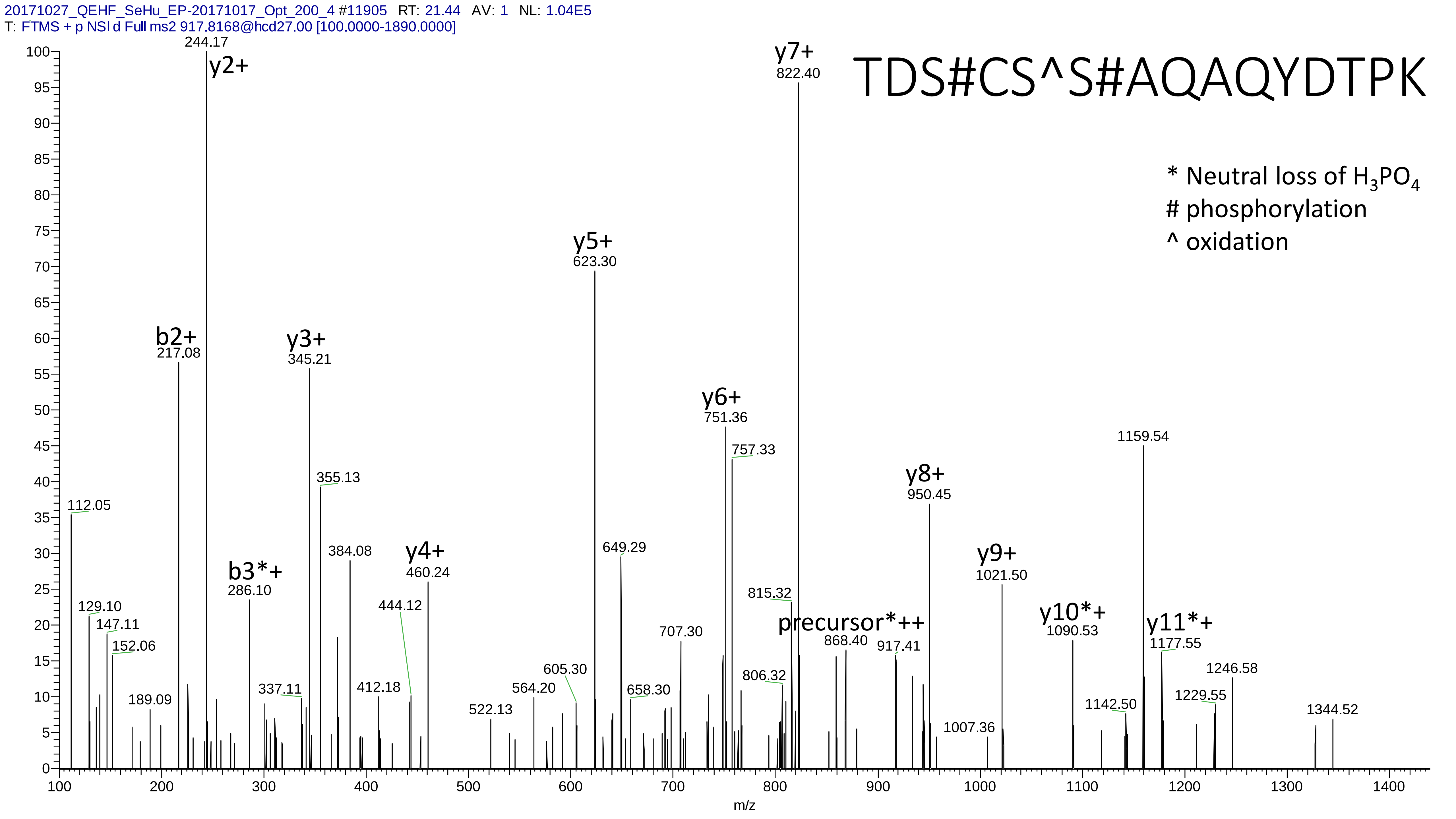

20171027_QEHF_SeHu_EP-20171017_Opt_200_4 #11905 RT: 21.44 AV: 1 NL: 1.04E5
T: FTMS + p NSI d Full ms2 917.8168@hcd27.00 [100.0000-1890.0000]
TDS#CS^S#AQAQYDTPK
* Neutral loss of $H_3PO_4$
### phosphorylation
^ oxidation
y2+ 244.17
y7+ 822.40
y5+ 623.30
b2+ 217.08
y3+ 345.21
y6+ 751.36
757.33
1159.54
355.13
y8+ 950.45
112.05
384.08
y4+ 460.24
649.29
b3*+ 286.10
y9+ 1021.50
815.32
129.10
444.12
147.11
precursor*++
y10*+ 1090.53
y11*+ 1177.55
152.06
707.30
868.40
917.41
605.30
1246.58
806.32
564.20
412.18
337.11
658.30
189.09
1142.50
1229.55
522.13
1344.52
1007.36
m/z

# 14

- Experimental mass: 0.9846
- Unimod information: Deamidated
- Possible composition: [H(-1)N(-1)O(1)]
- Sites in Unimod: Q;R;N;[F
- Sites observed:

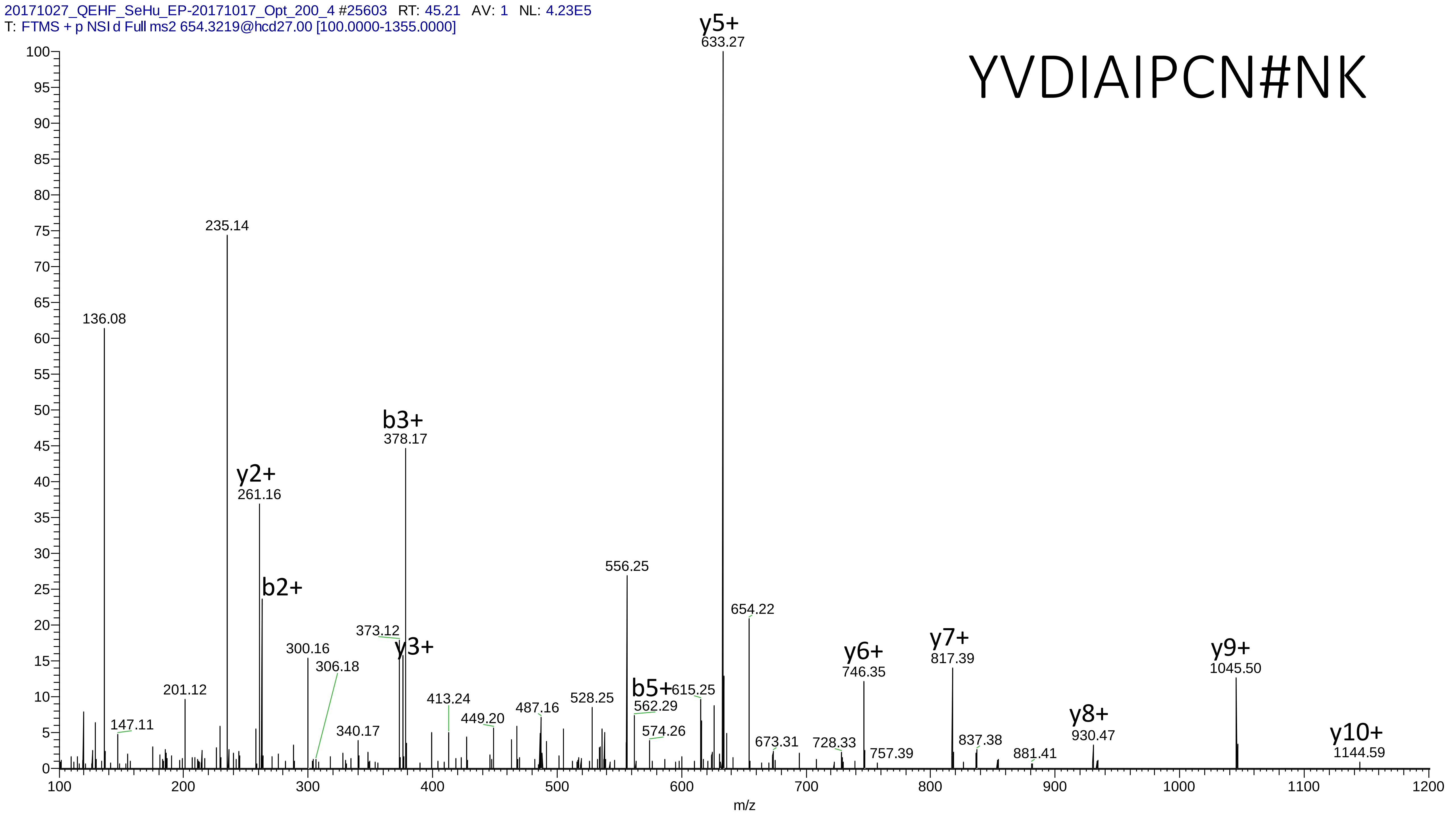

20171027_QEHF_SeHu_EP-20171017_Opt_200_4 #25603 RT: 45.21 AV: 1 NL: 4.23E5
T: FTMS + p NSI d Full ms2 654.3219@hcd27.00 [100.0000-1355.0000]
YVDIAIPCN#NK
y5+
633.27
235.14
136.08
b3+
378.17
y2+
261.16
b2+
556.25
654.22
373.12
y3+
300.16
306.18
y6+
746.35
y7+
817.39
y9+
1045.50
201.12
413.24
449.20
487.16
528.25
b5+
562.29
615.25
147.11
340.17
574.26
673.31
728.33
757.39
837.38
881.41
y8+
930.47
y10+
1144.59
m/z

# 15

- Experimental mass: 133.8856
- Unimod information:
- Possible composition: Phosphorylation + Cation:Fe[II], [H(1)O(3)P(1) + H(-2) Fe]
- Sites in Unimod:
- Sites observed: phosphorylation sites determined, cation modified sites are not observed

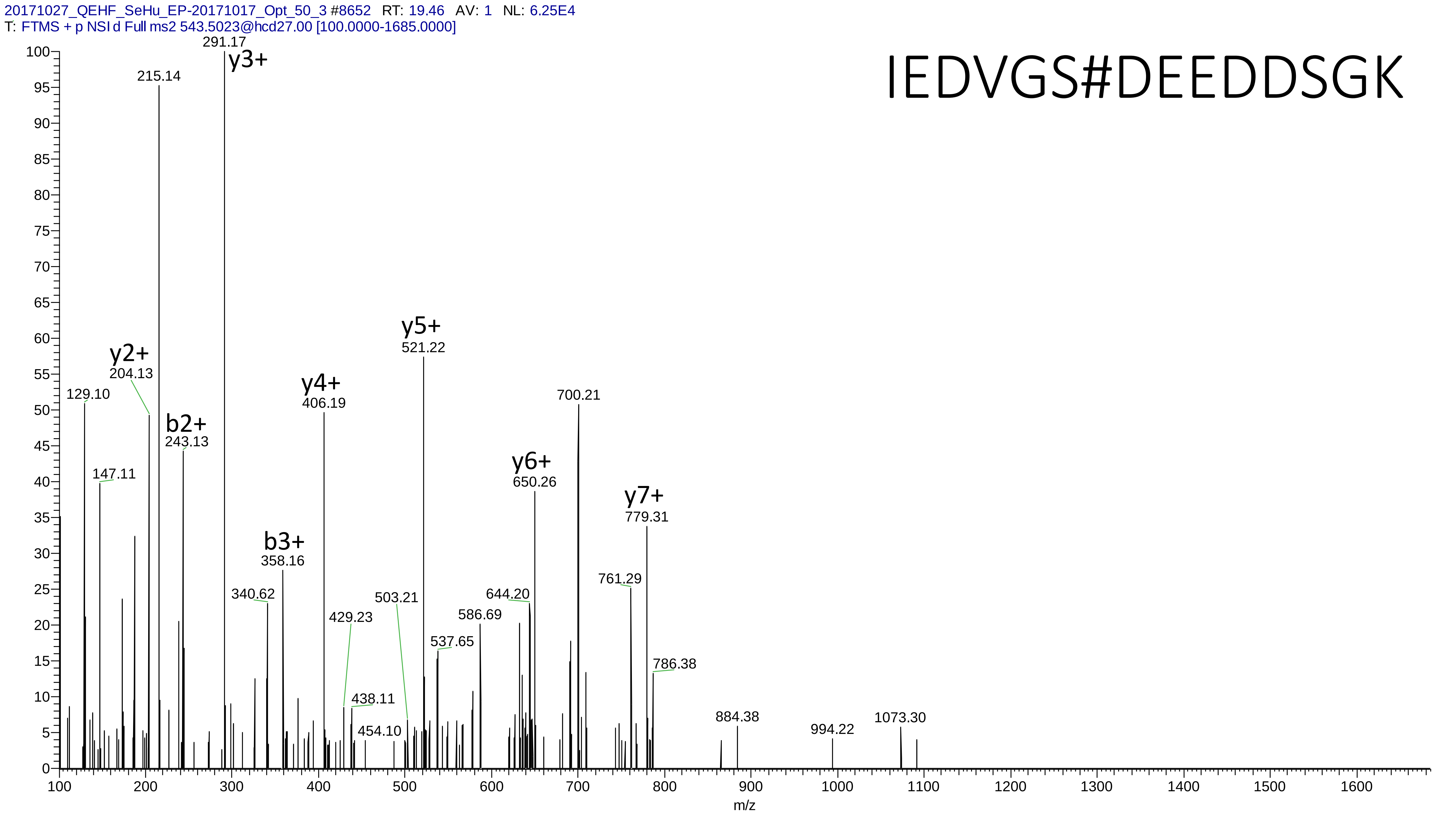

20171027_QEHF_SeHu_EP-20171017_Opt_50_3 #8652 RT: 19.46 AV: 1 NL: 6.25E4
T: FTMS + p NSI d Full ms2 543.5023@hcd27.00 [100.0000-1685.0000]
IEDVGS#DEEDDSGK
291.17
y3+
215.14
y2+
204.13
129.10
b2+
243.13
147.11
y4+
406.19
y5+
521.22
b3+
358.16
340.62
429.23
503.21
438.11
454.10
537.65
586.69
644.20
y6+
650.26
700.21
y7+
779.31
761.29
786.38
884.38
994.22
1073.30
m/z

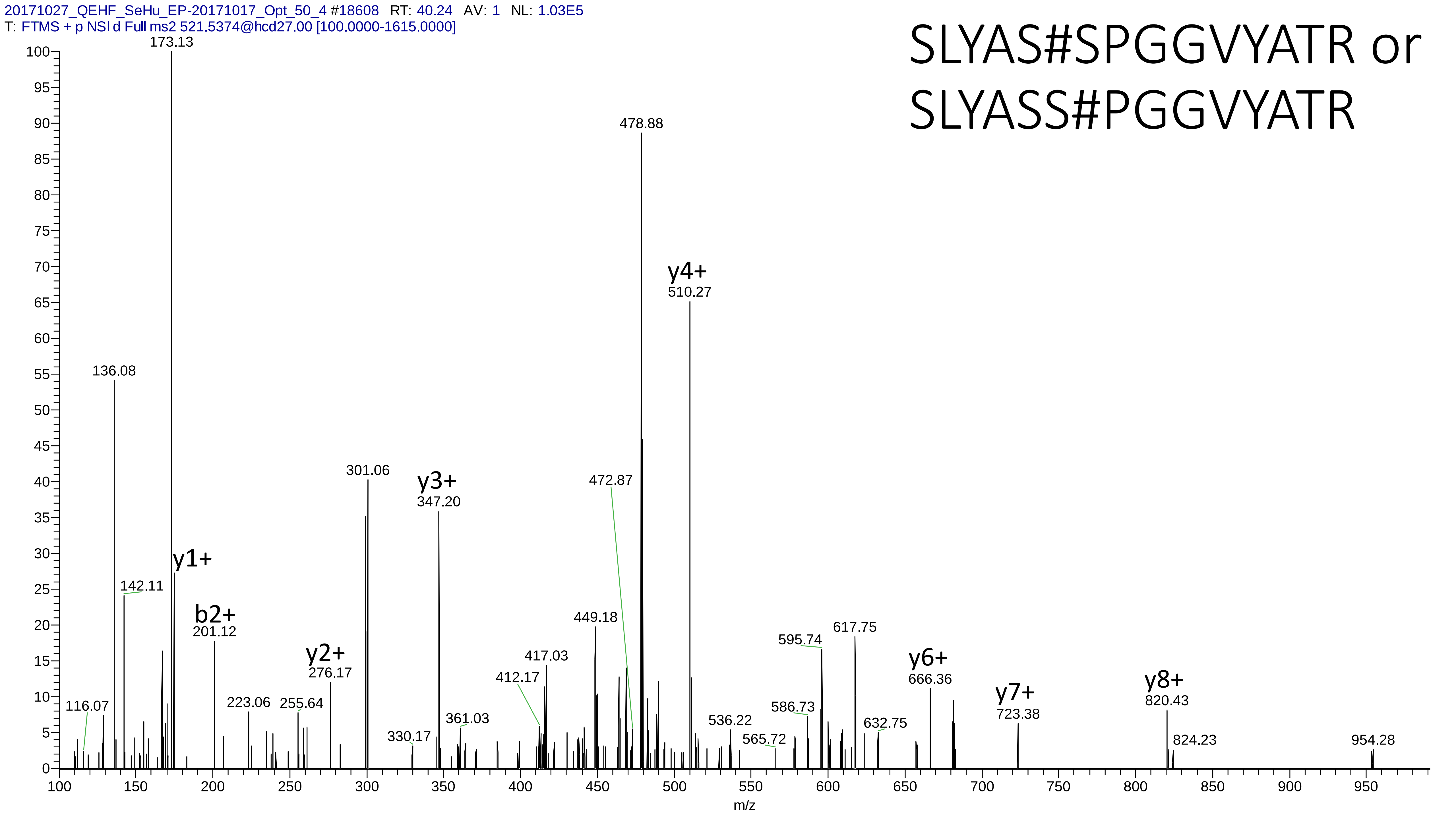
20171027_QEHF_SeHu_EP-20171017_Opt_50_4 #18608 RT: 40.24 AV: 1 NL: 1.03E5
T: FTMS + p NSI d Full ms2 521.5374@hcd27.00 [100.0000-1615.0000]
SLYAS#SPGGVYATR or
SLYASS#PGGVYATR
173.13
478.88
y4+
510.27
136.08
301.06
y3+
347.20
472.87
y1+
142.11
b2+
201.12
449.18
617.75
595.74
417.03
y6+
666.36
y2+
276.17
412.17
y7+
723.38
y8+
820.43
116.07
223.06
255.64
586.73
536.22
632.75
361.03
330.17
565.72
824.23
954.28
m/z

# 16

- Experimental mass: -76.1343
- Unimod information:
- Possible composition: Phosphorylation + Arg-loss, [H(1)O(3)P(1) + H(-12) C(-6) N(-4) O(-1)]
- Sites in Unimod:
- Sites observed:
- Features:  96.5% (137/142) peptides contain Arg at peptide n-term

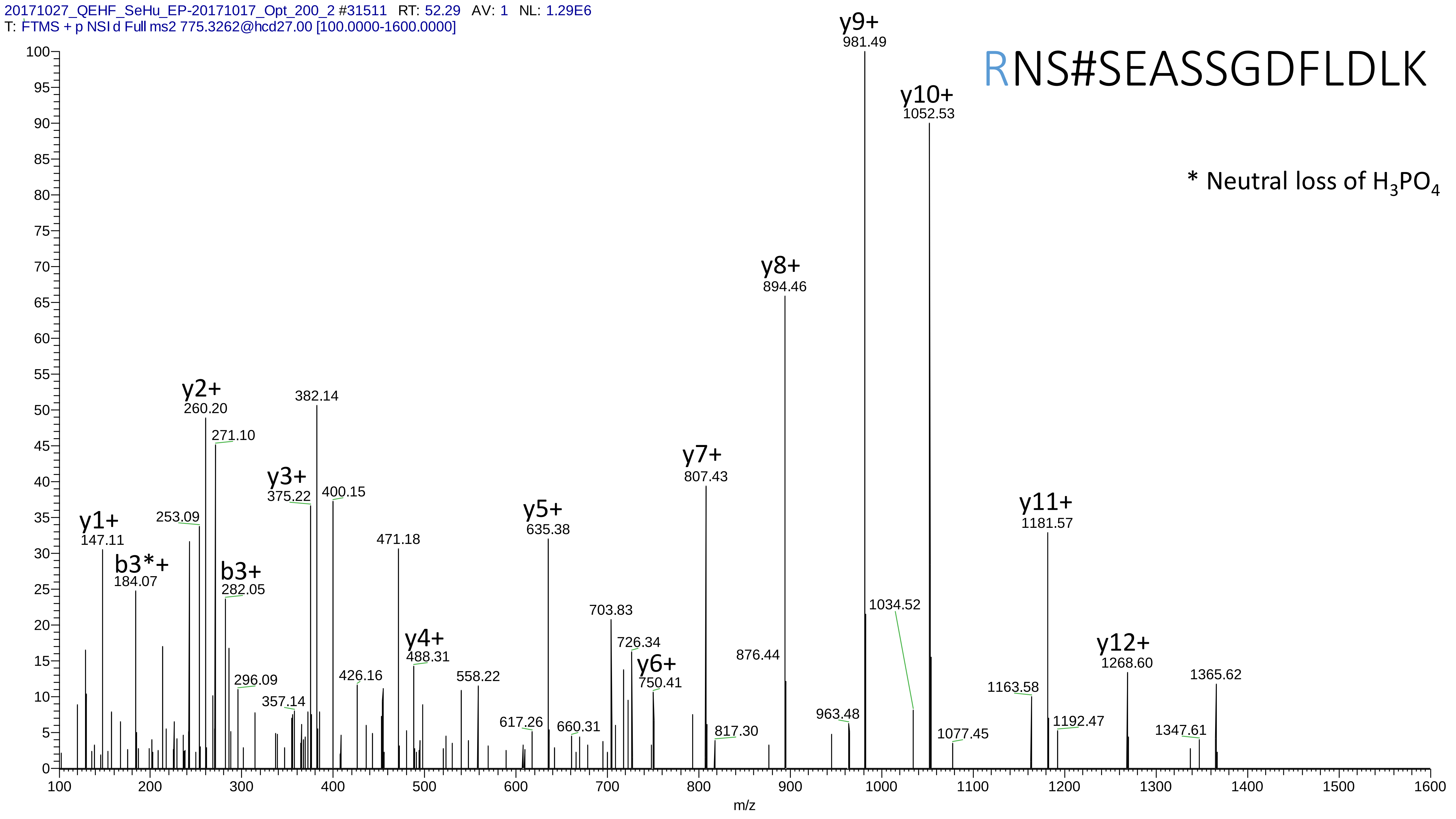

20171027_QEHF_SeHu_EP-20171017_Opt_200_2 #31511 RT: 52.29 AV: 1 NL: 1.29E6
T: FTMS + p NSI d Full ms2 775.3262@hcd27.00 [100.0000-1600.0000]
RNS#SEASSGDFLDLK
* Neutral loss of H3PO4
y1+ 147.11
b3*+ 184.07
253.09
y2+ 260.20
271.10
b3+ 282.05
296.09
357.14
y3+ 375.22
382.14
400.15
426.16
471.18
y4+ 488.31
558.22
617.26
y5+ 635.38
660.31
703.83
726.34
y6+ 750.41
y7+ 807.43
817.30
876.44
y8+ 894.46
963.48
y9+ 981.49
1034.52
y10+ 1052.53
1077.45
1163.58
y11+ 1181.57
1192.47
y12+ 1268.60
1347.61
1365.62
m/z

# 17

- Experimental mass: 122.0141
- Unimod information: Isopropylphospho
- Possible composition: H(7)C(3)O(3)P(1)
- Sites in Unimod: Y;T;S
- Sites observed:

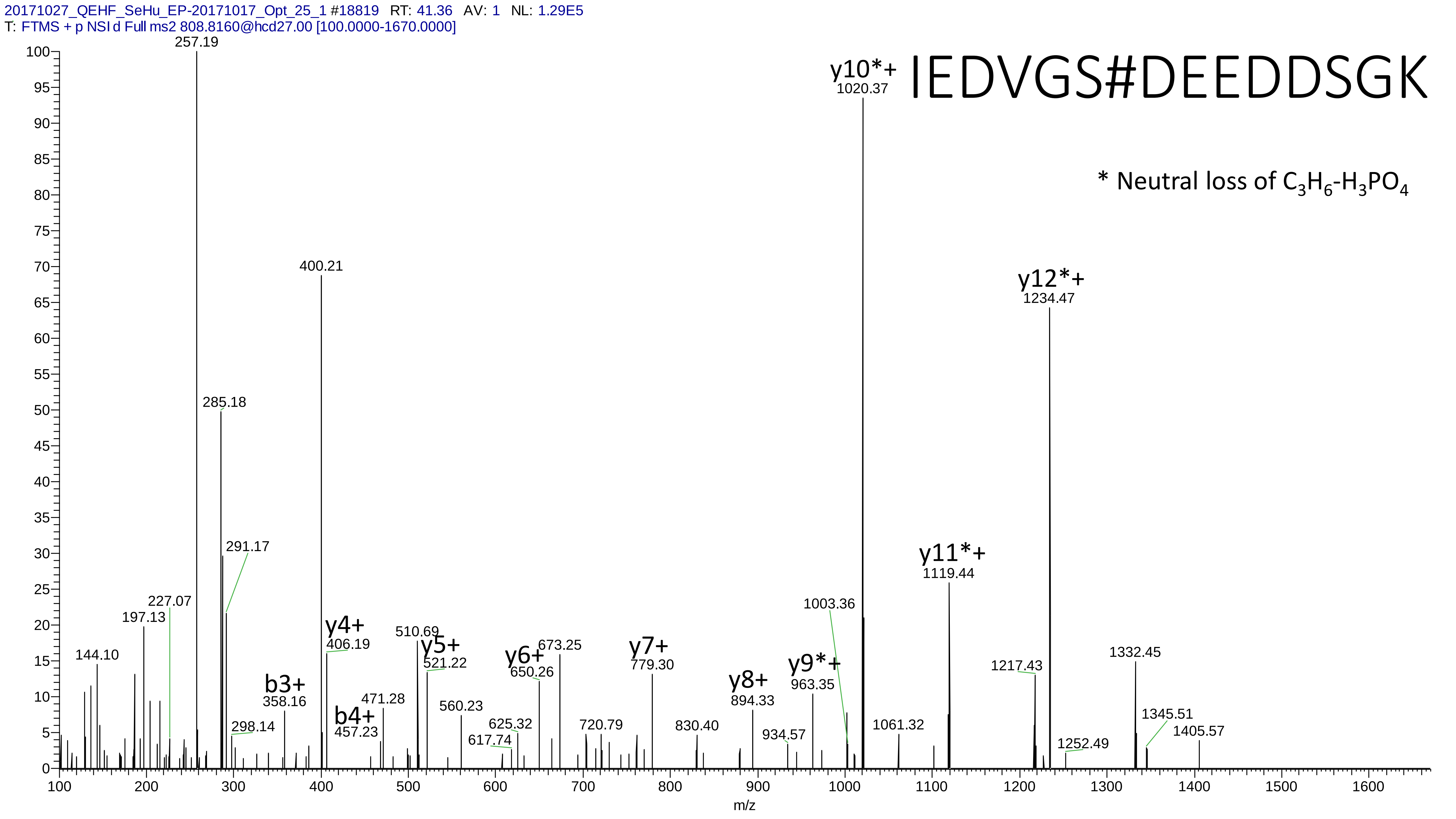
20171027_QEHF_SeHu_EP-20171017_Opt_25_1 #18819 RT: 41.36 AV: 1 NL: 1.29E5
T: FTMS + p NSI d Full ms2 808.8160@hcd27.00 [100.0000-1670.0000]
IEDVGS#DEEDDSGK
* Neutral loss of $C_3H_6$-$H_3PO_4$
257.19
400.21
285.18
291.17
227.07
197.13
144.10
298.14
b3+
358.16
b4+
457.23
y4+
406.19
471.28
510.69
y5+
521.22
560.23
617.74
625.32
y6+
650.26
673.25
720.79
y7+
779.30
830.40
y8+
894.33
934.57
y9*+
963.35
1003.36
y10*+
1020.37
1061.32
y11*+
1119.44
1217.43
y12*+
1234.47
1252.49
1332.45
1345.51
1405.57
m/z

# 18

- Experimental mass: 160.9203
- Unimod information:
- Possible composition: Phosphorylation * 2 + Deamidation, [H(1)O(3)P(1) * 2 + H(-1) N(-1) O]
- Sites in Unimod:
- Sites observed: 99.7% (302/303) peptides contain equal or more than two Ser/Thr

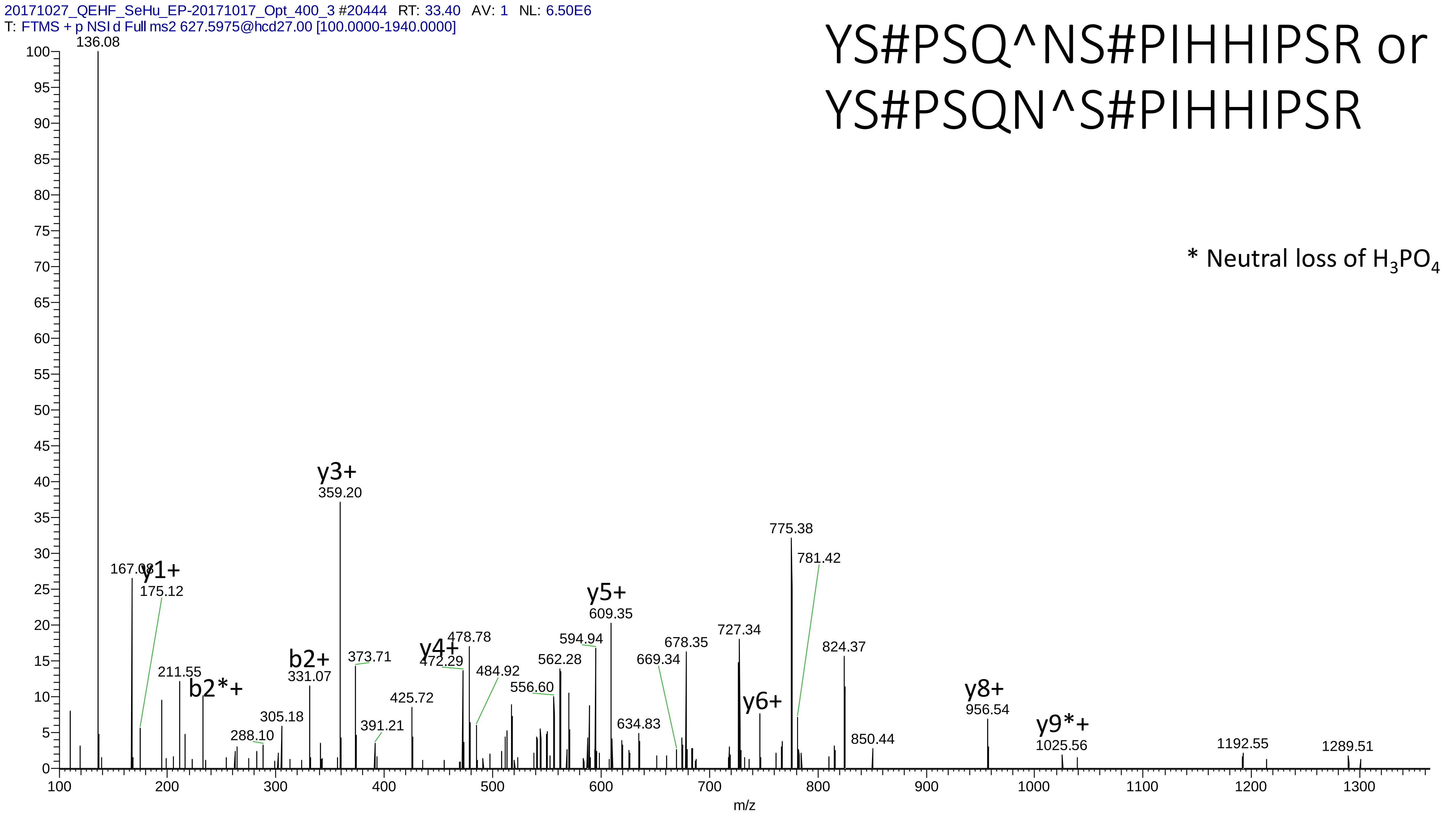
20171027_QEHF_SeHu_EP-20171017_Opt_400_3 #20444 RT: 33.40 AV: 1 NL: 6.50E6
T: FTMS + p NSI d Full ms2 627.5975@hcd27.00 [100.0000-1940.0000]
YS#PSQ^NS#PIHHIPSR or
YS#PSQN^S#PIHHIPSR
* Neutral loss of H3PO4
136.08
167.08
y1+
175.12
211.55
b2*+
288.10
305.18
b2+
331.07
y3+
359.20
373.71
391.21
425.72
y4+
472.29
478.78
484.92
556.60
562.28
594.94
y5+
609.35
634.83
669.34
678.35
727.34
y6+
775.38
781.42
824.37
850.44
y8+
956.54
y9*+
1025.56
1192.55
1289.51
m/z

# 19

- Experimental mass: 197.8807
- Unimod information:
- Possible composition: Phosphorylation * 2 + Cation:Ca[II], [H(1)O(3)P(1) * 2 + H(-2) Ca]
- Sites in Unimod:
- Sites observed:
- Features: because of the high charge state, the fragment information is insufficient for the determination of the modification sites

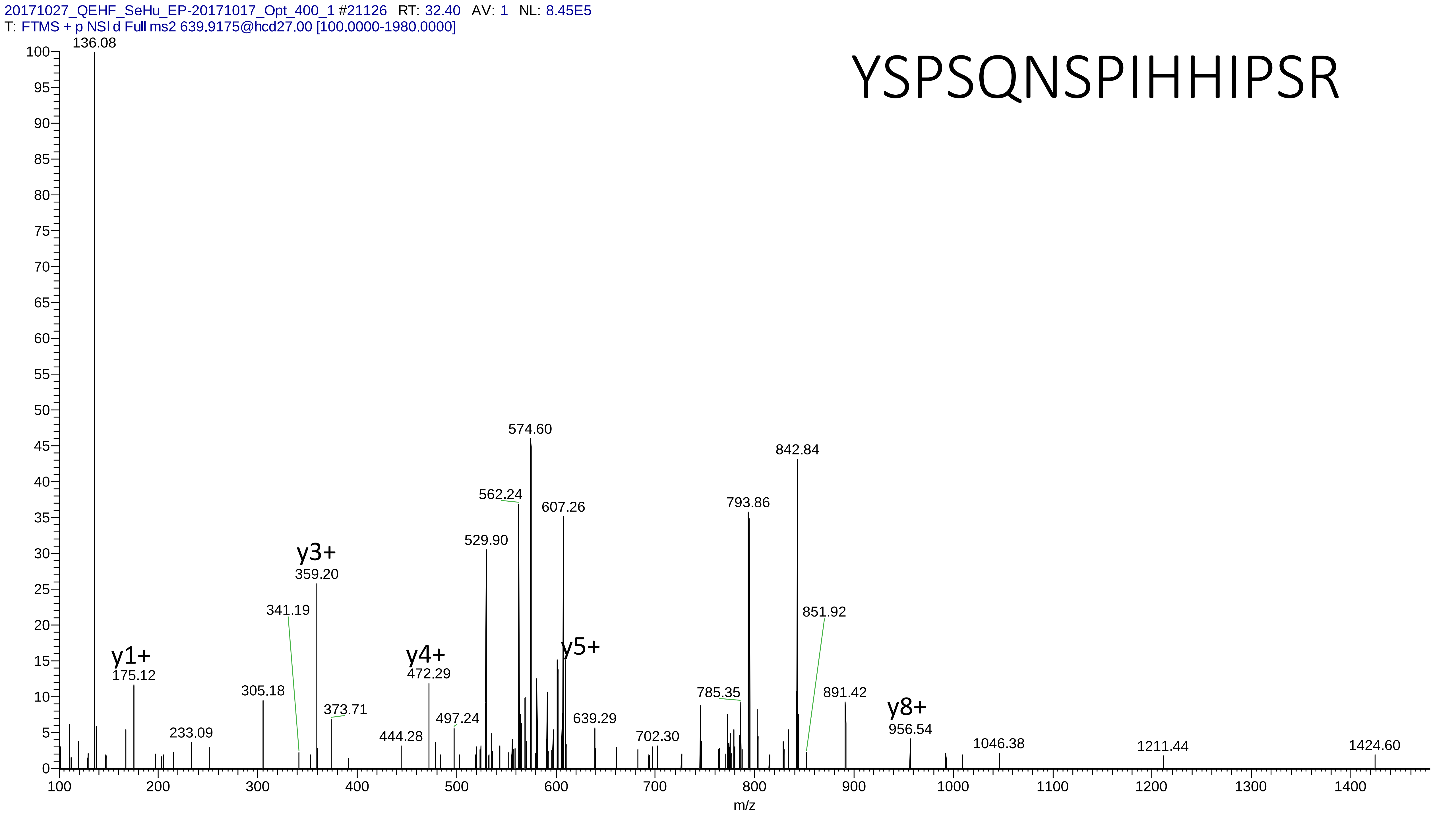

20171027_QEHF_SeHu_EP-20171017_Opt_400_1 #21126 RT: 32.40 AV: 1 NL: 8.45E5
T: FTMS + p NSI d Full ms2 639.9175@hcd27.00 [100.0000-1980.0000]
YSPSQNSPIHHIPSR
136.08
574.60
842.84
562.24
607.26
793.86
529.90
y3+
359.20
341.19
851.92
y1+
175.12
y4+
472.29
y5+
305.18
785.35
891.42
373.71
y8+
497.24
639.29
956.54
233.09
444.28
702.30
1046.38
1211.44
1424.60
m/z

# 20

- Experimental mass: 97.9865
- Unimod information:
- Possible composition:
- Sites in Unimod:
- Sites observed:
- Features: because of long peptide sequence and high charge state, the fragment information is insufficient for the determination of the modification

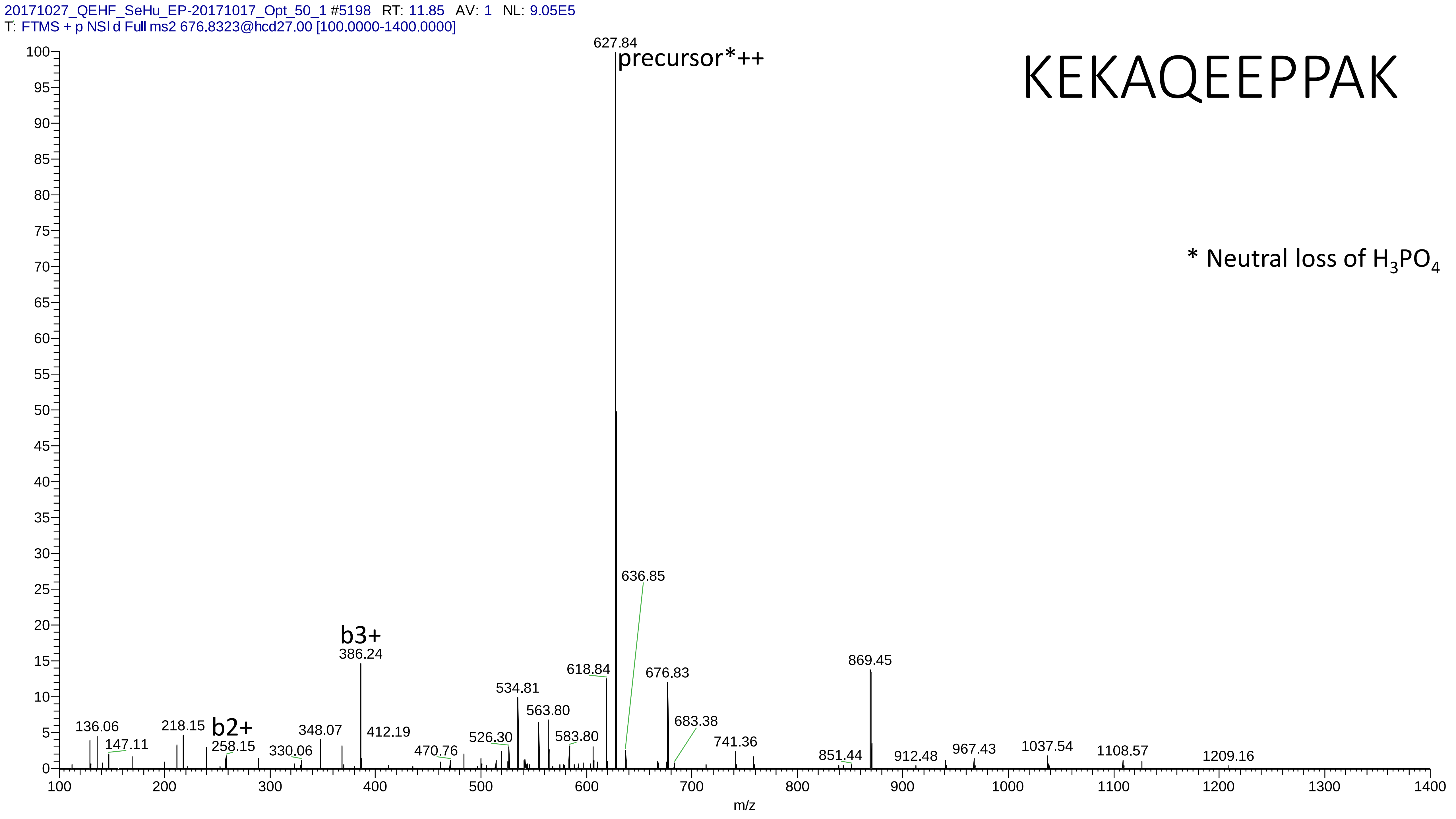

20171027_QEHF_SeHu_EP-20171017_Opt_50_1 #5198 RT: 11.85 AV: 1 NL: 9.05E5
T: FTMS + p NSI d Full ms2 676.8323@hcd27.00 [100.0000-1400.0000]
KEKAQEEPPAK
* Neutral loss of H3PO4
627.84
precursor*++
636.85
b3+
386.24
618.84
869.45
676.83
534.81
563.80
683.38
b2+
136.06
218.15
348.07
412.19
147.11
258.15
330.06
470.76
526.30
583.80
741.36
851.44
912.48
967.43
1037.54
1108.57
1209.16
m/z

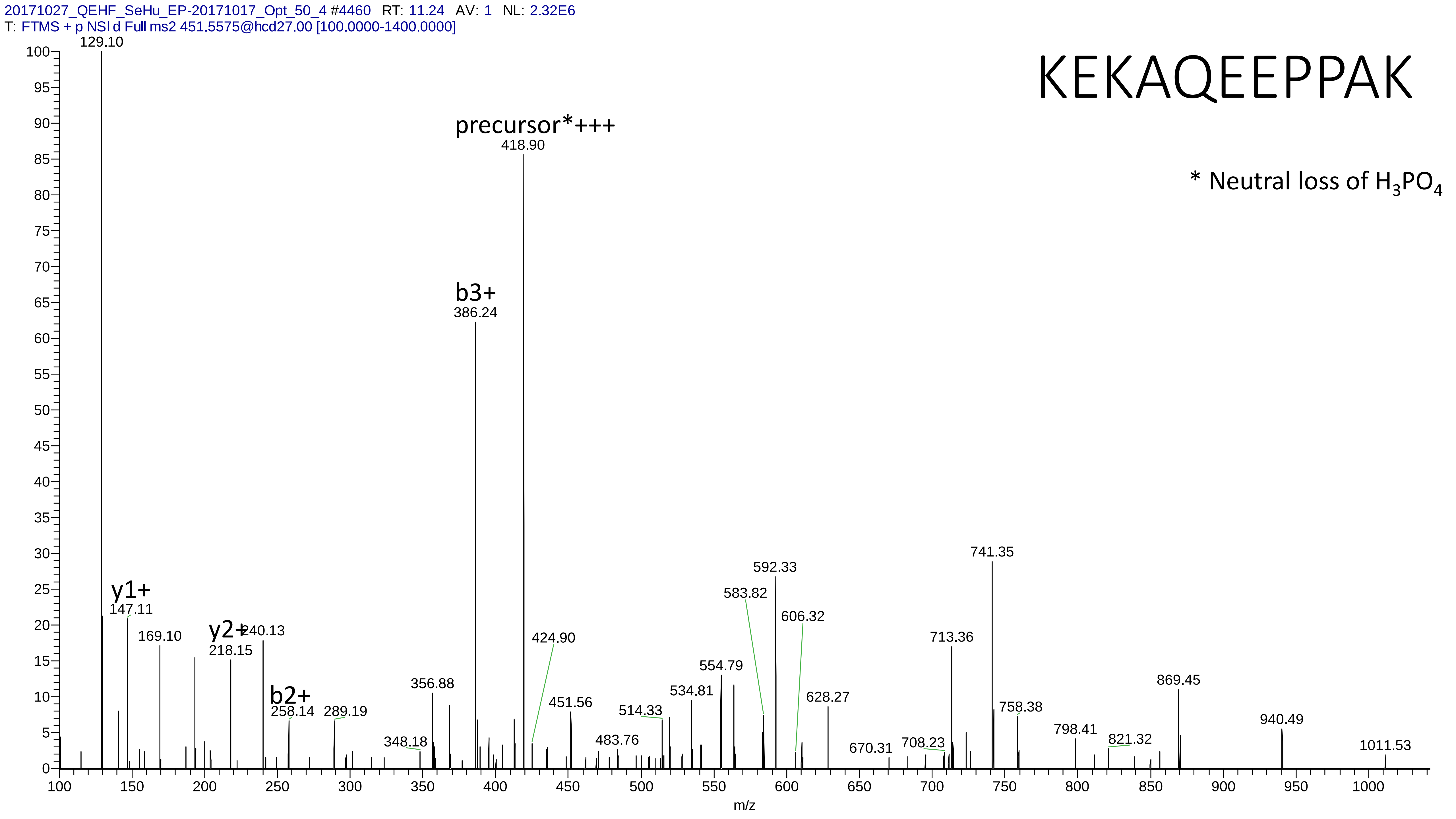

20171027_QEHF_SeHu_EP-20171017_Opt_50_4 #4460 RT: 11.24 AV: 1 NL: 2.32E6
T: FTMS + p NSI d Full ms2 451.5575@hcd27.00 [100.0000-1400.0000]
KEKAQEEPPAK
* Neutral loss of H3PO4
precursor*+++
418.90
b3+
386.24
129.10
y1+
147.11
169.10
y2+
218.15
240.13
b2+
258.14
289.19
348.18
356.88
424.90
451.56
483.76
514.33
534.81
554.79
583.82
592.33
606.32
628.27
670.31
708.23
713.36
741.35
758.38
798.41
821.32
869.45
940.49
1011.53
m/z

# 21

- Experimental mass: 117.9433
- Unimod information:
- Possible composition: Phosphorylation + Oxidation + Cation:Na, [H(1)O(3)P(1) + O(1) + H(-1) Na]
- Sites in Unimod:
- Sites observed:

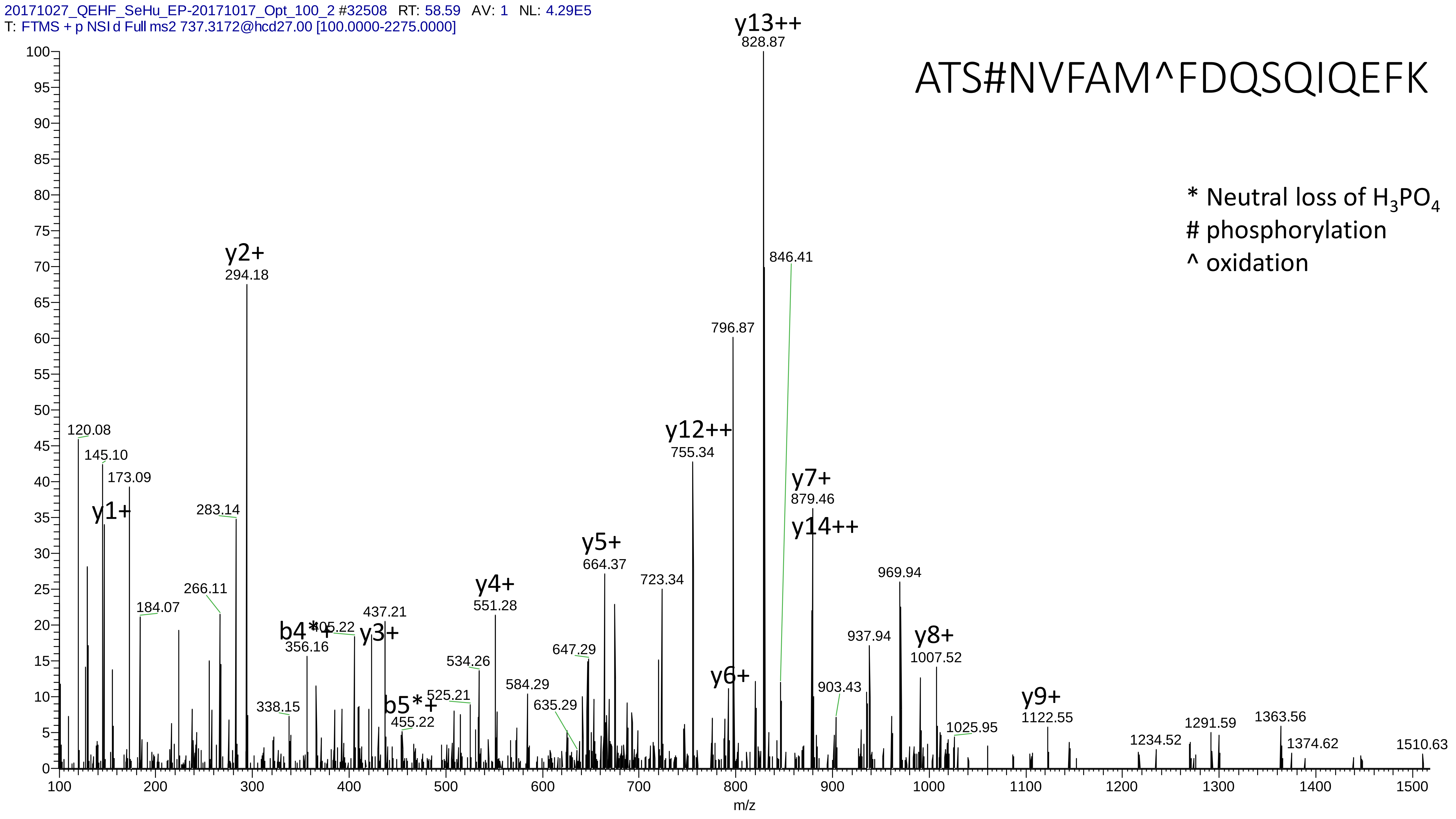

20171027_QEHF_SeHu_EP-20171017_Opt_100_2 #32508 RT: 58.59 AV: 1 NL: 4.29E5
T: FTMS + p NSI d Full ms2 737.3172@hcd27.00 [100.0000-2275.0000]
ATS#NVFAM^FDQSQIQEFK
* Neutral loss of H3PO4
### phosphorylation
^ oxidation
y13++
828.87
846.41
y2+
294.18
796.87
y12++
755.34
120.08
145.10
173.09
y1+
283.14
y7+
879.46
y14++
y5+
664.37
723.34
969.94
266.11
184.07
y4+
551.28
437.21
b4*+
405.22
y3+
356.16
937.94
y8+
1007.52
534.26
647.29
584.29
y6+
903.43
b5*+
525.21
455.22
338.15
635.29
y9+
1122.55
1025.95
1291.59
1363.56
1234.52
1374.62
1510.63
m/z

# 22

- Experimental mass: 257.7182
- Unimod information:
- Possible composition:
- Sites in Unimod:
- Sites observed:
- Features: because of long peptide sequence and high charge state, the fragment information is insufficient for the determination of the modification

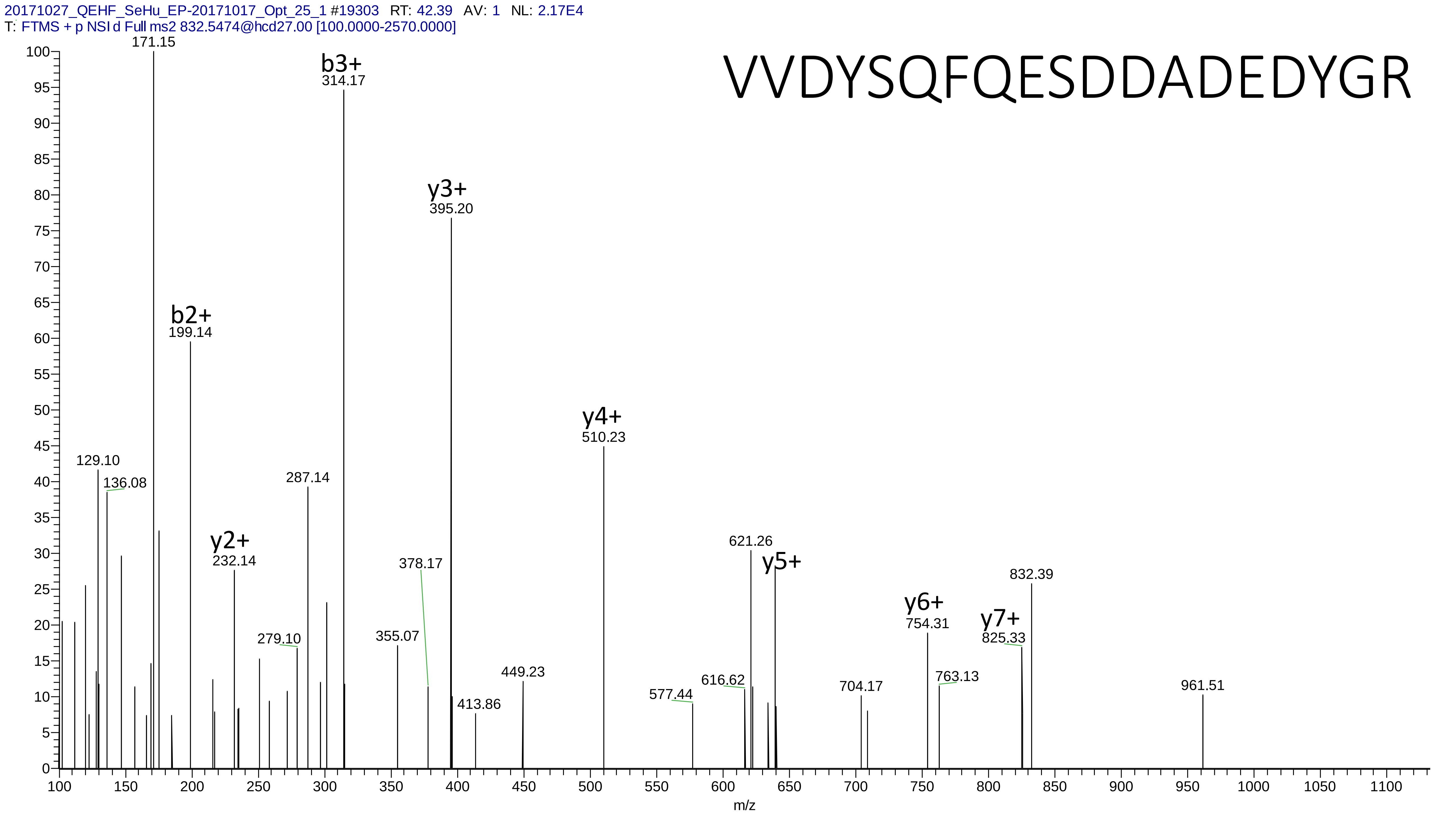
20171027_QEHF_SeHu_EP-20171017_Opt_25_1 #19303 RT: 42.39 AV: 1 NL: 2.17E4
T: FTMS + p NSI d Full ms2 832.5474@hcd27.00 [100.0000-2570.0000]
VVDYSQFQESDDADEDYGR
171.15
b3+
314.17
y3+
395.20
b2+
199.14
y4+
510.23
129.10
136.08
287.14
y2+
232.14
378.17
621.26
y5+
832.39
y6+
754.31
y7+
825.33
279.10
355.07
449.23
763.13
616.62
577.44
704.17
961.51
413.86
m/z

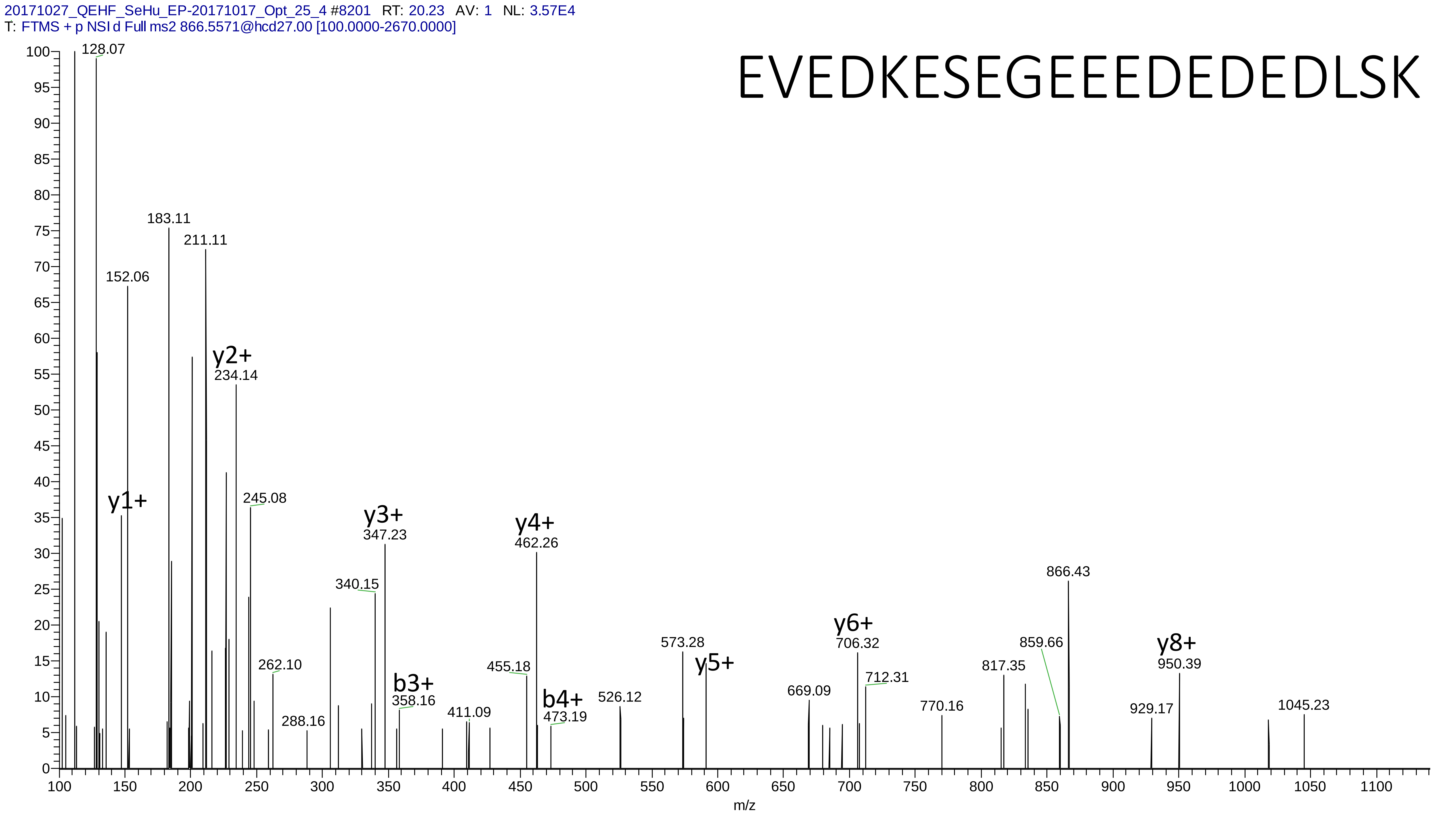
20171027_QEHF_SeHu_EP-20171017_Opt_25_4 #8201 RT: 20.23 AV: 1 NL: 3.57E4
T: FTMS + p NSI d Full ms2 866.5571@hcd27.00 [100.0000-2670.0000]
EVEDKESEGEEEDEDEDLSK
128.07
183.11
211.11
152.06
y2+
234.14
y1+
245.08
y3+
347.23
y4+
462.26
340.15
866.43
y6+
706.32
573.28
859.66
y8+
950.39
262.10
b3+
358.16
455.18
y5+
712.31
817.35
669.09
b4+
473.19
526.12
411.09
288.16
770.16
929.17
1045.23
m/z

# 23

- Experimental mass: 213.8526
- Unimod information:
- Possible composition: Phosphorylation * 2 + Cation:Fe[II], [H(1)O(3)P(1) * 2 + H(-2) Fe]
- Sites in Unimod:
- Sites observed:
- Features: 100.0% (129/129) peptides contain equal or more than two Ser/Thr, cation modified sites are not observed

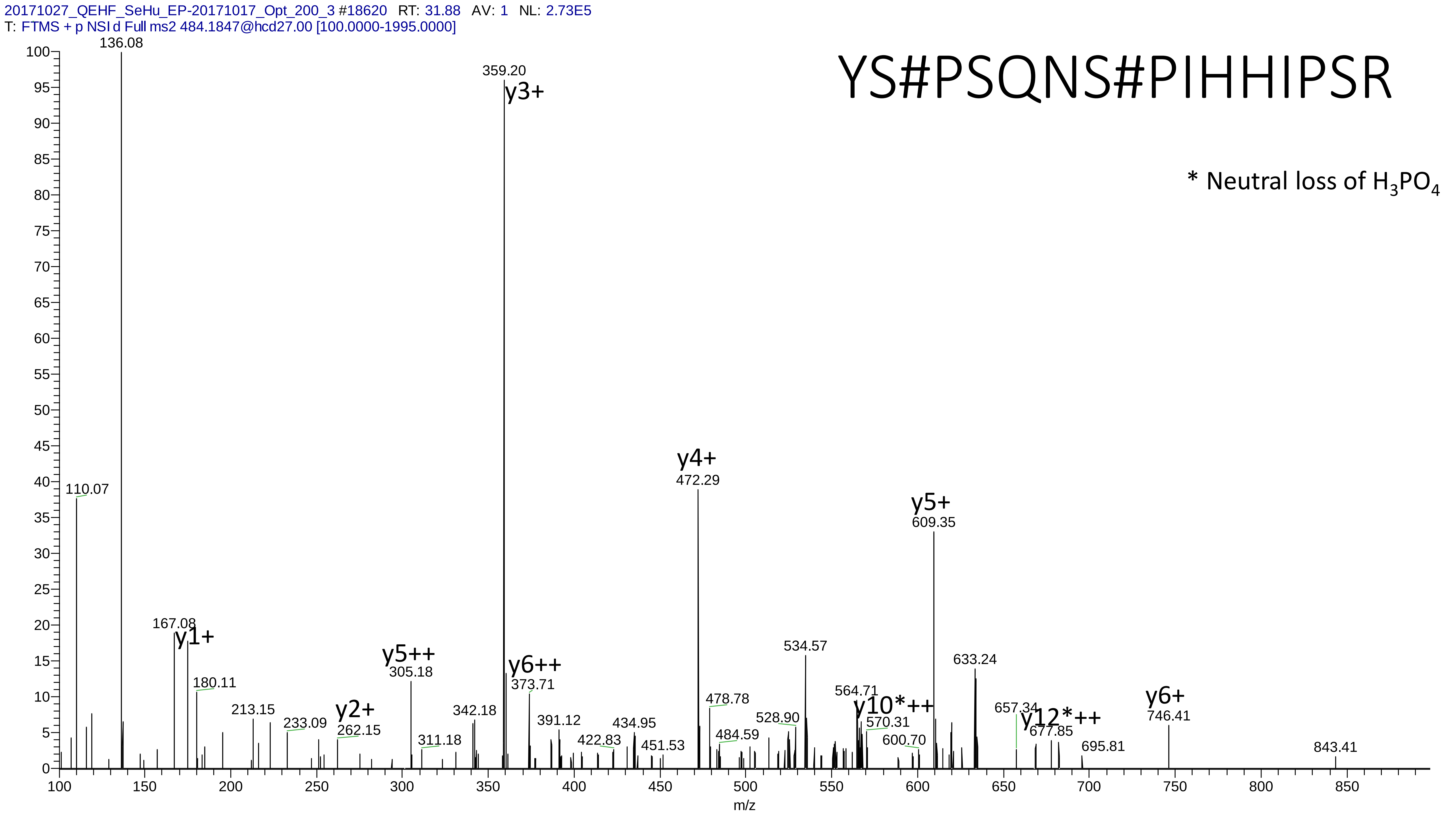

20171027_QEHF_SeHu_EP-20171017_Opt_200_3 #18620 RT: 31.88 AV: 1 NL: 2.73E5
T: FTMS + p NSI d Full ms2 484.1847@hcd27.00 [100.0000-1995.0000]
YS#PSQNS#PIHHIPSR
* Neutral loss of H3PO4
136.08
359.20
y3+
110.07
167.08
y1+
180.11
213.15
233.09
y2+
262.15
y5++
305.18
311.18
342.18
y6++
373.71
391.12
422.83
434.95
451.53
y4+
472.29
478.78
484.59
528.90
534.57
564.71
y10*++
570.31
600.70
y5+
609.35
633.24
657.34
y12*++
677.85
695.81
y6+
746.41
843.41
m/z

# 24

- Experimental mass: 236.0677
- Unimod information:
- Possible composition: Phosphorylation + Arg, [H(1)O(3)P(1) + H(12) C(6) N(4) O]
- Sites in Unimod:
- Sites observed:

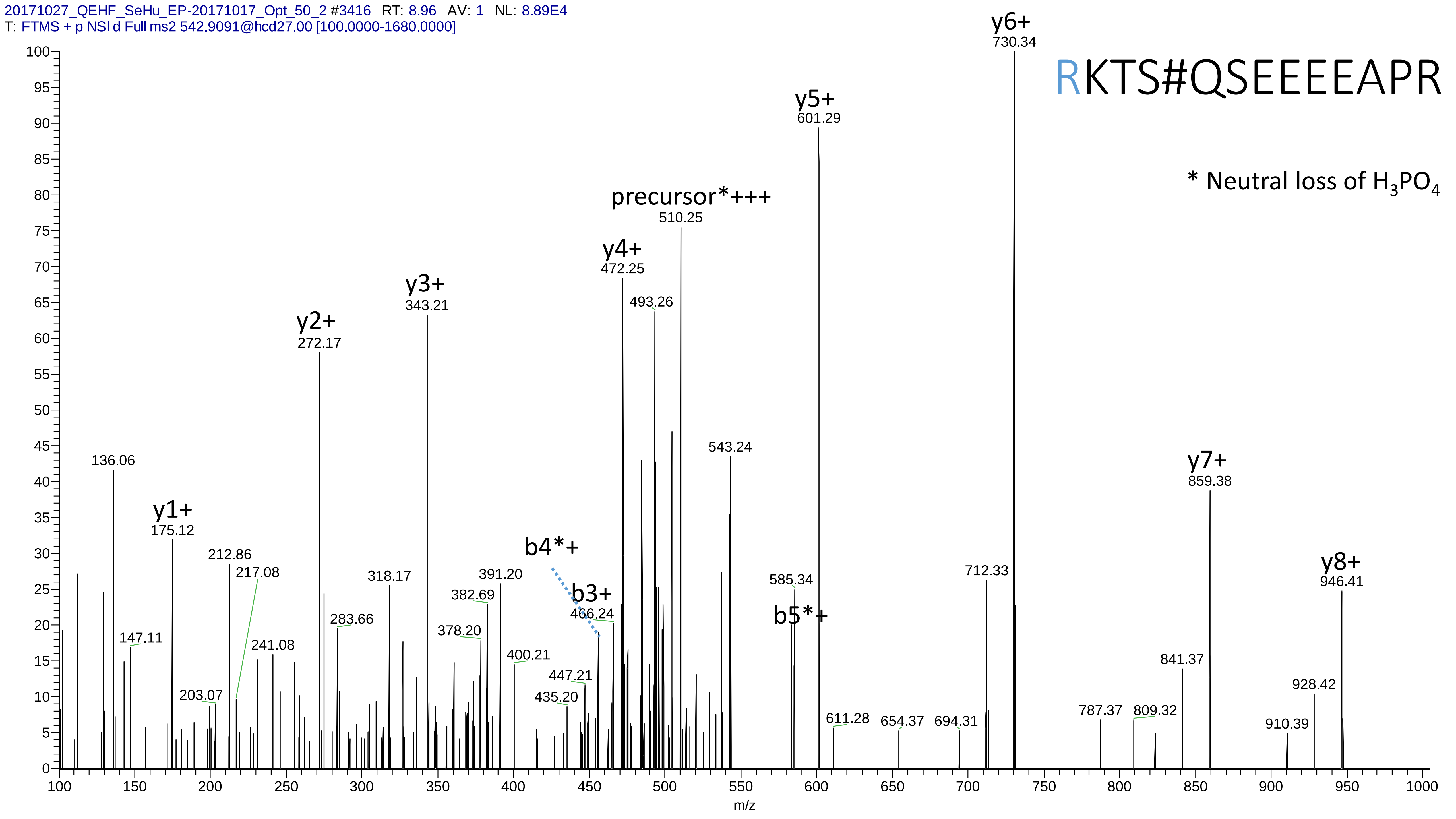

20171027_QEHF_SeHu_EP-20171017_Opt_50_2 #3416 RT: 8.96 AV: 1 NL: 8.89E4
T: FTMS + p NSI d Full ms2 542.9091@hcd27.00 [100.0000-1680.0000]
RKTS#QSEEEEAPR
* Neutral loss of $H_3PO_4$
y6+ 730.34
y5+ 601.29
precursor*+++ 510.25
y4+ 472.25
493.26
y3+ 343.21
y2+ 272.17
543.24
136.06
y7+ 859.38
y1+ 175.12
212.86
217.08
b4*+
318.17
391.20
382.69
b3+ 466.24
585.34
b5*+
712.33
y8+ 946.41
283.66
378.20
147.11
241.08
400.21
447.21
841.37
203.07
435.20
928.42
611.28
654.37
694.31
787.37
809.32
910.39
m/z

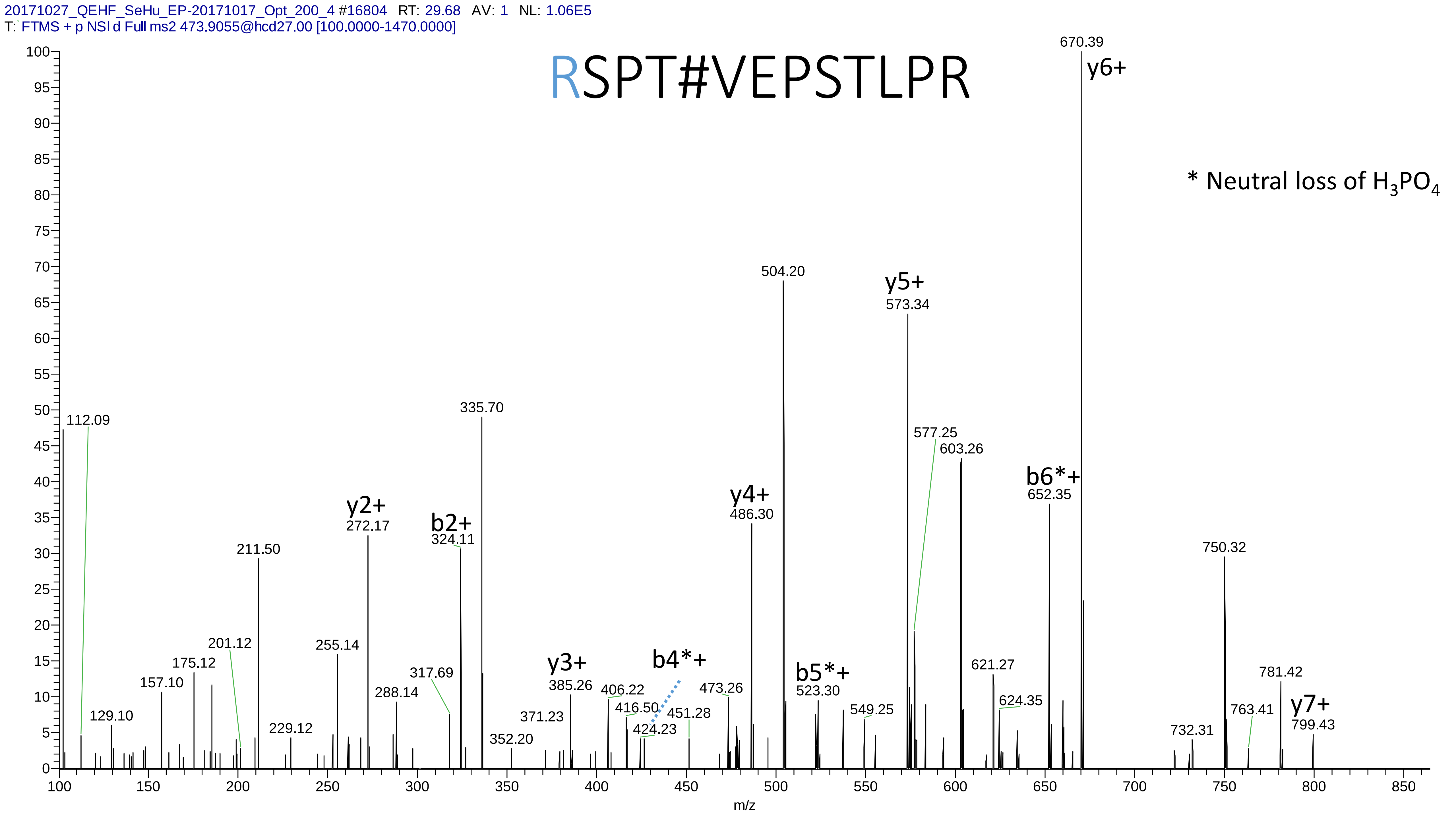
20171027_QEHF_SeHu_EP-20171017_Opt_200_4 #16804 RT: 29.68 AV: 1 NL: 1.06E5
T: FTMS + p NSI d Full ms2 473.9055@hcd27.00 [100.0000-1470.0000]
RSPT#VEPSTLPR
* Neutral loss of $H_3PO_4$
670.39
y6+
504.20
y5+
573.34
577.25
603.26
b6*+
652.35
750.32
781.42
763.41
y7+
799.43
732.31
621.27
624.35
549.25
b5*+
523.30
y4+
486.30
473.26
b4*+
451.28
424.23
416.50
406.22
y3+
385.26
371.23
352.20
335.70
b2+
324.11
317.69
y2+
272.17
288.14
255.14
229.12
211.50
201.12
175.12
157.10
129.10
112.09
m/z

# 25

- Experimental mass: 15.9955
- Unimod information: Oxidation
- Possible composition: could also be Dioxidation replace of oxidation, or Cys->Dha replace of Carbamidomethyl on Cys
- Sites in Unimod: W;H;C;M;R;Y;F;P;N;D;K;cG;U
- Sites observed: W;C

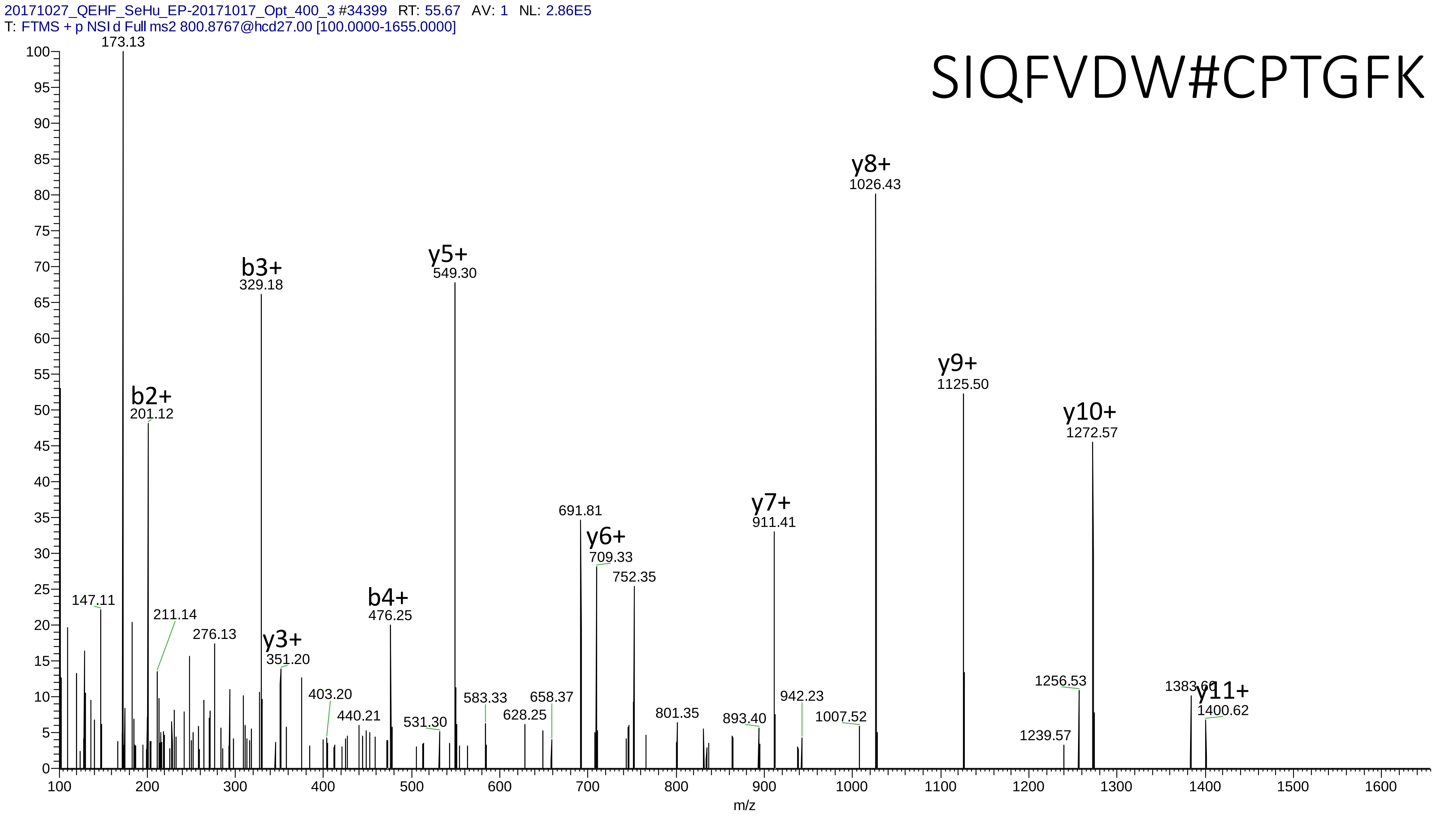

20171027_QEHF_SeHu_EP-20171017_Opt_400_3 #34399 RT: 55.67 AV: 1 NL: 2.86E5
T: FTMS + p NSI d Full ms2 800.8767@hcd27.00 [100.0000-1655.0000]
SIQFVDW#CPTGFK
173.13
y8+
1026.43
b3+
329.18
y5+
549.30
y9+
1125.50
b2+
201.12
y10+
1272.57
691.81
y6+
709.33
y7+
911.41
752.35
147.11
211.14
276.13
y3+
351.20
b4+
476.25
403.20
440.21
531.30
583.33
628.25
658.37
801.35
893.40
942.23
1007.52
1239.57
1256.53
1383.60
y11+
1400.62
m/z

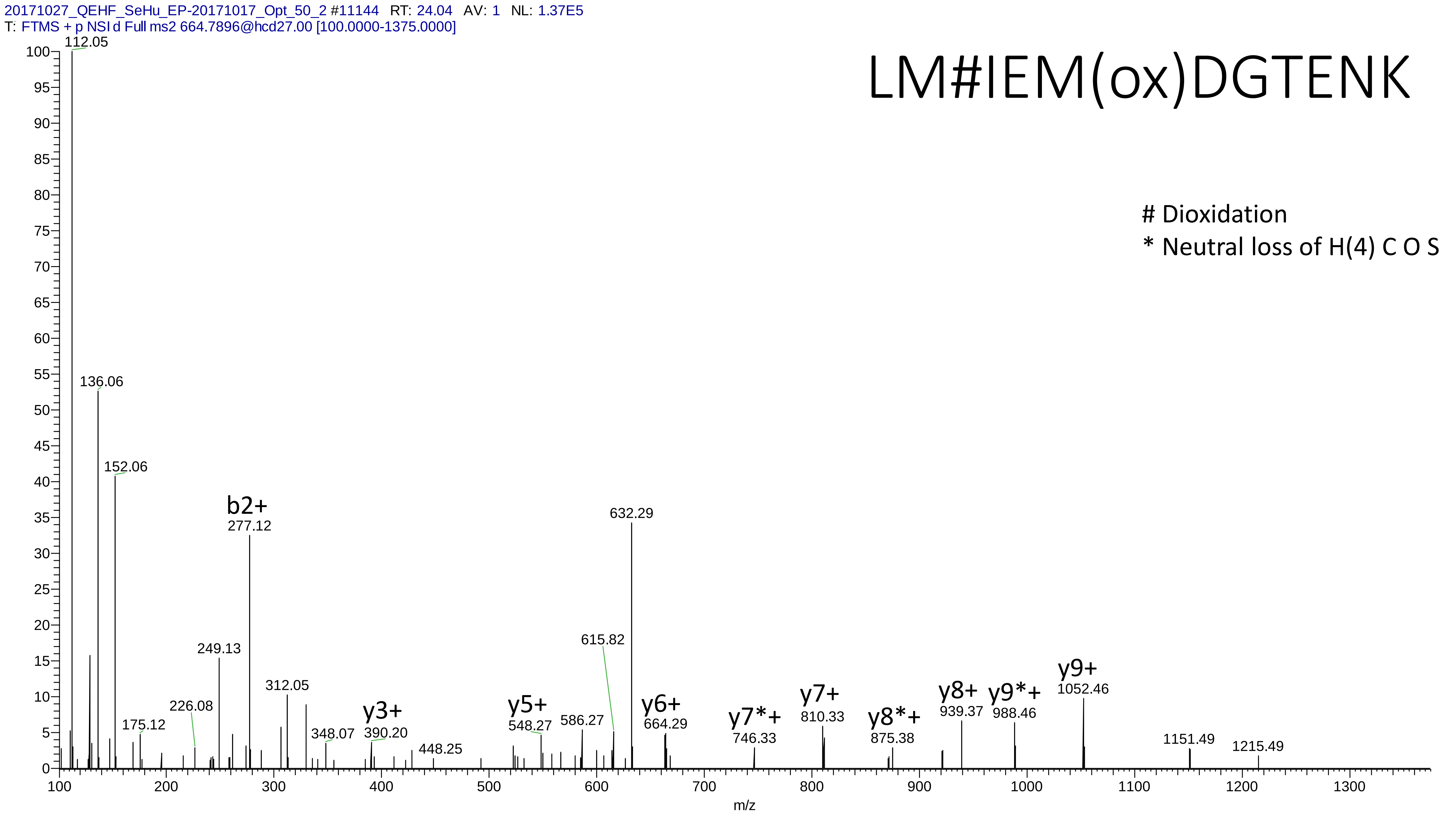
20171027_QEHF_SeHu_EP-20171017_Opt_50_2 #11144 RT: 24.04 AV: 1 NL: 1.37E5
T: FTMS + p NSI d Full ms2 664.7896@hcd27.00 [100.0000-1375.0000]
LM#IEM(ox)DGTENK
### Dioxidation
* Neutral loss of H(4) C O S
112.05
136.06
152.06
b2+
277.12
632.29
615.82
249.13
312.05
226.08
175.12
348.07
y3+
390.20
448.25
y5+
548.27
586.27
y6+
664.29
y7*+
746.33
y7+
810.33
y8*+
875.38
y8+
939.37
y9*+
988.46
y9+
1052.46
1151.49
1215.49
m/z

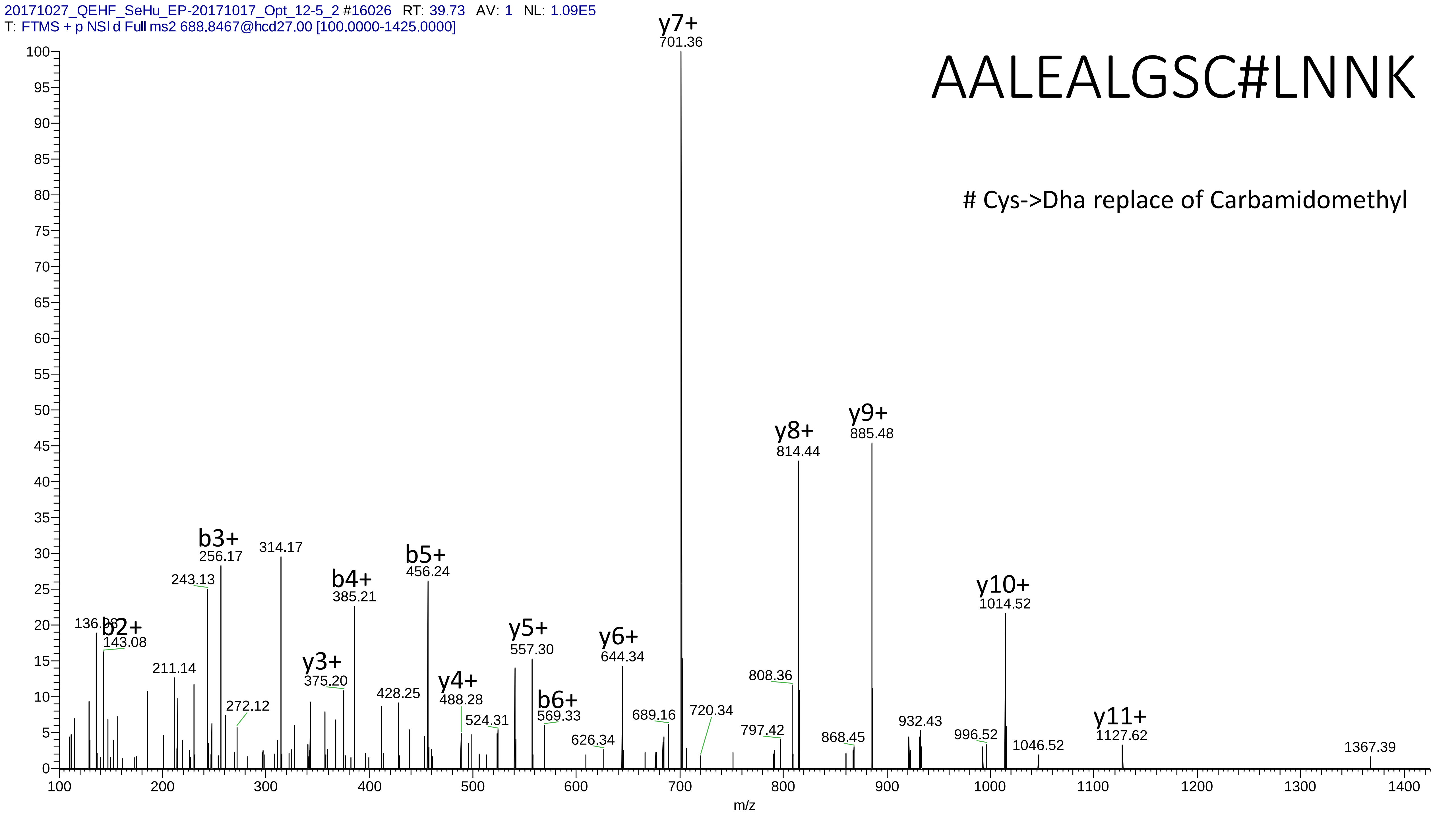
20171027_QEHF_SeHu_EP-20171017_Opt_12-5_2 #16026 RT: 39.73 AV: 1 NL: 1.09E5
T: FTMS + p NSI d Full ms2 688.8467@hcd27.00 [100.0000-1425.0000]
AALEALGSC#LNNK
### Cys->Dha replace of Carbamidomethyl
y7+ 701.36
y8+ 814.44
y9+ 885.48
y10+ 1014.52
y11+ 1127.62
b2+ 143.08
b3+ 256.17
b4+ 385.21
b5+ 456.24
b6+ 569.33
y3+ 375.20
y4+ 488.28
y5+ 557.30
y6+ 644.34
136.08
211.14
243.13
272.12
314.17
428.25
524.31
626.34
689.16
720.34
797.42
808.36
868.45
932.43
996.52
1046.52
1367.39
m/z

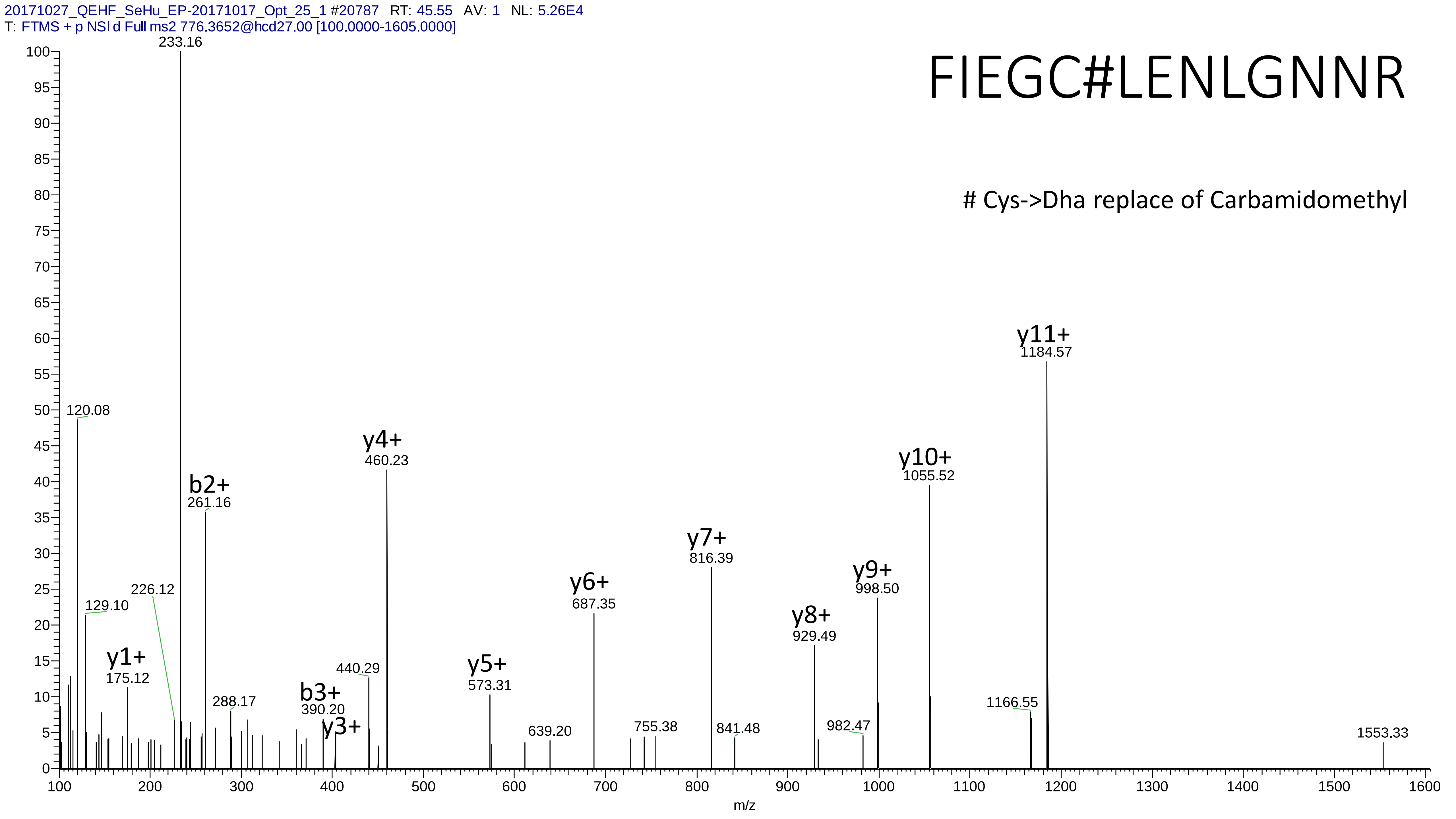
20171027_QEHF_SeHu_EP-20171017_Opt_25_1 #20787 RT: 45.55 AV: 1 NL: 5.26E4
T: FTMS + p NSI d Full ms2 776.3652@hcd27.00 [100.0000-1605.0000]
FIEGC#LENLGNNR
### Cys->Dha replace of Carbamidomethyl
233.16
120.08
129.10
226.12
y1+ 175.12
b2+ 261.16
288.17
b3+ 390.20
y3+
440.29
y4+ 460.23
y5+ 573.31
639.20
y6+ 687.35
755.38
y7+ 816.39
841.48
y8+ 929.49
982.47
y9+ 998.50
y10+ 1055.52
1166.55
y11+ 1184.57
1553.33
m/z

# 26

- Experimental mass: 132.8796
- Unimod information:
- Possible composition: Phosphorylation + Cation:Fe[III], [H(1)O(3)P(1) + H(-3) Fe]
- Sites in Unimod:
- Sites observed: phosphorylation sites determined, cation modified sites are not observed

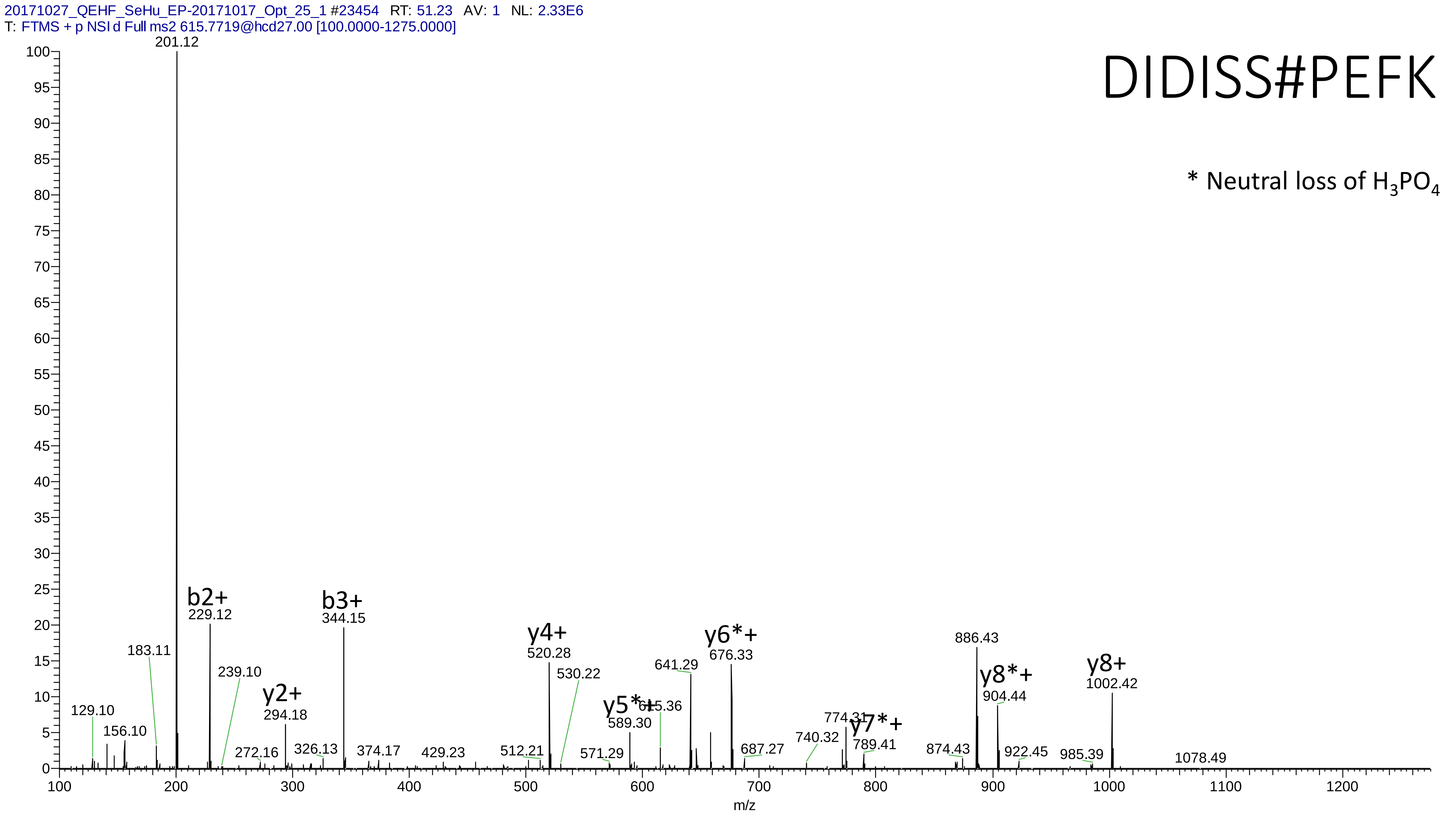
20171027_QEHF_SeHu_EP-20171017_Opt_25_1 #23454 RT: 51.23 AV: 1 NL: 2.33E6
T: FTMS + p NSI d Full ms2 615.7719@hcd27.00 [100.0000-1275.0000]
DIDISS#PEFK
* Neutral loss of $H_3PO_4$
201.12
b2+
229.12
b3+
344.15
y2+
294.18
y4+
520.28
y5*+
589.30
y6*+
676.33
y7*+
789.41
y8*+
904.44
y8+
1002.42
129.10
156.10
183.11
239.10
272.16
326.13
374.17
429.23
512.21
530.22
571.29
615.36
641.29
687.27
740.32
774.31
874.43
886.43
922.45
985.39
1078.49
m/z

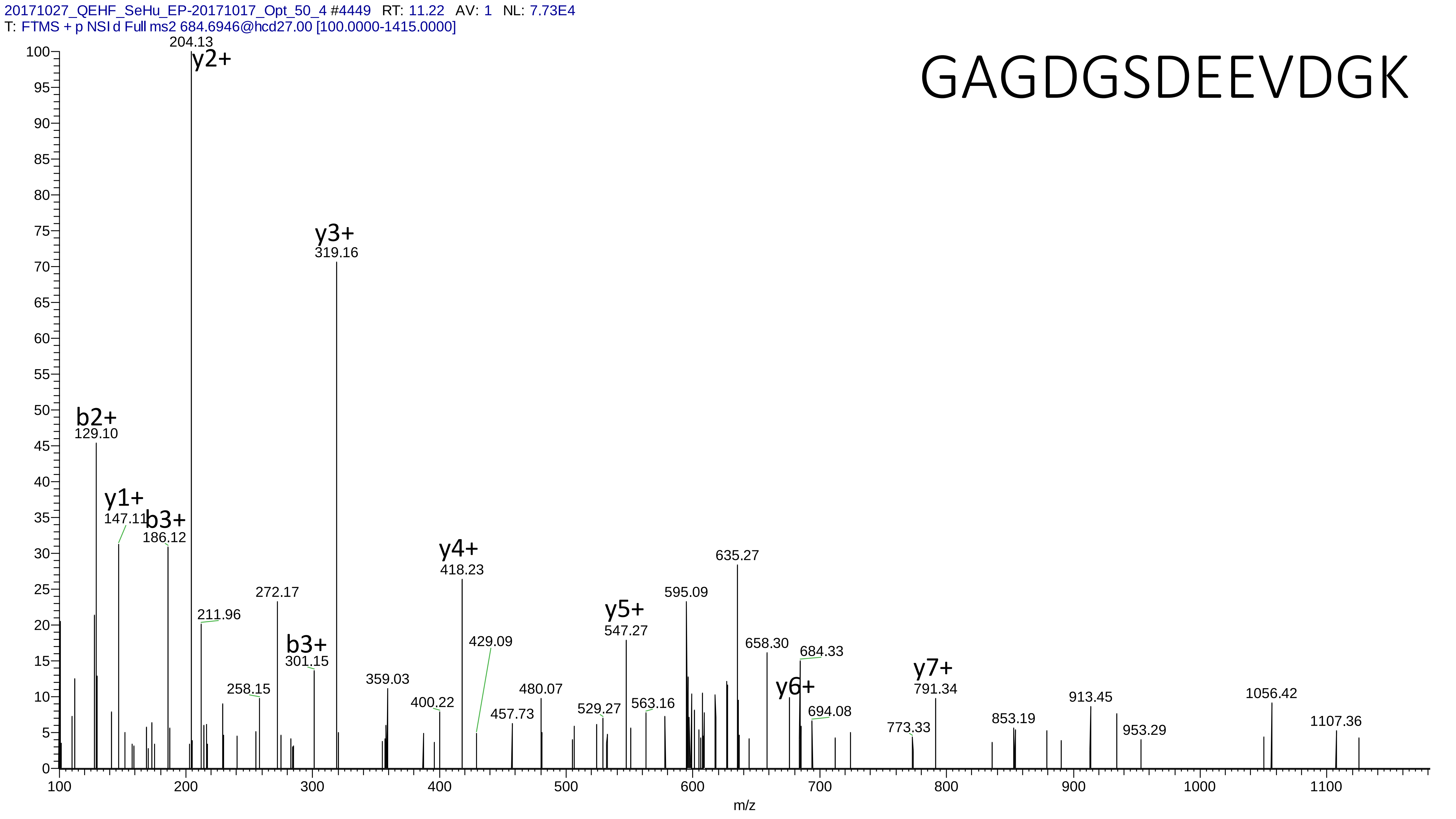
20171027_QEHF_SeHu_EP-20171017_Opt_50_4 #4449 RT: 11.22 AV: 1 NL: 7.73E4
T: FTMS + p NSI d Full ms2 684.6946@hcd27.00 [100.0000-1415.0000]
GAGDGSDEEVDGK
204.13
y2+
y3+
319.16
b2+
129.10
y1+
147.11
b3+
186.12
y4+
418.23
635.27
272.17
595.09
211.96
y5+
547.27
b3+
301.15
429.09
658.30
684.33
y7+
791.34
y6+
359.03
480.07
258.15
1056.42
400.22
529.27
563.16
913.45
694.08
457.73
853.19
1107.36
773.33
953.29
m/z

# 27

- Experimental mass: 31.9914
- Unimod information: Dioxidation
- Possible composition:
- Sites in Unimod: Y;W;F;M;K;R;P;C;U
- Sites observed: W;M
- Features: 41.2% (14/34) peptides contain Trp

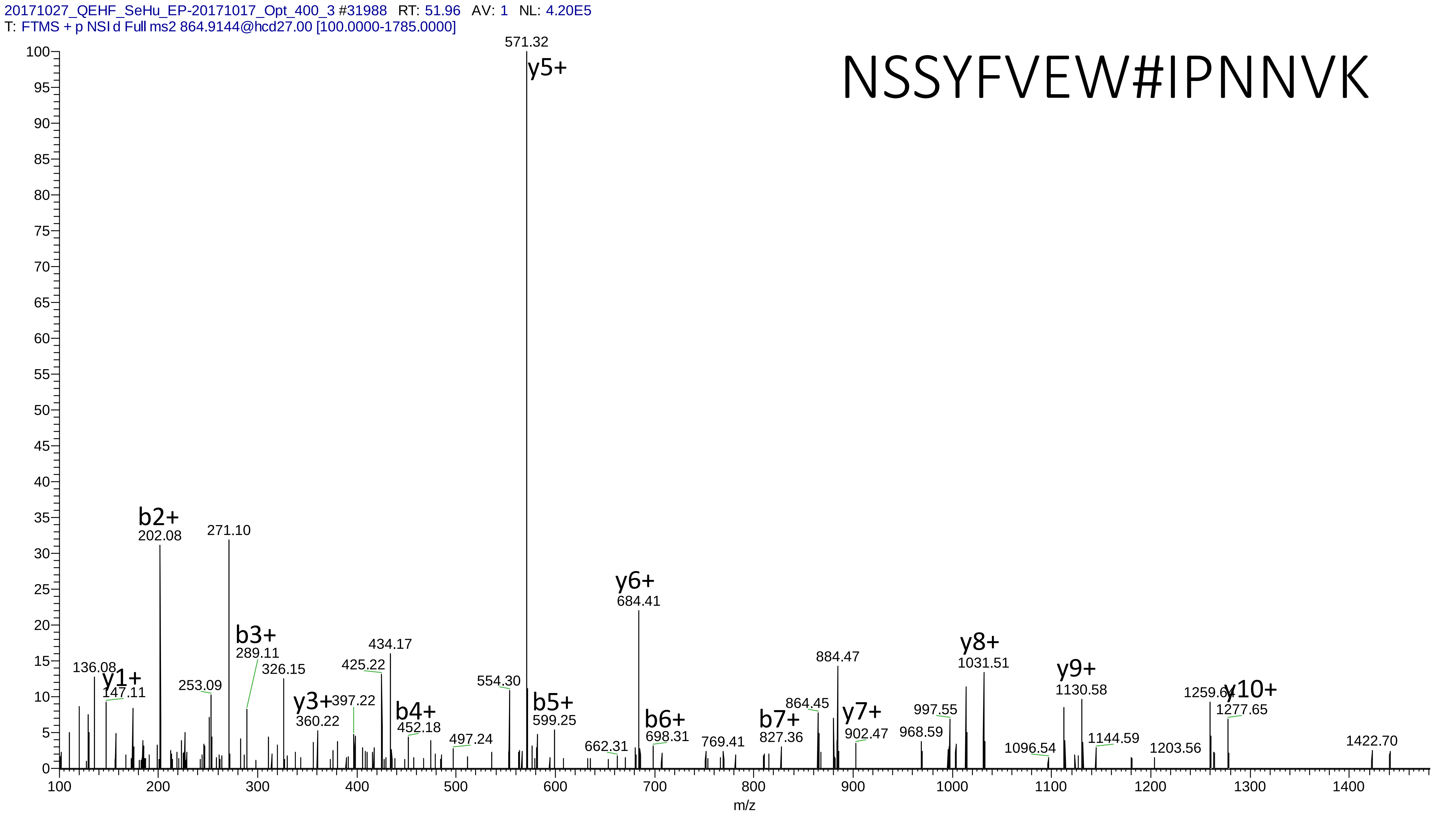
20171027_QEHF_SeHu_EP-20171017_Opt_400_3 #31988 RT: 51.96 AV: 1 NL: 4.20E5
T: FTMS + p NSI d Full ms2 864.9144@hcd27.00 [100.0000-1785.0000]
NSSYFVEW#IPNNVK
571.32
y5+
b2+
202.08
271.10
b3+
289.11
y6+
684.41
136.08
y1+
147.11
253.09
326.15
y3+
360.22
397.22
425.22
434.17
b4+
452.18
497.24
554.30
b5+
599.25
662.31
b6+
698.31
769.41
b7+
827.36
864.45
884.47
y7+
902.47
968.59
997.55
y8+
1031.51
1096.54
y9+
1130.58
1144.59
1203.56
1259.64
y10+
1277.65
1422.70
m/z

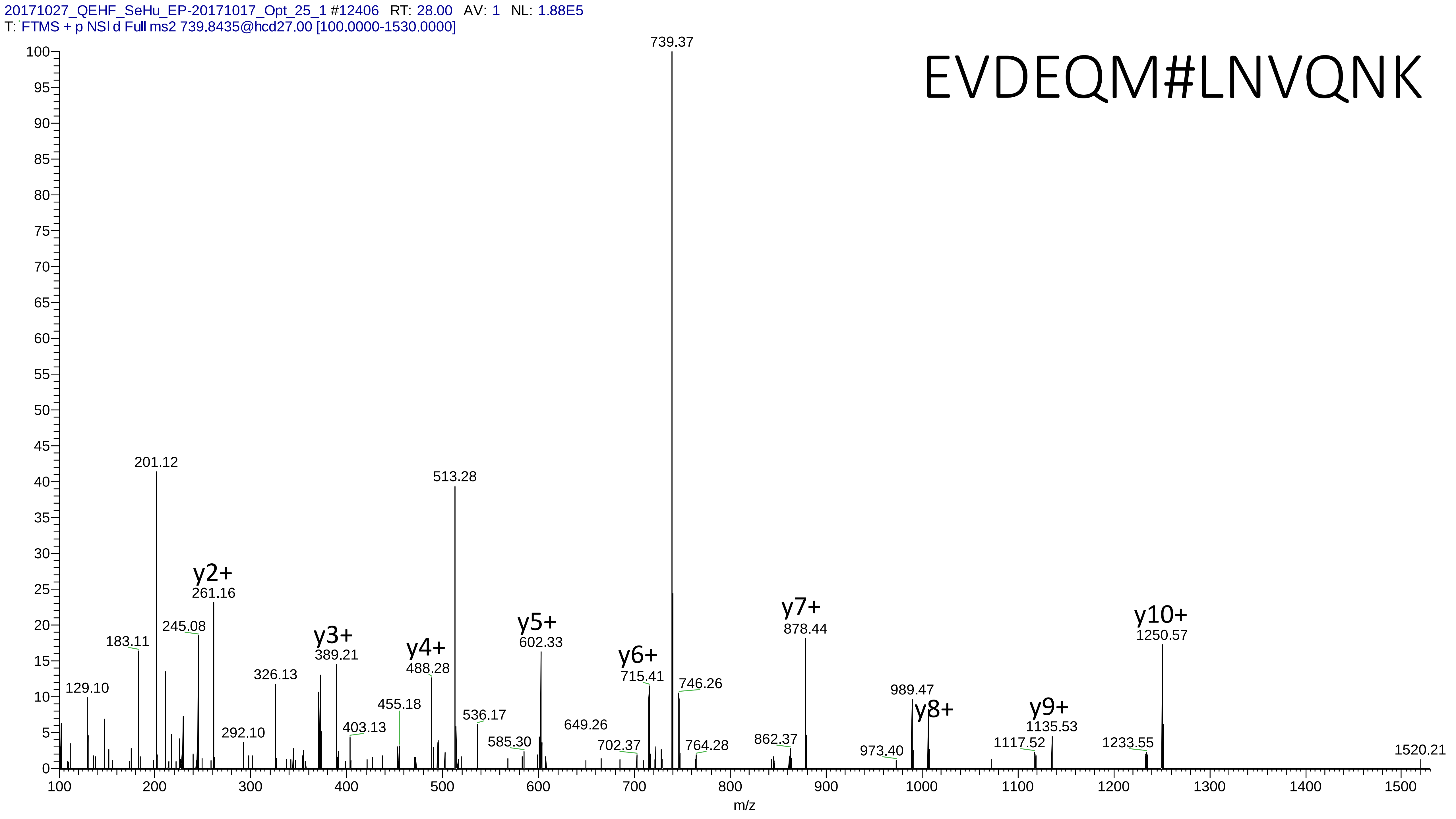

20171027_QEHF_SeHu_EP-20171017_Opt_25_1 #12406 RT: 28.00 AV: 1 NL: 1.88E5
T: FTMS + p NSI d Full ms2 739.8435@hcd27.00 [100.0000-1530.0000]
EVDEQM#LNVQNK
739.37
201.12
513.28
y2+
261.16
245.08
183.11
129.10
326.13
292.10
y3+
389.21
403.13
455.18
y4+
488.28
536.17
y5+
602.33
585.30
649.26
y6+
715.41
702.37
746.26
764.28
y7+
878.44
862.37
989.47
y8+
973.40
y9+
1135.53
1117.52
y10+
1250.57
1233.55
1520.21
m/z

# 28

- Experimental mass: 78.9355
- Unimod information:
- Possible composition:
- Sites in Unimod:
- Sites observed:
- Features: neutral loss of $H_3PO_4$ are observed

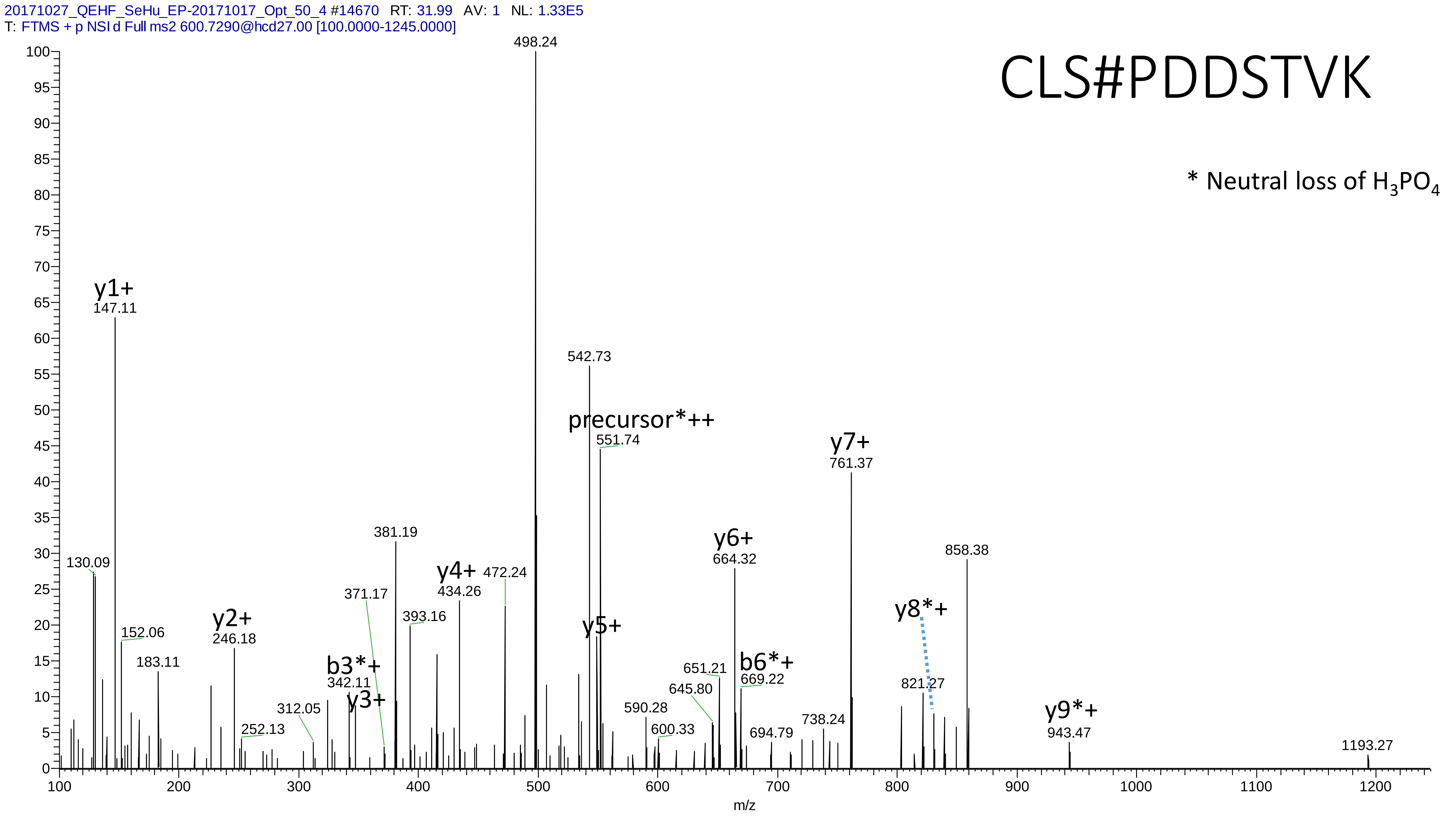
20171027_QEHF_SeHu_EP-20171017_Opt_50_4 #14670 RT: 31.99 AV: 1 NL: 1.33E5
T: FTMS + p NSI d Full ms2 600.7290@hcd27.00 [100.0000-1245.0000]
CLS#PDDSTVK
* Neutral loss of $H_3PO_4$
498.24
y1+
147.11
130.09
152.06
183.11
y2+
246.18
252.13
312.05
b3*+
342.11
y3+
371.17
381.19
393.16
y4+
434.26
472.24
542.73
precursor*++
551.74
y5+
590.28
600.33
645.80
651.21
y6+
664.32
b6*+
669.22
694.79
738.24
y7+
761.37
y8*+
821.27
858.38
y9*+
943.47
1193.27
m/z

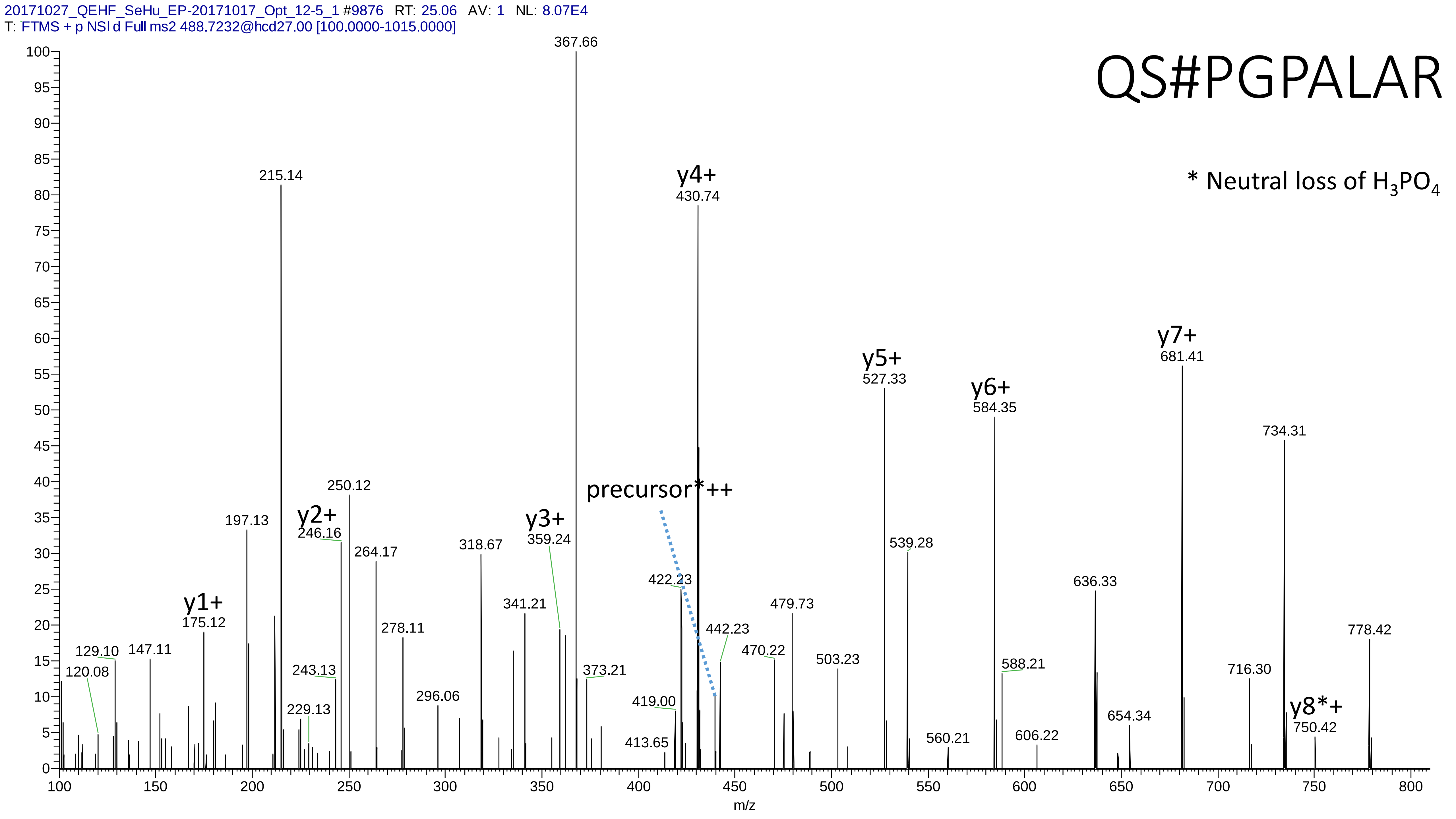
20171027_QEHF_SeHu_EP-20171017_Opt_12-5_1 #9876 RT: 25.06 AV: 1 NL: 8.07E4
T: FTMS + p NSI d Full ms2 488.7232@hcd27.00 [100.0000-1015.0000]
QS#PGPALAR
* Neutral loss of H3PO4
y1+
y2+
y3+
y4+
y5+
y6+
y7+
y8*+
precursor*++
m/z

# 29

- Experimental mass: 135.8869
- Unimod information:
- Possible composition: Phosphorylation + Cation:Ni[II], [H(1)O(3)P(1) + H(-2) Ni]
- Sites in Unimod:
- Sites observed:
- Features: neutral loss of $H_3PO_4$ are observed

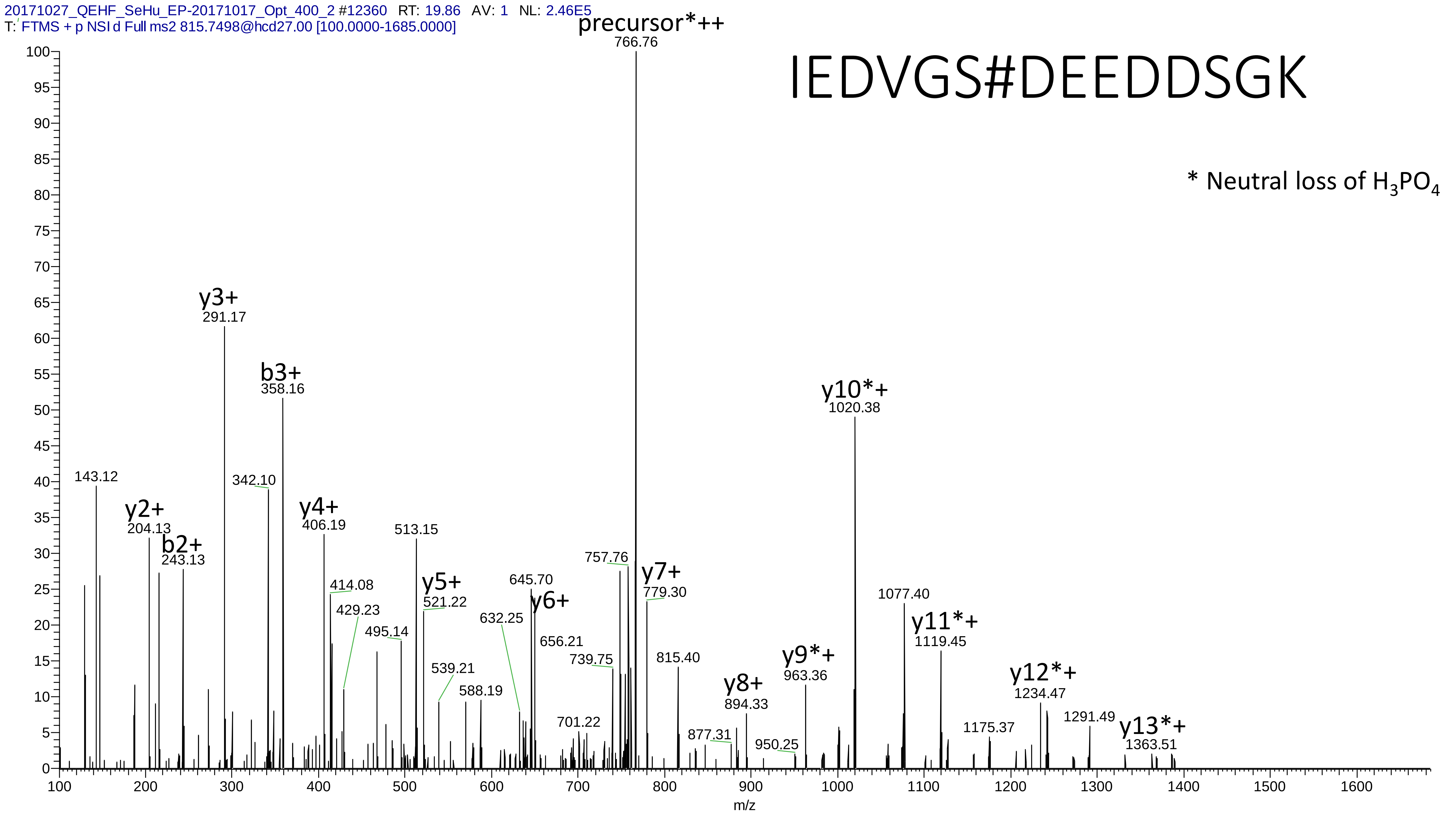

20171027_QEHF_SeHu_EP-20171017_Opt_400_2 #12360 RT: 19.86 AV: 1 NL: 2.46E5
T: FTMS + p NSI d Full ms2 815.7498@hcd27.00 [100.0000-1685.0000]
IEDVGS#DEEDDSGK
* Neutral loss of $H_3PO_4$
precursor*++
766.76
y3+
291.17
b3+
358.16
y10*+
1020.38
143.12
342.10
y2+
204.13
b2+
243.13
y4+
406.19
513.15
414.08
429.23
495.14
y5+
521.22
539.21
588.19
632.25
645.70
y6+
656.21
701.22
739.75
757.76
y7+
779.30
815.40
877.31
y8+
894.33
950.25
y9*+
963.36
1077.40
y11*+
1119.45
1175.37
y12*+
1234.47
1291.49
y13*+
1363.51
m/z

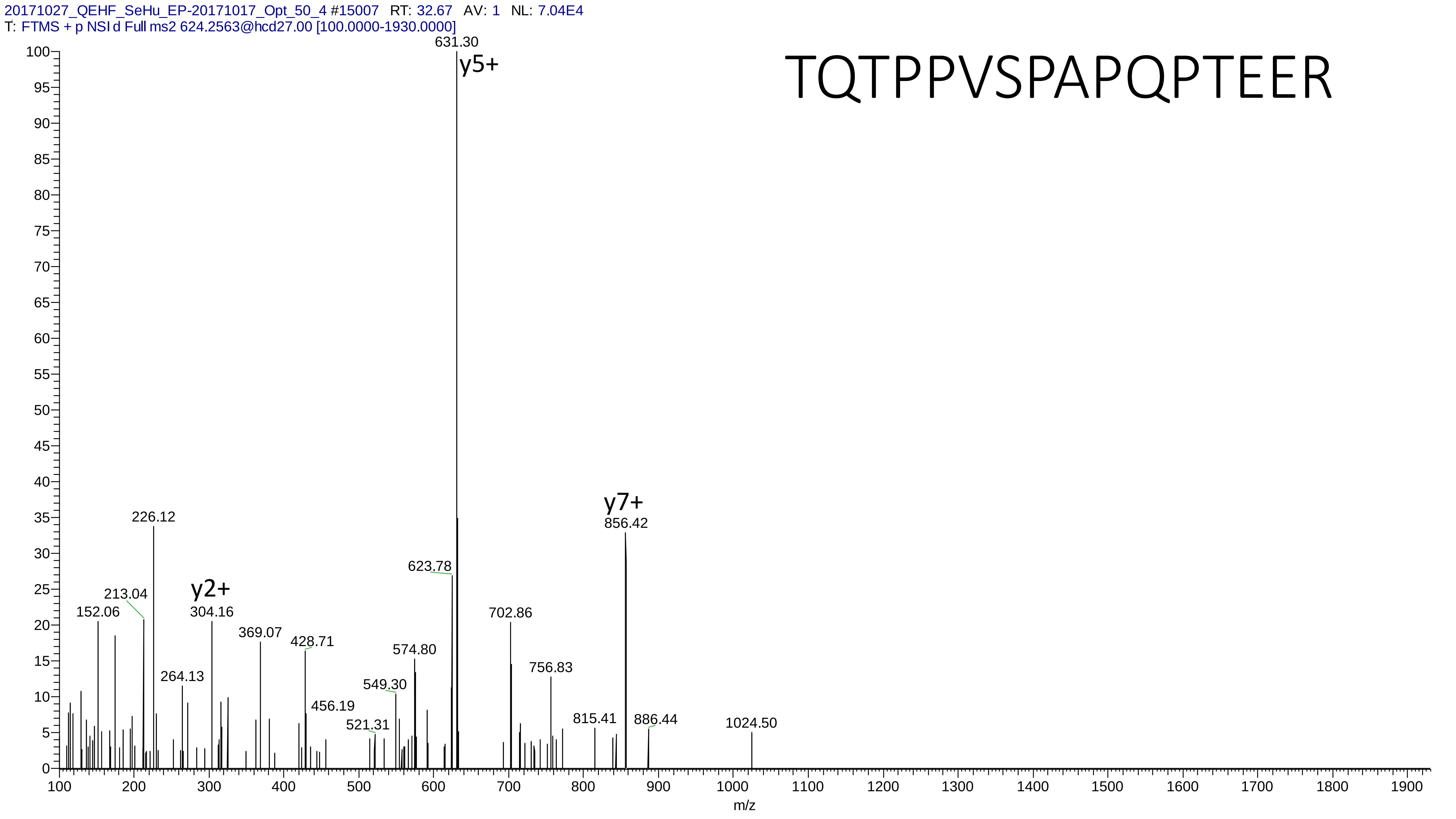
20171027_QEHF_SeHu_EP-20171017_Opt_50_4 #15007 RT: 32.67 AV: 1 NL: 7.04E4
T: FTMS + p NSI d Full ms2 624.2563@hcd27.00 [100.0000-1930.0000]
TQTPPVSPAPQPTEER
631.30
y5+
y7+
856.42
226.12
623.78
y2+
213.04
304.16
152.06
702.86
369.07
428.71
574.80
264.13
756.83
549.30
456.19
521.31
815.41
886.44
1024.50
m/z

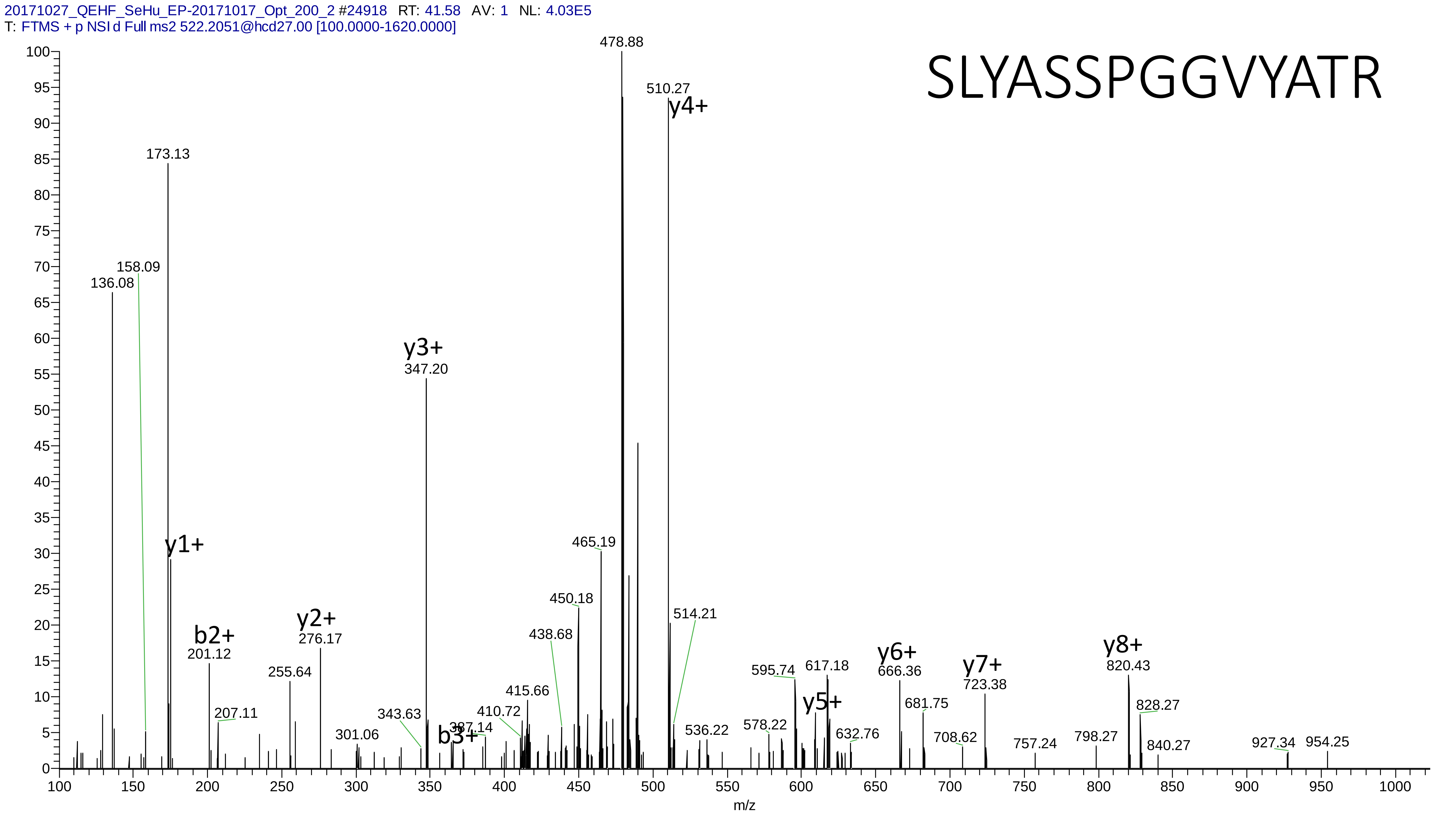

20171027_QEHF_SeHu_EP-20171017_Opt_200_2 #24918 RT: 41.58 AV: 1 NL: 4.03E5
T: FTMS + p NSI d Full ms2 522.2051@hcd27.00 [100.0000-1620.0000]
SLYASSPGGVYATR
478.88
510.27
y4+
173.13
158.09
136.08
y3+
347.20
y1+
465.19
450.18
514.21
438.68
b2+
201.12
y2+
276.17
255.64
207.11
415.66
410.72
343.63
387.14
b3+
301.06
595.74
617.18
y5+
578.22
536.22
632.76
y6+
666.36
681.75
708.62
y7+
723.38
757.24
798.27
y8+
820.43
828.27
840.27
927.34
954.25
m/z

# 30

- Experimental mass: 216.9552
- Unimod information:
- Possible composition: Phosphorylation * 2 + Carbamidomethyl, [H(1)O(3)P(1) * 2 + H(3) C(2) N O]
- Sites in Unimod:
- Sites observed:
- Features: 99.0% (100/101) peptides contain equal or more than two Ser/Thr, because of long peptide sequence and high charge state, the fragment information is insufficient for the determination of the modification sites

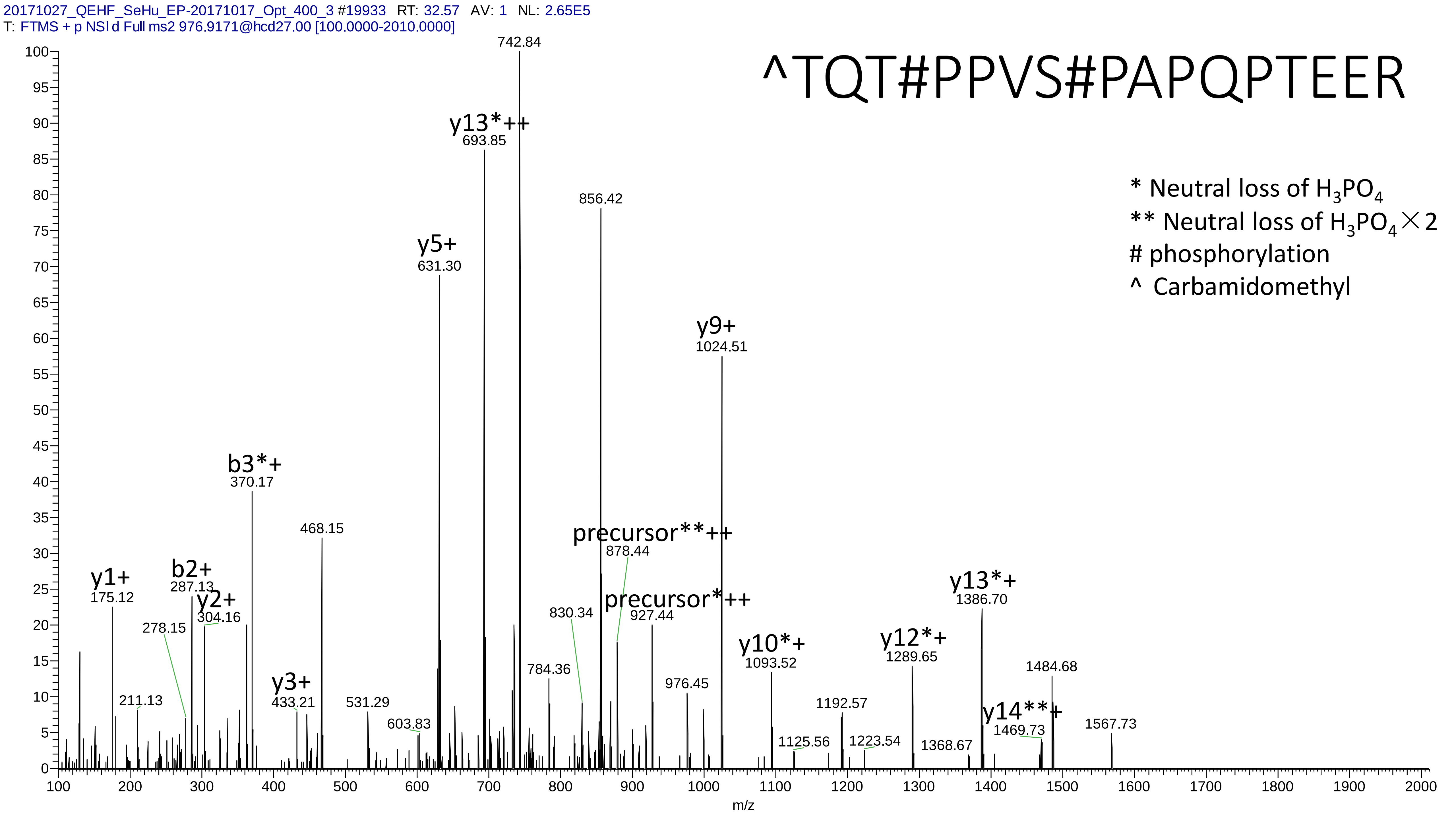
20171027_QEHF_SeHu_EP-20171017_Opt_400_3 #19933 RT: 32.57 AV: 1 NL: 2.65E5
T: FTMS + p NSI d Full ms2 976.9171@hcd27.00 [100.0000-2010.0000]
^TQT#PPVS#PAPQPTEER
* Neutral loss of $H_3PO_4$
** Neutral loss of $H_3PO_4$ × 2
### phosphorylation
^ Carbamidomethyl
742.84
y13*++
693.85
856.42
y5+
631.30
y9+
1024.51
b3*+
370.17
468.15
precursor**++
878.44
precursor*++
927.44
y1+
175.12
b2+
287.13
y2+
304.16
278.15
830.34
y13*+
1386.70
y10*+
1093.52
y12*+
1289.65
1484.68
784.36
976.45
211.13
y3+
433.21
531.29
603.83
1192.57
y14**+
1469.73
1567.73
1125.56
1223.54
1368.67
m/z

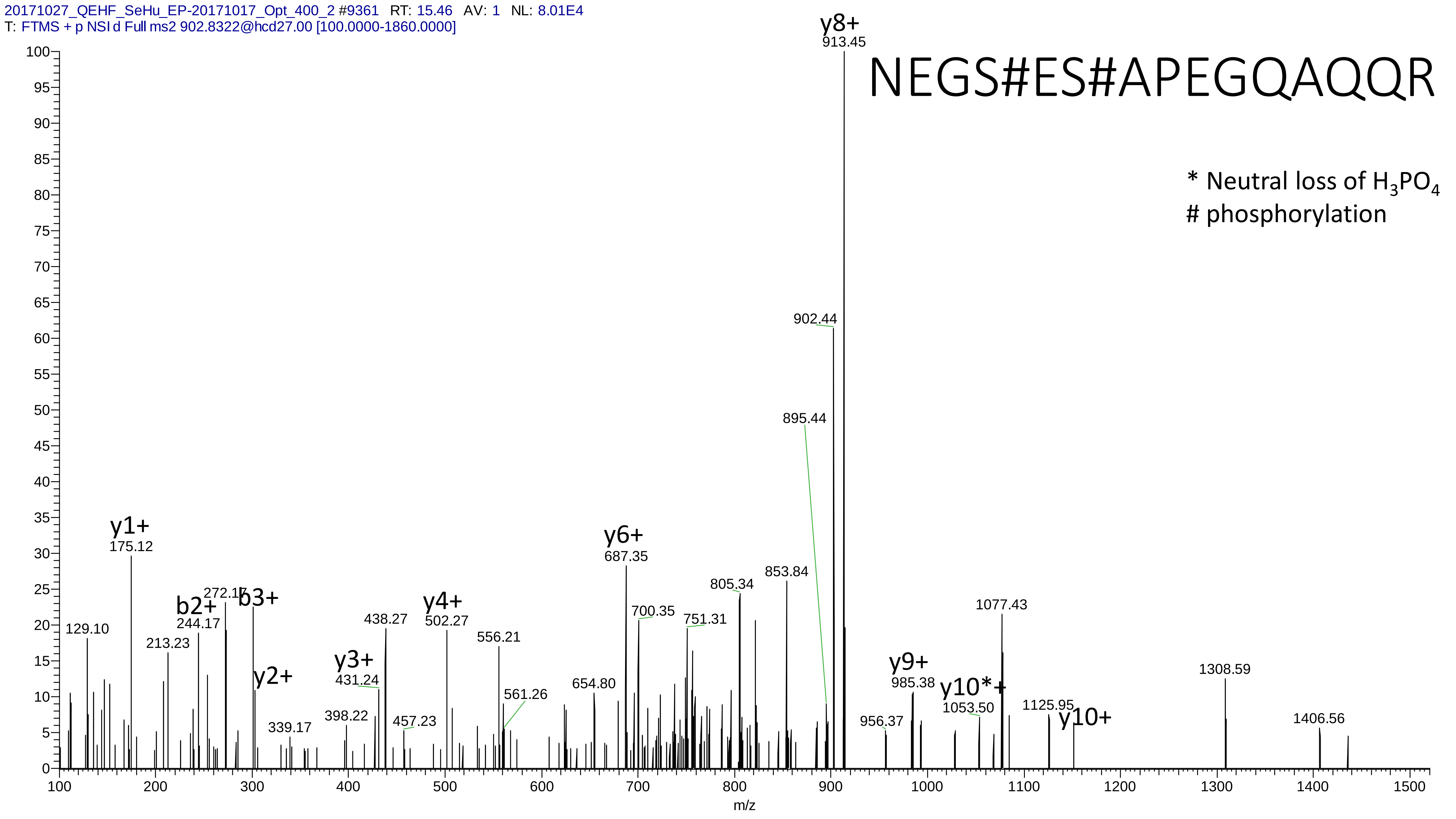
20171027_QEHF_SeHu_EP-20171017_Opt_400_2 #9361 RT: 15.46 AV: 1 NL: 8.01E4
T: FTMS + p NSI d Full ms2 902.8322@hcd27.00 [100.0000-1860.0000]
NEGS#ES#APEGQAQQR
* Neutral loss of $H_3PO_4$
### phosphorylation
y8+ 913.45
902.44
895.44
y1+ 175.12
y6+ 687.35
b2+ 244.17
272.17
b3+
y2+
y3+ 431.24
y4+ 502.27
129.10
213.23
438.27
556.21
561.26
339.17
398.22
457.23
654.80
700.35
751.31
805.34
853.84
1077.43
y9+ 985.38
y10*+ 1053.50
956.37
1125.95
y10+
1308.59
1406.56
m/z
