## Supplementary material 5 for "MetaLab 2.0 enables accurate post-translational modifications profiling in metaproteomics"

### Supplementary material 5: Manually annotated spectra of the glycosylation

Dataset: human_gut

# 1

- Experimental mass: 349.1375
- Unimod information: HexNAc(1)dHex(1)
- Possible composition: dHex HexNAc
- Sites in Unimod: N;S;T
- Sites observed: N
- Features:  81.5% (265/325) peptides contain N-glycan motif NX[S/T]

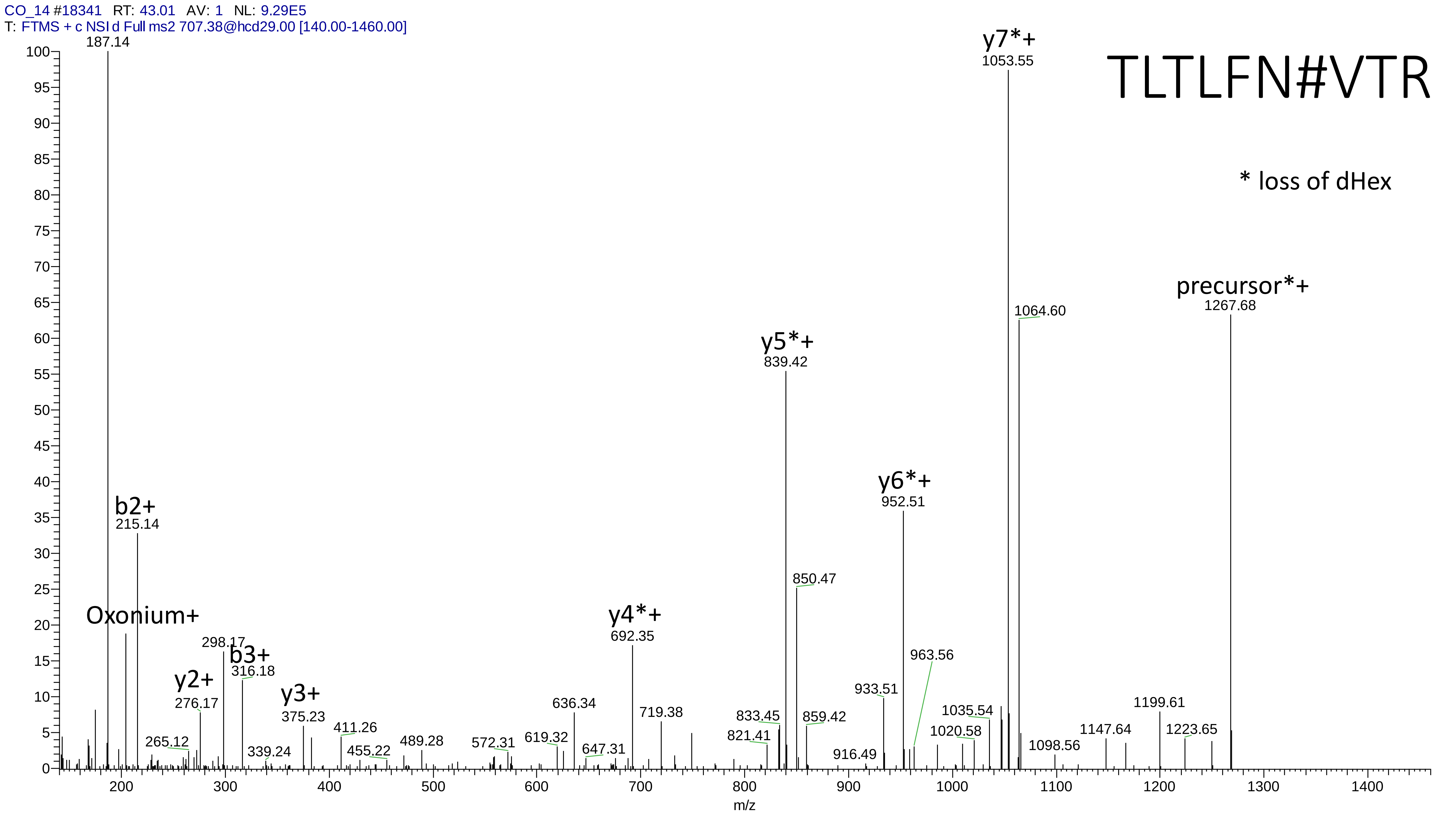
CO_14 #18341 RT: 43.01 AV: 1 NL: 9.29E5
T: FTMS + c NSI d Full ms2 707.38@hcd29.00 [140.00-1460.00]
TLTLFN#VTR
* loss of dHex
187.14
Oxonium+
b2+
215.14
y2+
276.17
298.17
b3+
316.18
y3+
375.23
y4*+
692.35
y5*+
839.42
850.47
y6*+
952.51
963.56
y7*+
1053.55
1064.60
precursor*+
1267.68
265.12
339.24
411.26
455.22
489.28
572.31
619.32
636.34
647.31
719.38
821.41
833.45
859.42
916.49
933.51
1020.58
1035.54
1098.56
1147.64
1199.61
1223.65
m/z

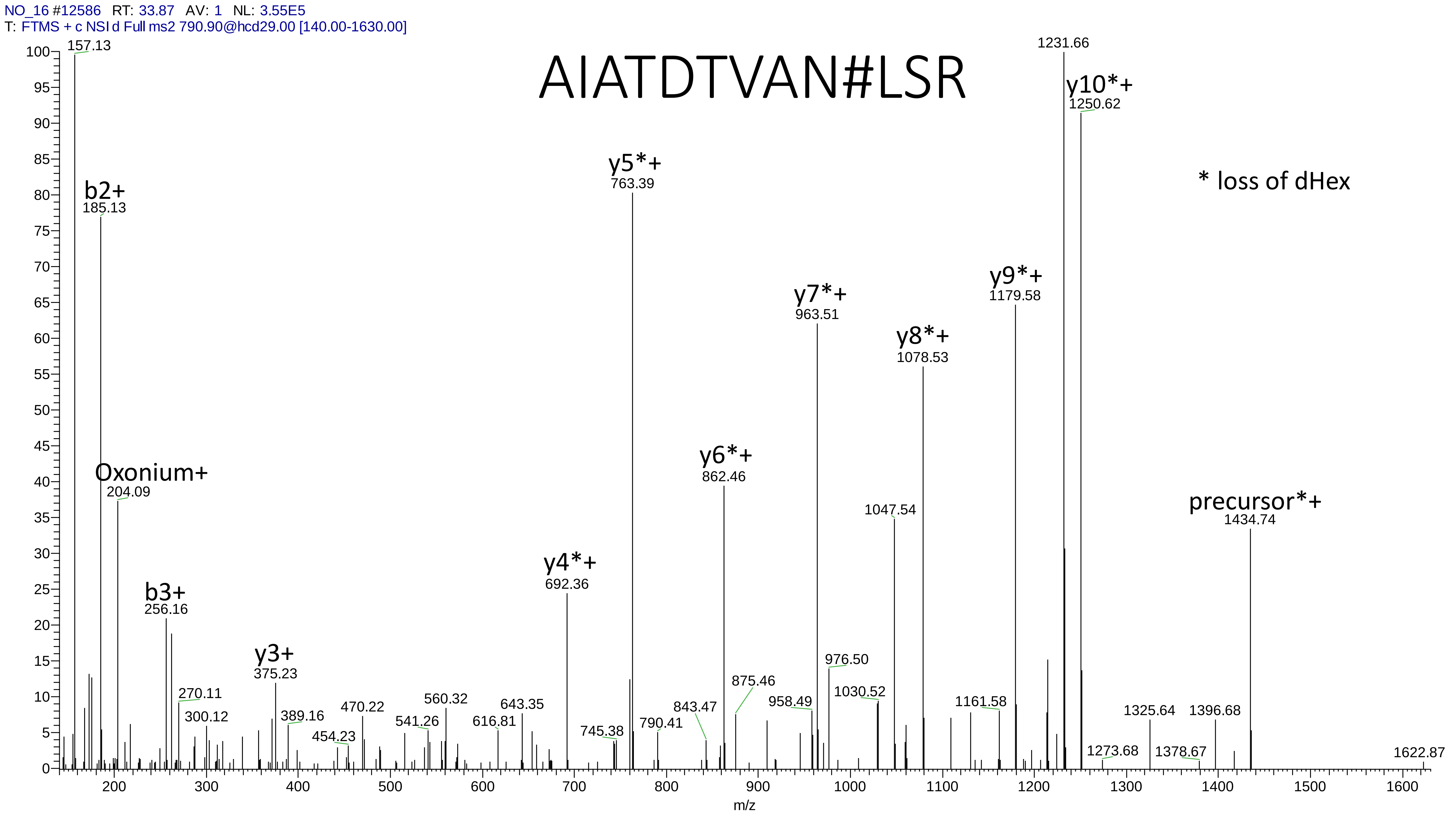

NO_16 #12586 RT: 33.87 AV: 1 NL: 3.55E5
T: FTMS + c NSI d Full ms2 790.90@hcd29.00 [140.00-1630.00]
AIATDTVAN#LSR
* loss of dHex
157.13
b2+
185.13
Oxonium+
204.09
b3+
256.16
270.11
300.12
y3+
375.23
389.16
454.23
470.22
541.26
560.32
616.81
643.35
y4*+
692.36
745.38
y5*+
763.39
790.41
843.47
y6*+
862.46
875.46
958.49
y7*+
963.51
976.50
1030.52
1047.54
y8*+
1078.53
1161.58
y9*+
1179.58
1231.66
y10*+
1250.62
1273.68
1325.64
1378.67
1396.68
precursor*+
1434.74
1622.87
m/z

# 2

- Experimental mass: 203.0796
- Unimod information: HexNAc
- Possible composition: HexNAc
- Sites in Unimod: N;S;T;C
- Sites observed: N
- Features:  84.6% (297/351) peptides contain N-glycan motif NX[S/T]

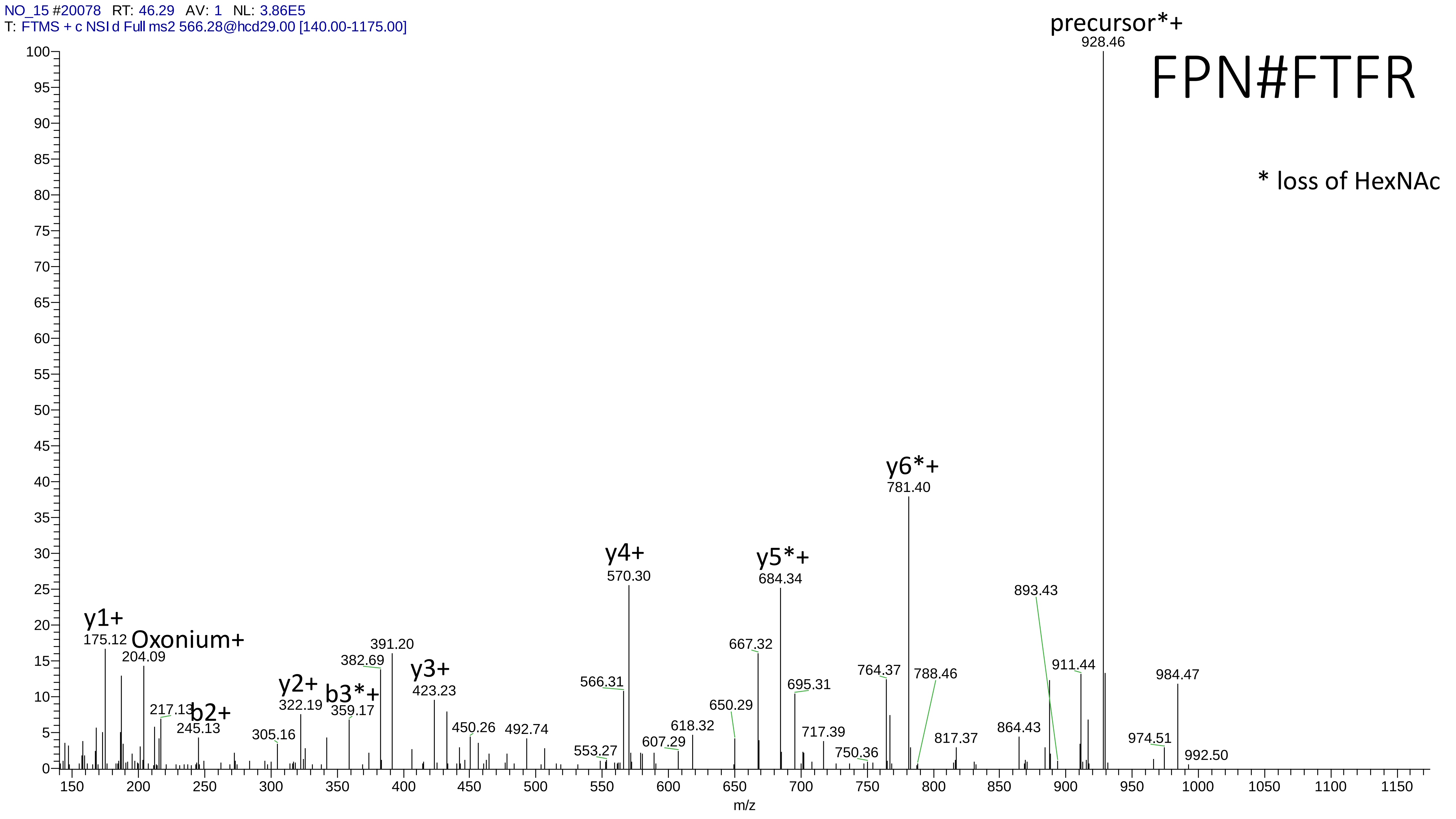

NO_15 #20078 RT: 46.29 AV: 1 NL: 3.86E5
T: FTMS + c NSI d Full ms2 566.28@hcd29.00 [140.00-1175.00]
FPN#FTFR
* loss of HexNAc
precursor*+
928.46
y6*+
781.40
y5*+
684.34
y4+
570.30
y3+
423.23
y2+
322.19
b3*+
359.17
b2+
245.13
y1+
175.12
Oxonium+
204.09
217.13
305.16
382.69
391.20
450.26
492.74
553.27
566.31
607.29
618.32
650.29
667.32
695.31
717.39
750.36
764.37
788.46
817.37
864.43
893.43
911.44
974.51
984.47
992.50
m/z

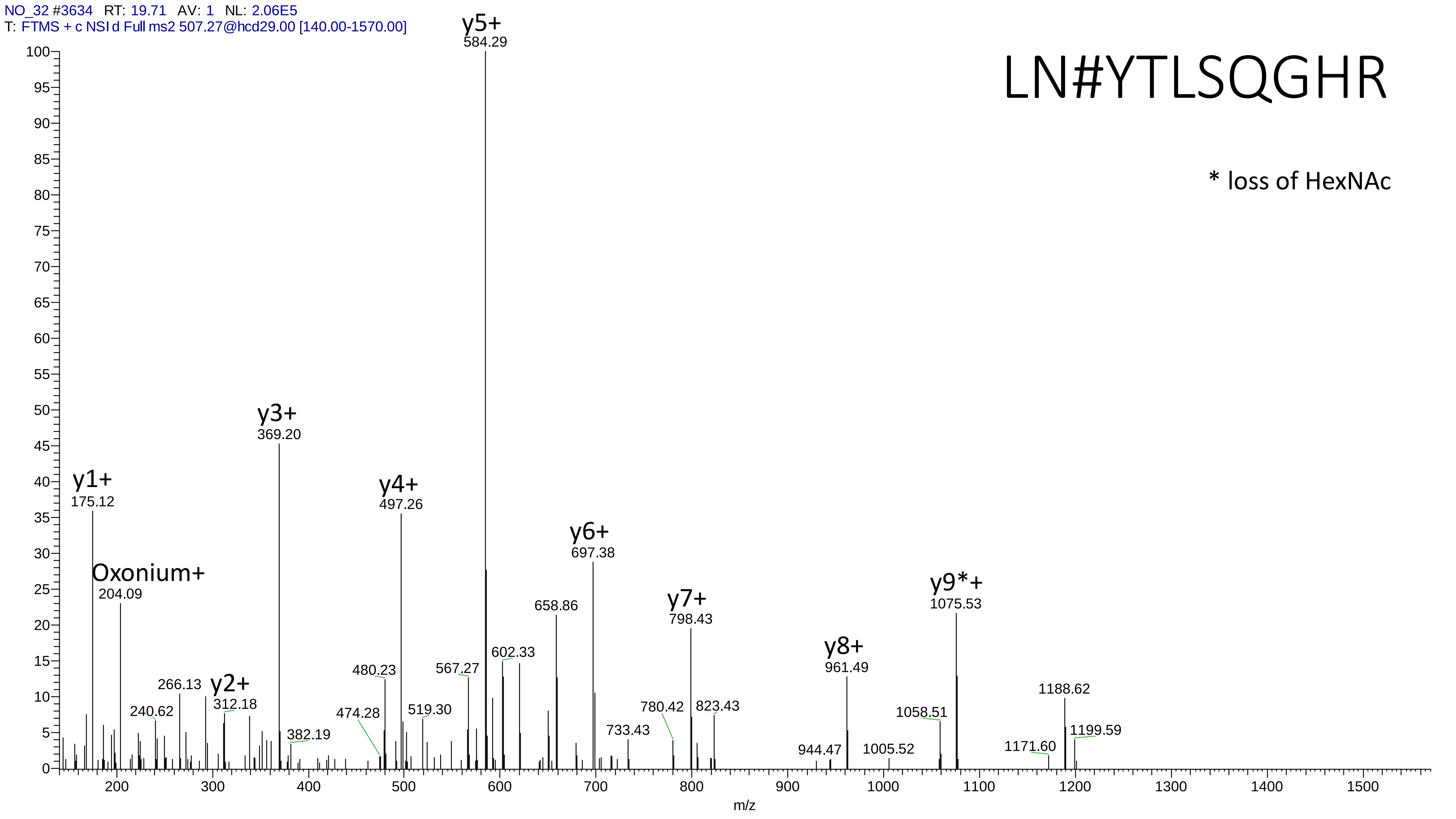
NO_32 #3634 RT: 19.71 AV: 1 NL: 2.06E5
T: FTMS + c NSI d Full ms2 507.27@hcd29.00 [140.00-1570.00]
LN#YTLSQGHR
* loss of HexNAc
y1+ 175.12
Oxonium+ 204.09
y2+ 312.18
y3+ 369.20
y4+ 497.26
y5+ 584.29
y6+ 697.38
y7+ 798.43
y8+ 961.49
y9*+ 1075.53
240.62
266.13
382.19
474.28
480.23
519.30
567.27
602.33
658.86
733.43
780.42
823.43
944.47
1005.52
1058.51
1171.60
1188.62
1199.59
m/z

# 3

- Experimental mass: 406.1593
- Unimod information: HexNAc(2)
- Possible composition: HexNAc(2)
- Sites in Unimod: N;S;T
- Sites observed: unknown
- Features:  66.7% (28/42) peptides contain N-glycan motif NX[S/T]

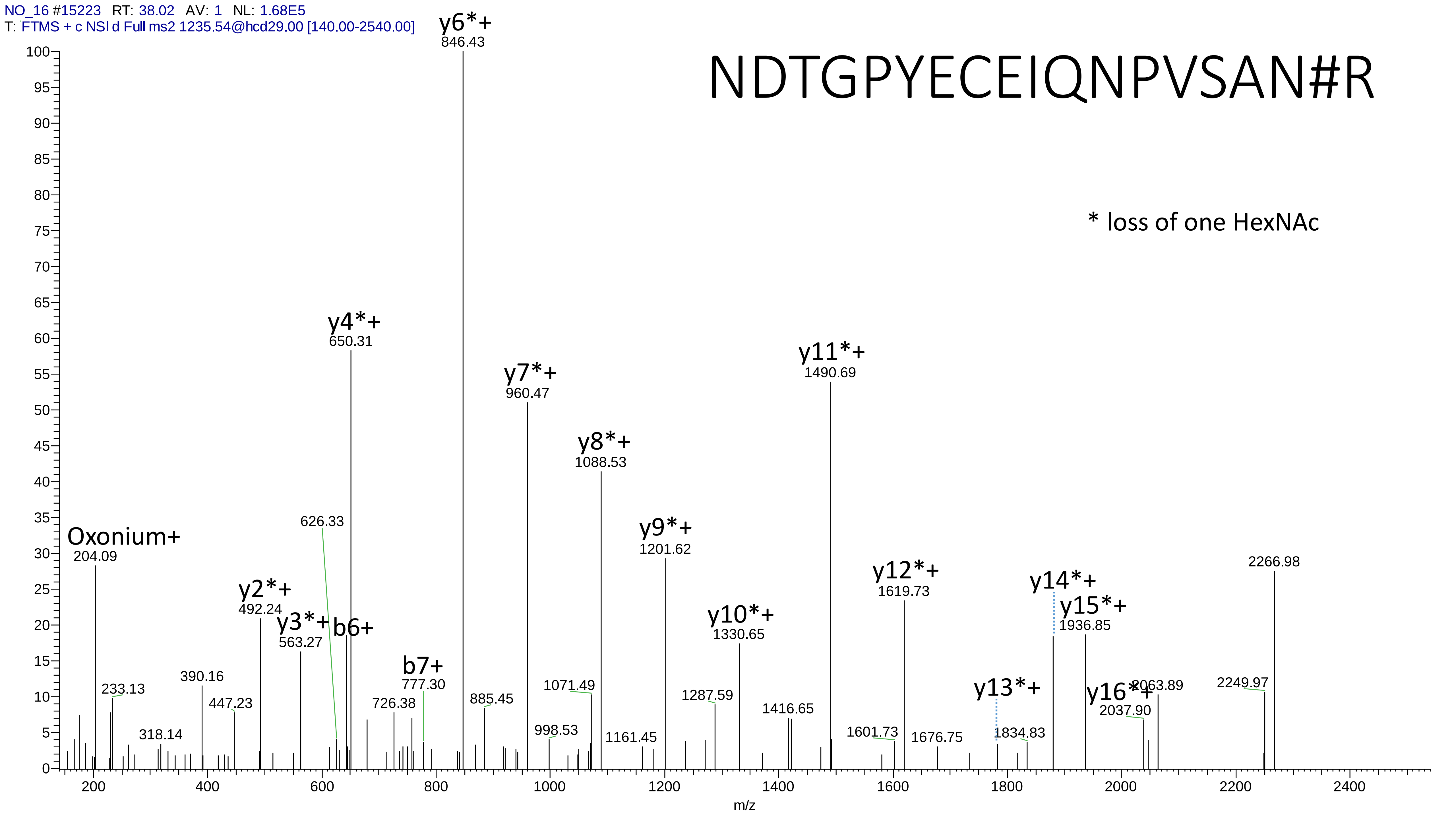

NO_16 #15223 RT: 38.02 AV: 1 NL: 1.68E5
T: FTMS + c NSI d Full ms2 1235.54@hcd29.00 [140.00-2540.00]
NDTGPYECEIQNPVSAN#R
* loss of one HexNAc
Oxonium+
204.09
233.13
318.14
390.16
447.23
y2*+
492.24
y3*+
563.27
b6+
626.33
y4*+
650.31
726.38
b7+
777.30
y6*+
846.43
885.45
y7*+
960.47
998.53
1071.49
y8*+
1088.53
1161.45
y9*+
1201.62
1287.59
y10*+
1330.65
1416.65
y11*+
1490.69
1601.73
y12*+
1619.73
1676.75
y13*+
1834.83
y14*+
y15*+
1936.85
y16*+
2037.90
2063.89
2249.97
2266.98
m/z

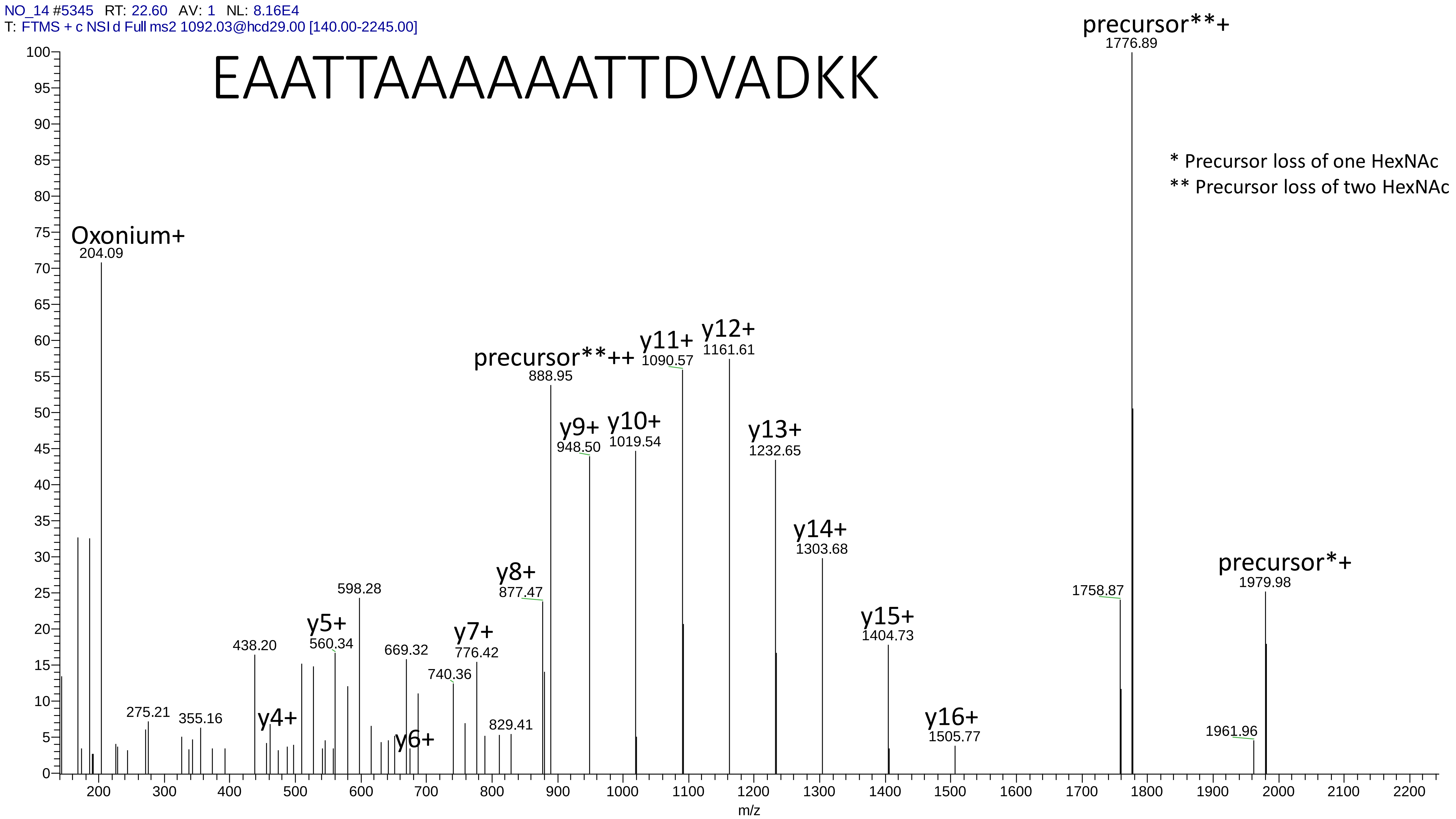
NO_14 #5345 RT: 22.60 AV: 1 NL: 8.16E4
T: FTMS + c NSI d Full ms2 1092.03@hcd29.00 [140.00-2245.00]
EAATTAAAAAATTDVADKK
* Precursor loss of one HexNAc
** Precursor loss of two HexNAc
Oxonium+
204.09
275.21
355.16
438.20
y4+
y5+
560.34
598.28
y6+
669.32
740.36
y7+
776.42
829.41
y8+
877.47
precursor**++
888.95
y9+
948.50
y10+
1019.54
y11+
1090.57
y12+
1161.61
y13+
1232.65
y14+
1303.68
y15+
1404.73
y16+
1505.77
1758.87
precursor**+
1776.89
1961.96
precursor*+
1979.98
m/z

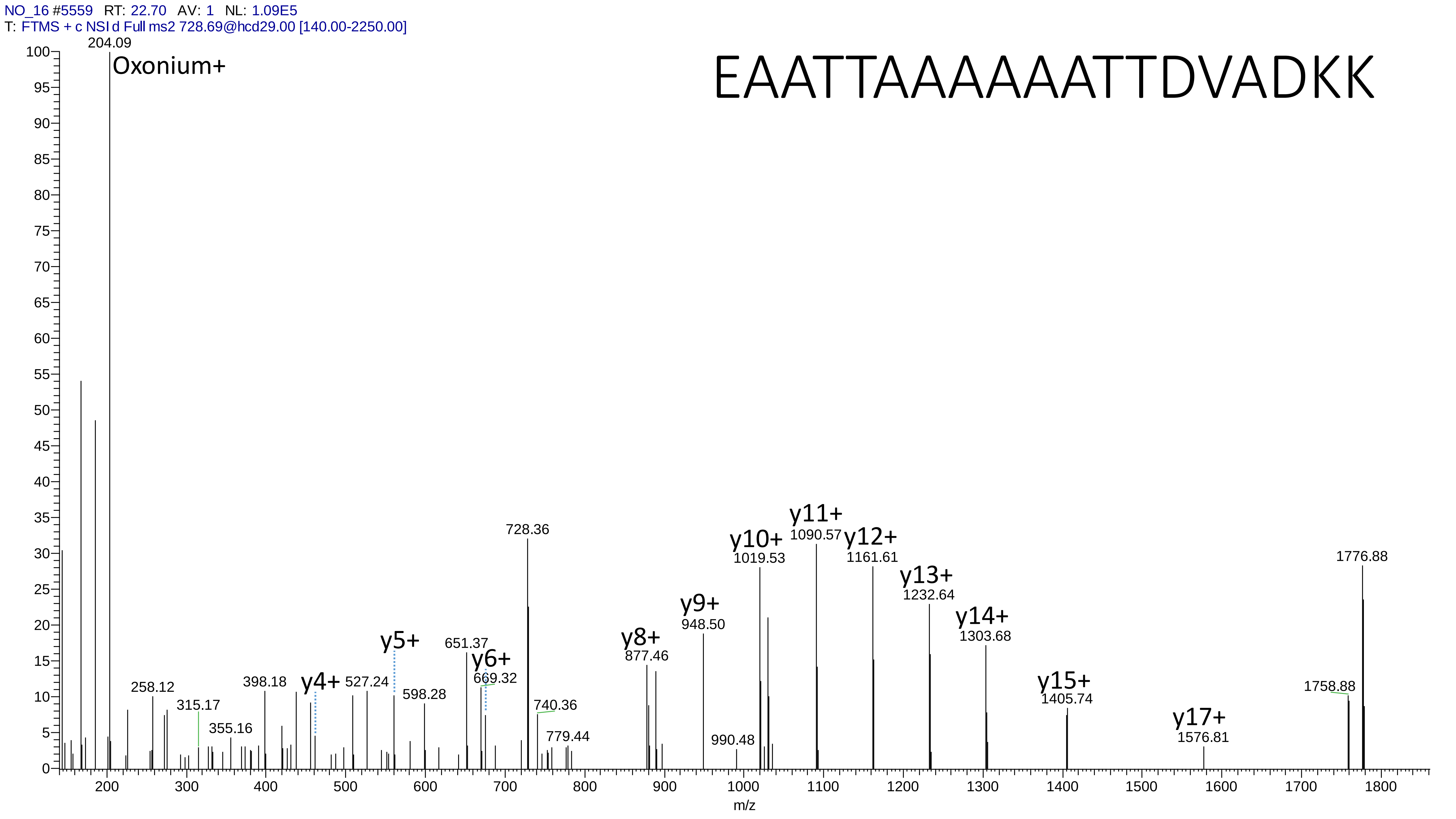
NO_16 #5559 RT: 22.70 AV: 1 NL: 1.09E5
T: FTMS + c NSI d Full ms2 728.69@hcd29.00 [140.00-2250.00]
EAATTAAAAAATTDVADKK
204.09
Oxonium+
258.12
315.17
355.16
398.18
y4+
527.24
y5+
598.28
651.37
y6+
669.32
728.36
740.36
779.44
y8+
877.46
y9+
948.50
990.48
y10+
1019.53
y11+
1090.57
y12+
1161.61
y13+
1232.64
y14+
1303.68
y15+
1405.74
y17+
1576.81
1758.88
1776.88
m/z

# 4

- Experimental mass: 162.053
- Unimod information: Hexose
- Possible composition: Hex
- Sites in Unimod: K;R;N;S;T;W;C;Y; peptide n-term
- Sites observed: unknown

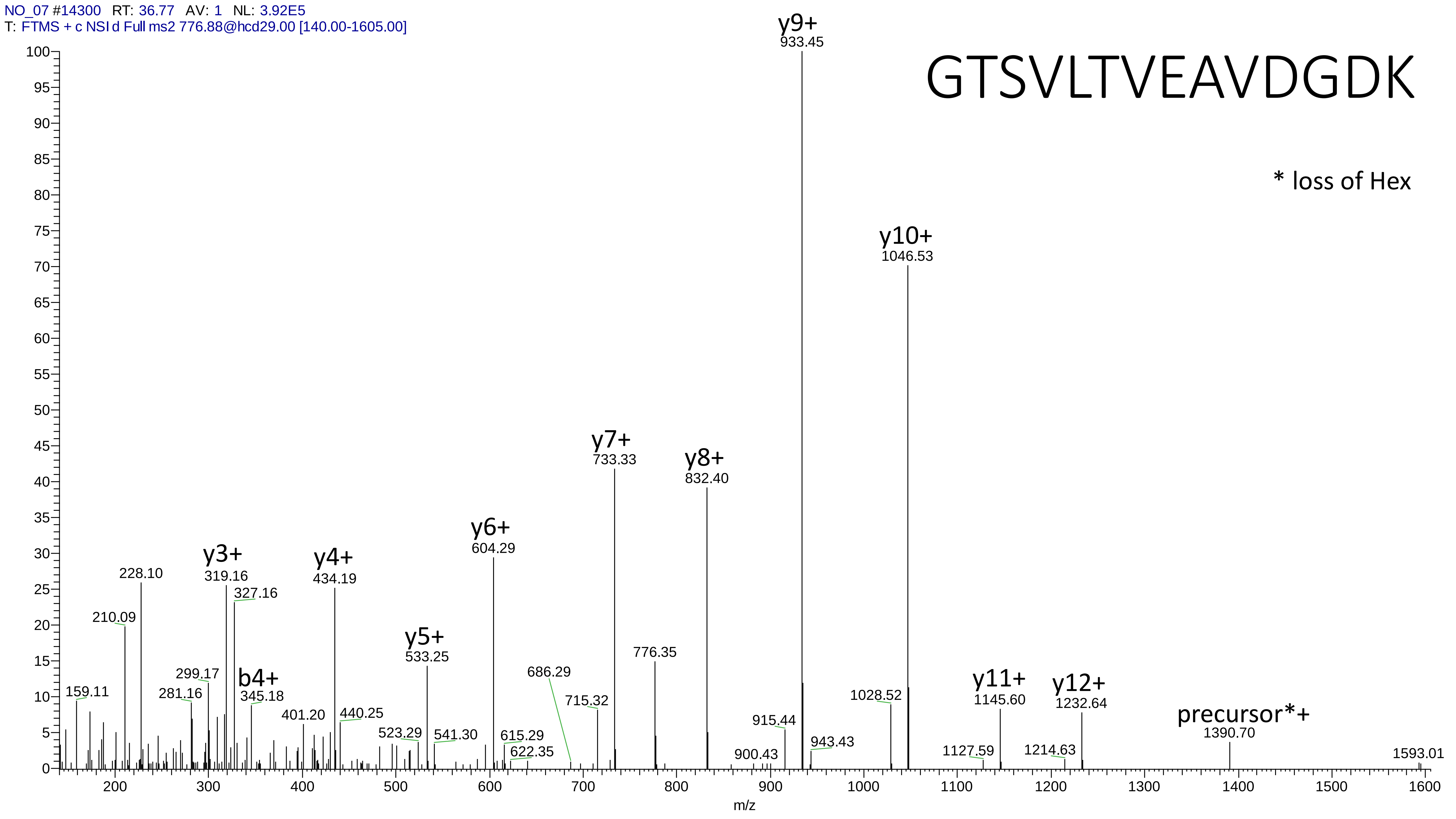
NO_07 #14300 RT: 36.77 AV: 1 NL: 3.92E5
T: FTMS + c NSI d Full ms2 776.88@hcd29.00 [140.00-1605.00]
GTSVLTVEAVDGDK
* loss of Hex
y9+
933.45
y10+
1046.53
y7+
733.33
y8+
832.40
y6+
604.29
y3+
319.16
y4+
434.19
228.10
327.16
210.09
y5+
533.25
776.35
299.17
b4+
345.18
159.11
281.16
686.29
715.32
401.20
440.25
523.29
541.30
615.29
622.35
900.43
915.44
943.43
1028.52
y11+
1145.60
y12+
1232.64
precursor*+
1390.70
1127.59
1214.63
1593.01
m/z

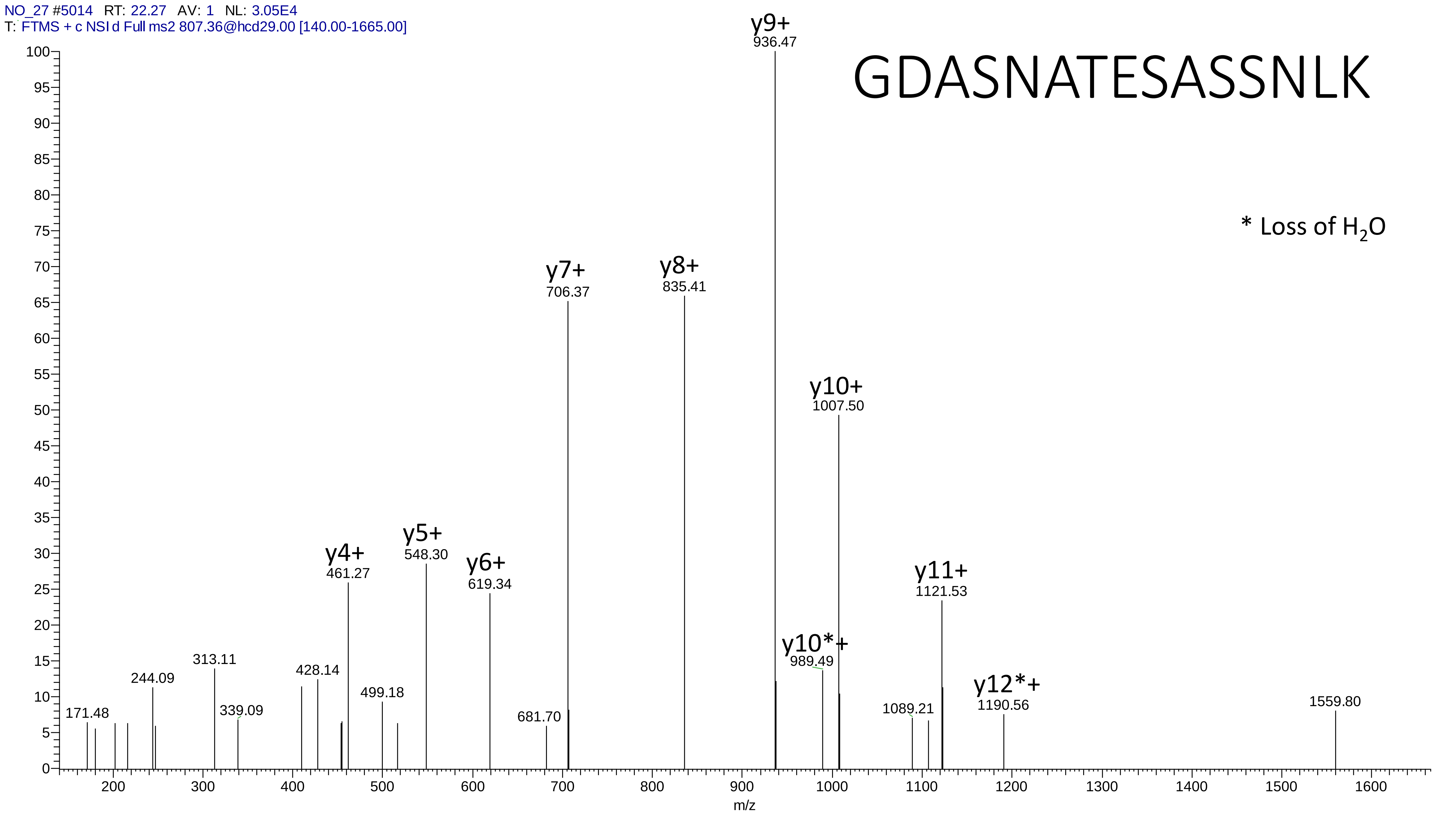
NO_27 #5014 RT: 22.27 AV: 1 NL: 3.05E4
T: FTMS + c NSI d Full ms2 807.36@hcd29.00 [140.00-1665.00]
GDASNATESASSNLK
* Loss of $H_2O$
y4+ 461.27
y5+ 548.30
y6+ 619.34
y7+ 706.37
y8+ 835.41
y9+ 936.47
y10*+ 989.49
y10+ 1007.50
y11+ 1121.53
y12*+ 1190.56
171.48
244.09
313.11
339.09
428.14
499.18
681.70
1089.21
1559.80
m/z

# 5

- Experimental mass: 228.1107
- Unimod information: Bacillosamine
- Possible composition: H(6) C(4) N(2) dHex
- Sites in Unimod: N
- Sites observed: unknown

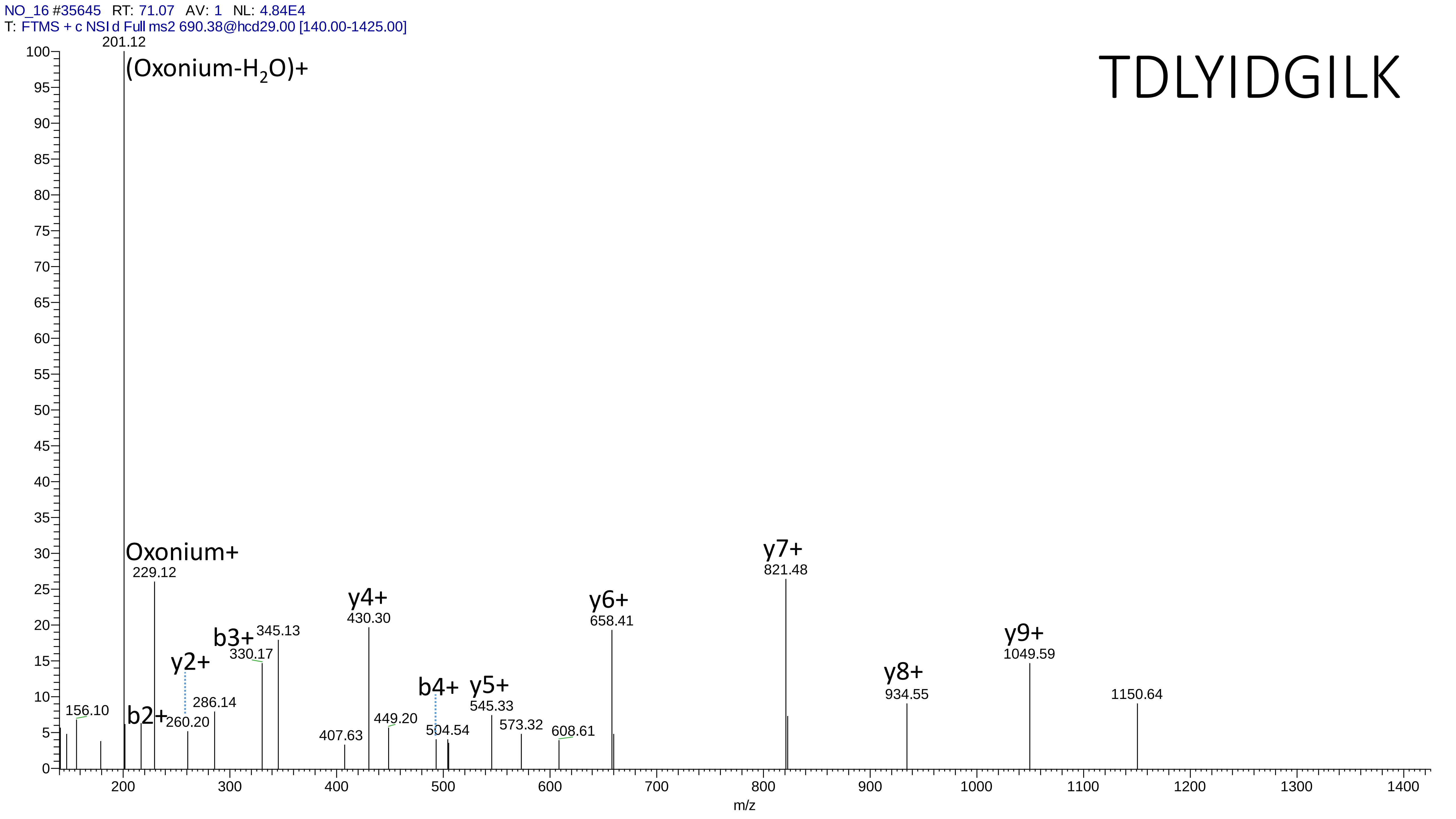
NO_16 #35645 RT: 71.07 AV: 1 NL: 4.84E4
T: FTMS + c NSI d Full ms2 690.38@hcd29.00 [140.00-1425.00]
TDLYIDGILK
201.12
(Oxonium-$H_2O$)+
Oxonium+
229.12
156.10
b2+
y2+
260.20
286.14
b3+
330.17
345.13
407.63
y4+
430.30
449.20
b4+
504.54
y5+
545.33
573.32
608.61
y6+
658.41
y7+
821.48
y8+
934.55
y9+
1049.59
1150.64
m/z

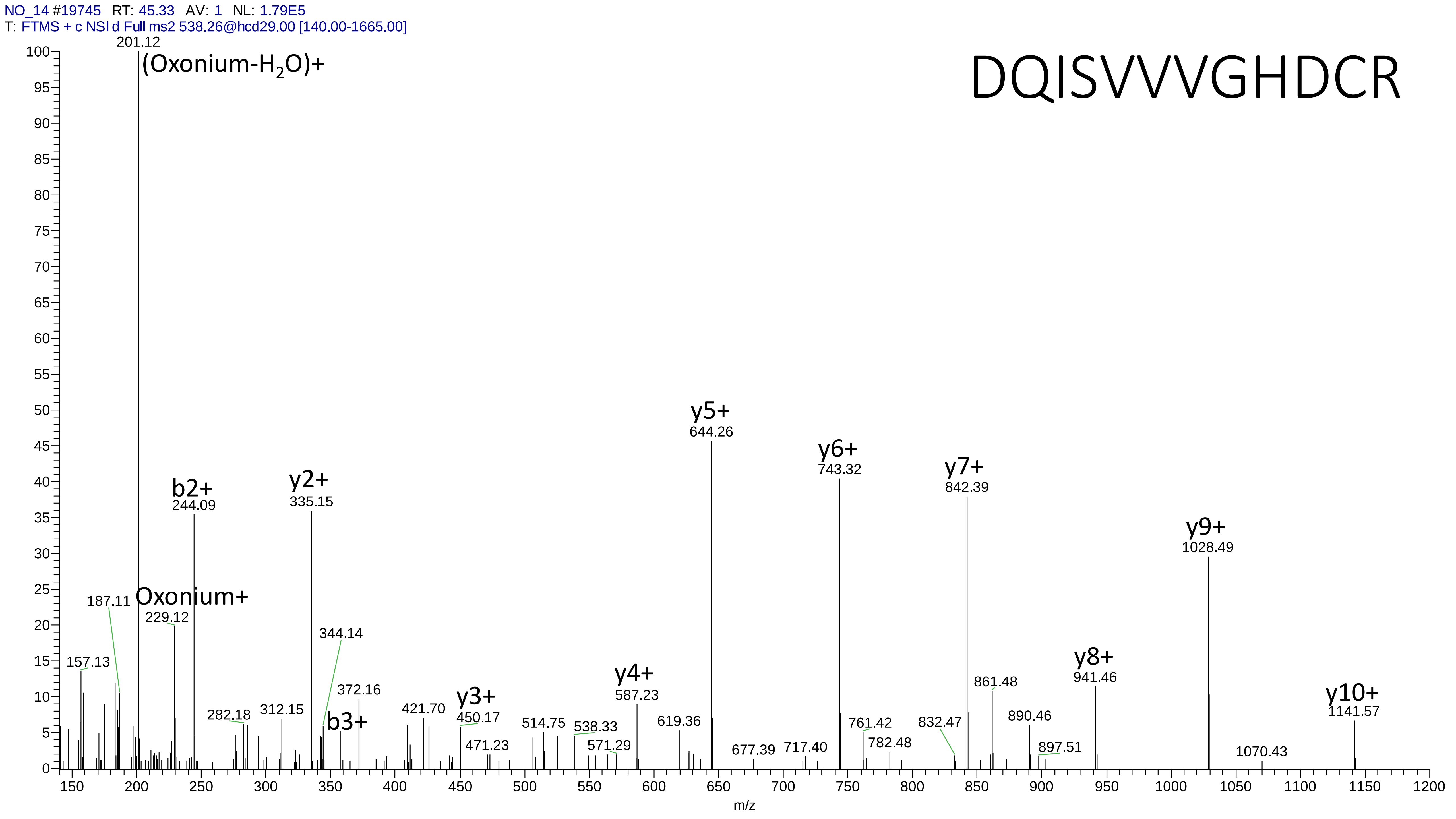
NO_14 #19745 RT: 45.33 AV: 1 NL: 1.79E5
T: FTMS + c NSI d Full ms2 538.26@hcd29.00 [140.00-1665.00]
DQISVVVGHDCR
201.12
(Oxonium-H2O)+
b2+
244.09
y2+
335.15
Oxonium+
229.12
187.11
157.13
344.14
372.16
b3+
282.18
312.15
421.70
y3+
450.17
471.23
514.75
538.33
571.29
y4+
587.23
619.36
y5+
644.26
677.39
717.40
y6+
743.32
761.42
782.48
y7+
842.39
832.47
861.48
890.46
897.51
y8+
941.46
y9+
1028.49
1070.43
y10+
1141.57
m/z
