## Supplementary notes for "MetaLab 2.0 enables accurate post-translational modifications profiling in metaproteomics"

### Supplementary figures and tables

| Supplementary figure 1 | Portion of the decoy PSMs in closed search that was recovered in open search |
| --- | --- |
| Supplementary figure 2 | Comparison of peptide identifications from open search and conventional closed search in a Homo_HEK293 dataset |
| Supplementary figure 3 | Comparison of peptide identifications from open search and conventional closed search in a Mock_micro dataset |
| Supplementary figure 4 | Quantitative analysis of a mock microbiome dataset |
| Supplementary figure 5 | The comparison of the identified (a) peptides and (b) species by closed search and open search from mouse gut microbiota samples. |
| Supplementary figure 6 | The sequence motif of the potential modification ∆Mass=71.0368 in the Mouse_gut dataset |
| Supplementary figure 7 | The comparison of peptide identifications from open search and conventional narrow search in Human_gut dataset |
| Supplementary figure 8 | The PSM counts and intensities of unmodified, oxidation/acetylation and modified peptides from human and microbial proteins |
| Supplementary figure 9 | The PSM count ratio of the identified potential modifications from Human_gut |
| Supplementary figure 10 | The network of HexNAc modified human glycoproteins |
| Supplementary figure 11 | The significantly changed (P<=0.05 by two-sample, two-tailed t-test) GO biological progresses of HexNAc modified human glycoproteins |
| Supplementary figure 12 | The taxon of Hex modified glycopeptides from CO and NO samples |
| Supplementary figure 13 | The functions of the microbial proteins modified by (a) Hex and (b) HexNAc |
| Supplementary figure 14 | The taxon of HexNAc2 modified glycopeptides from CO and NO samples |
| Supplementary figure 15 | The taxon of Bacillosamine modified glycopeptides from CO and NO samples |
| Supplementary table 1 | The features used for the machine learning classification |
| Supplementary table 2 | The 40 unknown ∆Mass which can be constructed from two known modifications already identified from this dataset |
| Supplementary table 3 | The glycan oxonium ions used for the determination of glycopeptides spectra |
| Supplementary table 4 | The information of the analyzed benchmark dataset |
| Supplementary table 5 | The Enriched Biological Process (GO) from the HexNAc modified human proteins |
| Supplementary table 6 | The glycopeptides and their taxons and functions |

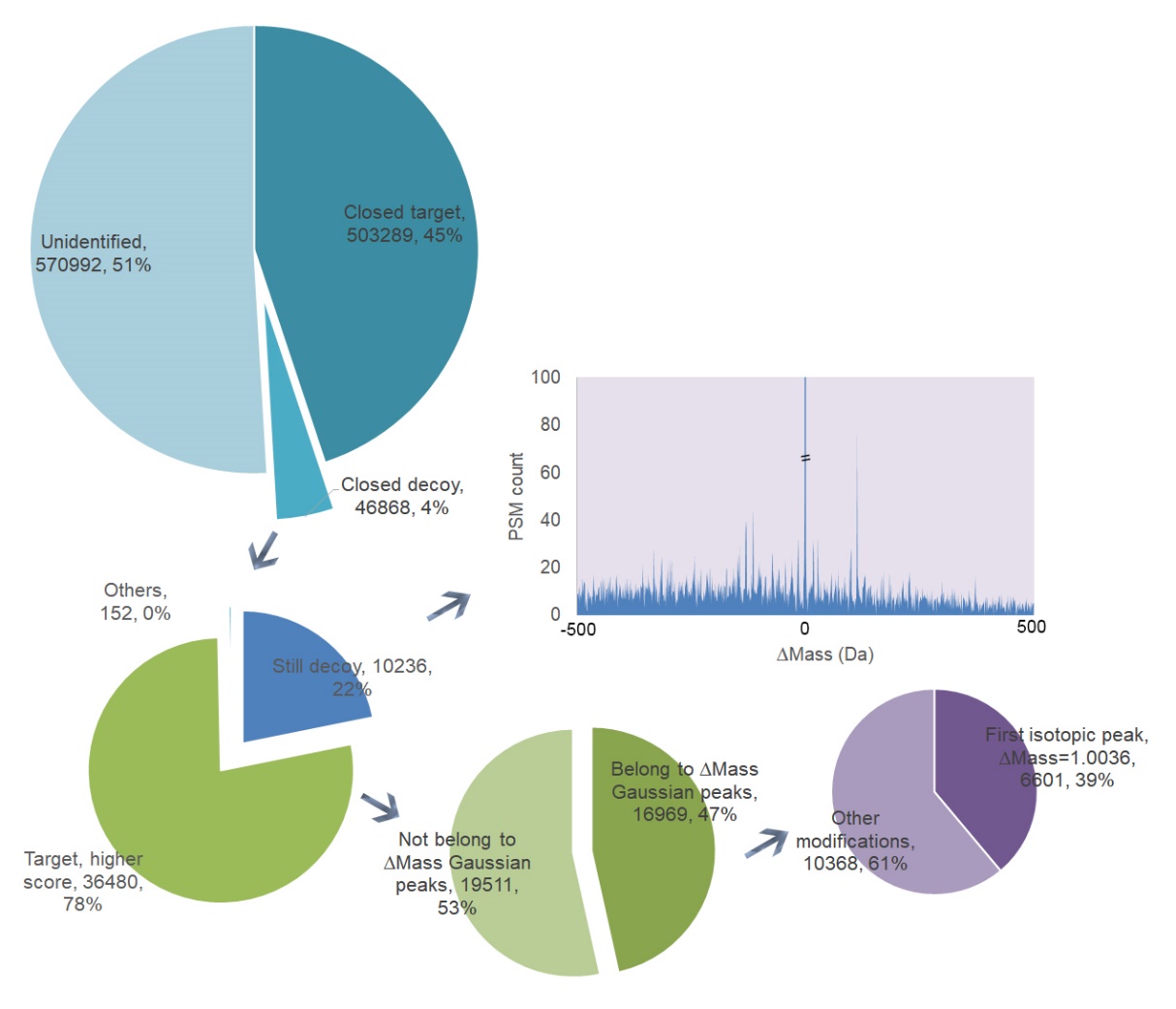

Supplementary Figure 1**. Portion of the decoy PSMs in closed search that was recovered in open search.** In the closed search 45% of the MS2 spectra were identified as target PSMs, 4% were identified as decoys and 51% of the MS2 spectra were unidentified. It was found that 78% of the decoy PSMs in closed search were matched to the target sequences and received higher scores in open search. The remaining 22% were decoy and evenly distributed in the ∆Mass range, except that the PSMs concentrated on the 0 Da position. For the PSMs that recieved higher scores with target sequences, 47% belonged to the determined ∆Mass peaks, which meant this part was recovered by the open search strategy.

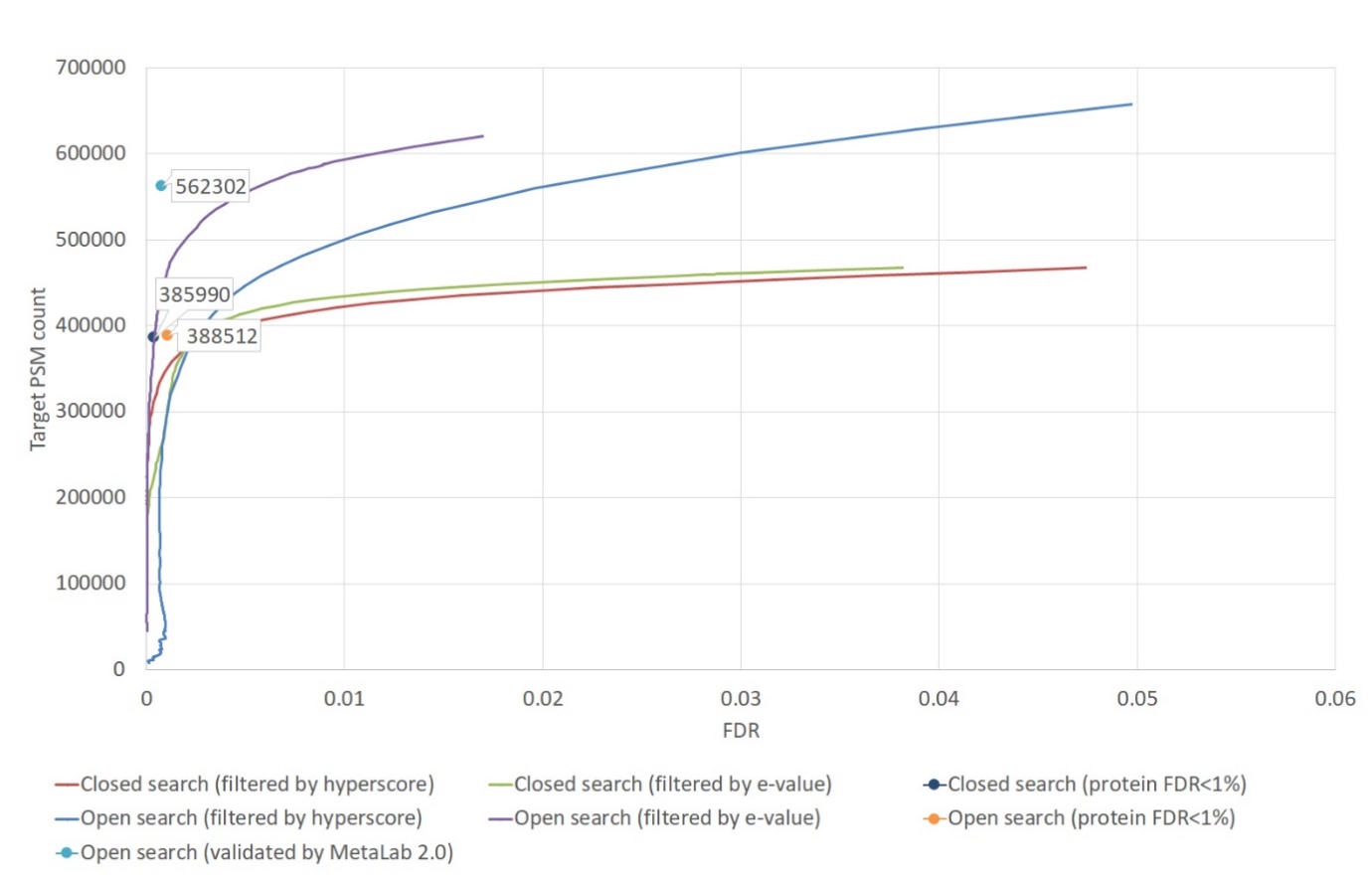
Supplementary Figure 2. **Comparison of peptide identifications from open search and conventional closed search in a Homo_HEK293 dataset.** The curves were obtained by increasing the score thresholds (hyperscore or e-value) gradually to get the corresponding FDR and target PSM number. The dots were obtained by restricting the FDRs in both of the peptide and protein levels below 0.01 then got the FDR and the target PSM number. In MetaLab 2.0 workflow a developed multistage filtering strategy was utilized. As a comparison, if this method was not used, a protein inference method was applied directly based on the search result to get the scores of proteins and peptides.
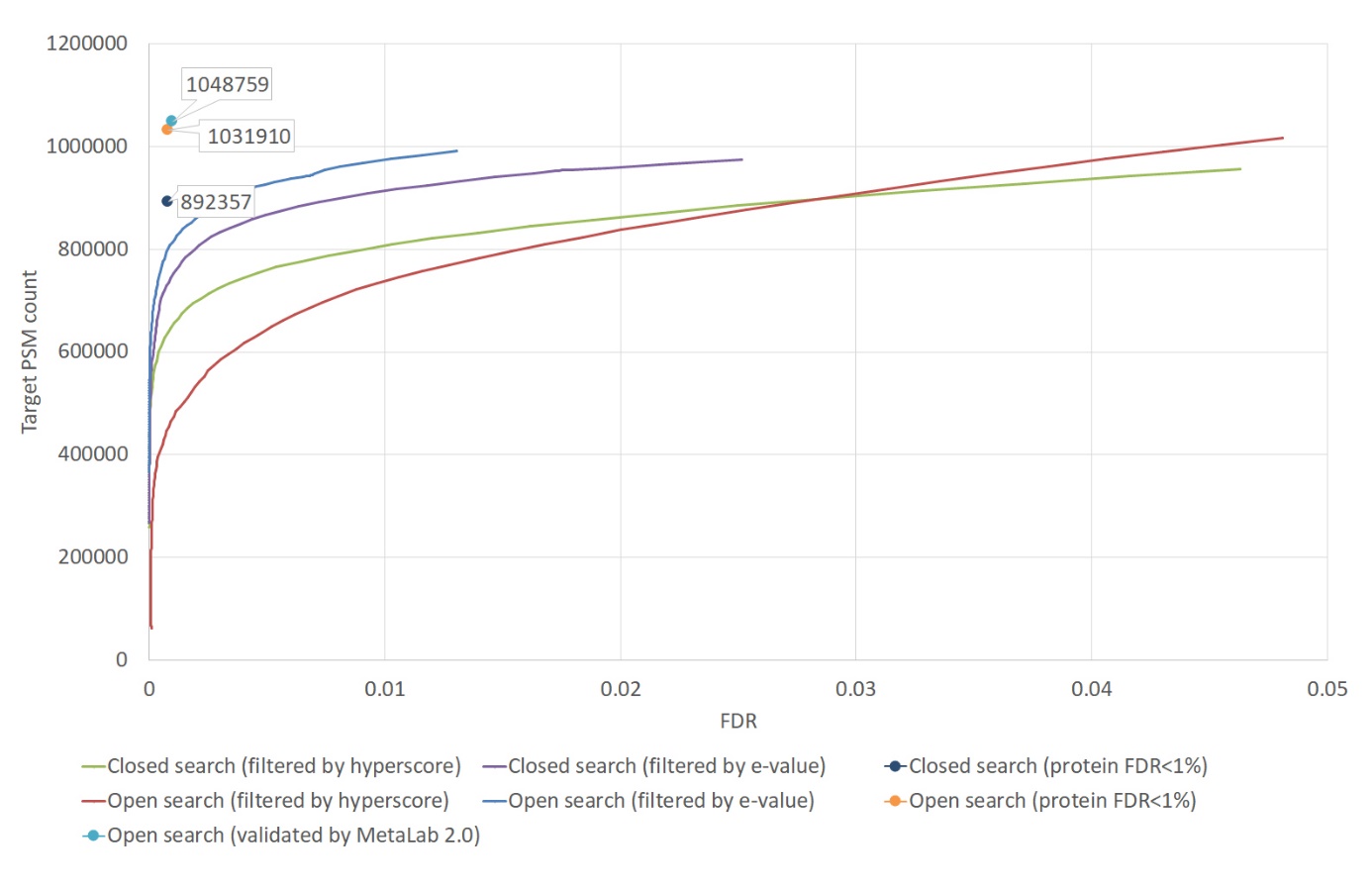

**Supplementary Figure 3. Comparison of peptide identifications from open search and conventional closed search in a Mock_micro dataset.** The curves were obtained by increasing the score thresholds (hyperscore or e-value) gradually to get the corresponding FDR and target PSM number. The dots were obtained by restricting the FDRs in both of the peptide and protein levels below 0.01 then got the FDR and the target PSM number. In MetaLab 2.0 workflow a developed multistage filtering strategy was utilized. As a comparison, if this method was not used, a protein inference method was applied directly based on the search result to get the scores of proteins and peptides.

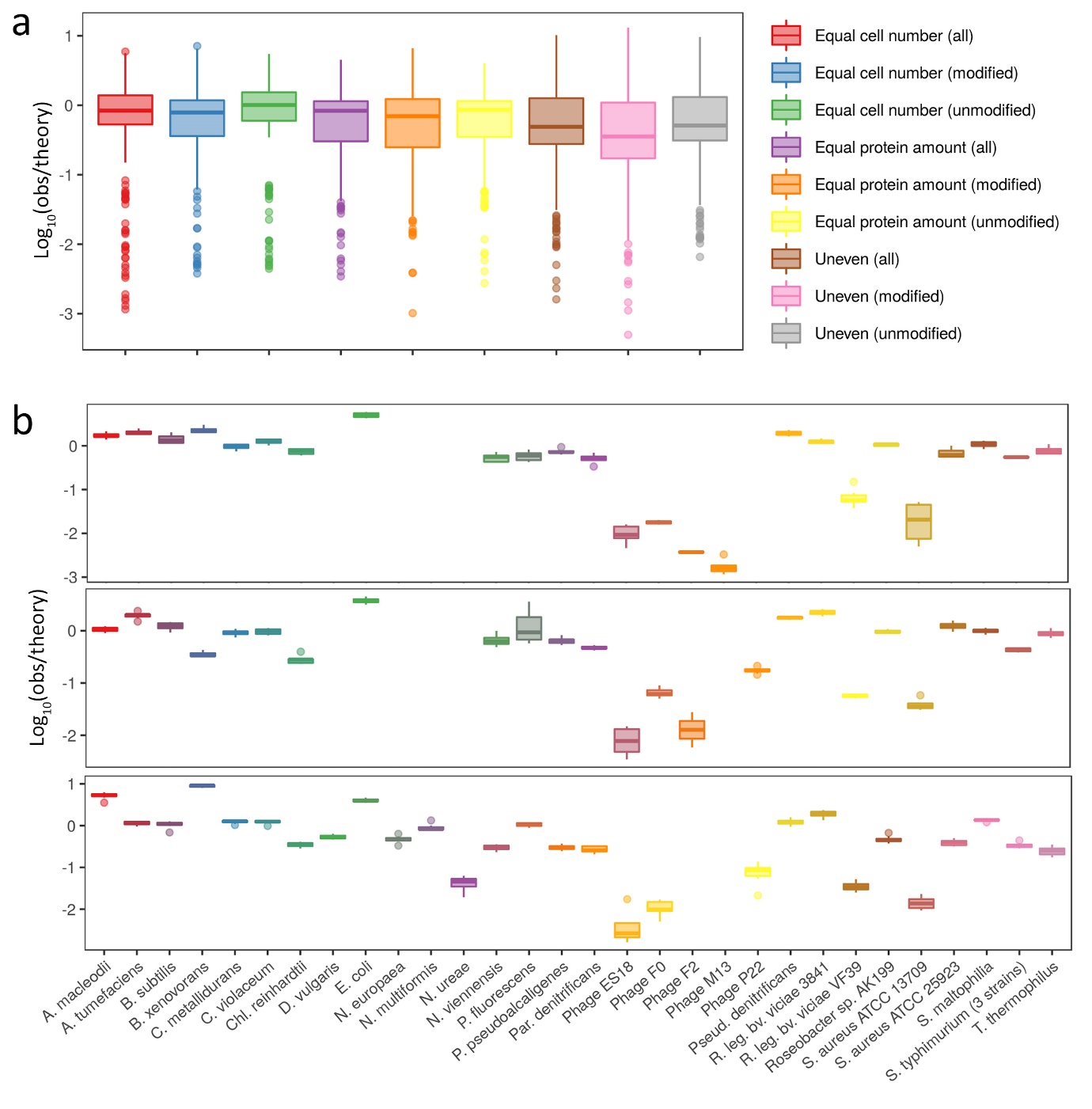

**Supplementary Figure 4. Quantitative analysis of a mock microbiome dataset.** (a) Deviation between the observed and theoretic relative abundance of species in different samples. The abundance of species were estimated by all the peptides, only unmodified peptides and only modified peptides, respectively. (b) Quantification of the mock communities at the species levels. The bottom and top of the boxes are the first and third quartiles, respectively, the middle lines represent the sample median.

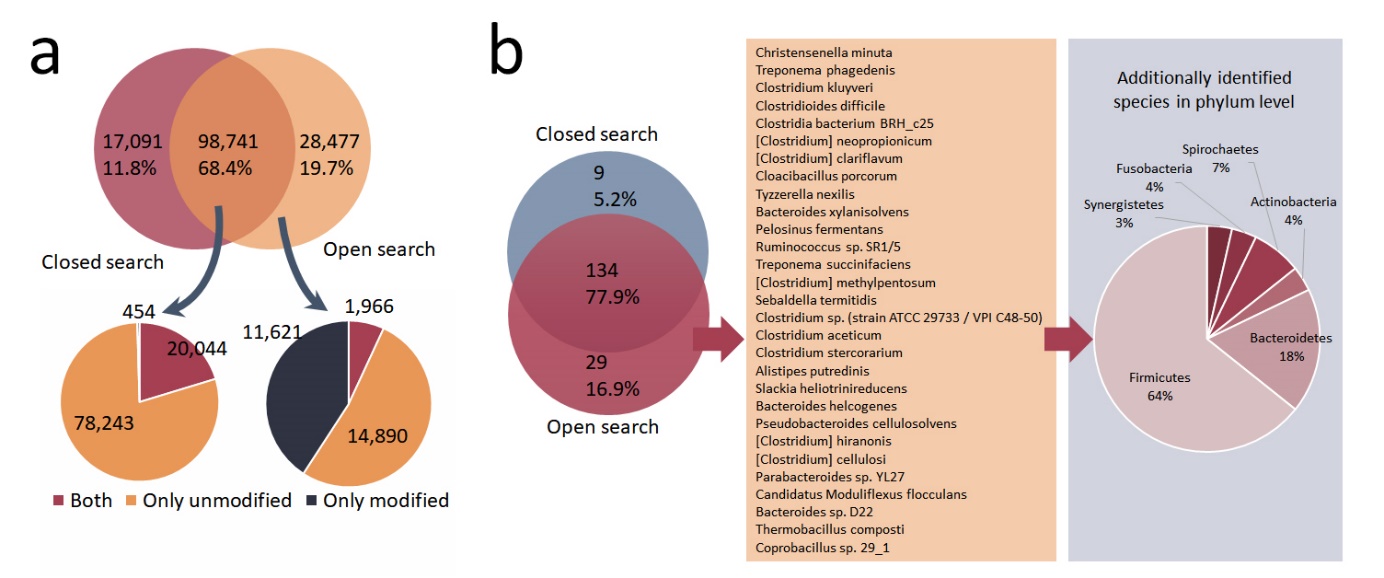

**Supplementary Figure 5. The comparison of the identified (a) peptides and (b) species by closed search and open search from mouse gut microbiota samples.**

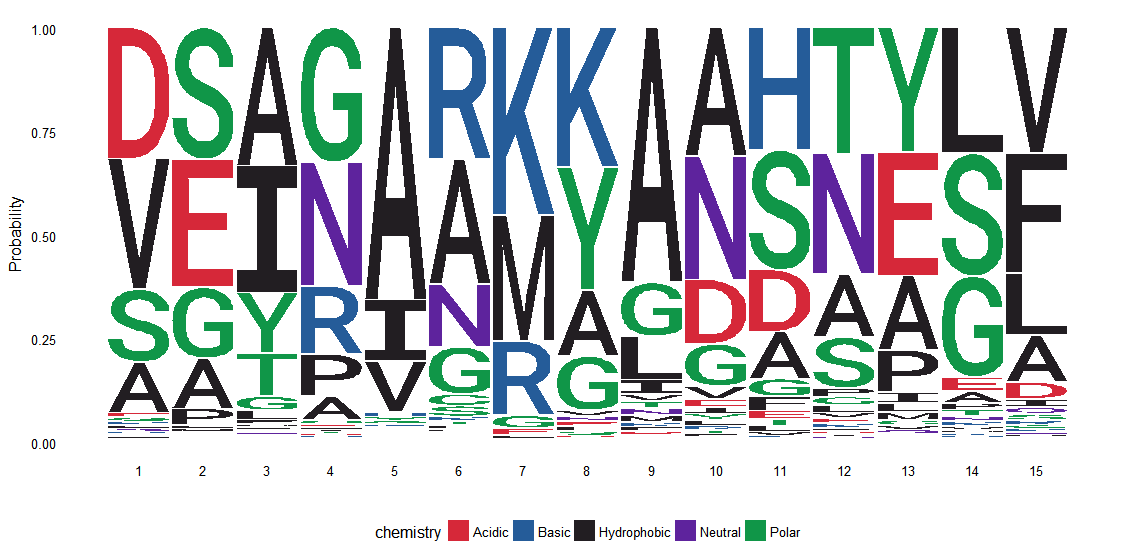
**Supplementary Figure 6. The sequence motif of the potential modification ∆Mass=71.0368 in the Mouse_gut dataset.** The length of the sequence window was 7 so the “8” in the horizontal axis represents the localization site.

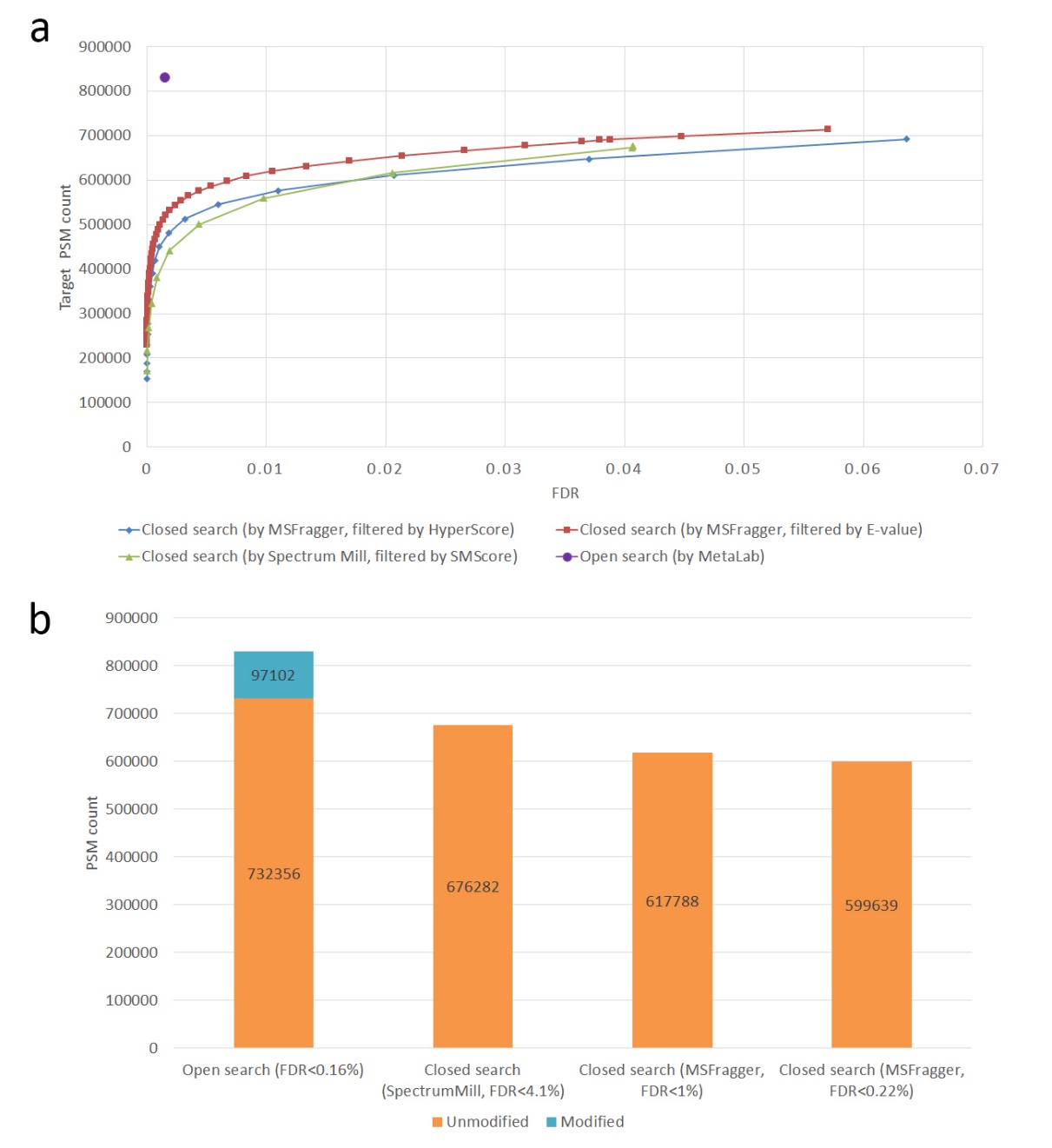

**Supplementary Figure 7. The comparison of peptide identifications from open search and conventional narrow search in Human_gut dataset.** (a) At the same FDR, far more PSMs were obtained using our strategy. The Spectrum Mill search result was downloaded from <https://www.ebi.ac.uk/pride/archive/projects/PXD008870/files;jsessionid=3C8E74D24253F9EABDBE1E885B89394C> (b) More PSMs were identified by open search, even only considering the unmodified peptides (peptides with oxidation on Met and/or acetylation on protein n-term were also included in this part, the modified part represented the peptides with other modifications identified by open search). The second column: the results were downloaded from the above link directly and no filtering was performed. Less PSMs were obtained at a much higher FDR. The third column: filtered PSMs by e-value to restrict the FDR<0.01. The fourth column: performed filtering at protein, peptide and PSMs levels simultaneously to keep the FDRs at all of the three levels below 0.01.

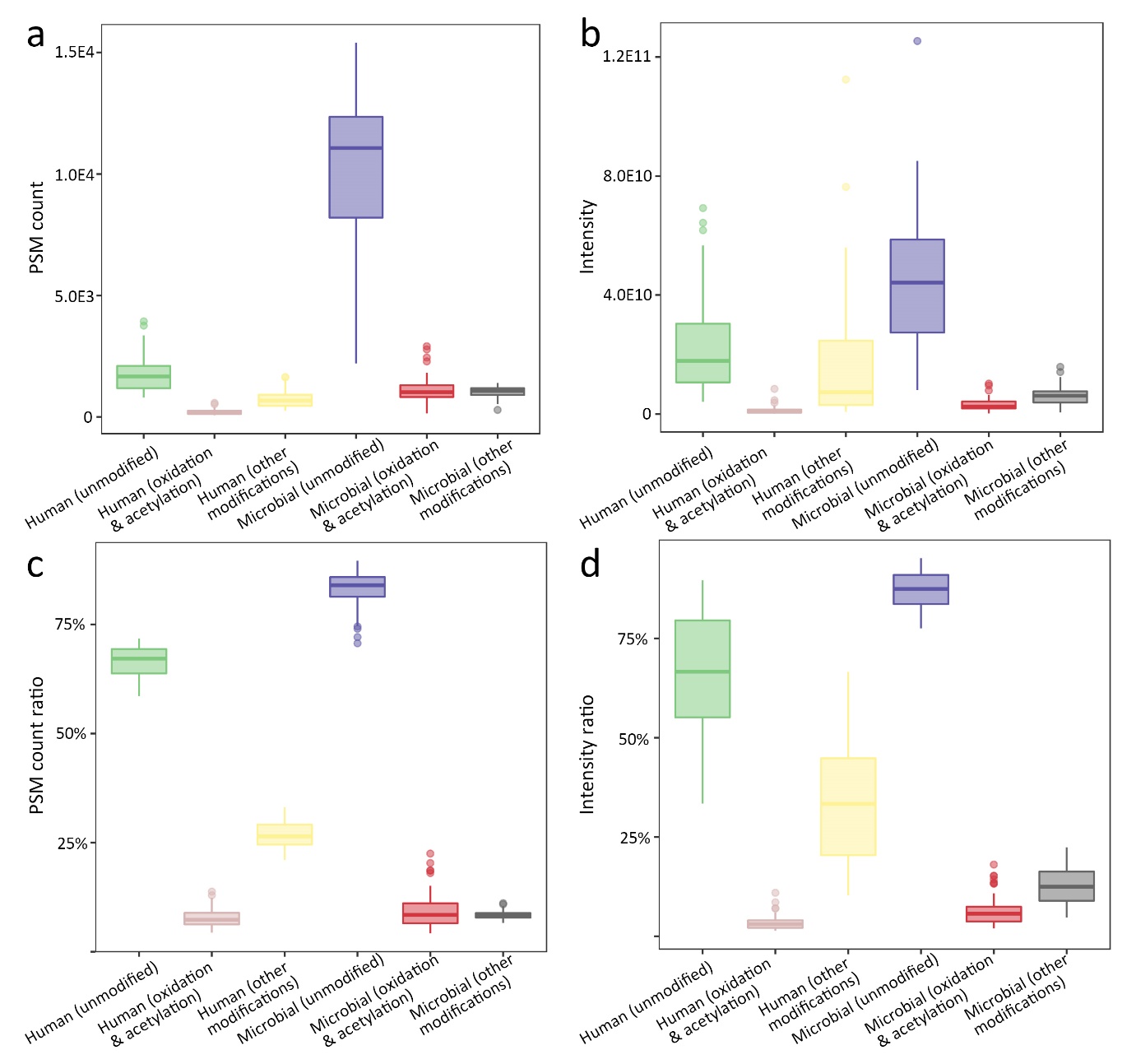

**Supplementary Figure 8. The PSM counts and intensities of unmodified, oxidation/acetylation and modified peptides from human and microbial proteins.** (a)(b) showed the absolute values of the PSM count and intensity. (c)(d) showed the relative values, i.e., the ratios of unmodified, oxidation/acetylation and modified peptides of the total amount from human and microbial proteins. The bottom and top of the boxes are the first and third quartiles, respectively, the middle lines represent the sample median.

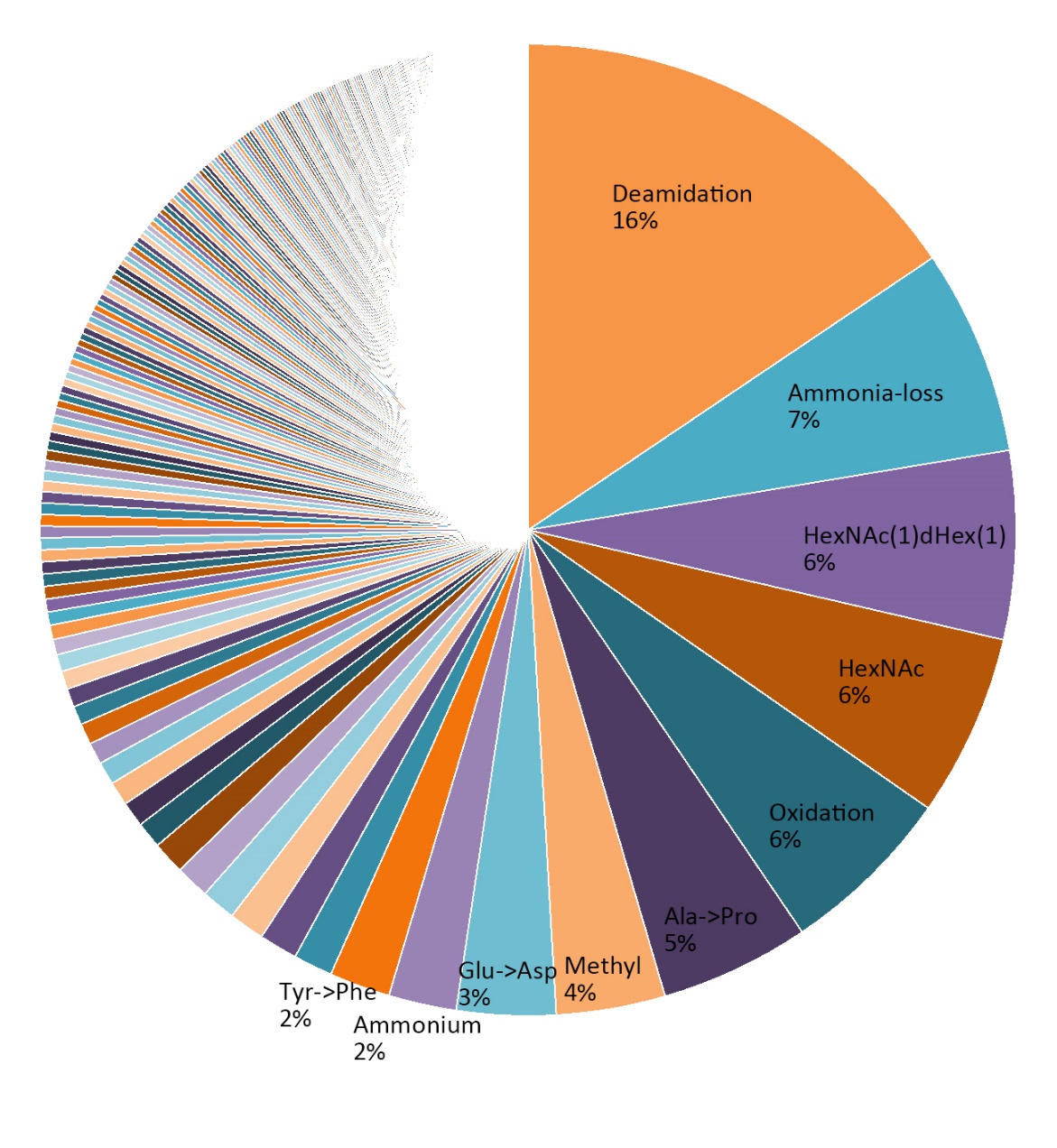

**Supplementary Figure 9. The PSM count ratio of the identified potential modifications from Human_gut.**

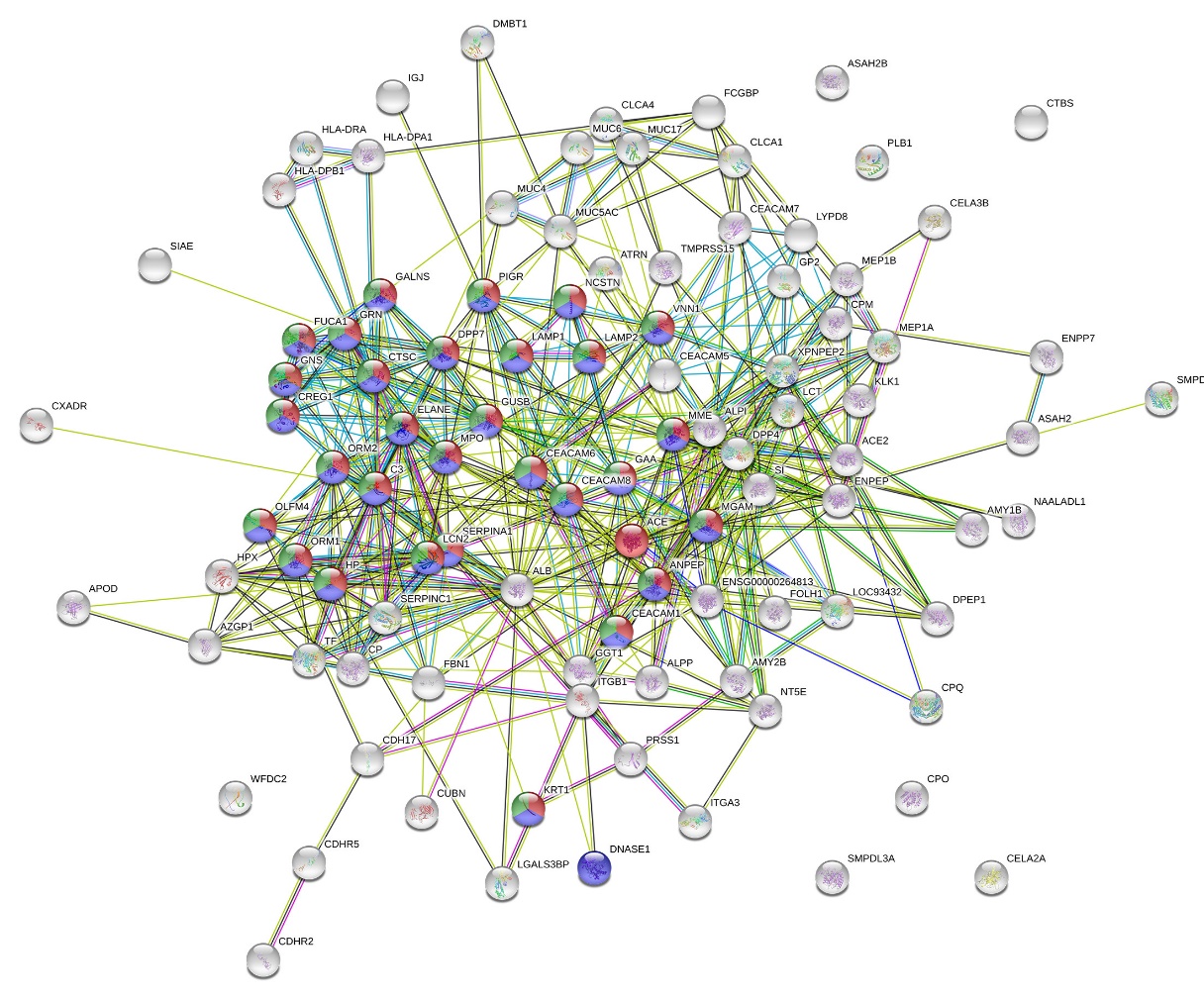
**Supplementary Figure 10. The network of HexNAc modified human glycoproteins.** Red: neutrophil mediated immunity (GO:0002446); blue: neutrophil activation involved in immune response (GO:0002283); green: neutrophil degranulation (GO:0043312).

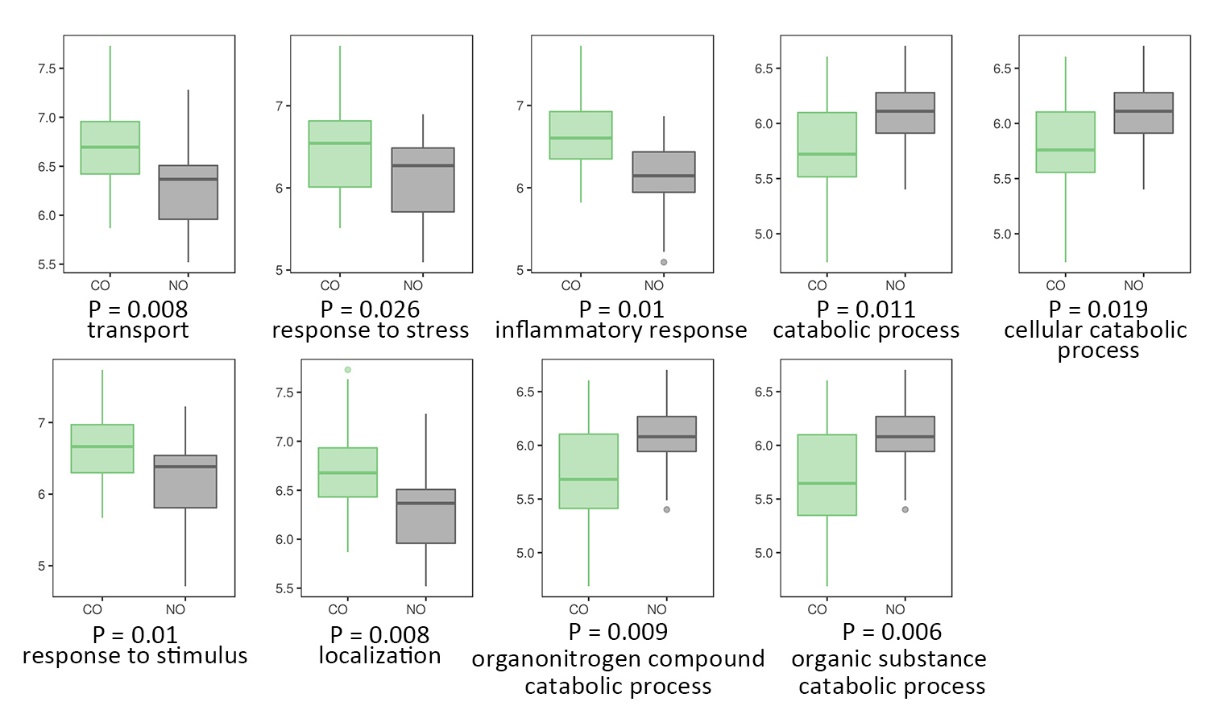

**Supplementary Figure 11. The significantly changed (P<=0.05 by two-sample, two-tailed t-test) GO biological progresses of HexNAc modified human glycoproteins.** The bottom and top of the boxes are the first and third quartiles, respectively, the middle lines represent the sample median.

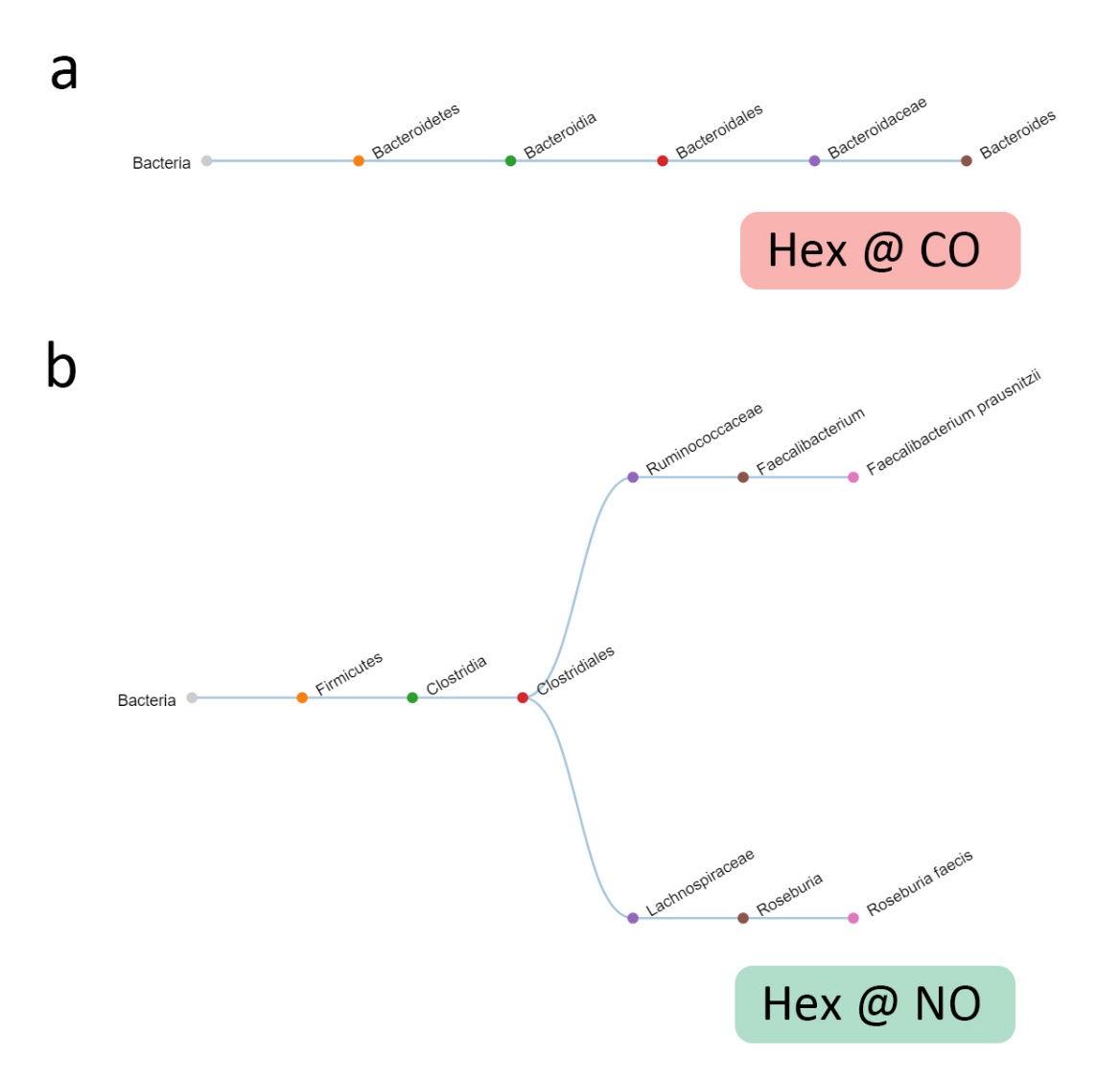

**Supplementary Figure 12. The taxon of Hex modified glycopeptides from CO and NO samples.**

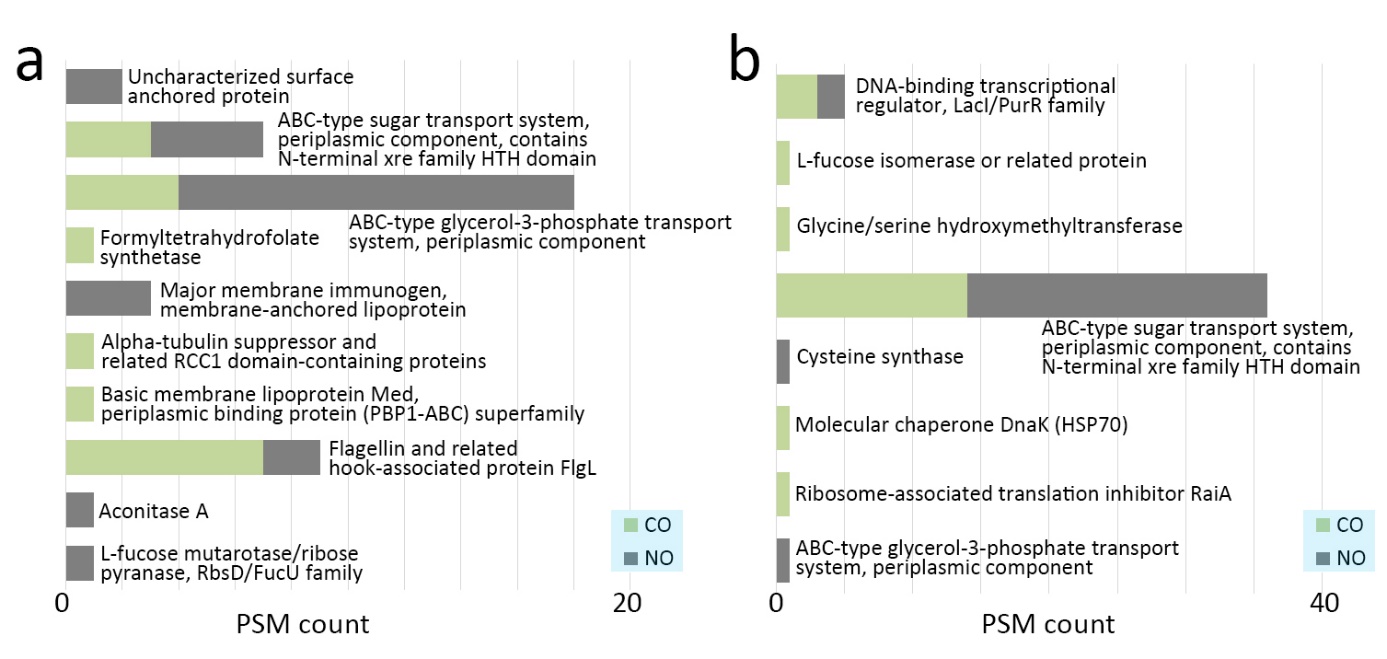

**Supplementary Figure 13. The functions of the microbial proteins modified by (a) Hex and (b) HexNAc**. CO: control subjects; NO; new-onset patients.

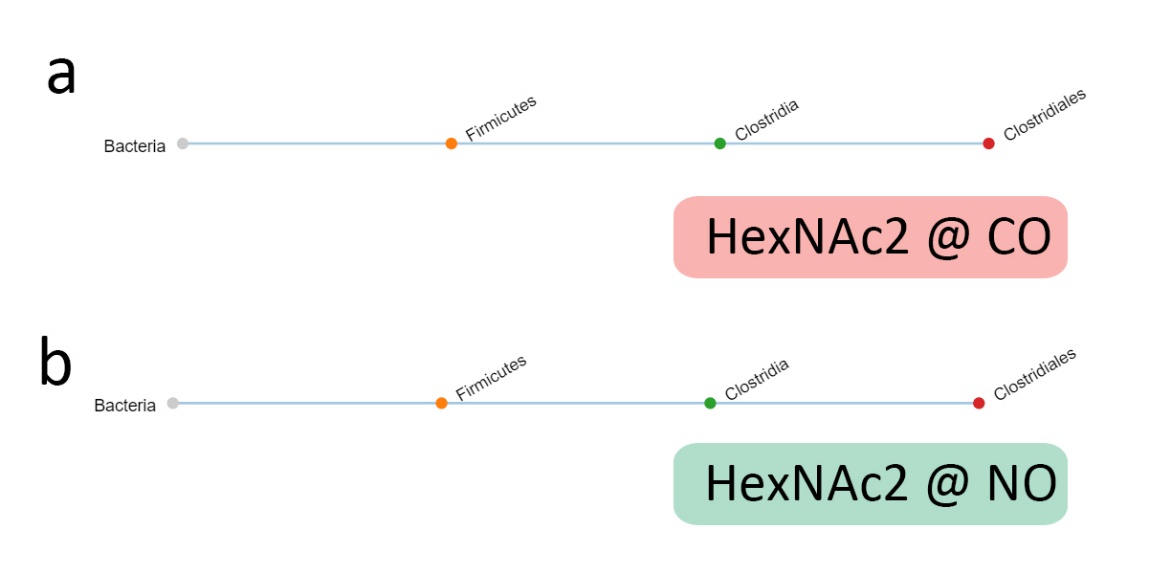

**Supplementary Figure 14. The taxon of HexNAc2 modified glycopeptides from CO and NO samples.**

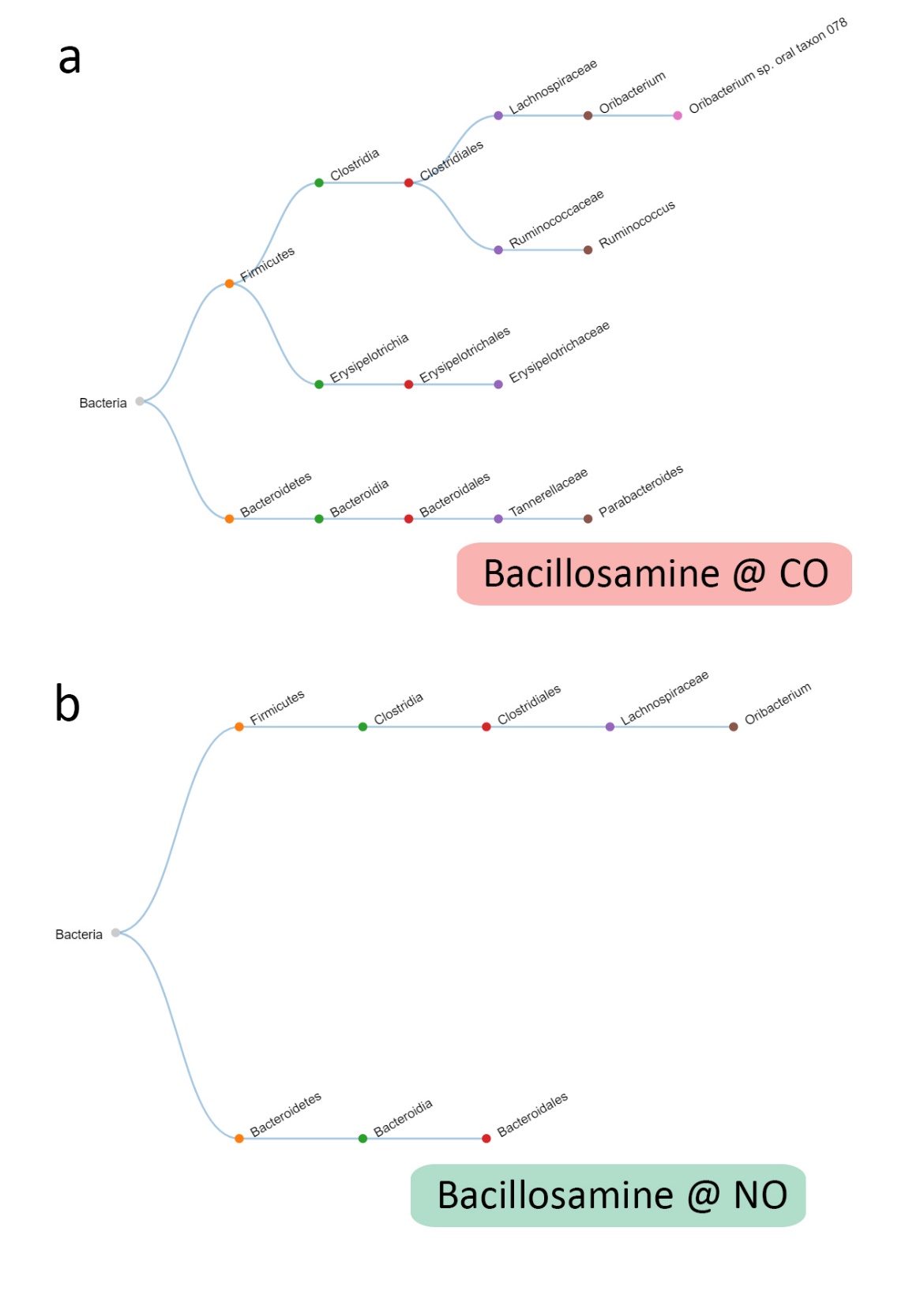

**Supplementary Figure 15. The taxon of Bacillosamine modified glycopeptides from CO and NO samples.**

**Supplementary Table 1. The features used for the machine learning classification.**

| Name | Description |
| --- | --- |
| charge | Charge state of the precursor ion |
| pepMass | The molecular weight of the peptide |
| pepLen | The length of the peptide |
| premr | The mass of the precursor ion |
| massDiff | The mass difference (Da) |
| massDiffPPM | The mass difference (PPM) |
| miss | The number of miss cleavages |
| num_tol_term | The number of tryptic termini |
| tot_num_ions | The number of peptide fragment ions predicted for peptide |
| num_matched_ions | The number of matched ions |
| hyperscore | The hyperscore of the PSM |
| nextscore | The hyperscore of the second ranked match |
| deltascore | The difference between the top hyperscore and second hyperscore |
| _logExpect | -Log_10_(Expect) |
| bintensity | The total intensity of the matched b ions |
| yintensity | The total intensity of the matched y ions |
| brankscore | The total rank score of the matched b ions (if there are N peaks in a MS2 spectra, and for one peak its intensity is higher than M peaks (max value of M is N-1), the rank score is calculated as (M+1)/N) |
| yrankscore | The total rank score of the matched y ions |
| bcount | The number of matched b ions |
| ycount | The number of matched y ions |
| fragDeltaMass | The average ∆Mass of matched fragment ions |
| modMass | The experimental mass of the potential modification |
| bmodIntensity | The total intensity of the matched b ions with the modification part |
| ymodIntensity | The total intensity of the matched y ions with the modification part |
| bmodrankscore | The total rank score of the matched b ions with the modification part |
| ymodrankscore | The total rank score of the matched y ions with the modification part |
| fragModDeltaMass | The average ∆Mass of matched fragment ions with the modification part |
| gaussian_a | The parameter a of the ∆Mass Gaussian peak (f_(x)_=ae^-(x-b)(x-b)/c^) |
| gaussian_b | The parameter b of the ∆Mass Gaussian peak (f_(x)_=ae^-(x-b)(x-b)/c^) |
| gaussian_r2 | The coefficient of the curving fitting (1-SS_reg_/SS_tot_) |
| sameModCount | The total number of PSMs with the same modification |
| samePepCount | The total number of PSMs with the same sequence |
| sameProCount | The total number of PSMs with the same protein |

**Supplementary Table 2. The 40 unknown ∆Mass which can be constructed from two known modifications already identified from this dataset.**

| Total mass | Mass 1 | Name 1 | Mass 2 | Name 2 |
| --- | --- | --- | --- | --- |
| -113.085 | 43.0058 | Ala->Asn substitution | -156.102 | Loss of arginine due to transpeptidation |
| -85.0898 | 43.0058 | Ala->Asn substitution | -128.096 | Loss of C-terminal K from Heavy Chain of MAb |
| -85.0898 | -14.0158 | Ala->Gly substitution | -71.0743 | Lys->Gly substitution |
| -145.122 | -17.0259 | Pyro-glu from Q | -128.096 | Loss of C-terminal K from Heavy Chain of MAb |
| -99.0688 | 57.022 | Addition of Glycine | -156.102 | Loss of arginine due to transpeptidation |
| -170.106 | -14.0158 | Ala->Gly substitution | -156.102 | Loss of arginine due to transpeptidation |
| 171.1008 | 43.0058 | Ala->Asn substitution | 128.0951 | Addition of lysine due to transpeptidation |
| 241.1791 | 128.0951 | Addition of lysine due to transpeptidation | 113.0833 | Acetylhypusine |
| -146.105 | -128.096 | Loss of C-terminal K from Heavy Chain of MAb | -18.0106 | Dehydration |
| 355.9057 | 301.9881 | Unidentified modification of 301.9864 found in open search | 53.9191 | Replacement of 2 protons by iron |
| -199.133 | -128.096 | Loss of C-terminal K from Heavy Chain of MAb | -71.0375 | Gln->Gly substitution |
| -128.059 | -71.0375 | Gln->Gly substitution | -57.0221 | Gln->Ala substitution |
| -128.059 | 28.0317 | di-Methylation | -156.102 | Loss of arginine due to transpeptidation |
| -87.0327 | -71.0375 | Gln->Gly substitution | -15.9946 | Tyr->Phe substitution |
| -87.0327 | -57.0221 | Gln->Ala substitution | -30.0113 | Thr->Ala substitution |
| -114.044 | -57.0221 | Gln->Ala substitution | -57.0221 | Gln->Ala substitution |
| 311.1296 | 183.0357 | Aminoethylbenzenesulfonylation | 128.0951 | Addition of lysine due to transpeptidation |
| -112.101 | 15.9953 | Phe->Tyr substitution | -128.096 | Loss of C-terminal K from Heavy Chain of MAb |
| -112.101 | 43.9908 | Ala->Asp substitution | -156.102 | Loss of arginine due to transpeptidation |
| 199.1075 | 43.0058 | Ala->Asn substitution | 156.1014 | Addition of arginine due to transpeptidation |
| 199.1075 | 113.0833 | Acetylhypusine | 86.0383 | 2-hydroxyisobutyrylation |
| -96.1053 | 31.9903 | Pro->Glu substitution | -128.096 | Loss of C-terminal K from Heavy Chain of MAb |
| 242.1378 | 156.1014 | Addition of arginine due to transpeptidation | 86.0383 | 2-hydroxyisobutyrylation |
| -101.049 | -71.0375 | Gln->Gly substitution | -30.0113 | Thr->Ala substitution |
| 269.1851 | 156.1014 | Addition of arginine due to transpeptidation | 113.0833 | Acetylhypusine |
| -227.164 | -156.102 | Loss of arginine due to transpeptidation | -71.0743 | Lys->Gly substitution |
| -74.1762 | 53.9191 | Replacement of 2 protons by iron | -128.096 | Loss of C-terminal K from Heavy Chain of MAb |
| 284.1968 | 128.0951 | Addition of lysine due to transpeptidation | 156.1014 | Addition of arginine due to transpeptidation |
| 69.9146 | 53.9191 | Replacement of 2 protons by iron | 15.9953 | Phe->Tyr substitution |
| -227.127 | -71.0375 | Gln->Gly substitution | -156.102 | Loss of arginine due to transpeptidation |
| -204.09 | -156.102 | Loss of arginine due to transpeptidation | -48.0043 | Prompt loss of side chain from oxidised Met |
| -186.101 | -156.102 | Loss of arginine due to transpeptidation | -30.0113 | Thr->Ala substitution |
| -172.085 | -15.9946 | Tyr->Phe substitution | -156.102 | Loss of arginine due to transpeptidation |
| -100.101 | 27.9954 | Formylation | -128.096 | Loss of C-terminal K from Heavy Chain of MAb |
| -100.101 | -71.0743 | Lys->Gly substitution | -29.028 | Lys->Val substitution |
| 212.1645 | 113.0833 | Acetylhypusine | 99.0671 | N-isopropylcarboxamidomethyl |
| -312.216 | -156.102 | Loss of arginine due to transpeptidation | -156.102 | Loss of arginine due to transpeptidation |
| 256.191 | 128.0951 | Addition of lysine due to transpeptidation | 128.0951 | Addition of lysine due to transpeptidation |
| 39.9035 | 53.9191 | Replacement of 2 protons by iron | -14.0158 | Ala->Gly substitution |
| 181.0395 | -15.9946 | Tyr->Phe substitution | 197.0491 | glycerylphosphorylethanolamine |
| -144.055 | -156.102 | Loss of arginine due to transpeptidation | 12.0376 | Thr->Leu/Ile substitution |
| 215.1274 | 203.0814 | N-Acetylhexosamine | 12.0376 | Thr->Leu/Ile substitution |
| 147.0344 | 31.9903 | Pro->Glu substitution | 115.0265 | Cleavage product of EGS protein crosslinks by hydroylamine treatment |
| 312.2021 | 156.1014 | Addition of arginine due to transpeptidation | 156.1014 | Addition of arginine due to transpeptidation |
| 36.893 | -17.0259 | Pyro-glu from Q | 53.9191 | Replacement of 2 protons by iron |
| -213.112 | -57.0221 | Gln->Ala substitution | -156.102 | Loss of arginine due to transpeptidation |

**Supplementary Table 3. The glycan oxonium ions used for the determination of glycopeptides spectra.**

| Name | Composition | MW | Glycan oxonium ion 1 | Glycan oxonium ion 2 |
| --- | --- | --- | --- | --- |
| HexNAc | C_8_O_5_NH_13_ | 203.0794 | 204.0867 | 186.0761 |
| HexNAc1dHex1 | C_14_O_9_NH_23_ | 349.1373 | 350.1444 | 332.1338‬‬‬‬‬‬‬‬‬‬‬ |
| HexNAc2 | C_16_O_10_N_2_H_26_ | 406.1588‬‬‬‬‬‬‬‬‬‬‬ | 204.0867 | 186.0761 |
| bacillosamine | C_10_O_4_N_2_H_16_ | 228.1110 | 229.1183‬‬‬‬‬‬‬‬‬‬‬ | 211.1077‬‬‬‬‬‬‬‬‬‬‬ |

**Supplementary Table 4. The information of the analyzed benchmark dataset.**

| Dataset ID | Supplementary Data ID | PRIDE id | Species | Tissues/Cell | Enrichment information | Raw file count |
| --- | --- | --- | --- | --- | --- | --- |
| Homo_HEK293^1^ | 1 | PXD001468 | Homo sapiens | HEK293 | - | 32 |
| Ecoli_phos^2^ | 2 | PXD008289 | Escherichia coli | - | Phosphorylation | 9 |
| Mouse_phos^3^ | 3 | PXD001792 | Mus musculus | Kidney | Phosphorylation | 6 |
| Hela_phos^4^ | 4 | PXD004940 | Homo sapiens | HeLa | Phosphorylation | 6 |
| U87_phos^5^ | 5 | PXD009227 | Homo sapiens | U-87 glioblastoma cells | Phosphorylation | 24 |
| Mock_micro^6^ | 6 | PXD006118 | Mock microbiome | - | - | 24 |
| Mouse_gut^7^ | 7 | PXD003527 | Mouse gut microbial community | - | - | 32 |
| Human_gut^8^ | 8 | PXD008870 | Human gut microbial community | - | - | 55 |

**Supplementary Table 5. The enriched Biological Process (GO) from the HexNAc modified human proteins.**

| #term ID | term description | observed gene count | background gene count | false discovery rate |
| --- | --- | --- | --- | --- |
| GO:0002283 | neutrophil activation involved in immune response | 32 | 489 | 5.96E-22 |
| GO:0002446 | neutrophil mediated immunity | 32 | 498 | 5.96E-22 |
| GO:0043312 | neutrophil degranulation | 31 | 485 | 1.06E-21 |
| GO:0002366 | leukocyte activation involved in immune response | 33 | 616 | 4.09E-21 |
| GO:0045055 | regulated exocytosis | 34 | 691 | 7.65E-21 |
| GO:0045321 | leukocyte activation | 36 | 894 | 1.61E-19 |
| GO:0002252 | immune effector process | 36 | 927 | 4.48E-19 |
| GO:0046903 | secretion | 36 | 1070 | 3.77E-17 |
| GO:0002376 | immune system process | 50 | 2370 | 9.78E-17 |
| GO:0006955 | immune response | 40 | 1560 | 2.04E-15 |
| GO:0016192 | vesicle-mediated transport | 40 | 1699 | 3.45E-14 |
| GO:0006508 | proteolysis | 34 | 1203 | 6.35E-14 |
| GO:0050435 | amyloid-beta metabolic process | 7 | 16 | 2.13E-09 |
| GO:0035333 | Notch receptor processing, ligand-dependent | 6 | 7 | 2.93E-09 |
| GO:0016485 | protein processing | 12 | 142 | 3.21E-09 |
| GO:1901564 | organonitrogen compound metabolic process | 61 | 5281 | 3.25E-09 |
| GO:0006810 | transport | 52 | 4130 | 1.42E-08 |
| GO:0002682 | regulation of immune system process | 29 | 1391 | 1.59E-08 |
| GO:0043603 | cellular amide metabolic process | 21 | 732 | 3.29E-08 |
| GO:0051179 | localization | 57 | 5233 | 2.38E-07 |
| GO:0042987 | amyloid precursor protein catabolic process | 5 | 9 | 4.59E-07 |
| GO:0006952 | defense response | 25 | 1234 | 5.31E-07 |
| GO:0019538 | protein metabolic process | 49 | 4194 | 7.31E-07 |
| GO:0006518 | peptide metabolic process | 16 | 497 | 8.20E-07 |
| GO:0022610 | biological adhesion | 20 | 849 | 1.78E-06 |
| GO:0034205 | amyloid-beta formation | 4 | 5 | 4.74E-06 |
| GO:0007155 | cell adhesion | 19 | 843 | 7.34E-06 |
| GO:0071704 | organic substance metabolic process | 76 | 9135 | 7.89E-06 |
| GO:0006509 | membrane protein ectodomain proteolysis | 5 | 20 | 9.09E-06 |
| GO:0002526 | acute inflammatory response | 7 | 73 | 1.10E-05 |
| GO:0050776 | regulation of immune response | 19 | 873 | 1.14E-05 |
| GO:0006898 | receptor-mediated endocytosis | 10 | 209 | 1.27E-05 |
| GO:0008152 | metabolic process | 77 | 9569 | 2.55E-05 |
| GO:0031347 | regulation of defense response | 16 | 676 | 3.65E-05 |
| GO:0002003 | angiotensin maturation | 4 | 11 | 4.15E-05 |
| GO:0043171 | peptide catabolic process | 5 | 31 | 5.24E-05 |
| GO:0005975 | carbohydrate metabolic process | 13 | 457 | 5.58E-05 |
| GO:0007586 | digestion | 7 | 104 | 8.52E-05 |
| GO:0050896 | response to stimulus | 66 | 7824 | 0.00012 |
| GO:0098609 | cell-cell adhesion | 12 | 416 | 0.00012 |
| GO:0044238 | primary metabolic process | 71 | 8808 | 0.00015 |
| GO:0050778 | positive regulation of immune response | 14 | 589 | 0.00015 |
| GO:0098742 | cell-cell adhesion via plasma-membrane adhesion molecules | 9 | 230 | 0.0002 |
| GO:0002576 | platelet degranulation | 7 | 129 | 0.00029 |
| GO:0002253 | activation of immune response | 11 | 393 | 0.00037 |
| GO:0044245 | polysaccharide digestion | 3 | 6 | 0.00039 |
| GO:0006954 | inflammatory response | 12 | 482 | 0.00044 |
| GO:0046718 | viral entry into host cell | 6 | 92 | 0.00046 |
| GO:0009056 | catabolic process | 25 | 1859 | 0.00052 |
| GO:1901565 | organonitrogen compound catabolic process | 17 | 958 | 0.00052 |
| GO:0002768 | immune response-regulating cell surface receptor signaling pathway | 9 | 266 | 0.00053 |
| GO:0002684 | positive regulation of immune system process | 16 | 882 | 0.00067 |
| GO:0006897 | endocytosis | 12 | 510 | 0.00067 |
| GO:0051701 | interaction with host | 7 | 156 | 0.00077 |
| GO:0098657 | import into cell | 13 | 609 | 0.00079 |
| GO:0007156 | homophilic cell adhesion via plasma membrane adhesion molecules | 7 | 158 | 0.00081 |
| GO:0002764 | immune response-regulating signaling pathway | 10 | 365 | 0.0009 |
| GO:0050900 | leukocyte migration | 9 | 296 | 0.001 |
| GO:0006026 | aminoglycan catabolic process | 5 | 67 | 0.0011 |
| GO:0002429 | immune response-activating cell surface receptor signaling pathway | 8 | 234 | 0.0012 |
| GO:1901575 | organic substance catabolic process | 22 | 1609 | 0.0012 |
| GO:0044248 | cellular catabolic process | 22 | 1646 | 0.0016 |
| GO:0080134 | regulation of response to stress | 19 | 1299 | 0.0017 |
| GO:0043170 | macromolecule metabolic process | 60 | 7453 | 0.002 |
| GO:0002757 | immune response-activating signal transduction | 9 | 332 | 0.0022 |
| GO:0006953 | acute-phase response | 4 | 45 | 0.0031 |
| GO:0090675 | intermicrovillar adhesion | 2 | 2 | 0.0037 |
| GO:0045088 | regulation of innate immune response | 9 | 361 | 0.0038 |
| GO:0006807 | nitrogen compound metabolic process | 64 | 8349 | 0.004 |
| GO:0006950 | response to stress | 33 | 3267 | 0.004 |
| GO:2000811 | negative regulation of anoikis | 3 | 18 | 0.004 |
| GO:0007157 | heterophilic cell-cell adhesion via plasma membrane cell adhesion molecules | 4 | 54 | 0.0056 |
| GO:0000023 | maltose metabolic process | 2 | 3 | 0.0058 |
| GO:0016266 | O-glycan processing | 4 | 57 | 0.0066 |
| GO:0046466 | membrane lipid catabolic process | 3 | 23 | 0.0071 |
| GO:0010669 | epithelial structure maintenance | 3 | 24 | 0.0079 |
| GO:1903596 | regulation of gap junction assembly | 2 | 4 | 0.0083 |
| GO:0002223 | stimulatory C-type lectin receptor signaling pathway | 4 | 62 | 0.0085 |
| GO:0006027 | glycosaminoglycan catabolic process | 4 | 62 | 0.0085 |
| GO:1901136 | carbohydrate derivative catabolic process | 6 | 175 | 0.0085 |
| GO:0006879 | cellular iron ion homeostasis | 4 | 66 | 0.0102 |
| GO:0050727 | regulation of inflammatory response | 8 | 338 | 0.0104 |
| GO:0006685 | sphingomyelin catabolic process | 2 | 5 | 0.0107 |
| GO:0042445 | hormone metabolic process | 6 | 186 | 0.0107 |
| GO:0045089 | positive regulation of innate immune response | 7 | 259 | 0.0107 |
| GO:0072338 | cellular lactam metabolic process | 2 | 6 | 0.0138 |
| GO:0044403 | symbiont process | 11 | 650 | 0.0151 |
| GO:0031349 | positive regulation of defense response | 8 | 365 | 0.016 |
| GO:0043687 | post-translational protein modification | 8 | 365 | 0.016 |
| GO:0032532 | regulation of microvillus length | 2 | 7 | 0.017 |
| GO:0006979 | response to oxidative stress | 8 | 373 | 0.0178 |
| GO:0051186 | cofactor metabolic process | 9 | 467 | 0.0181 |
| GO:0045785 | positive regulation of cell adhesion | 8 | 375 | 0.0182 |
| GO:0050818 | regulation of coagulation | 4 | 81 | 0.0189 |
| GO:0007229 | integrin-mediated signaling pathway | 4 | 84 | 0.0211 |
| GO:0044281 | small molecule metabolic process | 20 | 1779 | 0.0219 |
| GO:0002016 | regulation of blood volume by renin-angiotensin | 2 | 9 | 0.0243 |
| GO:0002438 | acute inflammatory response to antigenic stimulus | 2 | 9 | 0.0243 |
| GO:0008354 | germ cell migration | 2 | 9 | 0.0243 |
| GO:0097205 | renal filtration | 2 | 9 | 0.0243 |
| GO:0016477 | cell migration | 12 | 812 | 0.0246 |
| GO:0050852 | T cell receptor signaling pathway | 4 | 93 | 0.0285 |
| GO:0065008 | regulation of biological quality | 32 | 3559 | 0.0287 |
| GO:0006022 | aminoglycan metabolic process | 5 | 160 | 0.0299 |
| GO:0022409 | positive regulation of cell-cell adhesion | 6 | 238 | 0.0306 |
| GO:0001878 | response to yeast | 2 | 11 | 0.0316 |
| GO:0030155 | regulation of cell adhesion | 10 | 623 | 0.0316 |
| GO:0032101 | regulation of response to external stimulus | 11 | 732 | 0.0316 |
| GO:0097242 | amyloid-beta clearance | 2 | 11 | 0.0316 |
| GO:0051181 | cofactor transport | 3 | 46 | 0.0328 |
| GO:0002758 | innate immune response-activating signal transduction | 5 | 168 | 0.0341 |
| GO:0019731 | antibacterial humoral response | 3 | 47 | 0.0341 |
| GO:0042340 | keratan sulfate catabolic process | 2 | 12 | 0.0343 |
| GO:1904469 | positive regulation of tumor necrosis factor secretion | 2 | 12 | 0.0343 |
| GO:0042742 | defense response to bacterium | 6 | 250 | 0.0356 |
| GO:0001822 | kidney development | 6 | 251 | 0.0361 |
| GO:0043062 | extracellular structure organization | 7 | 339 | 0.0361 |
| GO:0006959 | humoral immune response | 6 | 252 | 0.0363 |
| GO:0002703 | regulation of leukocyte mediated immunity | 5 | 176 | 0.0379 |
| GO:0006684 | sphingomyelin metabolic process | 2 | 13 | 0.0379 |
| GO:0009251 | glucan catabolic process | 2 | 13 | 0.0379 |
| GO:0044130 | negative regulation of growth of symbiont in host | 2 | 13 | 0.0379 |
| GO:1901678 | iron coordination entity transport | 2 | 13 | 0.0379 |
| GO:0008217 | regulation of blood pressure | 5 | 177 | 0.0385 |
| GO:0031348 | negative regulation of defense response | 5 | 180 | 0.0411 |
| GO:0044247 | cellular polysaccharide catabolic process | 2 | 14 | 0.0411 |
| GO:0044273 | sulfur compound catabolic process | 3 | 53 | 0.0421 |
| GO:0006575 | cellular modified amino acid metabolic process | 5 | 185 | 0.0447 |
| GO:0002922 | positive regulation of humoral immune response | 2 | 15 | 0.0448 |
| GO:0009311 | oligosaccharide metabolic process | 3 | 55 | 0.0449 |
| GO:0051704 | multi-organism process | 22 | 2222 | 0.0449 |
| GO:0045087 | innate immune response | 10 | 676 | 0.046 |
| GO:0022407 | regulation of cell-cell adhesion | 7 | 366 | 0.0476 |
| GO:0016032 | viral process | 9 | 571 | 0.0478 |
| GO:0033013 | tetrapyrrole metabolic process | 3 | 57 | 0.048 |
| GO:0050728 | negative regulation of inflammatory response | 4 | 117 | 0.048 |
| GO:0051883 | killing of cells in other organism involved in symbiotic interaction | 2 | 16 | 0.048 |
| GO:1903037 | regulation of leukocyte cell-cell adhesion | 6 | 278 | 0.0488 |
| GO:0008015 | blood circulation | 7 | 373 | 0.0498 |

### Supplementary note

#### Application on analyzing an artificial microbiota community consisted with pure cultures

To evaluate the performance of open search in metaproteomics studies, we analyzed a microbiome dataset which was constructed by assembling 32 species/ and strains of Archaea, Bacteria, Eukaryotes and Bacteriophages^6^. 1,048,759 PSMs were identified from 24 raw files with FDR about 0.1%. Totally 70,108 unique peptide sequences and 16,444 proteins were found with FDR at peptide and protein levels 0.24% and 1%, respectively (**Supplementary Data 6**). 31 .4% (22,043/70,108) of the unique peptides contained at least one modified forms. The ratio reached 80% (9,061/11,328) at the protein level, which meant four-fifths of proteins contained at least one modification. This result clearly illustrated that PTMs widely occured on proteins of microbes.

For this dataset, the mock communities were mixed with three different ways, i.e., equal cell number mixture, equal protein amount mixture and uneven mixture. For each type the relative abundance of individual bacterial members were known. Therefore we used this dataset to benchmark the quantitative accuracy of our method. Firstly, we used all the peptides, only modified peptides and only unmodified peptides, respectively, to calculate the relative abundance of each species from each mixture types (**Supplementary Fig. 4a**). The distribution of relative abundance, determined in different ways, was consistent which also suggested the equal confidence of the modified and unmodified peptide identifications.

The species-level quantification result was showed in **Supplementary Fig. 4b**. As expected, for most of the species very good representations of the relative abundances were observed. For the equal cell number mixture and equal protein amount mixture accurate quantitative results were obtained for most of the species. Bacteriophages were observed with apparent deviations from the theory values because their abundance were ten times lower than other species. In the uneven mixture, the deviation was larger, may be because differences between individual species became larger and the theoretical abundances for some species were low.

#### Taxonomy analsyis and functional annotation of the glycopeptides identified in human gut microbiome dataset

Glycosylation plays a critical role in the regulation of protein functions. The alterations of glycosylation could affect the immune system of mammals^9^. The relationship between abnormal glycosylation and the pathogenesis of diabetes has also been reported^10^. Functional enrichment analysis of HexNAc modified proteins was performed (**Supplementary Table 5**). Neutrophil related immune response was the most significantly enriched biological processes (**Supplementary Fig. 10**). Among the enriched Gene Ontology (GO) terms, nine of them were changed (*p*<0.05 by two-sample, two-tailed t-test) (**Supplementary Fig. 11**). Five of them (transport, response to stress, inflammatory response, response to stimulus and localization) were also downregulated in NO samples which were consistent with the general trend. But four catabolic processes (catabolic process, cellular catabolic process, organonitrogen compound catabolic process and organic substance catabolic process) related functions were upregulated.

Besides high abundant human protein glycosylation, we also identified three types of glycosylation from microbial proteins. For hexose, 21 glycopeptides were identified, only one peptide was from *Bacteroidetes* and 16 peptides were from *Firmicutes* (**Supplementary Fig. 12, Supplementary Table 6**). In this dataset, 86.7% (39/45) of the hexose modified PSMs were from *Clostridiales*. From the functional annotation, we found that the most frequently observed function related to hexose was ABC-type glycerol-3-phosphate transport system, periplasmic component (**Supplementary Fig. 13**). For HexNAc2, one peptide EAATTAAAAAATTDVADKK was identified from eight CO and seven NO samples, respectively, which was the most widely identified glycopeptide from bacteria in Human_gut (**Supplementary Fig. 14, Supplementary Table 6**). The lowest common ancestor (LCA) of this peptide was *Clostridia* (order level). The corresponding protein was ABC-type sugar transport system, periplasmic component, contains N-terminal xre family HTH domain, which was also an ATP-binding cassette (ABC) transporter. It could be seen that the ABC transporter families were the most frequently identified glycoproteins from bacteria in Human_gut. We also identified bacteria specific monosaccharide Bacillosamine from 48 glycopeptides. Bacillosamine was described as an “Asn-linked glycan from Gram-negative Bacterium” in Unimod. However, we noticed that only 45.8% (22/48) glycopeptides contained Asn, which suggested the existence of other types such as O-linked glycopeptides in these samples. Bacillosamine was involved in various functions in this dataset (**Supplementary Table 6**). We identified the LCA for 21 glycopeptides (**Supplementary Fig. 15**). Three peptides determined in superkingdom level as bacteria were from the protein Elongation factor Tu (EF-Tu), which was a highly conserved proteins in prokaryotes. EF-Tu was a type of GTPase, which was also an important antibiotic target^11^. For the remainder of the glycopeptides, seven were from Bacteroidetes and eight were from Firmicutes. The main COG categories were carbohydrate transport and metabolism (G) and energy production and conversion (C). The characterization of bacteria glycosylation will greatly improve our understanding of gut microbiome-host interactions.

1.	Chick, J.M. et al. A mass-tolerant database search identifies a large proportion of unassigned spectra in shotgun proteomics as modified peptides. *Nat. Biotechnol.* **33**, 743-749 (2015).

2.	Potel, C.M., Lin, M.H., Heck, A.J.R. & Lemeer, S. Defeating major contaminants in Fe(3+)-IMAC phosphopeptide enrichment. *Mol Cell Proteomics* (2018).
